# Benchmarked sequencing methods reveal hidden mosaic variation in the human brain

**DOI:** 10.64898/2026.10.11.758380

**Authors:** Jiaming Weng, Do Hyeon Cha, Jinhui Chun, Pablo Santamarina-Ojeda, Yuhe Cheng, Robyn Howarth, Yixuan Xu, Xiangyu Huangfu, Pei-Jhen Wang, Saehoon Jung, Jinhee Ryu, Yunhye Noh, Sangmoon Lee, Xiaoxu Yang, Young Seok Ju, Ludmil B. Alexandrov, Changuk Chung, Joseph G. Gleeson

## Abstract

Mosaic variants (MVs) record cellular lineages, but their recovery is strongly method-dependent. Here, we benchmarked four duplex sequencing methods alongside bulk and single-neuron genome sequencing in matched human brain tissue. The methods differed in genome representation, MV yield and mutational profiles, with limited caller-level concordance. However, read-level interrogation revealed substantially greater cross-method molecular support. To systematically recover this evidence, we developed Read-level Integration for Variant Evidence Recovery (RIVER), expanding the pool of cross-method-supported candidates by approximately two orders of magnitude relative to caller-level overlap. Orthogonal sequencing at ∼10,000× across brain and peripheral tissues validated approximately half of the tested RIVER-rescued candidates. MVs recovered from a single brain region resolved broadly distributed and regionally restricted clones and provided lineage information among individual neurons. These findings show that RIVER substantially expands the pool of endogenous lineage markers available to investigate human development, aging, disease-associated mosaicism and clonal evolution.

## Introduction

Mosaic variants (MVs) accumulate throughout development to generate genetically distinct cell populations within an individual ^1^. When shared across multiple cells, MVs can provide endogenous lineage markers to reconstruct developmental relationships, clonal dynamics, and tissue remodeling ^2–5^. In the human brain, MVs have revealed clonal lateralization of the forebrain, regional restriction and long-range dispersion of progenitor-derived populations, providing insights into human-specific developmental processes *in vivo* ^6,7^. However, low variant allelic fraction (VAF) MVs may not validate across different methods and remain difficult to distinguish from technical artifacts using conventional sequencing, leaving an unexplored reservoir of MVs.

Duplex sequencing independently interrogates both complementary strands of the same DNA molecule, suppressing errors and enabling confident detection of rare MVs at VAFs orders of magnitude below those accessible by non-duplex methods. Current approaches include NanoSeq, CODEC, HiDEF-seq, ppmSeq, and UDSeq ^8–12^ differing in library construction, molecular tagging, sequencing platform (Illumina, PacBio, Ultima), strand-pair recovery approaches, genome coverage, and analytical processing. However, whether these methods capture distinct MV subsets, differ in their representation of clonal versus non-clonal MVs, or exhibit unique mutational signatures remains unclear.

Recent studies have benchmarked MV detection methods and callers, primarily using engineered reference mixtures ^13,14^. However, benchmarking and optimization of duplex sequencing methods remain limited, and how assay choice and read-support criteria shape orthogonal validation and clonal geography across human tissues remains unresolved. Therefore, an empirical basis for selecting a sequencing method, depth, and analysis strategy for a given biological question remains lacking.

Here, we systematically benchmarked HiDEF-seq, NanoSeq, ppmSeq, and UDSeq using an identical sample from a neurotypical human brain. We compared method-specific genomic coverage, sequence-context representation, and integrated duplex-derived MV candidates with standard Illumina whole-genome sequencing (IlluminaSeq) and single-neuron primary template-directed amplification-based whole-genome sequencing (PTA-Seq) datasets. We then stratified MVs according to cross-method concordance and read-level evidence, and evaluated 465 MV candidates at ∼10,000× targeted sequencing across eight bilateral brain regions and three peripheral tissues from the same donor. We discovered hidden concordance within the BAM/CRAM files that substantially increased confidence in the call sets. To advance this approach, we developed RIVER as a high-throughput, statistically rigorous approach to rescue MVs with cross-method evidence. RIVER expanded the cross-method supported set of MVs by approximately 100-fold relative to caller-level overlap, of which approximately 50% were validated in at least one sampled tissue. These findings demonstrate that caller-level concordance substantially underestimates shared mosaic variation and establish RIVER as a tool for recovering informative MVs often overlooked in existing sequencing datasets.

## Results

### Multiple sequencing approaches capture distinct features of the mosaic genome

We utilized the previously reported cadaveric donor ID01, from whom deep IlluminaSeq data (350x) across multiple bodily tissues were available ^15,16^. To benchmark multiple duplex sequencing methods in the same specimen, we selected a 2 × 1 × 1 cm block of neocortical gray matter from the left temporal cortex of donor ID01 as the reference site and isolated genomic DNA from this tissue (**Fig. 1a**). This cortical region had previously been analyzed by single-neuron PTA-Seq (∼30×), enabling evaluation of detected MVs at single-cell resolution. The extracted DNA was subjected to deep duplex sequencing using ppmSeq (∼350×), UDSeq (∼100×), NanoSeq (∼30×), and HiDEF-seq (∼30×) (**Fig. 1b**). In addition, sperm and blood samples from unrelated donors were included to estimate false-positive rates in the benchmarking analyses (**Methods**).

**Fig. 1.**
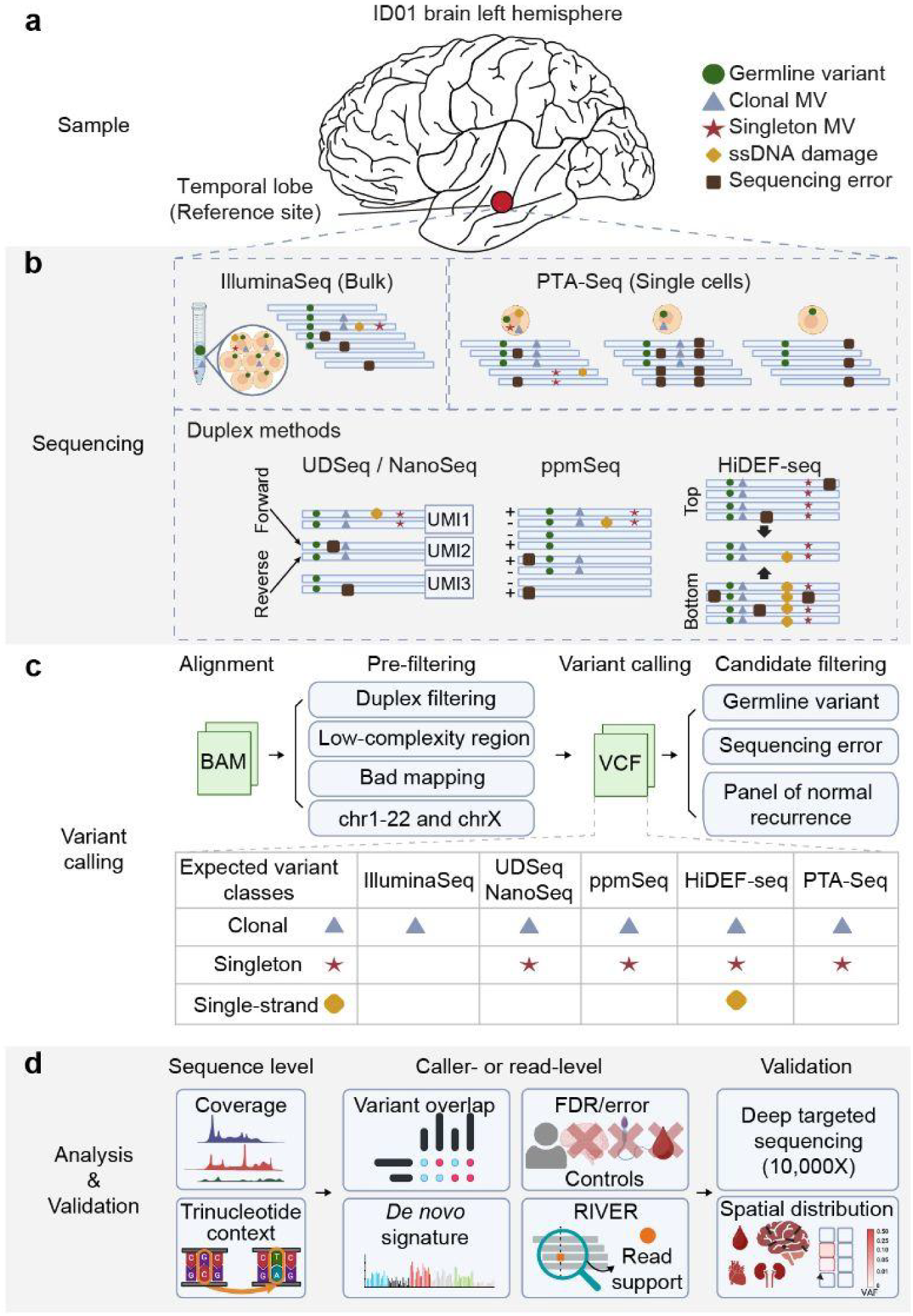
Experimental design and analytical framework for cross-method benchmarking of mosaic variants. **a,** Matched DNA and nuclei from the left temporal cortex of an aged neurotypical donor. **b,** Profiled by deep bulk Illumina whole-genome sequencing (IlluminaSeq), UDSeq, NanoSeq, ppmSeq, HiDEF-seq and single-neuron WGS after fluorescence-activated nuclei sorting (FANS) and primary template-directed amplification-based whole genome sequencing (PTA-Seq; *n* = 16 nuclei). Different sequencing methods capture distinct classes of genomic variation, including germline, clonal mosaic, singleton, single-strand, and generate platform-selective sequencing errors. **c,** Method-specific alignment, duplex filtering and variant calling were followed by harmonized filters excluding low-complexity or poorly mappable regions, germline-like calls, sequencing artifacts and variants recurrent in method-matched panels of normals (PoN). In duplex data, variants supported by multiple independent DNA molecules or one molecule were classified as clonal or singleton, respectively; HiDEF-seq additionally enabled detection of single-strand events. **d,** Benchmarking assessed coverage, trinucleotide-context representation, caller- and read-level variant support (RIVER), *de novo* mutational signatures, error and false-discovery rate estimates. Candidate variants validated by approximately 10,000× targeted amplicon sequencing across eight brain regions and three peripheral tissues, together with an unrelated negative control, to classify variants and quantify cross-tissue variant allele fractions. BAM, binary alignment map; FDR, false-discovery rate; UMI, unique molecular identifier; VAF, variant allele fraction; VCF, variant call format. RIVER, Read-level Integration for Variant Evidence Recovery.

Data were processed using method-specific pipelines, followed by harmonized filtering to remove germline-like variants, calls in error-prone genomic regions, and recurrent artifacts identified using a panel of normal samples according to best practices (**Methods**) ^14,17^. Germline SNV calls were highly concordant between IlluminaSeq and HiDEF-seq, with 99.76% and 98.17% reciprocal recovery, respectively. The final call set of duplex sequencing-derived MVs were classified as either multiread or singleton, according to whether they were supported by multiple or single DNA molecules, respectively (**Fig. 1c, Supplementary Table 1**). This annotated call set was used to assess the technical and biological properties of MVs at the sequence-, caller-, and read-levels, together with orthogonal validation and spatial distribution analysis (**Fig. 1d**).

### Sequencing methods differ in genome representation and mutational context

We first characterized the coverage overdispersion profiles (zero-inflated Poisson distribution; ZIP vs. zero-inflated negative binomial distribution; ZINB) of each duplex method (**Fig. 2a**), defined as the excess variability of genomic coverage across fixed genomic bins relative to that expected under a ZIP sampling model, quantified as the variance-to-mean ratio of coverage ^18^. NanoSeq, which uses restriction enzyme-based fragmentation, showed the highest overdispersion among the duplex sequencing methods compared with IlluminaSeq, as expected. In contrast, HiDEF-seq, UDSeq, and ppmSeq exhibited results comparable to or lower than those of IlluminaSeq. Repeating the analysis in independent samples and across sequencing depths yielded similar method-specific patterns (**Extended Data Fig. 1a**). Notably, ppmSeq maintained consistently lower overdispersion across samples and depths, whereas UDSeq showed greater variation. At equivalent mean coverage (30×), UDSeq and ppmSeq showed the most even genome-wide coverage distributions among the duplex sequencing approaches (**Fig. 2b**).

**Fig. 2.**
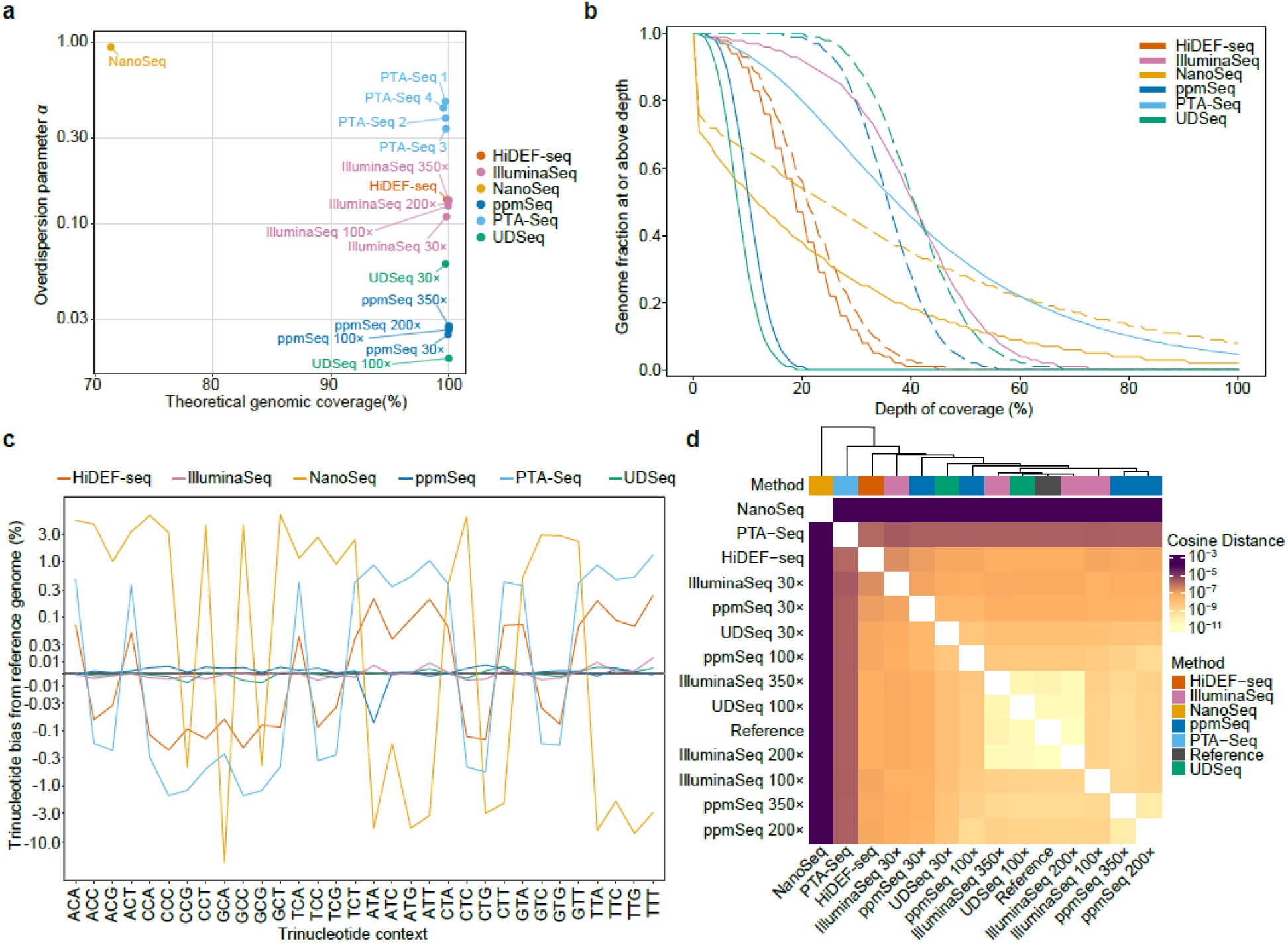
Method-specific read landscapes define the observable space for mosaic variant discovery. **a,** Genome-wide coverage breadth and read-depth overdispersion. X-axis: % GRCh38 positions with depth ≥ 1; Y-axis: zero-inflated negative-binomial (ZINB) overdispersion parameter *α*, plotted log10 scale. Larger *α* values indicate less uniform genomic sampling relative to the ideal zero-inflated Poisson distribution. Points: full-depth or down-sampled sequencing profiles. **b,** Complementary cumulative distributions of genome-wide coverage after down-sampling raw alignments to a mean 30× depth, before (solid line) and after (dash line) applying method-specific duplex filtering. Proportion of analyzed positions covered at or above indicated depth. PTA-Seq curve: mean coverage distribution across *n* = 16 single-neuron profiles. **c,** Differences in relative abundance of the 32 pyrimidine-first trinucleotide contexts between sequencing profiles and the GRCh38 reference. Difference between profile and reference proportion in percentage points on a signed pseudo-logarithmic scale; positive and negative values: enrichment and depletion, respectively. **d,** Pair-wise cosine-distance heatmap and hierarchical clustering of trinucleotide-context profiles across methods and GRCh38 reference. Cosine distance calculated as 1−cosine similarity; smaller values: greater similarity.

Trinucleotide composition analysis can reveal biases in amplification, sequencing chemistry, or read capture preferences, which can impact MV calling ^19–21^. Therefore, we next evaluated sequence-context representation across methods by comparing trinucleotide composition of the BAM files. Quantitative comparison against the reference genome showed minimal sequence-context distortion in IlluminaSeq, ppmSeq, and UDSeq, while HiDEF-seq exhibited only subtle deviations that may reflect characteristics of the PacBio chemistry (**Fig. 2c**). In contrast, NanoSeq displayed the largest deviation, likely due to its relatively shallow genome sampling, whereas PTA-Seq showed a similar bias trend to HiDEF-seq.

Datasets generated using the same sequencing technology consistently clustered together regardless of tissue source or sequencing depth (cosine similarity >99.99%), indicating that sequence-context representation is an intrinsic and highly reproducible property of each method (**Fig. 2d and Extended Data Fig. 1b**). Particularly, UDSeq and ppmSeq clustered more closely with the reference genome and IlluminaSeq than NanoSeq and PTA-Seq. Taken together, these findings indicate that duplex sequencing methods, by virtue of their unique methodologies, differ substantially in assessment of the underlying genome, a property likely to influence the spectrum of recovered MVs.

### MV callers recover distinct mosaic landscapes from the same brain

We next investigated MV discovery profiles of each duplex method. Because each sequencing technology requires distinct analytical strategies, we first optimized MV calling pipelines to maximize sensitivity while minimizing technical artifacts (**Fig. 3a and Methods**). Following method-specific MV calling and harmonized filtering, 0.01–16.68% and 4.23–26.46% of MV candidates were classified as germline-like calls and recurrent technical artifacts, respectively (**Fig. 3b**). Surprisingly, germline variants accounted for the largest fraction of calls removed in ppmSeq (16.68%), whereas other duplex sequencing datasets retained most of MV candidates, suggesting that subtraction using a germline variant call set may be particularly crucial for accurate MV profiling with ppmSeq.

**Fig. 3.**
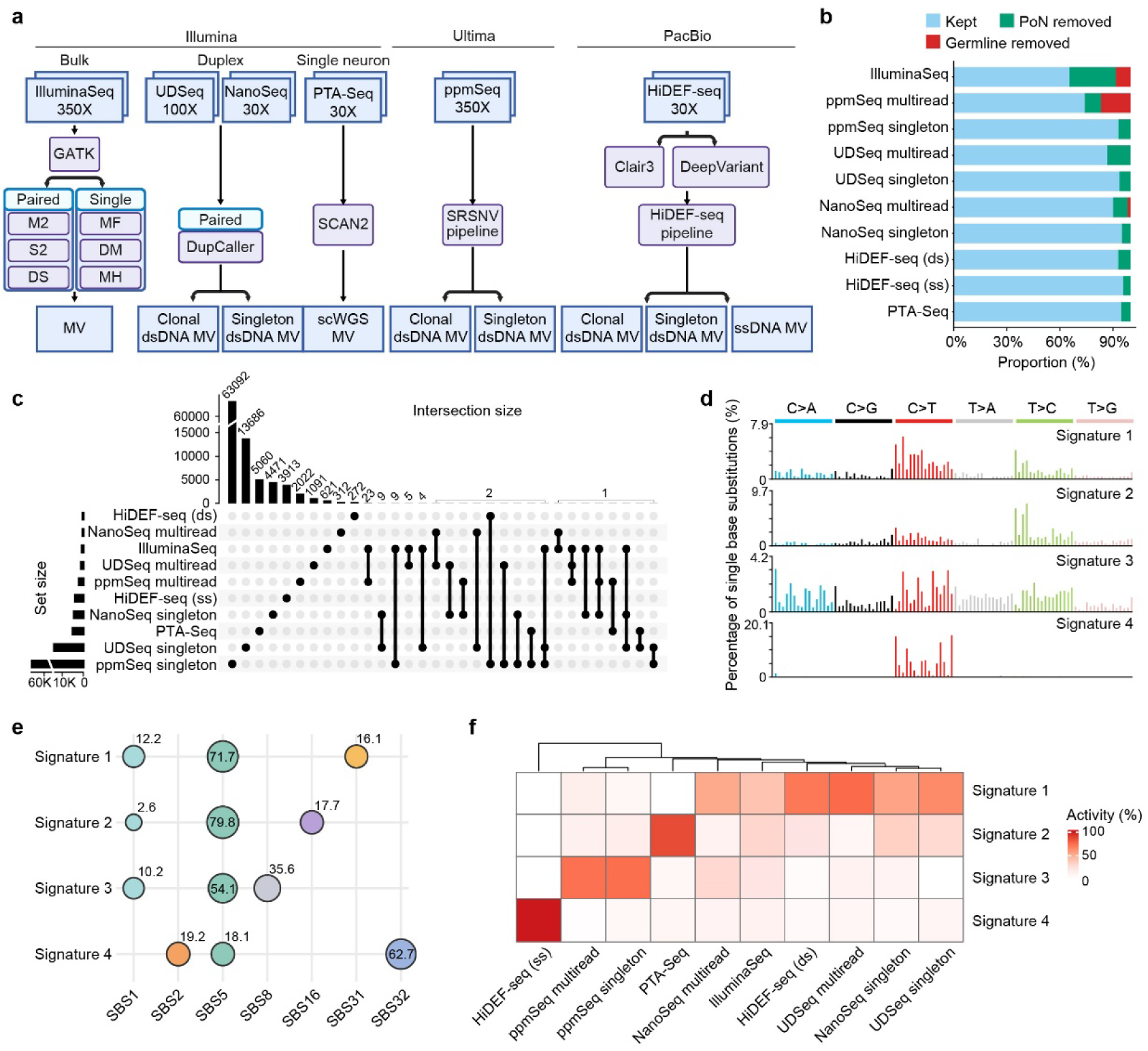
Method-specific mosaic variant calling reveals limited caller-level concordance and distinct mutational profiles. **a,** Method-specific MV calling workflows for IlluminaSeq, duplex sequencing and PTA-Seq of the left temporal cortex. IlluminaSeq was analyzed in paired- and single-sample modes, whereas duplex and single-neuron datasets were processed using the indicated method-specific callers. Method-specific outputs comprised clonal or singleton double-strand DNA (dsDNA) calls and, for HiDEF-seq, single-strand DNA (ssDNA) events. **b,** Proportions of candidates remaining after genomic prefilters that were retained, removed as germline or germline-like, or removed owing to recurrence in an unrelated PoN, normalized to the prefilter total within each call set. **c,** UpSet plot of caller-level concordance among retained full-depth SNV call sets. Horizontal bars: set sizes; vertical bars: intersection sizes; connected dots: contributing methods. **d,** Four *de novo* SBS signatures (Signatures 1-4) extracted from the combined MV call sets by non-negative matrix factorization. Bars: percentage of substitutions in each trinucleotide channel. **e,** Decomposition of *de novo* signatures into COSMIC SBS signatures. Circle size and labels: percentage contributions. Circle sizes were rescaled to a fixed display range for visibility and are therefore not directly proportional to percentages. **f,** Relative activities of the four *de novo* signatures in each MV call set. Cell values and shading: percentage contribution. Columns hierarchically clustered by signature composition. For duplex methods, multiread and singleton denote variants supported by multiple independent molecules or a single molecule, respectively. COSMIC, Catalogue of Somatic Mutations in Cancer; DM, DeepMosaic; DS, DeepSomatic; DupCaller, Duplex-based Variant Caller; GATK, Genome Analysis Toolkit; HC, high confidence; M2, Mutect2; MF, MosaicForecast; MH, MosaicHunter; S2, Strelka2; SBS, single-base substitution; SCAN2, Single Cell Analysis 2; SNV, single-nucleotide variant; SRSNV, Single Read SNV.

We next compared the numbers and types of MV candidates called by each method. At full depth, ppmSeq produced the largest multiread and singleton call sets (2,050 multiread and 63,112 MV candidates), respectively, followed by UDSeq (1,103 multiread and 13,706 singleton MV candidates) and NanoSeq (317 multiread and 4,489 singleton MV candidates). IlluminaSeq identified 669 putative multiread MV candidates, whereas HiDEF-seq yielded 274 double-strand and 3,913 single-strand calls at 30× depth. Among multiread calls, mean VAFs were relatively similar for ppmSeq, UDSeq, NanoSeq and IlluminaSeq (2.03–2.98%), whereas HiDEF-seq double-strand calls showed a somewhat higher mean VAF of 5.02%.

To assess the contribution of sequencing depth, we downsampled IlluminaSeq, ppmSeq and UDSeq to a nominal depth of 30×, to compare with native 30× NanoSeq and HiDEF-seq datasets (**Methods**). At 30×, NanoSeq yielded the largest multiread call set (317 MVs), followed by IlluminaSeq (190), UDSeq (51), and ppmSeq (30), whereas ppmSeq retained the largest singleton call set (9,010 MVs). Downsampling also shifted the retained multiread calls from IlluminaSeq, ppmSeq, and UDSeq toward higher VAFs, indicating preferential retention of higher-frequency MVs as sequencing depth decreased. Together, these results show that sequencing depth substantially influences both MV yield and the allele-frequency range accessible to multiread detection, while method-specific differences remain at comparable nominal depth.

### Little evidence of overlap between duplex sequencing MV callsets

Duplex sequencing is powered to reliably detect non-clonal MVs, potentially resulting in limited concordance among methods. To assess caller-level concordance, we compared MV call sets (**Fig. 3c**). Notably, overlap among duplex methods was limited, with the majority of MVs detected by only a single method. This limited overlap likely reflects not only differences in sequencing chemistry and genome representation, but also the method-specific variant-calling algorithms. Even the absolute number of recurrent MV calls shared across duplex methods remained lower than the number of clonal MVs reported through IlluminaSeq, indicating that cross-method recurrence at caller level is insufficient to comprehensively recover biologically meaningful clonal MVs.

To investigate the biological basis of mutational spectra of duplex sequencing methods, we extracted four *de novo* mutational signatures from the combined MV call set across all methods (**Fig. 3d**) ^22^. Comparison with established COSMIC SBS signatures showed that Signature 1 to 3 were predominantly composed of SBS1-and SBS5-like processes, likely arising from canonical clock-like mutational mechanisms associated with development and aging, whereas Signature 4 showed enrichment for distinct mutational processes associated with strand asymmetry (**Fig. 3e**).

We then quantified the contribution of each *de novo* signature to individual method-derived MV candidates (**Fig. 3f**). NanoSeq, HiDEF-seq double-strand and UDSeq were all dominated by Signature 1, while ppmSeq formed a distinct cluster characterized by enrichment of Signature 3, suggesting preferential recovery of a unique subset of MV candidates or the influence of method-specific processes. Within each duplex method, singleton and multiread MV candidates showed similar mutational signatures, suggesting that a substantial fraction of singleton MVs may share the same underlying mutational processes as multiread MVs.

Notably, HiDEF-seq single-strand MVs were dominated by the distinct Signature 4, which was enriched for strand-asymmetric SBS signatures, consistent with the strand-specific mutational processes. We further observed method-specific enrichment of MV candidates across genomic sites with distinct functional annotations (**Extended Data Fig. 2**). Together, these analyses demonstrate that differences among duplex sequencing methods arise from both method-specific MV recovery and mutational spectrum composition.

### RIVER reveals MV candidates missed by individual callers

Given the ability of duplex sequencing to detect MVs at very low VAF and the distinct MV-calling algorithms used across methods, we reasoned that MV candidates called by only one method might nevertheless be supported by reads from other methods. Evaluating cross-sample reads was proposed as a method to rescue putative MVs ^23^. We therefore comprehensively examined whether multiread-supported and singleton MVs from each method showed pileup evidence in the other sequencing datasets, within a defined statistical framework with a 5% FDR, which we call Read-level Integration for Variant Evidence Recovery (RIVER; **Fig. 4a, Extended Data Fig. 3a** and **Methods**). Notably, a subset of method-specific MV candidates was supported by reads from other methods through RIVER despite not being identified by their respective MV callers (**Fig. 4b and Extended Data Fig. 3b**). Using unrelated control datasets, we estimated method-specific false-discovery rates of 1.1–8.19% for RIVER support **(Supplementary Table 2)**. Cross-method support increased with VAF measured by IlluminaSeq, indicating that MVs with higher VAFs were more consistently recovered across independent duplex datasets through RIVER (**Fig. 4c**).

**Fig. 4.**
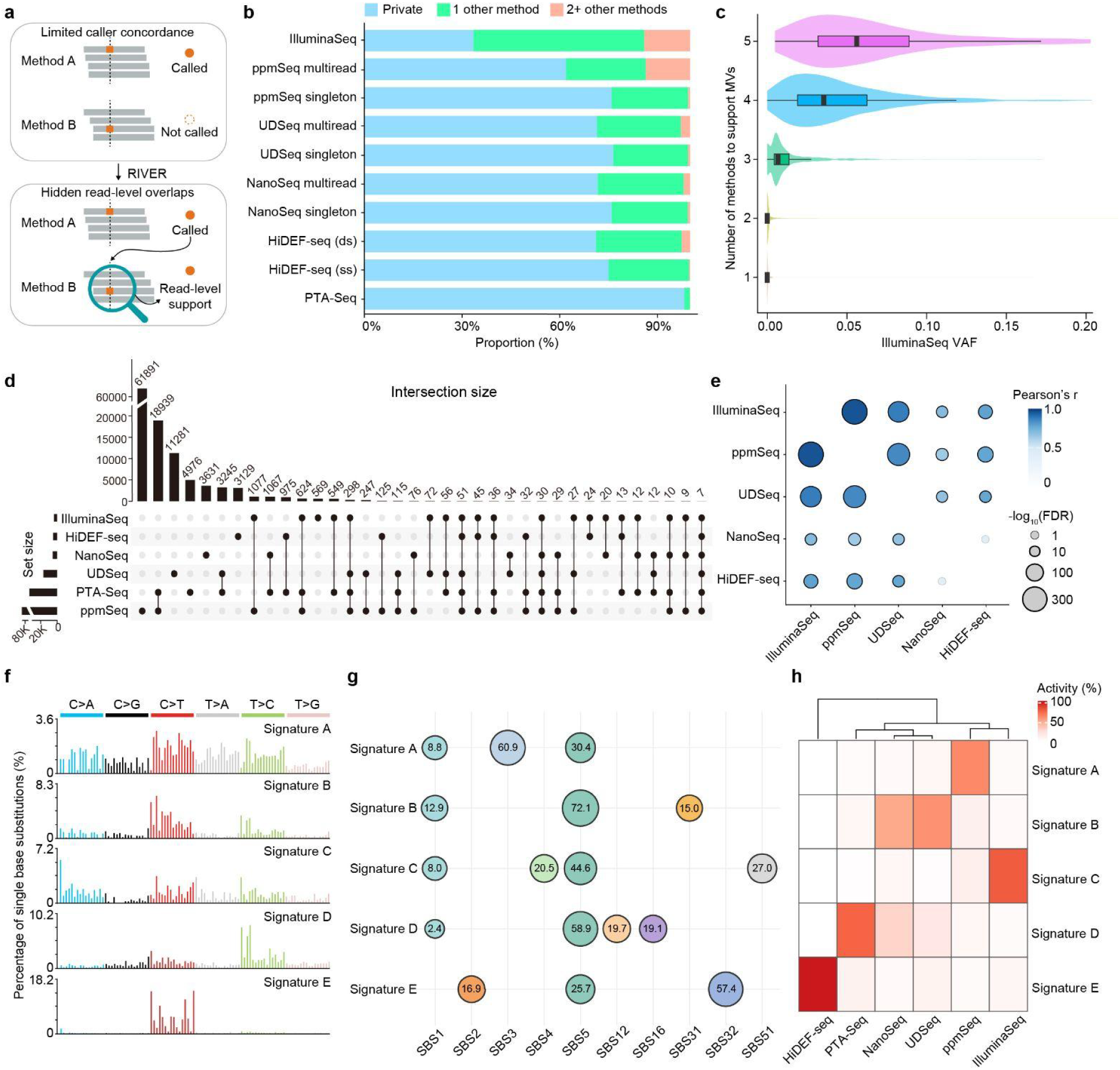
RIVER reveals shared mosaic profiles beyond caller outputs. **a,** RIVER interrogates MV candidates identified by one sequencing method to recover supporting read-level evidence missed by callers in other methods. **b,** Proportion of each retained full-depth call set with read-level support in no other method (private), one other method or ≥ 2 other methods. Rows: method and molecular-support class. **c,** IlluminaSeq VAF distributions stratified by the number of methods with read-level support. Violins: distributions; boxes: median and interquartile range. **d**, UpSet plot of MV sets defined by read-level evidence in each method. Horizontal bars: set sizes; vertical bars: intersection sizes; connected dots: contributing methods. **e,** Pairwise Pearson correlations of method-specific VAFs for MVs with read-level support in both methods. Circle color: Pearson’s *r*; size: −log10(FDR), after Benjamini–Hochberg adjustment across the 10 unique method pairs; bold outlines: FDR < 0.05. Numerically zero FDR values were floored at 10^−300^ for plotting. **f,** Five *de novo* SBS signatures (Signatures A-E) extracted from read-level-supported SNVs by non-negative matrix factorization. Bars: percentage of substitutions in each trinucleotide channel. **g,** Decomposition of the *de novo* signatures into COSMIC SBS signatures. Circle size: rescaled contribution; labels: exact percentage contributions. Circle sizes are not directly proportional to the printed percentages. **h,** Relative activities of Signature A–E across methods, normalized to 100% within each method. Color intensity: percentage activity. Columns: average-linkage hierarchical clustering of method-specific SBS96 spectra using cosine distance. For duplex methods, multiread and singleton denote variants supported by multiple independent molecules or a single molecule, respectively. RIVER, Read-level Integration for Variant Evidence Recovery.

To evaluate the performance of each method combination, we further dissected cross-method overlap after read-level rescue (**Fig. 4d**). Among comparisons, aggregated PTA-Seq showed the greatest overlap with ppmSeq and UDSeq, whereas among bulk-tissue datasets, ppmSeq and IlluminaSeq showed the highest degree of overlap. Because these two datasets were sequenced at relatively high depth, we randomly downsampled each dataset to 30× and repeated the cross-method comparison. This analysis again showed the greatest overlap between ppmSeq and IlluminaSeq (**Extended Data Fig. 4a**). Furthermore, comparison of VAFs for MV candidates recovered by RIVER revealed the strongest correlation between ppmSeq and IlluminaSeq even after accounting for sequencing depth (**Fig. 4e and Extended Data Fig. 4b**).

We further examined the mutational spectrum profiles of MVs identified through RIVER. We identified five de novo mutational signatures (Signatures A–E, **Fig. 4f**), which were subsequently decomposed into their constituent COSMIC SBS signatures (**Fig. 4g**). Read-supported MVs showed markedly different signature activities across methods, indicating strong method-specific mutational characteristics (**Fig. 4h**). These results suggest that RIVER expands MV discovery beyond caller-defined overlap, but also captures distinct subsets of MVs depending on the sequencing method.

### Orthogonal validation reveals a large pool of *bona fide* low-frequency clones

To determine which MV candidates are the most reliable and informative for clonal distribution analysis, we first established an evidence-based tiering strategy according to the level of support for each MV (**Fig. 5a**). Tier1 MV candidates were supported at the caller level by multiple sequencing methods, Tier2 MV candidates were called by a single method but supported at the read level by at least one additional sequencing method, and Tier3 MV candidates were detected exclusively by a single sequencing method. We further annotated each MV candidate according to whether it was represented by multiple independent reads or a singleton read. We then designed a panel targeting 465 MV candidates and performed ultra-deep amplicon sequencing (∼10,000x) for orthogonal validation (**Extended Data Fig. 5a and Supplementary Table 3**). Validation included the reference site used for MV discovery and multiple anatomically distributed brain regions from the same donor, enabling assessment of whether validated MVs exhibited spatial patterns consistent with developmental clonal distributions previously reported ^15,16^.

**Fig. 5.**
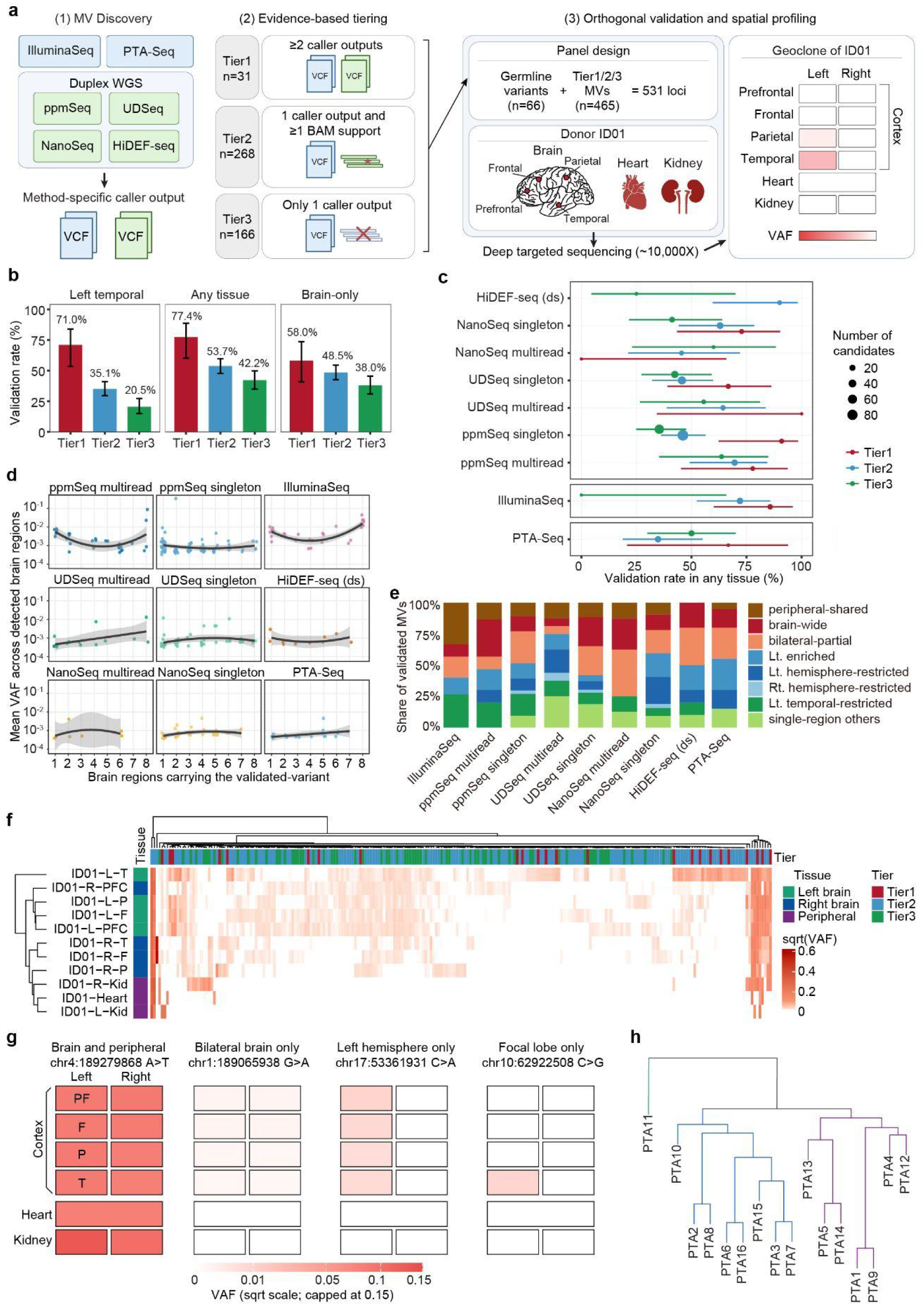
Orthogonal validation reveals method-specific detection and spatial patterns of mosaic variants. **a,** MV discovery, evidence-based tiering and orthogonal validation. Tier1: ≥ 2 caller outputs (n = 31); Tier2: one caller plus ≥ 1 cross-method BAM support (n = 268); Tier3: one caller without cross-method BAM support (n = 166). Target panel: 465 MVs and 66 germline variants (531 loci), sequenced at approximately 10,000× across eight brain and three peripheral tissues. **b,** Validation rates by tier in the reference site, any tissue and brain only (no peripheral-tissue detection). Bars: rates; error bars: Wilson 95% CIs; n: validated MVs. **c,** Validation rates in any tissue across tiers. Points: rates; horizontal lines: Wilson 95% CIs; size: tested candidates; color: tier. **d,** Spatial extent versus VAF for validated MVs. X-axis: number of brain regions carrying the variant; Y-axis: mean VAF across positive brain regions, plotted on a log10 scale. Points: MVs; curves and shading: fitted trends and confidence bands. **e,** Spatial classes of validated MVs. Hierarchical definitions: peripheral-shared, detected in any peripheral tissue; brain-wide, otherwise brain-restricted, bilateral and detected in ≥ 6 of 8 brain regions; left-enriched, other bilateral variants detected in ≥ 2 more left- than right-hemisphere regions; bilateral-partial, remaining bilateral variants; left- or right-hemisphere-restricted, detected in ≥ 2 regions of one hemisphere only; left-temporal-restricted or other single-region, detected in one brain region. Bars: proportions within call sets. **f,** VAF heatmap across eight brain and three peripheral tissues. Rows: tissues; columns: validated MVs; color: square-root-transformed VAF. Dendrograms: hierarchical clustering of tissues and variants by VAF profiles. **g,** Representative geoclones of four validated MVs showing brain-and-peripheral (Tier2), bilateral brain-only (Tier2), left brain-only (Tier2) and focal brain-only (Tier1) distributions. Rows: prefrontal (PF), frontal (F), parietal (P), and temporal (T) cortex, heart, and kidneys; columns: left and right for paired tissues. Color intensity: VAF on square-root scale, capped at 0.15; white: variant not detected by targeted amplicon sequencing. **h,** Jaccard-distance hierarchical clustering of n = 16 single-neuron PTA-Seq profiles in the left temporal lobe of ID01 using binary support for Tier2 ppmSeq singleton MV candidates. Shorter branches indicate more similar MV profiles. CI, confidence interval.

Of the targeted MVs, 32.2% (150/465) were positively validated at the reference discovery site (**Methods and Fig. 5b**). When we expanded the validation criteria to include MVs detected in any tissue from the same donor, validation rates increased to 51.2% (238/465), due to the significantly increased recovery of Tier2 and 3 MVs. This finding indicates that MVs not validated at the original discovery site may nevertheless represent *bona fide* MVs in other anatomical regions, reflecting spatially restricted or uneven clonal distributions. We therefore defined validated MVs as those detected in at least one tissue from the same donor. With the exception of NanoSeq multiread and PTA-Seq MV candidates, validation rates generally decreased from Tier 1 to Tier 3 (**Fig. 5c**). Thus, independent caller-or read-level support from an additional sequencing method substantially increased the likelihood that an MV candidate could be orthogonally validated. Further stratification within each tier revealed that validation rates varied with the combination of supporting sequencing methods and whether the MV was supported by multiple reads or a single read (**Extended Data Fig. 5b**).

Notably, Tier3 MVs still showed validation rates of approximately 30–50%, depending on the sequencing method (**Fig. 5c**). Thus, even MVs detected exclusively by a single duplex sequencing method contained a substantial fraction of *bona fide* clonal MVs. Their recovery by deep targeted sequencing indicates that at least a subset of these apparently singleton observations represent ultra-low-VAF clonal MVs rather than private ones confined to individual cells. These results expand the range of duplex MVs that can be considered for clonal analysis and demonstrate that stringent reliance on cross-method recurrence would discard a substantial fraction of genuine ultra-low-VAF clonal MVs.

### Validated MVs resolve spatial and single-cell lineage relationships

We next examined the spatial distributions of orthogonally validated MVs across the brain. Except for NanoSeq, multiread MVs generally showed increasing mean VAF as they were detected across a greater number of brain regions (**Fig. 5d**). Notably, IlluminaSeq, HiDEF-seq and ppmSeq also captured locally enriched MVs with relatively high VAFs despite being restricted to a single brain region.

Classification of validated MVs according to their spatial distributions revealed that the relative representation of clonal distribution patterns varied across sequencing methods, suggesting that different duplex methods capture distinct components of the clonal landscape (**Fig. 5e and Supplementary Data 1**). Given the well-established hemispheric restriction of developmental clones in the human brain ^15,16^ we next asked whether the validated MVs recapitulated this known developmental organization. Clustering of their VAFs across bilateral brain regions revealed a clear separation between left-and right-hemisphere tissues, demonstrating that validated MVs retain spatial patterns consistent with developmental clonal architecture (**Fig. 5f and g**).

Identifying shared low-VAF MVs across individual cells remains challenging. Therefore, we finally asked whether MVs missed by conventional single-cell calling could nevertheless be recovered directly from PTA-Seq reads. Among ppmSeq singleton MV calls, 3,633 MV candidates were shared by two or more neurons, including 2,609 shared by two neurons and 1,024 shared by three or more neurons, showing two major lineages of neurons (**Fig. 5h**). These results demonstrate that duplex MVs rescued through RIVER may recover lineage information already present in single-cell sequencing reads but missed by conventional MV calling.

## Discussion

MV discovery is shaped not only by the rarity of the variants, but also by how sequencing technologies sample DNA molecules, represent the genome, and translate molecular observations into MV calls. By benchmarking four duplex sequencing approaches alongside deep bulk whole-genome and single-neuron sequencing in matched human brain tissue, we found that caller-level concordance captures only a fraction of the mosaic evidence shared across datasets. Direct interrogation of aligned reads revealed extensive cross-method support for MVs called by only one method, indicating that much of the apparent method specificity reflects algorithmic non-detection rather than an absence of molecular evidence.

RIVER addresses this gap by testing MV candidates for statistically defined read-level evidence across independently generated datasets. This strategy expanded the set of MVs supported by more than one method by approximately two orders of magnitude compared with MVs independently called by multiple methods. The recovered evidence was biologically meaningful: approximately half of the RIVER-rescued candidates assessed by ∼10,000× targeted sequencing was detected in at least one tissue from the donor. The validation rate remained approximately 50% even when the analysis was restricted to MV candidates initially called as singletons by one method, but for which RIVER recovered additional evidence in another method. Thus, evidence that falls below the reporting threshold of an individual caller can still contribute to the reliable identification of clonal MVs when evaluated across complementary datasets.

A practical benefit of RIVER is that a focused duplex experiment could increase the information obtainable from sequencing datasets. Previous studies used deep whole-genome sequencing and targeted validation across multiple anatomical sites to establish sets of hundreds of informative MVs for clonal analysis ^2,24^. Our results suggest that even duplex sequencing of a single tissue region can provide a much larger pool of MV candidates whose supporting evidence can be recovered from existing datasets, with approximately half of the rescued candidates tested and independently validated. These MV candidates discovered at one site could be queried in existing sequencing data from other tissues to map clonal distributions. The same strategy could connect bulk duplex sequencing with existing single-cell datasets, recovering potential lineage markers missed by individual callers ^20^. Once validated, these additional MVs could improve spatial and single-cell lineage analyses without requiring duplex sequencing of every sample.

Current duplex sequencing methods are complementary rather than interchangeable, recovering broadly similar clock-like mutational signatures while differing in genome representation, mutational profiles, and MV recovery ^25^. ppmSeq showed the greatest sharing of read-level evidence with IlluminaSeq and the strongest cross-method VAF correlation, despite differences in library design and sequencing technology. HiDEF-seq, by contrast, produced call sets with limited caller-level agreement with other methods but the highest proportion of Tier 2 candidates, indicating that many of its calls had supporting evidence elsewhere that did not meet other callers’ thresholds. Its single-strand calls were also distinguished by a profile consistent with DNA mismatches or damage^10^. Together, these findings show that method-specific sampling and calling criteria can create apparent disagreement despite compatible molecular evidence, emphasizing the need to evaluate both final call sets and subthreshold read support when benchmarking duplex sequencing methods.

The tier framework translates caller- and read-level evidence into a practical hierarchy. Tier 1 MVs, independently called by multiple methods, provide the strongest support for downstream analysis. Tier 2 MVs, called by one method but supported by reads from another, broaden the discovery set while showing higher validation rates than method-private calls. Although Tier 3 MVs lack cross-method support, 30–50% were validated, depending on the discovery method, likely reflecting ultra-low allele fractions, spatially restricted clones, or incomplete molecular sampling. This hierarchy can guide follow-up studies: Tier 1 and Tier 2 MVs can be prioritized for targeted sequencing, spatial mapping, and single-cell genotyping, whereas Tier 3 MVs can be evaluated using allele fraction, genomic context, recurrence, and biological relevance.

Our study has some limitations. First, our analyses focused on SNVs, and the complete cross-method comparison and spatial validation were performed in a single healthy donor. However, this donor was representative of neurotypical individuals ^15,16^ and the matched design enabled direct comparison within a common biological background. Second, although pileup review has been used in the MV field ^14,23,26^ application to duplex sequencing has not been widely evaluated. Unrelated controls for all methods except HiDEF-seq provided estimates of background false-positive rates for read-level support. Finally, although only a subset of MV candidates was assessed by targeted amplicon sequencing, selection across evidence tiers and sequencing methods enabled evaluation of candidates with varying levels of support. Future studies should extend these findings to additional individuals and variant classes.

Together, these findings position read-level integration as a means of extracting additional biological information from complementary sequencing datasets. RIVER does not replace method-specific callers or orthogonal validation; rather, it connects their outputs by distinguishing the absence of a reported call from the absence of molecular evidence. As duplex, spatial, and single-cell sequencing datasets become increasingly available, this framework could expand the number of informative MVs available for studying somatic mosaicism, developmental lineage relationships, and clonal dynamics in human tissues.

## Supporting information

supplementary_table_1_bioRxiv.xlsx

supplementary_table_2_bioRxiv.xlsx

supplementary_table_3_bioRxiv.xlsx

supplementary_data_1_bioRxiv.pdf

## Acknowledgements

Authors thank Gilad D. Evrony lab for advice on HiDEF-seq protocols, Dan-Avi Landau and Alexandre Pellan Cheng for advice on ppmSeq, BioSkryb Genomics, Inc for advice on PTA methods and Rady Genomics Institute for sequencing support. We thank Stem Cell Genomics Core at SCRM for sequencing services. This research was supported by the Korean Ministry of Science and Information and Communication Technology (RS-2024-00411768, RS-2026-25470079, RS-2024-00407383, RS-2026-25501101 to C.C.); InnoCORE program of the Ministry of Science and ICT (N10260098 to C.C.); KAIST Creative & Challenging Project (N11260025 to C.C.); Medical Scientist Training Program from the Korean Ministry of Science and Information and Communication Technology (to D.H.C); Center for Genomic Medicine and the Lawrence T. & Janet T. Dee Foundation Innovation Award (18638 to X.Y.), Research to Prevent Blindness Stein Innovation Award (51007425 to X.Y.), Collaborative Research Catalyst Award (43719 to X.Y.), Margolis Foundation Transformational Research Project (to X.Y.); Larry L. Hillbloom Foundation (313915 to J.G.G.), the Korea-US Grant (to J.G.G.), and CIRM Grant (EDUC4-12804, to P.S.-O.). We used AI tools to polish language in early drafts.

## Contributions

J.W., D.H.C., C.C. and J.G.G. designed the study. J.C., P.S.O., Y.C. organized, handled and sequenced human samples. J.W., D.H.C., R.H., Y.X., X.H., and P.J.W. performed the bioinformatics and data analyses for RIVER. D.H.C., C.C., S.J., J.R., Y.N., S.L. and Y.S.J. established ppmSeq variant calling pipeline. X.Y. advised benchmarking analysis and deep targeted sequencing experiment design. Y.C. and L.B.A. analyzed UDSeq datasets.

## Conflict of interest

S.L. is an employee of Inocras Inc., and Y.S.J. is a co-founder of the company. X.Y. is an advisory board member and shareholder of Paterna Biosciences, Inc. All other authors declare no competing interests.

## Online Methods

### Sample preparation

For donor ID01, we used genomic DNA extracted in our previous study, briefly, a neurotypical human whole brain from a 70-year-old female ^16^. Sperm samples were obtained from a healthy 19-year-old male enrolled under University of California Institutional Review Board (IRB) protocols approved for blood, saliva, and semen collection (protocol nos. 140028 and 161115) ^27^. Genomic DNA was extracted using the DNeasy Blood & Tissue Kit (QIAGEN, cat. no. 69506) according to the manufacturer’s instructions.

### Deep targeted sequencing

A customized AmpliSeq Custom DNA Panel for Illumina (20020495, Illumina) was used for ID01 (Illumina Design Studio Solution ID: IAA40112_166). The list of candidate MVs from the mosaic variant detection pipeline in ID01 described above was subjected to the AmpliSeq design system. We randomly selected 45 high-confidence heterozygous variants as positive controls. These heterozygous variants presented with estimated allele fractions between 48% and 52% for all 11 sequenced bulk tissues, and with read depths between 270 and 330×. We also randomly selected 21 reference homozygous variants as negative controls. These reference homozygous variants presented with approximately 0% allele fraction across all sequenced samples, with an average depth of 270–330×, and a gnomAD (v2.0.1) allele frequency of more than 0.5 to exclude any potential contamination or amplification bias. DNA from extracted tissue, amplified single nuclei, and a duplicate unrelated control sample was diluted to 5 ng µl−1 in low-TE provided in the AmpliSeq Library PLUS (384 reactions) kit (20019103, Illumina). AmpliSeq was carried out following the manufacturer’s protocol (document 1000000036408v07). For amplification in bulk samples, 18 cycles each for 4min were used. After amplification and fragmentase universal primer assay (FUPA) treatment, libraries were barcoded with AmpliSeq CD Indexes (20031676, Illumina) and pooled with similar molecular numbers based on measurements made with a Qubit dsDNA High Sensitivity kit (Q32854, Thermo Fisher Scientific).

### Data analysis for deep targeted sequencing

Raw reads from deep targeted sequencing were mapped to the GRCh38 genome with BWA’s (v.0.7.17-r1188; -t 1 -M) mem command. Alignments were coordinate-sorted and mate information was corrected using Picard (v.2.18.27) SortSam and FixMateInformation, respectively, with SORT_ORDER=coordinate, VALIDATION_STRINGENCY=LENIENT, CREATE_INDEX=true and MAX_RECORDS_IN_RAM=2500000. BAM files were processed without removing PCR duplicates. Reads near indels were realigned with GATK’s (v.3.8) RealignerTargetCreator and IndelRealigner, using the GRCh38 Mills and 1000 Genomes gold-standard indels and assembly38 known-indel resource. The final BAM files were parsed by SAMtools’s (v.1.18) mpileup and the 95% binomial confidence intervals of the measured allele fraction of all the candidate mosaic variants, together with the homozygous (negative control) and heterozygous (positive control) variants, were estimated based on a Wilson score estimation (https://github.com/shishenyxx/Human_Inhibitory_Neurons). Following depth calculation, regions of 465 mosaic candidates, 45 heterozygous variants (positive controls), and 21 homozygous variants (negative controls) were detected and subjected to the next genotyping steps. The genotypes of candidate mosaic variants from deep targeted sequencing were determined by comparing them to the allele fraction distribution of the reference homozygous and heterozygous variants. The lower bounds of the 75% Wilson score confidence intervals of all reference homozygous variants with more than 30 read depth were estimated, and the lower-bound threshold derived from this distribution was 0.0001859. The distribution of the upper bounds of the 95% Wilson score confidence intervals of all heterozygous variants was calculated, and 0.4744415 was the threshold for the upper bound based on an approximately 25% false discovery rate based on the built-in heterozygous and alternative homozygous genomic positions. Mosaic candidates from WGS were considered positive if variants met the following criteria at the same DNA samples: (1) the lower bound of the 75% Wilson score confidence interval was more than 0.0001859, (2) the lower bound of the 75% Wilson score confidence interval in the unrelated control sample was less than 0.0001859, (3) the upper bound of the 95% Wilson score confidence interval was less than 0.4744415, (4) the sequencing depth was more than 30, and (5) variants were positively detected in any of tissue sequenced. These criteria ensured that the false discovery rate for each variant was under 5%. After quantification, 238 MV candidates were considered positively validated in the sample where the variant was detected in any tissue within ID01. The total validation rate was 51.2% (238 of 465).

### Illumina bulk WGS processing and MV calling

Paired-end Illumina WGS reads from the ID01 brain and matched-normal specimens were aligned to the GRCh38 full analysis set plus decoy and HLA reference using BWA-MEM (v.0.7.18; ‘-M -’ for the brain and ‘-K 100000000 -Y’ for the matched normal), coordinate-sorted using SAMtools (v.1.21) and Sambamba (v.0.7.0), and duplicate-marked using Sambamba. Base quality scores were recalibrated using GATK BaseRecalibrator and ApplyBQSR (v.4.6.1.0) with dbSNP build 146, GATK known indels and Mills and 1000 Genomes gold-standard indels. Germline SNVs and indels were called from the matched-normal BAM using GATK HaplotypeCaller, recalibrated separately by VQSR, with SNVs filtered at 99.5% truth sensitivity, and merged using BCFtools (v.1.17 and v.1.21).

Mosaic SNVs and small indels were called from the brain WGS data. Five candidate sets were generated: final-passed MosaicHunter single-sample calls (multithreaded version v.1.0.0; posterior mosaic probability >0.05); PASS tumour-only Mutect2 calls (GATK v.4.6.2.0) classified as mosaic by DeepMosaic or MosaicForecast using pretrained EfficientNet-B4 and 200× random-forest models, respectively; biallelic PASS DeepSomatic calls (v.1.9.0) in WGS tumor–normal mode; and the intersection of biallelic PASS calls from paired Mutect2 and Strelka2 (v.2.9.10). Tumour–normal analyses used the matched-normal BAM downsampled to approximately 50×.

DeepSomatic panel-of-normals filtering was disabled. Call sets were merged by exact chromosome, position, reference and alternate alleles. The paired Mutect2– Strelka2 intersection contributed one vote and each other workflow one vote; variants supported by at least two votes formed the strict consensus set. Variants were excluded if they overlapped RepeatMasker or segmental-duplication regions, occurred within homopolymer or dinucleotide repeats of at least four units, lay within 5 bp of a germline indel, had a gnomAD allele frequency >0.001, or had a lower 95% brain allele-fraction bound <0.001. Candidates were annotated using ANNOVAR and BEDTools. Brain and matched-normal allele counts were obtained using SAMtools mpileup, and allele fractions and confidence intervals were calculated as described under RIVER. Scripts for variant calling and filtering are provided on GitHub.

### NanoSeq and UDSeq library preparation

NanoSeq 2021 protocol was followed when generating the library as described ^8^. The complete step-by-step UDSeq protocol is provided in the preprint ^12^. Briefly, to minimize DNA damage during fragmentation, intact genomic DNA was enzymatically fragmented using NEBNext dsDNA Fragmentase (M0348S) or UltraShear (M7634L) to achieve an average fragment size of ∼350 bp. For human sperm, DNA was fragmented for 15 minutes, while human brain tissue required 20 minutes. Fragmented DNA was then used for UMI adapter ligation with the xGen™ cfDNA & FFPE DNA Library Preparation Kit. All steps were carried out on magnetic beads to reduce DNA loss during purification, thereby improving library conversion efficiency. In the final step, an appropriate femtomole input amount was used for PCR amplification to incorporate sample index sequences compatible with Illumina® sequencing platforms.

### NanoSeq and UDSeq data processing and somatic variant calling

Paired-end full-depth NanoSeq and UDSeq reads from the left cerebral cortex were analyzed using the corresponding liver and left kidney samples, respectively, as matched normals. Duplex molecular barcodes were retained in read names and the DB SAM tag. Reads were aligned to the GDC GRCh38.d1.vd1 reference using BWA-MEM (v.0.7.17) with -C -T 0, coordinate-sorted with SAMtools (v.1.19.2) and duplicate-marked with GATK MarkDuplicates (v.4.3.0.0) using --BARCODE_TAG DB, -- DUPLEX_UMI true and --TAGGING_POLICY OpticalOnly. Somatic SNVs and short indels were called with DupCaller (v.1.0.5; commit 918572d), which learned sample-specific error profiles from each cortical dataset. Calling used a germline VCF generated from matched-normal Illumina WGS with GATK HaplotypeCaller and ApplyVQSR (v.4.6.1.0) and a merged noise mask comprising RepeatMasker, segmental-duplication, and tandem-repeat regions; all other calling and filtering parameters were left at default values.

### HiDEF-seq library preparation

The HiDEF-seq library was generated by following the protocol previously described^10^. Input DNA amounts of 500 ng were used per library. With high-quality DNA, input amounts of 500 ng provide sufficient HiDEF-seq library yield for approximately one full Pacific Biosciences Revio instrument sequencing run. SMRT sequencing was performed without barcoded multiplexing (Sample is indexed: FALSE). Data was captured over a 30-hour movie acquisition time to maximize subread yield. For secondary analysis, circular consensus sequences were generated using the strand consensus mode to preserve strand-specific information. To ensure maximum data fidelity for downstream analysis, full-resolution base quality scores were retained (Full Resolution Base Qual: TRUE), and raw subreads were aligned and compiled against the final HiFi reads (Subread To HiFi Pileup: TRUE).

### HiDEF-seq data processing and somatic SNV detection

Full-depth PacBio Revio sequencing data from the left temporal cortex of donor ID01 were analyzed using the HiDEF-seq v.1.2 pipeline (‘gevrony/hidef-seq:1.2’). Strand-specific consensus reads were generated with CCS (v.8.2.0) in by-strand mode, demultiplexed with Lima (v.2.12.0), paired by zero-mode waveguide identifier, and aligned to the CHM13 draft v.1.0 reference using pbmm2 (v.1.16.0) with the CCS preset; alignments were coordinate-sorted and indexed using SAMtools (v.1.19).

Identical substitutions detected in both opposing strand consensus sequences were classified as double-stranded DNA (dsDNA) mutations, whereas substitutions confined to one strand were classified as single-stranded DNA (ssDNA) mismatch or damage calls. Candidate sites were filtered against germline variants using DeepVariant (v.1.2.0) and GATK HaplotypeCaller (v.4.6.1.0), and were subjected to the two standard HiDEF-seq filter sets with otherwise default parameters. Only SNVs on chromosomes 1–22 and X were retained for downstream analyses.

### ppmSeq library preparation and sequencing

ppmSeq library preparation and sequencing were performed at Inocras Inc. (San Diego, CA, USA). gDNA concentration was measured fluorometrically on a Qubit fluorometer (Thermo Fisher Scientific, Waltham, MA, USA). For each sample, 100 ng of gDNA was diluted to a final volume of 26 µL in 1× TE buffer (10 mM Tris-HCl pH 8.0, 1 mM EDTA). Because enzymatic fragmentation is sensitive to buffer composition and to contaminants carried over from DNA extraction, gDNA was diluted only in 1× TE, and samples supplied in other buffers were buffer-exchanged or purified before fragmentation where necessary. Sequencing libraries were prepared with the NEBNext UltraShear FFPE DNA Library Prep Kit (New England Biolabs, Ipswich, MA, USA) and UG ppmSeq™ Adapters v3 (Ultima Genomics, Fremont, CA, USA), following the Ultima Genomics ppmSeq workflow. In the ppmSeq assay, the adapter design allows the plus and minus strands of each original double-stranded DNA molecule to be clonally amplified together on the same sequencing bead, so that agreement between the two strands can be assessed for every read, enabling duplex-level suppression of sequencing and library-preparation errors. gDNA was enzymatically fragmented with NEBNext UltraShear (45 °C for 25 min, then 65 °C for 15 min), targeting a fragment size of 250–270 bp. Fragmented DNA was end-repaired and dA-tailed with the NEBNext Ultra II End Prep Enzyme Mix in the presence of DTT (20 °C for 30 min, then 65 °C for 30 min), and ligated to indexed UG ppmSeq v3 adapters using the NEBNext Ultra II Ligation Master Mix and Ligation Enhancer (20 °C for 60 min). Ligation products were purified with 0.7× magnetic beads (NGS Magnetic Beads; Bulldog Bio, Portsmouth, NH, USA) and then size-selected by a double-sided bead cleanup (0.65× followed by a cumulative 1.0× bead ratio), with two 80% ethanol washes at each step. Libraries were not PCR-amplified, so each library molecule retained both strands of the original genomic DNA duplex. Library concentration was measured by qPCR with the NEBNext Library Quant Kit for Ultima Genomics (New England Biolabs) on a QuantStudio™ 5 Real-Time PCR System (Thermo Fisher Scientific). Libraries were diluted 2,500-fold in NEBNext Library Dilution Buffer and assayed in duplicate against a five-point standard curve, together with a no-template control. The fragment-size distribution of the libraries was assessed on a 4200 TapeStation System using D1000 ScreenTape (Agilent Technologies, Santa Clara, CA, USA). Molar library concentrations were calculated from the qPCR results assuming a standard fragment size of 300 bp. Libraries were then pooled in equimolar amounts and diluted with 10 mM Tris-HCl to a final pool concentration. Sequencing templates were prepared using the Ultima Genomics Solaris™ Free workflow, with pools processed in pairs. Amplification reactions were assembled on an Opentrons OT-2 liquid handler (Opentrons, New York, NY, USA) and emulsified on a Thermomixer C (Eppendorf, Hamburg, Germany). Library molecules were then clonally amplified on sequencing beads by emulsion PCR on the UG Amplification Station (Ultima Genomics), followed by emulsion breaking and enrichment of template-bearing beads. Enriched template beads were sequenced on the Ultima Genomics UG 100™ sequencer (Ultima Genomics, Fremont, CA, USA). Base calling was performed by the on-instrument Ultima Genomics software, and reads were assigned to samples according to their ppmSeq adapter index using a sample sheet generated for each pool.

### ppmSeq data processing and somatic SNV discovery

Full-depth ppmSeq data from the left temporal cortex of donor ID01 were processed using a custom somatic SNV discovery framework developed downstream of the Ultima Genomics Single Read SNV (SRSNV) pipeline (v.1.22.1) ^28^. Candidate substitutions were extracted from aligned CRAMs with GATK FlowFeatureMapper, requiring a mapping quality of 60 and reference identity over the five bases flanking each substitution to keep compound homopolymer errors from being called as SNVs, while retaining flow-space, sequence-context, fragment-level and ppmSeq adapter features. SRSNV then trained an XGBoost classifier to separate true-like events (homozygous substitutions with variant allele fraction ≥0.7 at ≥20× coverage) from false-like events (substitutions supported by a single read at otherwise reference-concordant loci, enriched for sequencing artifacts), using features such as base-calling quality, local sequence context, cycle-skip status and ppmSeq adapter categories. Its output was calibrated into a PHRED-scaled SNVQ estimating the residual error rate of each read-level substitution. Only candidates passing SRSNV quality filtering (PASS; SNVQ ≥40) were retained. Because the ppmSeq start- and end-adapter signals report whether both strands of the original DNA molecule contributed to a read, we further required MIXED annotations at both ends, restricting the analysis to double-stranded-derived reads. Germline variants called from matched peripheral-tissue Illumina WGS were removed, together with candidates overlapping homopolymers ≥7 bp, simple tandem repeats, recurrent ppmSeq panel-of-normal sites and gnomAD variants with a minor allele frequency >0.001.

Ultima flow-based errors depend on sequence context, so candidates were then denoised with trinucleotide-context-specific thresholds. To account for strand bias, we evaluated all 192 substitution contexts in their sequenced orientation, treating purine- and pyrimidine-centered contexts separately rather than collapsing them into the conventional 96 pyrimidine-referenced classes ^11^. For each context, we applied the quality threshold maximizing Youden’s J in ROC analysis of true versus false events, with a minimum of 12.

Surviving read-level substitutions were reconstructed into site-resolved calls by grouping identical chromosome, position, reference and alternate alleles, discarding multiallelic sites at which more than one alternate allele survived filtering. Tracing each remaining locus back to the unfiltered SRSNV output, we quantified denoised duplex support, total raw evidence, SNVQ ≥40 support and competing alternate alleles in the raw data. This trace-back defines a tiered call set: variants with at least two independent duplex reads and no competing allele are multiread SNVs and variants observed exactly once at every level, with a single denoised read, raw record and SNVQ ≥40 read, are high-confidence singletons. Rather than discarding single-read evidence, this reconstruction converts SRSNV read-level candidates into explicit multiread and singleton somatic dsSNV call sets for cross-method benchmarking, recovering as many accurate variants as possible while keeping singletons confirmable by orthogonal sequencing.

### Brain Sample Sorting and PTA

Frozen, non-fixed brain tissue from the left temporal cortex of ID01 was homogenized in 1 ml ice-cold NIB (0.25M sucrose, 25 mM KCl, 5mM MgCl2, 10 mM Tris pH 7.5, 100 mM DTT, and 0.1% Triton X-100) and dounced five times in a 2 ml Wheaton Dounce Tissue Grinder (DWK Life Sciences). The homogenate was incubated on a rocker for 5 minutes at 4°C and centrifuged at 1000×g using the ‘soft’ setting in a swinging bucket centrifuge. The supernatant was removed, and the pellet was resuspended in 0.5 ml sorting buffer and filtered through a 70 um strainer. Pellets were stained for 30 minutes using NeuN Alexa Fluor 488 (1:2,500; Millipore Sigma, MAB377). After washing, nuclei were stained with 0.5 μg/ml DAPI. DAPI+/NeuN+ and DAPI+/NeuN-nuclei were sorted on a BD InFlux Cytometer (Becton-Dickinson) into a 96-well plate pre-filled with PBS. The 96-well plate with single nuclei in each well was quickly spun down and stored at - 80°C until further processing for PTA. Single-nucleus FANS data were visualized using FlowJo software. After sorting, single-nucleus nuclei were treated for 20 min at room temperature for lysis with lysis reagent. Single-cell whole-genome amplification kit (ResolveDNA, BioSkryb), 1 ng, 100 pg, and 10 pg DNA controls were diluted based on Qubit dsDNA HS (Cat Q32854, Thermofisher). After adding the reagent mix, PTA reactions were carried out for 10 hours at 30°C, 3 min at 65°C, and held at 4°C with a lid temperature of 70°C. After bead cleanup with beads and reagents from the kit, libraries were prepared following the ResolveDNA Library Prep Kit Protocol (BioSkryb). IDT for Illumina TruSeq UD Indexes (20020590, Illumina) were used to distinguish amplified single Neuron nuclei. Library concentrations were further determined by Qubit (Q33226 Invitrogen). Libraries were pooled at normalized molecular numbers for a 0.1× shallow WGS as quality control on a MiniSeq System (H3K7TG, Illumina). High-fidelity libraries were subjected to 20 × whole-genome sequencing on a NovaSeq 6000 system (Illumina).

### PTA-Seq data processing and somatic SNV detection

Paired-end reads from 16 PTA-amplified single neurons from donor ID01 were independently aligned to GRCh38 using BWA-MEM (v.0.7.17-r1188; -K 100000000 -Y), coordinate-sorted with SAMtools (v.1.12) and duplicate-marked with Picard MarkDuplicates (v.2.22.1). Base quality scores were recalibrated using GATK BaseRecalibrator and ApplyBQSR (v.4.1.4.1) with dbSNP build 138, Mills and 1000 Genomes gold-standard indels and assembly38 known indels. Somatic SNVs were called on autosomes 1–22 using SCAN2 (v.2.0) ^20^, pairing each neuron with the matched Illumina bulk sequencing. GATK HaplotypeCaller (v.4.2.6.1) jointly analysed each neuron–bulk pair in separate passes with minimum mapping qualities of 60 and 1, excluding soft-clipped bases. Bulk heterozygous SNPs were phased using SHAPEIT (v.2.r904) with the 1000 Genomes Phase 3 reference panel and used to estimate local allelic imbalance through SCAN2’s chromosome-specific spatial Gaussian-process model. Candidate allele-balance distributions were inferred from heterozygous SNPs within 100-kb flanks, with parsimony-based phase adjustment disabled. Candidates required ≥2 alternate reads and depth ≥6 in the neuron, bulk depth ≥11 with no alternate reads, absence from the supplied dbSNP common-variant resource, and an allele-balance-consistency P value >0.05. Alternate evidence in the low-mapping-quality bulk call set also caused exclusion. Empirical indel and clipping scores, calculated from overlapping paired reads with mapping quality ≥60, were required to exceed their respective fifth-percentile germline-training thresholds. SCAN2 calculated separate model-based false-discovery rates for pre-amplification and early-amplification artefacts, requiring bulk homozygous-reference status. Benchmark analyses retained only GRCh38-represented records annotated as muttype = snv and pass = TRUE, with both artefact-model false-discovery rates <0.01; indels and rescue-only records were excluded. Calls were retained separately by neuron and subjected to the shared benchmark-wide post-calling filters described below.

### Downsampling and molecule filtering

For cross-method comparisons, GRCh38 alignments were deterministically downsampled with random seed 42 using SAMtools view -s. Illumina and ppmSeq data were sampled to nominal 30×, 100× and 200× depths, with the unsampled data designated 350×; UDSeq was sampled to nominal 30×, with the unsampled data designated 100×. NanoSeq, HiDEF-seq and PTA-Seq data were not downsampled. NanoSeq and UDSeq molecular families were defined by alignment start and orientation-normalized paired unique molecular identifiers and retained when supported by both F1R2 and F2R1 reads after standard alignment-quality filtering. ppmSeq reads were retained when both st and et tags were MIXED. For HiDEF-seq, assay-filtered coverage and RIVER analyses required zero-mode waveguides (ZMWs) containing both forward- and reverse-strand consensus records after exclusion of secondary, supplementary, duplicate and vendor-quality-control-failed alignments. Coverage-uniformity modelling and trinucleotide-context analyses used the CCS-filtered alignment before this additional selection. No assay-specific molecule filtering was applied to Illumina or PTA-Seq data. Analyses were restricted to chromosomes 1–22 and X.

### Genome coverage and uniformity analysis

A common coverage exclusion mask combined segmental duplications, assembly gaps, satellite and ribosomal DNA regions, the ENCODE blacklist, 50-mer Umap regions with mappability <0.4, simple repeats, and homopolymers ≥7 bp using UCSC bigWigToBedGraph (v.469) and BEDTools (v.2.31.1). BigWig-derived intervals were selected at signal ≥0.1, except for Umap. The retained genome spanned 2.546 Gb and was used across methods. Complementary cumulative coverage distributions were calculated using mosdepth (v.0.3.12) in fast mode against the GRCh38 full analysis set plus decoy and HLA reference, without a minimum mapping-quality threshold and excluding secondary, vendor-quality-control-failed, duplicate, and supplementary alignments. PTA-Seq coverage was represented by the pointwise mean of the 16 single-neuron distributions. “Raw” denotes alignments before assay-specific molecule selection but after the shared genomic and alignment-flag exclusions.

Single-base depth was calculated separately using SAMtools depth with a minimum base quality of 13, no minimum mapping-quality threshold, and exclusion of unmapped, secondary, vendor-quality-control-failed, duplicate, and supplementary alignments. Zero-depth counts were inferred from the masked-reference denominator after excluding chromosome-terminal positions and positions lacking canonical A/C/G/T trinucleotide contexts. Depth distributions were fitted by maximum likelihood to the conventional zero-inflated Poisson (ZIP) and zero-inflated negative-binomial (ZINB) models to assess the data uniformity and to capture the real-world overdispersion parameter beyond the Poisson lambda, since negative binomial distributions may represent a Gamma-Poisson mixture distribution ^25^. Coverage breadth was the proportion of eligible positions with depth ≥1. Coverage nonuniformity was summarized by the ZINB overdispersion parameter α, whose negative-binomial component has mean μ and variance μ + αμ²; larger α values indicate less uniform genomic sampling. Four PTA neurons were included in this model-based comparison.

### Trinucleotide-context analysis

Reference trinucleotides at eligible covered positions were collapsed into 32 pyrimidine-oriented contexts by reverse-complementing A-or G-centered sequences. Full-depth sequencing profiles were normalized to relative abundance and compared with the corresponding masked GRCh38 reference profile. PTA-Seq counts were pooled across the 16 neurons for reference-composition comparisons. For cross-method similarity analysis, profiles were normalized independently, the 16 PTA-Seq profiles were averaged, and pairwise cosine distance (1-cosine similarity) was calculated, followed by average-linkage hierarchical clustering.

### Call-set integration and genomic filtering

Full-depth SNVs were merged by exact GRCh38 chromosome, position, reference, and alternate alleles across ten call sets: IlluminaSeq; multiread and singleton ppmSeq, UDSeq and NanoSeq; HiDEF-seq double-stranded and single-stranded DNA calls; and PTA-Seq singleton calls. Duplex multiread and singleton calls denote support from multiple independent molecules or one molecule, respectively; only high-confidence ppmSeq singletons were included. Candidates were excluded for segmental-duplication, assembly-gap or satellite/ribosomal DNA overlap (locus-level bigWig signal ≥0.1); mean 50-mer Umap mappability <0.4 or ENCODE blacklist v.2 signal ≥0.1 within a centred 20-bp window; UCSC simple-repeat or ≥7-bp homopolymer overlap; exact allele matches to GATK known-indels or Mills and 1000 Genomes gold-standard indel records; or gnomAD allele frequency ≥0.001. Repeat overlaps were determined using BEDTools.

### Read-level Integration for Variant Evidence Recovery (RIVER)

Detailed instructions for users, as well as the demo input and output, are provided on GitHub (https://github.com/Lola-W/RIVER). RIVER represents a series of steps to recover MVs from multiple duplex method discovery workflows. We developed RIVER to assess cross-method read support at candidate SNVs in the integrated GRCh38 matrix. Candidate alleles were queried in full-depth IlluminaSeq alignments, assay-selected full-depth ppmSeq, UDSeq, NanoSeq, and HiDEF-seq alignments prepared as described under “Downsampling and molecule filtering”, and 16 PTA-Seq single-neuron alignments. Candidate alleles were also queried in unrelated controls. Pileups were generated using SAMtools mpileup (-Q 13 -q 0 -A -B --ff SECONDARY,QCFAIL,DUP,SUPPLEMENTARY -d 1000000) against the GRCh38 full analysis set plus decoy and HLA reference. hg19-aligned controls were queried against human_g1k_v37_decoy after CrossMap conversion using the hg38ToHg19 chain.

For candidate i in dataset m, let *A_im_* and *R_im_* denote the retained alternate and reference observations, respectively, summed across read orientations. The allele-count denominator was *n_im_* = *A_im_* + *R_im_*, excluding observations of other alleles, and the estimated variant allele fraction was 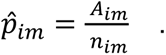 Sampling uncertainty was summarized using two-sided 95% Wilson score intervals for the binomial proportion. For *n_im_* > 0, the lower confidence bound was

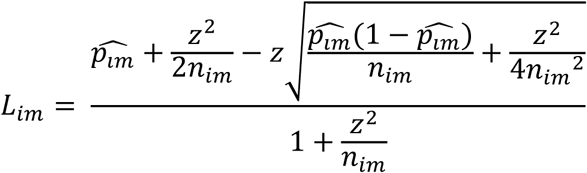

where *z* = *Φ*^−1^(0.975) ≈ 1.96. VAFs and confidence intervals were undefined when no reference or alternate observations were available.

### RIVER Statistical Framework

Read support in IlluminaSeq and the four duplex methods required a lower 95% Wilson VAF bound >0.001, whereas PTA-Seq support required ≥1 alternate observation in any of the 16 neurons; support in exactly one or ≥2 neurons was recorded separately.

Support did not require a call from the supporting method or per-site alternate-allele agreement between strands. No additional minimum-depth or fixed alternate-read-count threshold was applied to bulk-method support. Following the shared genomic, population, germline and unrelated-control exclusions, support was counted across sequencing methods, combining molecular-support classes within each method and treating all PTA neurons collectively as one method. For each originating call set, its own sequencing method was excluded, and variants were classified as supported by zero, one, or ≥2 other methods. Proportions were calculated over all filtered variants in that call set, irrespective of whether they met the originating method’s own read-support threshold. Zero cross-method support indicated absence of qualifying evidence, not confirmation of a false-positive call.

### Germline and unrelated-control filtering

Candidates matching the matched-normal HaplotypeCaller/VQSR call set were excluded as germline. Additional germline-like exclusions required ≥1 alternate read, REF + ALT depth ≥10, and an upper 95% Wilson VAF bound >0.52 in any of 12 bulk datasets. These datasets comprised IlluminaSeq and duplex-filtered ppmSeq at full depth and nominal 30×, 100× and 200×; duplex-filtered UDSeq at full depth and 30×; and full-depth duplex-filtered NanoSeq and HiDEF-seq. PTA neurons were excluded from this screen. The unrelated-donor panel comprised IlluminaSeq from donor 6566 blood and sperm and donor 7669; ppmSeq, UDSeq and NanoSeq from donor 6566; and two PTA libraries each from donors 6566 and 8420. Candidates were removed for a lower 95% Wilson VAF bound >0.001 in any bulk control or ≥1 alternate read in any PTA control, irrespective of discovery method. Within the pooled candidate matrix, genomic, population and germline filtering retained 123,270 of 149,382 candidates; subsequent unrelated-control filtering retained 113,452 variants for concordance analyses.

### Caller-level and cross-method concordance

Germline removal, subsequent unrelated-control removal and retention were summarized relative to the post-genomic-prefilter total within each call set. Caller-level intersections used membership in the ten call sets, whereas read-level intersections used the six method-level RIVER indicators. Missing membership and support indicators were treated as absent. IlluminaSeq VAF distributions were compared by the total number of supporting methods, excluding the zero- and six-method groups.

Pairwise VAF concordance among the five bulk methods was assessed using variants with qualifying support and non-missing VAFs in both datasets. Pearson correlations and two-sided tests were calculated using R ‘cor.test’ for pairs with ≥3 shared variants. P-values were adjusted across the ten method pairs using the Benjamini–Hochberg procedure, with FDR <0.05 considered significant.

### Mutational spectra and signature analysis

Signature analyses used separate caller-level and read-evidence exports rather than the 113,452-variant concordance set. Bulk support criteria were unchanged, but PTA support and unrelated-PTA exclusions required ≥1 alternate read, REF + ALT depth ≥10, and an upper 95% Wilson VAF bound >0.52 within an individual library. Caller-level exports were reservoir-sampled to ≤3,000 variants per call set (seed 42) before deduplication. Read-evidence duplicates were collapsed by retaining the maximum binary support indicator per method. SBS96 spectra included SNVs on chromosomes 1–22 and X with ‘PASS’ or unset VCF FILTER fields, canonical GRCh38 trinucleotide contexts and matching reference alleles; purine-reference substitutions were reverse-complemented.

*De novo* signatures were extracted using SigProfilerExtractor (v.1.2.6)^22^, evaluating 1–8 caller-level and 1–6 read-evidence signatures with 200 non-negative matrix-factorization replicates, GMM matrix normalization and resampling. Solution-estimation thresholds were set to 0.8 for stability, 0.2 for minimum stability and 1.0 for combined stability, with no stability drop allowed. The selected caller-level and read-evidence solutions contained four and five signatures, respectively. COSMIC SBS v.3.4 decomposition used SigProfilerAssignment (v.1.1.1), GRCh38 mutational opportunities and non-negative least squares, with add, remove and initial-remove penalties of 0.05, 0.01, and 0.05, respectively. Activities were normalized to 100% within each call set or method. Activity heatmaps reused the average-linkage dendrogram calculated from the corresponding SBS96 count spectra using cosine distance, rather than reclustering activities.

### Statistical software and visualization

Analyses used Python v.3.13.11 with SigProfilerExtractor v.1.2.6, SigProfilerPlotting v.1.4.2, SigProfilerMatrixGenerator v.1.3.6, SigProfilerAssignment v.1.1.1, pandas v.2.3.3, NumPy v.2.4.0, SciPy v.1.16.3 and scikit-learn v.1.8.0. Data manipulation and visualization were performed in R v.4.5.1 using readr v.2.1.6, dplyr v.1.1.4, tidyr v.1.3.2, stringr v.1.6.0, ggplot2 v.4.0.1, UpSetR v.1.4.0, ComplexHeatmap v.2.26.0, circlize v.0.4.17, ggdendro v.0.2.0 and patchwork v.1.3.2.

## Data availability

Sequencing datasets used in this study have been deposited in the NCBI Sequence Read Archive (SRA) and the European Nucleotide Archive (ENA). HiDEF-seq data from the brain of donor ID01 are available under NCBI BioProject accession PRJNA1537344, deep bulk Illumina whole-genome sequencing (IlluminaSeq) data from donor ID01 under PRJNA1537534, and targeted AmpliSeq validation data under PRJNA1537374. NanoSeq data from the brain and matched liver of donor ID01 are available through ENA under PRJEB127894, ppmSeq brain data under PRJEB127901, and UDSeq brain and matched left kidney data under PRJEB127902. Single-neuron PTA-based whole-genome sequencing data from donor ID01 are available under PRJEB127899 and PRJEB127900. Control datasets from donor ID06 are available under PRJNA1537641 (NanoSeq sperm), PRJEB127895 (UDSeq sperm), PRJEB127896 (ppmSeq sperm), PRJEB127897 (IlluminaSeq sequencing of sperm) and PRJEB127898 (IlluminaSeq of blood).

## Code availability

Details and codes for the data processing and annotation are provided on GitHub (https://github.com/Lola-W/Benchmark_RIVER_paper). RIVER is currently implemented in Python; the source code, documentation and demos are available on the GitHub (https://github.com/Lola-W/RIVER).

---

**Extended Data Fig. 1.**
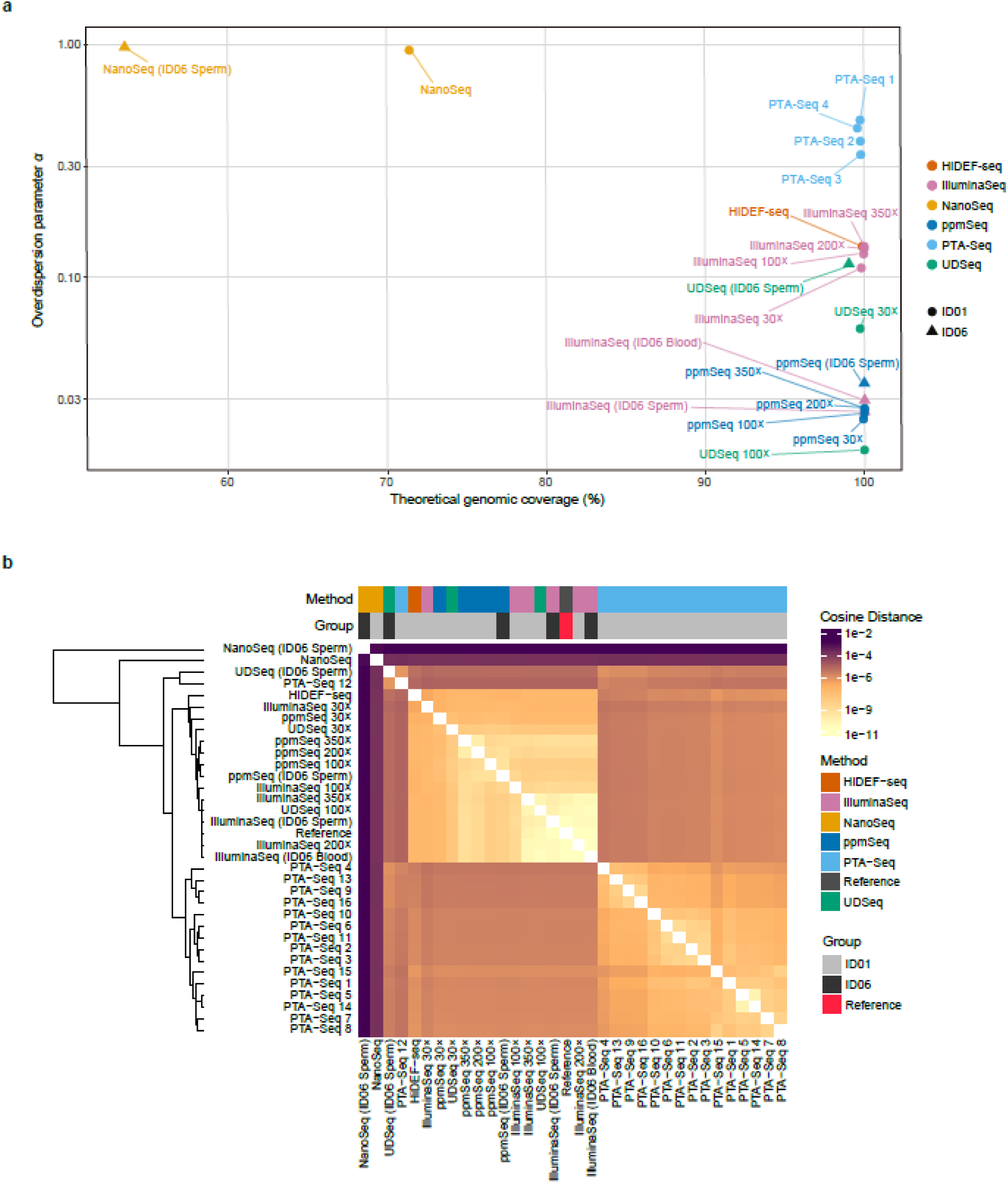
Method-specific read landscapes are consistent across donors and sequencing depths. **a,** Genome-wide coverage breadth and read-depth overdispersion for ID01 left temporal cortex and ID06 blood or sperm profiles. X-axis: % GRCh38 positions with depth ≥1; Y-axis: zero-inflated negative binomial (ZINB) overdispersion parameter, α, plotted on a log10 scale. Larger α values: less uniform genomic sampling. Points: full-depth or down-sampled profiles; colors: method; circles: ID01; triangles: ID06. PTA-Seq: four single-neuron profiles. **b,** Pairwise cosine-distance heatmap and hierarchical clustering of the 32 pyrimidine-first trinucleotide-context profiles across methods, donors, and the GRCh38 reference. Annotation bars: method and group. PTA-Seq: all n = 16 single-neuron profiles. Cosine distance calculated as 1 – cosine similarity; smaller values: greater similarity.

**Extended Data Fig. 2.**
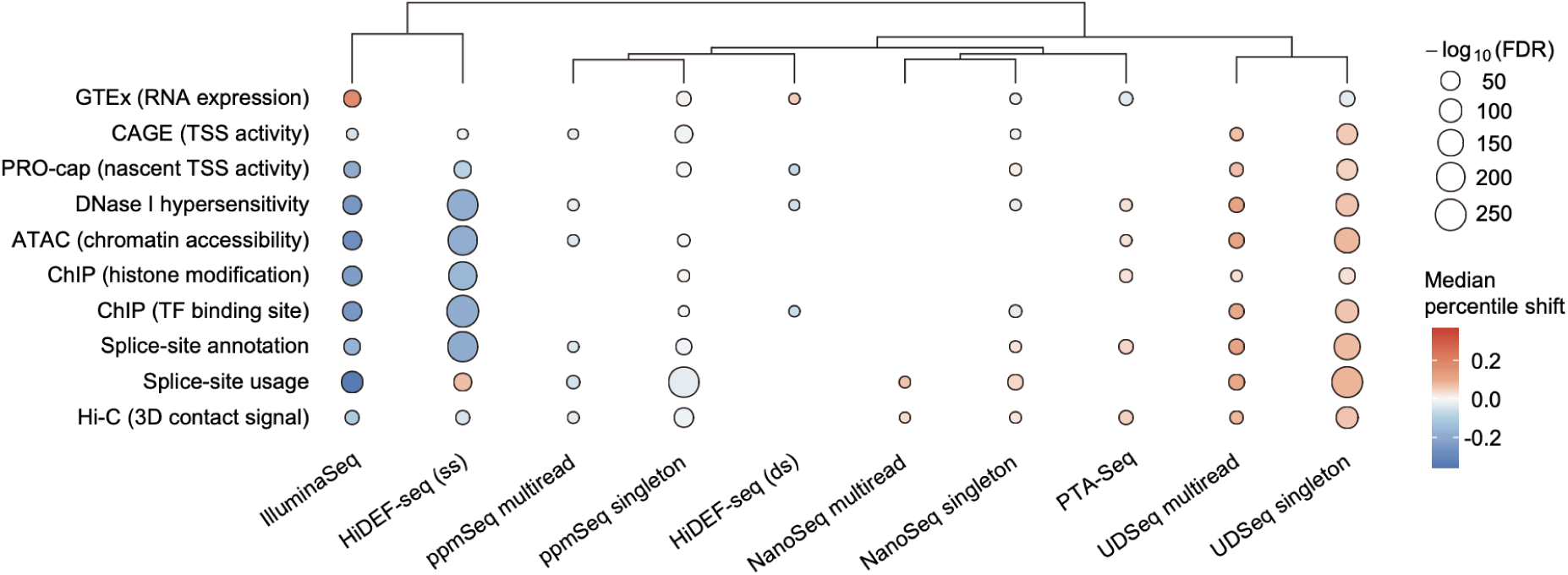
Method-specific functional annotation profiles of mosaic variants. Rows: genomic functional annotations; columns: method and molecular-support class. Circle color: median percentile shift, with positive and negative values indicating higher and lower annotation percentiles, respectively; circle size:−log10(FDR). Dendrogram: hierarchical clustering of call sets by functional annotation profiles. ds, double-stranded; ss, single-stranded; TF, transcription factor; TSS, transcription start site.

**Extended Data Fig. 3.**
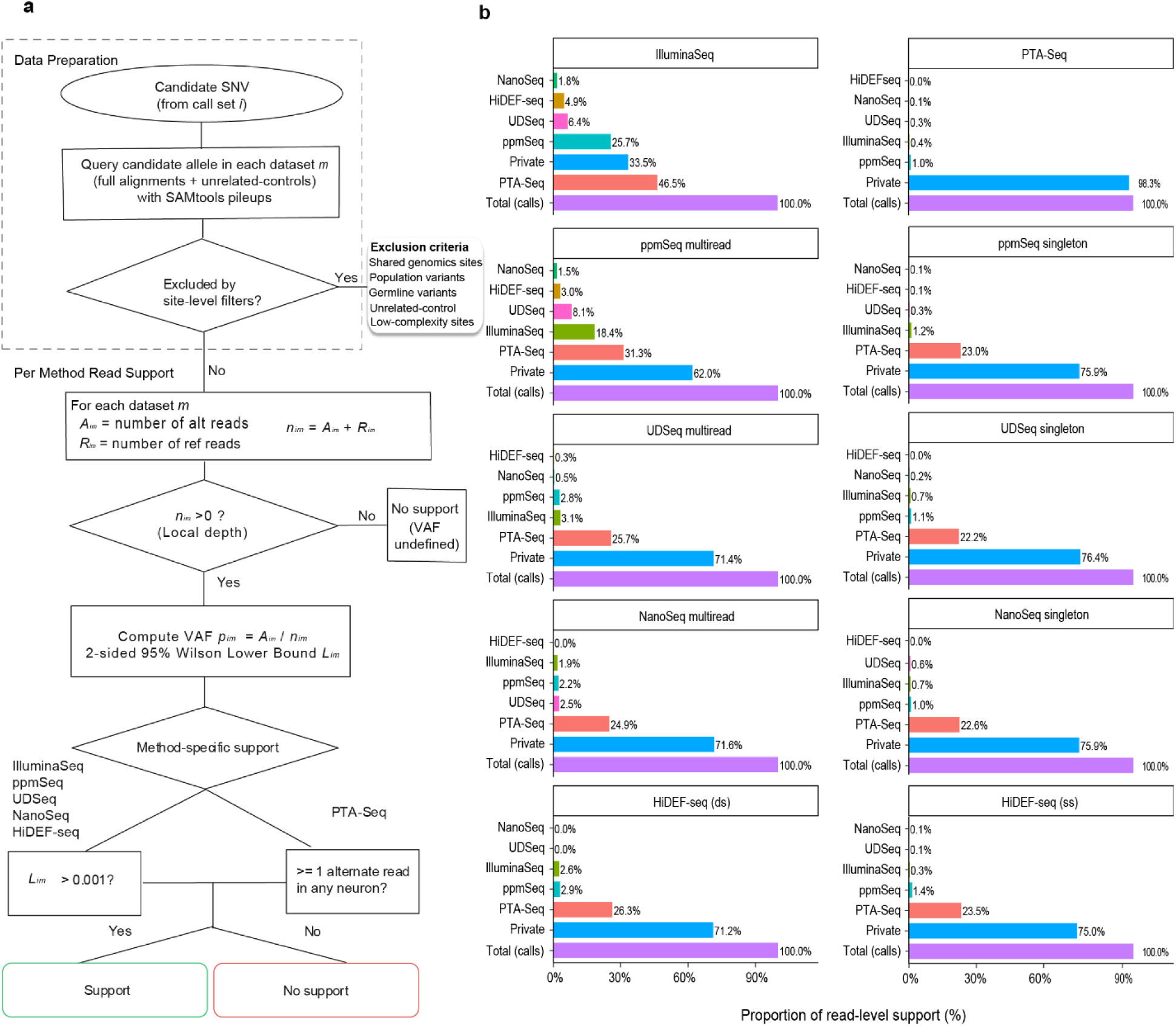
RIVER workflow and cross-method read-level support for mosaic variant call sets. **a,** Read-level Integration for Variant Evidence Recovery (RIVER) workflow. Candidate alleles are queried in full-depth sequencing alignments and unrelated controls using SAMtools pileups, followed by the indicated site-level exclusions. Variant allele fraction (VAF) is calculated as ALT/(REF + ALT), where ALT and REF are alternate- and reference-read counts. IlluminaSeq and duplex-method support requires a lower bound of the two-sided 95% Wilson VAF interval >0.001; PTA-Seq support requires ≥1 alternate read in any neuron. Sites with no reference or alternate reads have undefined VAF and no support. **b,** Cross-method support for retained full-depth call sets, stratified by discovery method and molecular-support class. Bars show the proportion of source calls with qualifying read-level support in each other method. ‘Private’ denotes no qualifying support in any other method; ‘Total (calls)’ denotes the source call set after filtering and RIVER assessment, normalized to 100%. Cross-method support categories are non-exclusive, and support does not require a call from the supporting method. For duplex call sets, multiread and singleton denote support by multiple independent molecules or one molecule in the discovery dataset, respectively. ds, double-stranded; ss, single-stranded; PTA, primary template-directed amplification.

**Extended Data Fig. 4.**
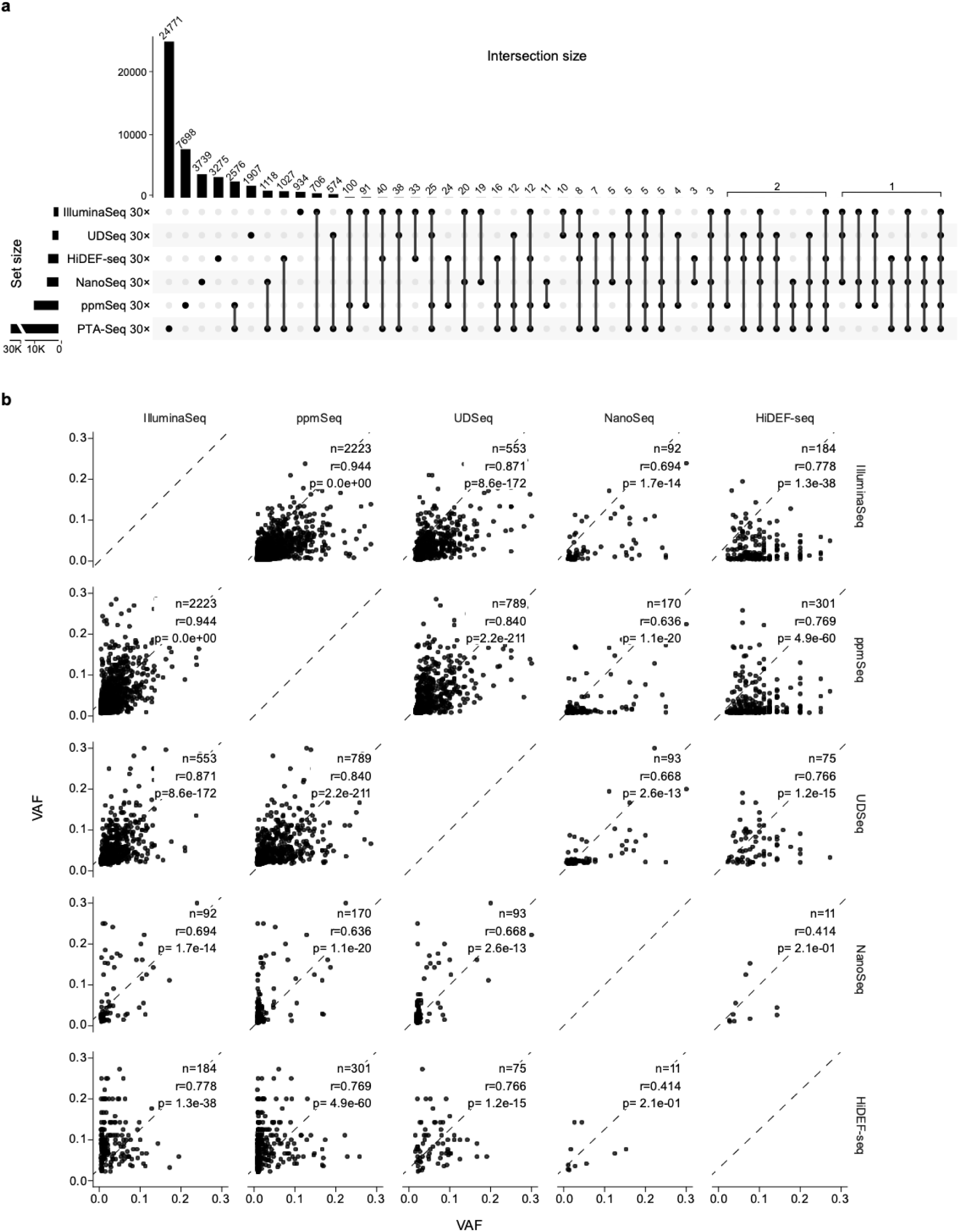
Cross-method read-level overlap and allele-fraction concordance of mosaic variants. **a,** UpSet plot of read-supported MV sets after random downsampling to 30×. Horizontal bars: set sizes; vertical bars and labels: intersection sizes; connected dots: contributing methods. PTA-Seq: aggregated single-neuron support. Set-size axis: discontinuous. **b,** Pairwise VAF comparisons for MVs with read-level support in both methods. Points: individual MVs; dashed lines: equal VAFs. Axes: square-root-transformed VAFs; tick labels: original VAFs. Labels: shared MV count (n), Pearson’s r, and FDR. Correlations and two-sided tests used untransformed VAFs; FDR: Benjamini–Hochberg adjustment across 10 unique method pairs. FDR = 0 indicates numerical underflow. FDR, false-discovery rate; MV, mosaic variant; PTA, primary template-directed amplification; VAF, variant allele fraction.

**Extended Data Fig. 5.**
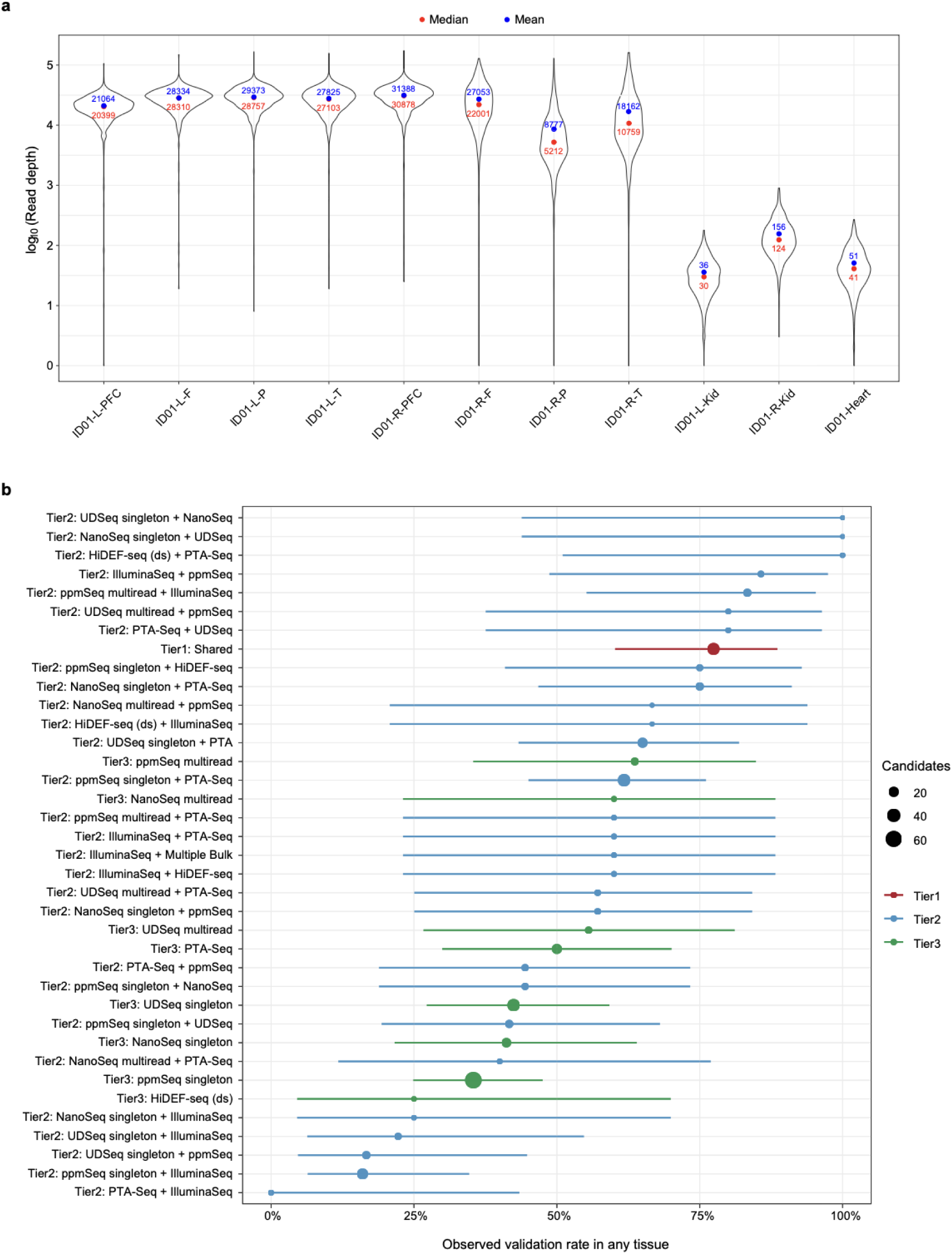
Targeted sequencing coverage and evidence-stratified validation of mosaic variants. **a,** Read-depth distributions across targeted loci in eight brain and three peripheral tissues from donor ID01. Violins: distributions on a log10 scale; red and blue points: median and mean depths, respectively; labels: untransformed read depths. **b,** Validation rates in any sampled tissue by evidence tier, discovery method, read-support class and cross-method support. Points: observed rates; horizontal lines: Wilson 95% CIs; size: number of tested candidates; color: tier. Tier 2 labels: calling method followed by method(s) providing additional read-level support. Shared: tier 1 candidates called by ≥2 methods; Multiple Bulk: read support from multiple bulk-tissue methods; tier 3: method-private candidates without cross-method support. CI, confidence interval; ds, double-stranded; PTA, primary template-directed amplification. For Tier 2 candidates, labels follow the format ‘calling method + read-supporting method’. Singleton and multiread refer to molecular support in the dataset of the calling method.

**Supplementary Table 1 | Integrated mosaic variant candidates, annotations and cross-method read-level evidence.** Rows: 149,382 candidate single-nucleotide variants (SNVs) before harmonized filtering, indexed by GRCh38 chromosome, position, reference and alternate alleles. Columns: caller membership by method and molecular-support class, genomic and functional annotations, filtering flags, and allele measurements from full-depth and downsampled datasets. MAF: variant allele fraction (VAF), calculated as ALT/(REF + ALT); LOWER_CI and UPPER_CI: Wilson 95% confidence bounds. Caller membership, qualifying cross-method read support, and alternate-read presence (is_existing_lower_ci_0001_DUPLEX_PPMSEQ or is_altge1_*, ≥1 alternate observation) are recorded separately. PTA-Seq support is summarized across 16 single-neuron profiles. *_IS_GREATER: within-BAM local-background flags; P/F indicate whether the candidate VAF’s lower 95% Wilson bound exceeds/does not exceed the upper bound for the most abundant non-reference event at each covered neighboring position within ±5 bp. The underscore separates upstream and downstream positions; these flags do not represent final variant classification. Candidates subsequently excluded remain in the matrix with filtering annotations. For duplex methods, multiread and singleton denote support by multiple independent molecules or one molecule, respectively. BAM, binary alignment map; CI, confidence interval; HC, high confidence; PTA, primary template-directed amplification.

**Supplementary Table 2 | Empirical RIVER false-positive recovery in unrelated controls.** Rows: originating call sets evaluated before panel-of-normals (PoN) exclusion, with the originating sequencing method excluded from the control recovery panel. Columns: candidate SNV counts, eligible control BAMs or PTA cells, recovered SNVs, recovery rates and Wilson 95% confidence intervals, reported separately for bulk and PTA controls. Recovery rate: proportion of tested candidates meeting RIVER support criteria in at least one eligible control. Bulk support requires a lower 95% Wilson VAF bound >0.001; PTA support requires ≥1 alternate observation. PTA-to-PTA recovery is not applicable because the originating method is excluded. RIVER, Read-level Integration for Variant Evidence Recovery; BAM, binary alignment map; CI, confidence interval; HC, high confidence; n.a., not applicable; PTA, primary template-directed amplification; SNV, single-nucleotide variant; VAF, variant allele fraction.

**Supplementary Table 3 | Targeted amplicon sequencing measurements and candidate selection for orthogonal validation.** Tab1_all_ampliseq_sites: 531 retained target loci, comprising 465 mosaic variant (MV) candidates and 66 germline control loci, across eight brain and three peripheral tissues from donor ID01. Rows: tissue–locus combinations. Columns: GRCh38 coordinates, genomic annotations, discovery source, reference and alternate allele counts, VAFs (MAF) and Wilson 95% confidence bounds. NORMAL_*: corresponding measurements in the unrelated control. CI_IS_GREATER and NORMAL_CI_IS_GREATER: local-background P/F flags defined in Supplementary Table 1, calculated separately within sample and control BAMs. Tab2_strata: candidate-selection counts by evidence tier and caller/read-support combination. n: available candidates; submitted_num: candidates submitted for panel design; amplicon_made_and_genome_filtered: targets retained after amplicon generation and genomic filtering. Tier 1: called by ≥2 methods; tier 2: called by one method with additional cross-method read support; tier 3: called by one method without cross-method support. Multiread and singleton refer to molecular support in the originating method. BAM, binary alignment map; CI, confidence interval; HC, high confidence; PTA, primary template-directed amplification; VAF, variant allele fraction; VCF, variant call format.

**Supplementary Data 1 | Measured allelic fractions for 238 validated mosaic variants by targeted amplicon sequencing.** Each page shows a single validated variant, with corresponding tier and method source. Upper dash: 95% CI for germline variants; Lower dash: 75% CI for detection limit. Abbreviation: left: L; right: R; prefrontal (PF); frontal (F); parietal (P); temporal (T) cortex; kidney (K); heart (H). y-axis: sqrt(AF). Red: variants: measured AF surpassed the lower bound of the confidence interval for validation; Blue: measured AF failed to surpass the lower bound of the confidence interval and were considered non-validated.

## References

1. Lodato, M. A. et al. Somatic mutation in single human neurons tracks developmental and transcriptional history. Science 350, 94–98 (2015).

2. Huang, A. Y. et al. Parallel RNA and DNA analysis after deep sequencing (PRDD-seq) reveals cell type-specific lineage patterns in human brain. Proc. Natl. Acad. Sci. U. S. A. 117, 13886–13895 (2020).

3. Park, S. et al. Clonal dynamics in early human embryogenesis inferred from somatic mutation. Nature 597, 393–397 (2021).

4. Spencer Chapman, M., et al. Lineage tracing of human development through somatic mutations. Nature 595, 85–90 (2021).

5. Ju, Y. S. et al. Somatic mutations reveal asymmetric cellular dynamics in the early human embryo. Nature 543, 714–718 (2017).

6. Coorens, T. H. H. et al. Extensive phylogenies of human development inferred from somatic mutations. Nature 597, 387–392 (2021).

7. Bizzotto, S. et al. Landmarks of human embryonic development inscribed in somatic mutations. Science 371, 1249–1253 (2021).

8. Abascal, F. et al. Somatic mutation landscapes at single-molecule resolution. Nature 593, 405–410 (2021).

9. Bae, J. H. et al. Single duplex DNA sequencing with CODEC detects mutations with high sensitivity. Nat. Genet. 55, 871–879 (2023).

10. Liu, M. H. et al. DNA mismatch and damage patterns revealed by single-molecule sequencing. Nature 630, 752–761 (2024).

11. Cheng, A. P. et al. Paired plus-minus sequencing is an ultra-high throughput and accurate method for dual strand sequencing of DNA molecules. Preprint at

12. 10.1101/2025.08.11.669689 (2025).

12. Nandi, S. P. et al. A Universal Duplex Sequencing Approach for Accurate Detection of Somatic Mutations. Preprint at 10.1101/2025.09.14.676103 (2025).

13. Guille, A. et al. A benchmarking study of individual somatic variant callers and voting-based ensembles for whole-exome sequencing. Brief. Bioinform. 26, bbae697 (2024).

14. Ha, Y.-J. et al. Comprehensive benchmarking and guidelines of mosaic variant calling strategies. Nat. Methods 20, 2058–2067 (2023).

15. Breuss, M. W. et al. Somatic mosaicism reveals clonal distributions of neocortical development. Nature 604, 689–696 (2022).

16. Chung, C. et al. Cell-type-resolved mosaicism reveals clonal dynamics of the human forebrain. Nature 629, 384–392 (2024).

17. Dou, Y. et al. Accurate detection of mosaic variants in sequencing data without matched controls. Nat. Biotechnol. 38, 314–319 (2020).

18. Lindner, M. S., Kollock, M., Zickmann, F. & Renard, B. Y. Analyzing genome coverage profiles with applications to quality control in metagenomics. Bioinformatics 29, 1260–1267 (2013).

19. Rogozin, I. B. et al. Mutational signatures and mutable motifs in cancer genomes. Brief. Bioinform. 19, 1085–1101 (2018).

20. Luquette, L. J. et al. Single-cell genome sequencing of human neurons identifies somatic point mutation and indel enrichment in regulatory elements. Nat. Genet. 54, 1564–1571 (2022).

21. Fu, B. J., et al. A recurrent sequencing artifact on Illumina sequencers with two-color fluorescent dye chemistry and its impact on somatic variant detection. Genome Biol. 27, 179 (2026).

22. Islam, S. M. A. et al. Uncovering novel mutational signatures by de novo extraction with SigProfilerExtractor. Cell Genomics 2, None (2022).

23. Kim, J. et al. The use of technical replication for detection of low-level somatic mutations in next-generation sequencing. Nat. Commun. 10, 1047 (2019).

24. Viswanadham, V. V. et al. Combined somatic mutation and transcriptome analysis reveals region-specific differences in clonal architecture in human cortex. Cell Rep. 44, 116458 (2025).

25. Zhang, Y. et al. Benchmarking of duplex sequencing approaches to reveal somatic mutation landscapes. Preprint at 10.64898/2025.12.12.692823 (2025).

26. Yang, X. et al. Control-independent mosaic single nucleotide variant detection with DeepMosaic. Nat. Biotechnol. 41, 870–877 (2023).

27. Yang, X. et al. Developmental and temporal characteristics of clonal sperm mosaicism. Cell 184, 4772–4783.e15 (2021).

28. Cheng, A. P. et al. Error-corrected flow-based sequencing at whole-genome scale and its application to circulating cell-free DNA profiling. Nat. Methods 22, 973–981 (2025).

