## supplementary_data_1_bioRxiv.pdf for "Benchmarked sequencing methods reveal hidden mosaic variation in the human brain"

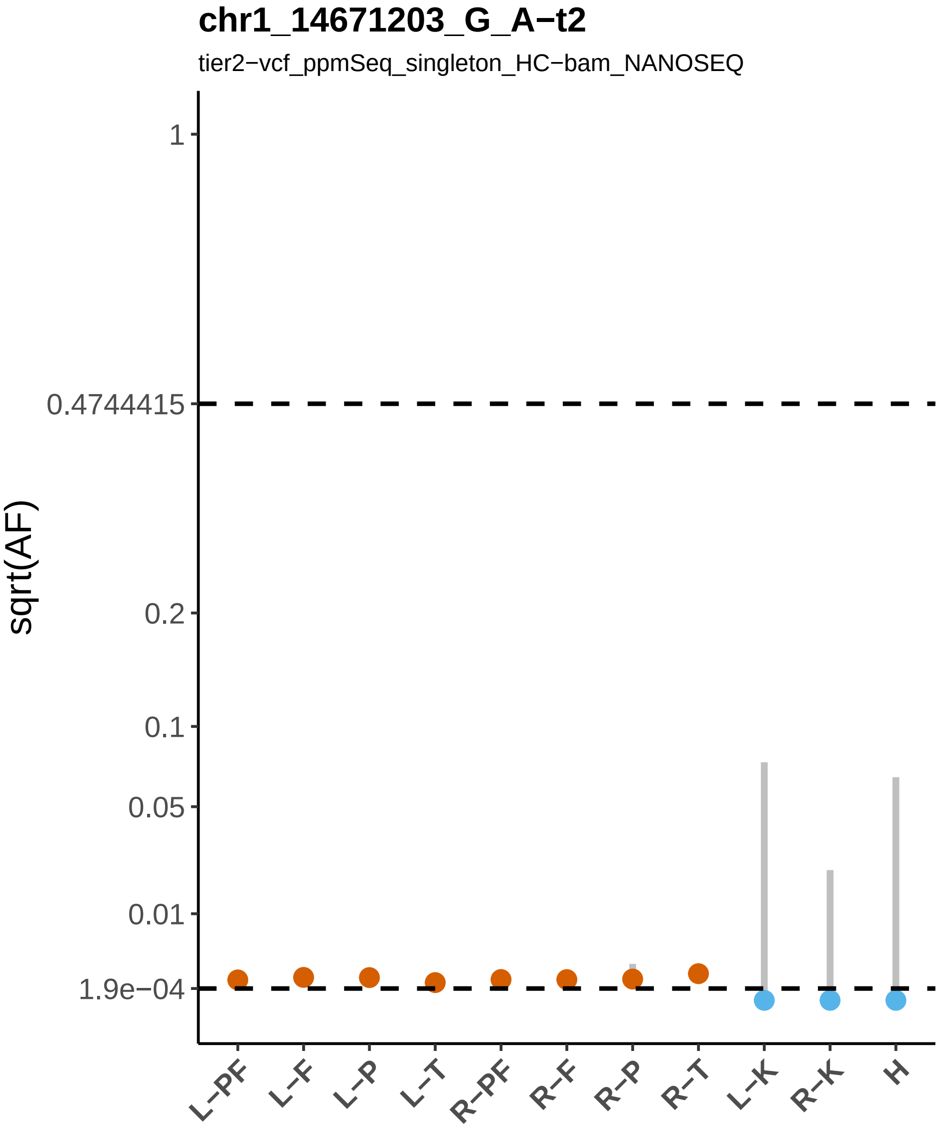

chr1_14671203_G_A−t2
tier2−vcf_ppmSeq_singleton_HC−bam_NANOSEQ
sqrt(AF)
1
0.4744415
0.2
0.1
0.05
0.01
1.9e−04
L−PF
L−F
L−P
L−T
R−PF
R−F
R−P
R−T
L−K
R−K
H

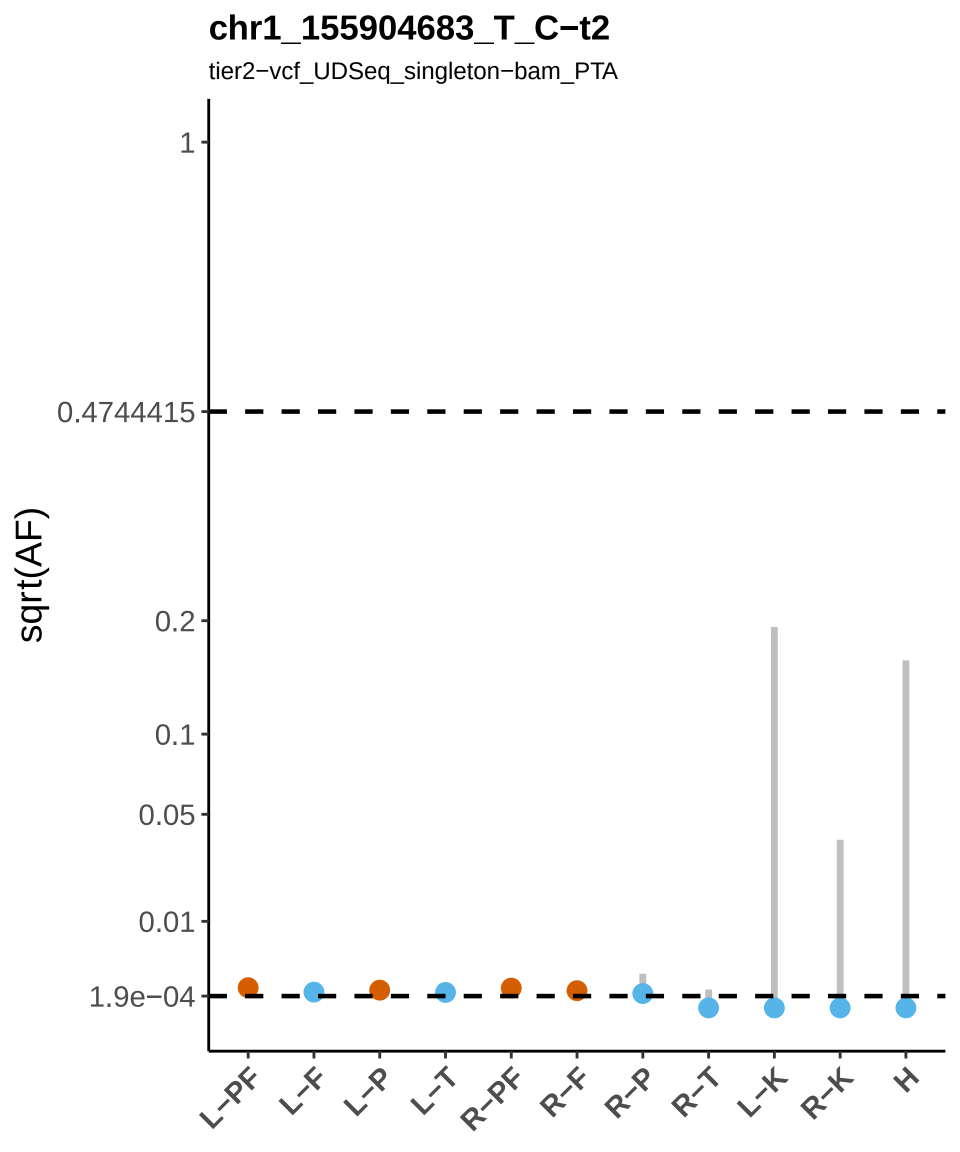
chr1_155904683_T_C−t2
tier2−vcf_UDSeq_singleton−bam_PTA
sqrt(AF)
1
0.4744415
0.2
0.1
0.05
0.01
1.9e−04
L−PF
L−F
L−P
L−T
R−PF
R−F
R−P
R−T
L−K
R−K
H

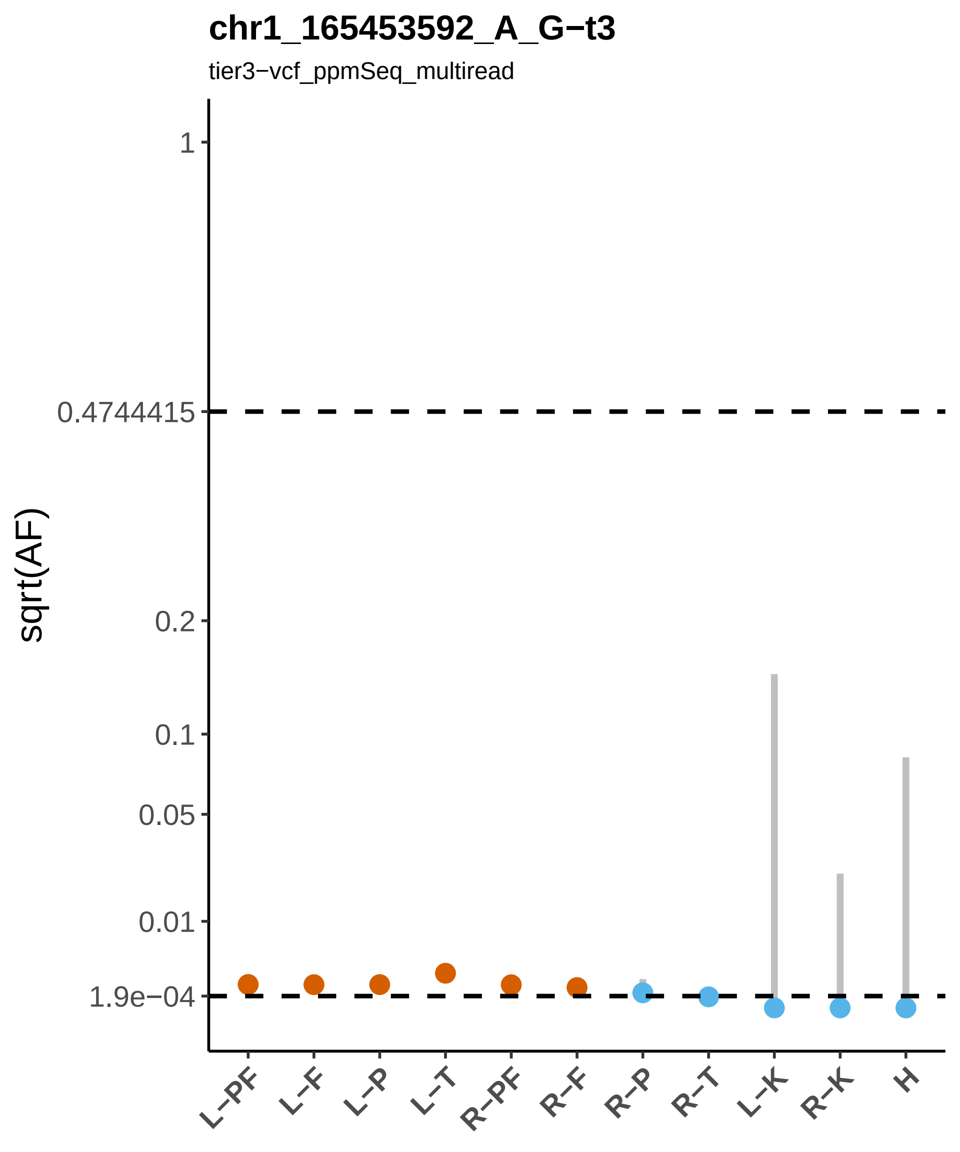

chr1_165453592_A_G−t3
tier3−vcf_ppmSeq_multiread
sqrt(AF)
1
0.4744415
0.2
0.1
0.05
0.01
1.9e−04
L−PF
L−F
L−P
L−T
R−PF
R−F
R−P
R−T
L−K
R−K
H

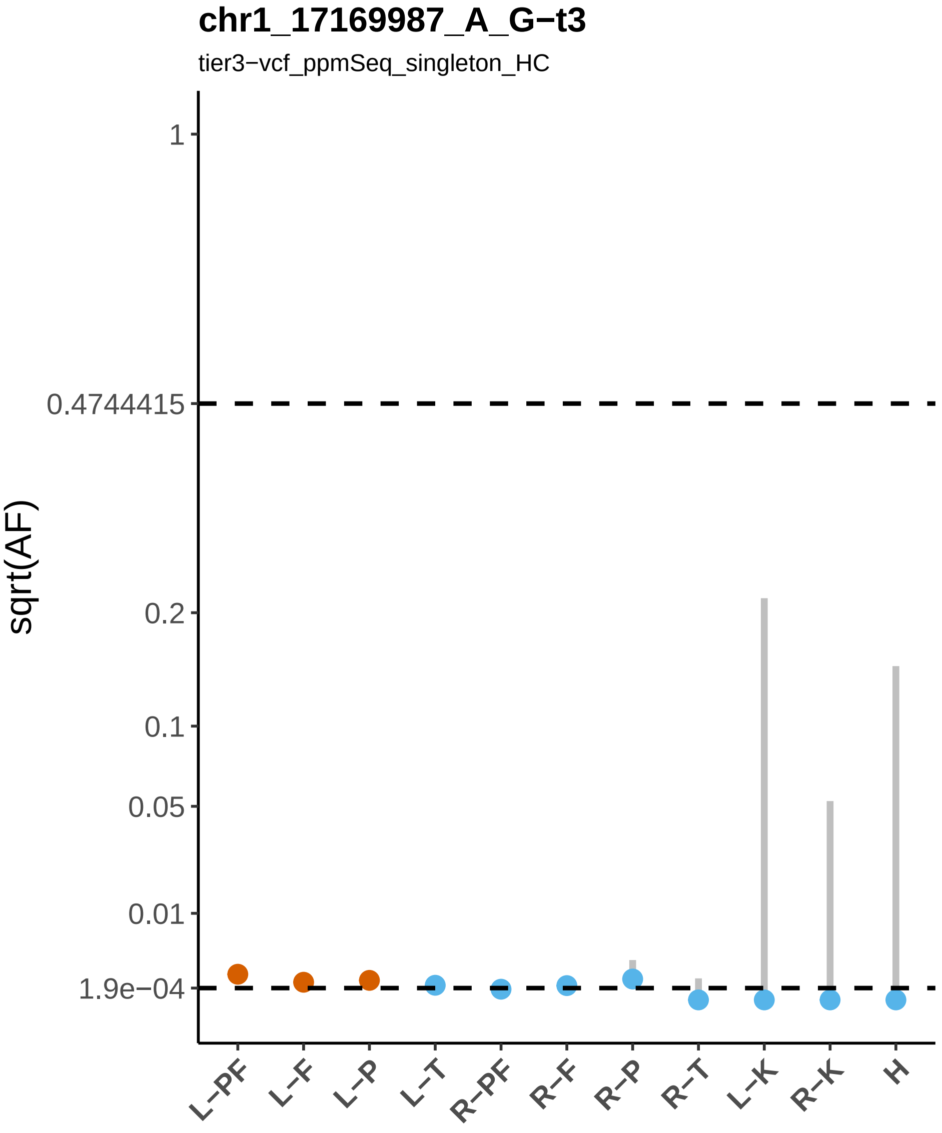
chr1_17169987_A_G−t3
tier3−vcf_ppmSeq_singleton_HC
sqrt(AF)
1
0.4744415
0.2
0.1
0.05
0.01
1.9e−04
L−PF
L−F
L−P
L−T
R−PF
R−F
R−P
R−T
L−K
R−K
H

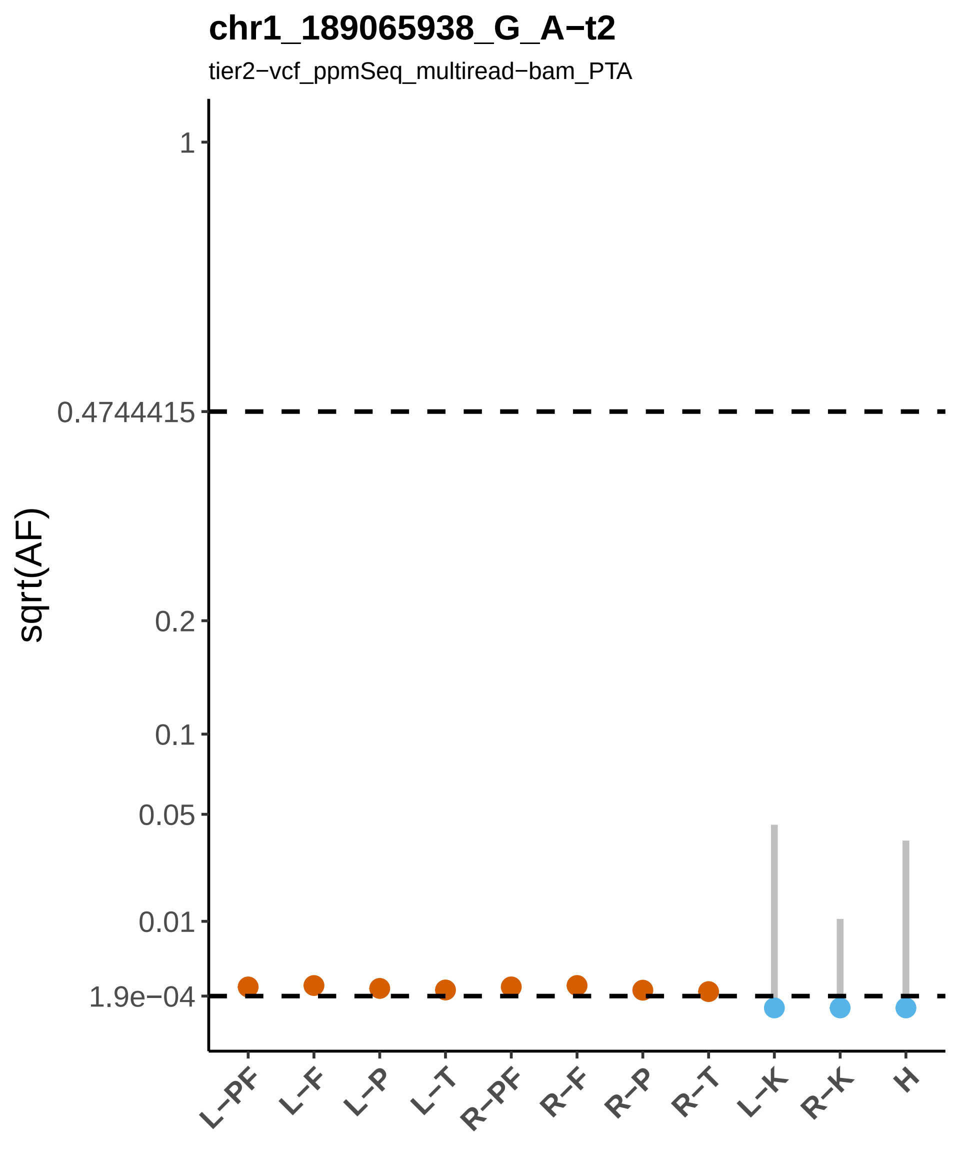

chr1_189065938_G_A−t2
tier2−vcf_ppmSeq_multiread−bam_PTA
sqrt(AF)
1
0.4744415
0.2
0.1
0.05
0.01
1.9e−04
L−PF
L−F
L−P
L−T
R−PF
R−F
R−P
R−T
L−K
R−K
H

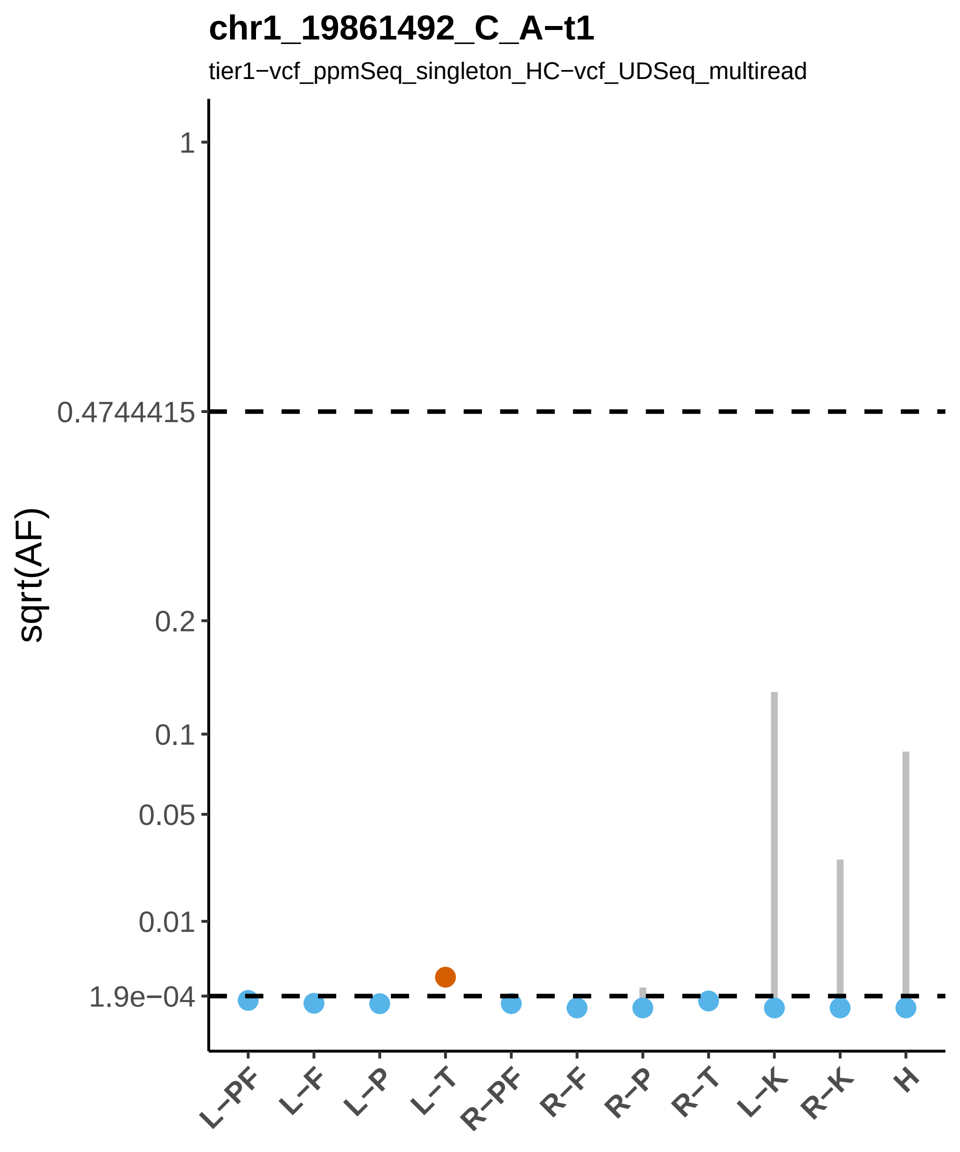
chr1_19861492_C_A−t1
tier1−vcf_ppmSeq_singleton_HC−vcf_UDSeq_multiread
sqrt(AF)
1
0.4744415
0.2
0.1
0.05
0.01
1.9e−04
L−PF
L−F
L−P
L−T
R−PF
R−F
R−P
R−T
L−K
R−K
H

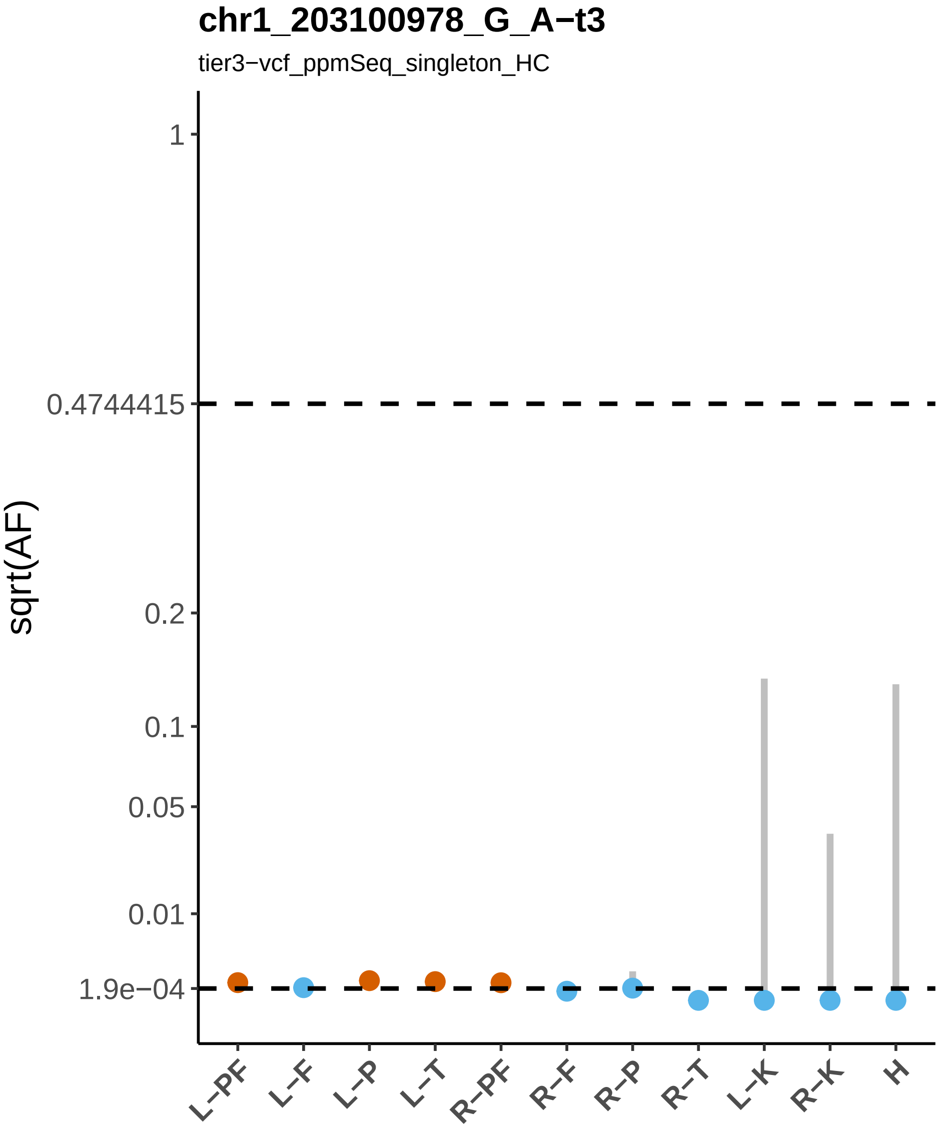
chr1_203100978_G_A−t3
tier3−vcf_ppmSeq_singleton_HC
sqrt(AF)
1
0.4744415
0.2
0.1
0.05
0.01
1.9e−04
L−PF
L−F
L−P
L−T
R−PF
R−F
R−P
R−T
L−K
R−K
H

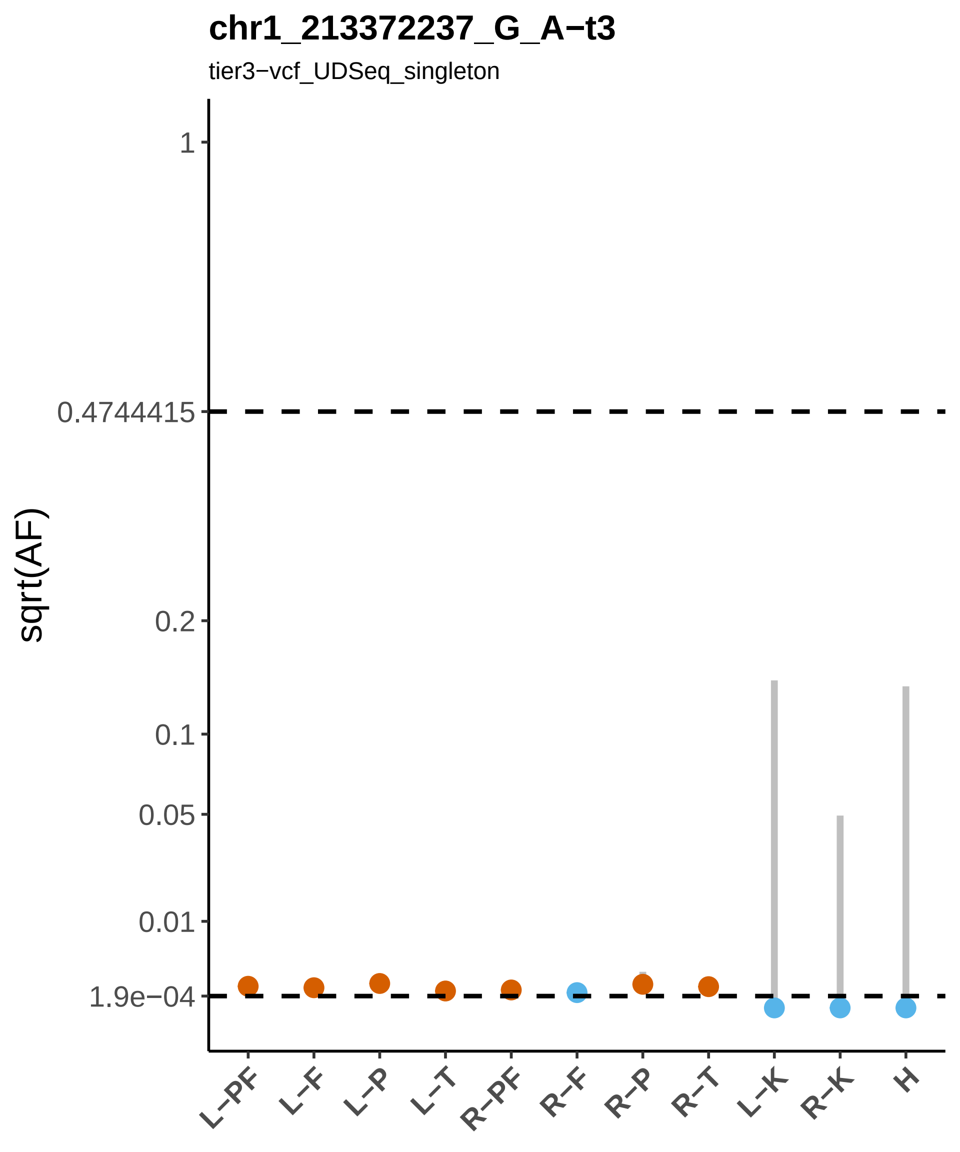
chr1_213372237_G_A−t3
tier3−vcf_UDSeq_singleton
sqrt(AF)
1
0.4744415
0.2
0.1
0.05
0.01
1.9e−04
L−PF
L−F
L−P
L−T
R−PF
R−F
R−P
R−T
L−K
R−K
H

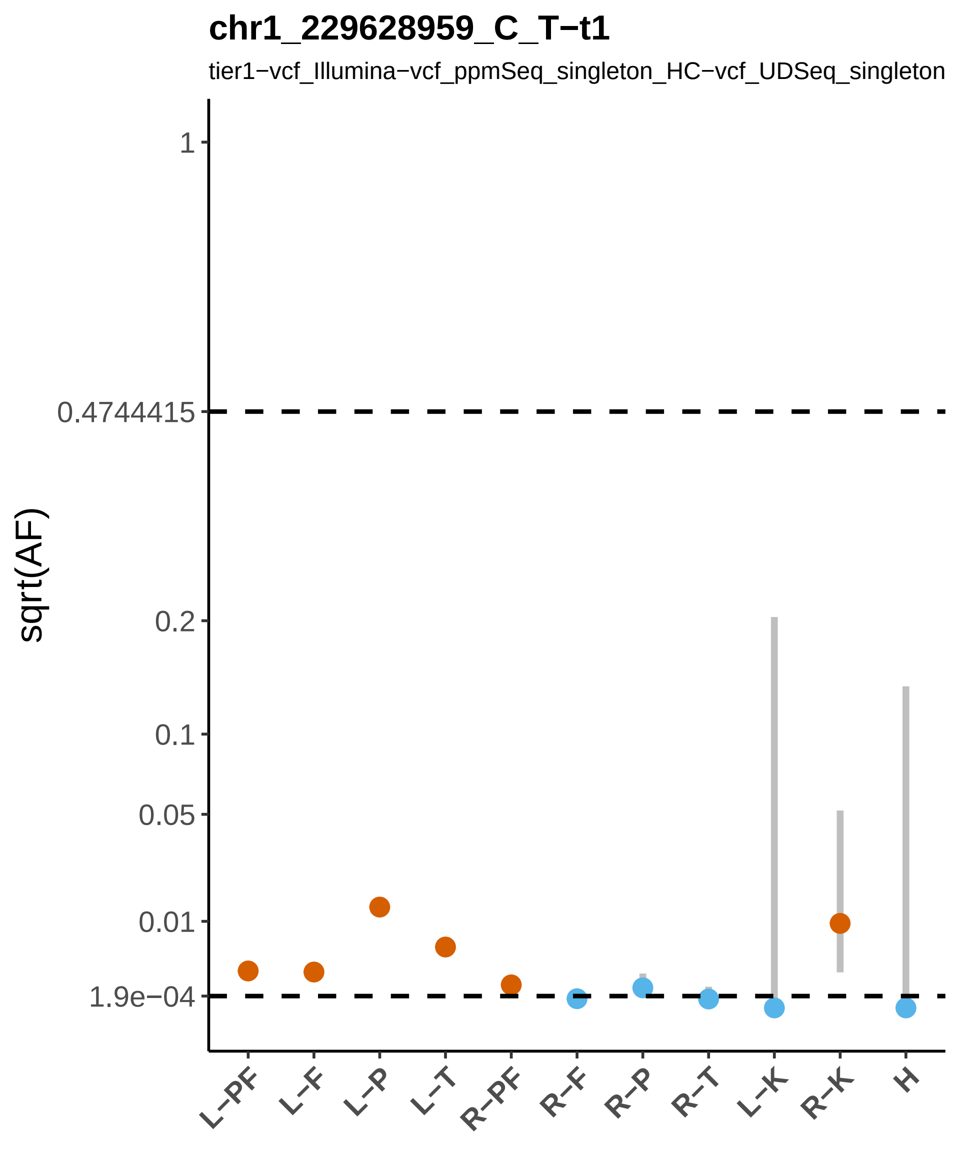
chr1_229628959_C_T−t1
tier1−vcf_Illumina−vcf_ppmSeq_singleton_HC−vcf_UDSeq_singleton
sqrt(AF)
1
0.4744415
0.2
0.1
0.05
0.01
1.9e−04
L−PF
L−F
L−P
L−T
R−PF
R−F
R−P
R−T
L−K
R−K
H

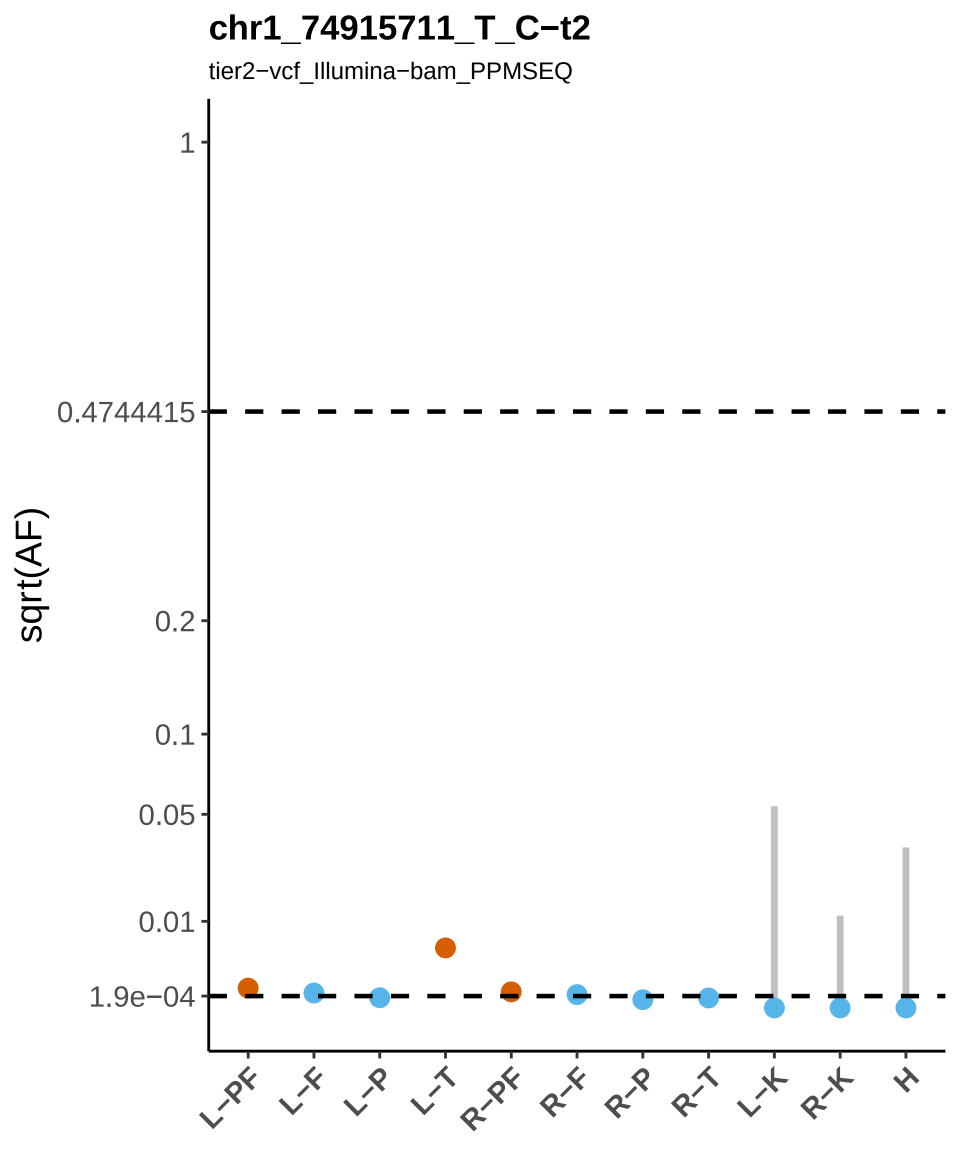

chr1_74915711_T_C−t2
tier2−vcf_Illumina−bam_PPMSEQ
sqrt(AF)
1
0.4744415
0.2
0.1
0.05
0.01
1.9e−04
L−PF
L−F
L−P
L−T
R−PF
R−F
R−P
R−T
L−K
R−K
H

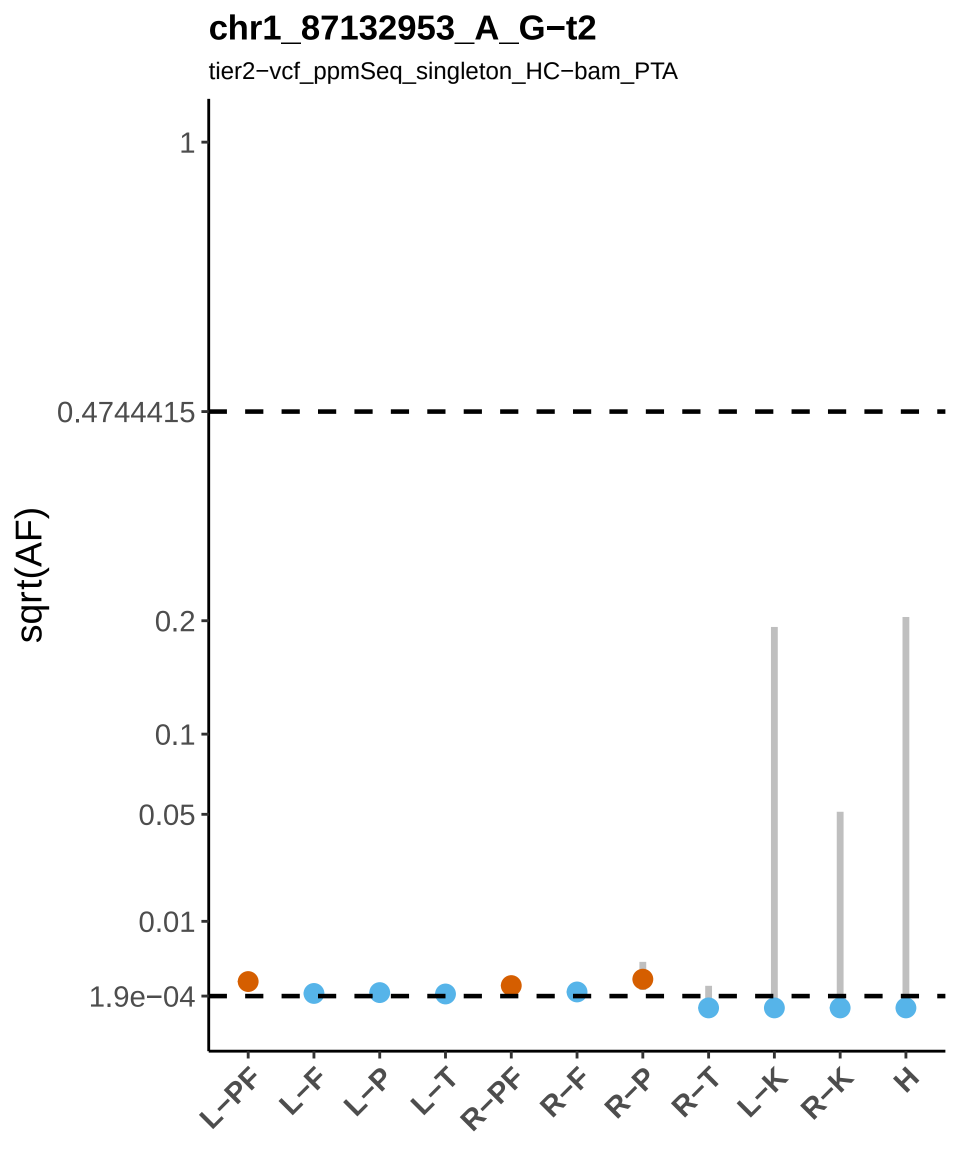
chr1_87132953_A_G−t2
tier2−vcf_ppmSeq_singleton_HC−bam_PTA
sqrt(AF)
1
0.4744415
0.2
0.1
0.05
0.01
1.9e−04
L−PF
L−F
L−P
L−T
R−PF
R−F
R−P
R−T
L−K
R−K
H

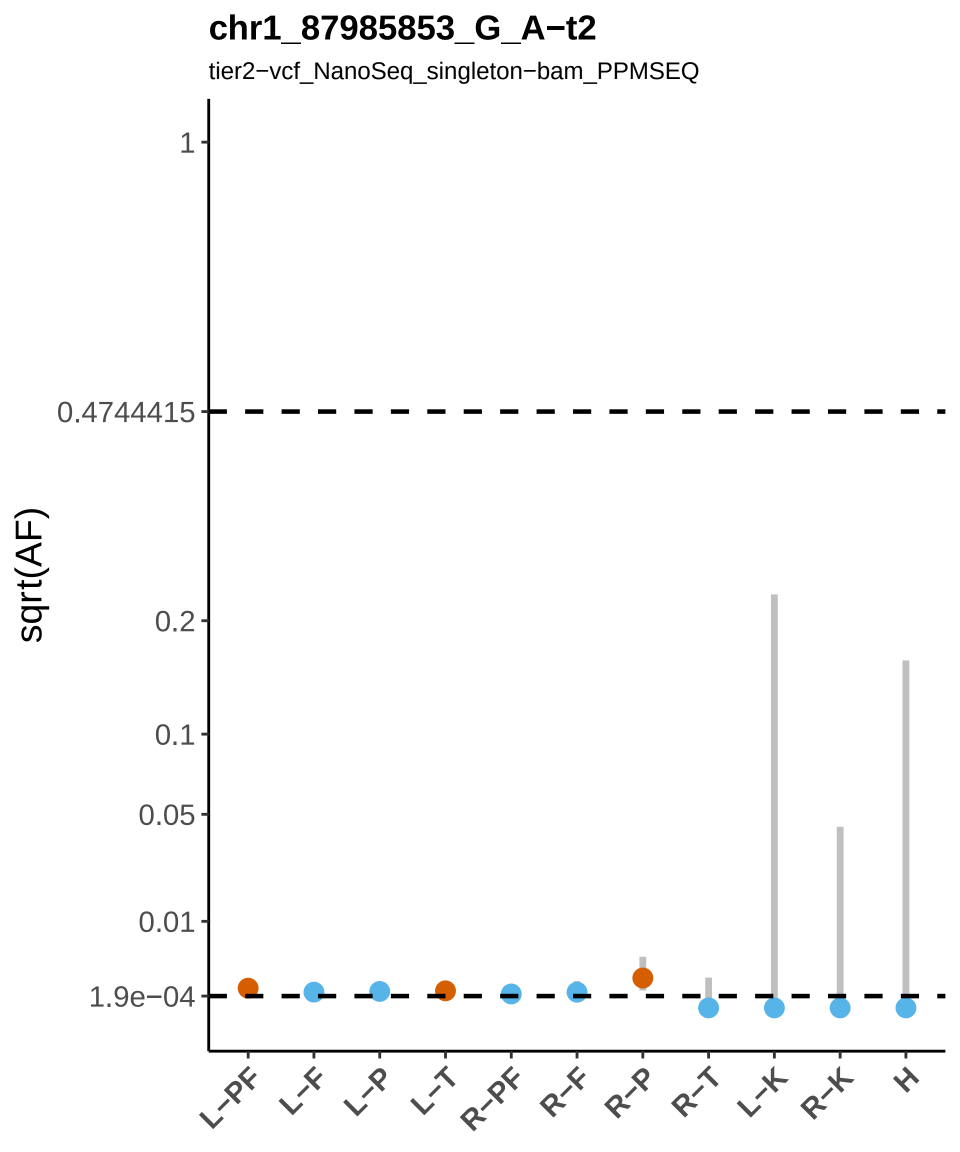

chr1_87985853_G_A−t2
tier2−vcf_NanoSeq_singleton−bam_PPMSEQ
sqrt(AF)
1
0.4744415
0.2
0.1
0.05
0.01
1.9e−04
L−PF
L−F
L−P
L−T
R−PF
R−F
R−P
R−T
L−K
R−K
H

#### chr1_88306738_G_A−t2

tier2−vcf_NanoSeq_singleton−bam_ILLUMINA

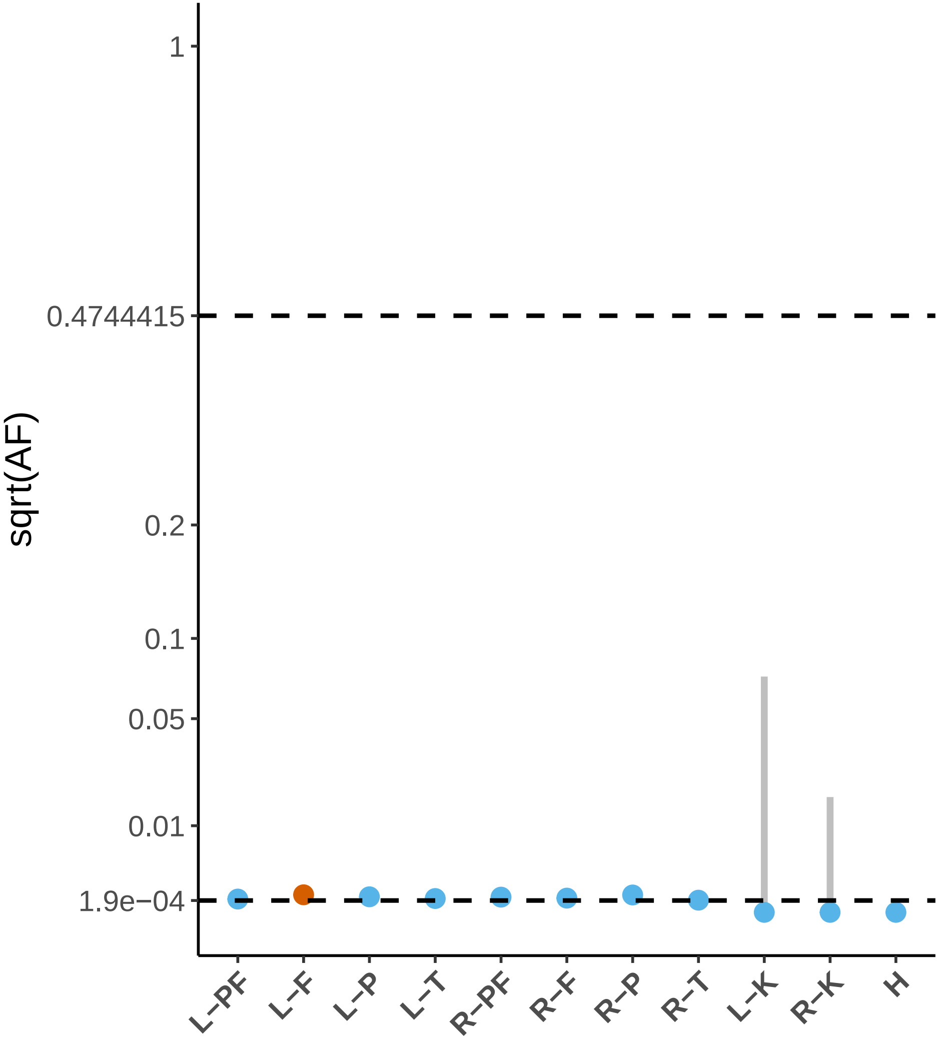

### chr1_88630096_C_T−t3

tier3−vcf_UDSeq_singleton

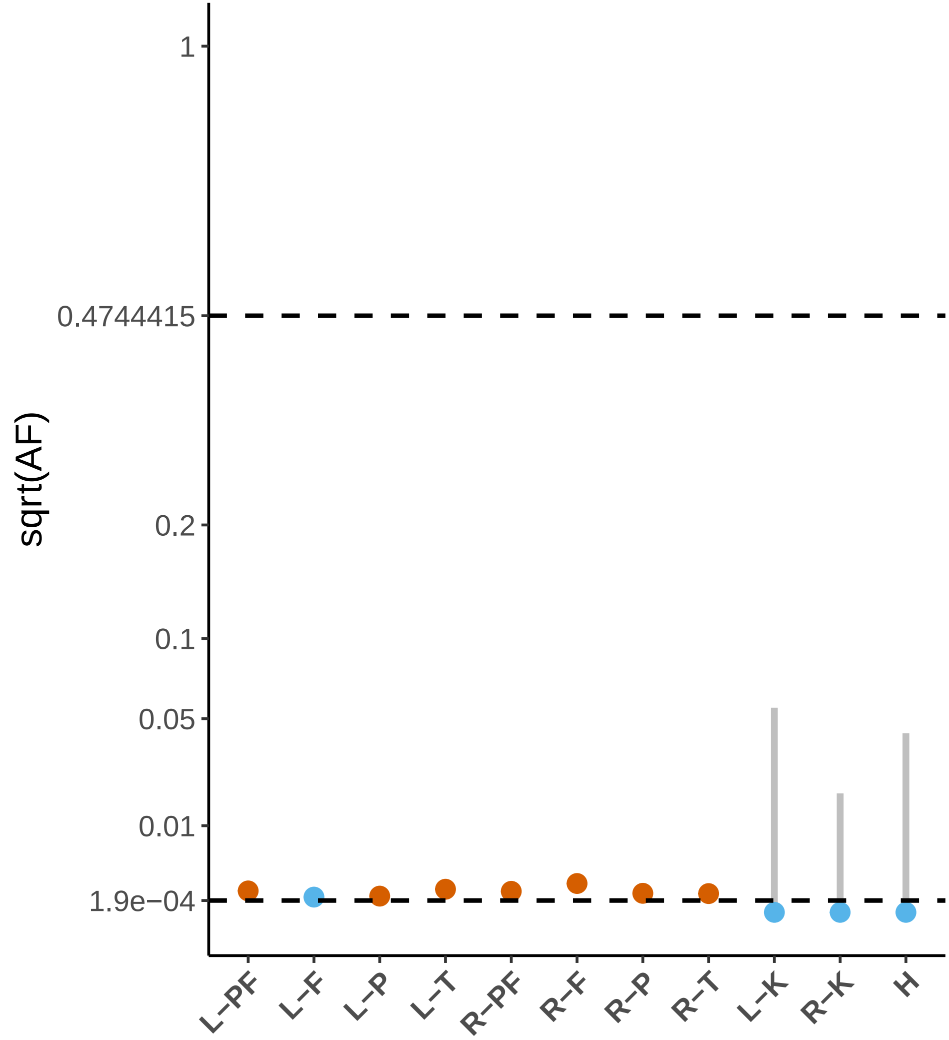

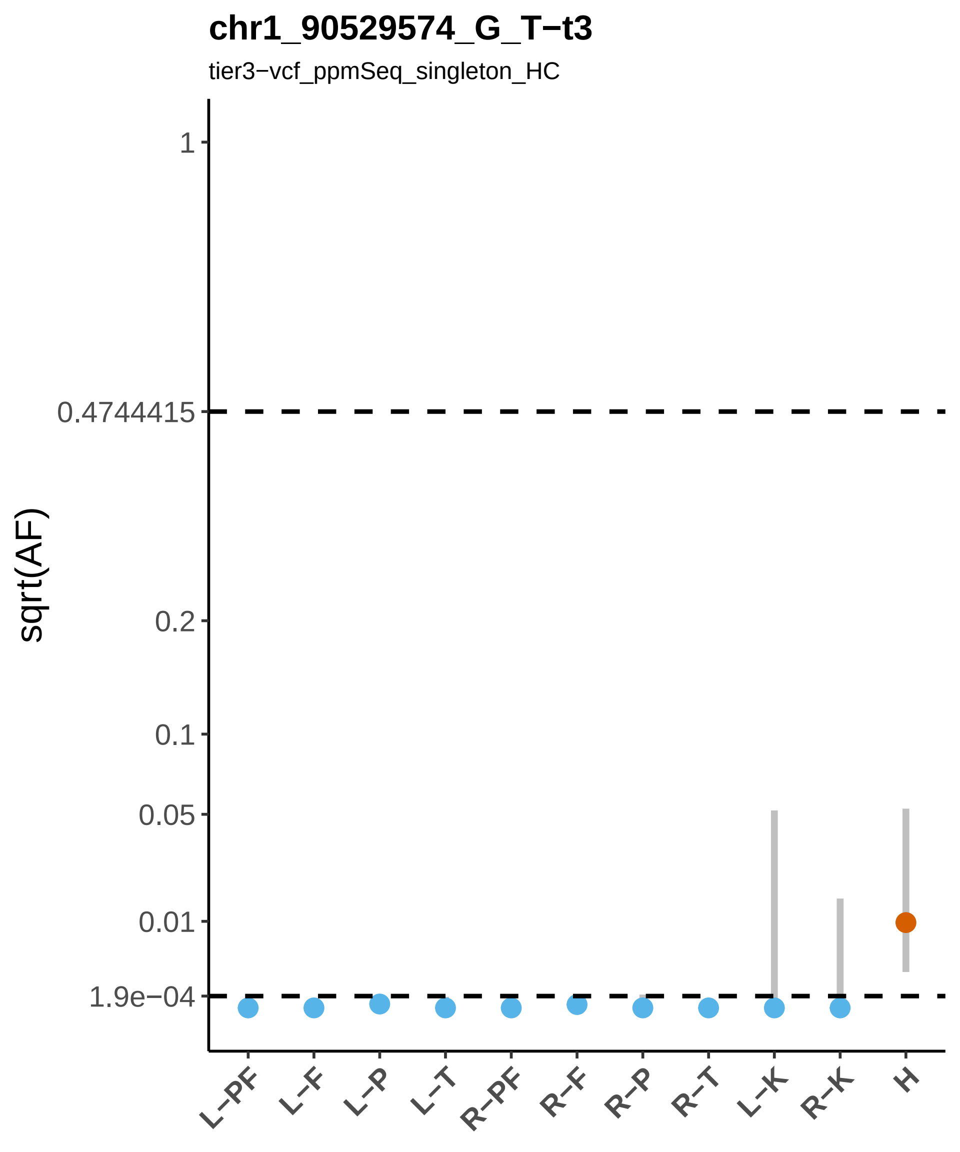
chr1_90529574_G_T−t3
tier3−vcf_ppmSeq_singleton_HC
sqrt(AF)
1
0.4744415
0.2
0.1
0.05
0.01
1.9e−04
L−PF
L−F
L−P
L−T
R−PF
R−F
R−P
R−T
L−K
R−K
H

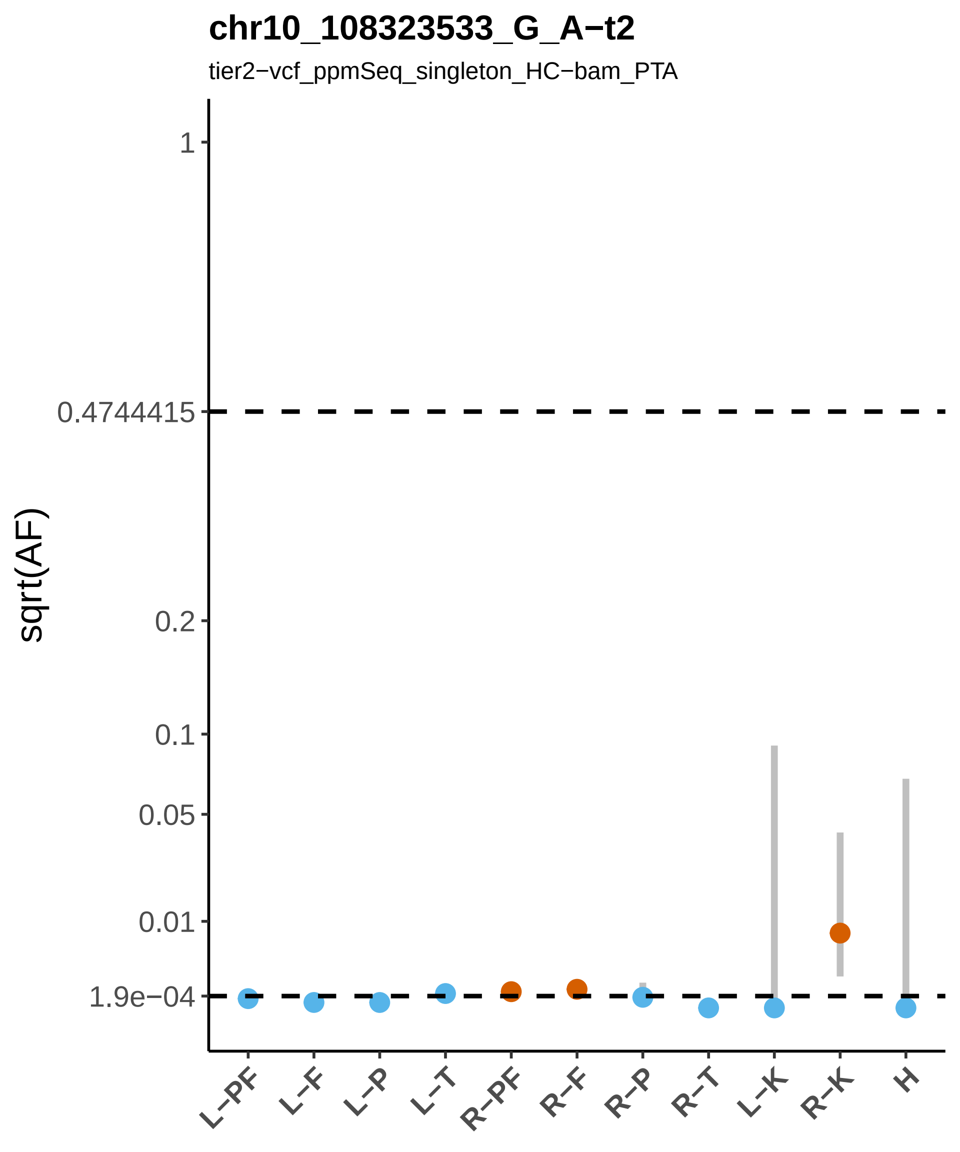

chr10_108323533_G_A−t2
tier2−vcf_ppmSeq_singleton_HC−bam_PTA
sqrt(AF)
1
0.4744415
0.2
0.1
0.05
0.01
1.9e−04
L−PF
L−F
L−P
L−T
R−PF
R−F
R−P
R−T
L−K
R−K
H

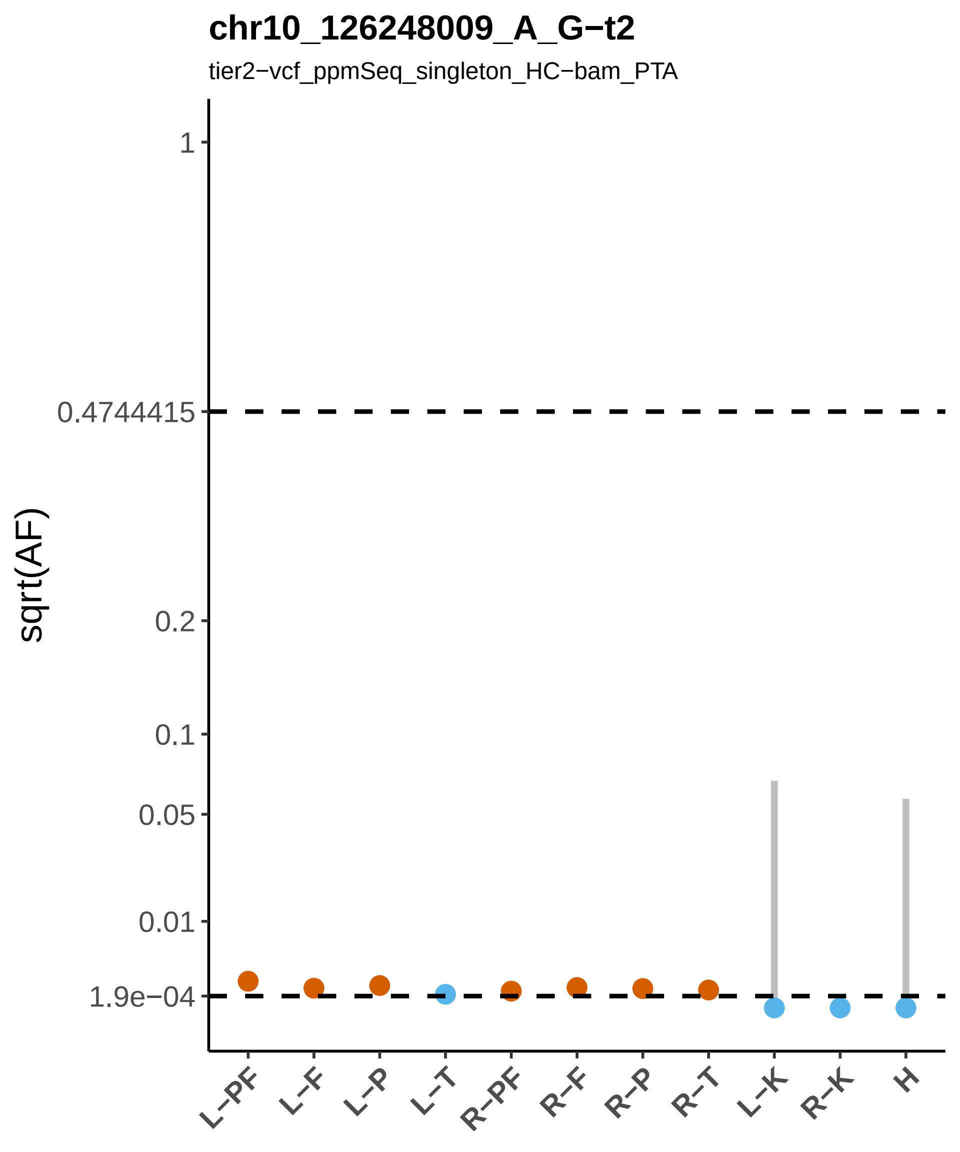
chr10_126248009_A_G−t2
tier2−vcf_ppmSeq_singleton_HC−bam_PTA
sqrt(AF)
1
0.4744415
0.2
0.1
0.05
0.01
1.9e−04
L−PF
L−F
L−P
L−T
R−PF
R−F
R−P
R−T
L−K
R−K
H

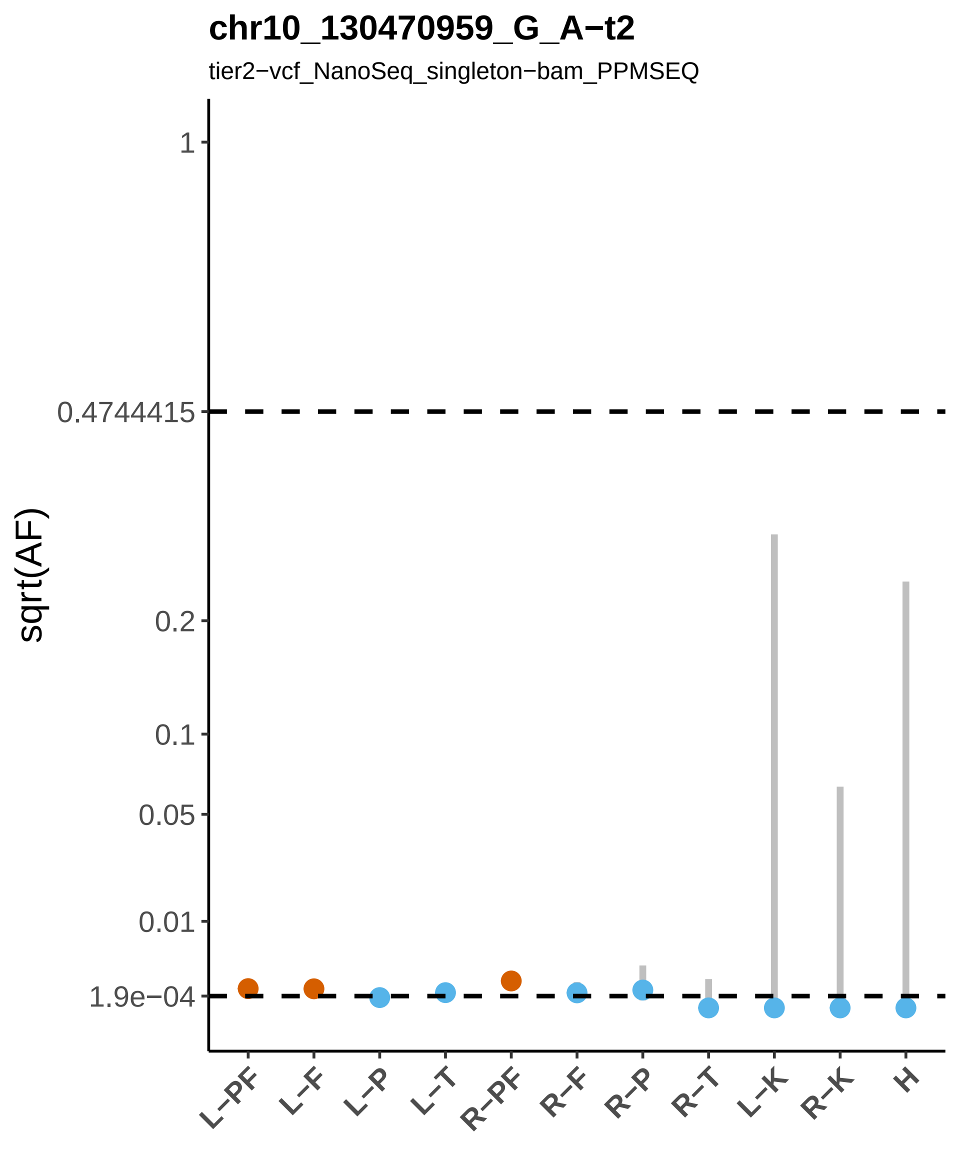
chr10_130470959_G_A−t2
tier2−vcf_NanoSeq_singleton−bam_PPMSEQ
sqrt(AF)
1
0.4744415
0.2
0.1
0.05
0.01
1.9e−04
L−PF
L−F
L−P
L−T
R−PF
R−F
R−P
R−T
L−K
R−K
H

**chr10_132043740_C_T−t2**

tier2−vcf_UDSeq_singleton−bam_PTA

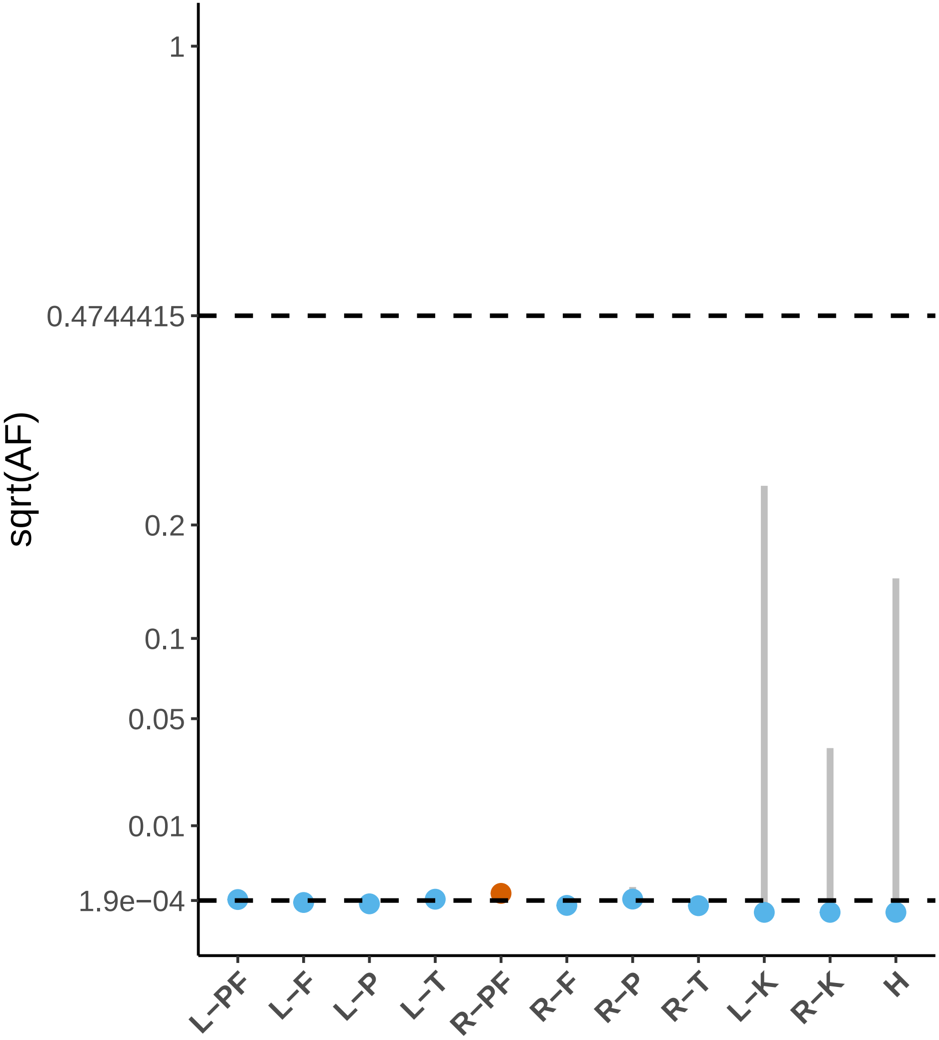

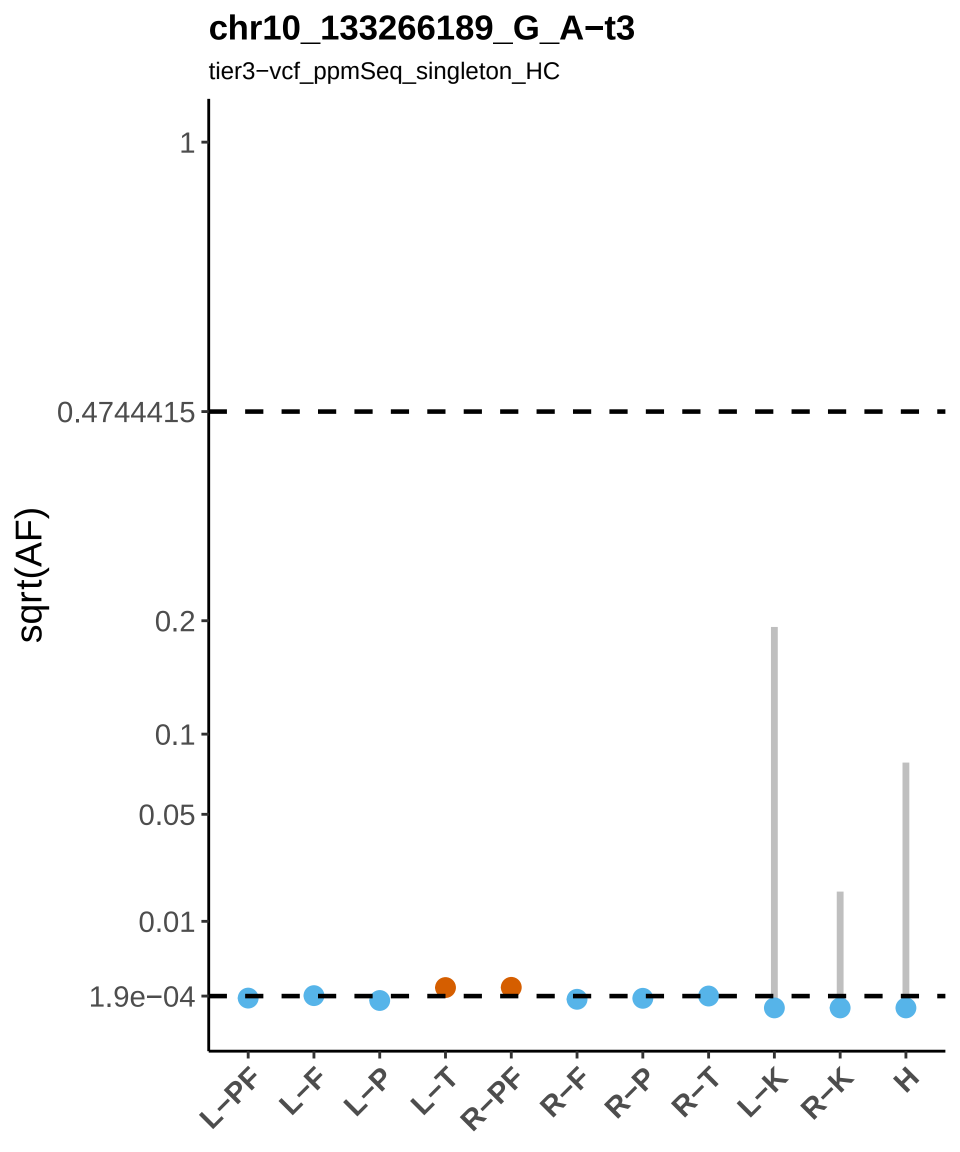
chr10_133266189_G_A−t3
tier3−vcf_ppmSeq_singleton_HC
sqrt(AF)
1
0.4744415
0.2
0.1
0.05
0.01
1.9e−04
L−PF
L−F
L−P
L−T
R−PF
R−F
R−P
R−T
L−K
R−K
H

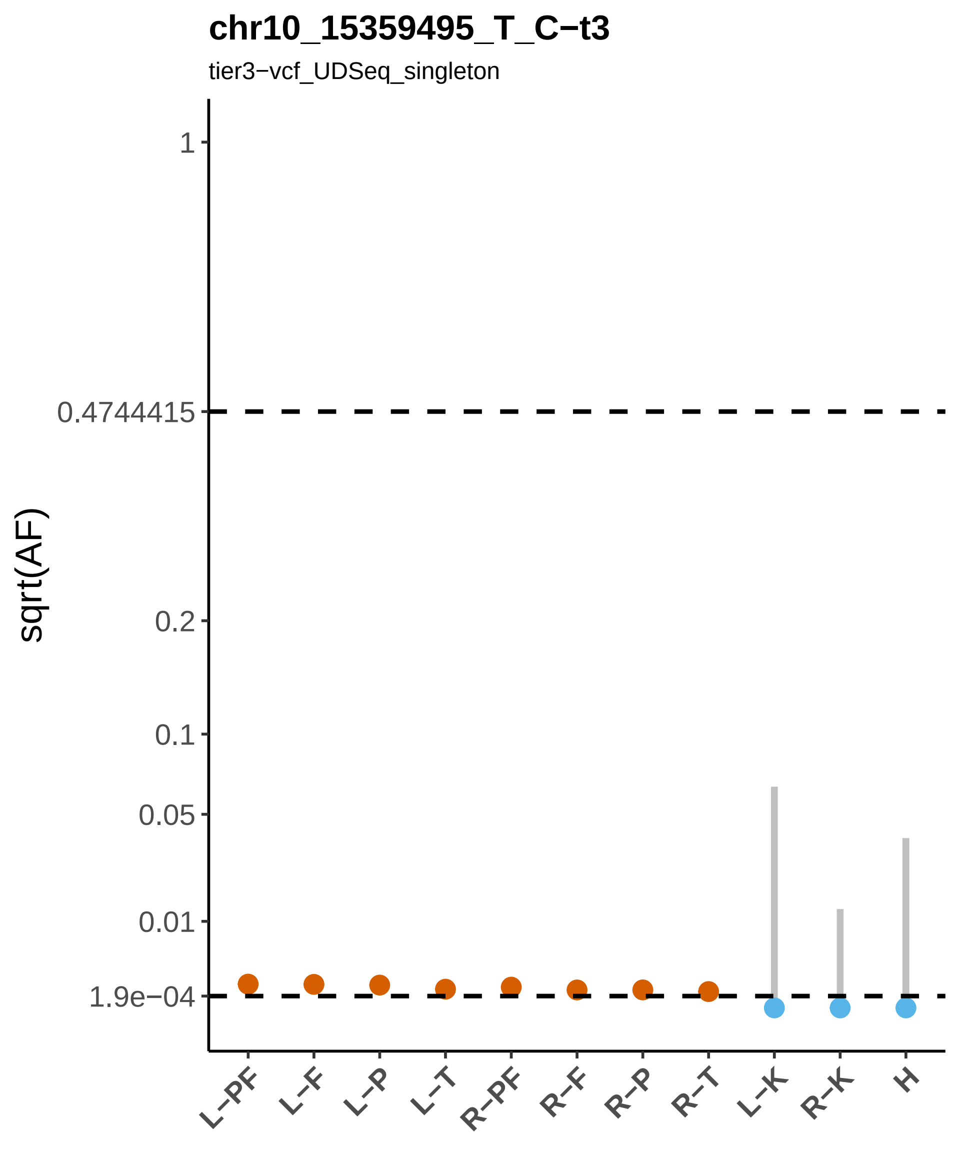

chr10_15359495_T_C−t3
tier3−vcf_UDSeq_singleton
sqrt(AF)
1
0.4744415
0.2
0.1
0.05
0.01
1.9e−04
L−PF
L−F
L−P
L−T
R−PF
R−F
R−P
R−T
L−K
R−K
H

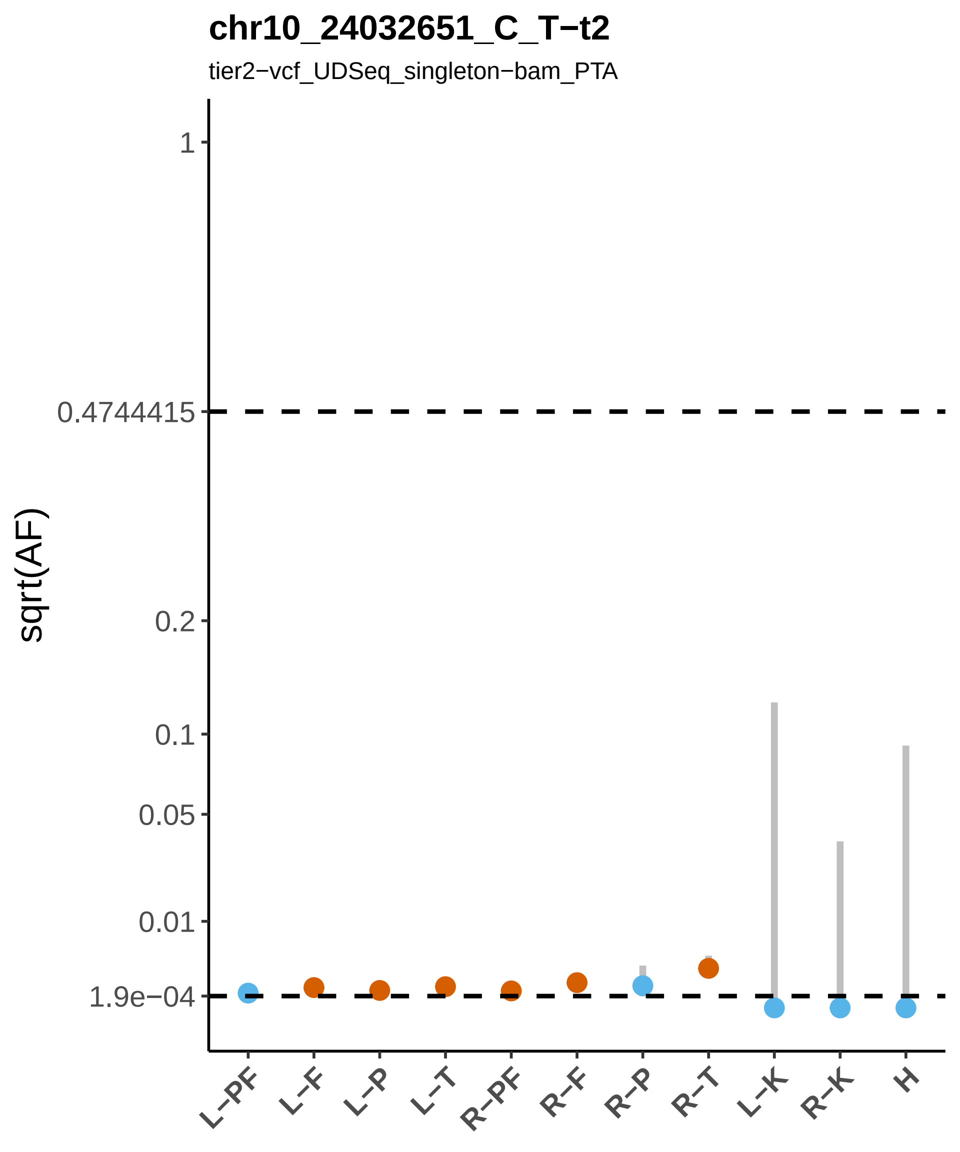

chr10_24032651_C_T−t2
tier2−vcf_UDSeq_singleton−bam_PTA
sqrt(AF)
1
0.4744415
0.2
0.1
0.05
0.01
1.9e−04
L−PF
L−F
L−P
L−T
R−PF
R−F
R−P
R−T
L−K
R−K
H

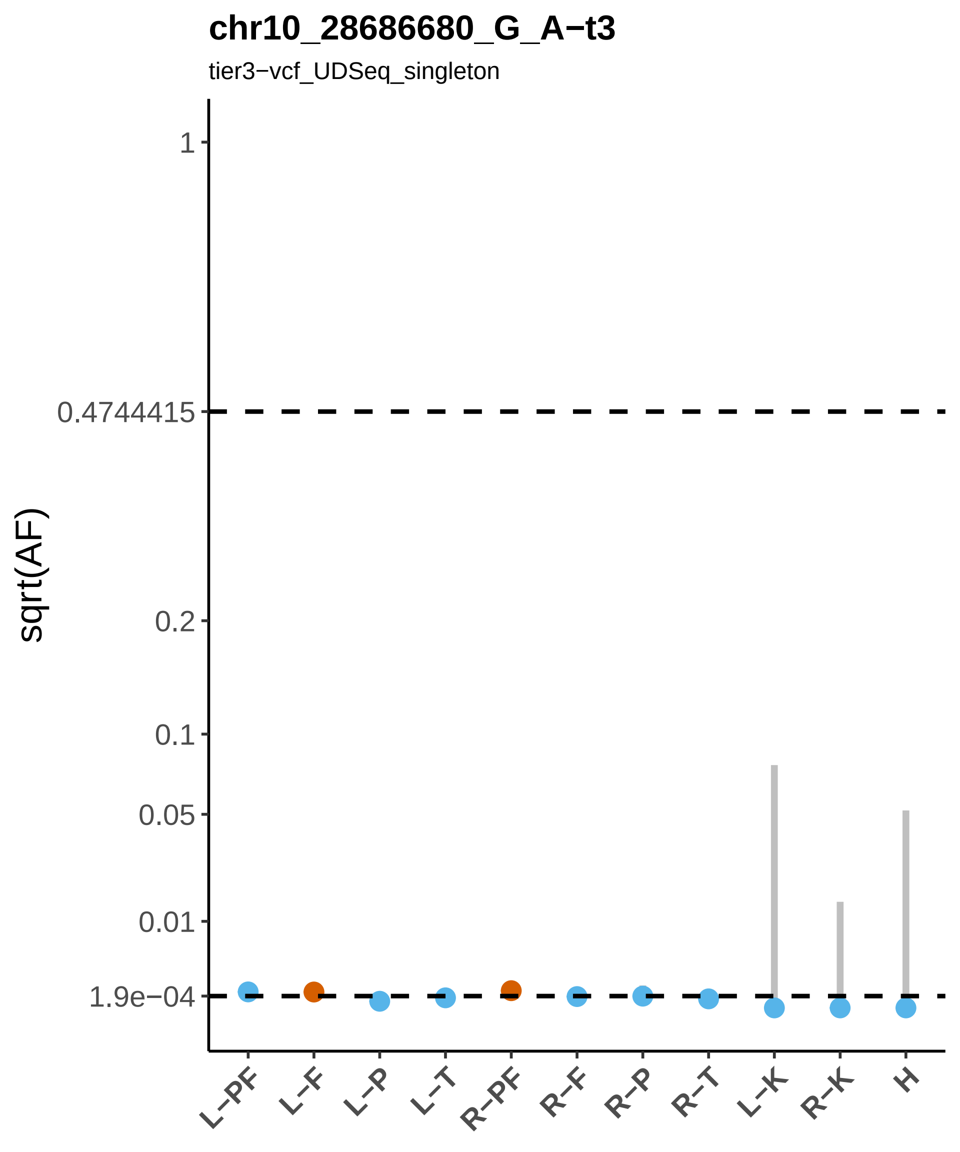

chr10_28686680_G_A−t3
tier3−vcf_UDSeq_singleton
sqrt(AF)
1
0.4744415
0.2
0.1
0.05
0.01
1.9e−04
L−PF
L−F
L−P
L−T
R−PF
R−F
R−P
R−T
L−K
R−K
H

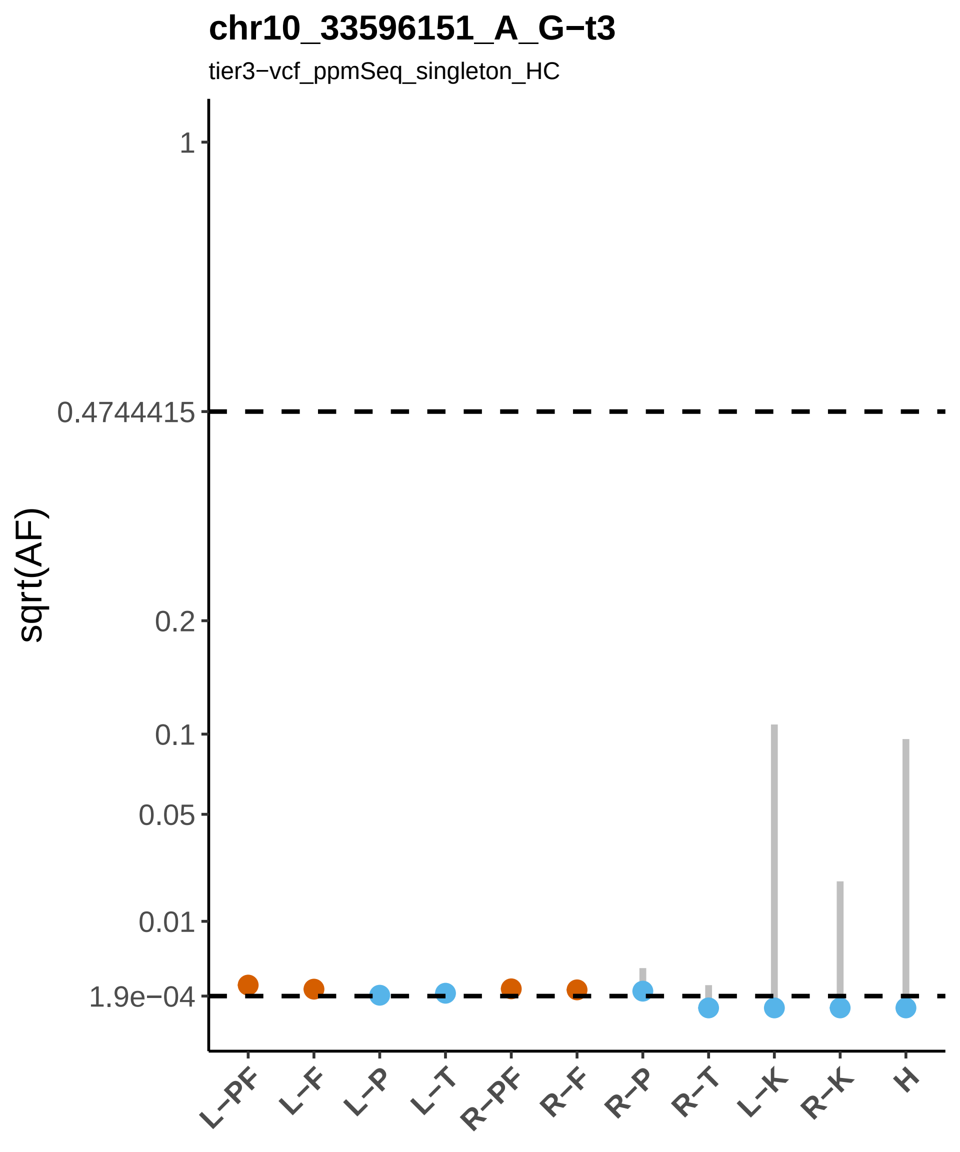
chr10_33596151_A_G−t3
tier3−vcf_ppmSeq_singleton_HC
sqrt(AF)
1
0.4744415
0.2
0.1
0.05
0.01
1.9e−04
L−PF
L−F
L−P
L−T
R−PF
R−F
R−P
R−T
L−K
R−K
H

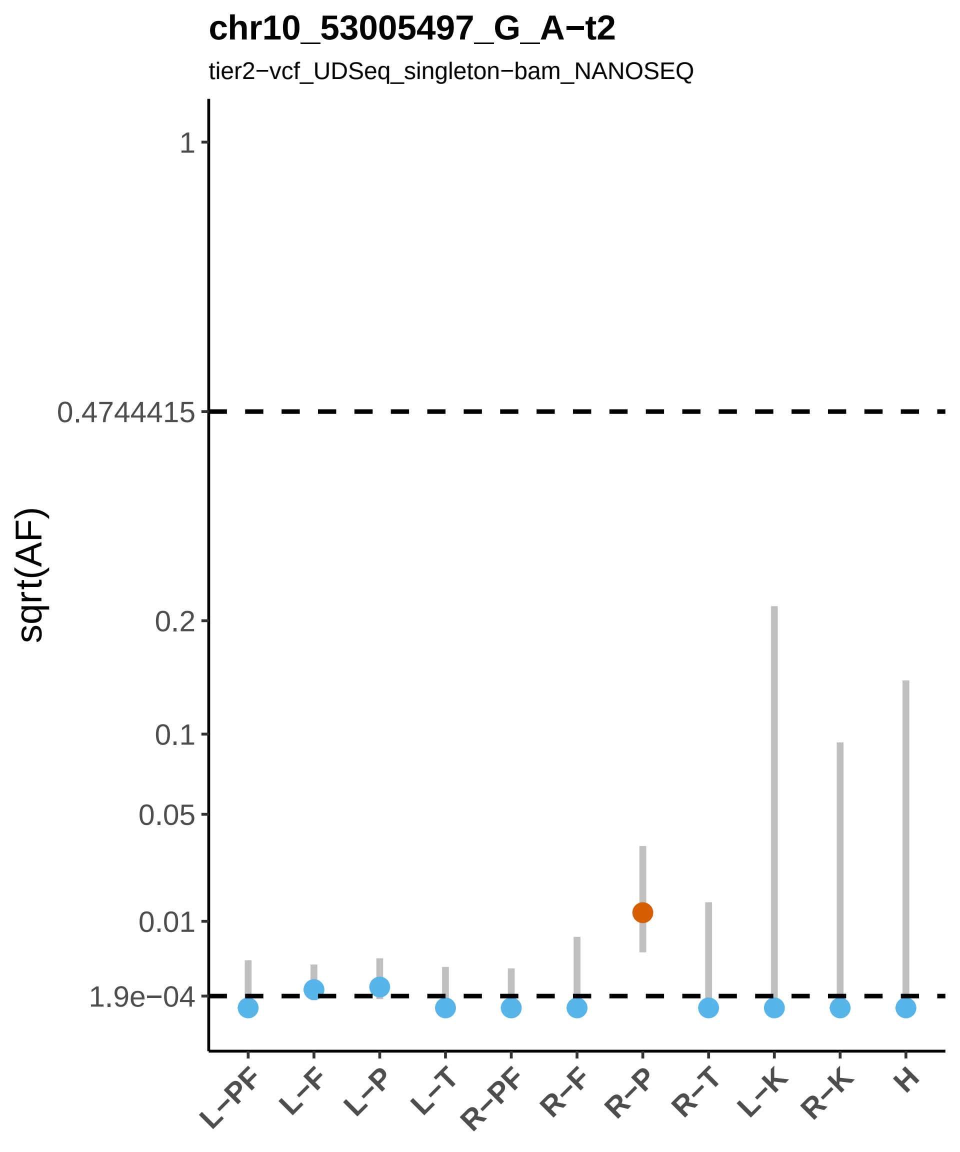

chr10_53005497_G_A−t2
tier2−vcf_UDSeq_singleton−bam_NANOSEQ
sqrt(AF)
1
0.4744415
0.2
0.1
0.05
0.01
1.9e−04
L−PF
L−F
L−P
L−T
R−PF
R−F
R−P
R−T
L−K
R−K
H

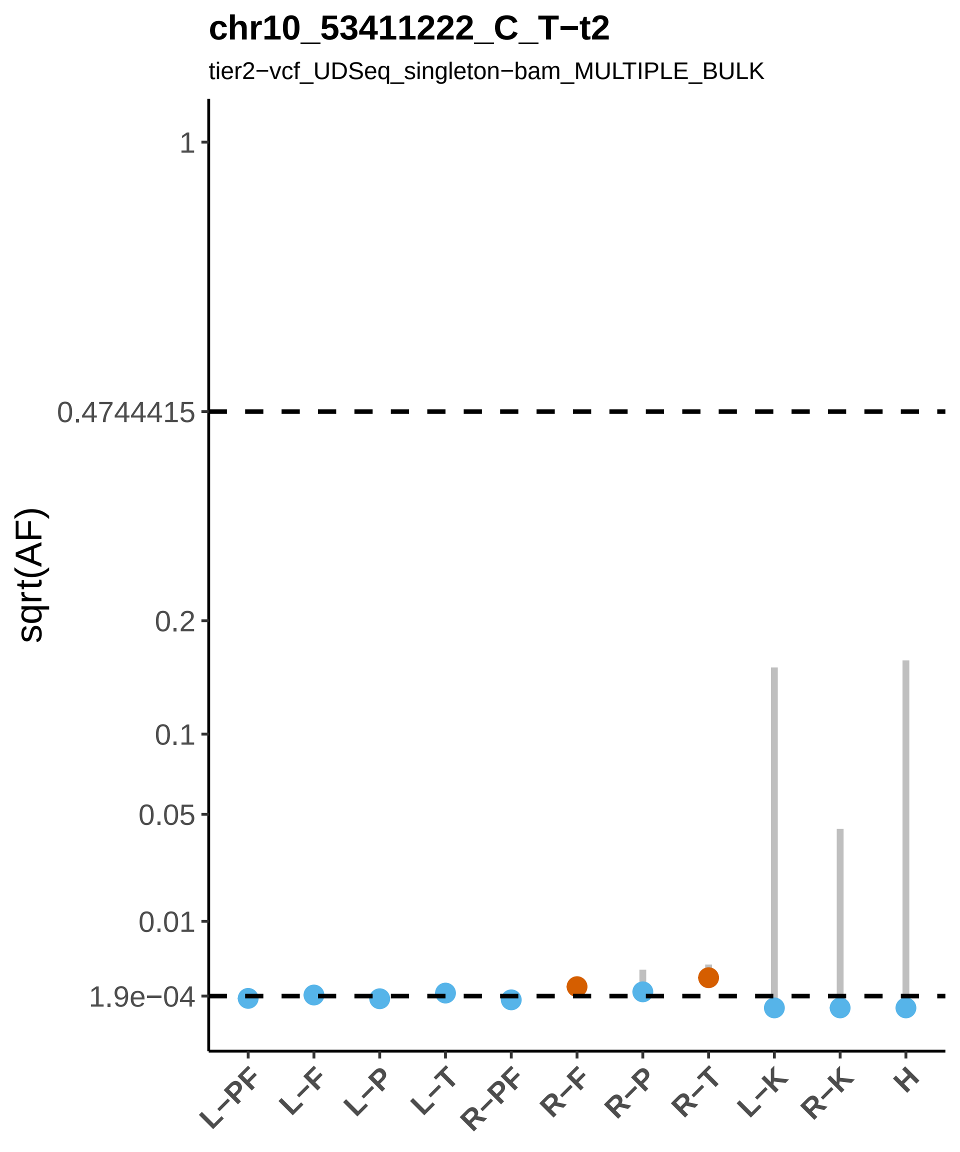

chr10_53411222_C_T−t2
tier2−vcf_UDSeq_singleton−bam_MULTIPLE_BULK
sqrt(AF)
1
0.4744415
0.2
0.1
0.05
0.01
1.9e−04
L−PF
L−F
L−P
L−T
R−PF
R−F
R−P
R−T
L−K
R−K
H

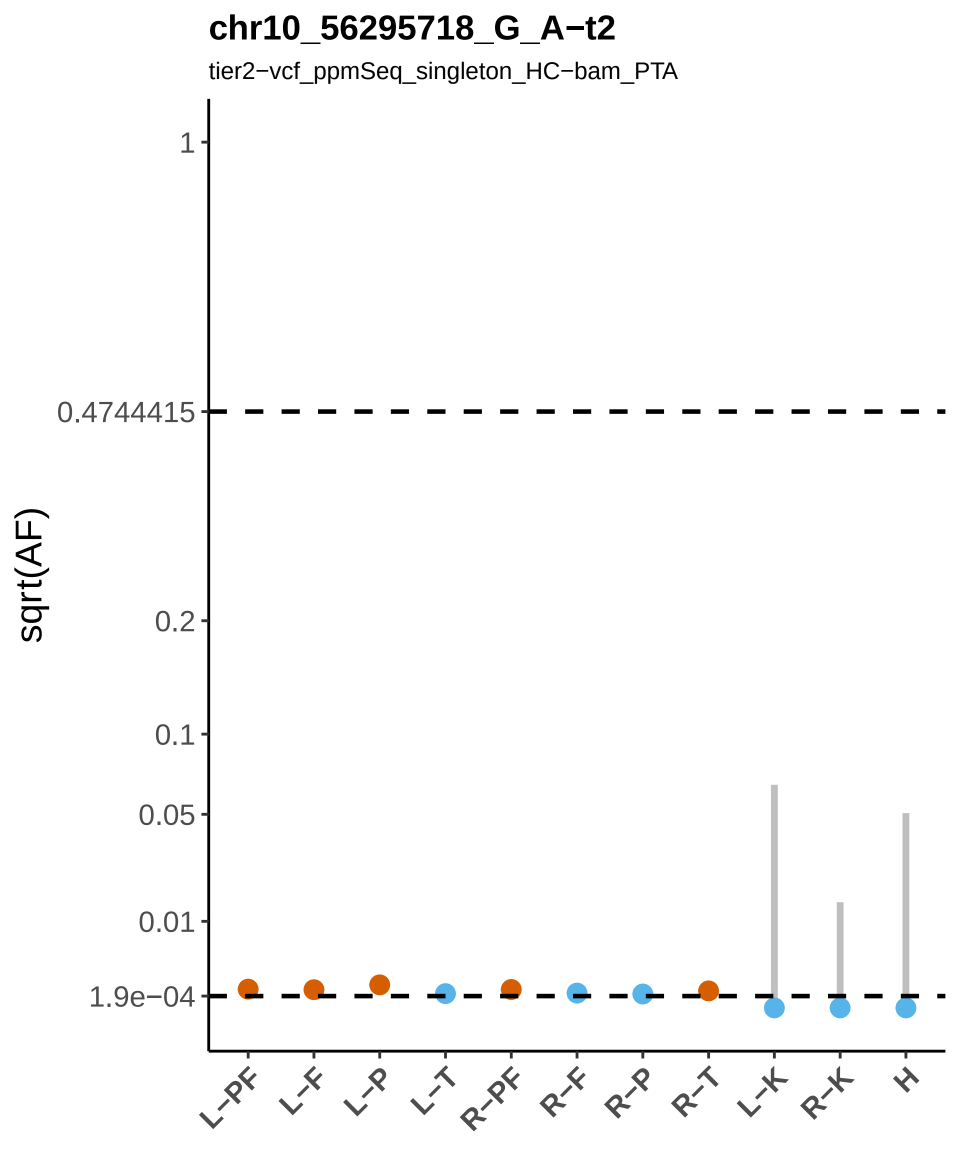

chr10_56295718_G_A−t2
tier2−vcf_ppmSeq_singleton_HC−bam_PTA
sqrt(AF)
1
0.4744415
0.2
0.1
0.05
0.01
1.9e−04
L−PF
L−F
L−P
L−T
R−PF
R−F
R−P
R−T
L−K
R−K
H

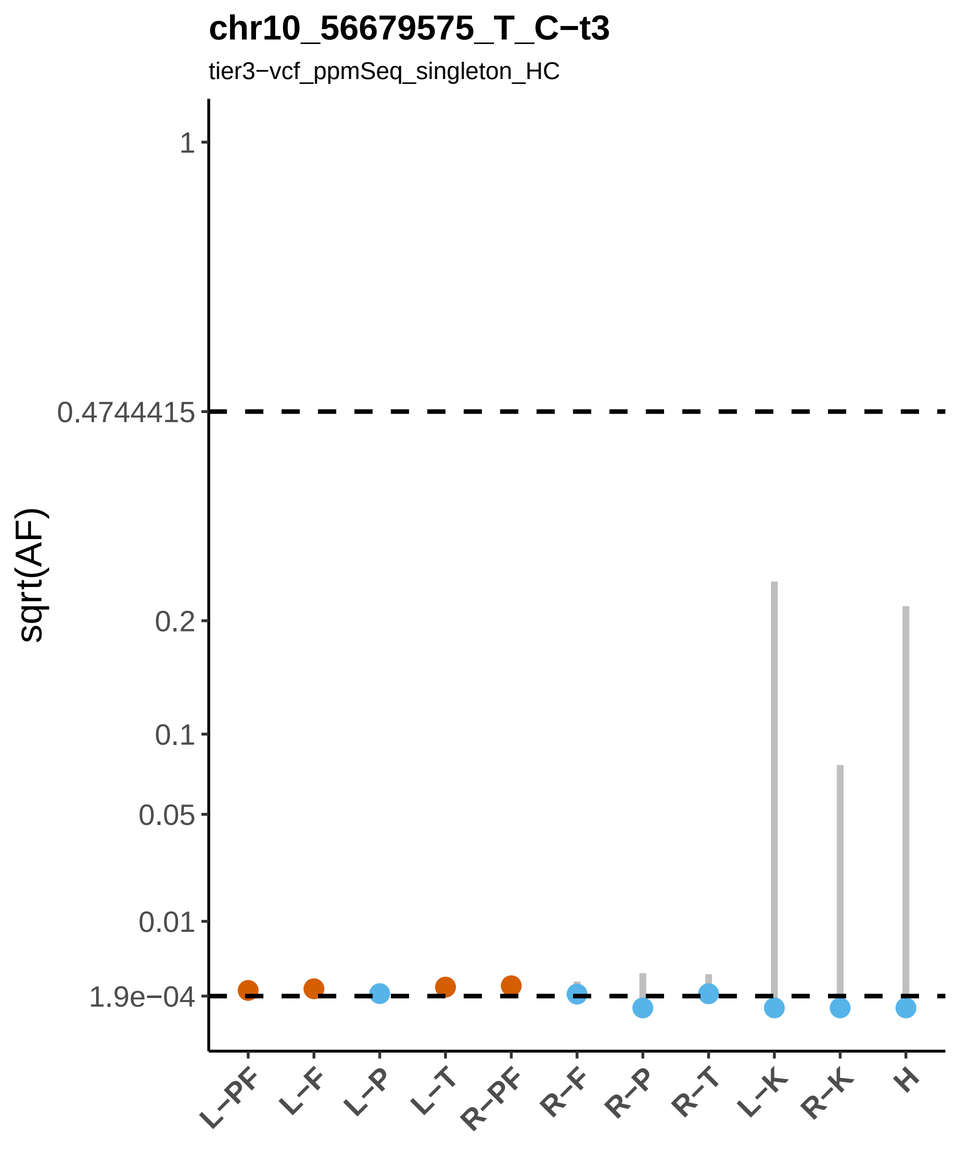
chr10_56679575_T_C−t3
tier3−vcf_ppmSeq_singleton_HC
sqrt(AF)
1
0.4744415
0.2
0.1
0.05
0.01
1.9e−04
L−PF
L−F
L−P
L−T
R−PF
R−F
R−P
R−T
L−K
R−K
H

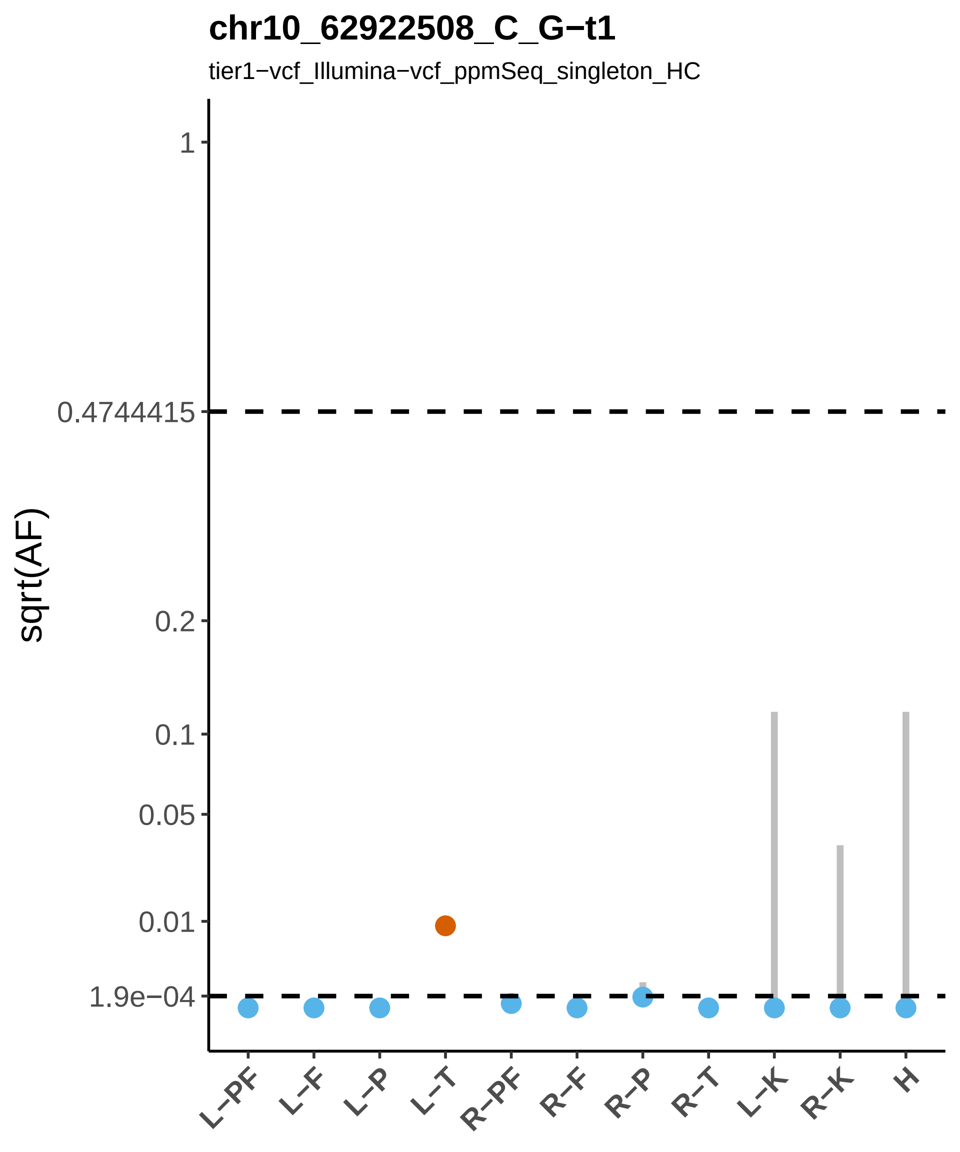
chr10_62922508_C_G−t1
tier1−vcf_Illumina−vcf_ppmSeq_singleton_HC
sqrt(AF)
1
0.4744415
0.2
0.1
0.05
0.01
1.9e−04
L−PF
L−F
L−P
L−T
R−PF
R−F
R−P
R−T
L−K
R−K
H

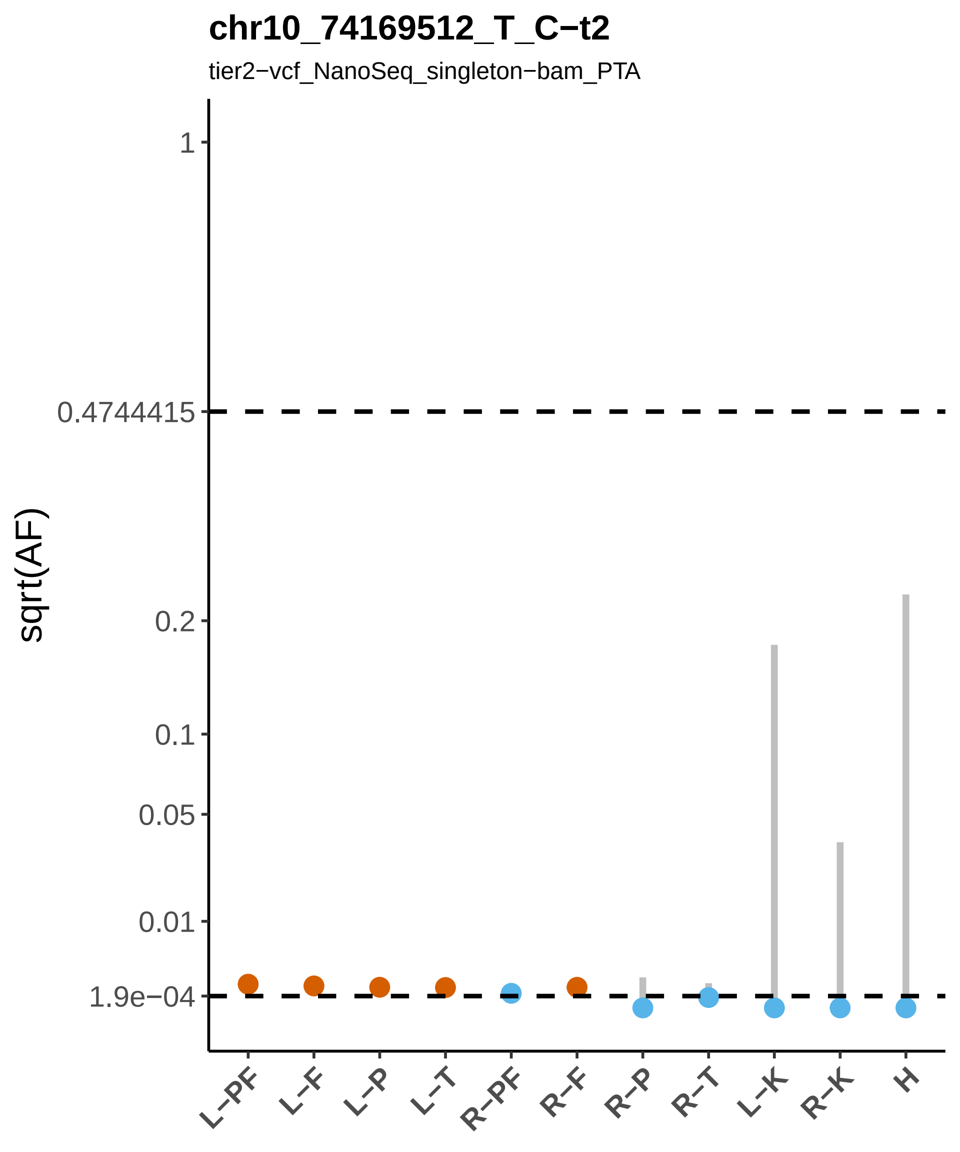

chr10_74169512_T_C−t2
tier2−vcf_NanoSeq_singleton−bam_PTA
sqrt(AF)
1
0.4744415
0.2
0.1
0.05
0.01
1.9e−04
L−PF
L−F
L−P
L−T
R−PF
R−F
R−P
R−T
L−K
R−K
H

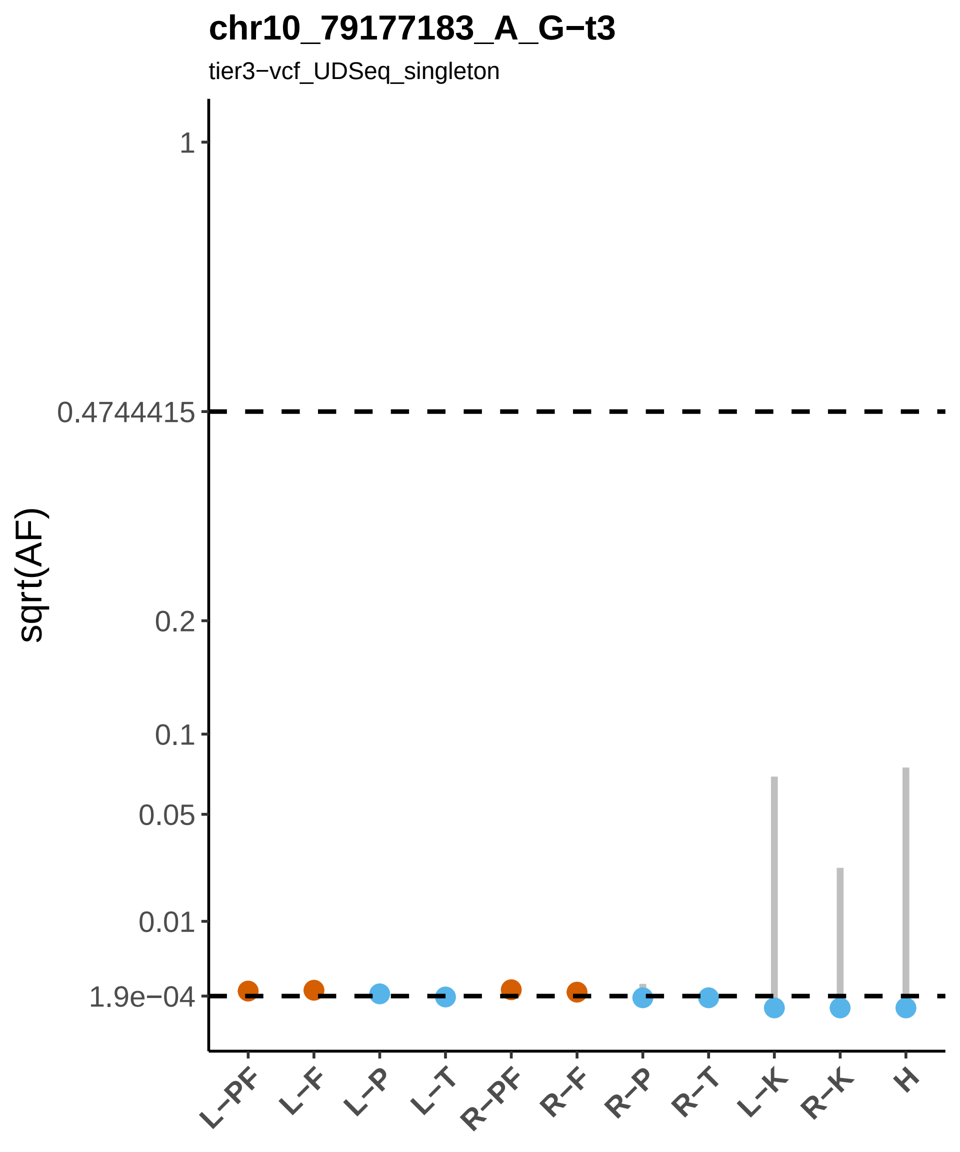

chr10_79177183_A_G−t3
tier3−vcf_UDSeq_singleton
sqrt(AF)
1
0.4744415
0.2
0.1
0.05
0.01
1.9e−04
L−PF
L−F
L−P
L−T
R−PF
R−F
R−P
R−T
L−K
R−K
H

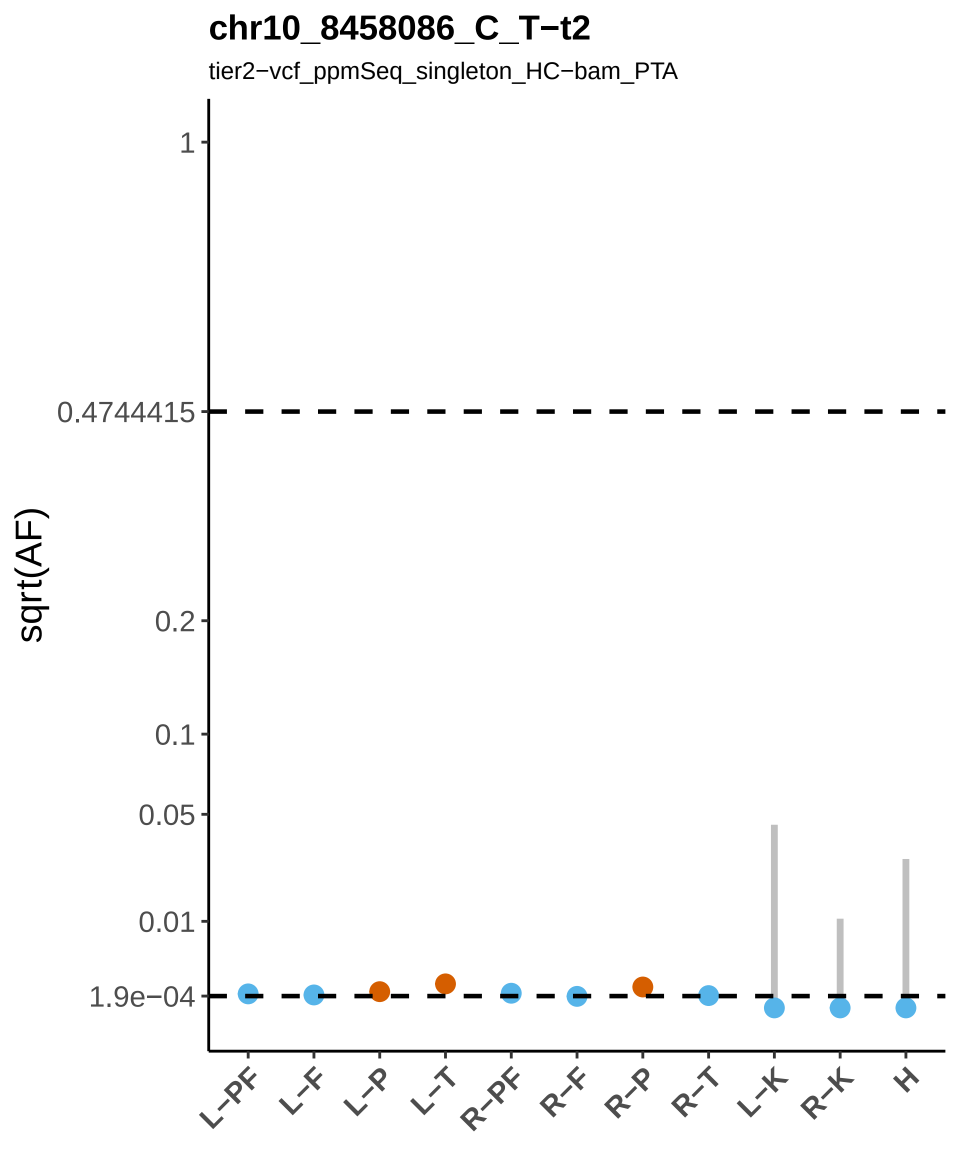

chr10_8458086_C_T−t2
tier2−vcf_ppmSeq_singleton_HC−bam_PTA
sqrt(AF)
1
0.4744415
0.2
0.1
0.05
0.01
1.9e−04
L−PF
L−F
L−P
L−T
R−PF
R−F
R−P
R−T
L−K
R−K
H

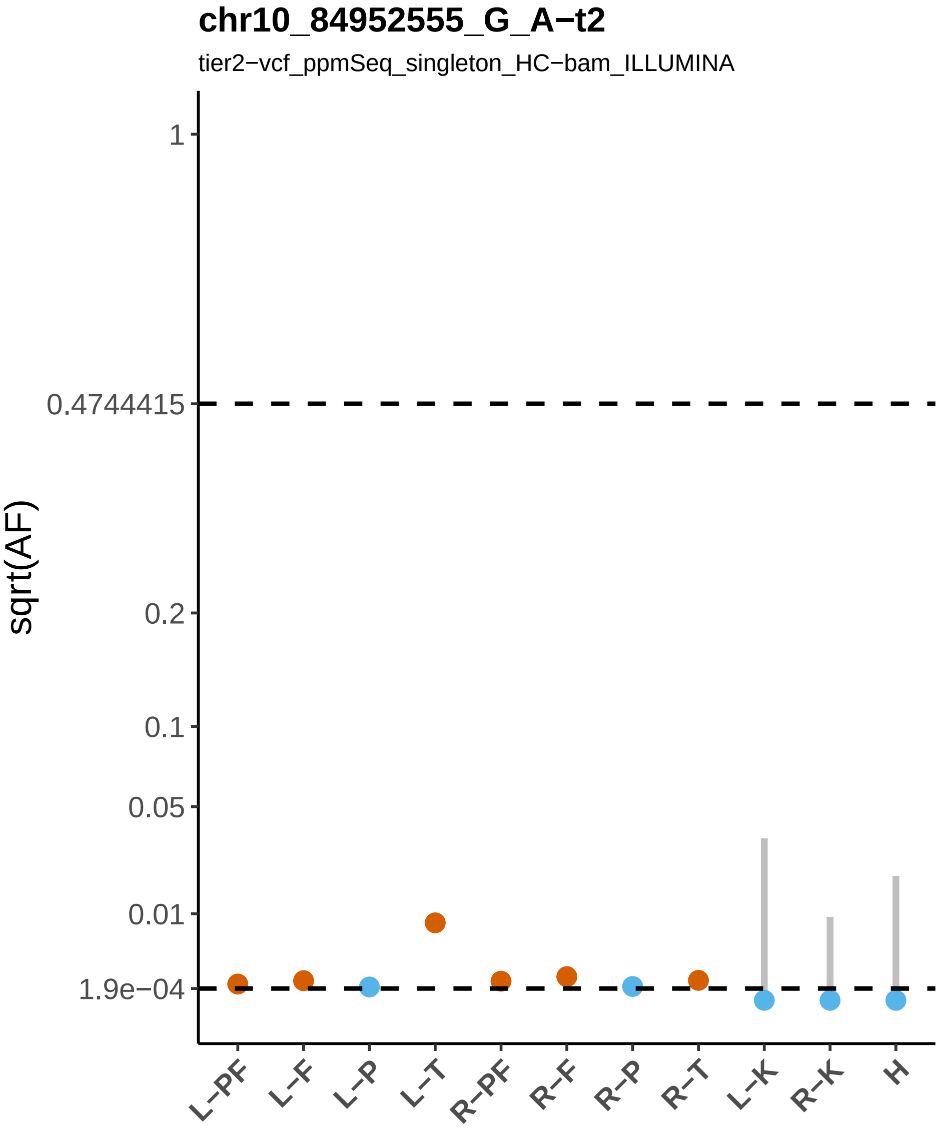

chr10_84952555_G_A−t2
tier2−vcf_ppmSeq_singleton_HC−bam_ILLUMINA
sqrt(AF)
1
0.4744415
0.2
0.1
0.05
0.01
1.9e−04
L−PF
L−F
L−P
L−T
R−PF
R−F
R−P
R−T
L−K
R−K
H

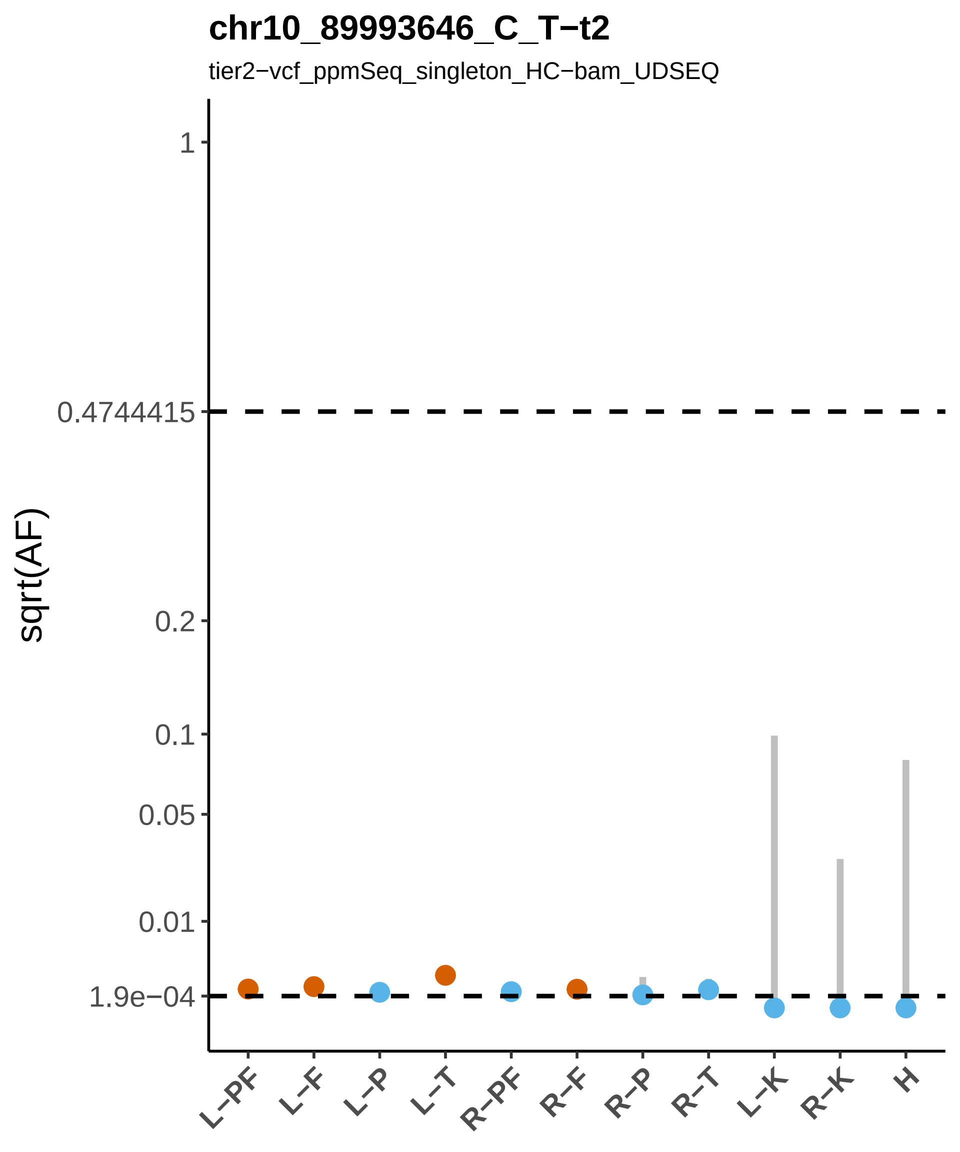

chr10_89993646_C_T−t2
tier2−vcf_ppmSeq_singleton_HC−bam_UDSEQ
sqrt(AF)
1
0.4744415
0.2
0.1
0.05
0.01
1.9e−04
L−PF
L−F
L−P
L−T
R−PF
R−F
R−P
R−T
L−K
R−K
H

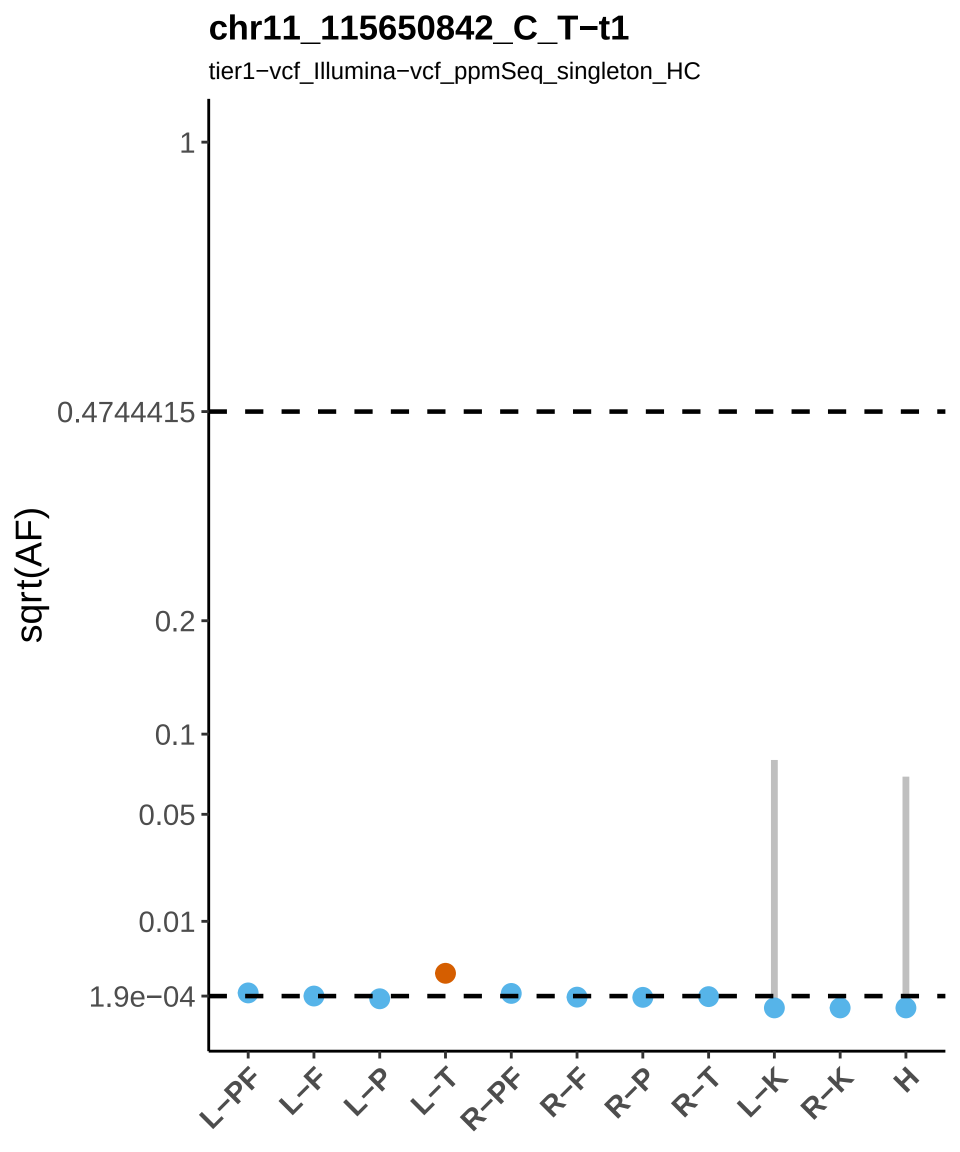

chr11_115650842_C_T−t1
tier1−vcf_Illumina−vcf_ppmSeq_singleton_HC
sqrt(AF)
1
0.4744415
0.2
0.1
0.05
0.01
1.9e−04
L−PF
L−F
L−P
L−T
R−PF
R−F
R−P
R−T
L−K
R−K
H

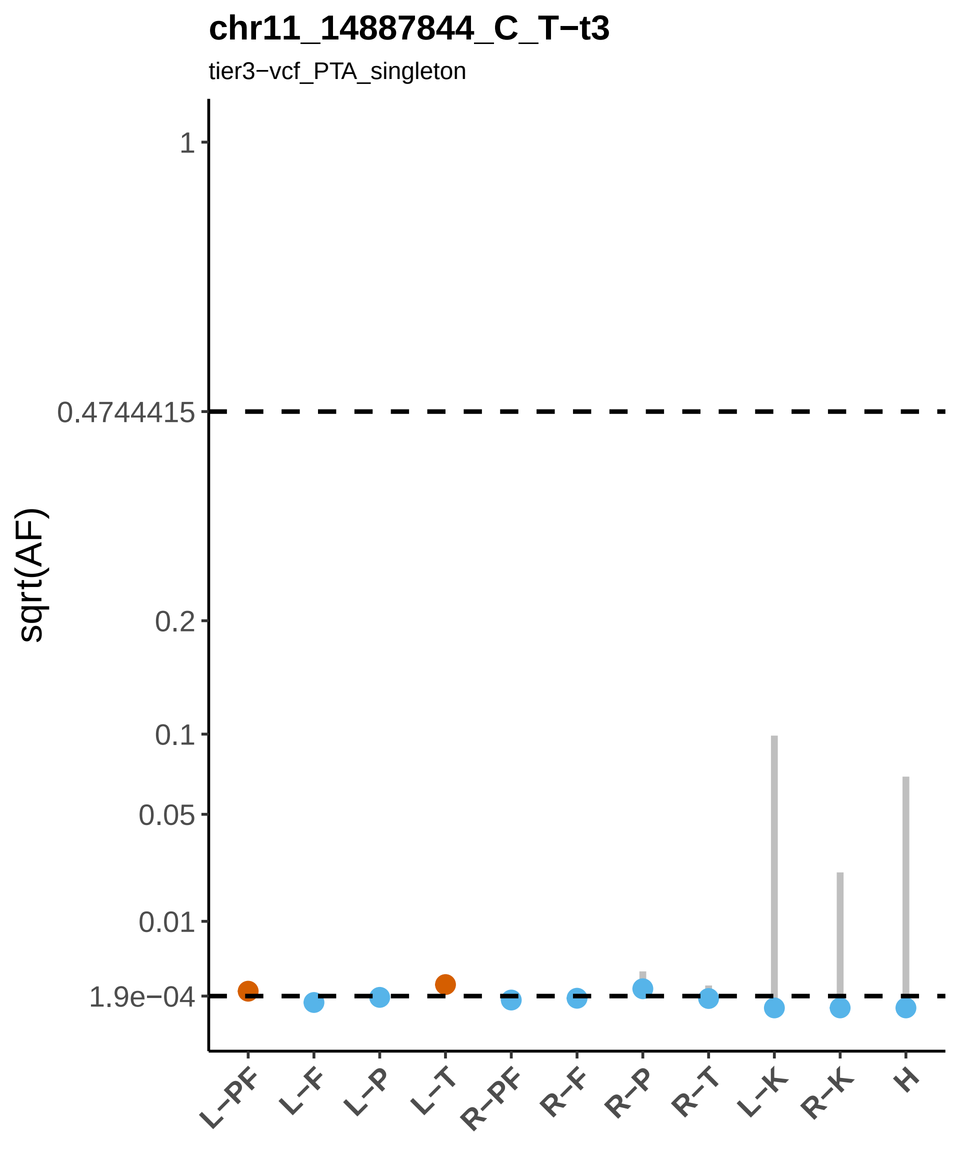

chr11_14887844_C_T−t3
tier3−vcf_PTA_singleton
sqrt(AF)
1
0.4744415
0.2
0.1
0.05
0.01
1.9e−04
L−PF
L−F
L−P
L−T
R−PF
R−F
R−P
R−T
L−K
R−K
H

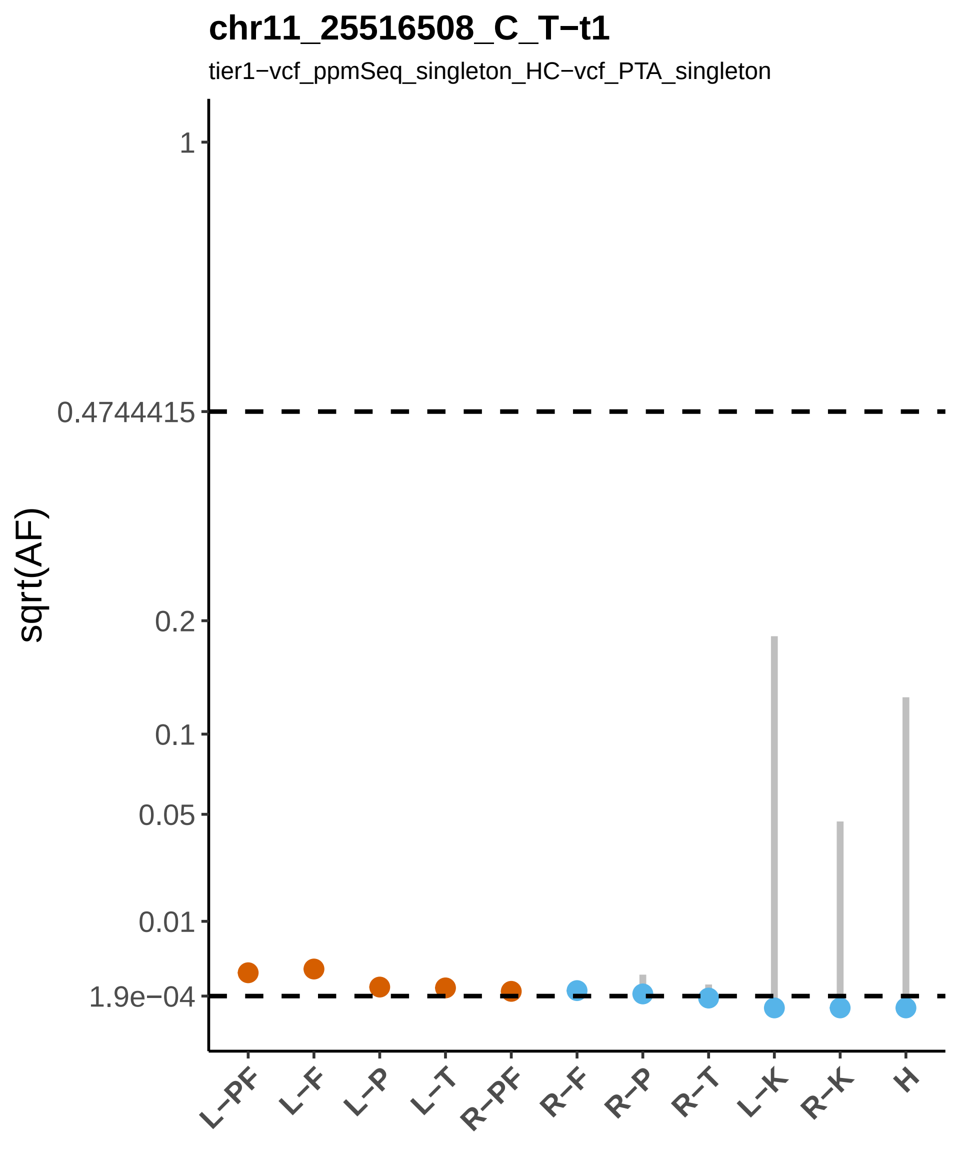

chr11_25516508_C_T−t1
tier1−vcf_ppmSeq_singleton_HC−vcf_PTA_singleton
sqrt(AF)
1
0.4744415
0.2
0.1
0.05
0.01
1.9e−04
L−PF
L−F
L−P
L−T
R−PF
R−F
R−P
R−T
L−K
R−K
H

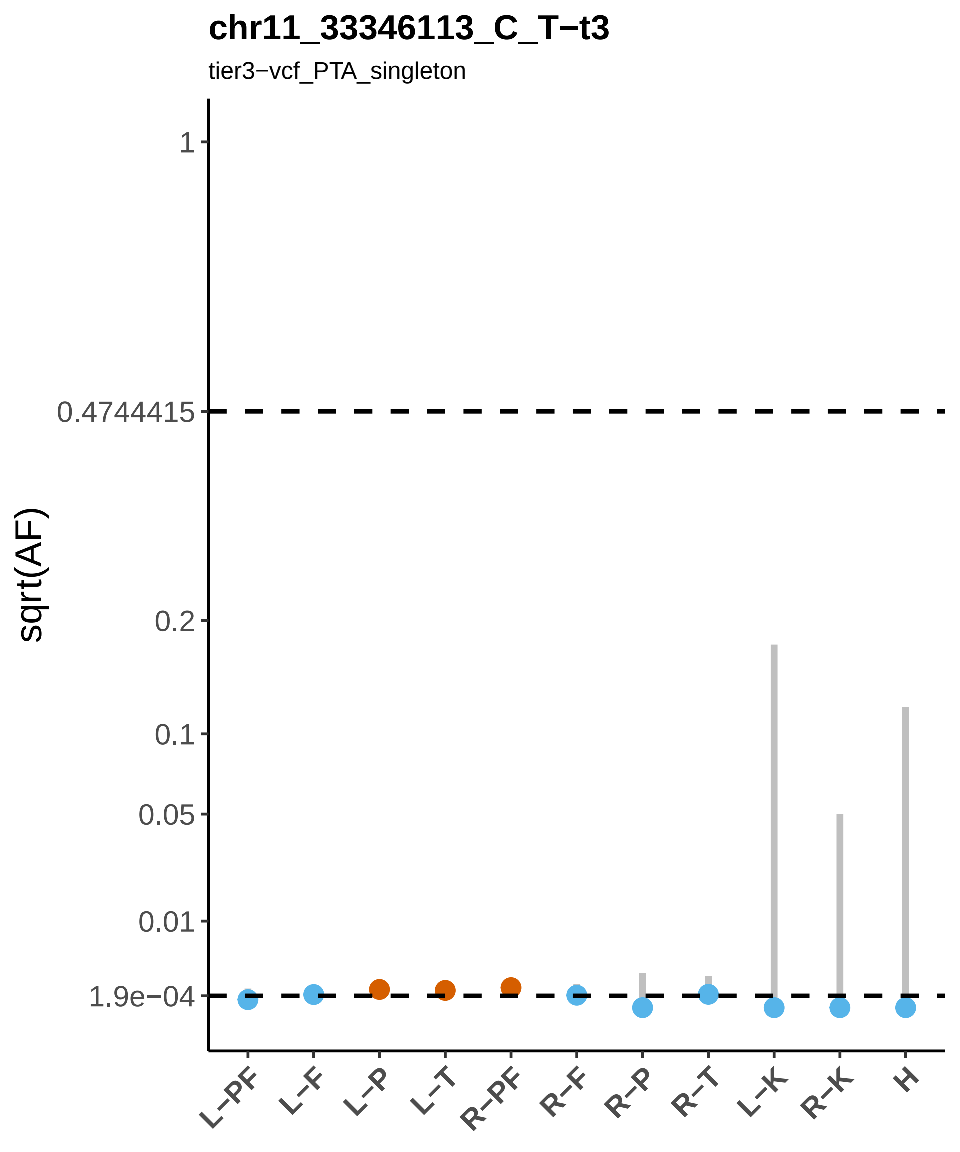

chr11_33346113_C_T−t3
tier3−vcf_PTA_singleton
sqrt(AF)
1
0.4744415
0.2
0.1
0.05
0.01
1.9e−04
L−PF
L−F
L−P
L−T
R−PF
R−F
R−P
R−T
L−K
R−K
H

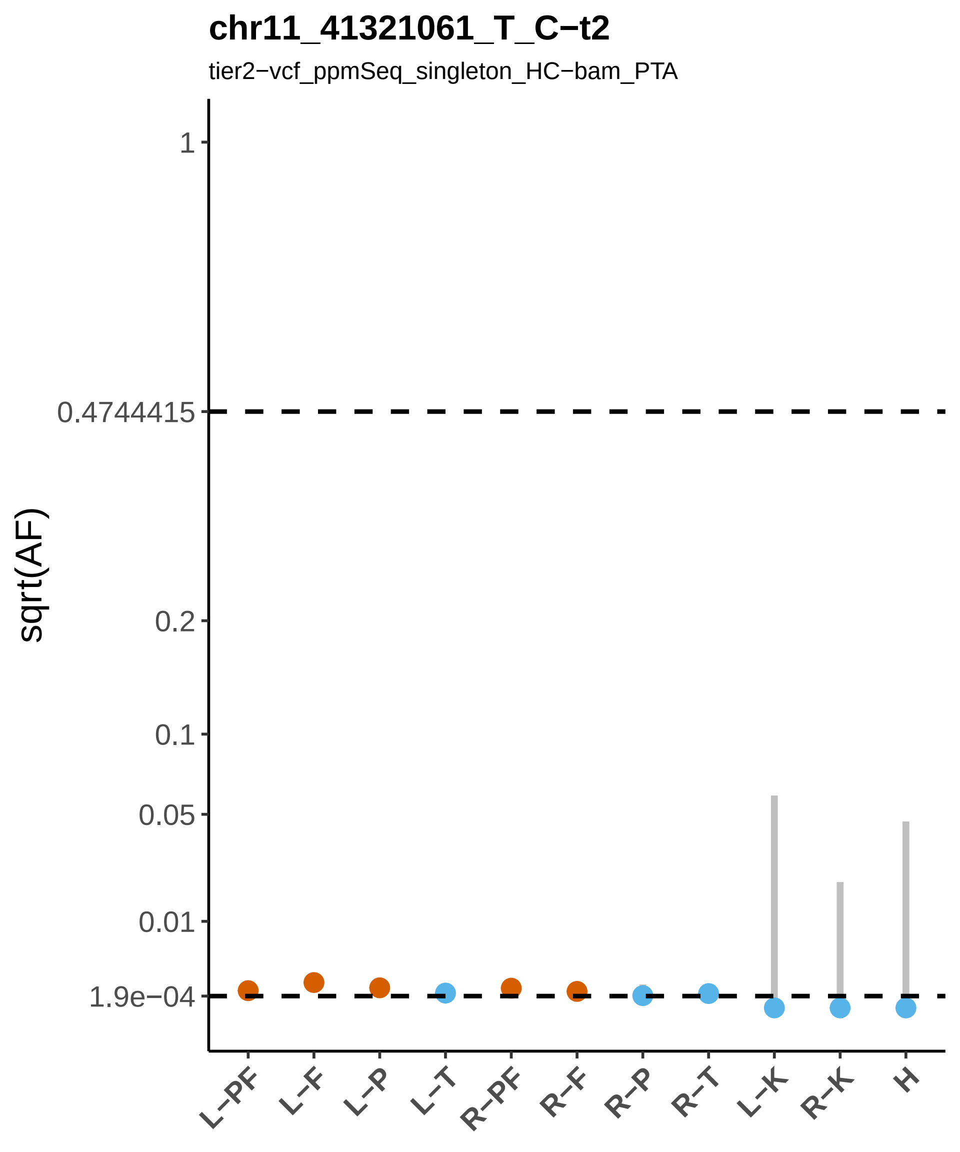

chr11_41321061_T_C−t2
tier2−vcf_ppmSeq_singleton_HC−bam_PTA
sqrt(AF)
1
0.4744415
0.2
0.1
0.05
0.01
1.9e−04
L−PF
L−F
L−P
L−T
R−PF
R−F
R−P
R−T
L−K
R−K
H

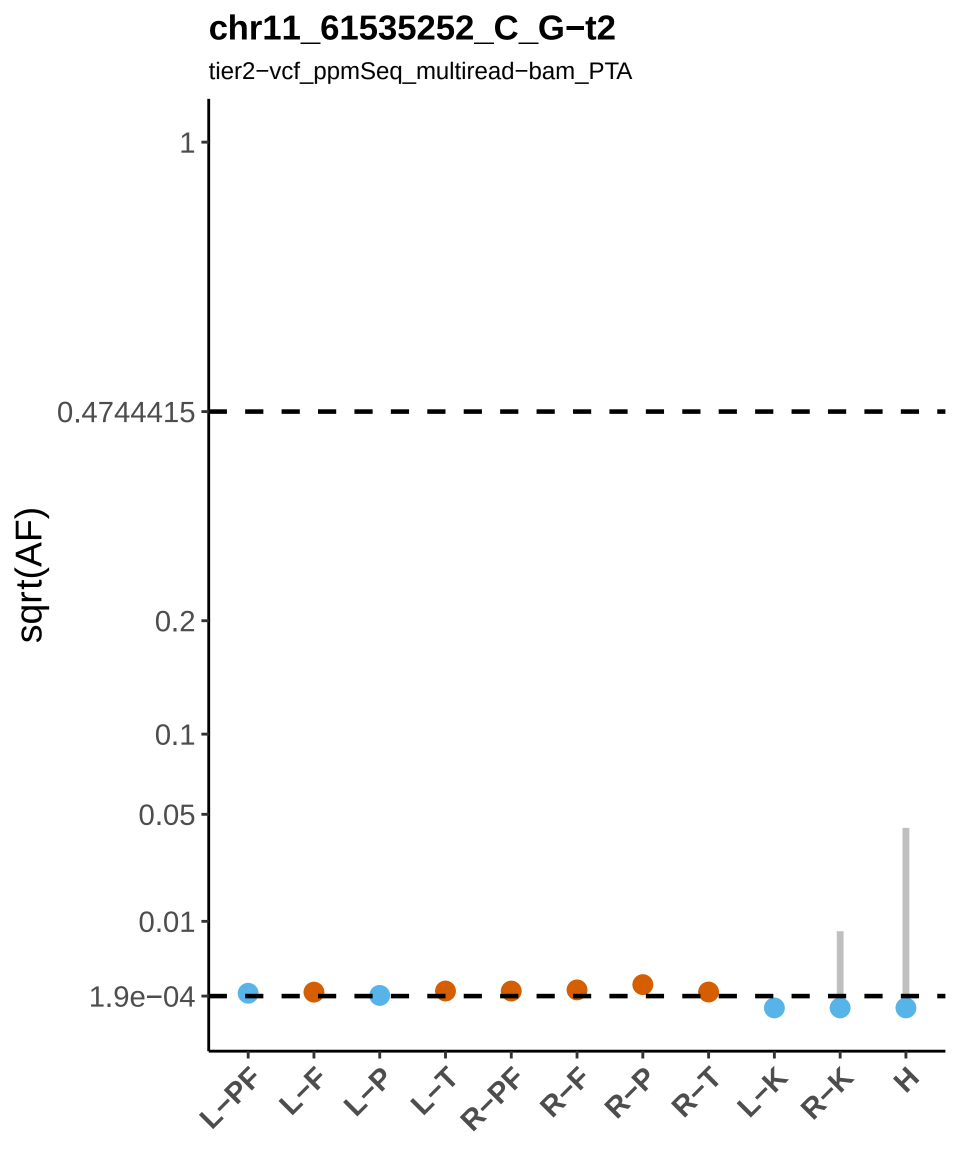

chr11_61535252_C_G−t2
tier2−vcf_ppmSeq_multiread−bam_PTA
sqrt(AF)
1
0.4744415
0.2
0.1
0.05
0.01
1.9e−04
L−PF
L−F
L−P
L−T
R−PF
R−F
R−P
R−T
L−K
R−K
H

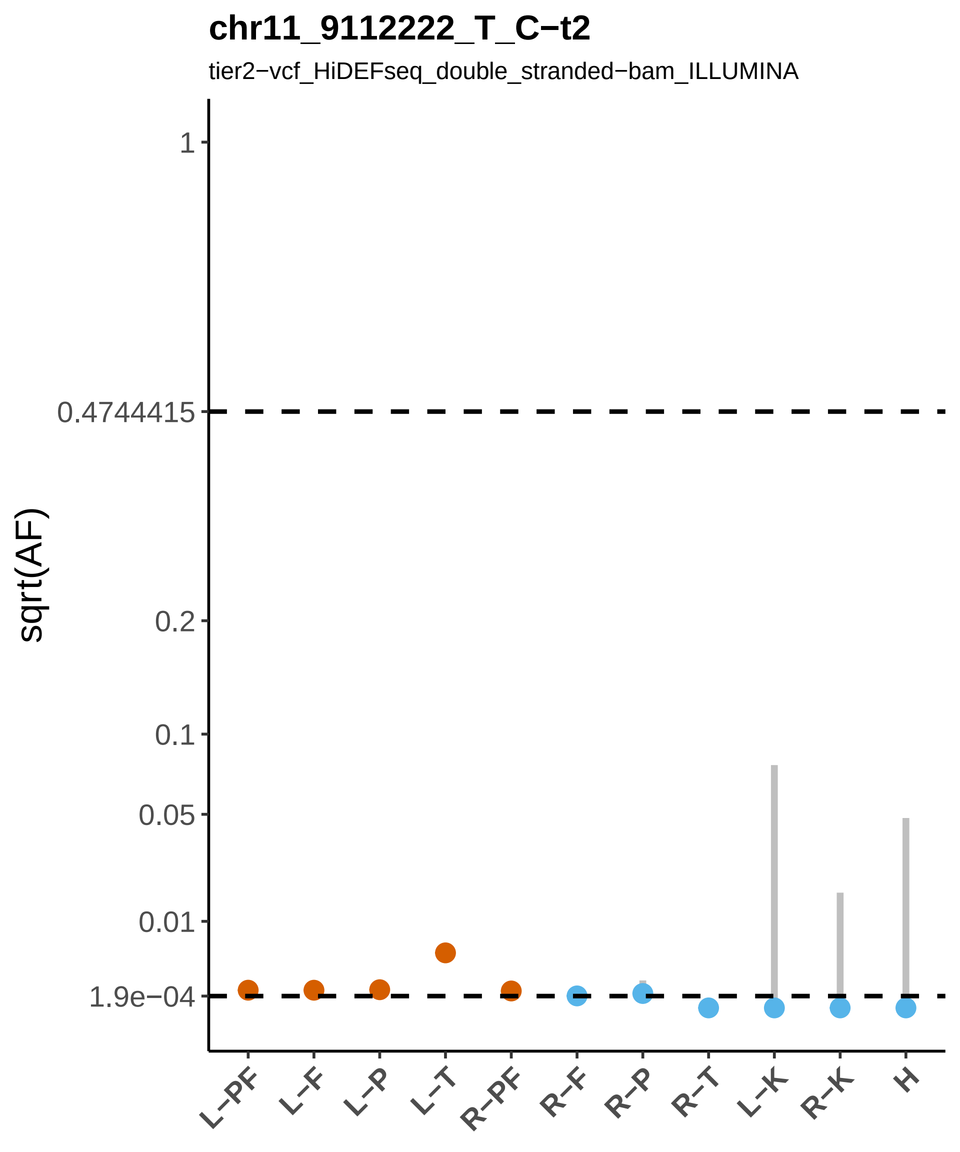

chr11_9112222_T_C−t2
tier2−vcf_HiDEFseq_double_stranded−bam_ILLUMINA
sqrt(AF)
1
0.4744415
0.2
0.1
0.05
0.01
1.9e−04
L−PF
L−F
L−P
L−T
R−PF
R−F
R−P
R−T
L−K
R−K
H

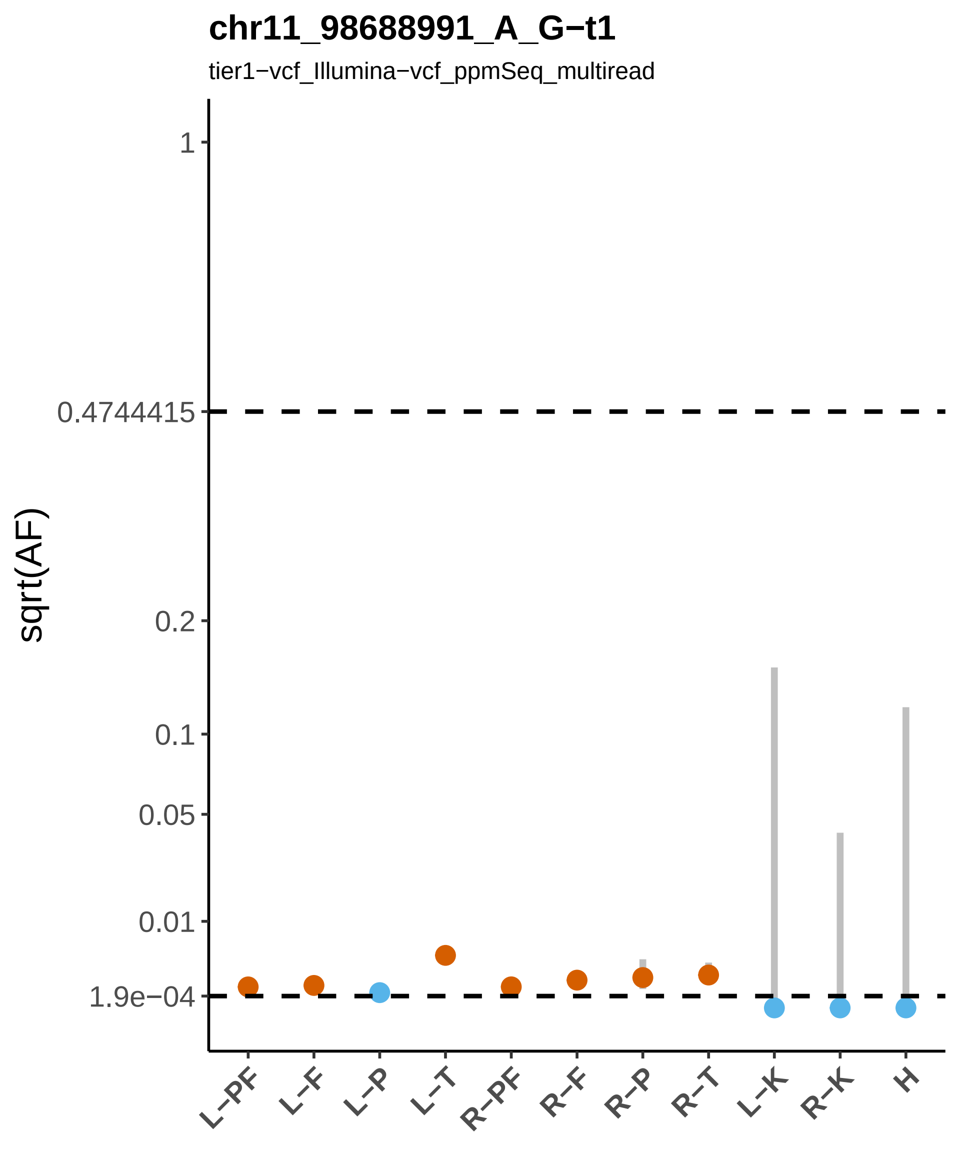
chr11_98688991_A_G−t1
tier1−vcf_Illumina−vcf_ppmSeq_multiread
sqrt(AF)
1
0.4744415
0.2
0.1
0.05
0.01
1.9e−04
L−PF
L−F
L−P
L−T
R−PF
R−F
R−P
R−T
L−K
R−K
H

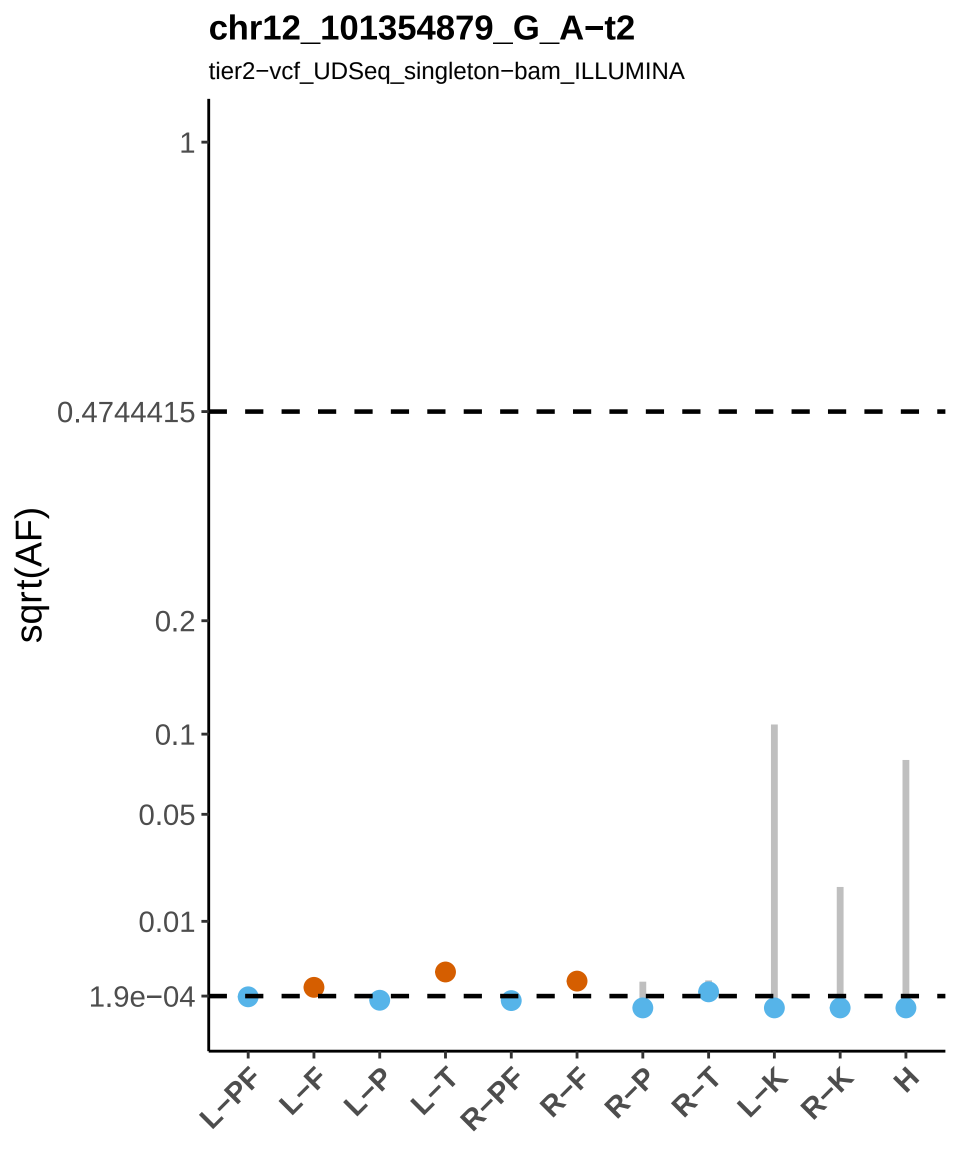
chr12_101354879_G_A−t2
tier2−vcf_UDSeq_singleton−bam_ILLUMINA
sqrt(AF)
1
0.4744415
0.2
0.1
0.05
0.01
1.9e−04
L−PF
L−F
L−P
L−T
R−PF
R−F
R−P
R−T
L−K
R−K
H

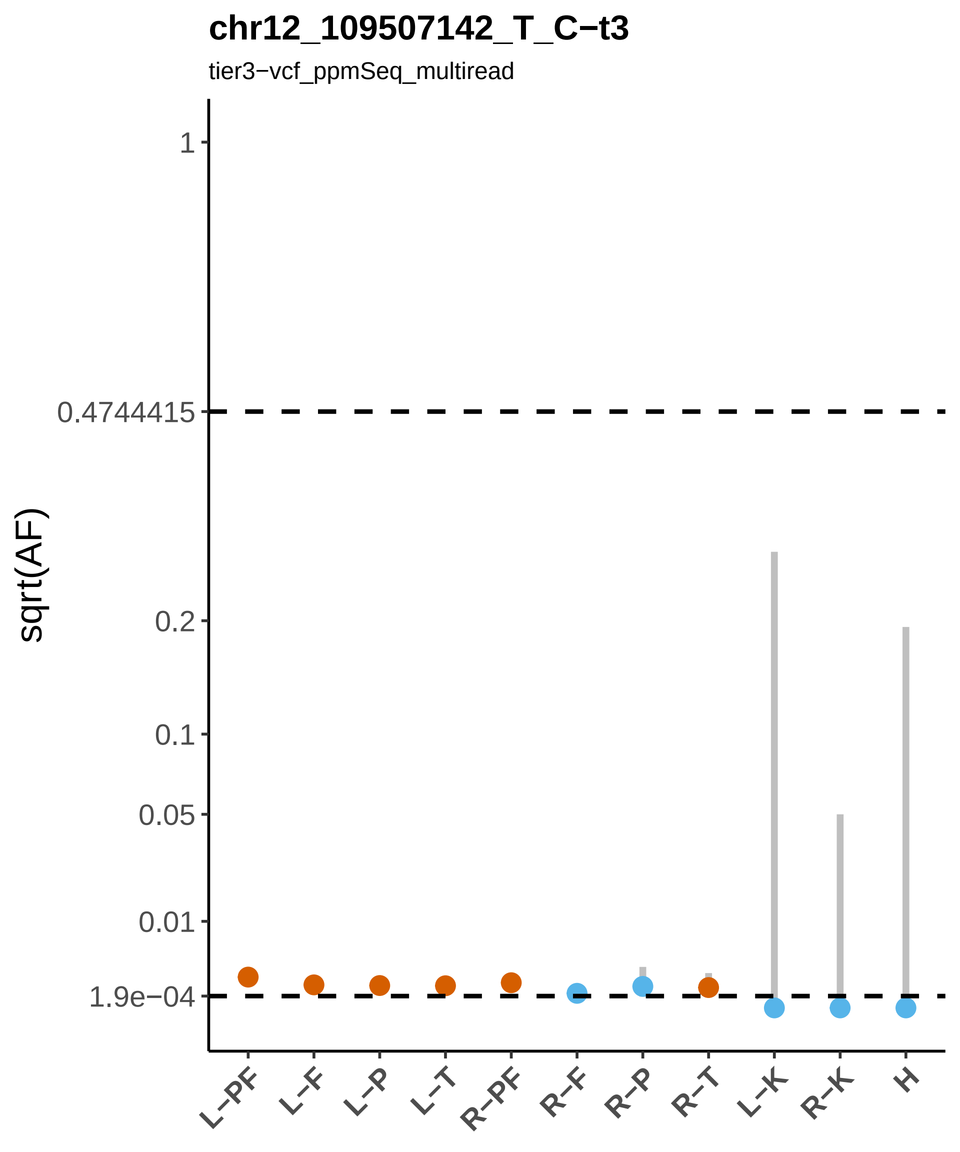
chr12_109507142_T_C−t3
tier3−vcf_ppmSeq_multiread
sqrt(AF)
1
0.4744415
0.2
0.1
0.05
0.01
1.9e−04
L−PF
L−F
L−P
L−T
R−PF
R−F
R−P
R−T
L−K
R−K
H

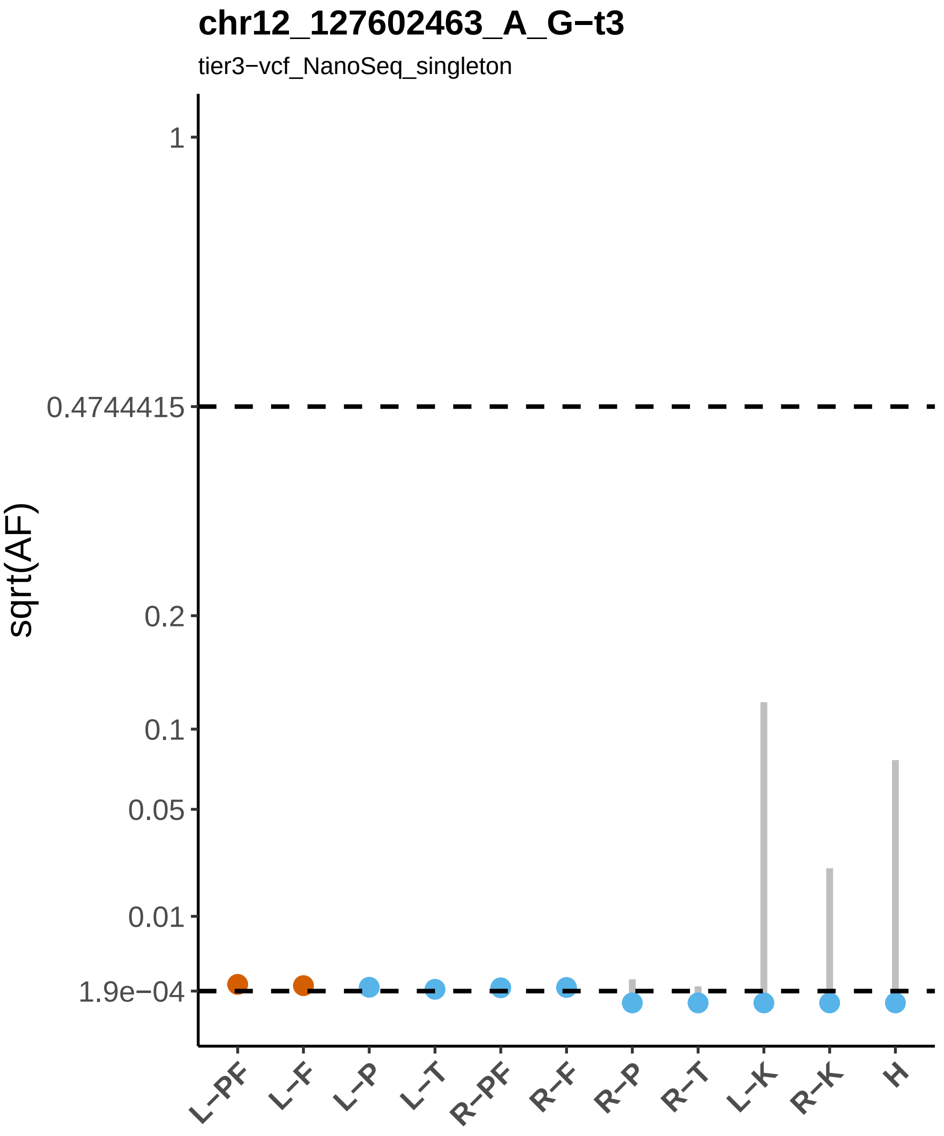
chr12_127602463_A_G−t3
tier3−vcf_NanoSeq_singleton
sqrt(AF)
1
0.4744415
0.2
0.1
0.05
0.01
1.9e−04
L−PF
L−F
L−P
L−T
R−PF
R−F
R−P
R−T
L−K
R−K
H

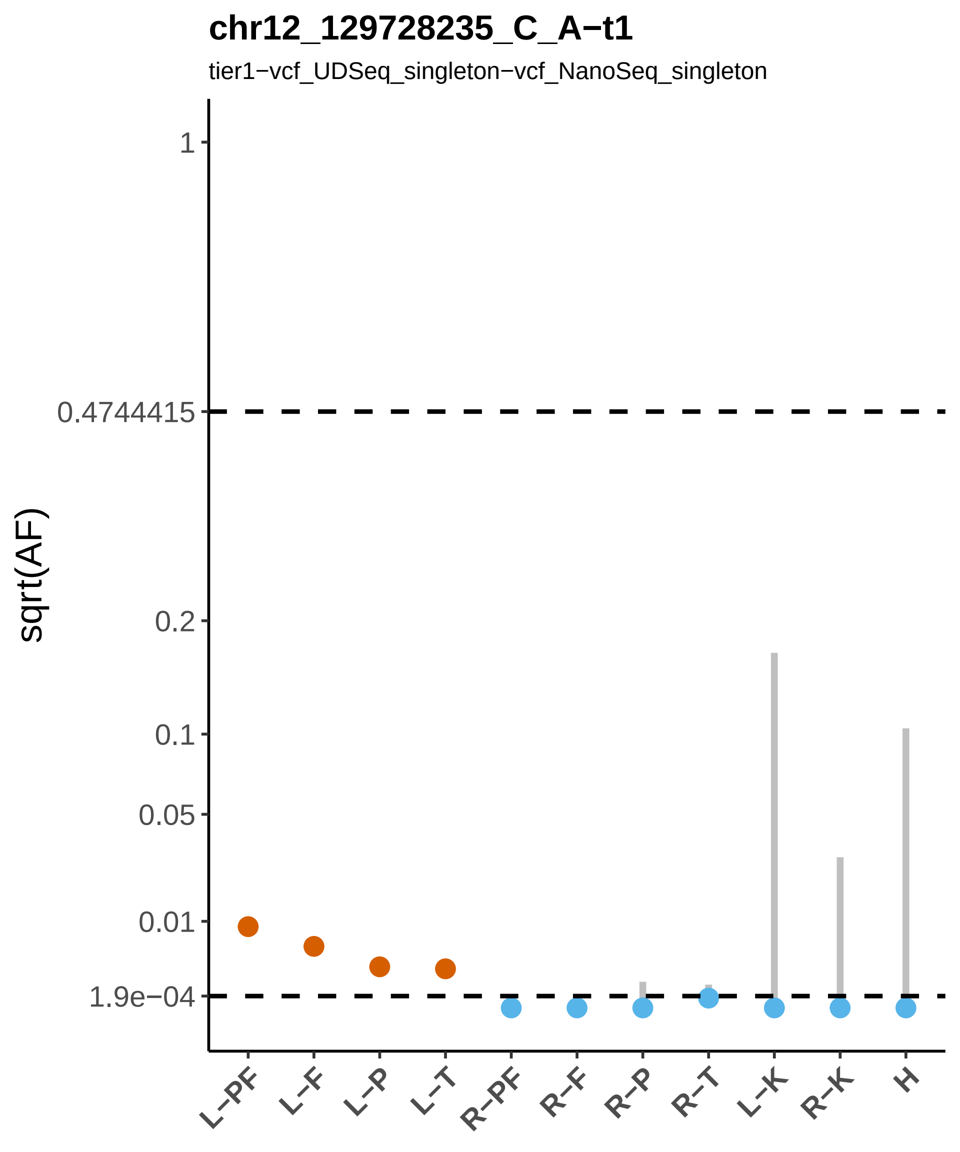
chr12_129728235_C_A−t1
tier1−vcf_UDSeq_singleton−vcf_NanoSeq_singleton
sqrt(AF)
1
0.4744415
0.2
0.1
0.05
0.01
1.9e−04
L−PF
L−F
L−P
L−T
R−PF
R−F
R−P
R−T
L−K
R−K
H

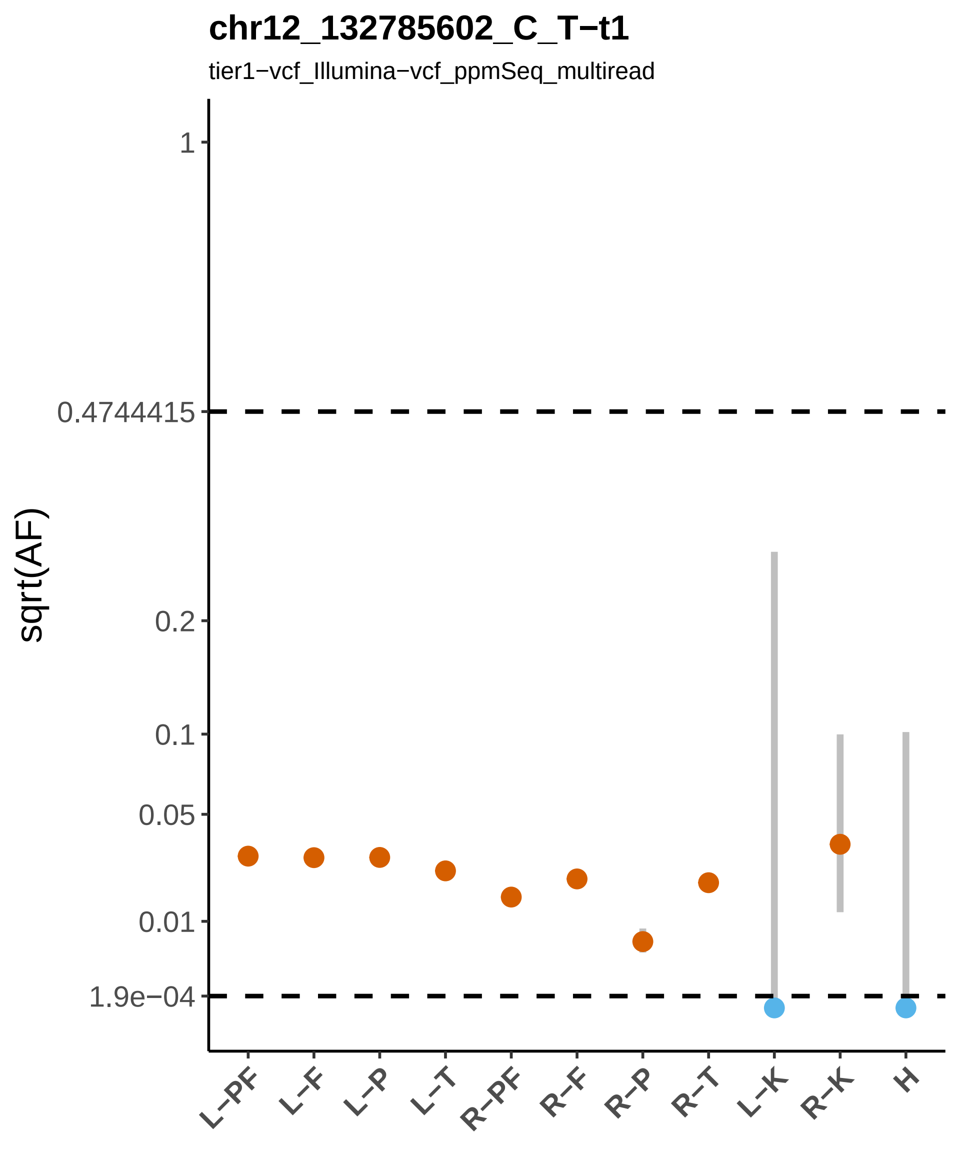
chr12_132785602_C_T−t1
tier1−vcf_Illumina−vcf_ppmSeq_multiread
sqrt(AF)
1
0.4744415
0.2
0.1
0.05
0.01
1.9e−04
L−PF
L−F
L−P
L−T
R−PF
R−F
R−P
R−T
L−K
R−K
H

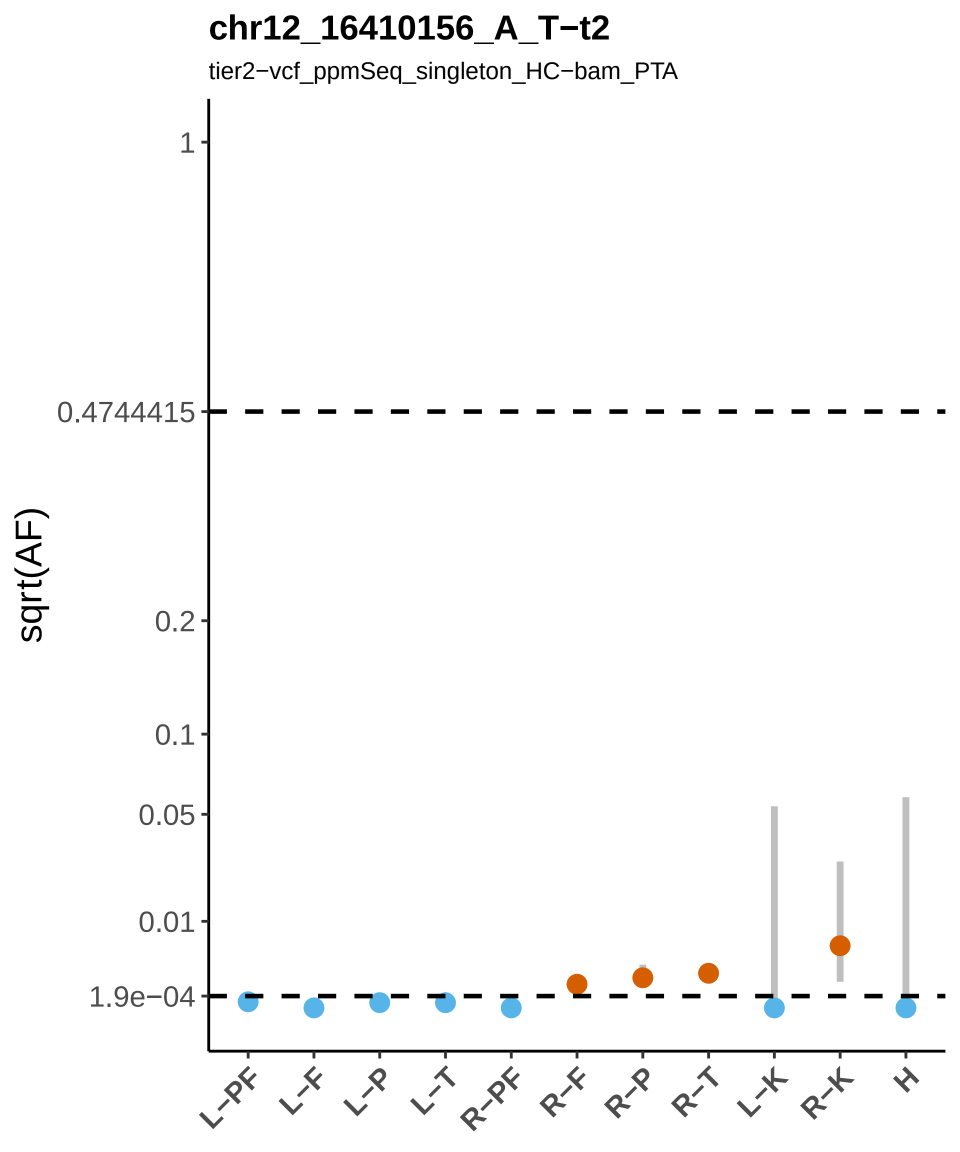
chr12_16410156_A_T−t2
tier2−vcf_ppmSeq_singleton_HC−bam_PTA
sqrt(AF)
1
0.4744415
0.2
0.1
0.05
0.01
1.9e−04
L−PF
L−F
L−P
L−T
R−PF
R−F
R−P
R−T
L−K
R−K
H

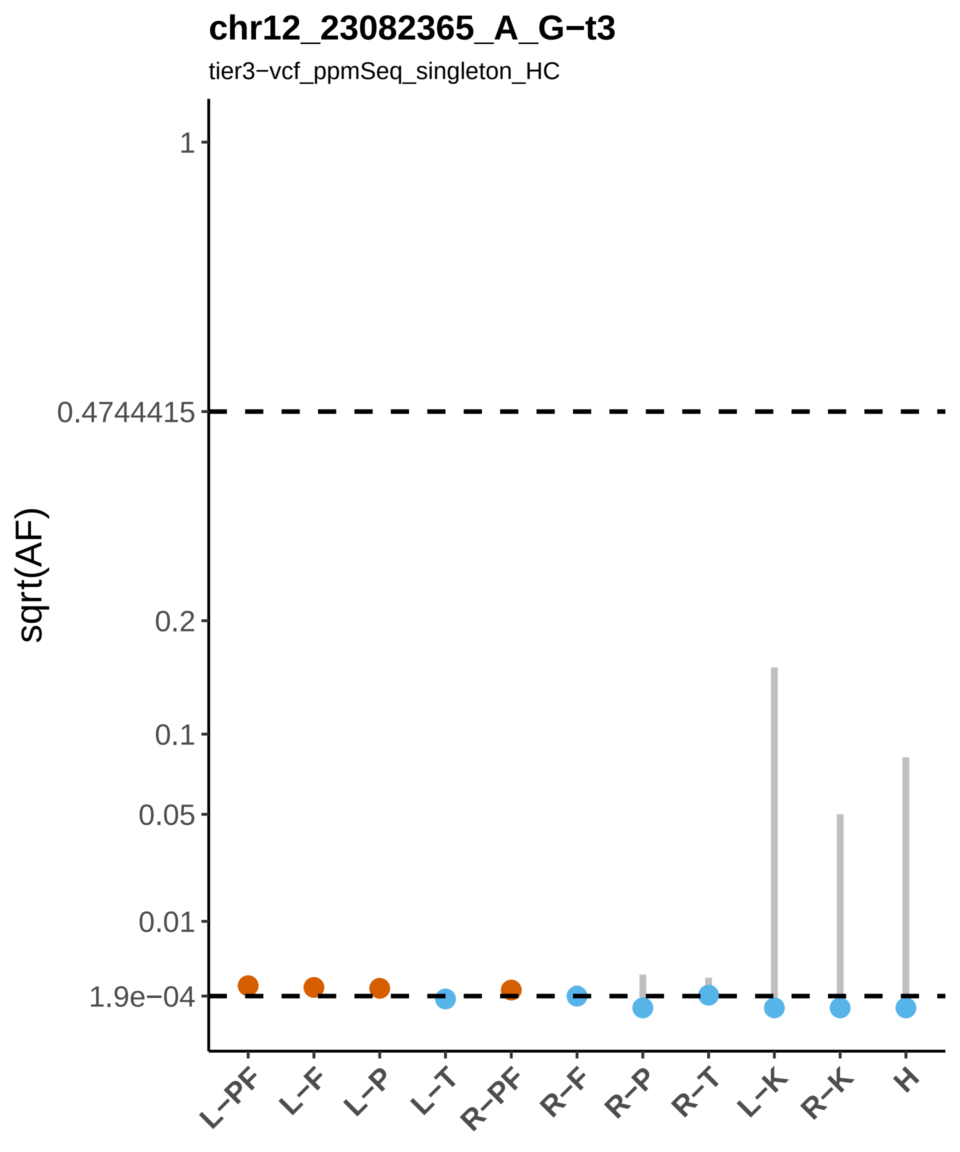

chr12_23082365_A_G−t3
tier3−vcf_ppmSeq_singleton_HC
sqrt(AF)
1
0.4744415
0.2
0.1
0.05
0.01
1.9e−04
L−PF
L−F
L−P
L−T
R−PF
R−F
R−P
R−T
L−K
R−K
H

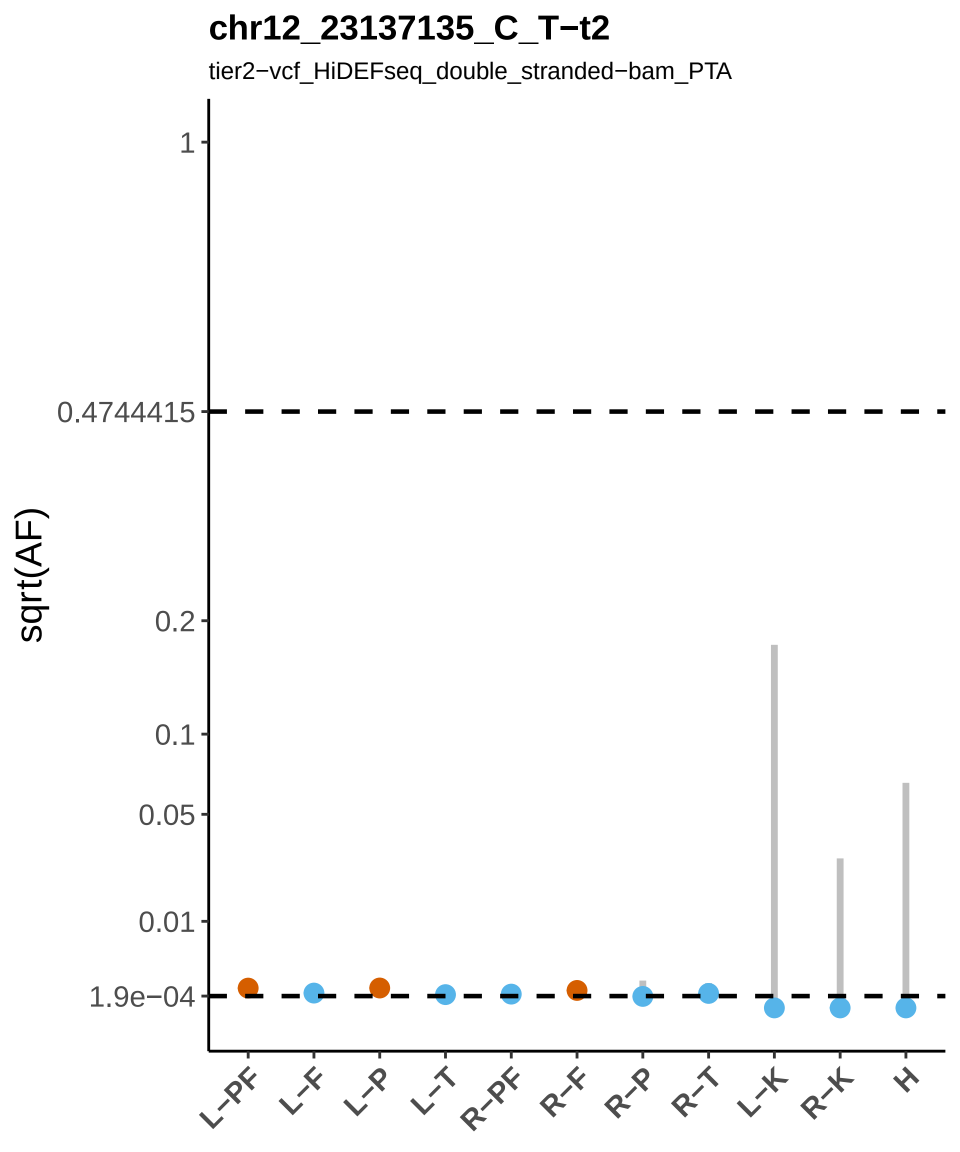

chr12_23137135_C_T−t2
tier2−vcf_HiDEFseq_double_stranded−bam_PTA
sqrt(AF)
1
0.4744415
0.2
0.1
0.05
0.01
1.9e−04
L−PF
L−F
L−P
L−T
R−PF
R−F
R−P
R−T
L−K
R−K
H

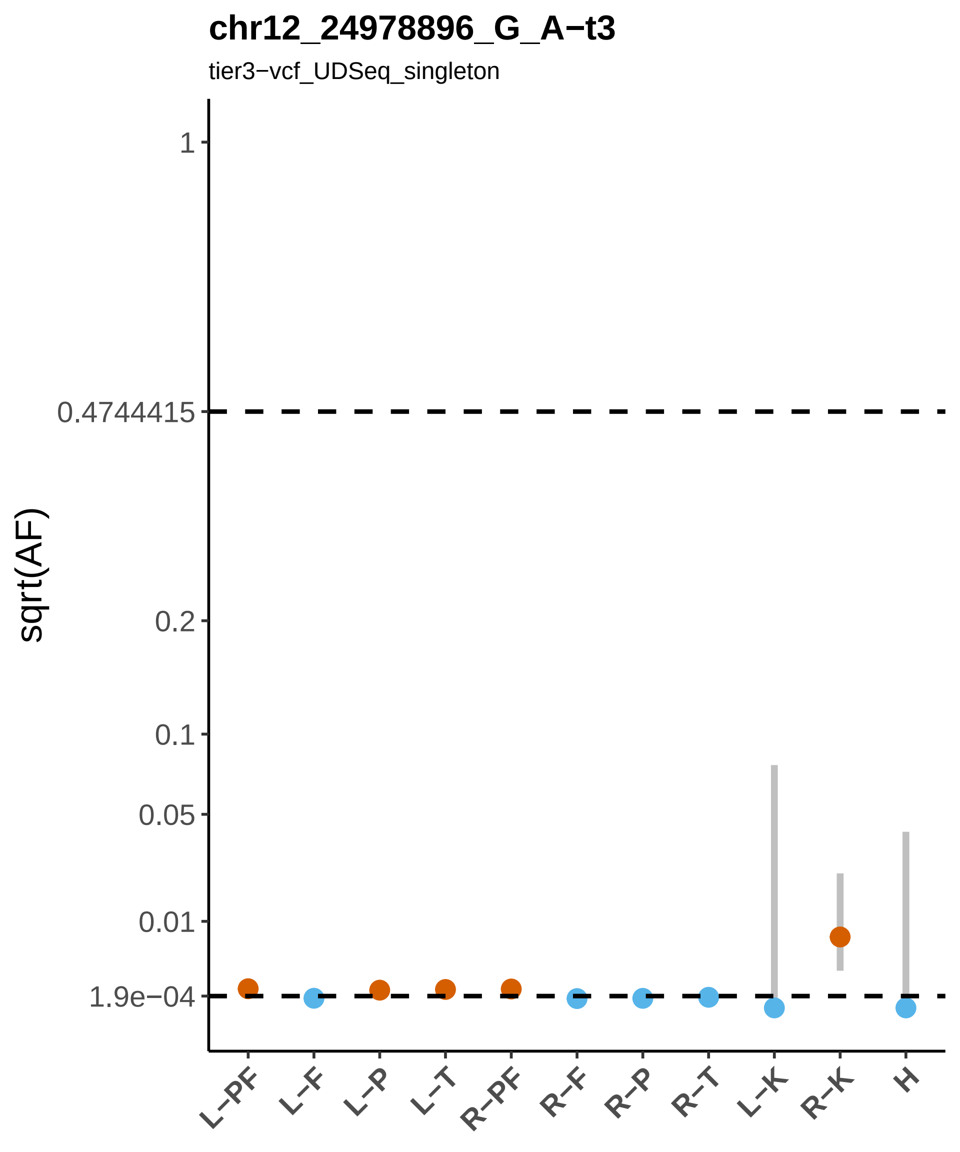
chr12_24978896_G_A−t3
tier3−vcf_UDSeq_singleton
sqrt(AF)
1
0.4744415
0.2
0.1
0.05
0.01
1.9e−04
L−PF
L−F
L−P
L−T
R−PF
R−F
R−P
R−T
L−K
R−K
H

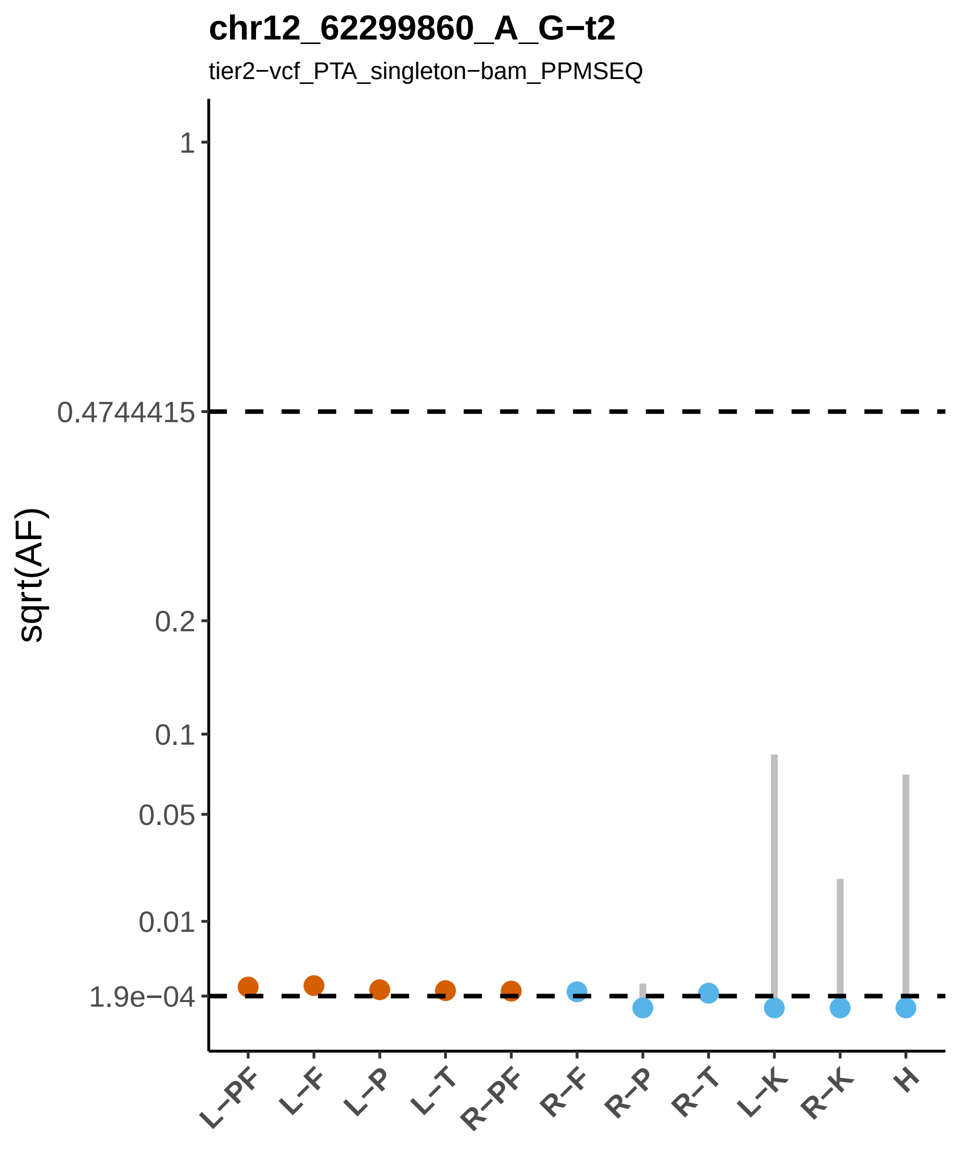
chr12_62299860_A_G−t2
tier2−vcf_PTA_singleton−bam_PPMSEQ
sqrt(AF)
1
0.4744415
0.2
0.1
0.05
0.01
1.9e−04
L−PF
L−F
L−P
L−T
R−PF
R−F
R−P
R−T
L−K
R−K
H

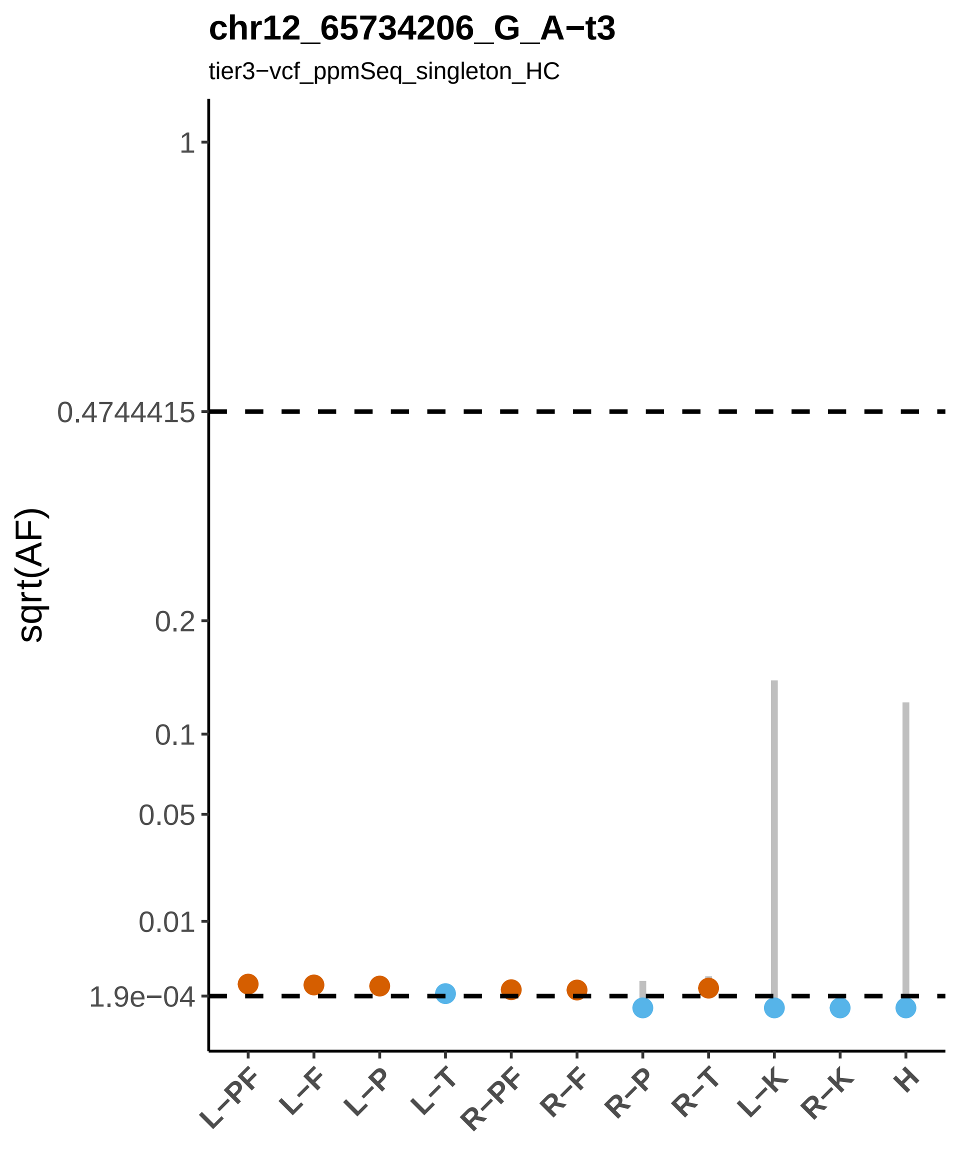
chr12_65734206_G_A−t3
tier3−vcf_ppmSeq_singleton_HC
sqrt(AF)
1
0.4744415
0.2
0.1
0.05
0.01
1.9e−04
L−PF
L−F
L−P
L−T
R−PF
R−F
R−P
R−T
L−K
R−K
H

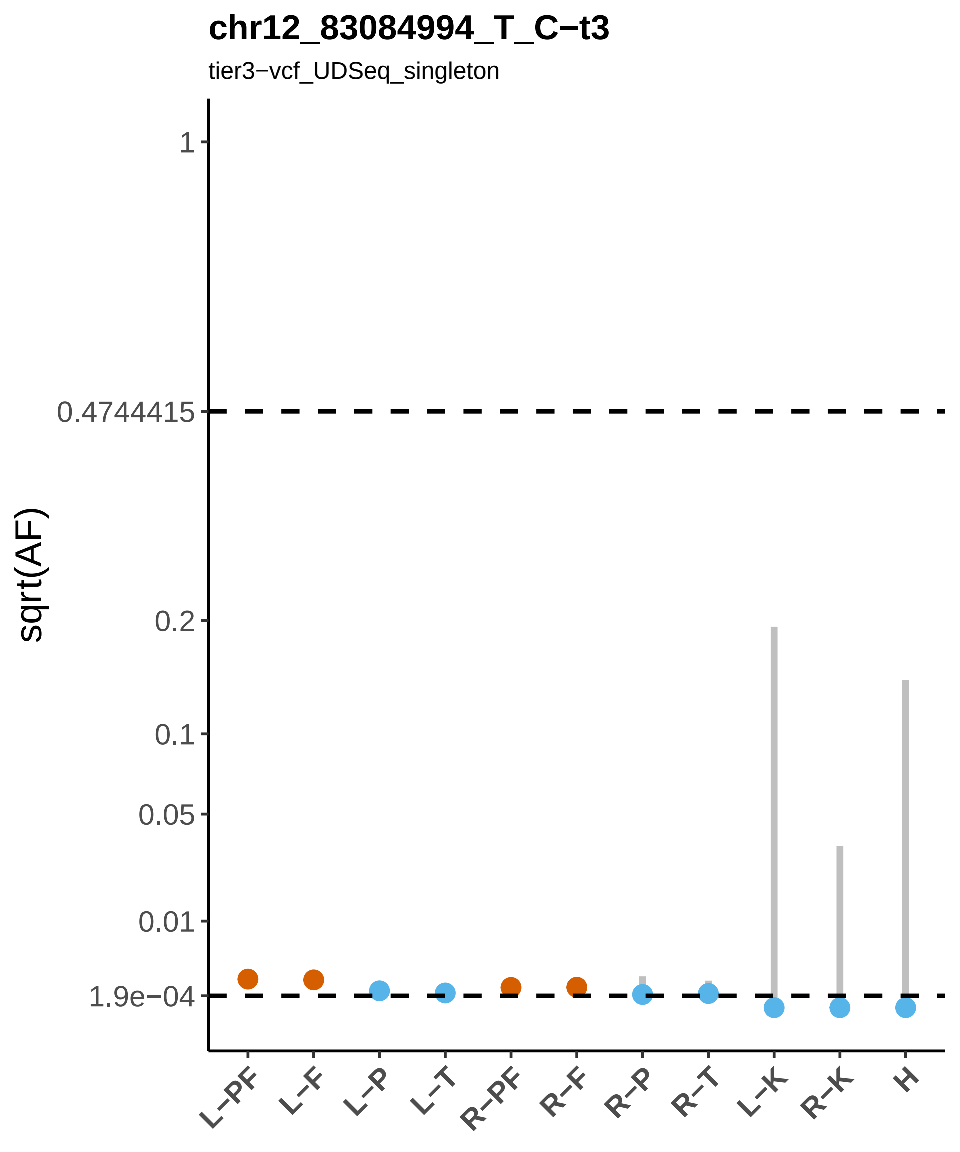
chr12_83084994_T_C−t3
tier3−vcf_UDSeq_singleton
sqrt(AF)
1
0.4744415
0.2
0.1
0.05
0.01
1.9e−04
L−PF
L−F
L−P
L−T
R−PF
R−F
R−P
R−T
L−K
R−K
H

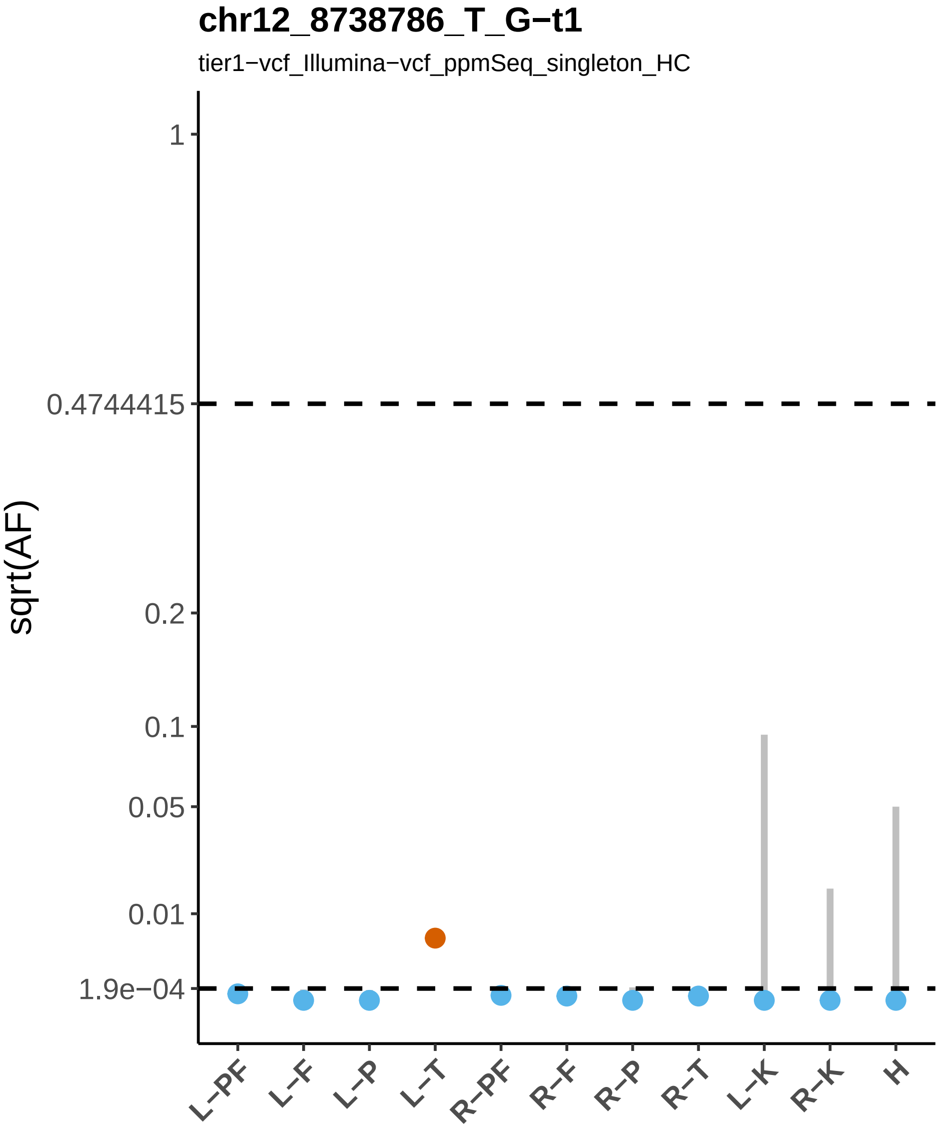

chr12_8738786_T_G−t1
tier1−vcf_Illumina−vcf_ppmSeq_singleton_HC
sqrt(AF)
1
0.4744415
0.2
0.1
0.05
0.01
1.9e−04
L−PF
L−F
L−P
L−T
R−PF
R−F
R−P
R−T
L−K
R−K
H

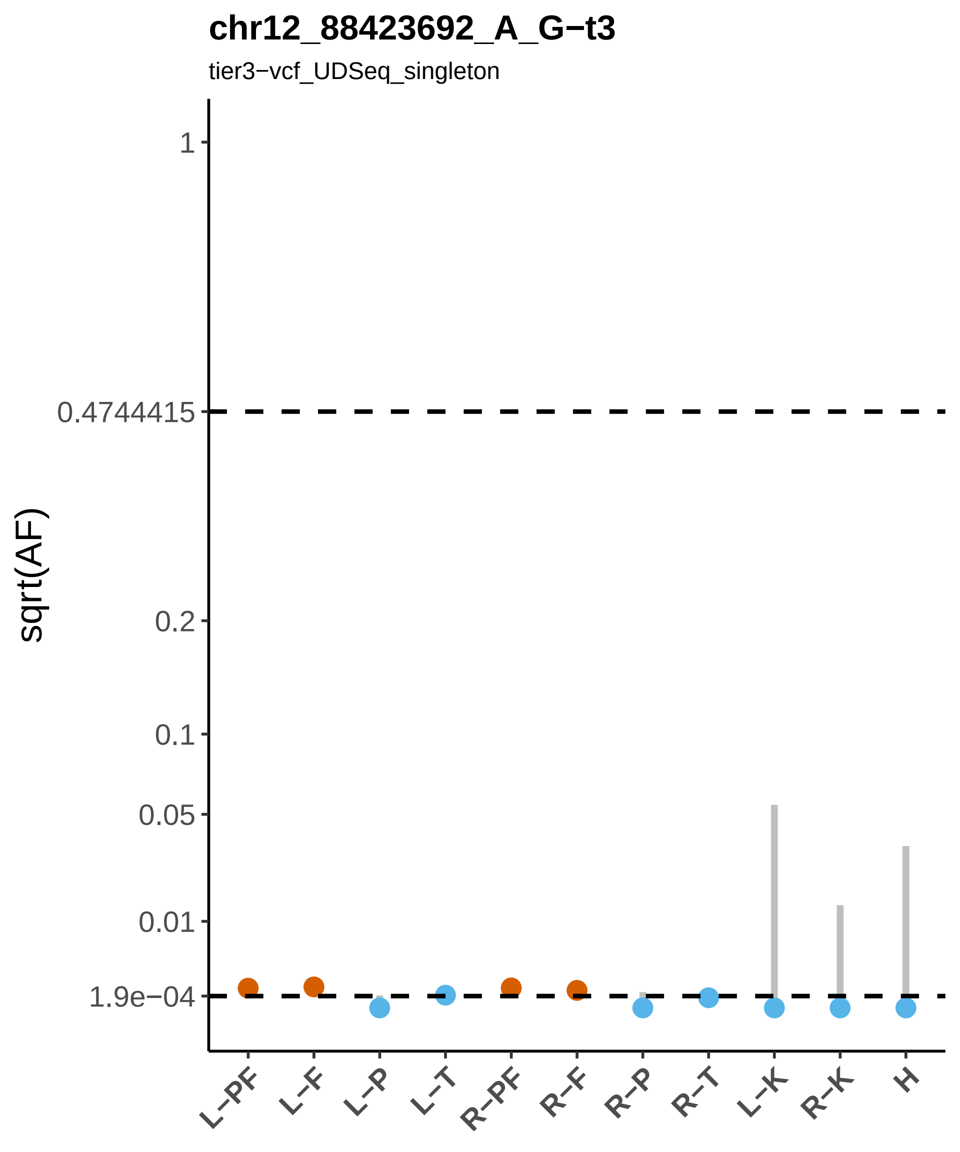

chr12_88423692_A_G−t3
tier3−vcf_UDSeq_singleton
sqrt(AF)
1
0.4744415
0.2
0.1
0.05
0.01
1.9e−04
L−PF
L−F
L−P
L−T
R−PF
R−F
R−P
R−T
L−K
R−K
H

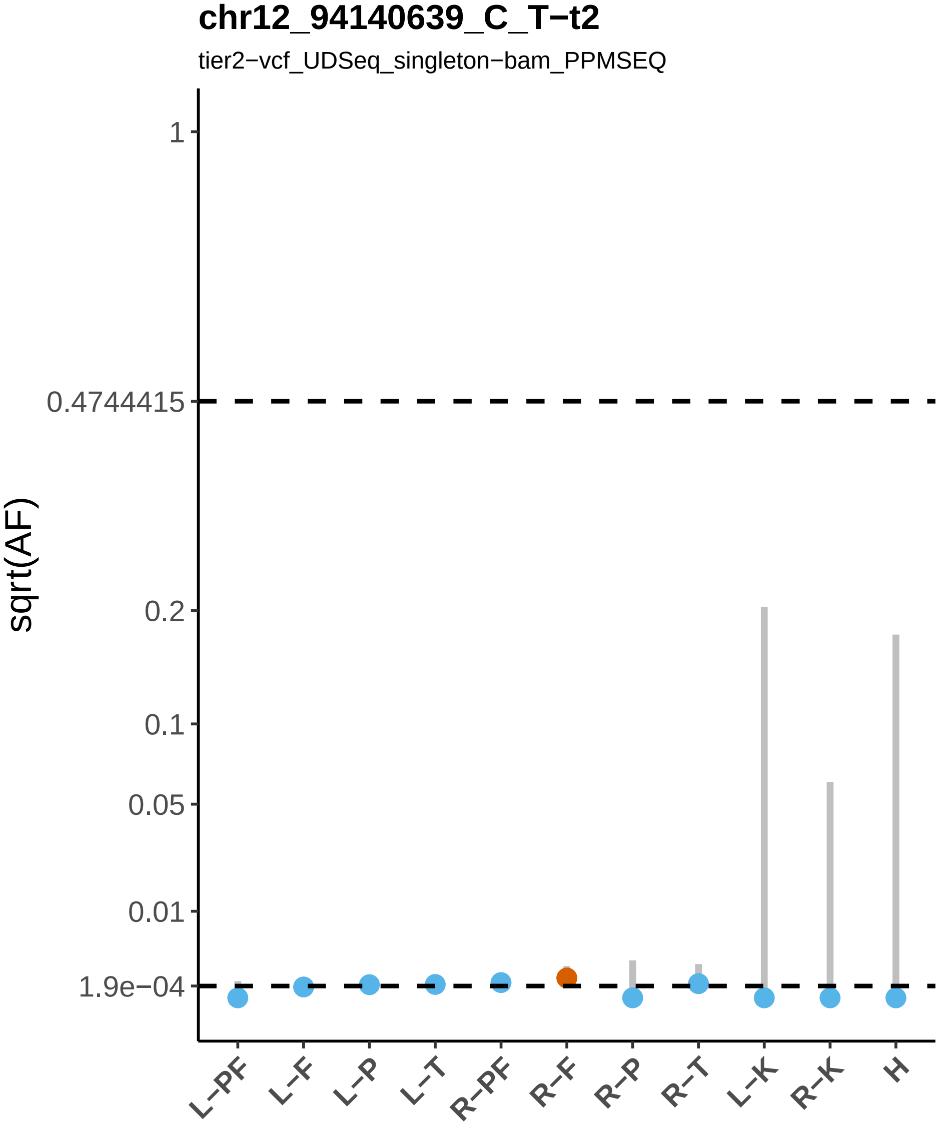
chr12_94140639_C_T−t2
tier2−vcf_UDSeq_singleton−bam_PPMSEQ
sqrt(AF)
1
0.4744415
0.2
0.1
0.05
0.01
1.9e−04
L−PF
L−F
L−P
L−T
R−PF
R−F
R−P
R−T
L−K
R−K
H

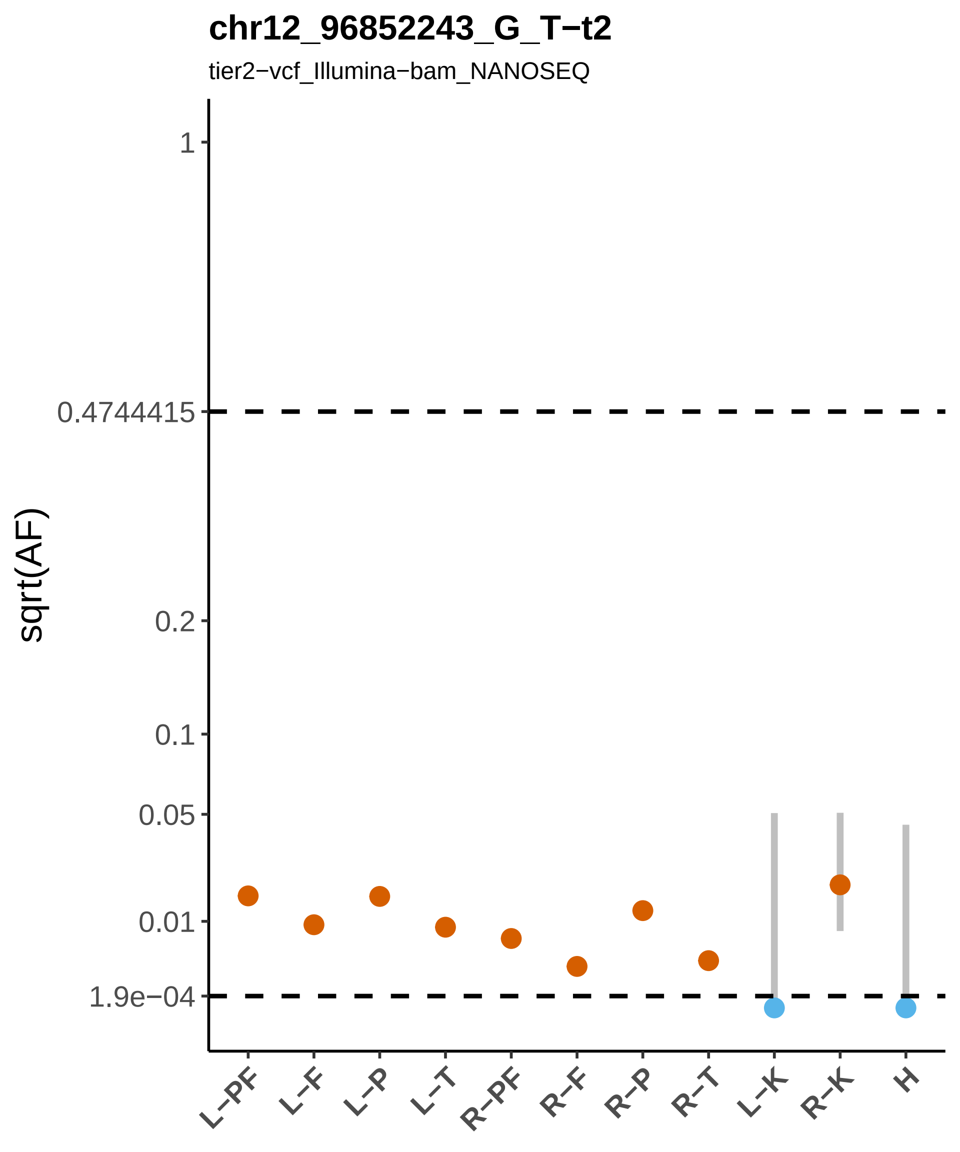

chr12_96852243_G_T−t2
tier2−vcf_Illumina−bam_NANOSEQ
sqrt(AF)
1
0.4744415
0.2
0.1
0.05
0.01
1.9e−04
L−PF
L−F
L−P
L−T
R−PF
R−F
R−P
R−T
L−K
R−K
H

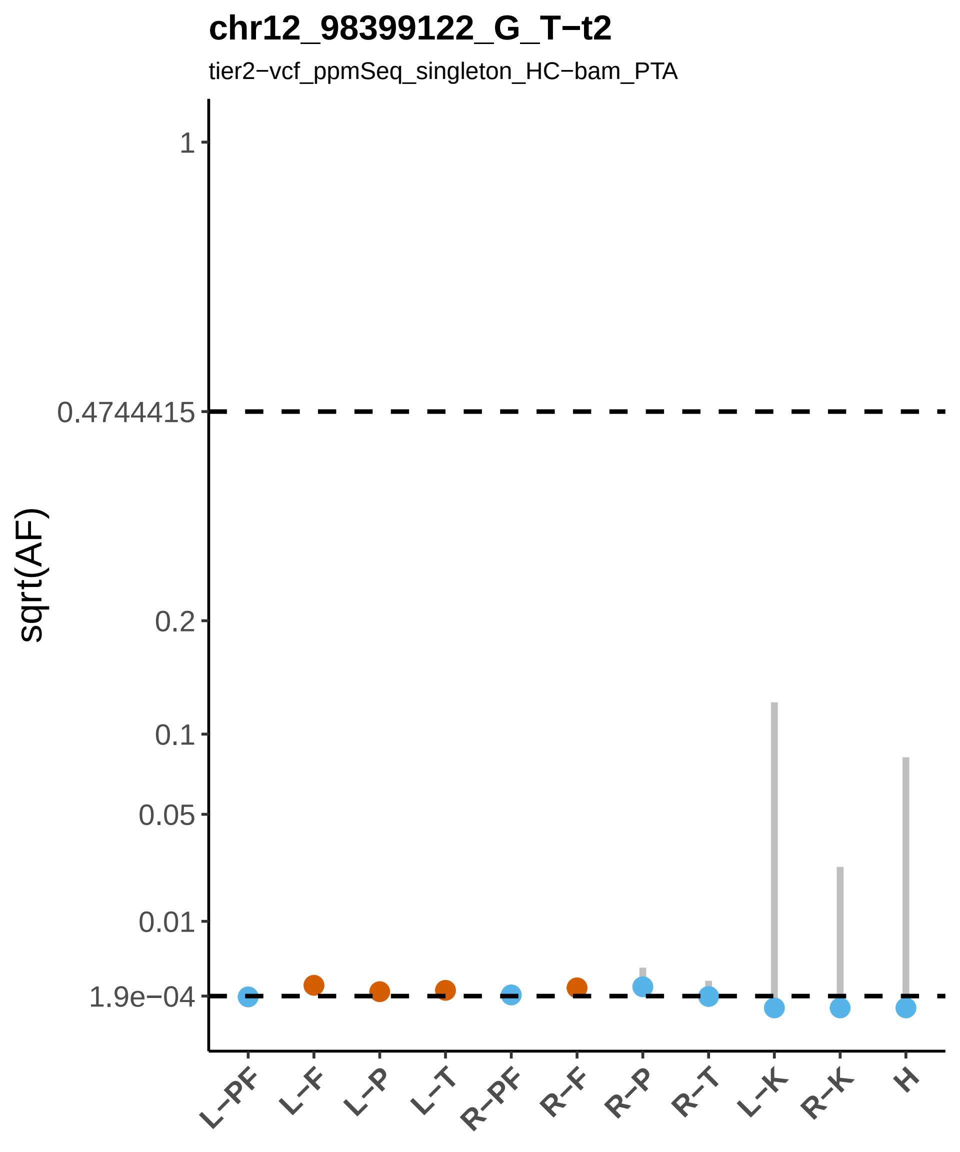

chr12_98399122_G_T−t2
tier2−vcf_ppmSeq_singleton_HC−bam_PTA
sqrt(AF)
1
0.4744415
0.2
0.1
0.05
0.01
1.9e−04
L−PF
L−F
L−P
L−T
R−PF
R−F
R−P
R−T
L−K
R−K
H

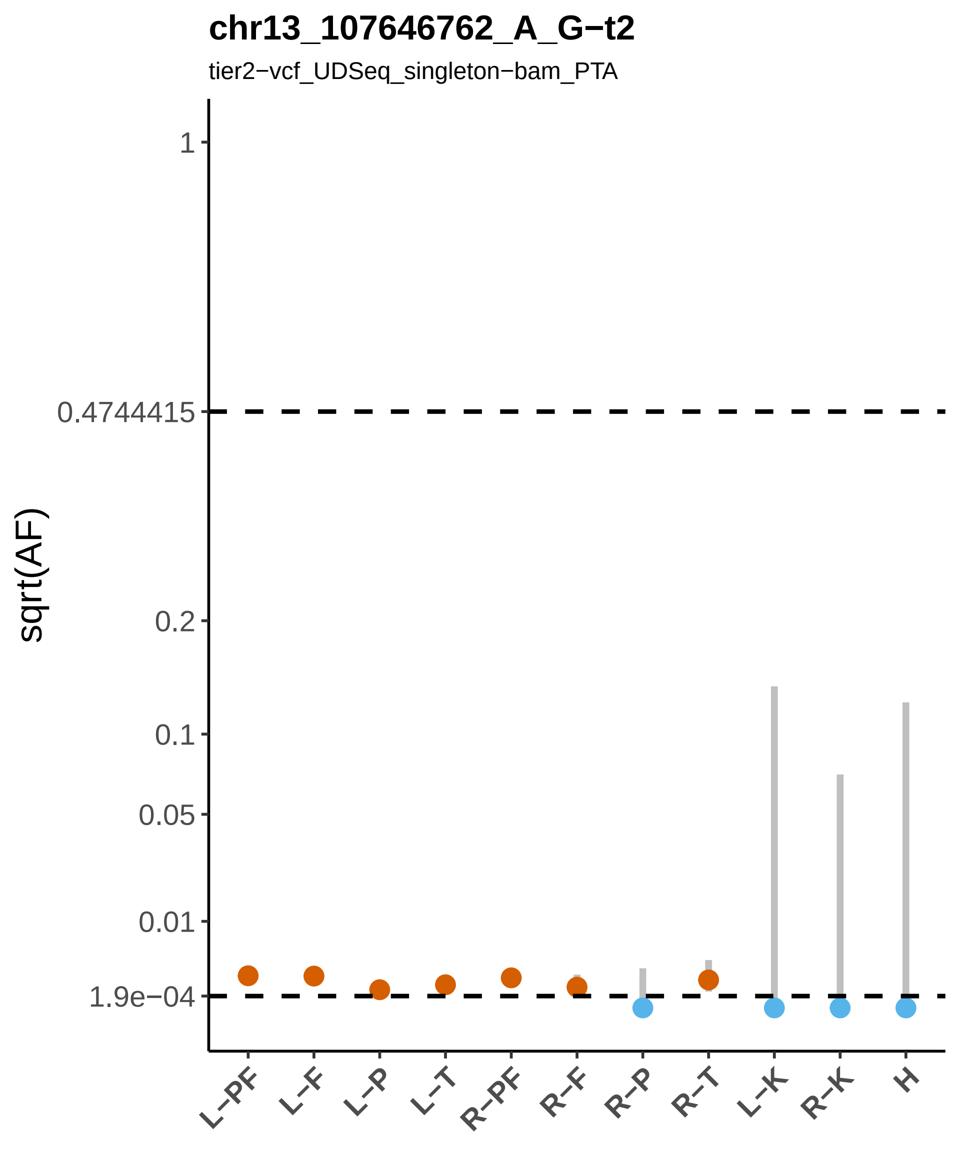

chr13_107646762_A_G−t2
tier2−vcf_UDSeq_singleton−bam_PTA
sqrt(AF)
1
0.4744415
0.2
0.1
0.05
0.01
1.9e−04
L−PF
L−F
L−P
L−T
R−PF
R−F
R−P
R−T
L−K
R−K
H

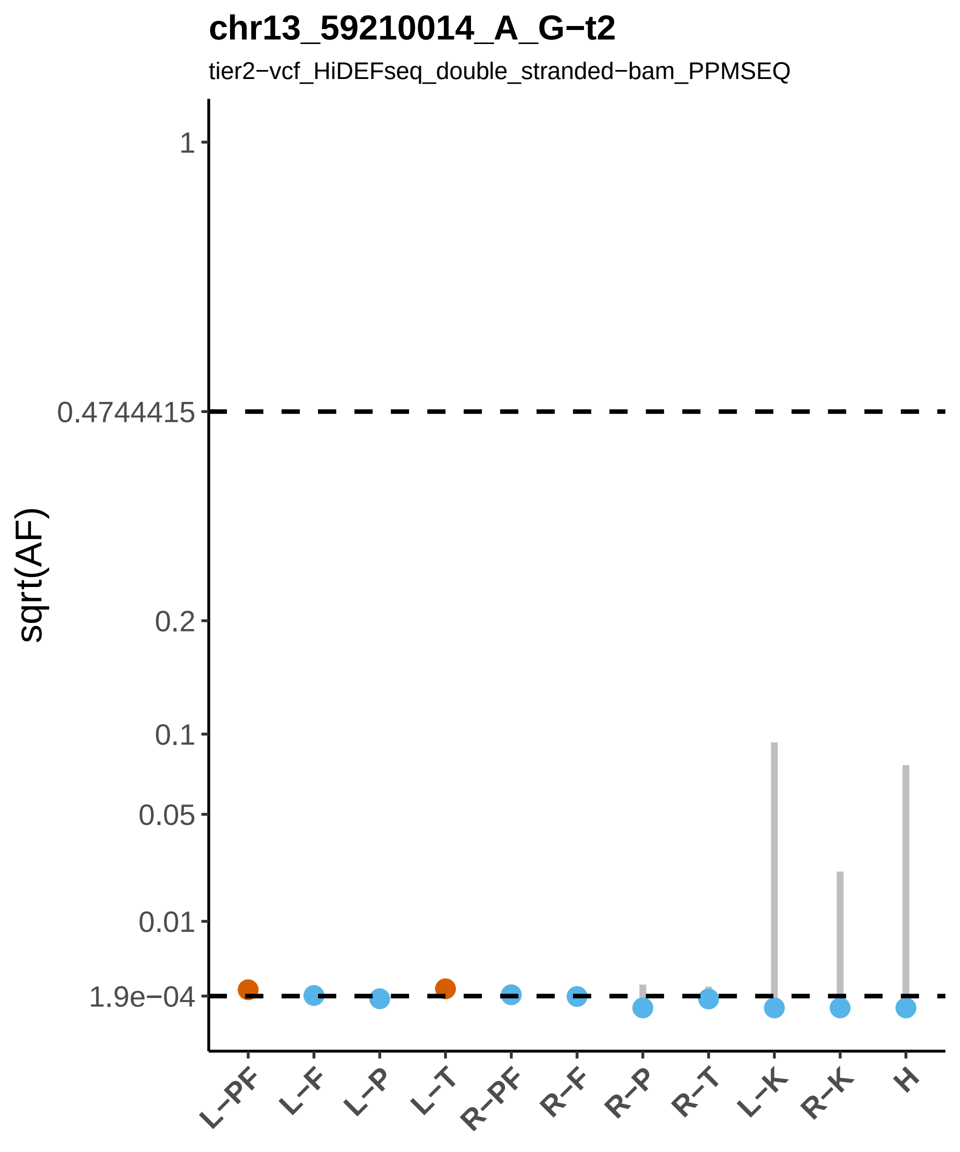
chr13_59210014_A_G−t2
tier2−vcf_HiDEFseq_double_stranded−bam_PPMSEQ
sqrt(AF)
1
0.4744415
0.2
0.1
0.05
0.01
1.9e−04
L−PF
L−F
L−P
L−T
R−PF
R−F
R−P
R−T
L−K
R−K
H

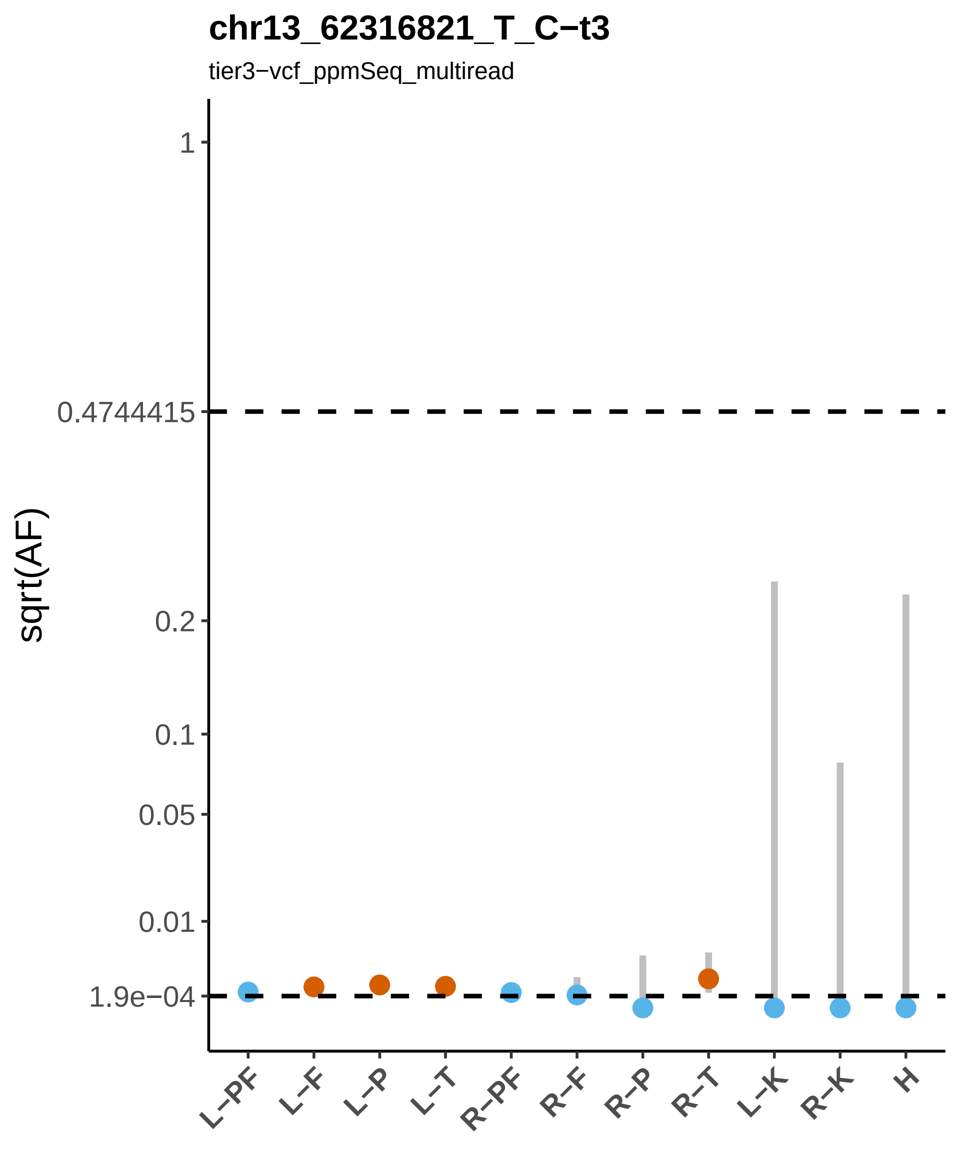
chr13_62316821_T_C−t3
tier3−vcf_ppmSeq_multiread
sqrt(AF)
1
0.4744415
0.2
0.1
0.05
0.01
1.9e−04
L−PF
L−F
L−P
L−T
R−PF
R−F
R−P
R−T
L−K
R−K
H

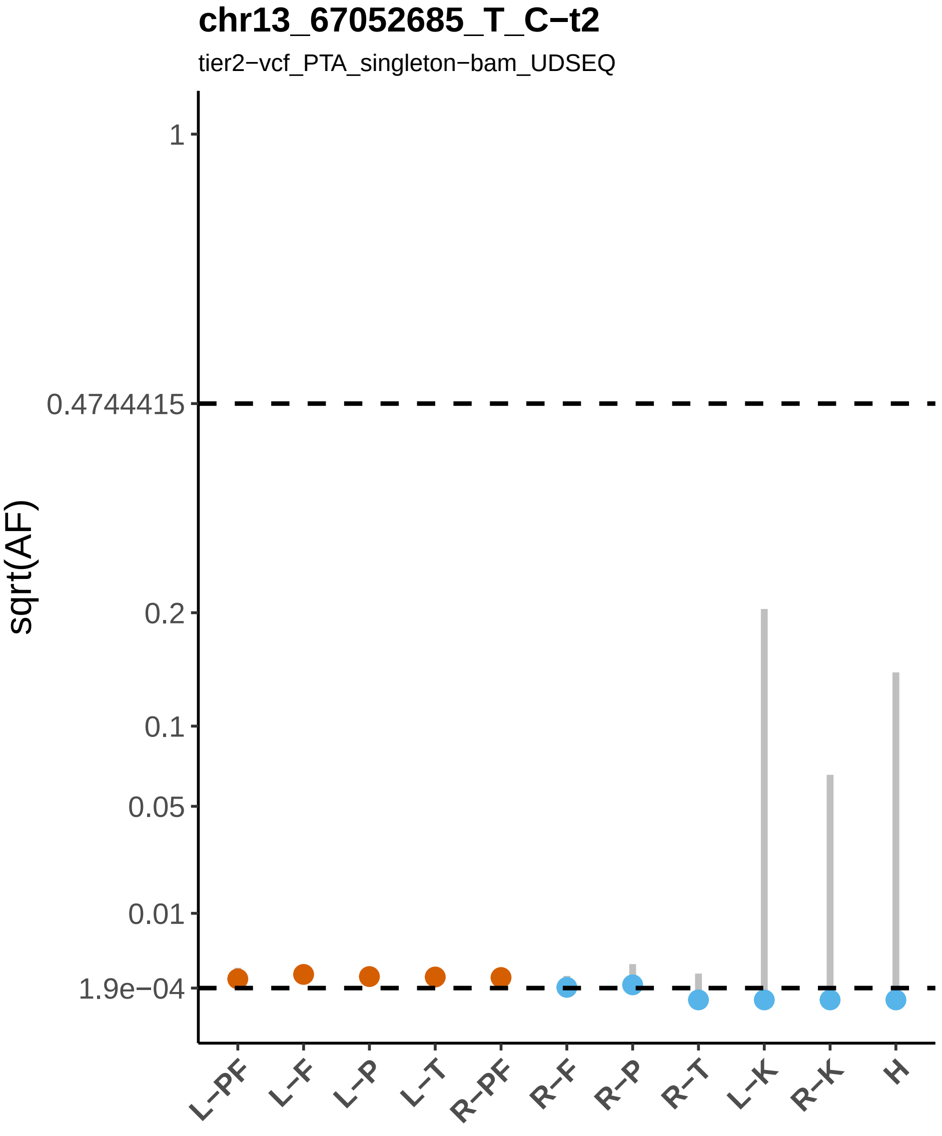
chr13_67052685_T_C−t2
tier2−vcf_PTA_singleton−bam_UDSEQ
sqrt(AF)
1
0.4744415
0.2
0.1
0.05
0.01
1.9e−04
L−PF
L−F
L−P
L−T
R−PF
R−F
R−P
R−T
L−K
R−K
H

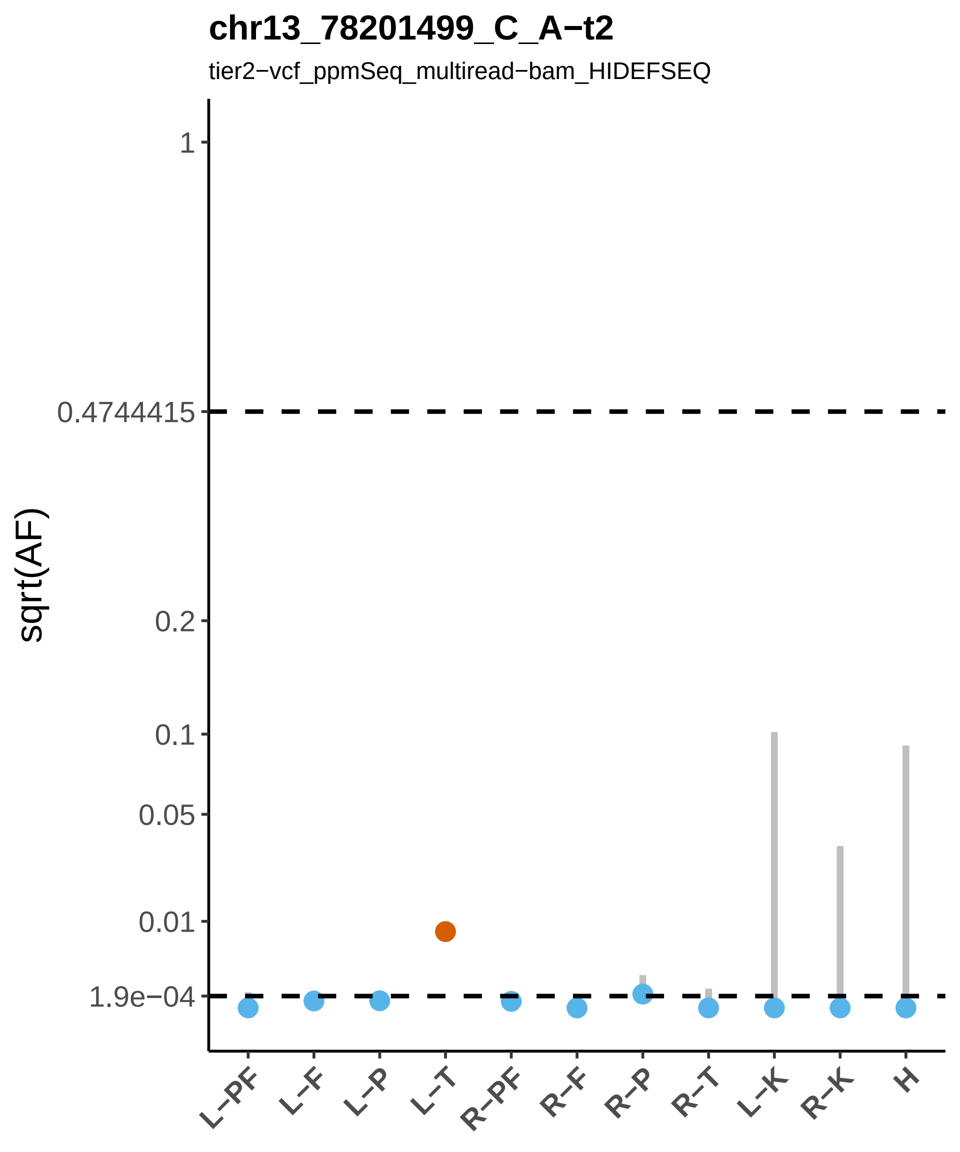

chr13_78201499_C_A−t2
tier2−vcf_ppmSeq_multiread−bam_HIDEFSEQ
sqrt(AF)
1
0.4744415
0.2
0.1
0.05
0.01
1.9e−04
L−PF
L−F
L−P
L−T
R−PF
R−F
R−P
R−T
L−K
R−K
H

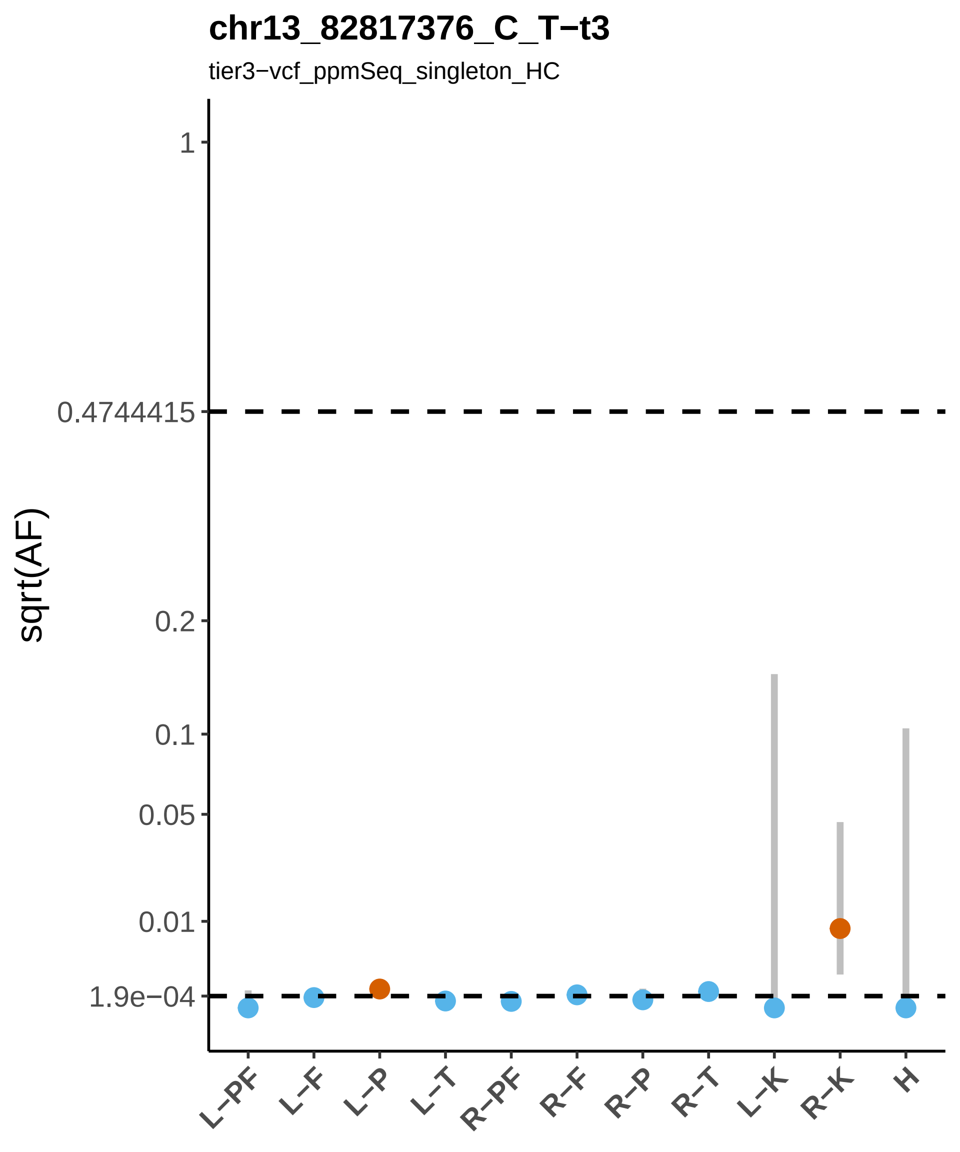

chr13_82817376_C_T−t3
tier3−vcf_ppmSeq_singleton_HC
sqrt(AF)
1
0.4744415
0.2
0.1
0.05
0.01
1.9e−04
L−PF
L−F
L−P
L−T
R−PF
R−F
R−P
R−T
L−K
R−K
H

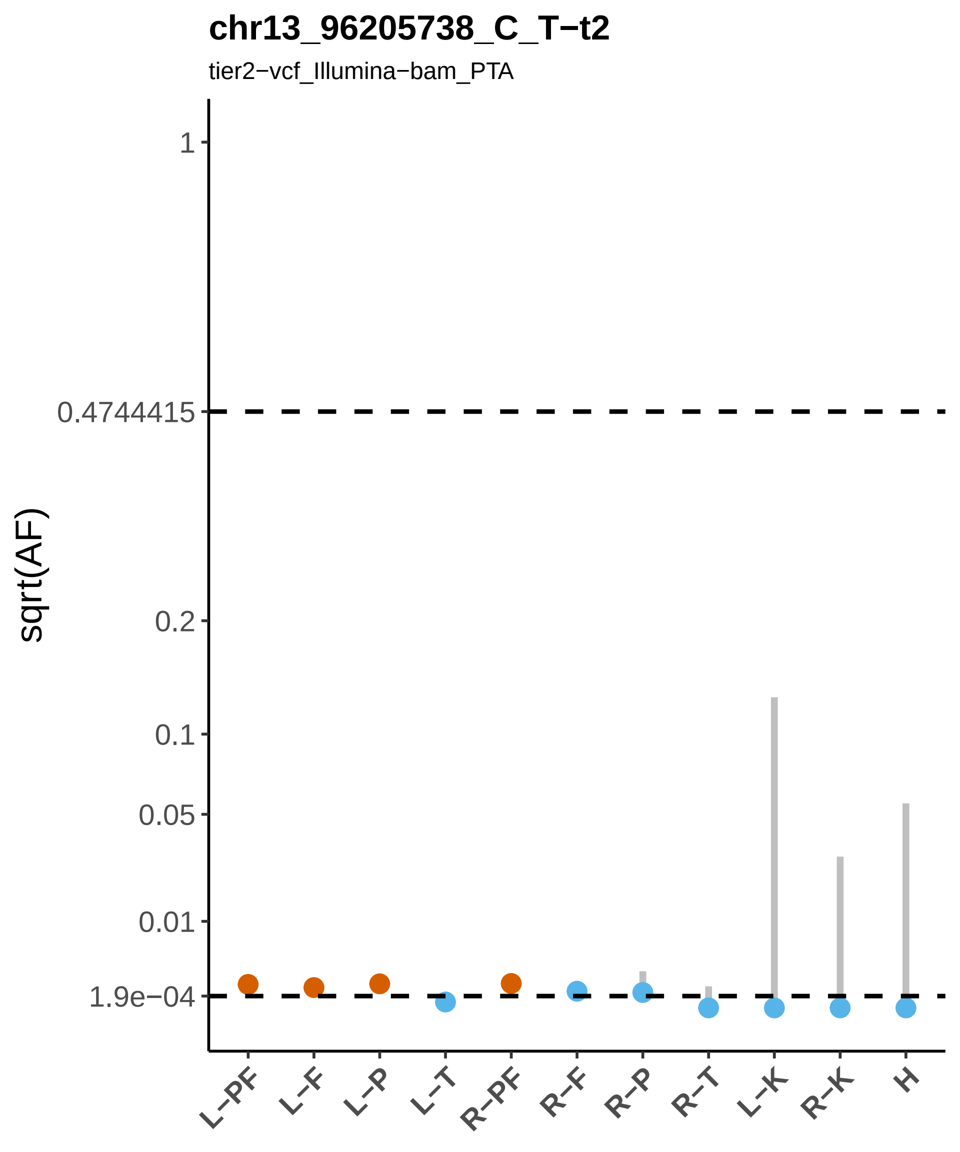

chr13_96205738_C_T−t2
tier2−vcf_Illumina−bam_PTA
sqrt(AF)
1
0.4744415
0.2
0.1
0.05
0.01
1.9e−04
L−PF
L−F
L−P
L−T
R−PF
R−F
R−P
R−T
L−K
R−K
H

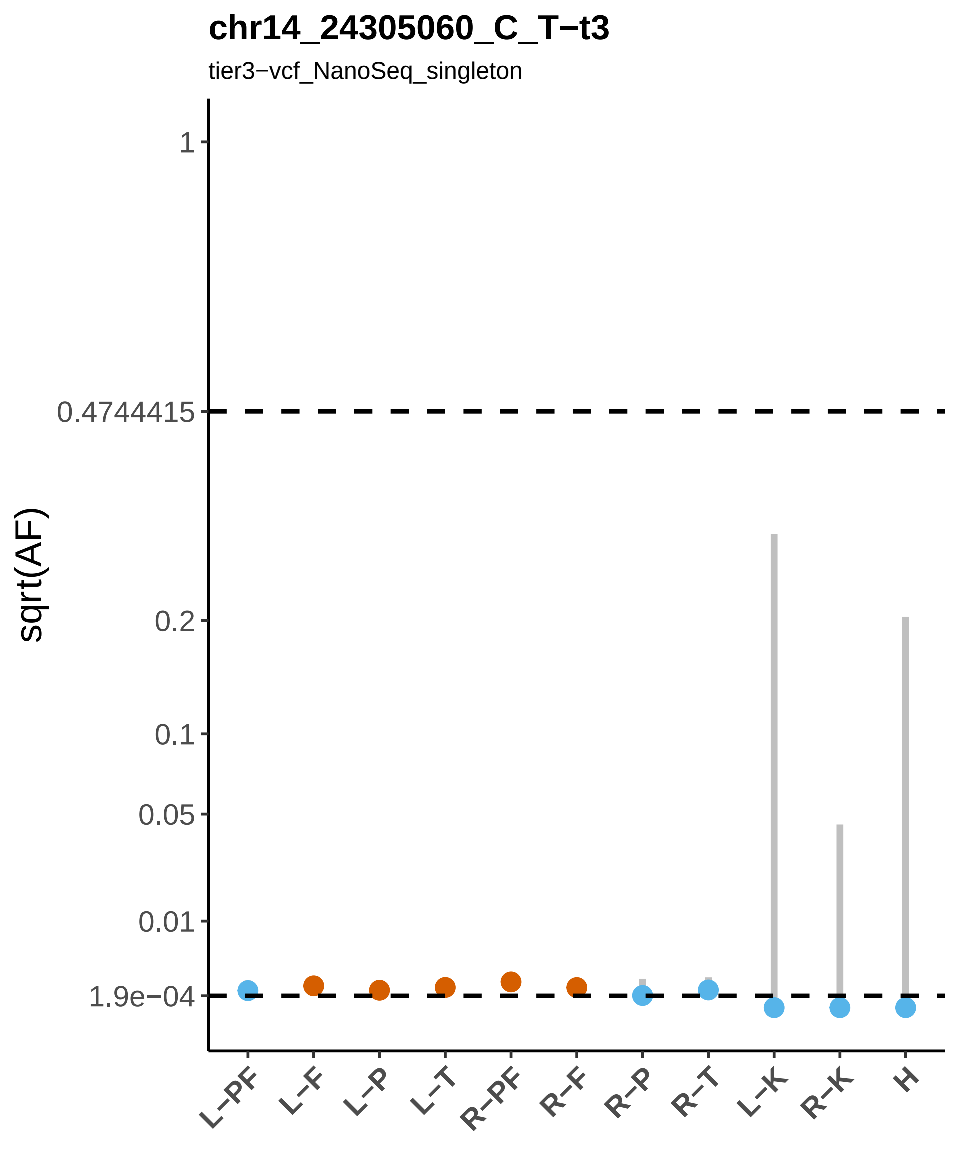

chr14_24305060_C_T−t3
tier3−vcf_NanoSeq_singleton
sqrt(AF)
1
0.4744415
0.2
0.1
0.05
0.01
1.9e−04
L−PF
L−F
L−P
L−T
R−PF
R−F
R−P
R−T
L−K
R−K
H

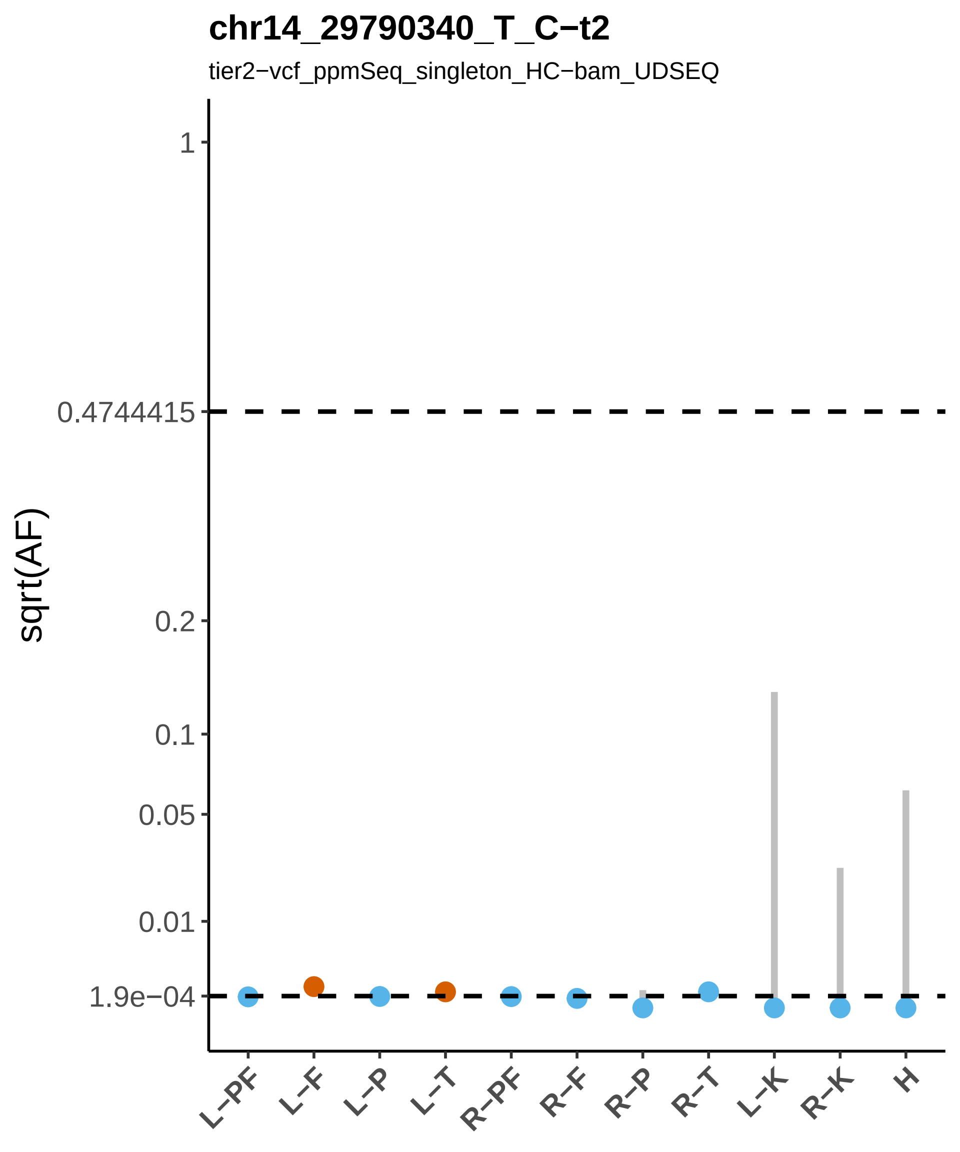

chr14_29790340_T_C−t2
tier2−vcf_ppmSeq_singleton_HC−bam_UDSEQ
sqrt(AF)
1
0.4744415
0.2
0.1
0.05
0.01
1.9e−04
L−PF
L−F
L−P
L−T
R−PF
R−F
R−P
R−T
L−K
R−K
H

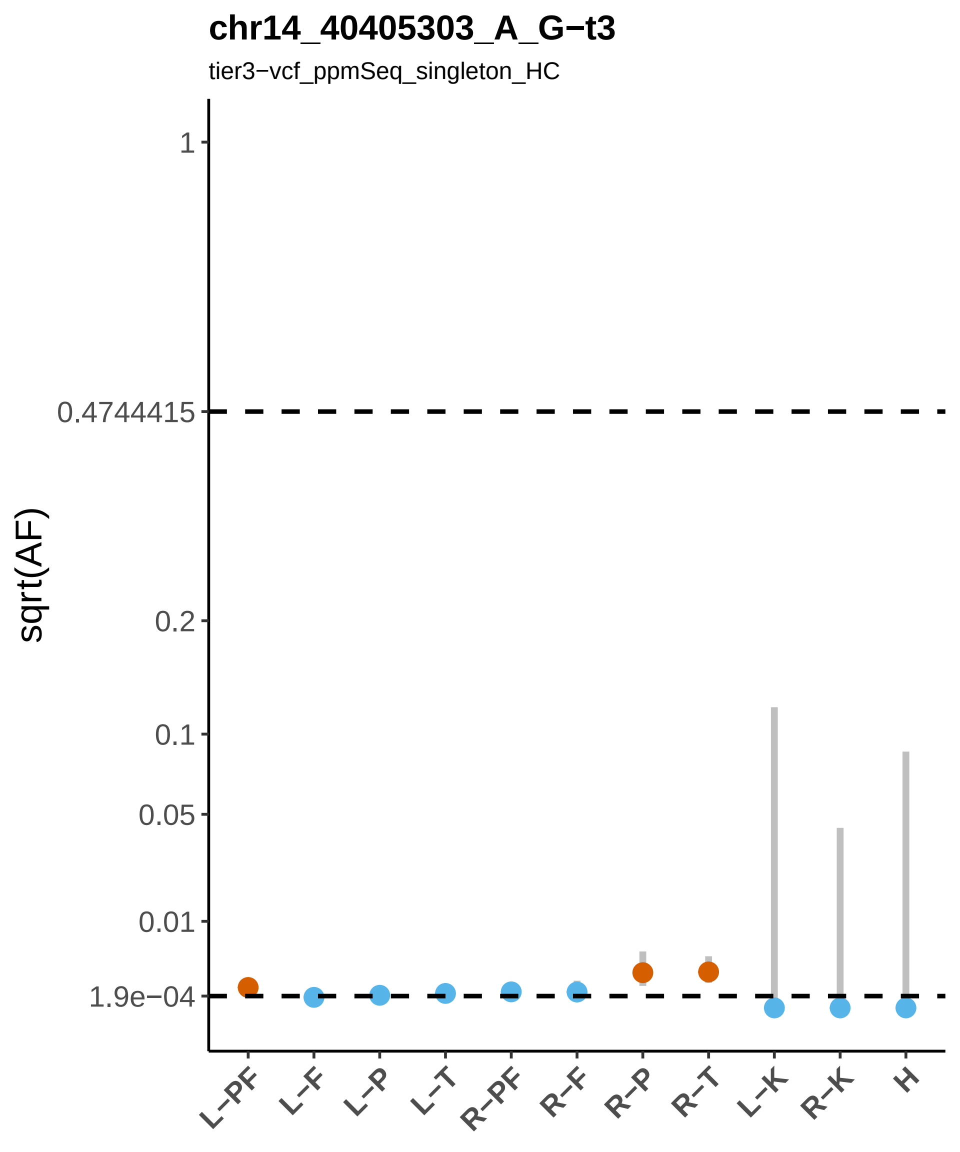

chr14_40405303_A_G−t3
tier3−vcf_ppmSeq_singleton_HC
sqrt(AF)
1
0.4744415
0.2
0.1
0.05
0.01
1.9e−04
L−PF
L−F
L−P
L−T
R−PF
R−F
R−P
R−T
L−K
R−K
H

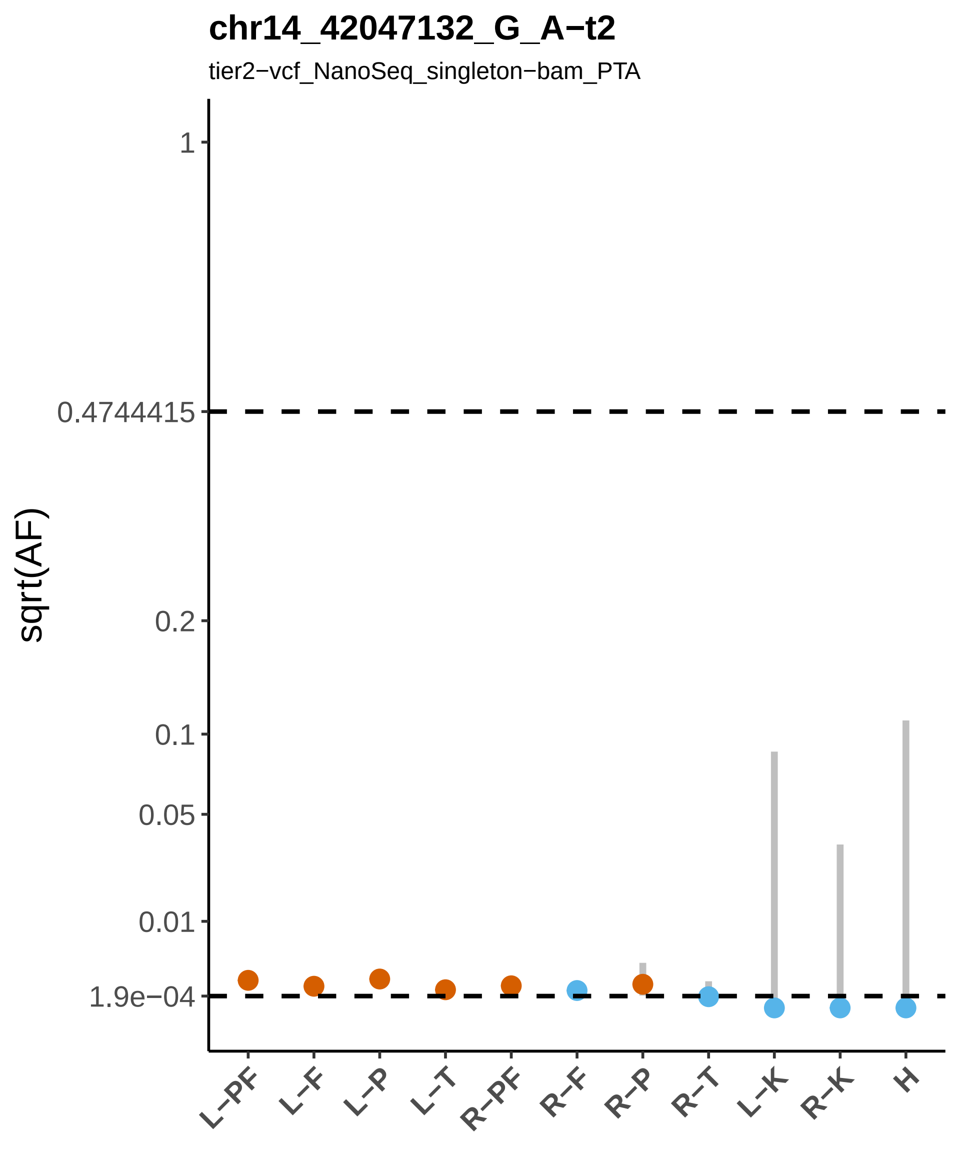
chr14_42047132_G_A−t2
tier2−vcf_NanoSeq_singleton−bam_PTA
sqrt(AF)
1
0.4744415
0.2
0.1
0.05
0.01
1.9e−04
L−PF
L−F
L−P
L−T
R−PF
R−F
R−P
R−T
L−K
R−K
H

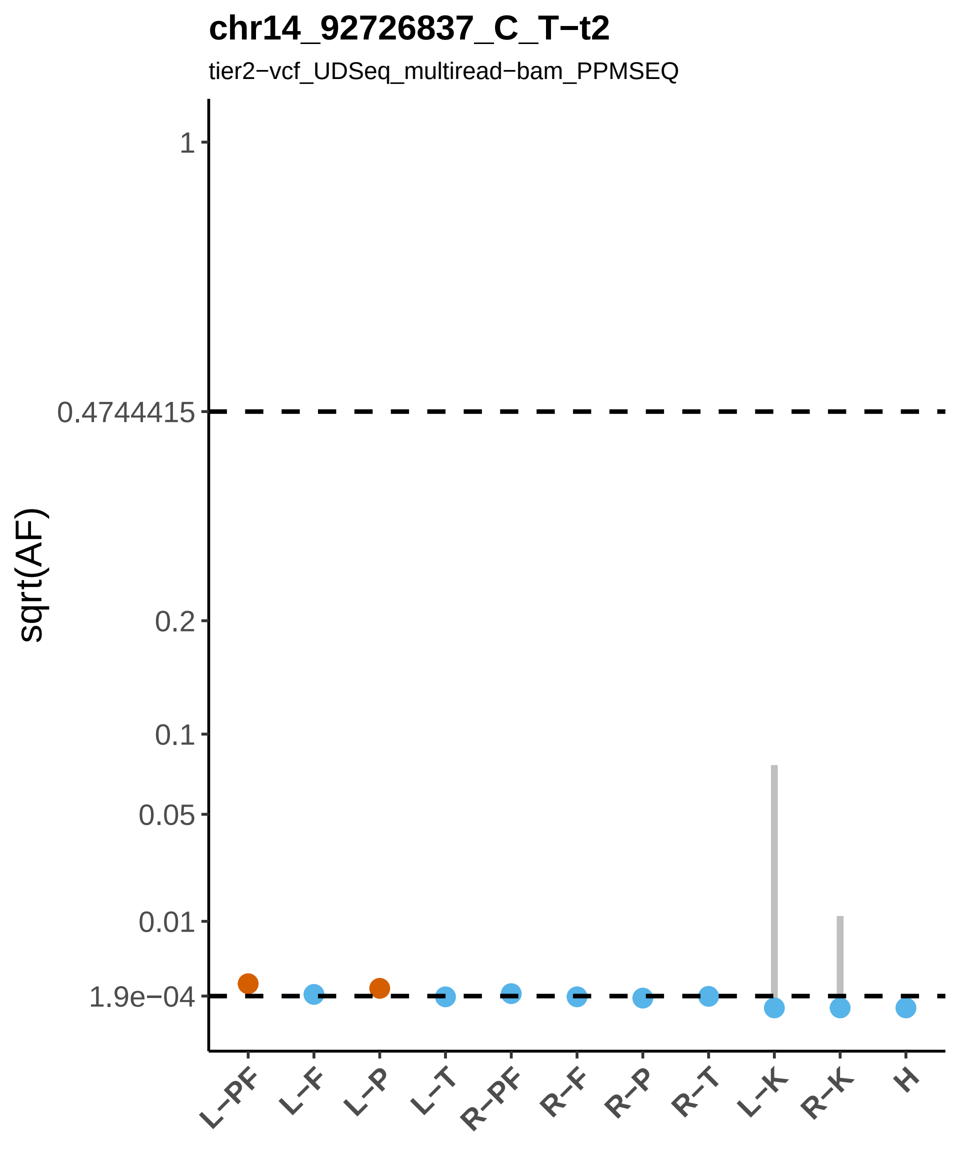

chr14_92726837_C_T−t2
tier2−vcf_UDSeq_multiread−bam_PPMSEQ
sqrt(AF)
1
0.4744415
0.2
0.1
0.05
0.01
1.9e−04
L−PF
L−F
L−P
L−T
R−PF
R−F
R−P
R−T
L−K
R−K
H

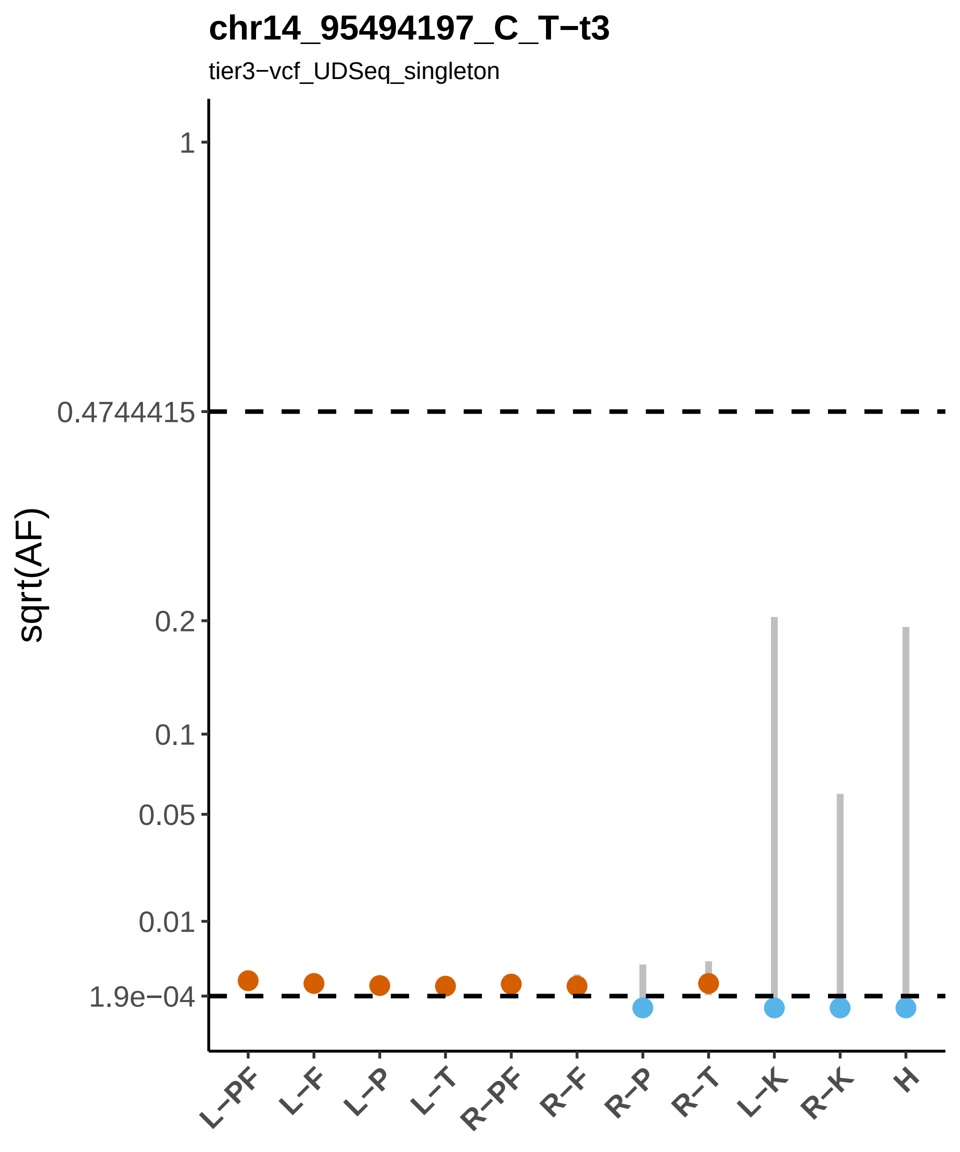

chr14_95494197_C_T−t3
tier3−vcf_UDSeq_singleton
sqrt(AF)
1
0.4744415
0.2
0.1
0.05
0.01
1.9e−04
L−PF
L−F
L−P
L−T
R−PF
R−F
R−P
R−T
L−K
R−K
H

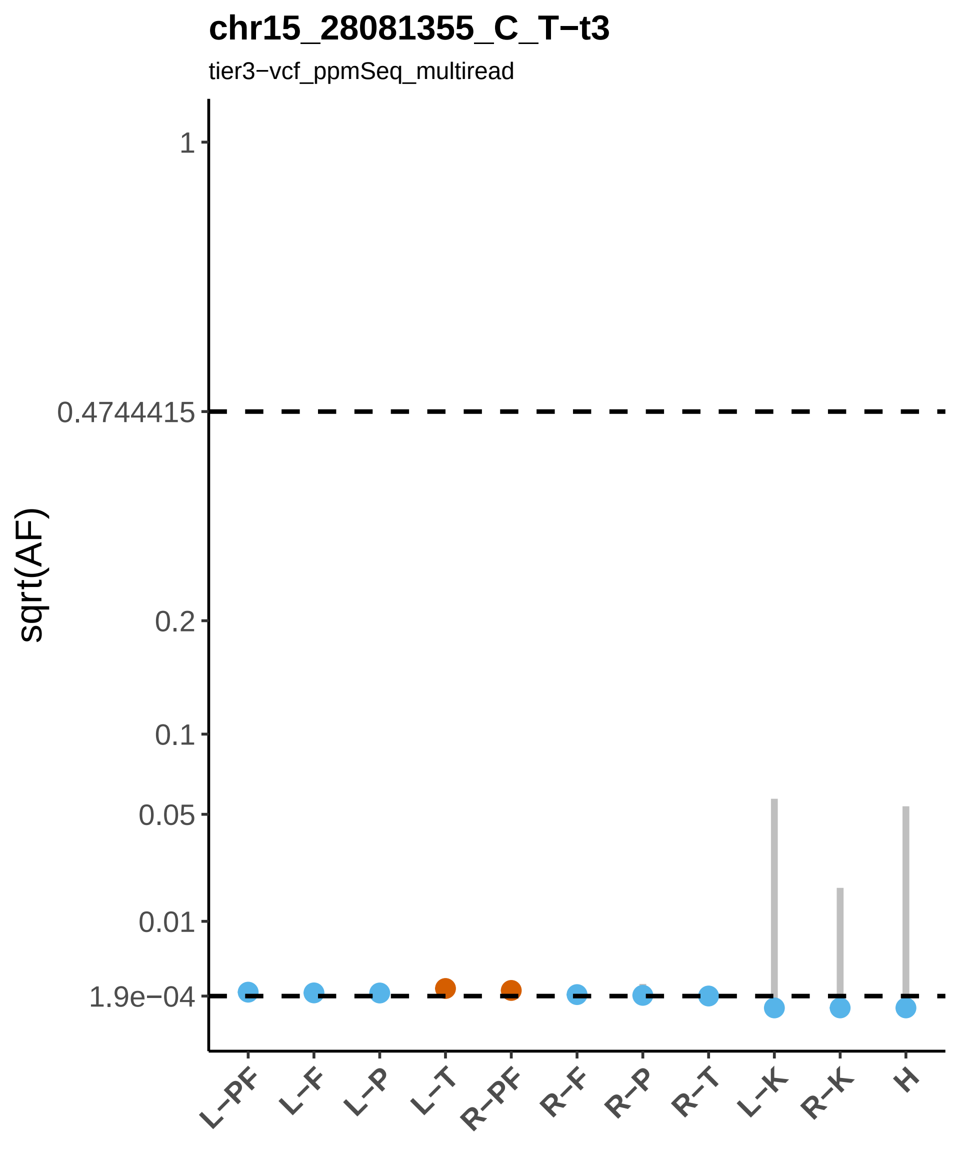

chr15_28081355_C_T−t3
tier3−vcf_ppmSeq_multiread
sqrt(AF)
1
0.4744415
0.2
0.1
0.05
0.01
1.9e−04
L−PF
L−F
L−P
L−T
R−PF
R−F
R−P
R−T
L−K
R−K
H

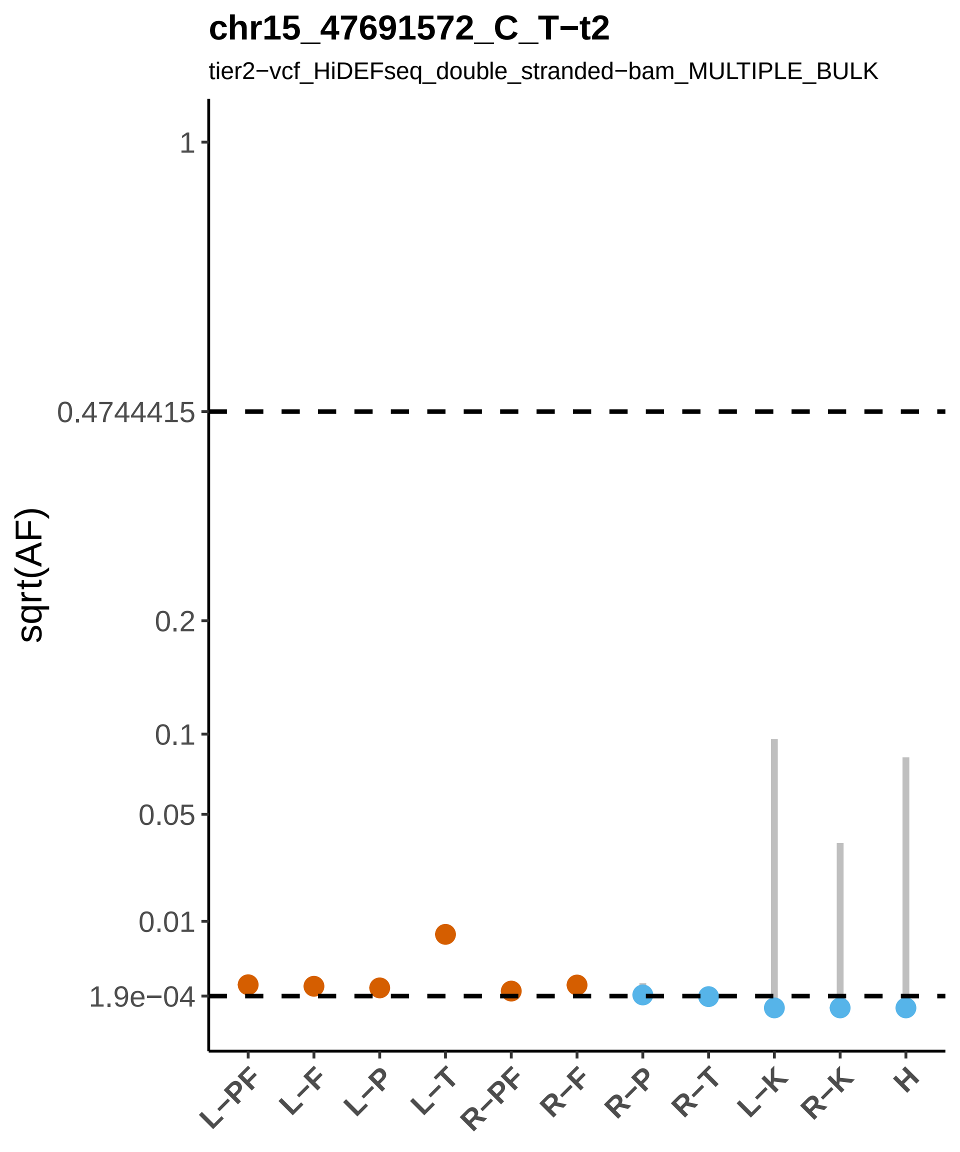
chr15_47691572_C_T−t2
tier2−vcf_HiDEFseq_double_stranded−bam_MULTIPLE_BULK
sqrt(AF)
1
0.4744415
0.2
0.1
0.05
0.01
1.9e−04
L−PF
L−F
L−P
L−T
R−PF
R−F
R−P
R−T
L−K
R−K
H

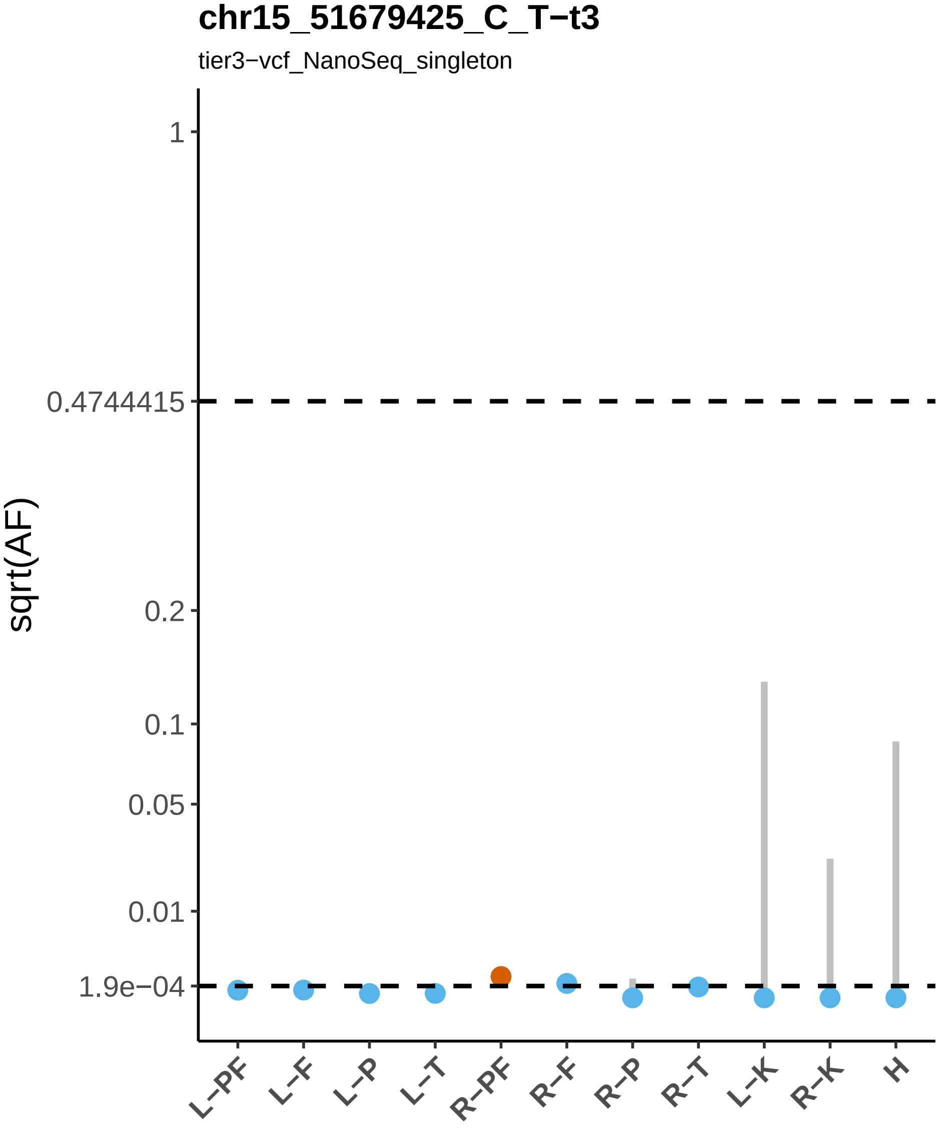
chr15_51679425_C_T−t3
tier3−vcf_NanoSeq_singleton
sqrt(AF)
1
0.4744415
0.2
0.1
0.05
0.01
1.9e−04
L−PF
L−F
L−P
L−T
R−PF
R−F
R−P
R−T
L−K
R−K
H

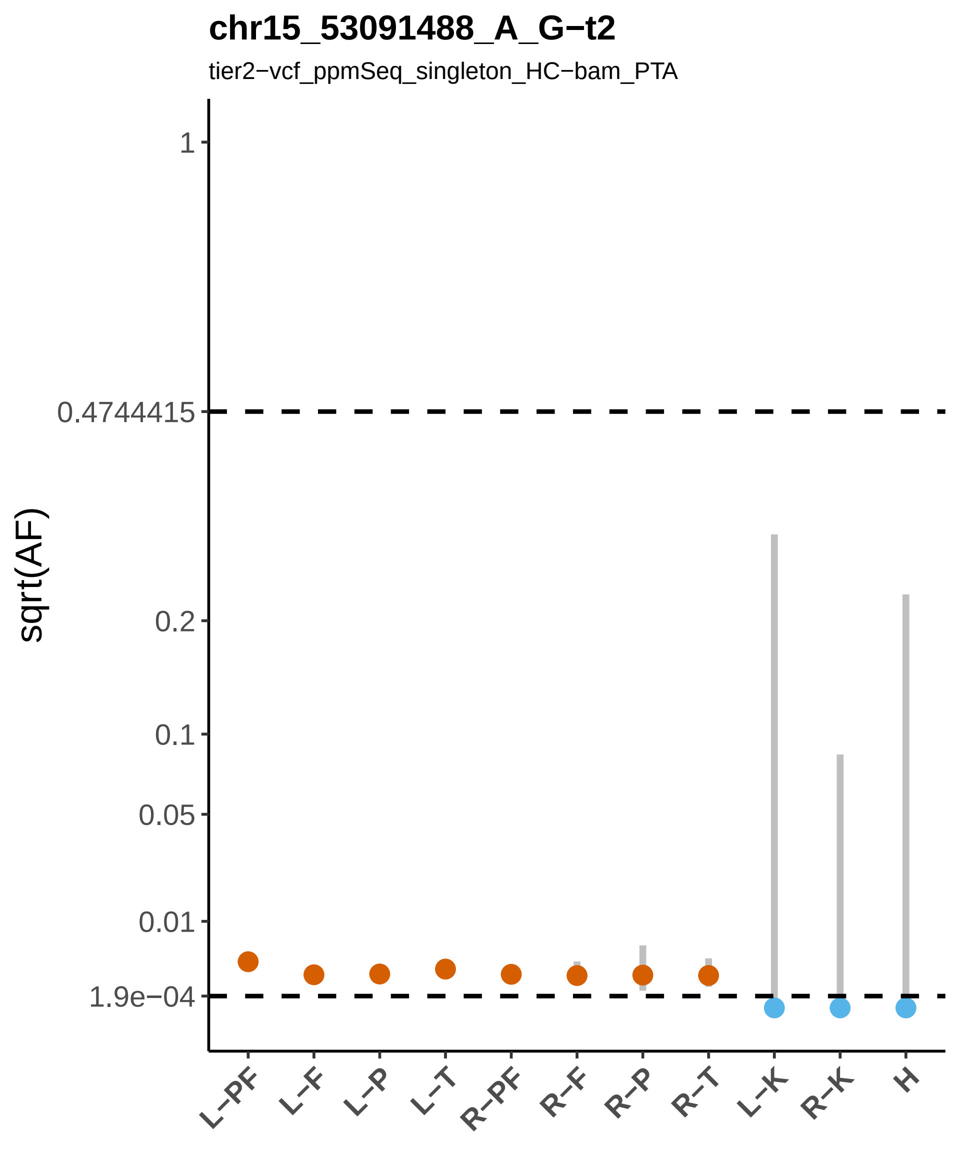

chr15_53091488_A_G−t2
tier2−vcf_ppmSeq_singleton_HC−bam_PTA
sqrt(AF)
1
0.4744415
0.2
0.1
0.05
0.01
1.9e−04
L−PF
L−F
L−P
L−T
R−PF
R−F
R−P
R−T
L−K
R−K
H

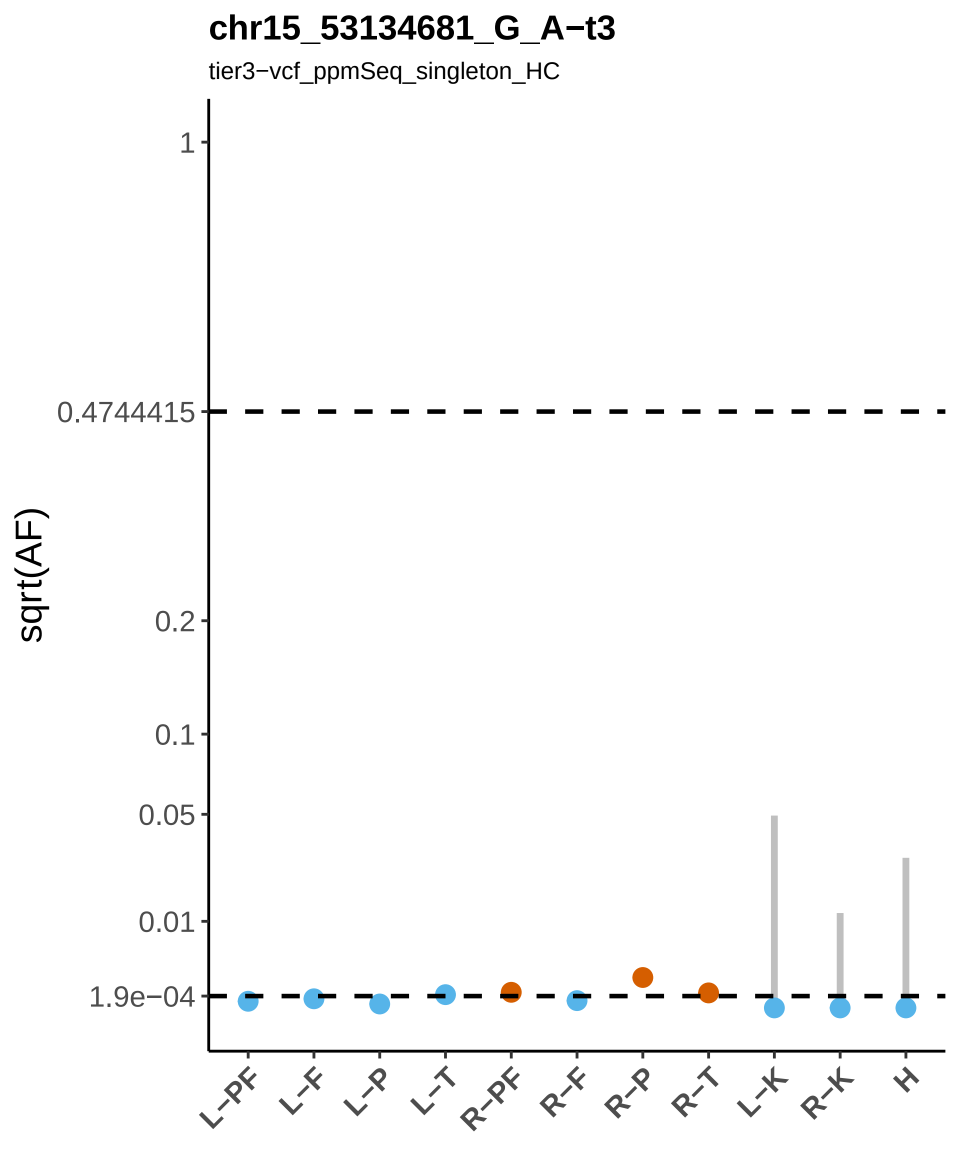
chr15_53134681_G_A−t3
tier3−vcf_ppmSeq_singleton_HC
sqrt(AF)
1
0.4744415
0.2
0.1
0.05
0.01
1.9e−04
L−PF
L−F
L−P
L−T
R−PF
R−F
R−P
R−T
L−K
R−K
H

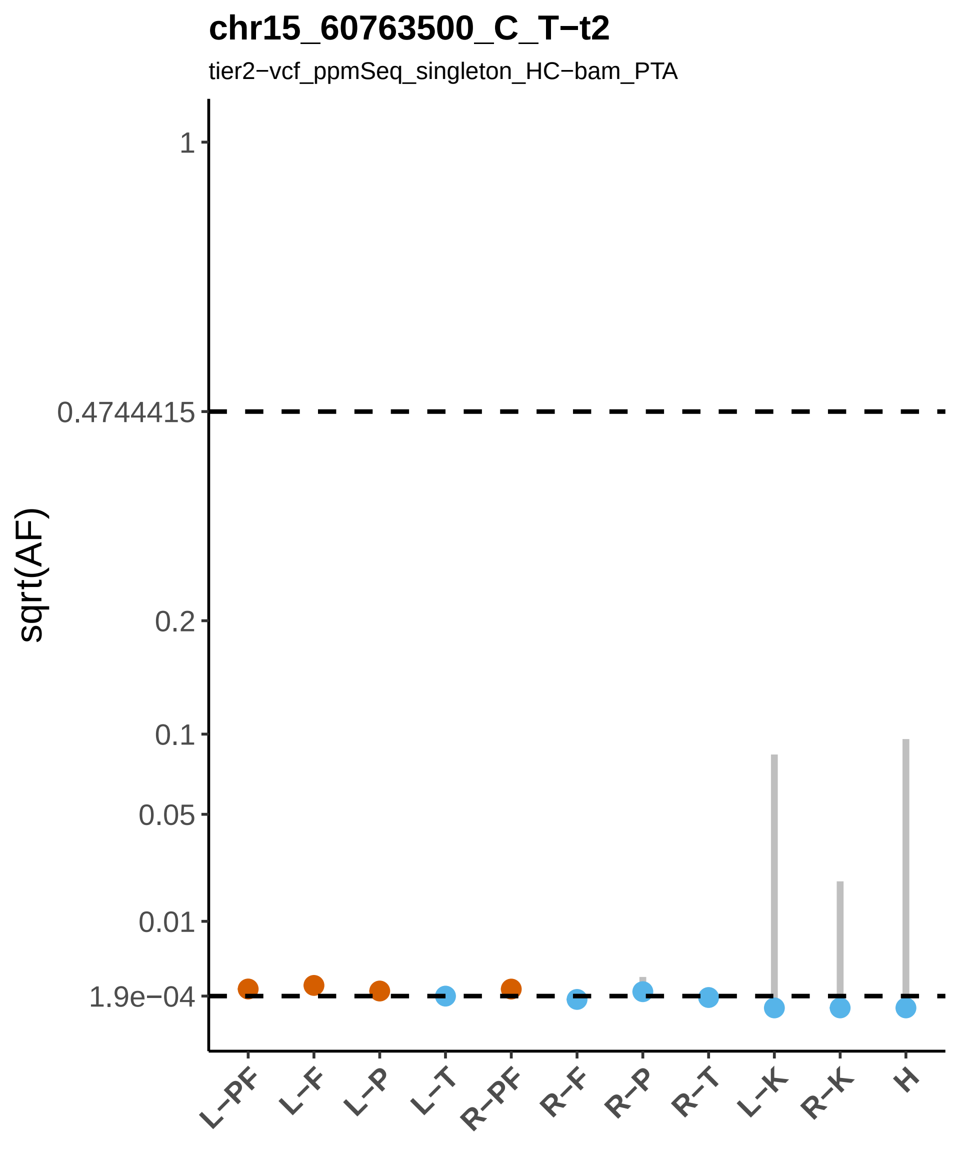

chr15_60763500_C_T−t2
tier2−vcf_ppmSeq_singleton_HC−bam_PTA
sqrt(AF)
1
0.4744415
0.2
0.1
0.05
0.01
1.9e−04
L−PF
L−F
L−P
L−T
R−PF
R−F
R−P
R−T
L−K
R−K
H

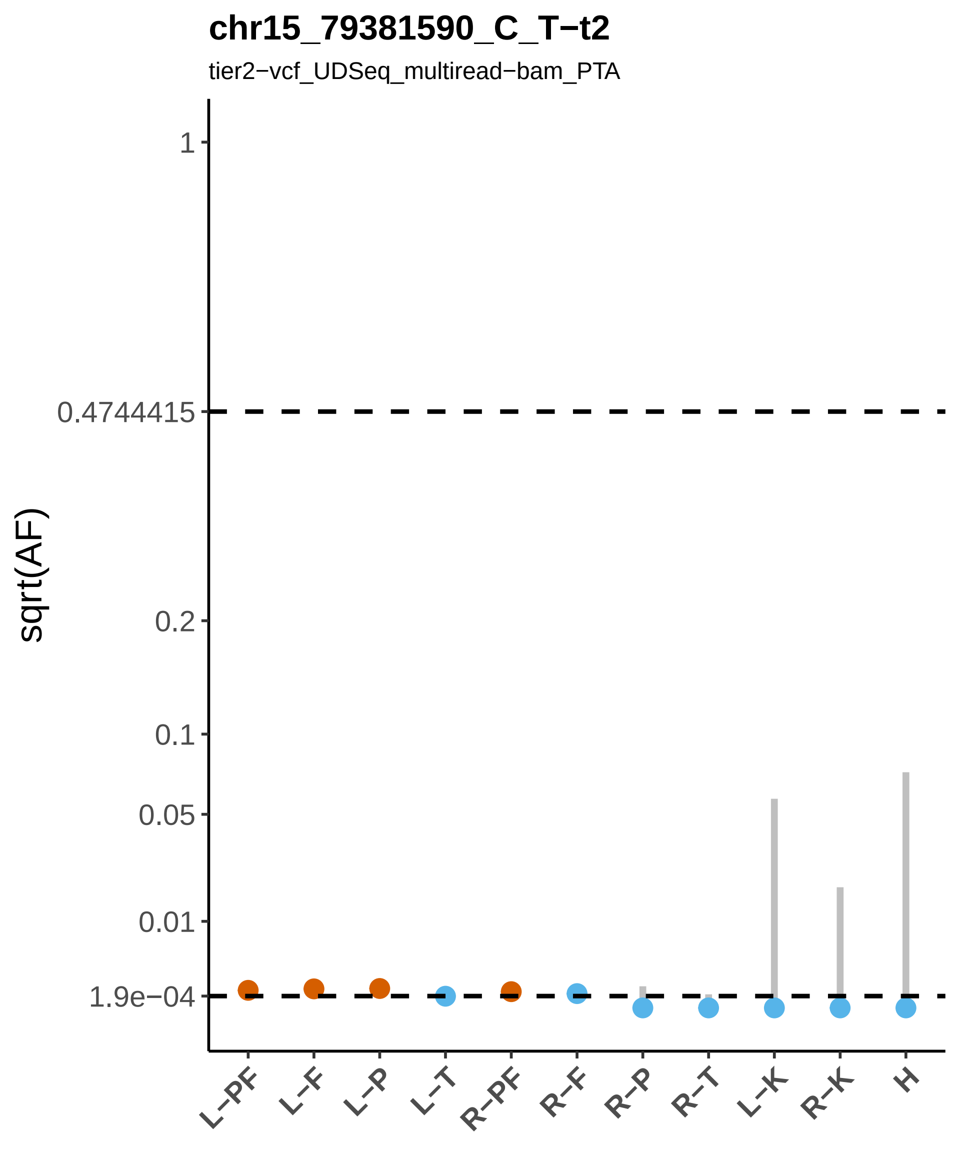
chr15_79381590_C_T−t2
tier2−vcf_UDSeq_multiread−bam_PTA
sqrt(AF)
1
0.4744415
0.2
0.1
0.05
0.01
1.9e−04
L−PF
L−F
L−P
L−T
R−PF
R−F
R−P
R−T
L−K
R−K
H

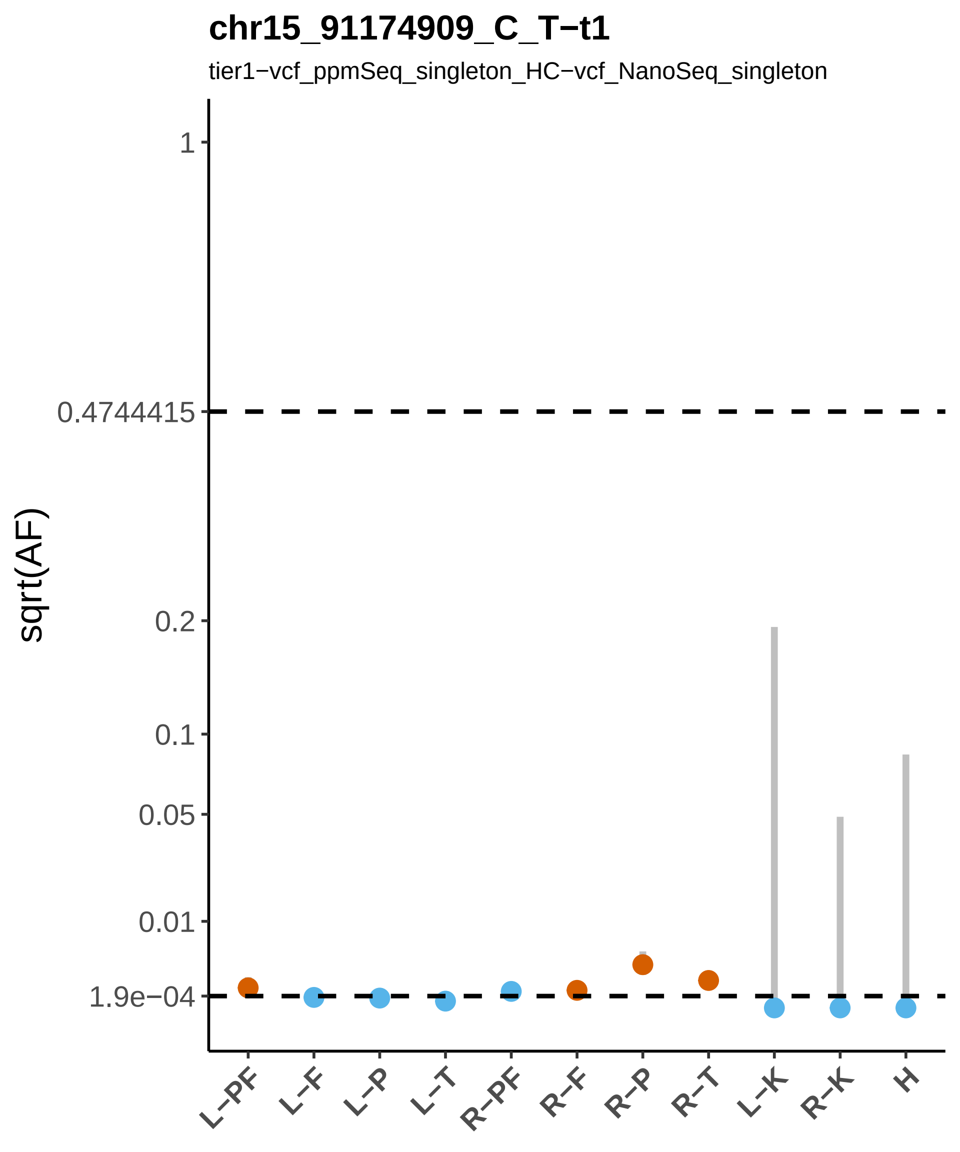

chr15_91174909_C_T−t1
tier1−vcf_ppmSeq_singleton_HC−vcf_NanoSeq_singleton
sqrt(AF)
1
0.4744415
0.2
0.1
0.05
0.01
1.9e−04
L−PF
L−F
L−P
L−T
R−PF
R−F
R−P
R−T
L−K
R−K
H

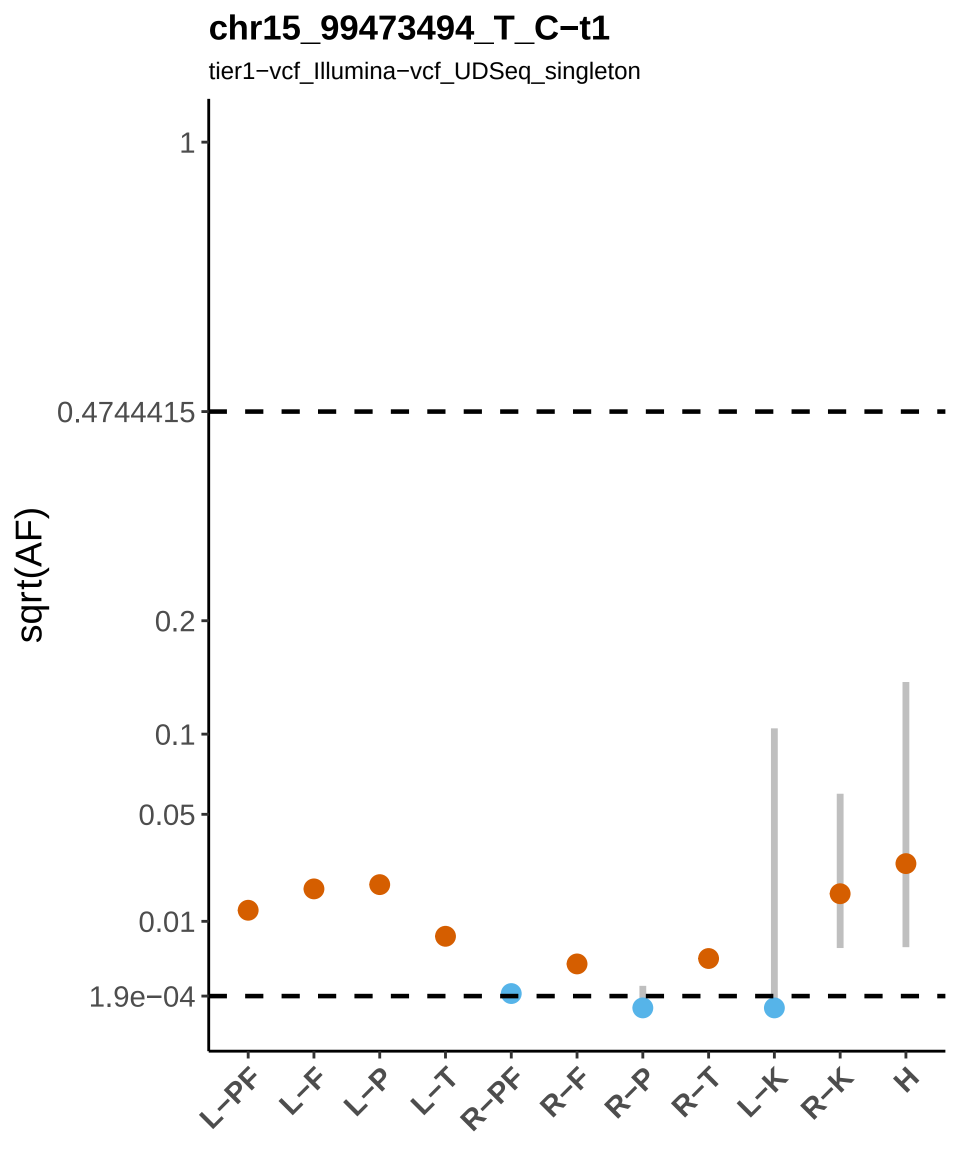

chr15_99473494_T_C−t1
tier1−vcf_Illumina−vcf_UDSeq_singleton
sqrt(AF)
1
0.4744415
0.2
0.1
0.05
0.01
1.9e−04
L−PF
L−F
L−P
L−T
R−PF
R−F
R−P
R−T
L−K
R−K
H

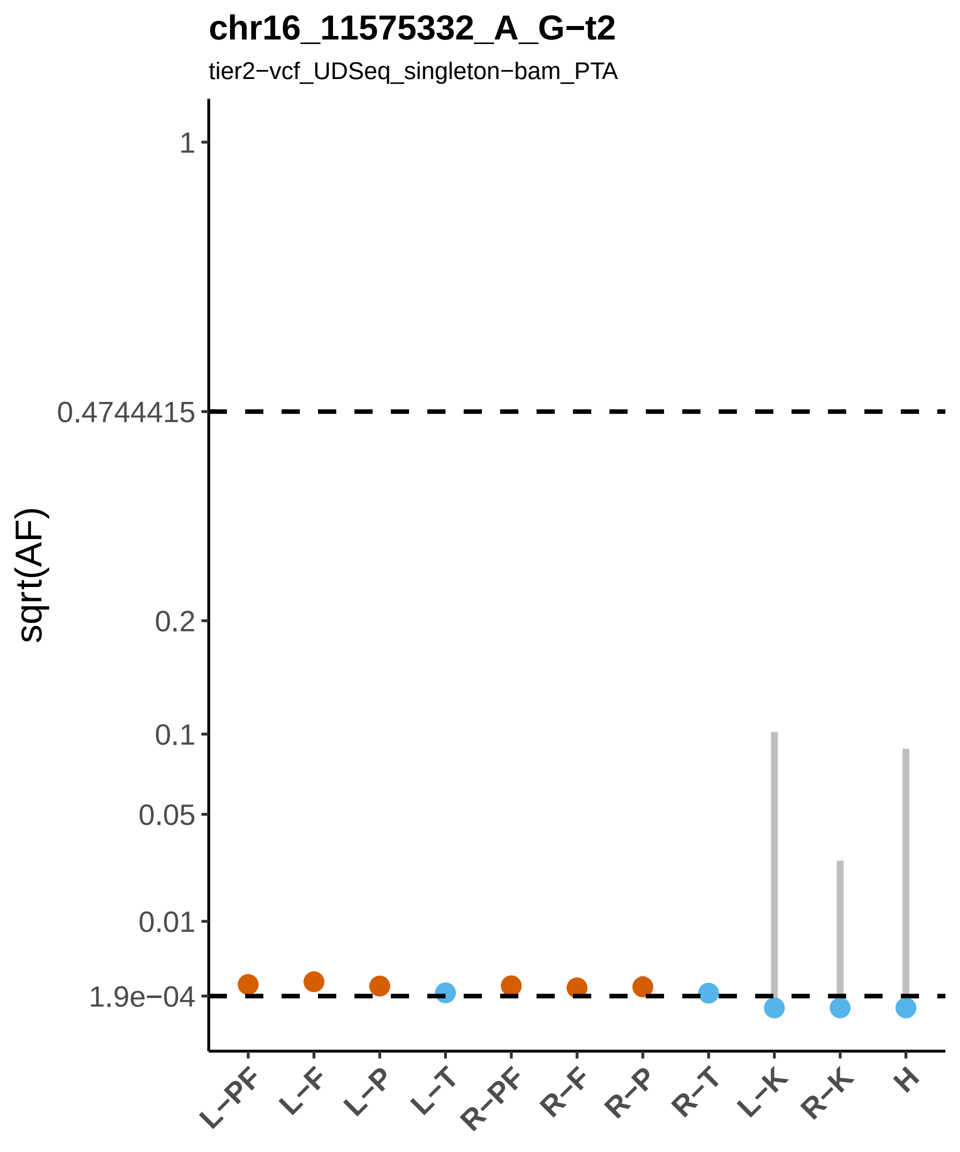
chr16_11575332_A_G−t2
tier2−vcf_UDSeq_singleton−bam_PTA
sqrt(AF)
1
0.4744415
0.2
0.1
0.05
0.01
1.9e−04
L−PF
L−F
L−P
L−T
R−PF
R−F
R−P
R−T
L−K
R−K
H

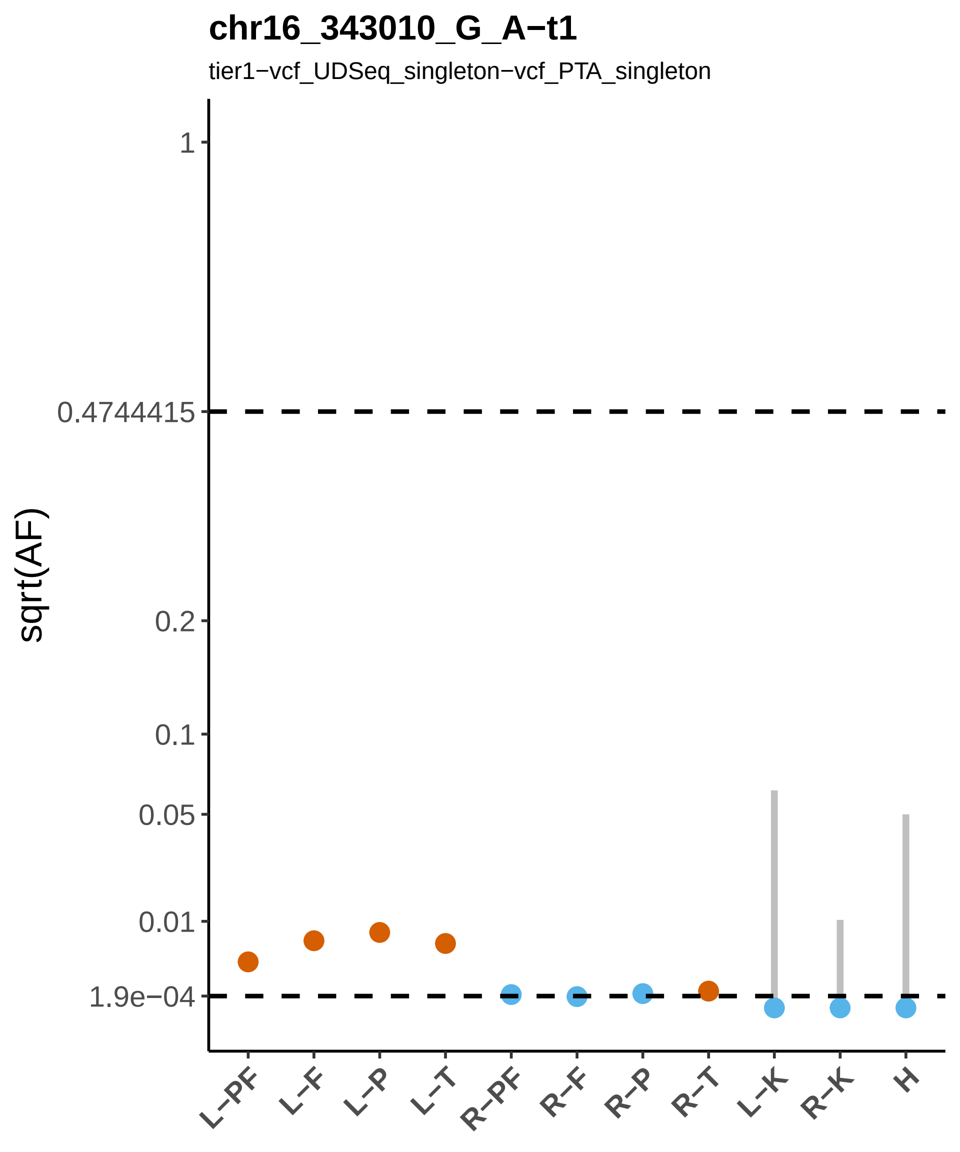

chr16_343010_G_A−t1
tier1−vcf_UDSeq_singleton−vcf_PTA_singleton
sqrt(AF)
1
0.4744415
0.2
0.1
0.05
0.01
1.9e−04
L−PF
L−F
L−P
L−T
R−PF
R−F
R−P
R−T
L−K
R−K
H

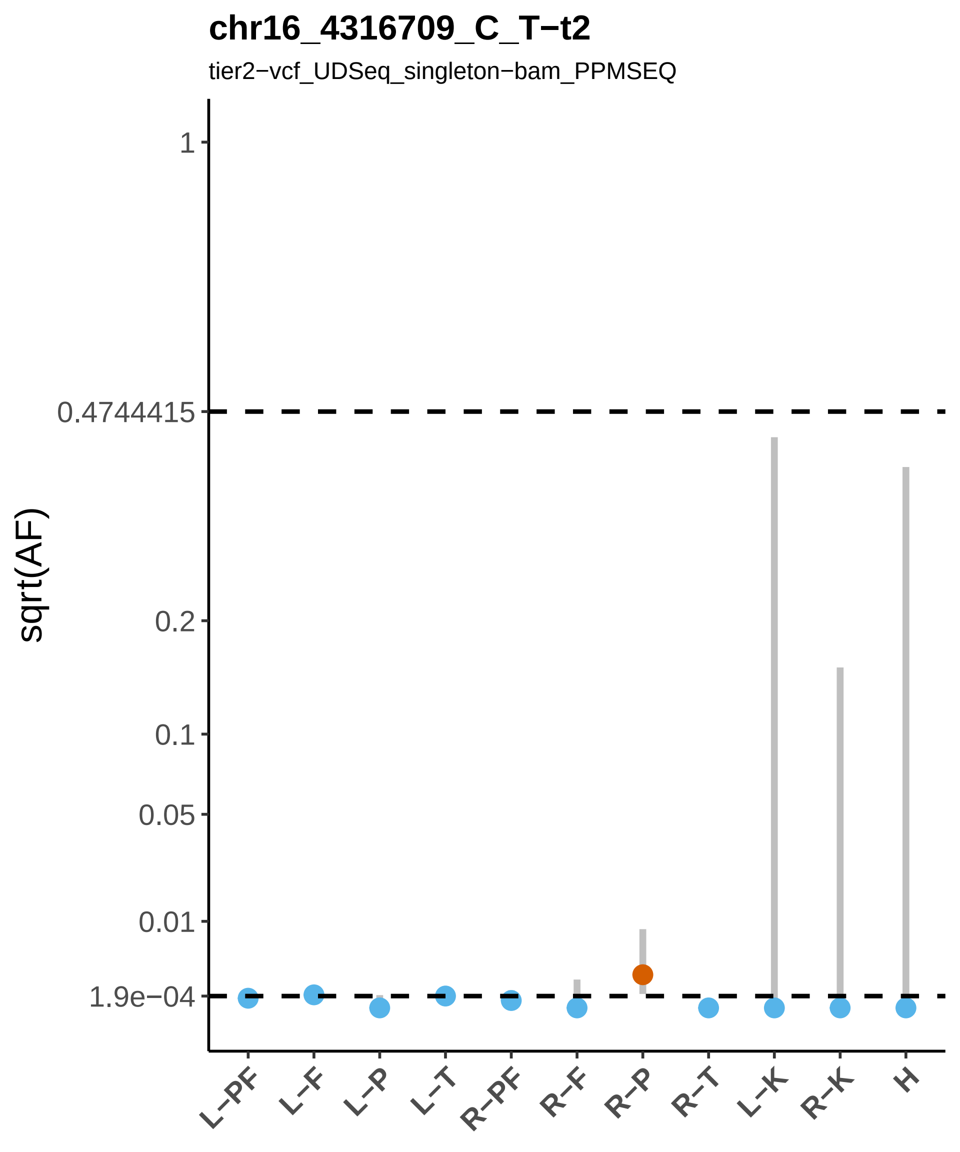

chr16_4316709_C_T−t2
tier2−vcf_UDSeq_singleton−bam_PPMSEQ
sqrt(AF)
1
0.4744415
0.2
0.1
0.05
0.01
1.9e−04
L−PF
L−F
L−P
L−T
R−PF
R−F
R−P
R−T
L−K
R−K
H

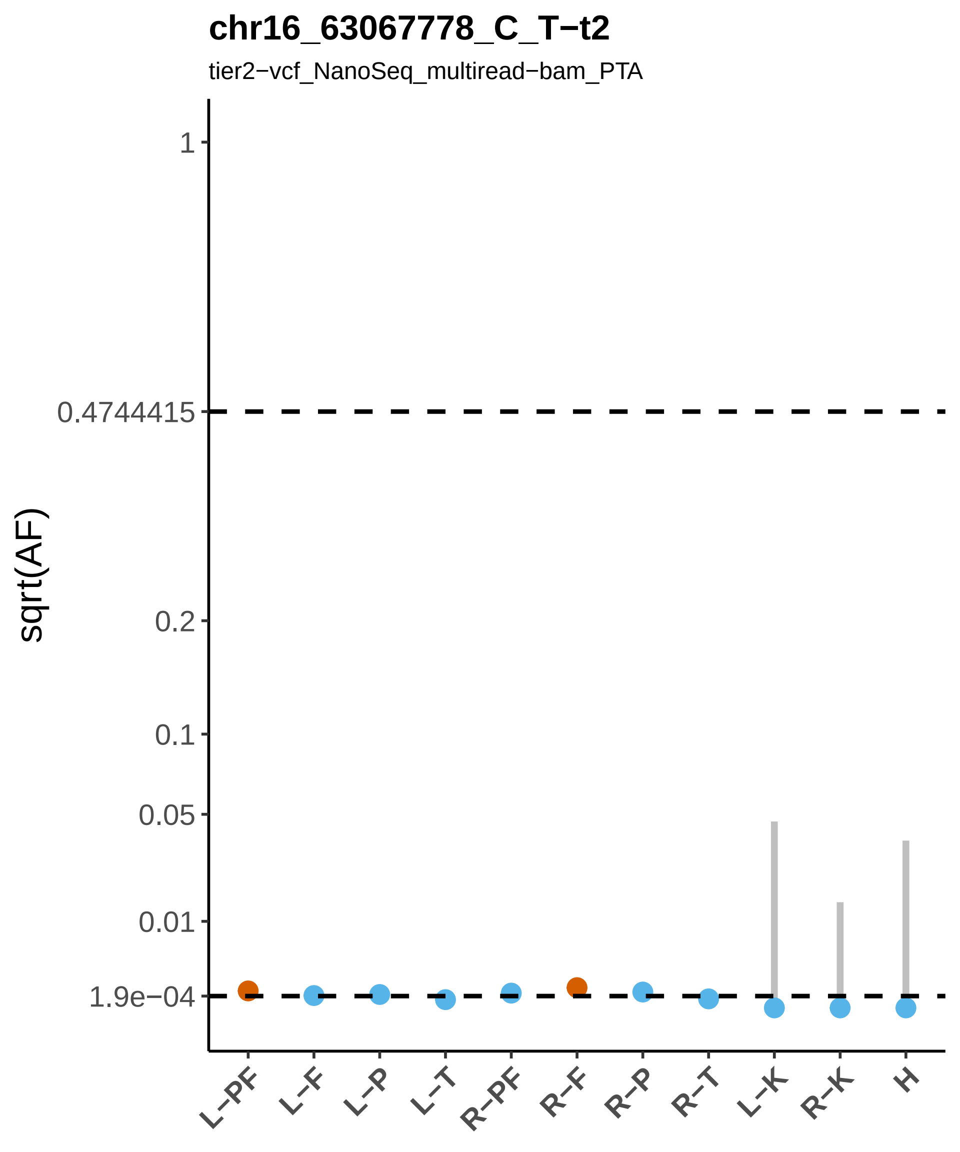

chr16_63067778_C_T−t2
tier2−vcf_NanoSeq_multiread−bam_PTA
sqrt(AF)
1
0.4744415
0.2
0.1
0.05
0.01
1.9e−04
L−PF
L−F
L−P
L−T
R−PF
R−F
R−P
R−T
L−K
R−K
H

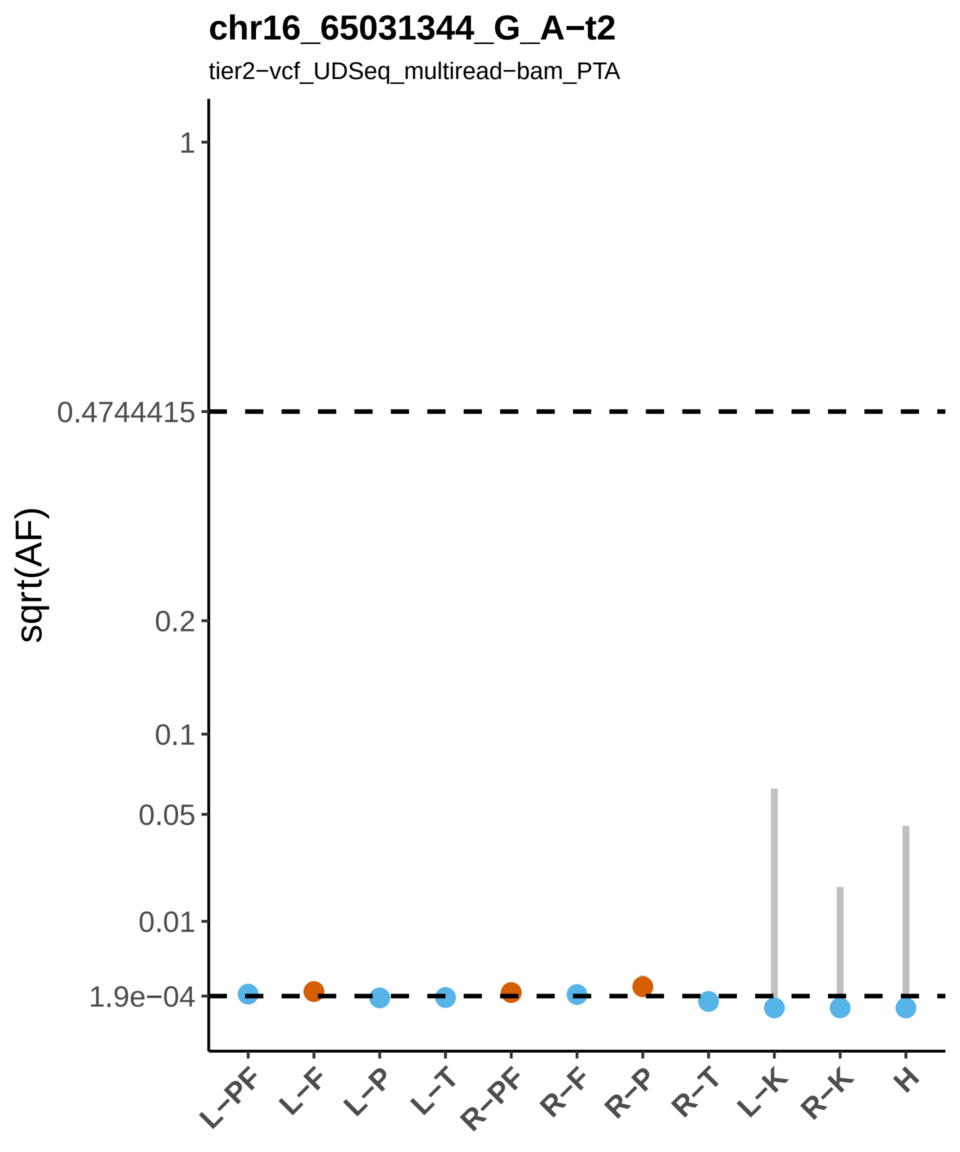
chr16_65031344_G_A−t2
tier2−vcf_UDSeq_multiread−bam_PTA
sqrt(AF)
1
0.4744415
0.2
0.1
0.05
0.01
1.9e−04
L−PF
L−F
L−P
L−T
R−PF
R−F
R−P
R−T
L−K
R−K
H

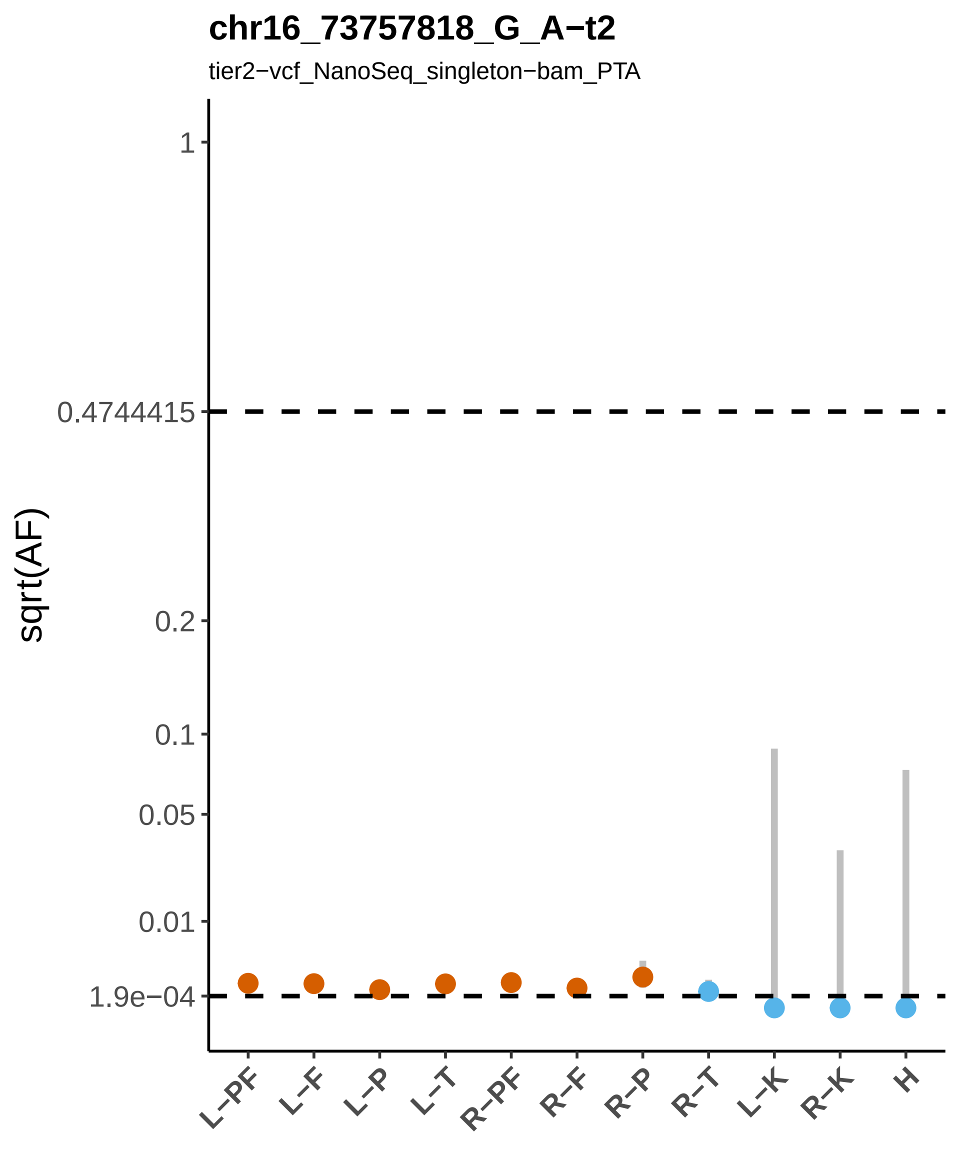
chr16_73757818_G_A−t2
tier2−vcf_NanoSeq_singleton−bam_PTA
sqrt(AF)
1
0.4744415
0.2
0.1
0.05
0.01
1.9e−04
L−PF
L−F
L−P
L−T
R−PF
R−F
R−P
R−T
L−K
R−K
H

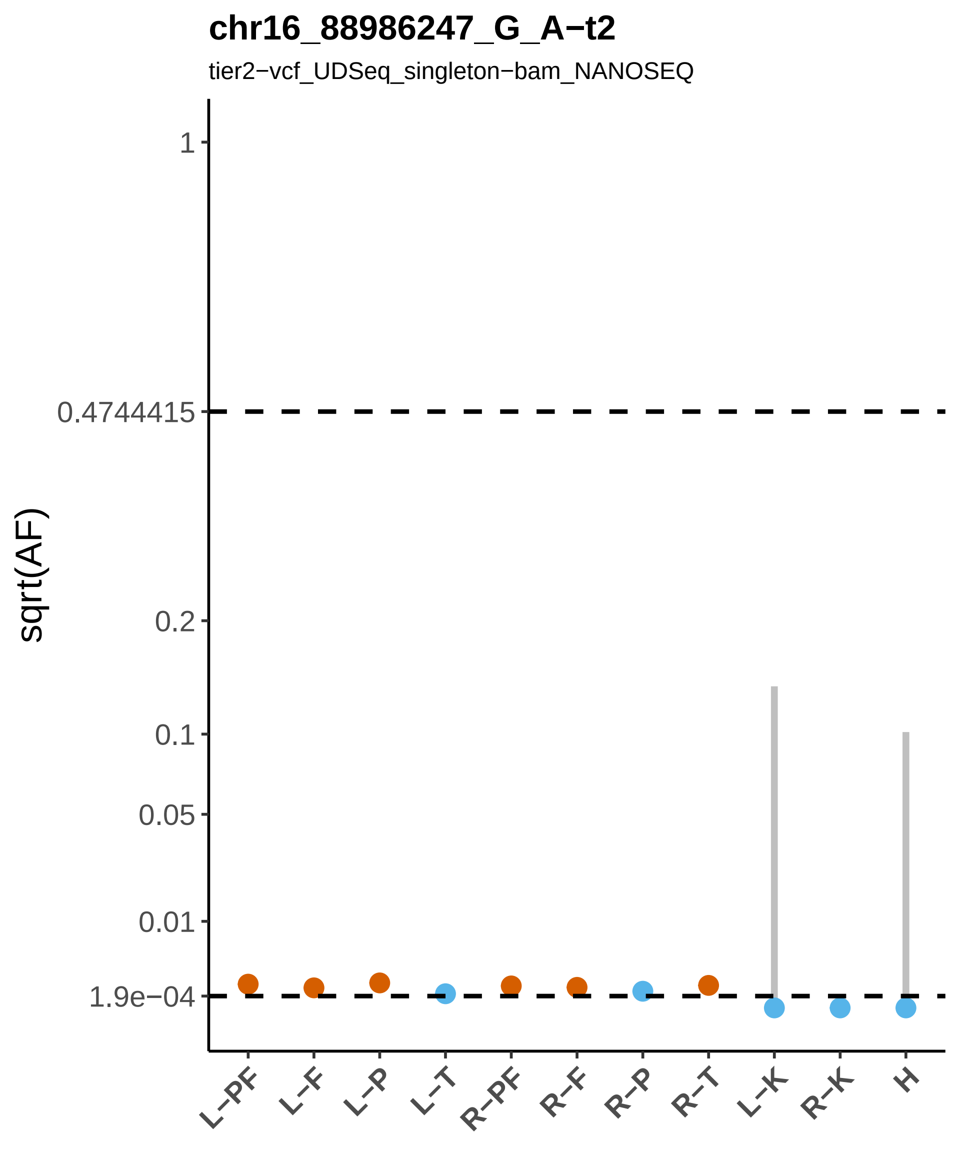
chr16_88986247_G_A−t2
tier2−vcf_UDSeq_singleton−bam_NANOSEQ
sqrt(AF)
1
0.4744415
0.2
0.1
0.05
0.01
1.9e−04
L−PF
L−F
L−P
L−T
R−PF
R−F
R−P
R−T
L−K
R−K
H

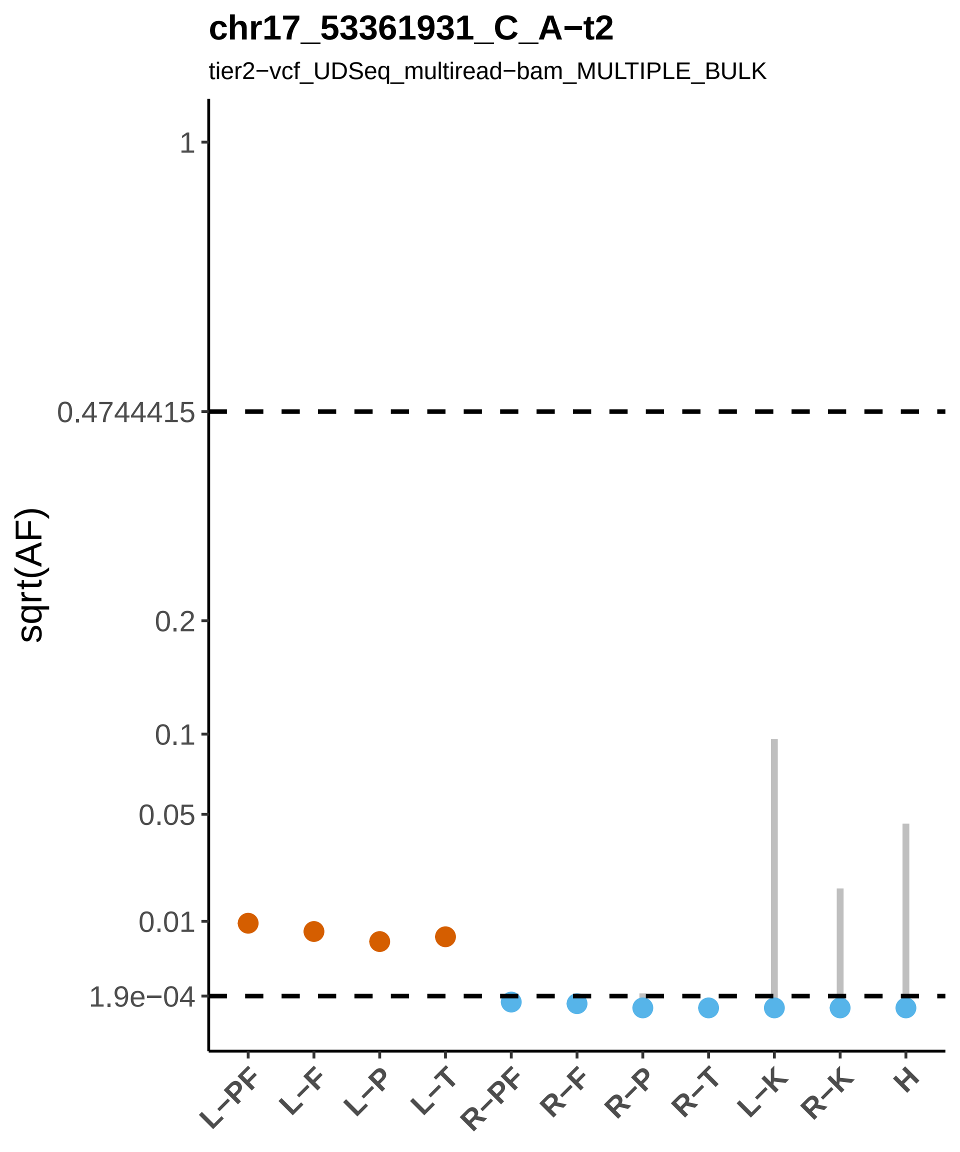

chr17_53361931_C_A−t2
tier2−vcf_UDSeq_multiread−bam_MULTIPLE_BULK
sqrt(AF)
1
0.4744415
0.2
0.1
0.05
0.01
1.9e−04
L−PF
L−F
L−P
L−T
R−PF
R−F
R−P
R−T
L−K
R−K
H

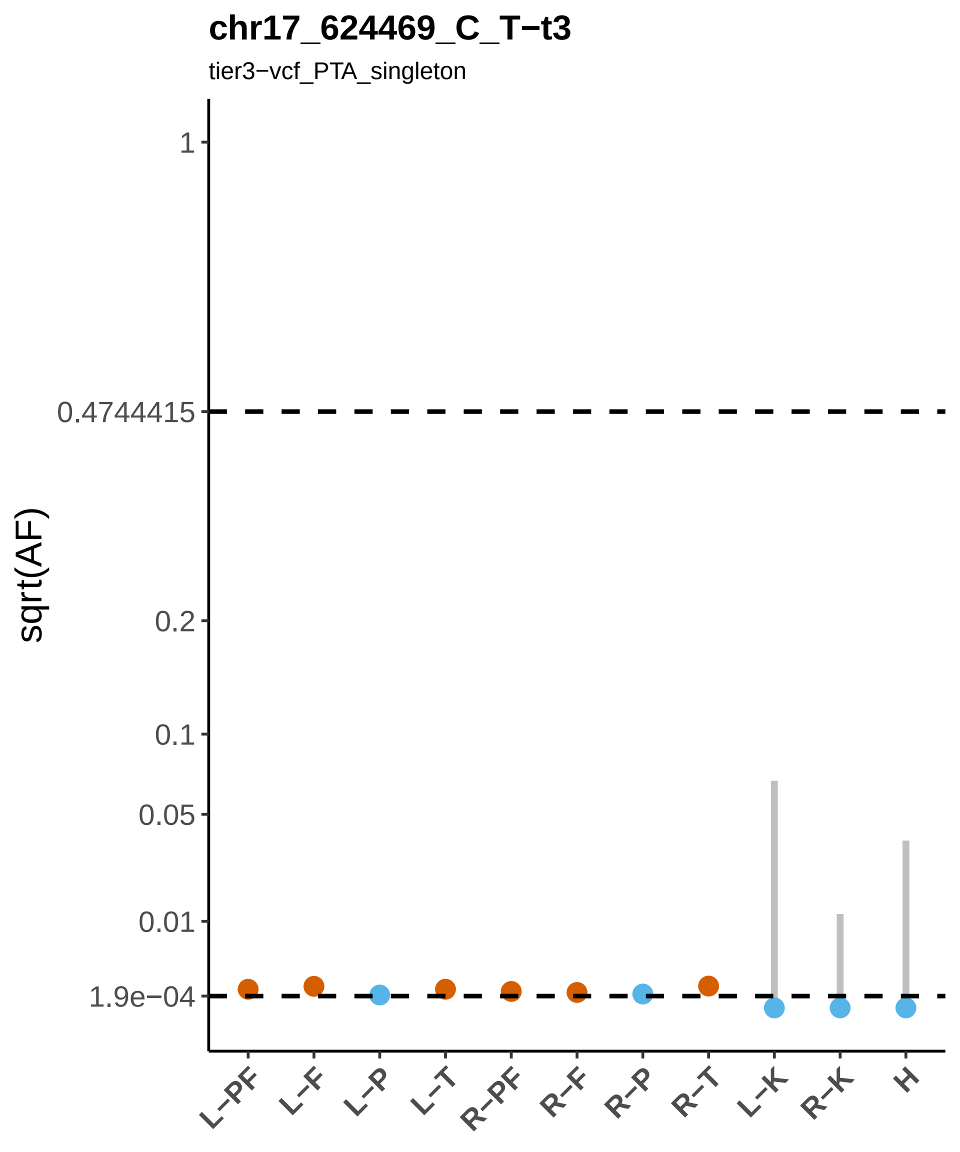
chr17_624469_C_T−t3
tier3−vcf_PTA_singleton
sqrt(AF)
1
0.4744415
0.2
0.1
0.05
0.01
1.9e−04
L−PF
L−F
L−P
L−T
R−PF
R−F
R−P
R−T
L−K
R−K
H

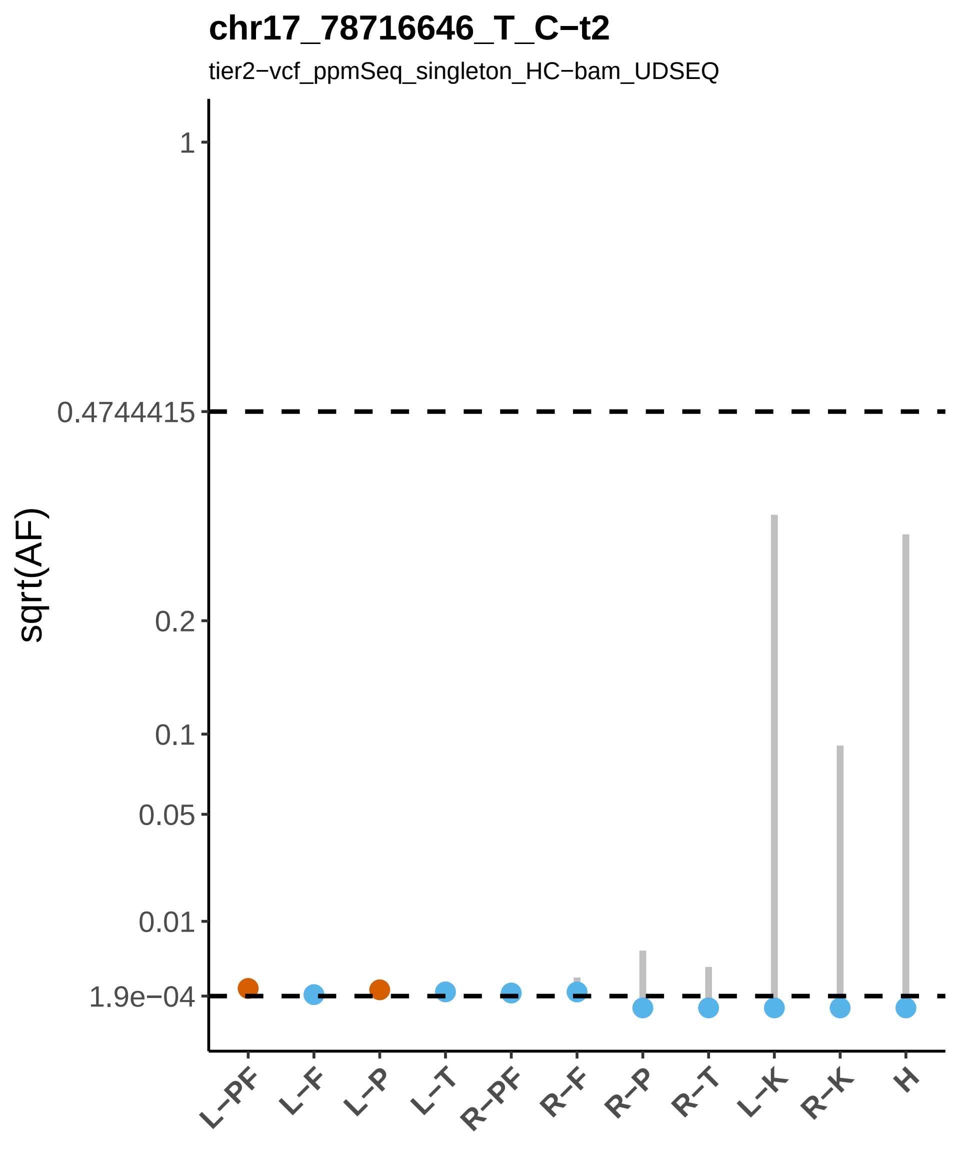
chr17_78716646_T_C−t2
tier2−vcf_ppmSeq_singleton_HC−bam_UDSEQ
sqrt(AF)
1
0.4744415
0.2
0.1
0.05
0.01
1.9e−04
L−PF
L−F
L−P
L−T
R−PF
R−F
R−P
R−T
L−K
R−K
H

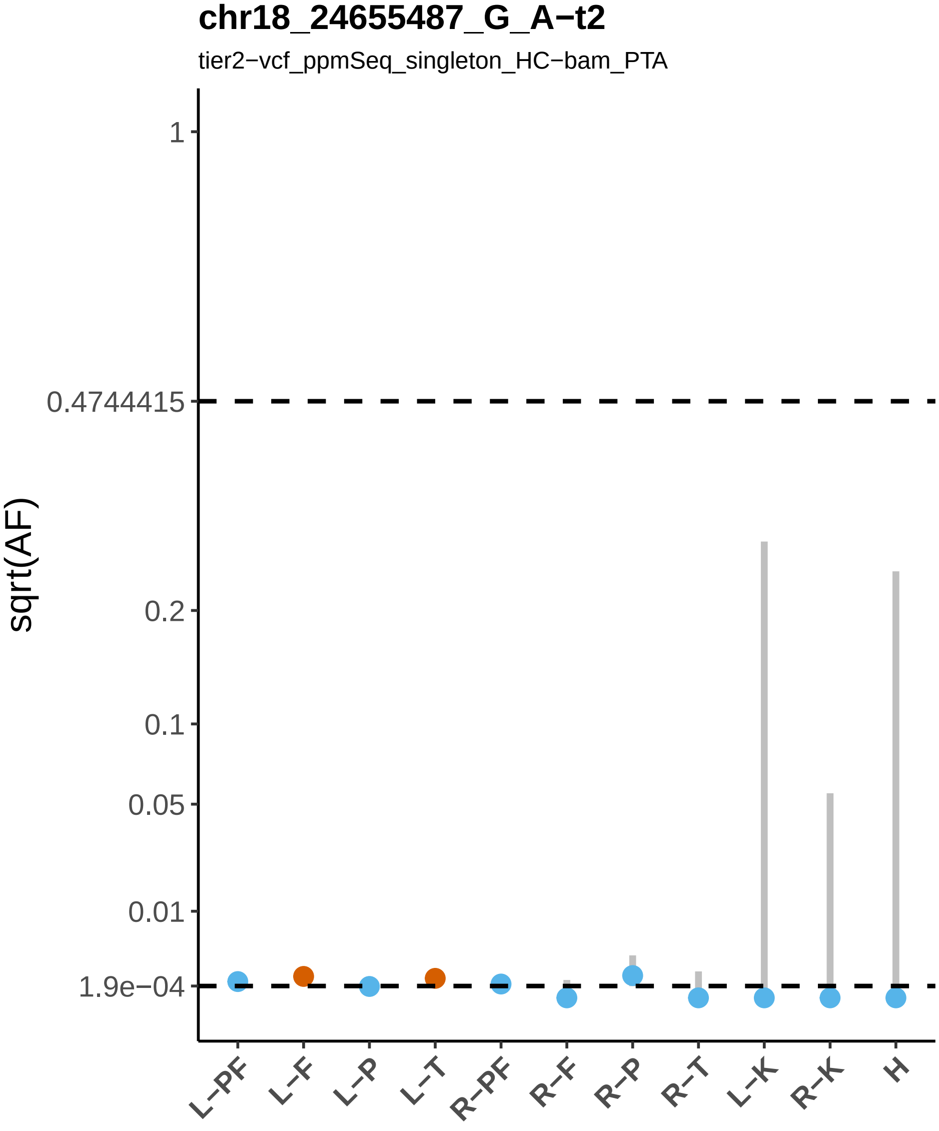

chr18_24655487_G_A−t2
tier2−vcf_ppmSeq_singleton_HC−bam_PTA
sqrt(AF)
1
0.4744415
0.2
0.1
0.05
0.01
1.9e−04
L−PF
L−F
L−P
L−T
R−PF
R−F
R−P
R−T
L−K
R−K
H

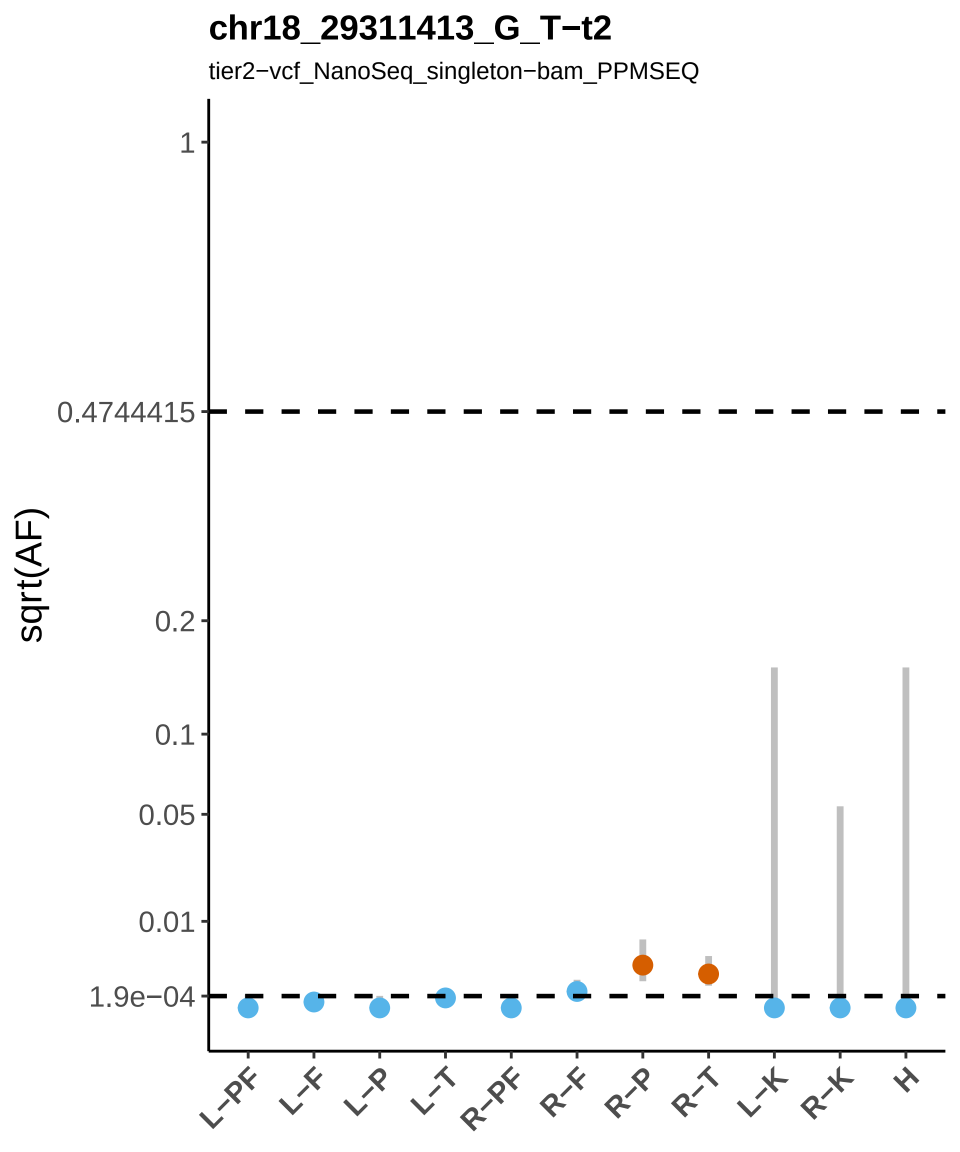

chr18_29311413_G_T−t2
tier2−vcf_NanoSeq_singleton−bam_PPMSEQ
sqrt(AF)
1
0.4744415
0.2
0.1
0.05
0.01
1.9e−04
L−PF
L−F
L−P
L−T
R−PF
R−F
R−P
R−T
L−K
R−K
H

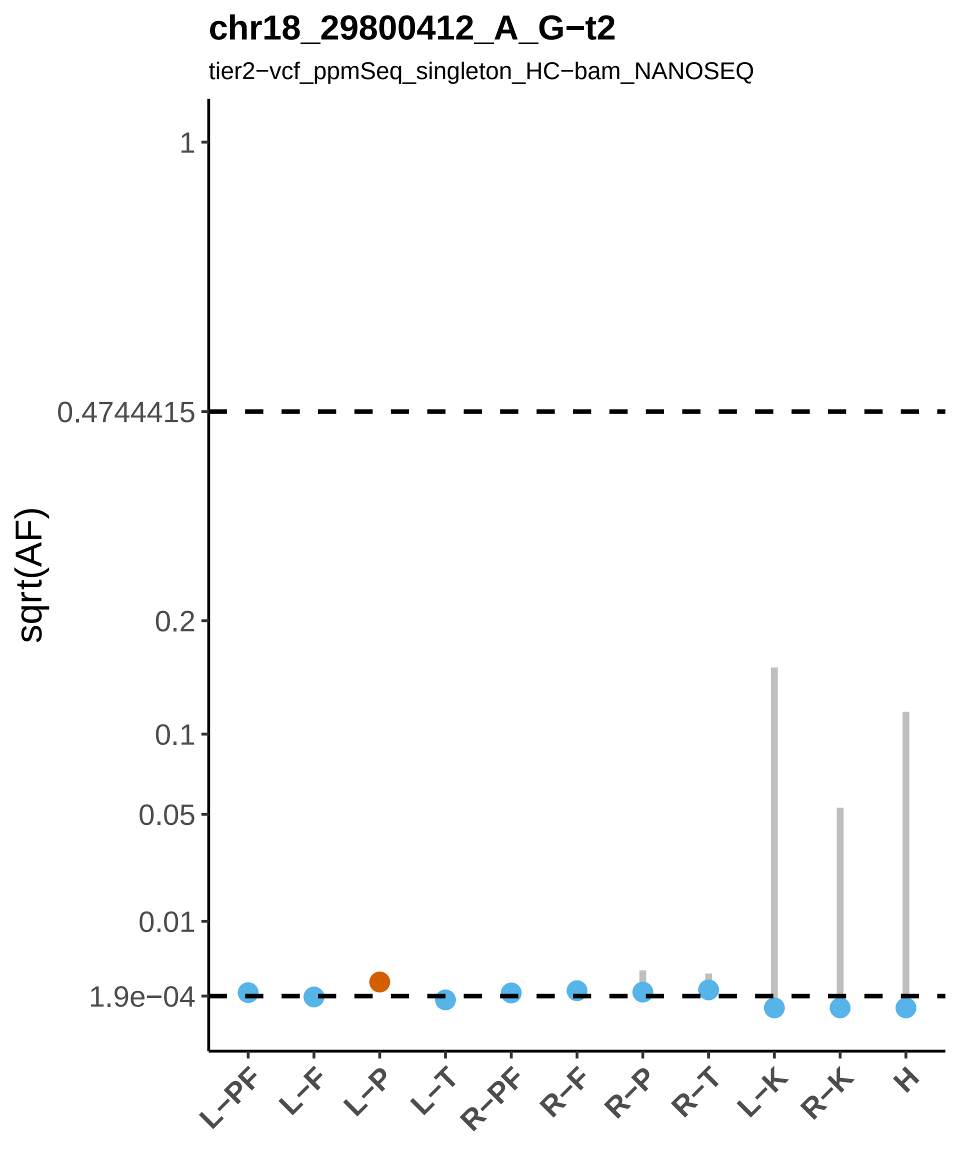

chr18_29800412_A_G−t2
tier2−vcf_ppmSeq_singleton_HC−bam_NANOSEQ
sqrt(AF)
1
0.4744415
0.2
0.1
0.05
0.01
1.9e−04
L−PF
L−F
L−P
L−T
R−PF
R−F
R−P
R−T
L−K
R−K
H

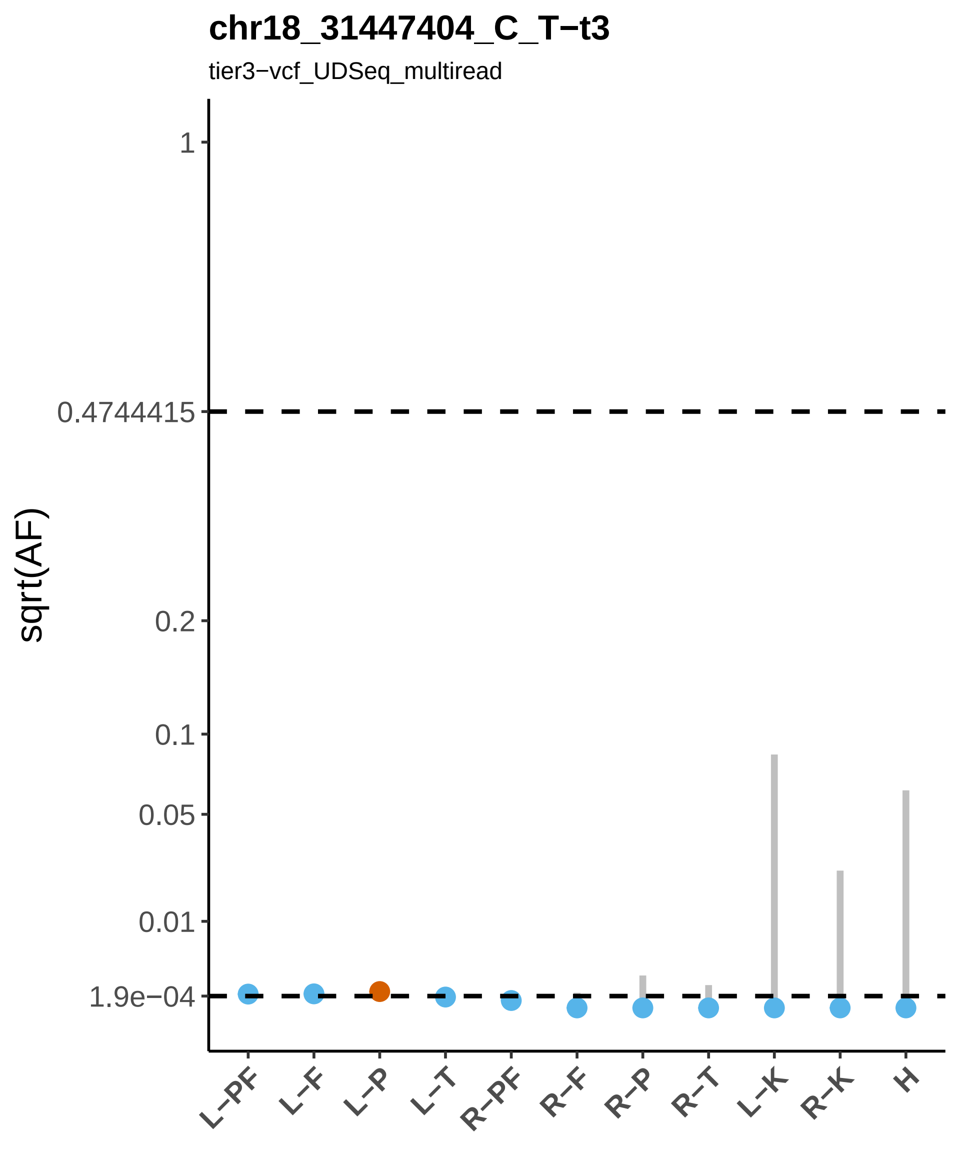

chr18_31447404_C_T−t3
tier3−vcf_UDSeq_multiread
sqrt(AF)
1
0.4744415
0.2
0.1
0.05
0.01
1.9e−04
L−PF
L−F
L−P
L−T
R−PF
R−F
R−P
R−T
L−K
R−K
H

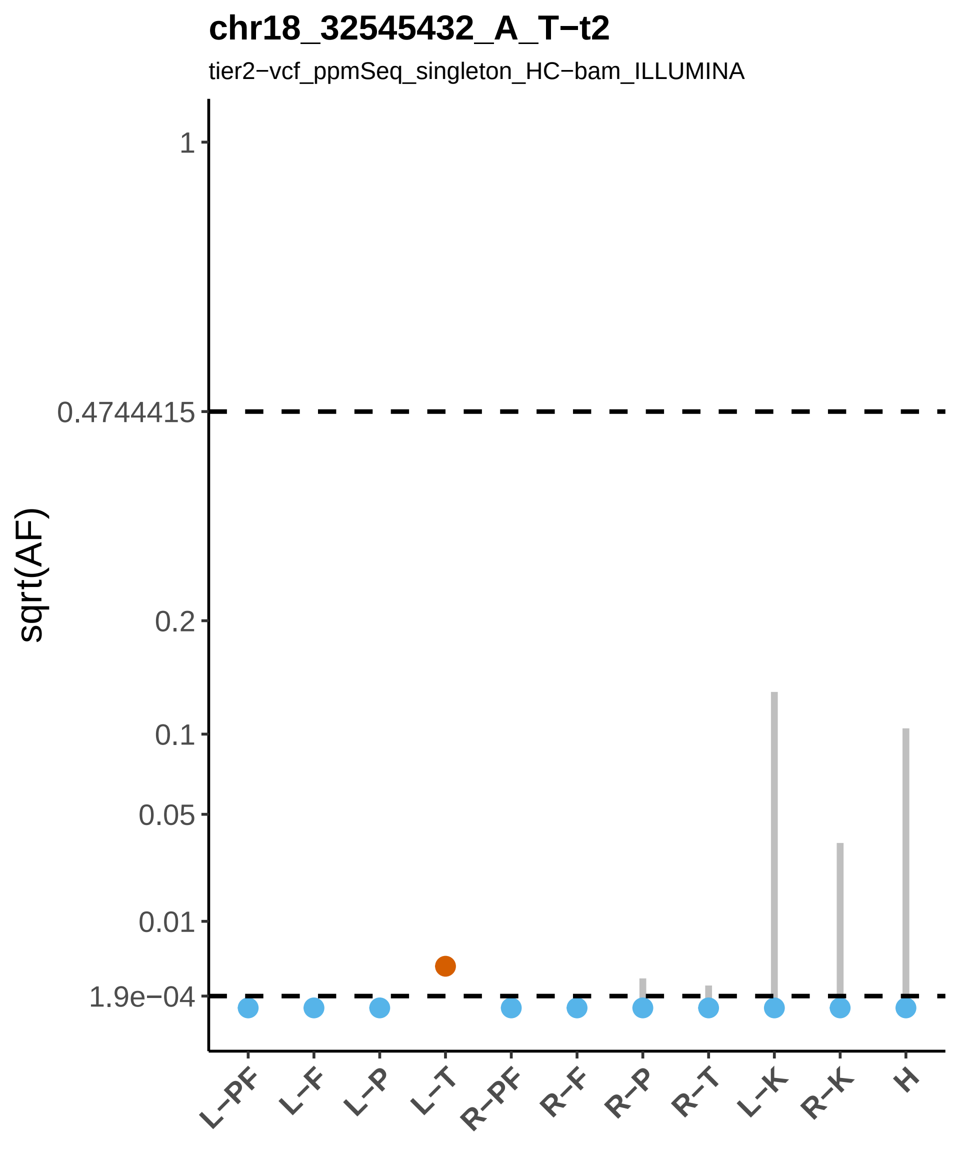
chr18_32545432_A_T−t2
tier2−vcf_ppmSeq_singleton_HC−bam_ILLUMINA
sqrt(AF)
1
0.4744415
0.2
0.1
0.05
0.01
1.9e−04
L−PF
L−F
L−P
L−T
R−PF
R−F
R−P
R−T
L−K
R−K
H

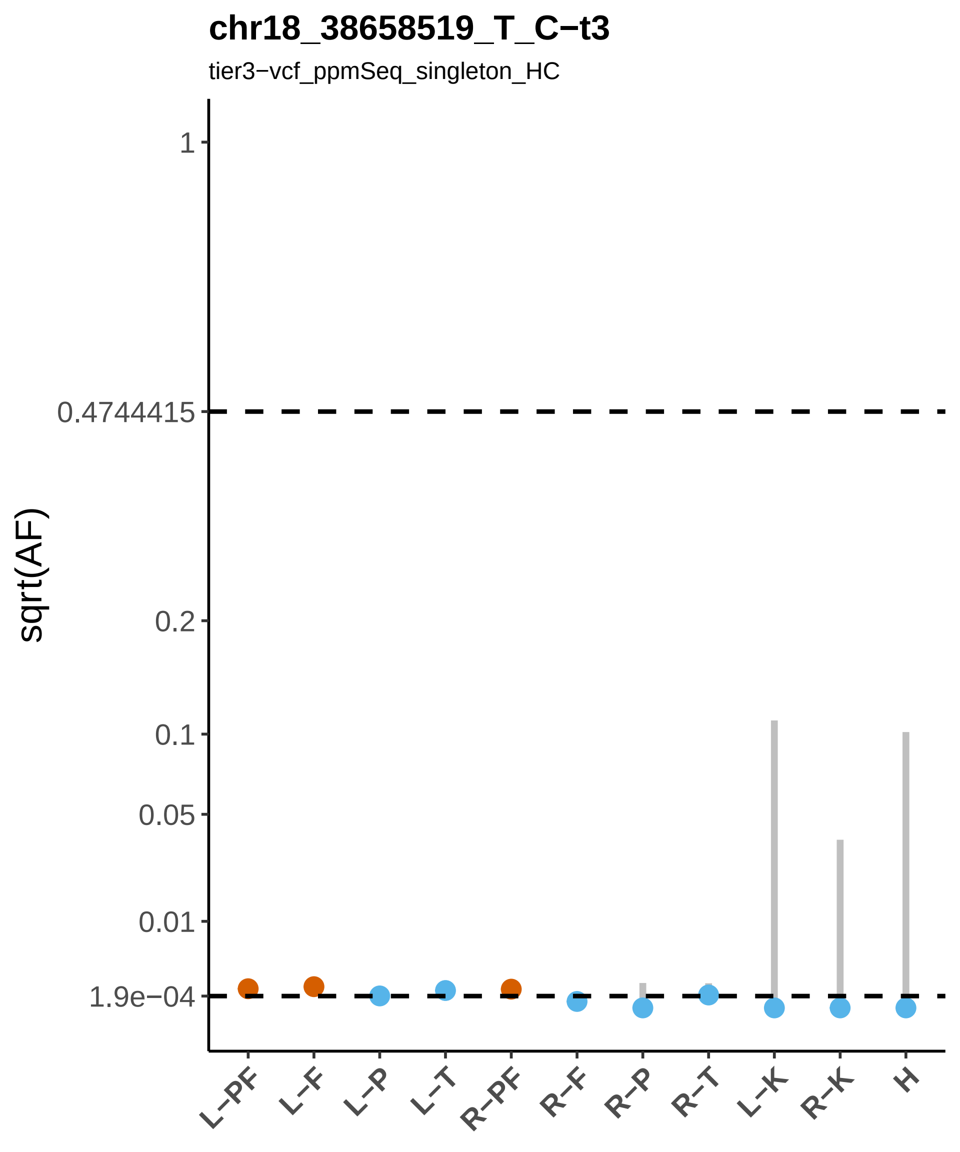

chr18_38658519_T_C−t3
tier3−vcf_ppmSeq_singleton_HC
sqrt(AF)
1
0.4744415
0.2
0.1
0.05
0.01
1.9e−04
L−PF
L−F
L−P
L−T
R−PF
R−F
R−P
R−T
L−K
R−K
H

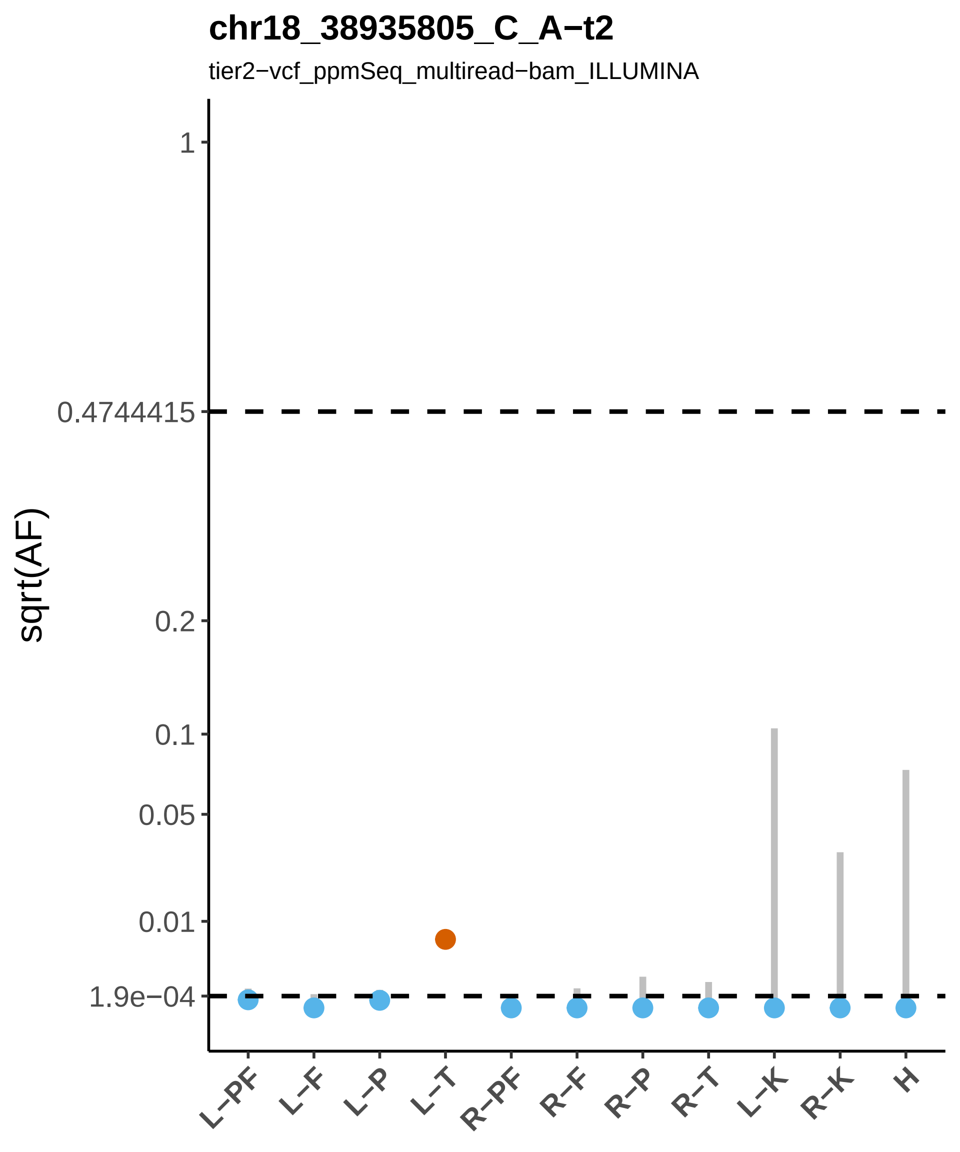

chr18_38935805_C_A−t2
tier2−vcf_ppmSeq_multiread−bam_ILLUMINA
sqrt(AF)
1
0.4744415
0.2
0.1
0.05
0.01
1.9e−04
L−PF
L−F
L−P
L−T
R−PF
R−F
R−P
R−T
L−K
R−K
H

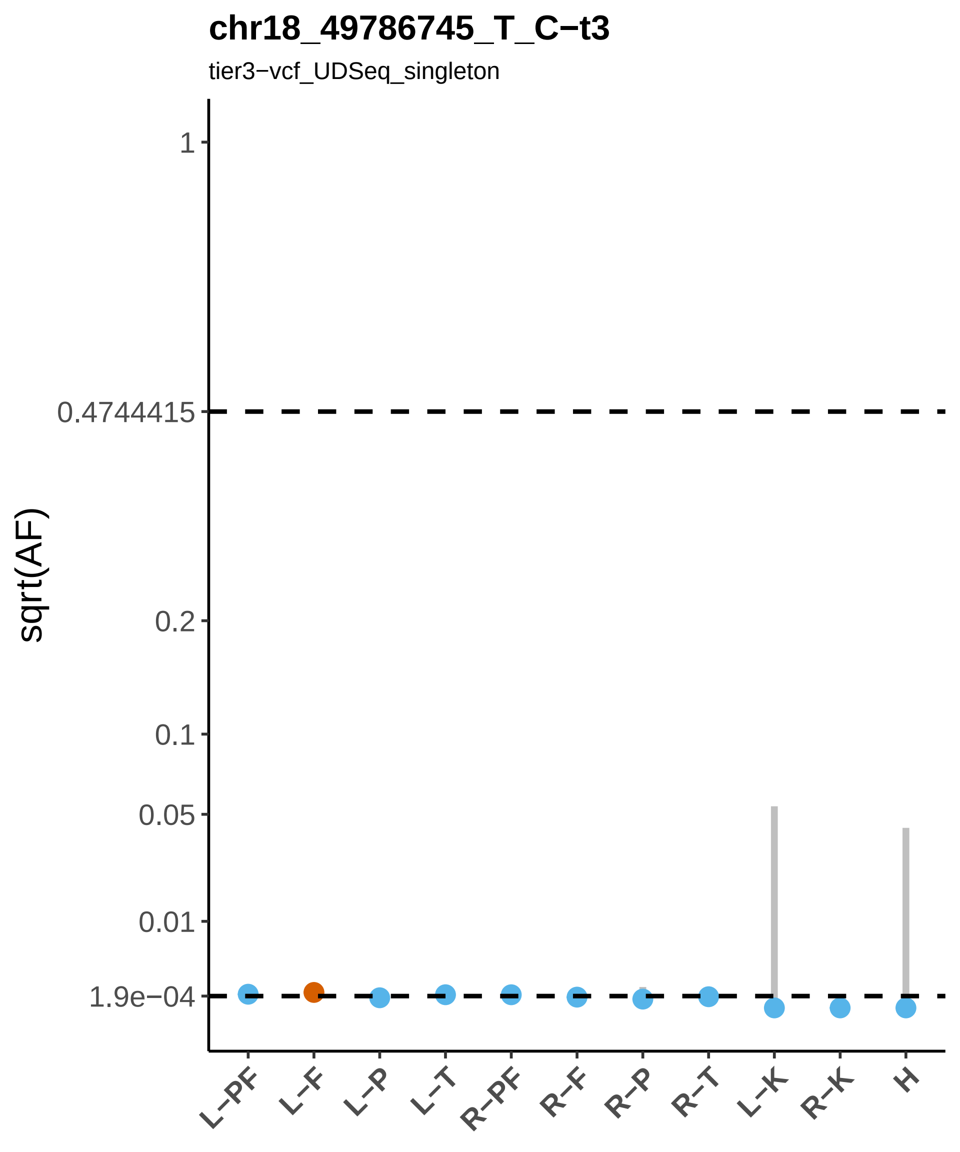

chr18_49786745_T_C−t3
tier3−vcf_UDSeq_singleton
sqrt(AF)
1
0.4744415
0.2
0.1
0.05
0.01
1.9e−04
L−PF
L−F
L−P
L−T
R−PF
R−F
R−P
R−T
L−K
R−K
H

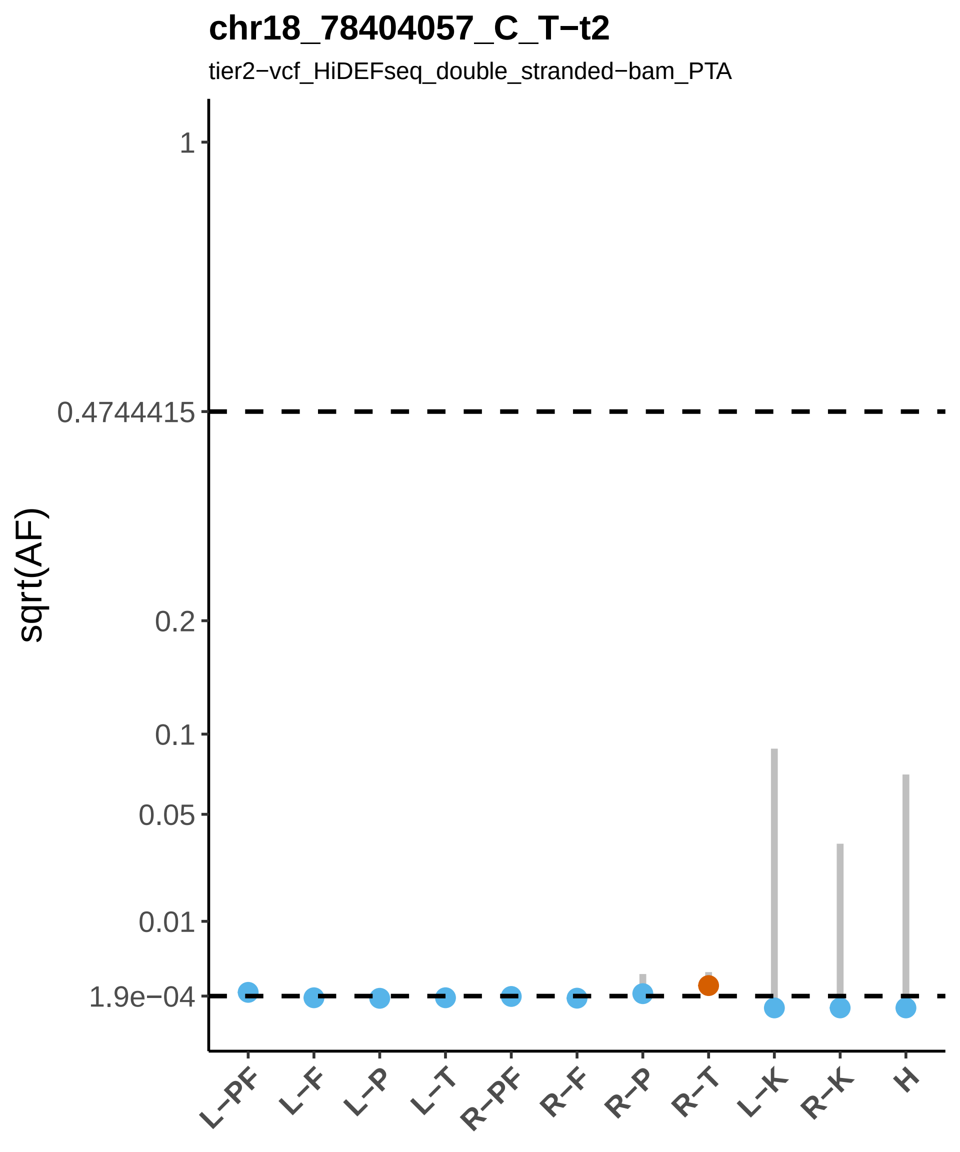

chr18_78404057_C_T−t2
tier2−vcf_HiDEFseq_double_stranded−bam_PTA
sqrt(AF)
1
0.4744415
0.2
0.1
0.05
0.01
1.9e−04
L−PF
L−F
L−P
L−T
R−PF
R−F
R−P
R−T
L−K
R−K
H

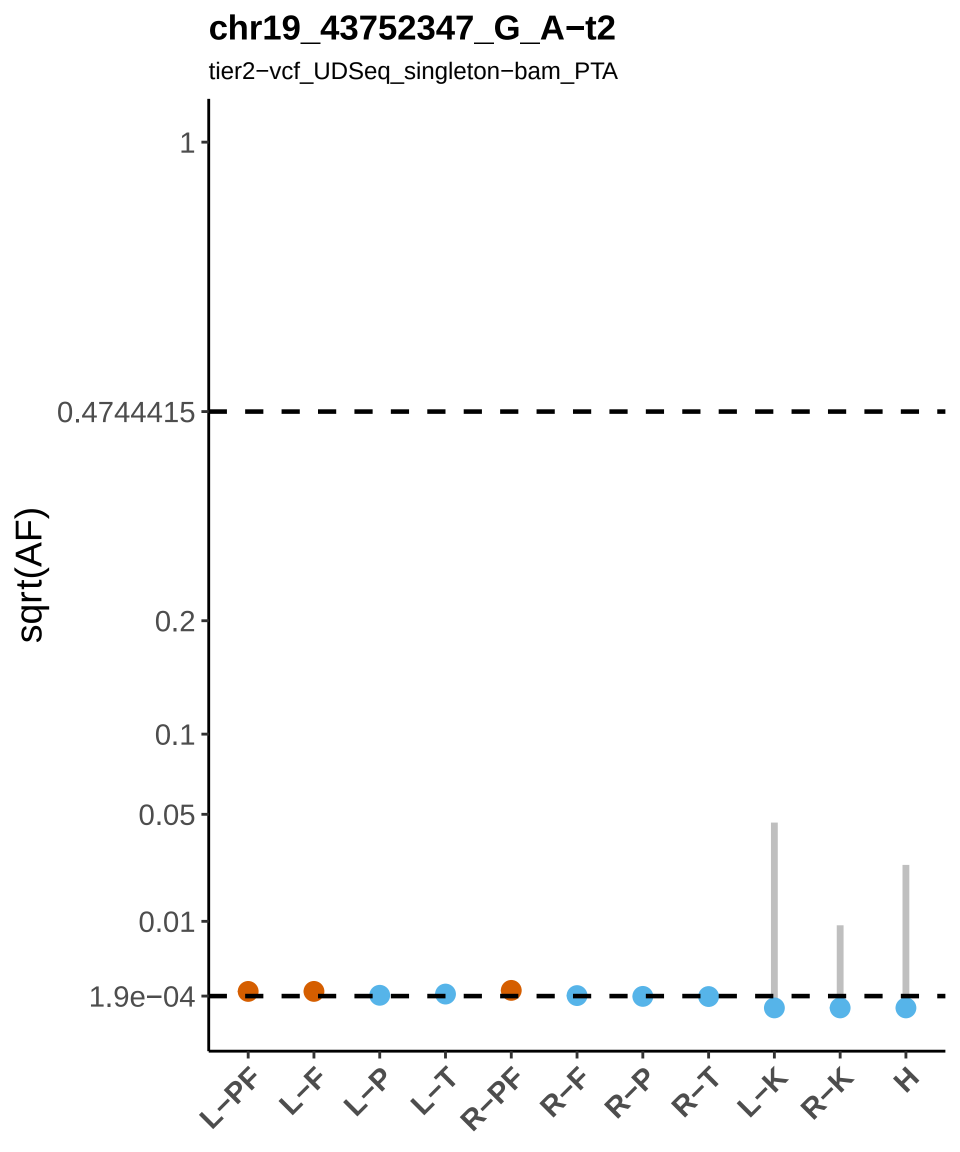

chr19_43752347_G_A−t2
tier2−vcf_UDSeq_singleton−bam_PTA
sqrt(AF)
1
0.4744415
0.2
0.1
0.05
0.01
1.9e−04
L−PF
L−F
L−P
L−T
R−PF
R−F
R−P
R−T
L−K
R−K
H

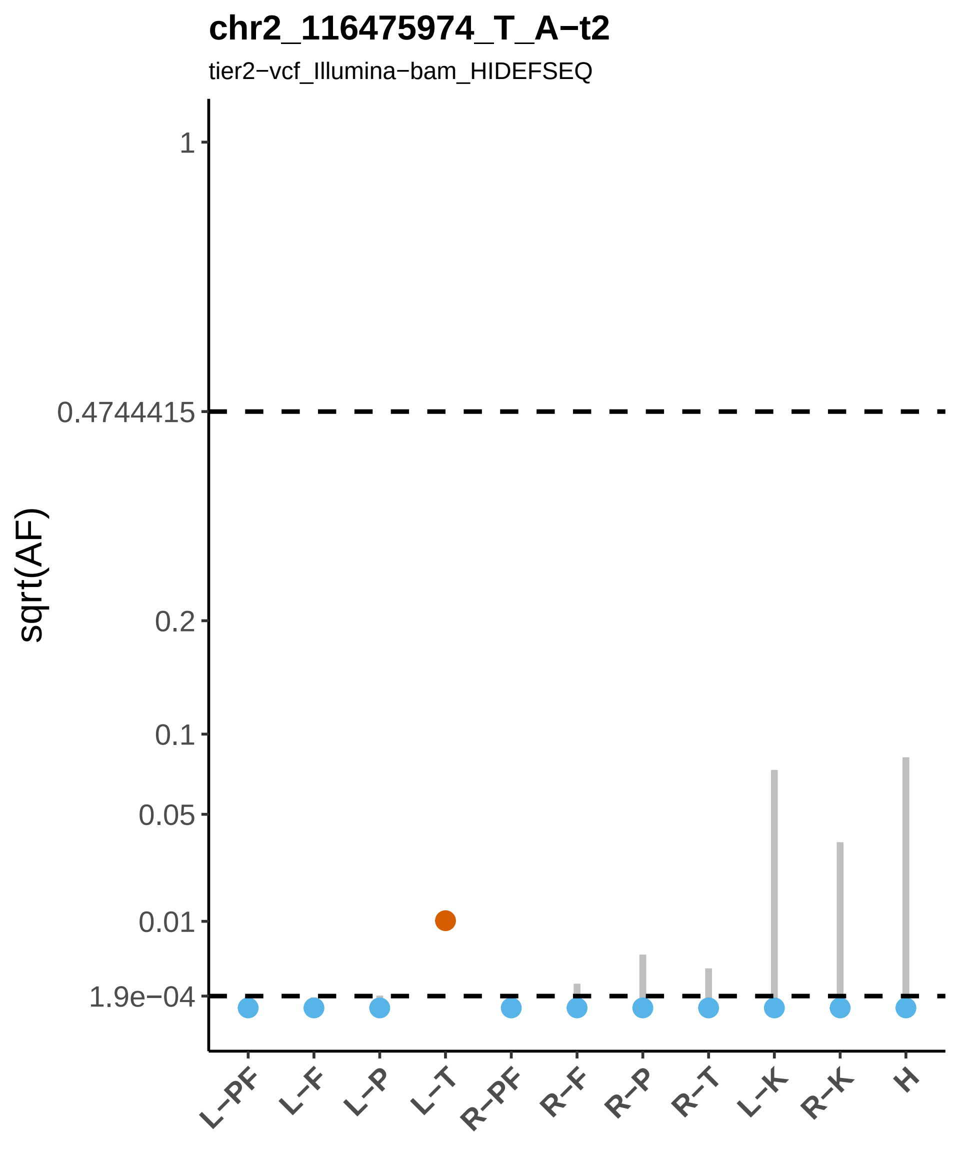
chr2_116475974_T_A−t2
tier2−vcf_Illumina−bam_HIDEFSEQ
sqrt(AF)
1
0.4744415
0.2
0.1
0.05
0.01
1.9e−04
L−PF
L−F
L−P
L−T
R−PF
R−F
R−P
R−T
L−K
R−K
H

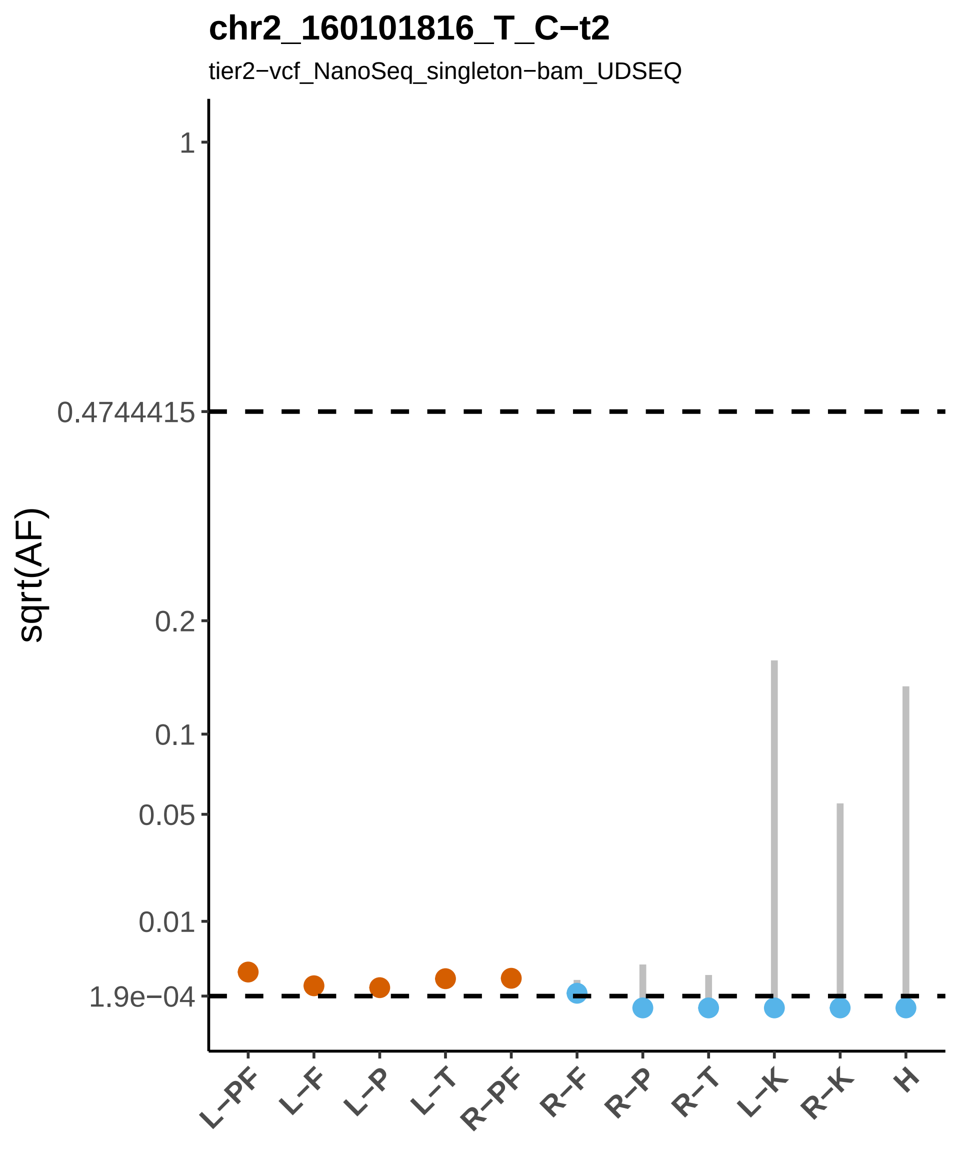

chr2_160101816_T_C−t2
tier2−vcf_NanoSeq_singleton−bam_UDSEQ
sqrt(AF)
1
0.4744415
0.2
0.1
0.05
0.01
1.9e−04
L−PF
L−F
L−P
L−T
R−PF
R−F
R−P
R−T
L−K
R−K
H

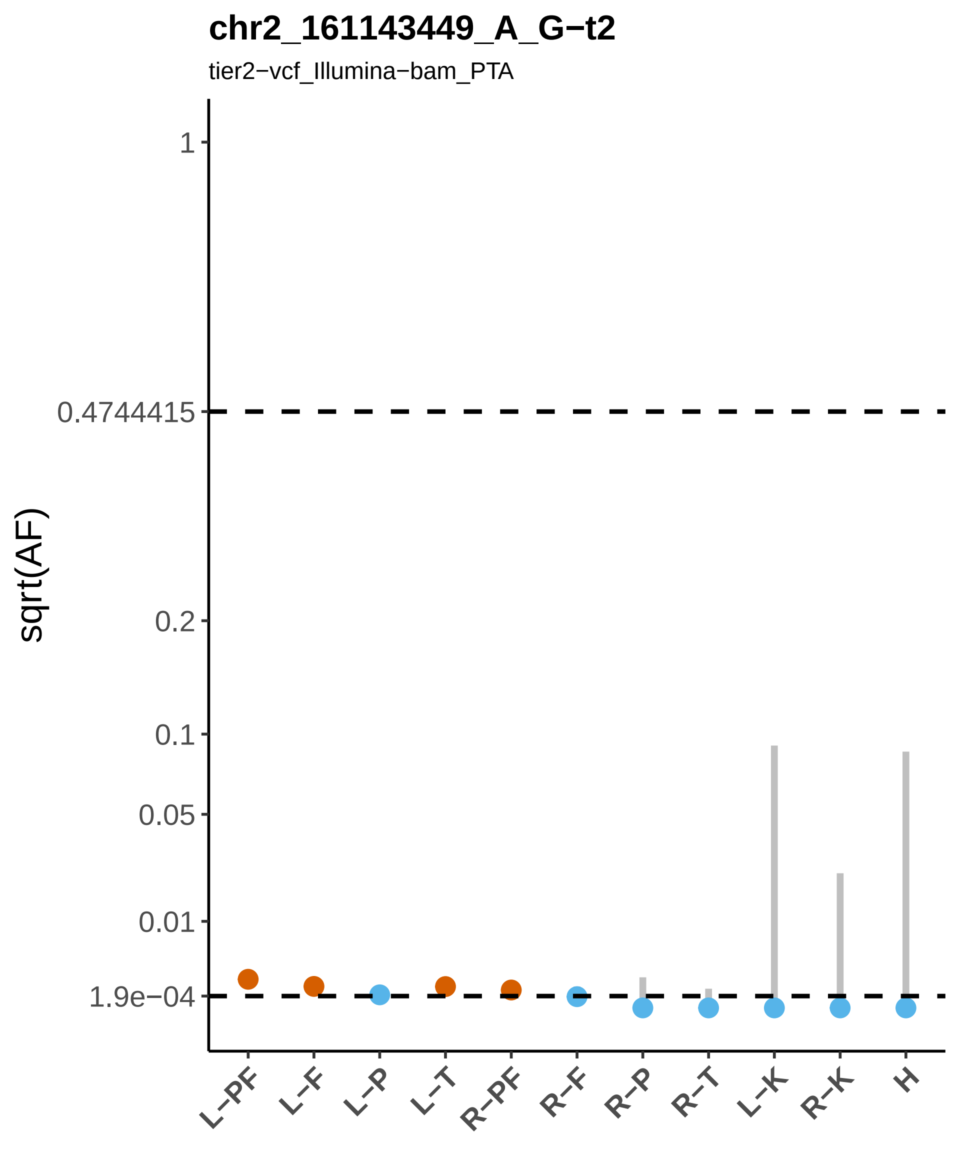
chr2_161143449_A_G−t2
tier2−vcf_Illumina−bam_PTA
sqrt(AF)
1
0.4744415
0.2
0.1
0.05
0.01
1.9e−04
L−PF
L−F
L−P
L−T
R−PF
R−F
R−P
R−T
L−K
R−K
H

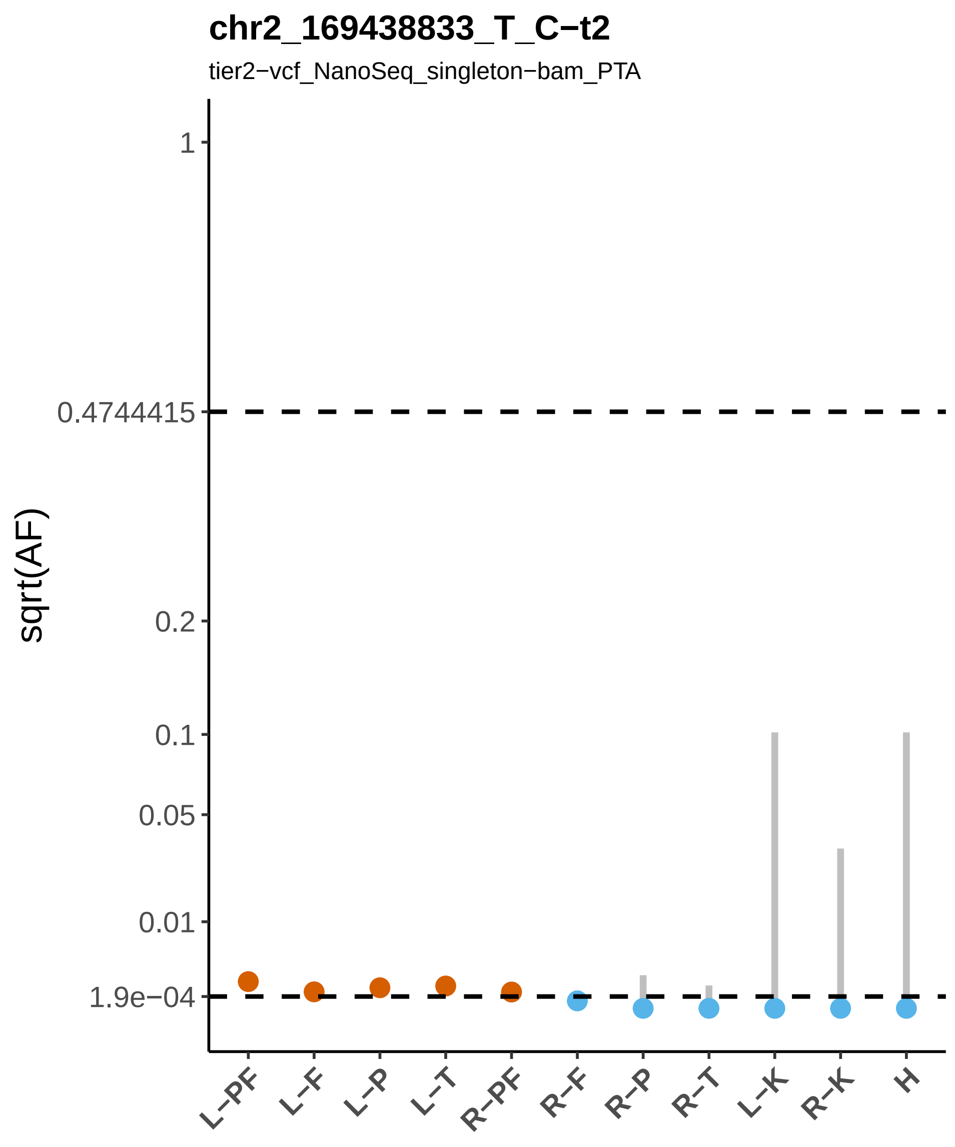

chr2_169438833_T_C−t2
tier2−vcf_NanoSeq_singleton−bam_PTA
sqrt(AF)
1
0.4744415
0.2
0.1
0.05
0.01
1.9e−04
L−PF
L−F
L−P
L−T
R−PF
R−F
R−P
R−T
L−K
R−K
H

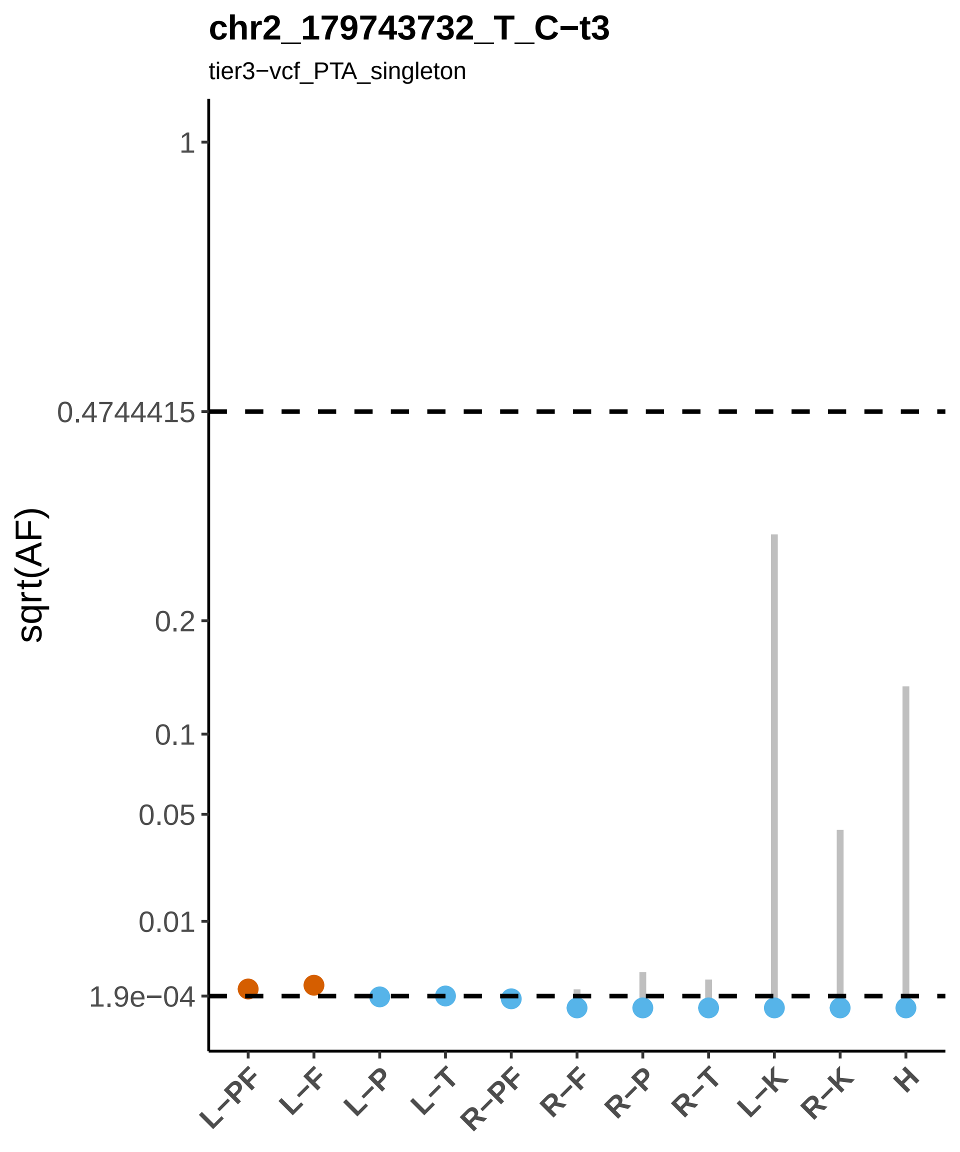
chr2_179743732_T_C−t3
tier3−vcf_PTA_singleton
sqrt(AF)
1
0.4744415
0.2
0.1
0.05
0.01
1.9e−04
L−PF
L−F
L−P
L−T
R−PF
R−F
R−P
R−T
L−K
R−K
H

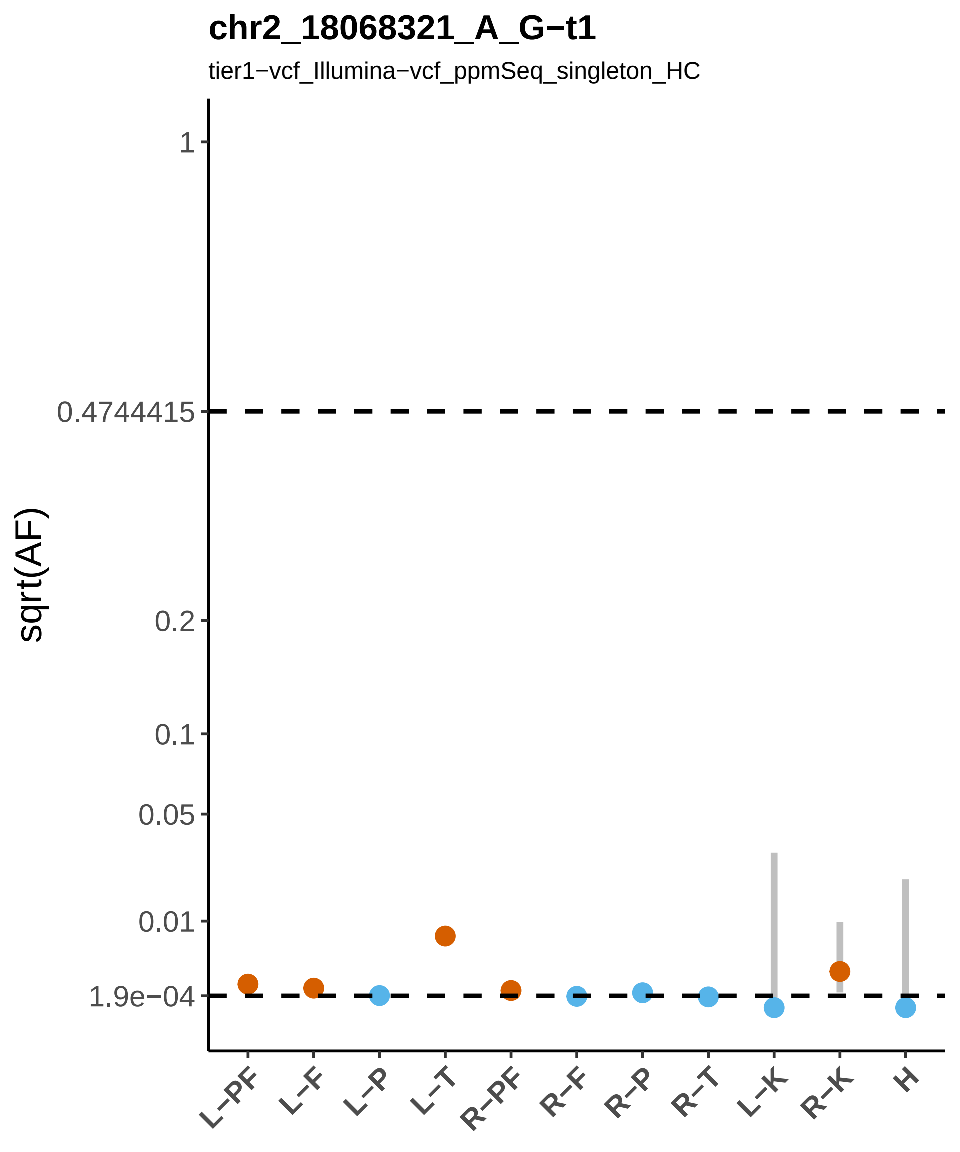

chr2_18068321_A_G−t1
tier1−vcf_Illumina−vcf_ppmSeq_singleton_HC
sqrt(AF)
1
0.4744415
0.2
0.1
0.05
0.01
1.9e−04
L−PF
L−F
L−P
L−T
R−PF
R−F
R−P
R−T
L−K
R−K
H

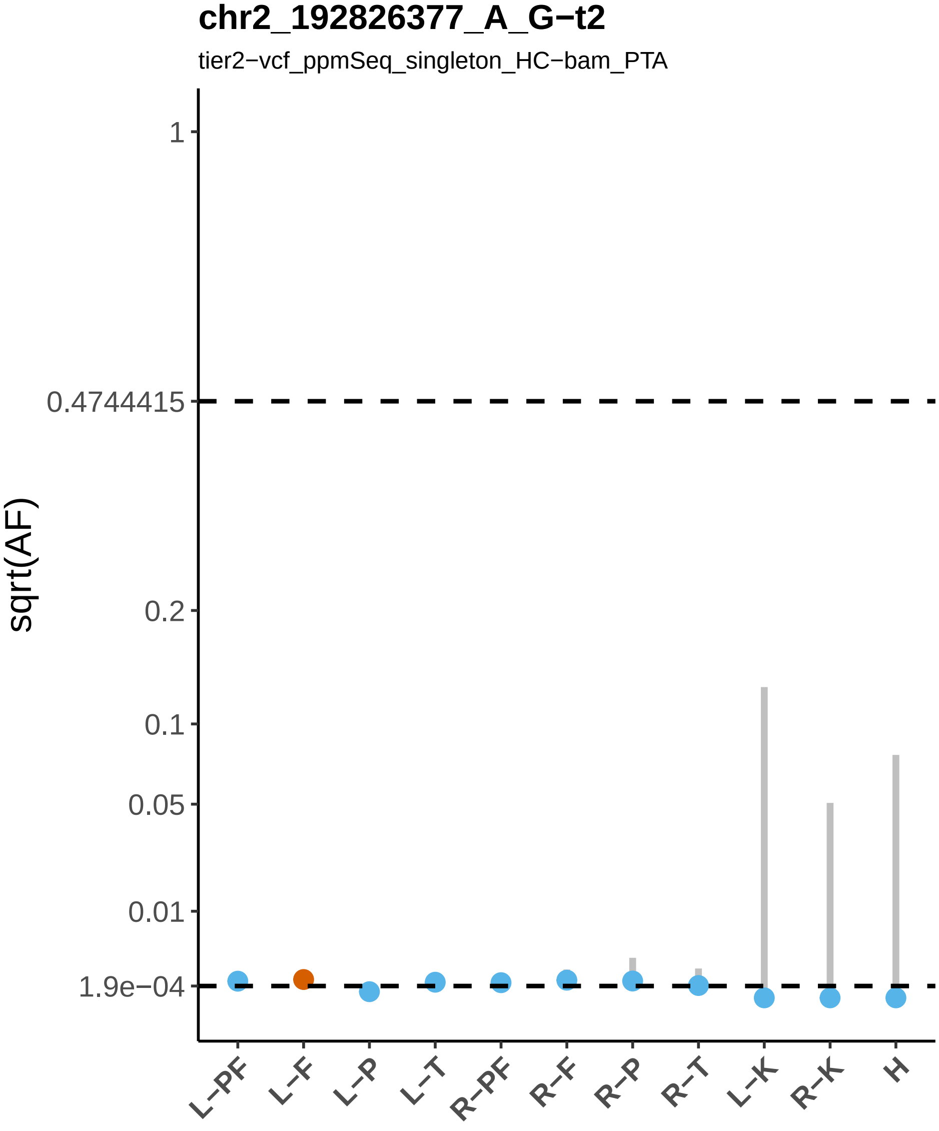
chr2_192826377_A_G−t2
tier2−vcf_ppmSeq_singleton_HC−bam_PTA
sqrt(AF)
1
0.4744415
0.2
0.1
0.05
0.01
1.9e−04
L−PF
L−F
L−P
L−T
R−PF
R−F
R−P
R−T
L−K
R−K
H

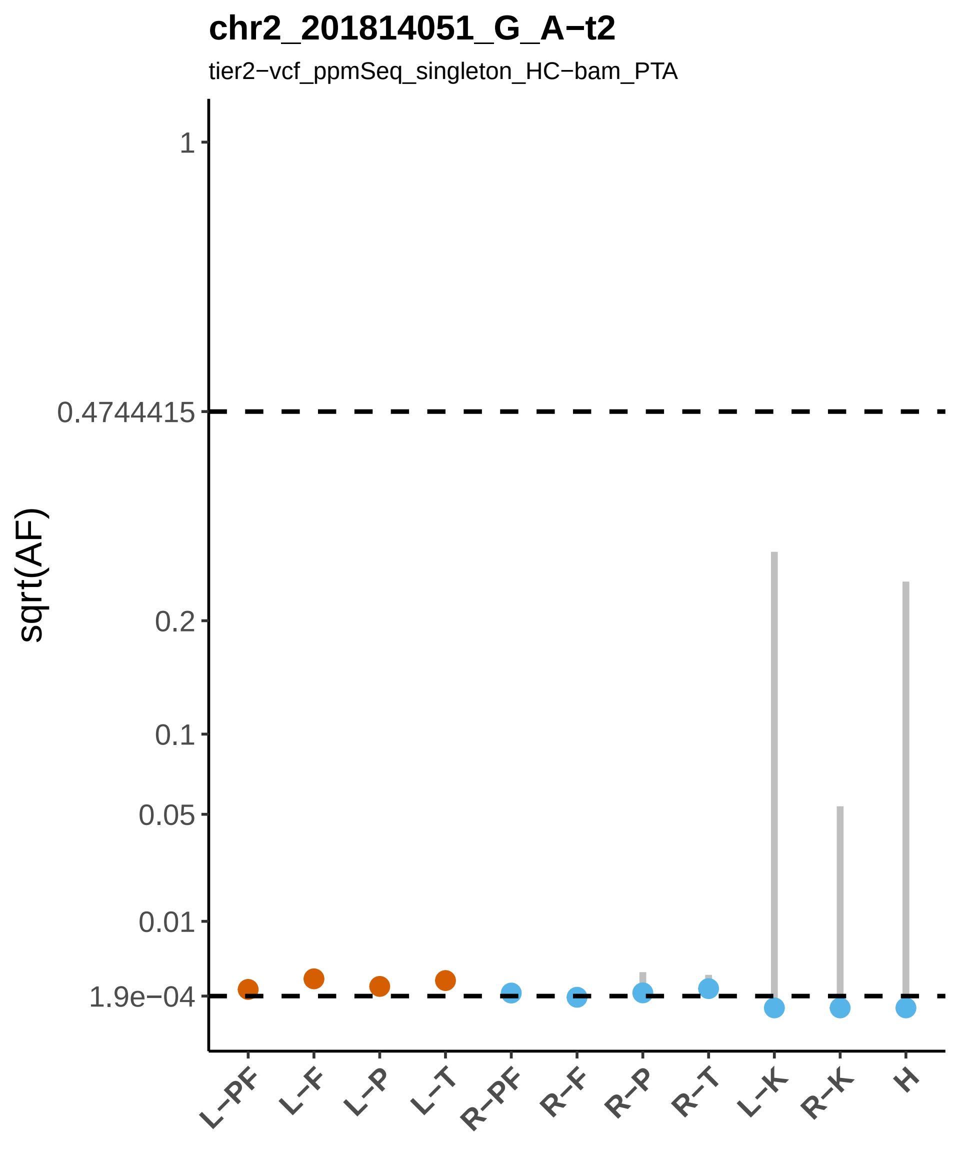
chr2_201814051_G_A−t2
tier2−vcf_ppmSeq_singleton_HC−bam_PTA
sqrt(AF)
1
0.4744415
0.2
0.1
0.05
0.01
1.9e−04
L−PF
L−F
L−P
L−T
R−PF
R−F
R−P
R−T
L−K
R−K
H

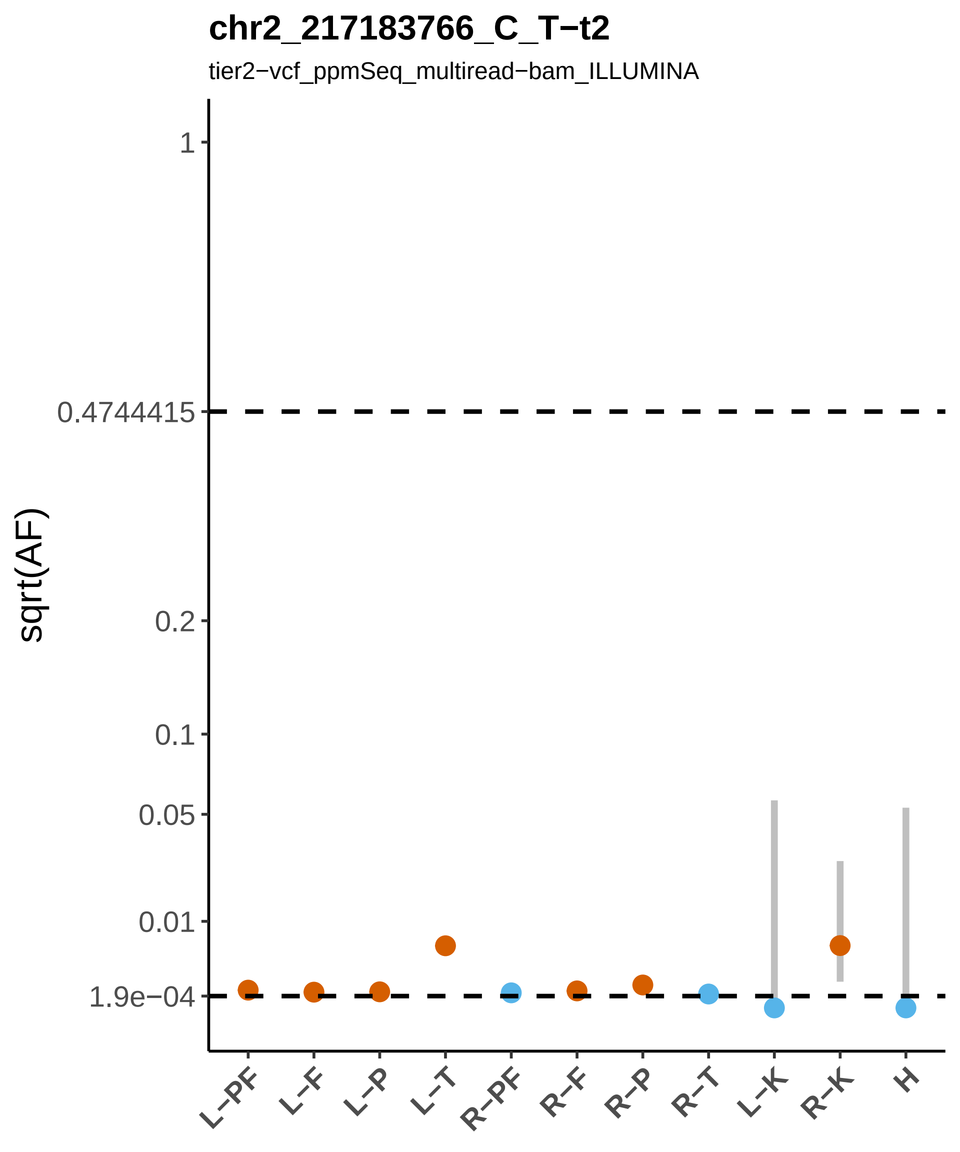
chr2_217183766_C_T−t2
tier2−vcf_ppmSeq_multiread−bam_ILLUMINA
sqrt(AF)
1
0.4744415
0.2
0.1
0.05
0.01
1.9e−04
L−PF
L−F
L−P
L−T
R−PF
R−F
R−P
R−T
L−K
R−K
H

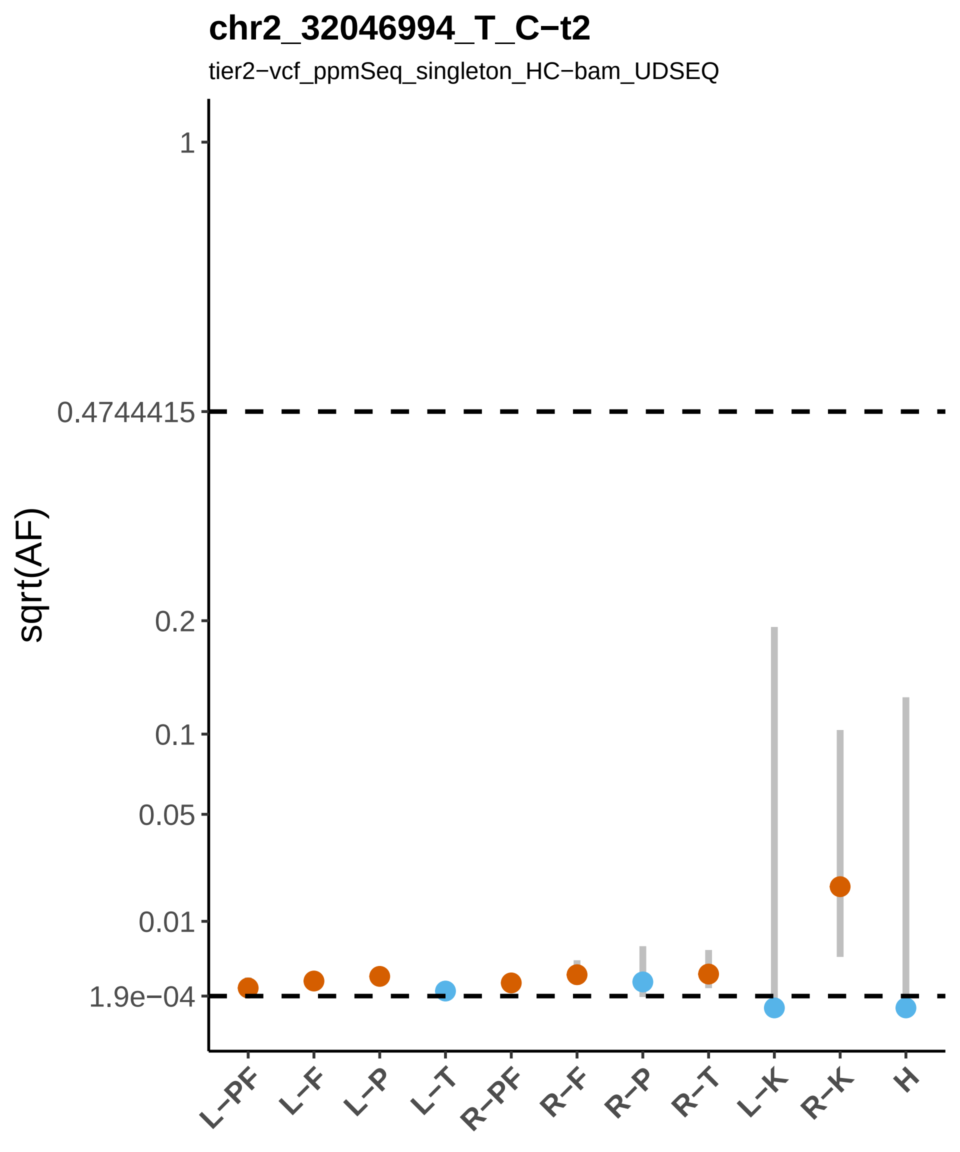
chr2_32046994_T_C−t2
tier2−vcf_ppmSeq_singleton_HC−bam_UDSEQ
sqrt(AF)
1
0.4744415
0.2
0.1
0.05
0.01
1.9e−04
L−PF
L−F
L−P
L−T
R−PF
R−F
R−P
R−T
L−K
R−K
H

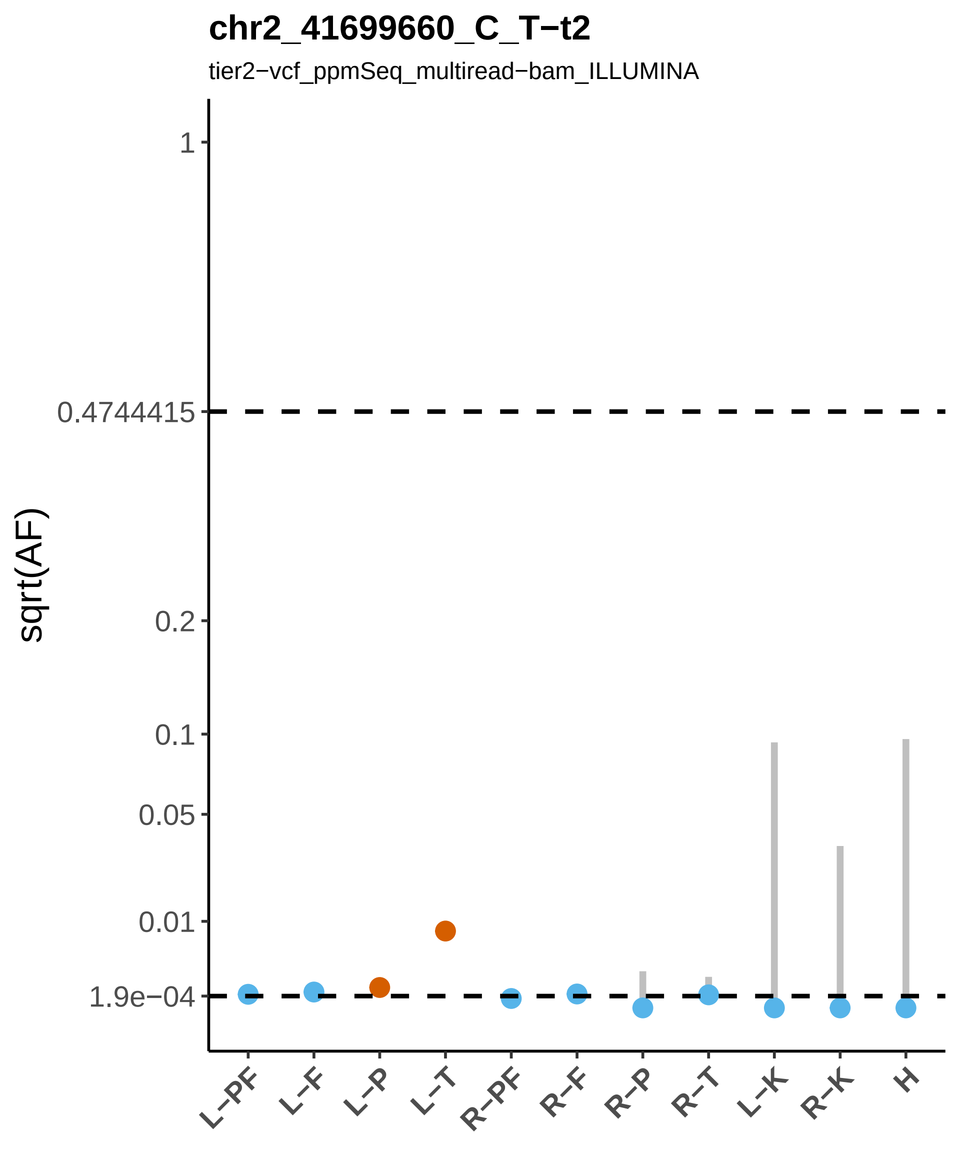
chr2_41699660_C_T−t2
tier2−vcf_ppmSeq_multiread−bam_ILLUMINA
sqrt(AF)
1
0.4744415
0.2
0.1
0.05
0.01
1.9e−04
L−PF
L−F
L−P
L−T
R−PF
R−F
R−P
R−T
L−K
R−K
H

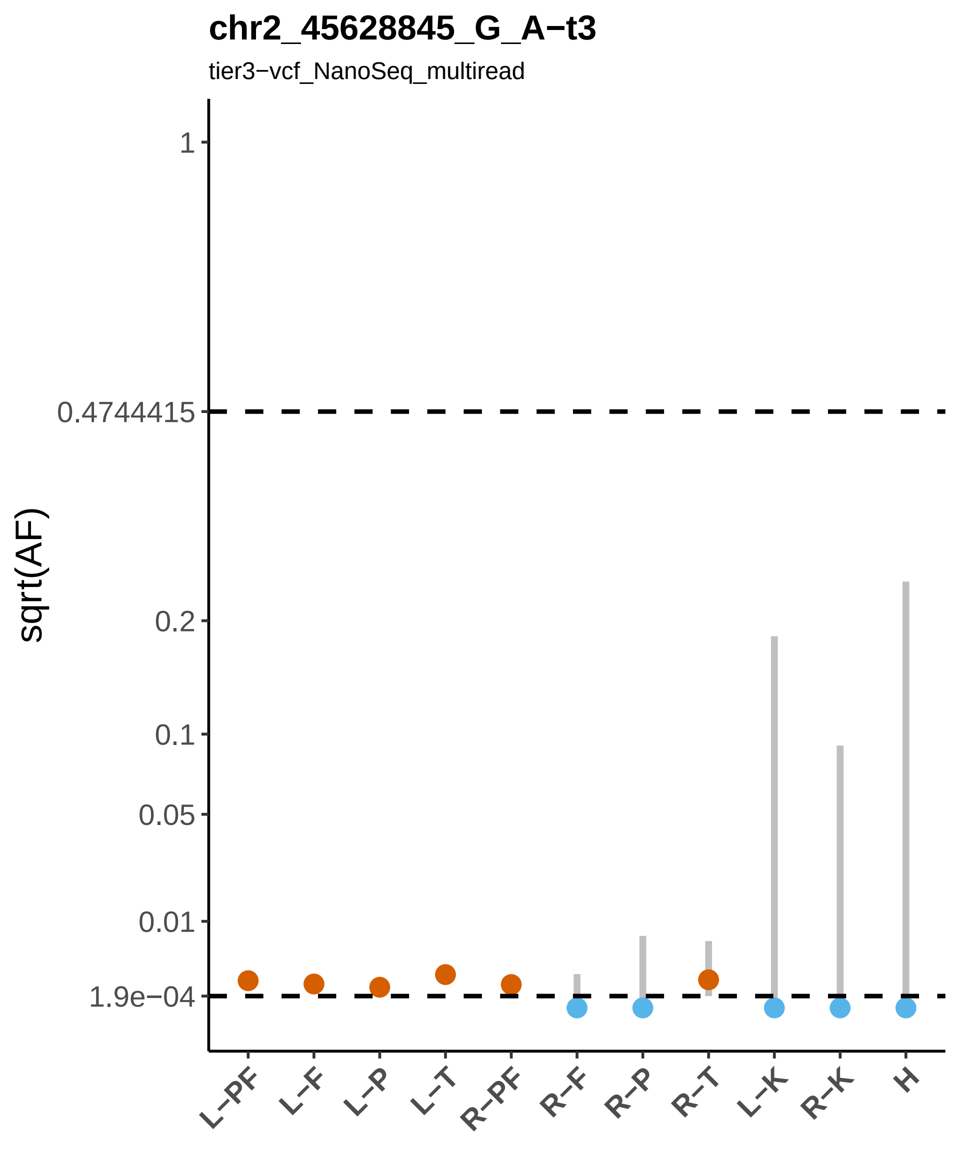
chr2_45628845_G_A−t3
tier3−vcf_NanoSeq_multiread
sqrt(AF)
1
0.4744415
0.2
0.1
0.05
0.01
1.9e−04
L−PF
L−F
L−P
L−T
R−PF
R−F
R−P
R−T
L−K
R−K
H

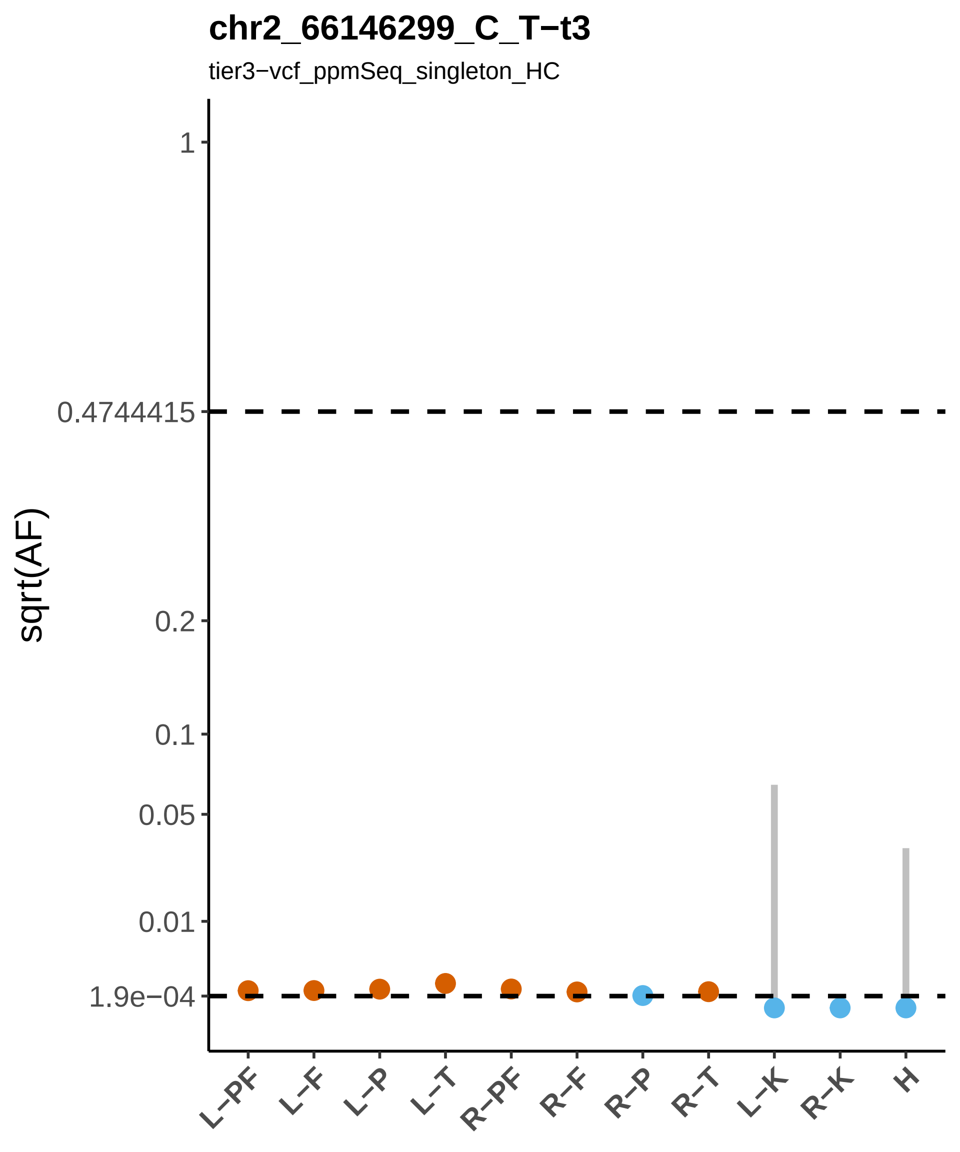

chr2_66146299_C_T−t3
tier3−vcf_ppmSeq_singleton_HC
sqrt(AF)
1
0.4744415
0.2
0.1
0.05
0.01
1.9e−04
L−PF
L−F
L−P
L−T
R−PF
R−F
R−P
R−T
L−K
R−K
H

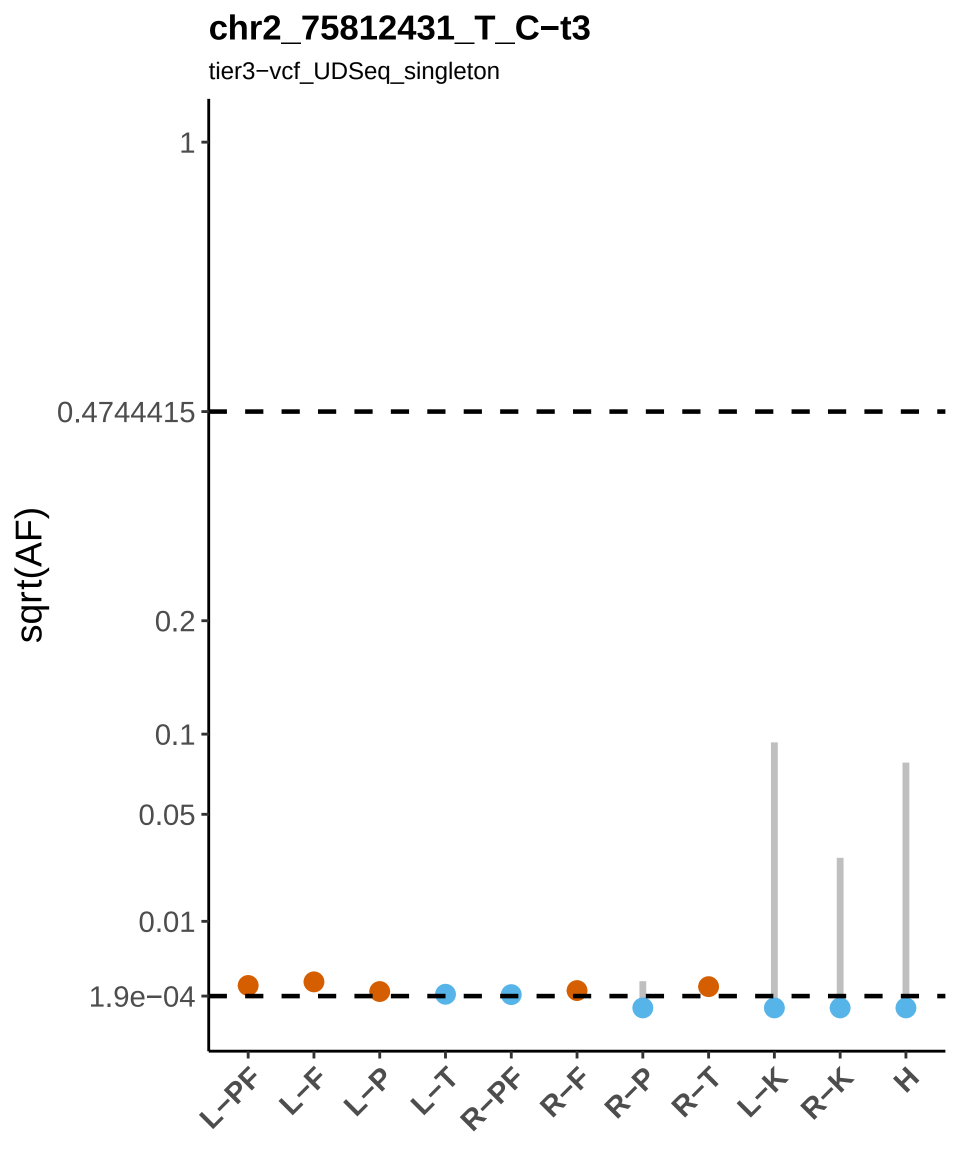
chr2_75812431_T_C−t3
tier3−vcf_UDSeq_singleton
sqrt(AF)
1
0.4744415
0.2
0.1
0.05
0.01
1.9e−04
L−PF
L−F
L−P
L−T
R−PF
R−F
R−P
R−T
L−K
R−K
H

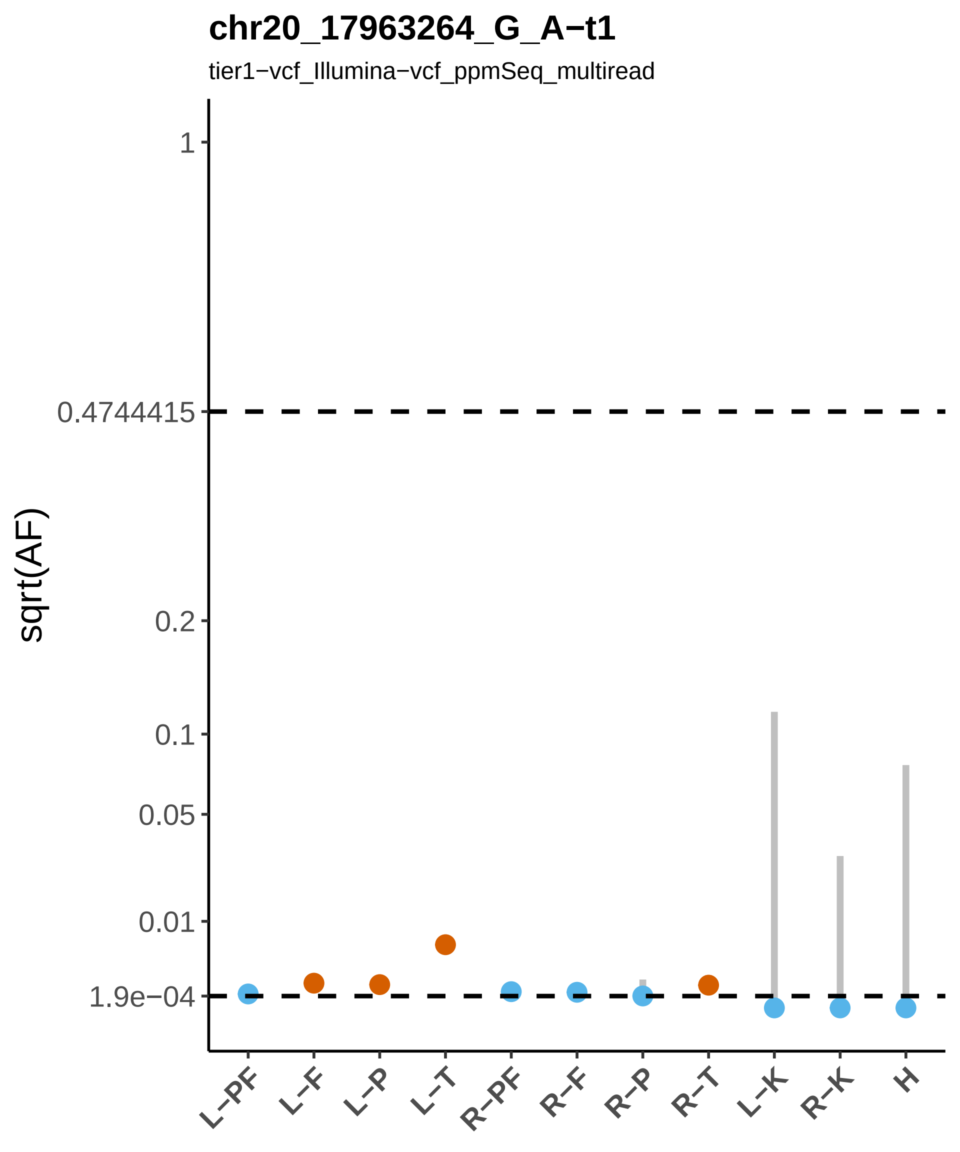
chr20_17963264_G_A−t1
tier1−vcf_Illumina−vcf_ppmSeq_multiread
sqrt(AF)
1
0.4744415
0.2
0.1
0.05
0.01
1.9e−04
L−PF
L−F
L−P
L−T
R−PF
R−F
R−P
R−T
L−K
R−K
H

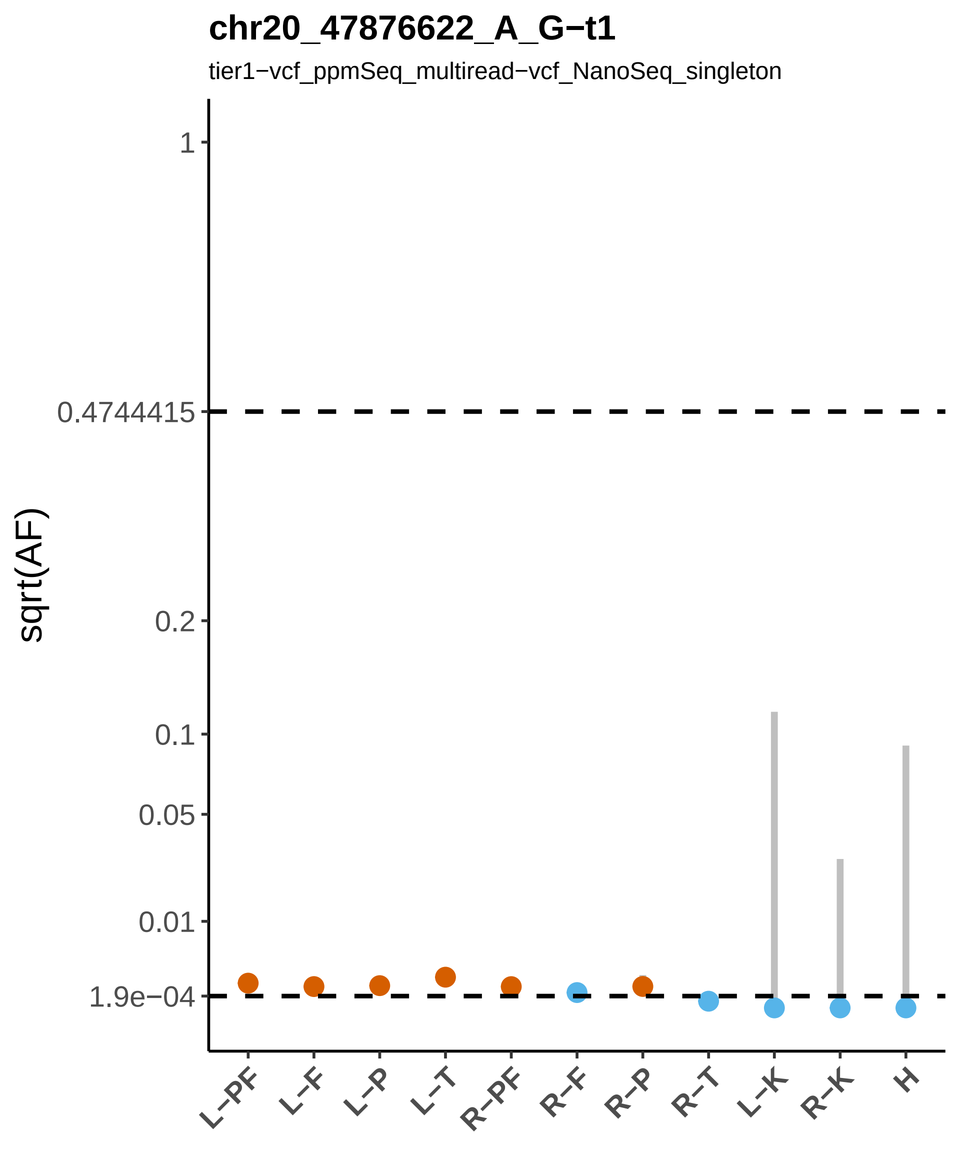

chr20_47876622_A_G−t1
tier1−vcf_ppmSeq_multiread−vcf_NanoSeq_singleton
sqrt(AF)
1
0.4744415
0.2
0.1
0.05
0.01
1.9e−04
L−PF
L−F
L−P
L−T
R−PF
R−F
R−P
R−T
L−K
R−K
H

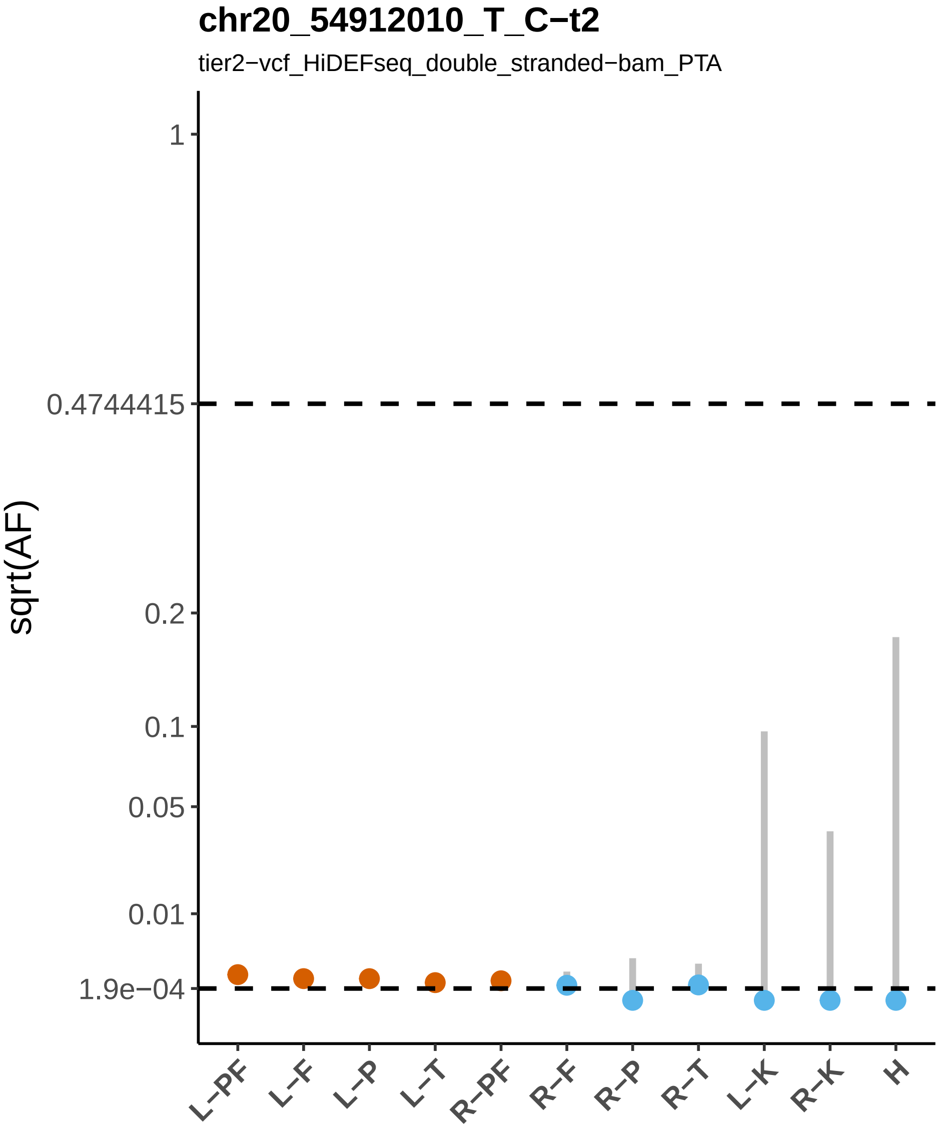

chr20_54912010_T_C−t2
tier2−vcf_HiDEFseq_double_stranded−bam_PTA
sqrt(AF)
1
0.4744415
0.2
0.1
0.05
0.01
1.9e−04
L−PF
L−F
L−P
L−T
R−PF
R−F
R−P
R−T
L−K
R−K
H

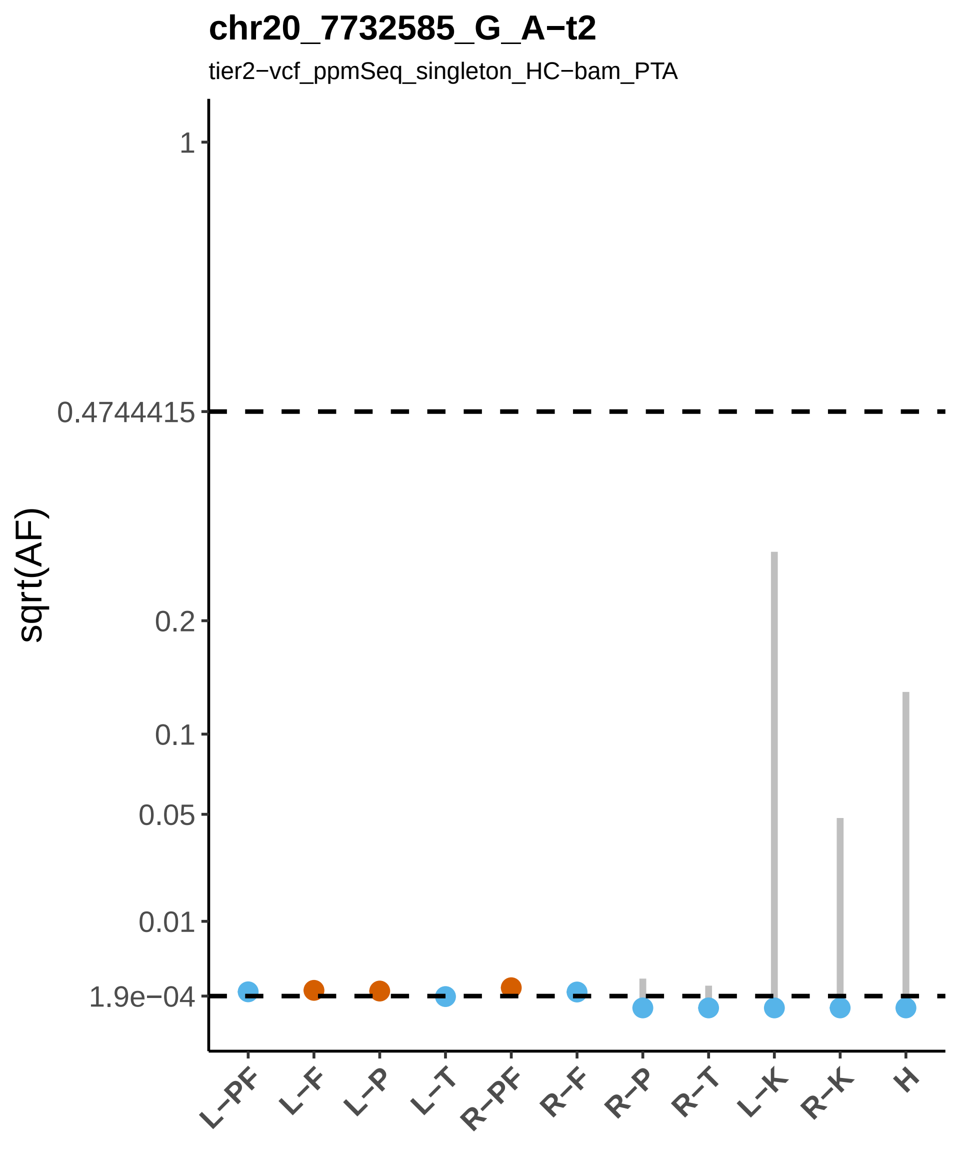
chr20_7732585_G_A−t2
tier2−vcf_ppmSeq_singleton_HC−bam_PTA
sqrt(AF)
1
0.4744415
0.2
0.1
0.05
0.01
1.9e−04
L−PF
L−F
L−P
L−T
R−PF
R−F
R−P
R−T
L−K
R−K
H

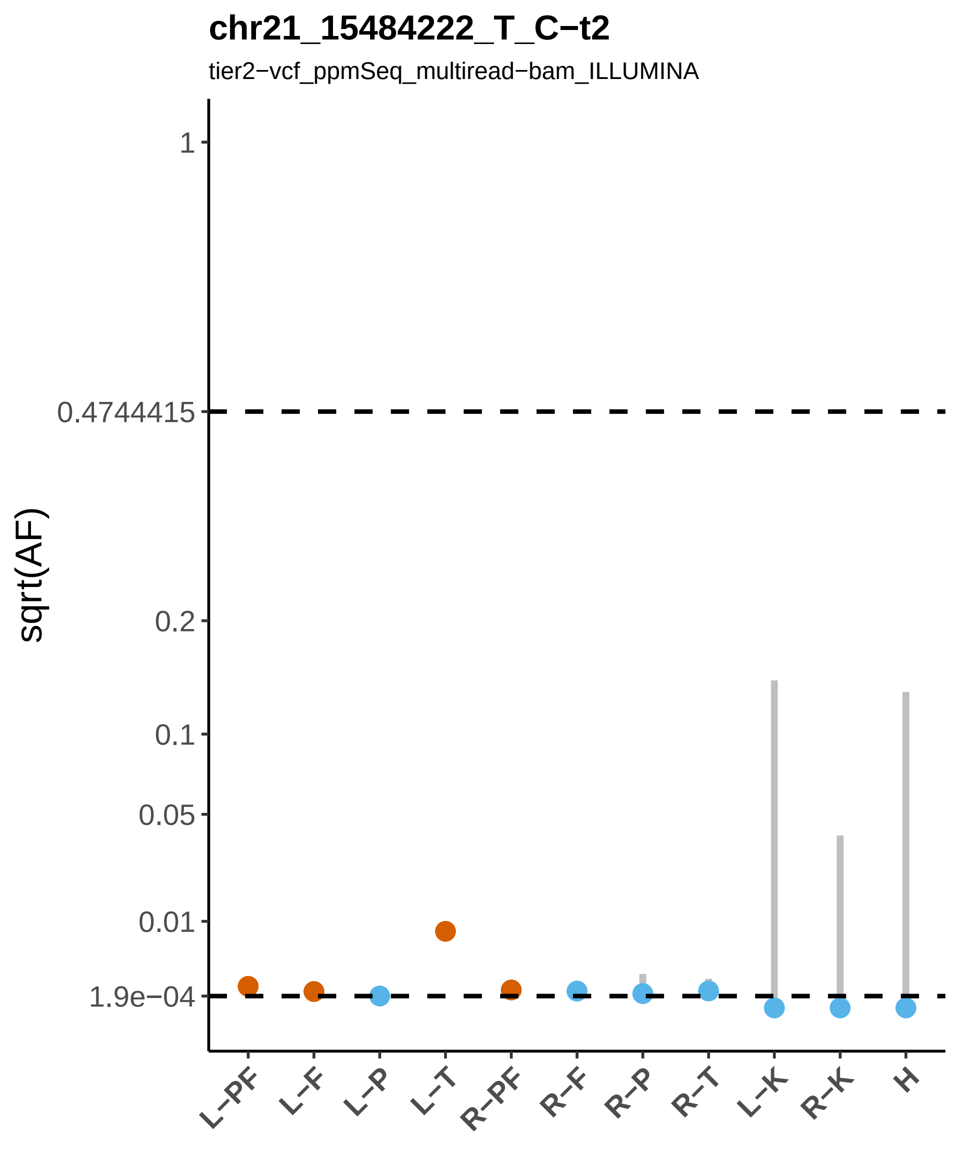

chr21_15484222_T_C−t2
tier2−vcf_ppmSeq_multiread−bam_ILLUMINA
sqrt(AF)
1
0.4744415
0.2
0.1
0.05
0.01
1.9e−04
L−PF
L−F
L−P
L−T
R−PF
R−F
R−P
R−T
L−K
R−K
H

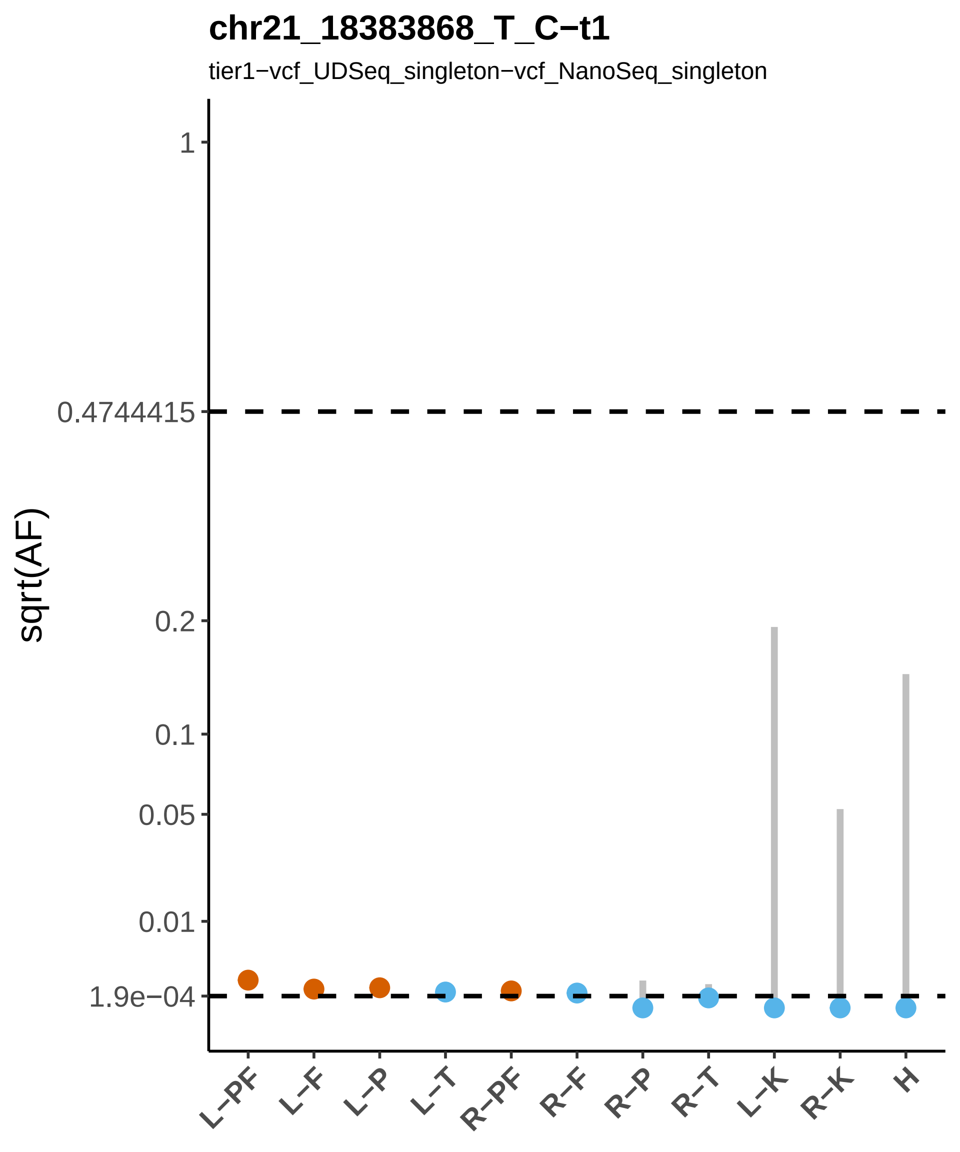

chr21_18383868_T_C−t1
tier1−vcf_UDSeq_singleton−vcf_NanoSeq_singleton
sqrt(AF)
1
0.4744415
0.2
0.1
0.05
0.01
1.9e−04
L−PF
L−F
L−P
L−T
R−PF
R−F
R−P
R−T
L−K
R−K
H

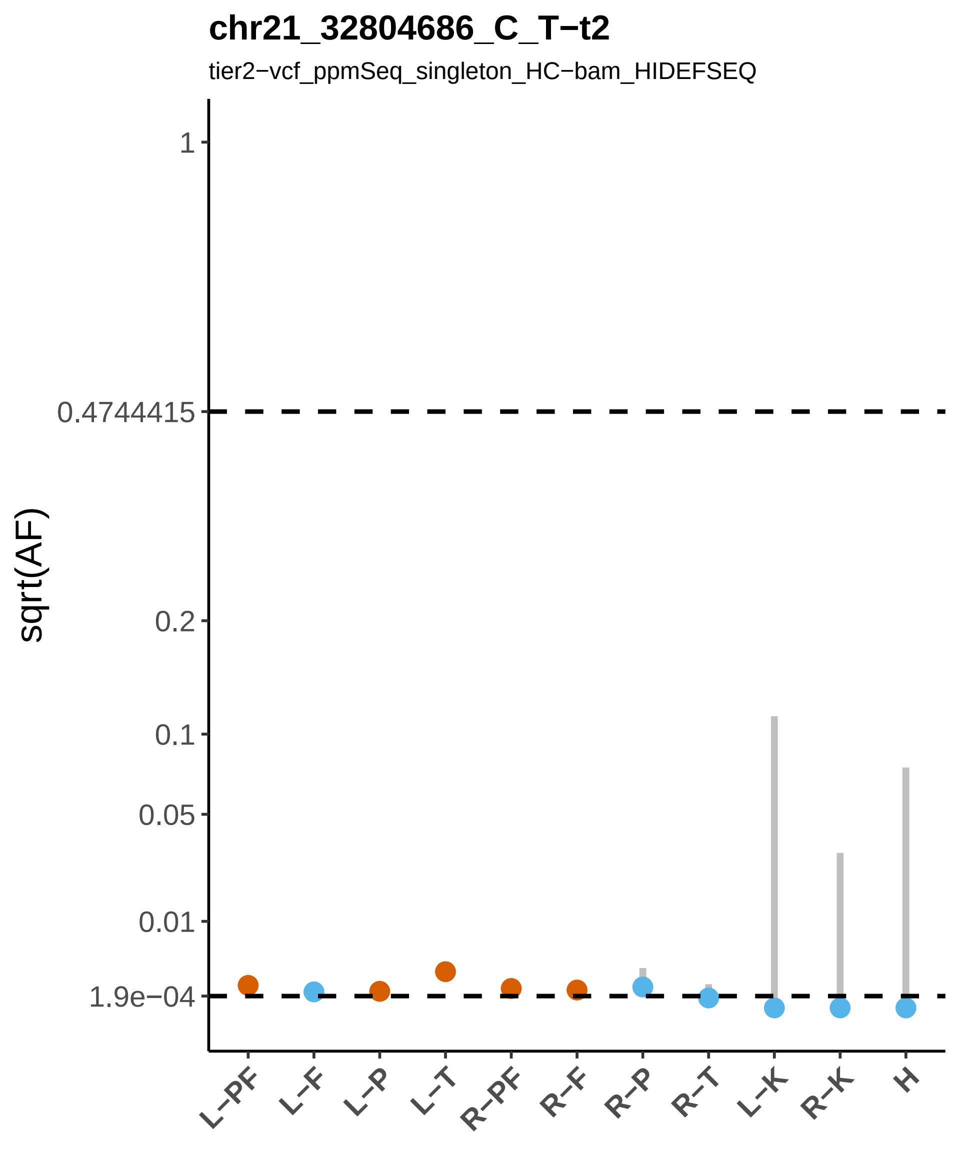
chr21_32804686_C_T−t2
tier2−vcf_ppmSeq_singleton_HC−bam_HIDEFSEQ
sqrt(AF)
1
0.4744415
0.2
0.1
0.05
0.01
1.9e−04
L−PF
L−F
L−P
L−T
R−PF
R−F
R−P
R−T
L−K
R−K
H

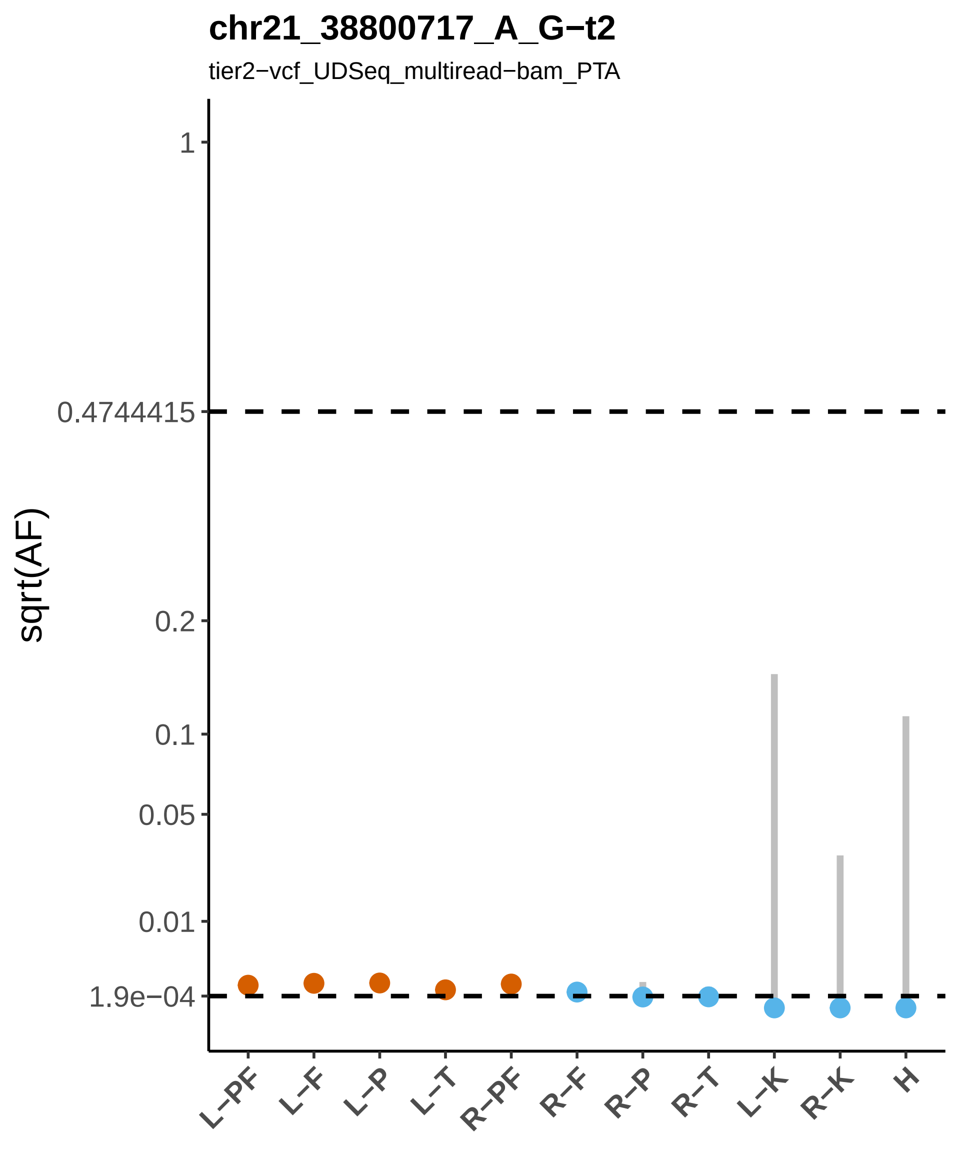
chr21_38800717_A_G−t2
tier2−vcf_UDSeq_multiread−bam_PTA
sqrt(AF)
1
0.4744415
0.2
0.1
0.05
0.01
1.9e−04
L−PF
L−F
L−P
L−T
R−PF
R−F
R−P
R−T
L−K
R−K
H

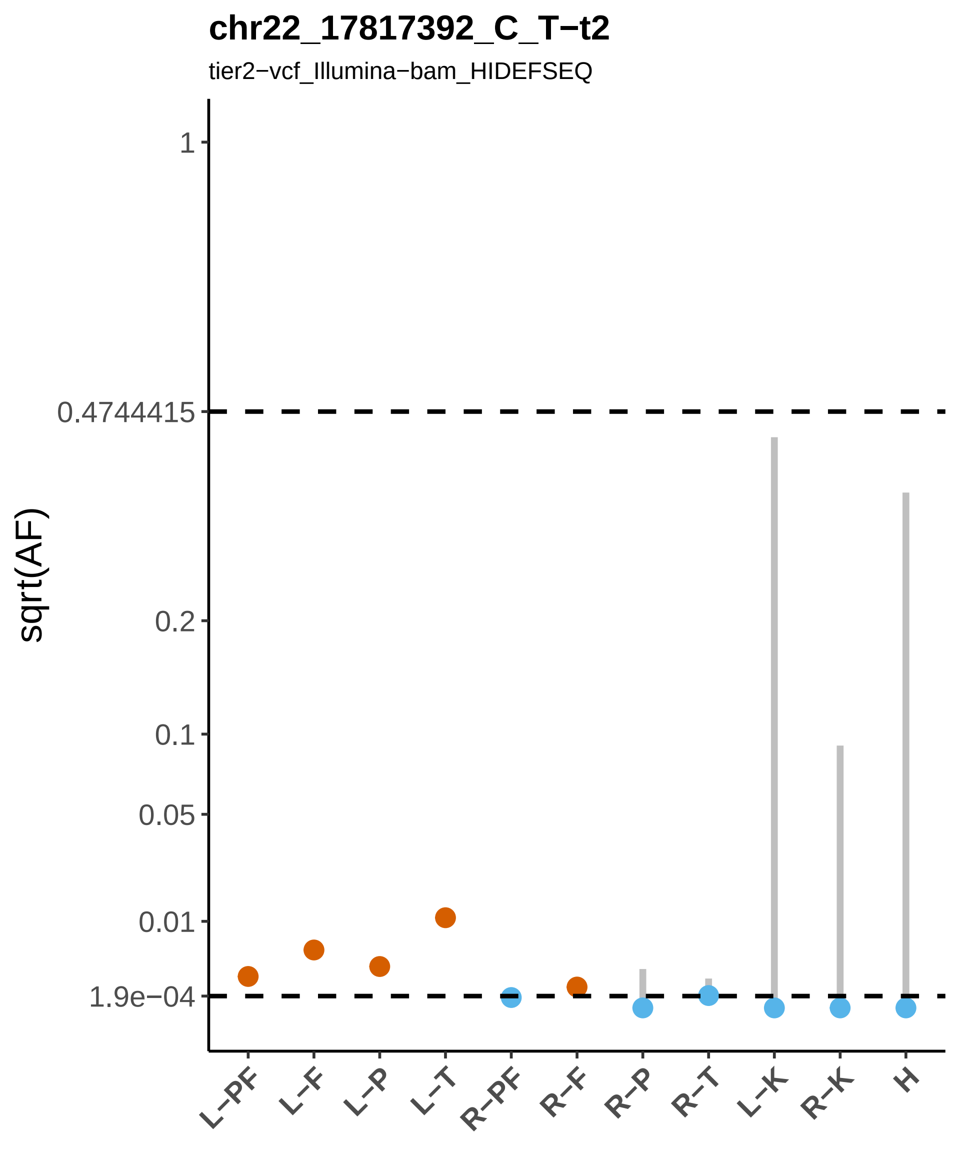
chr22_17817392_C_T−t2
tier2−vcf_Illumina−bam_HIDEFSEQ
sqrt(AF)
1
0.4744415
0.2
0.1
0.05
0.01
1.9e−04
L−PF
L−F
L−P
L−T
R−PF
R−F
R−P
R−T
L−K
R−K
H

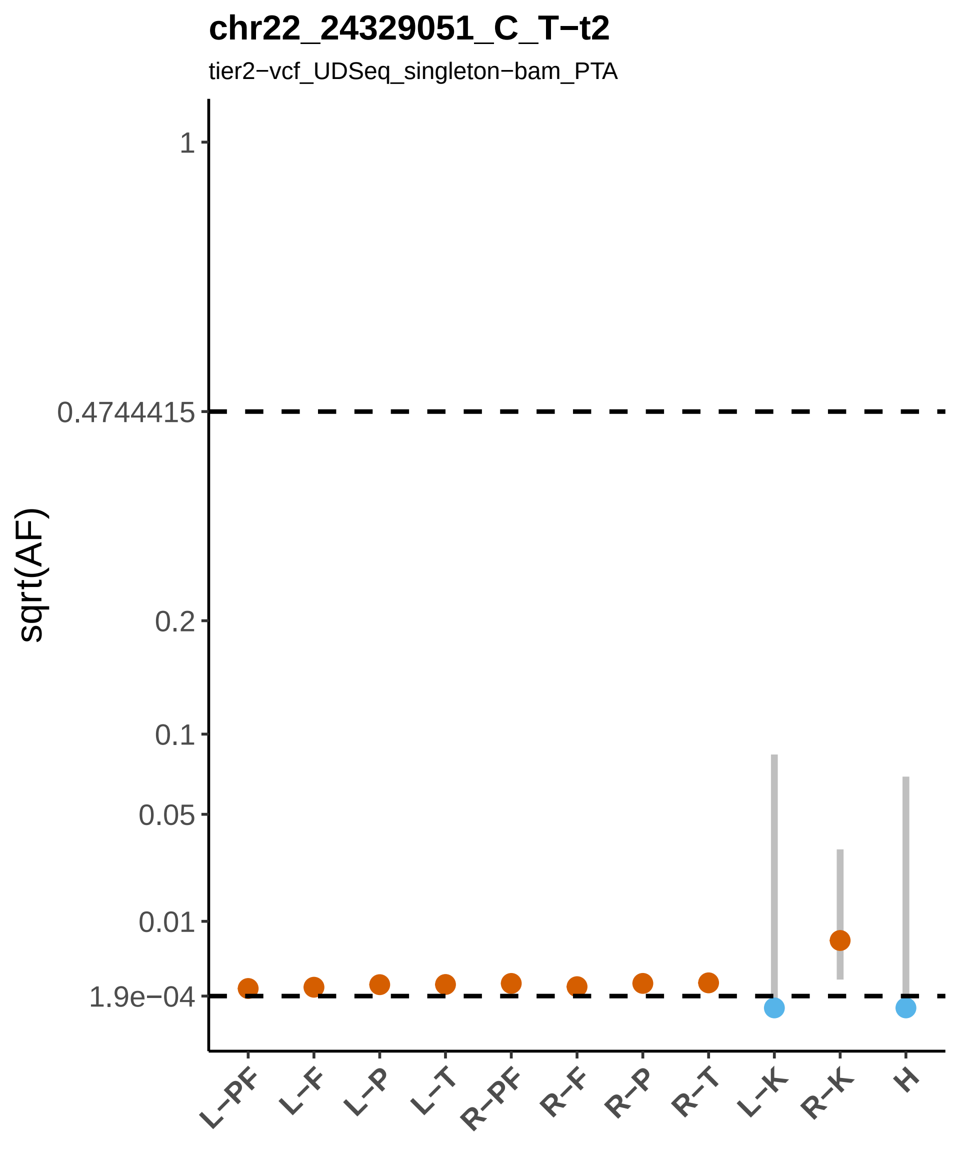
chr22_24329051_C_T−t2
tier2−vcf_UDSeq_singleton−bam_PTA
sqrt(AF)
1
0.4744415
0.2
0.1
0.05
0.01
1.9e−04
L−PF
L−F
L−P
L−T
R−PF
R−F
R−P
R−T
L−K
R−K
H

#### chr22_31918949_A_G−t1

tier1−vcf_ppmSeq_singleton_HC−vcf_NanoSeq_singleton

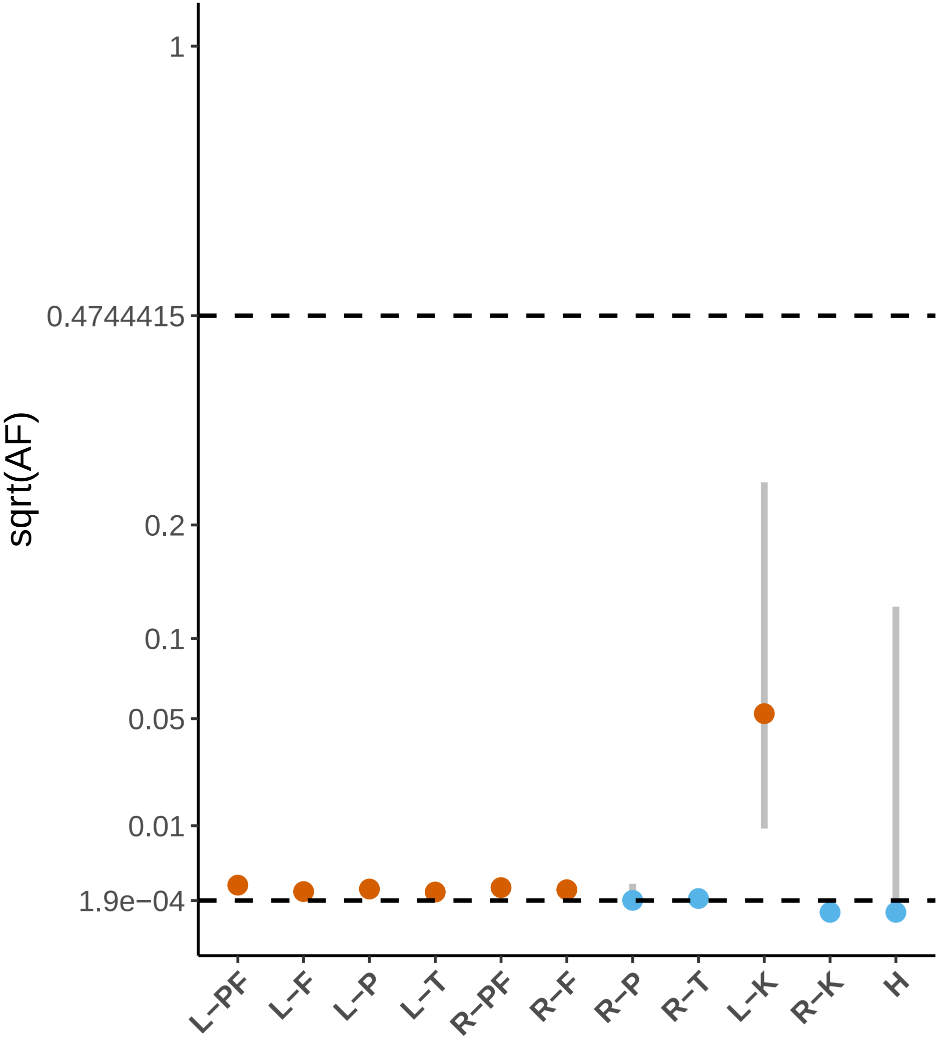

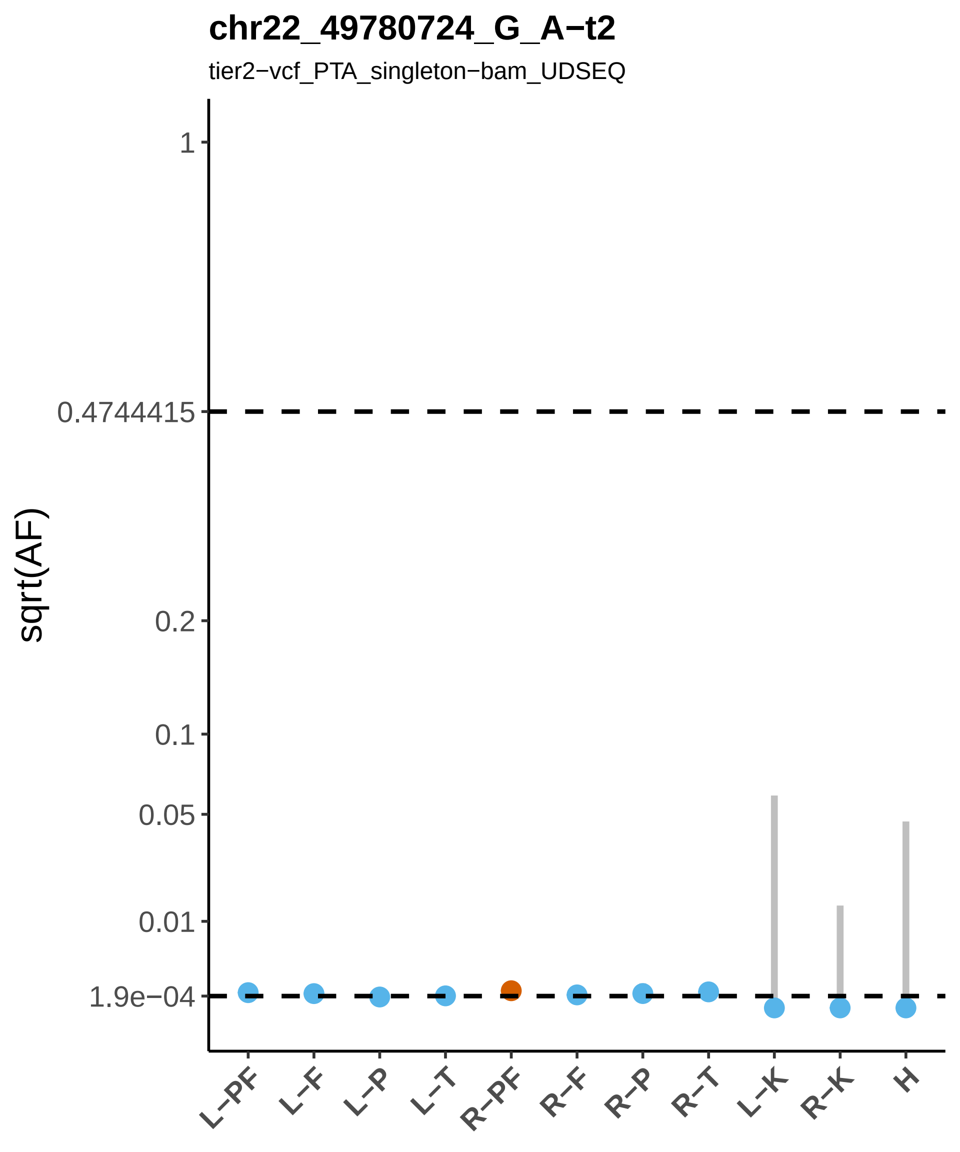
chr22_49780724_G_A−t2
tier2−vcf_PTA_singleton−bam_UDSEQ
sqrt(AF)
1
0.4744415
0.2
0.1
0.05
0.01
1.9e−04
L−PF
L−F
L−P
L−T
R−PF
R−F
R−P
R−T
L−K
R−K
H

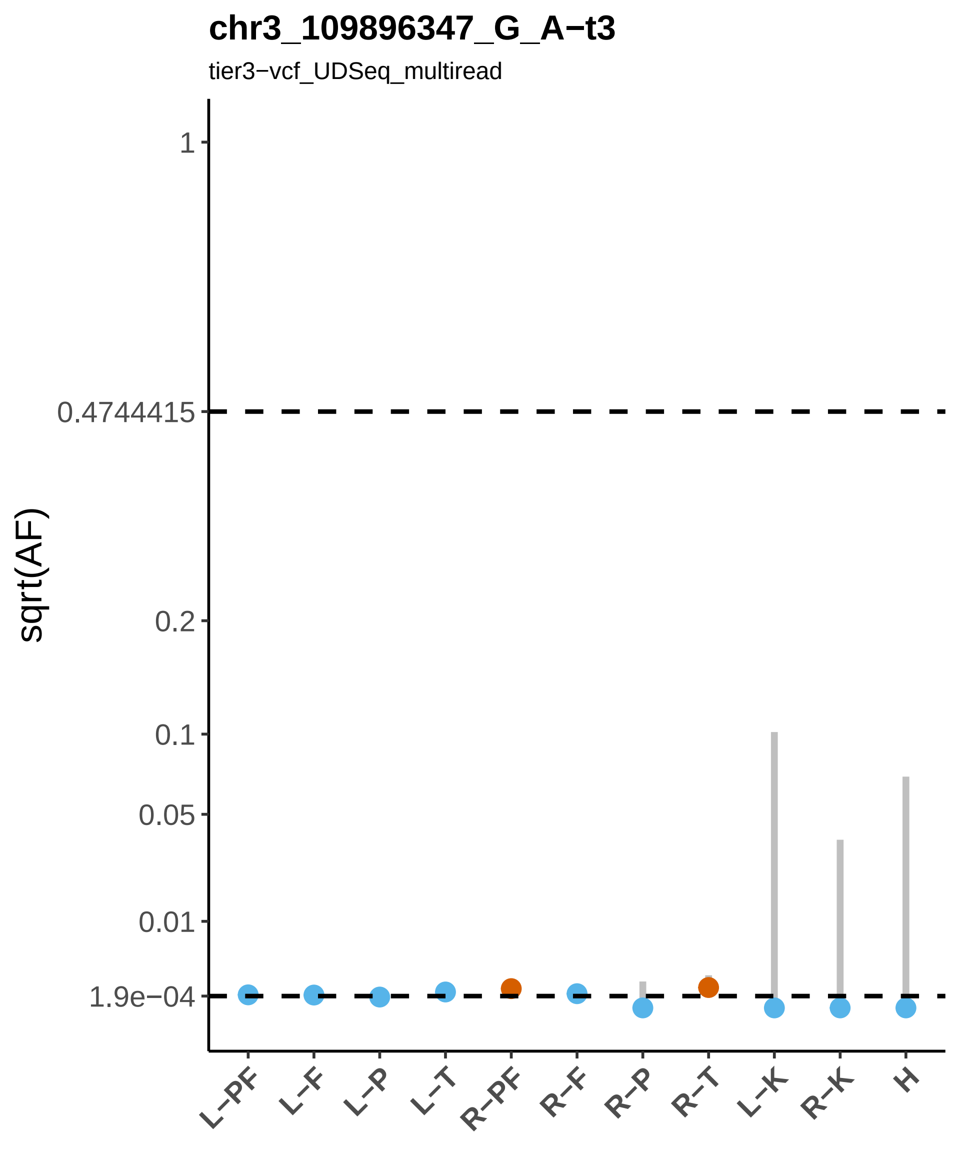

chr3_109896347_G_A−t3
tier3−vcf_UDSeq_multiread
sqrt(AF)
1
0.4744415
0.2
0.1
0.05
0.01
1.9e−04
L−PF
L−F
L−P
L−T
R−PF
R−F
R−P
R−T
L−K
R−K
H

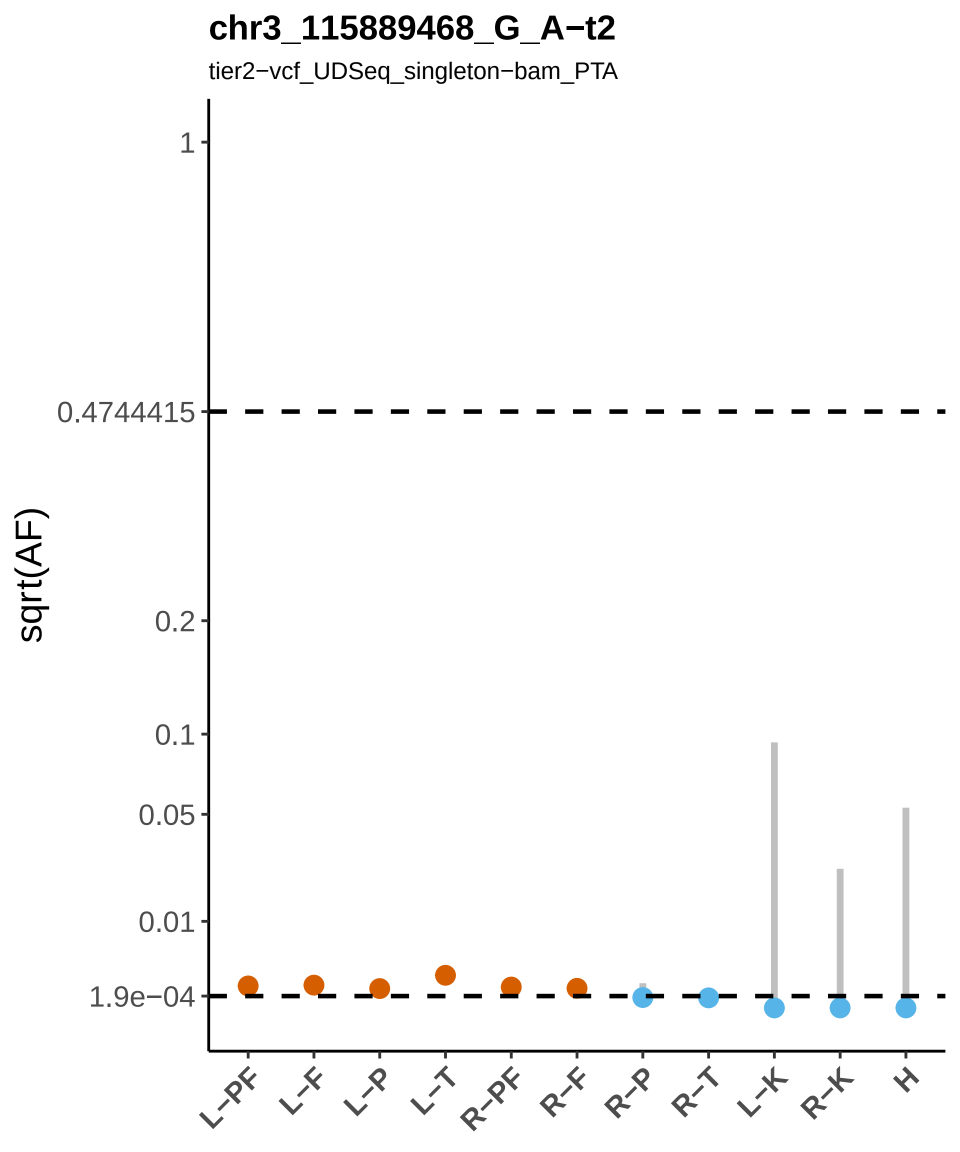

chr3_115889468_G_A−t2
tier2−vcf_UDSeq_singleton−bam_PTA
sqrt(AF)
1
0.4744415
0.2
0.1
0.05
0.01
1.9e−04
L−PF
L−F
L−P
L−T
R−PF
R−F
R−P
R−T
L−K
R−K
H

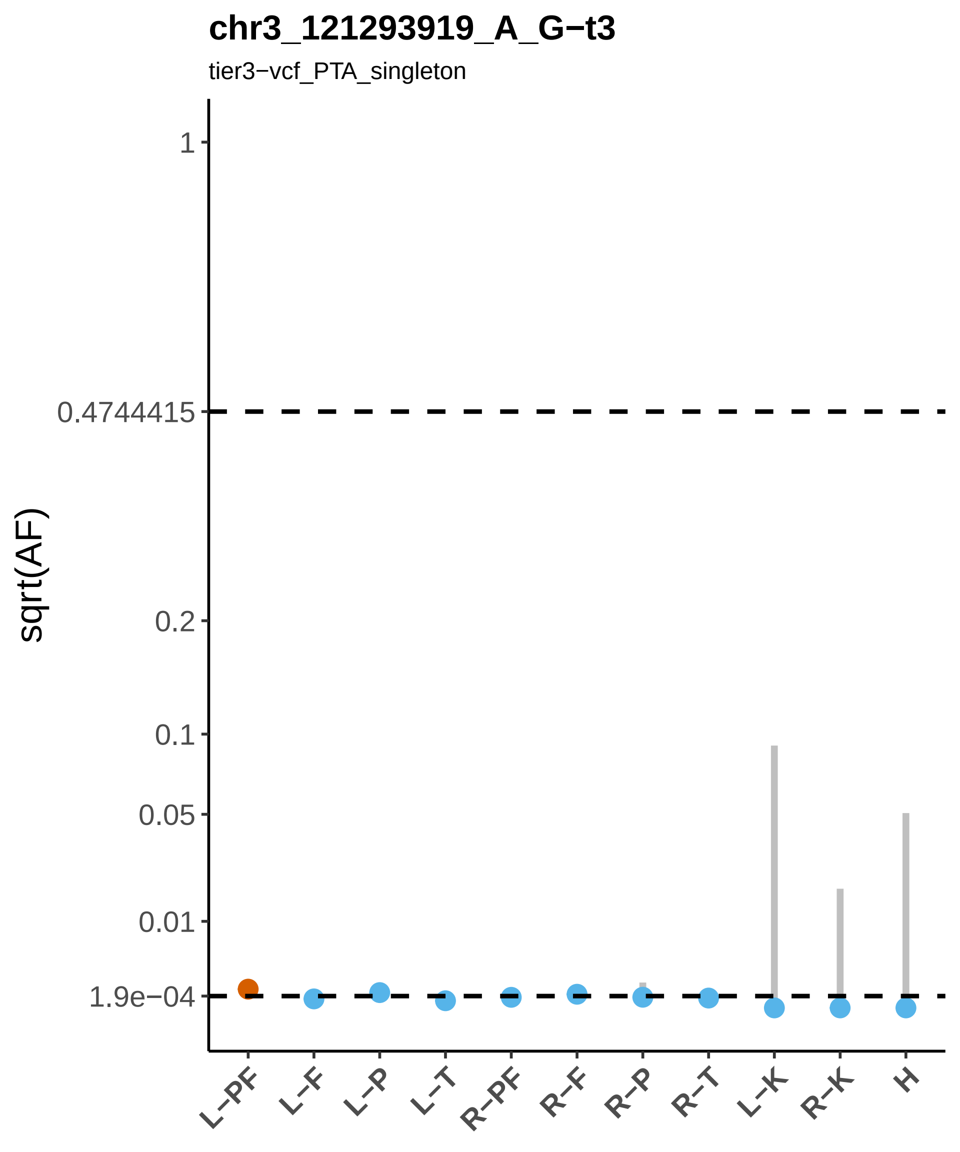

chr3_121293919_A_G−t3
tier3−vcf_PTA_singleton
sqrt(AF)
1
0.4744415
0.2
0.1
0.05
0.01
1.9e−04
L−PF
L−F
L−P
L−T
R−PF
R−F
R−P
R−T
L−K
R−K
H

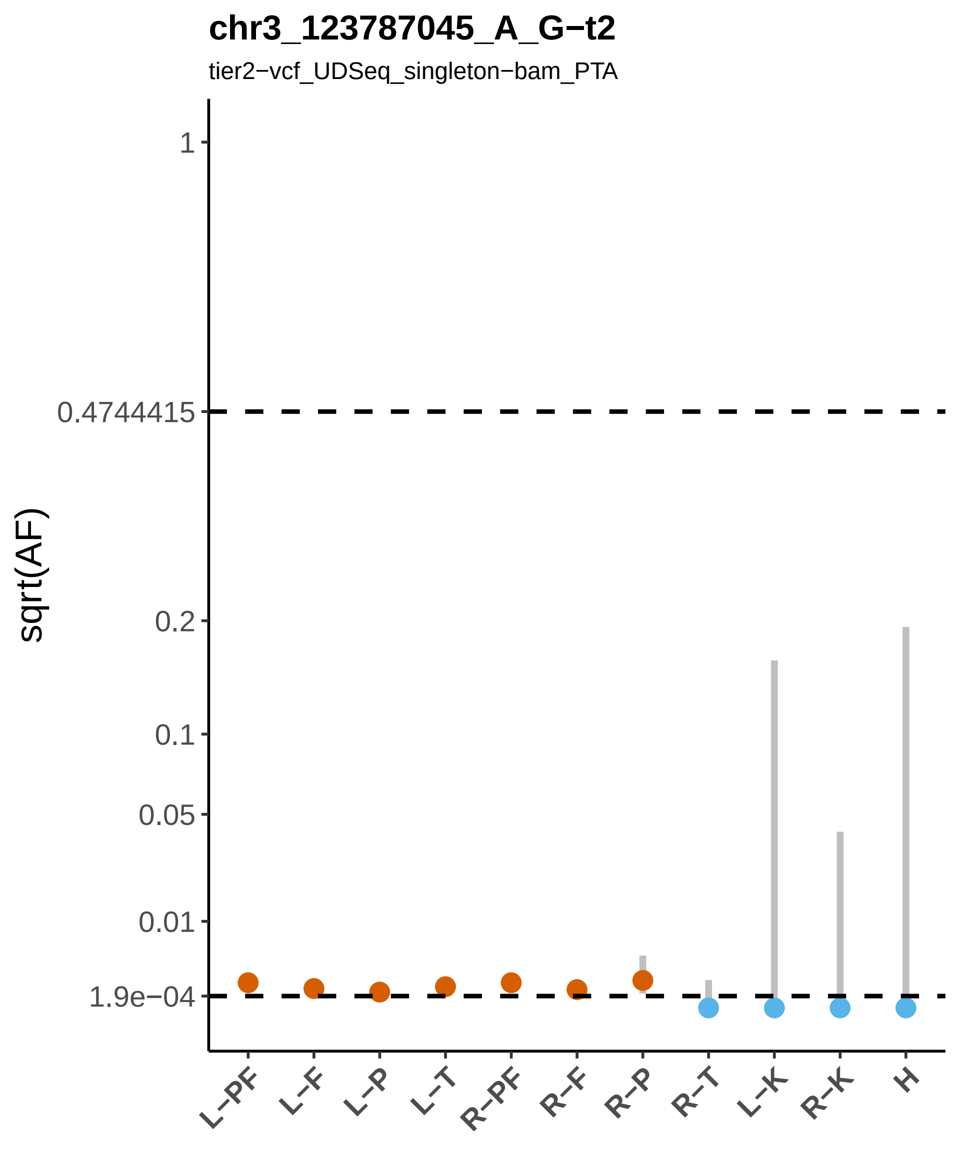
chr3_123787045_A_G−t2
tier2−vcf_UDSeq_singleton−bam_PTA
sqrt(AF)
1
0.4744415
0.2
0.1
0.05
0.01
1.9e−04
L−PF
L−F
L−P
L−T
R−PF
R−F
R−P
R−T
L−K
R−K
H

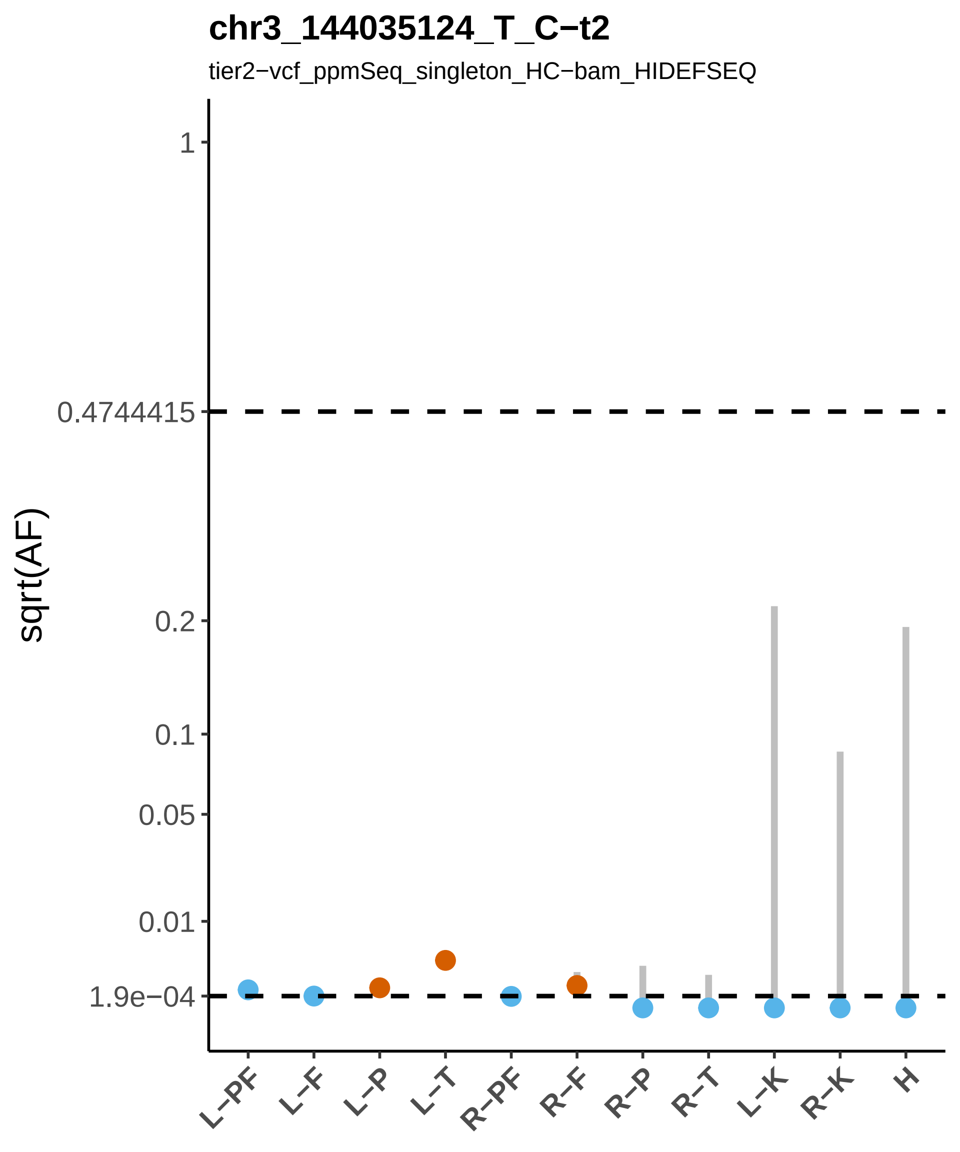
chr3_144035124_T_C−t2
tier2−vcf_ppmSeq_singleton_HC−bam_HIDEFSEQ
sqrt(AF)
1
0.4744415
0.2
0.1
0.05
0.01
1.9e−04
L−PF
L−F
L−P
L−T
R−PF
R−F
R−P
R−T
L−K
R−K
H

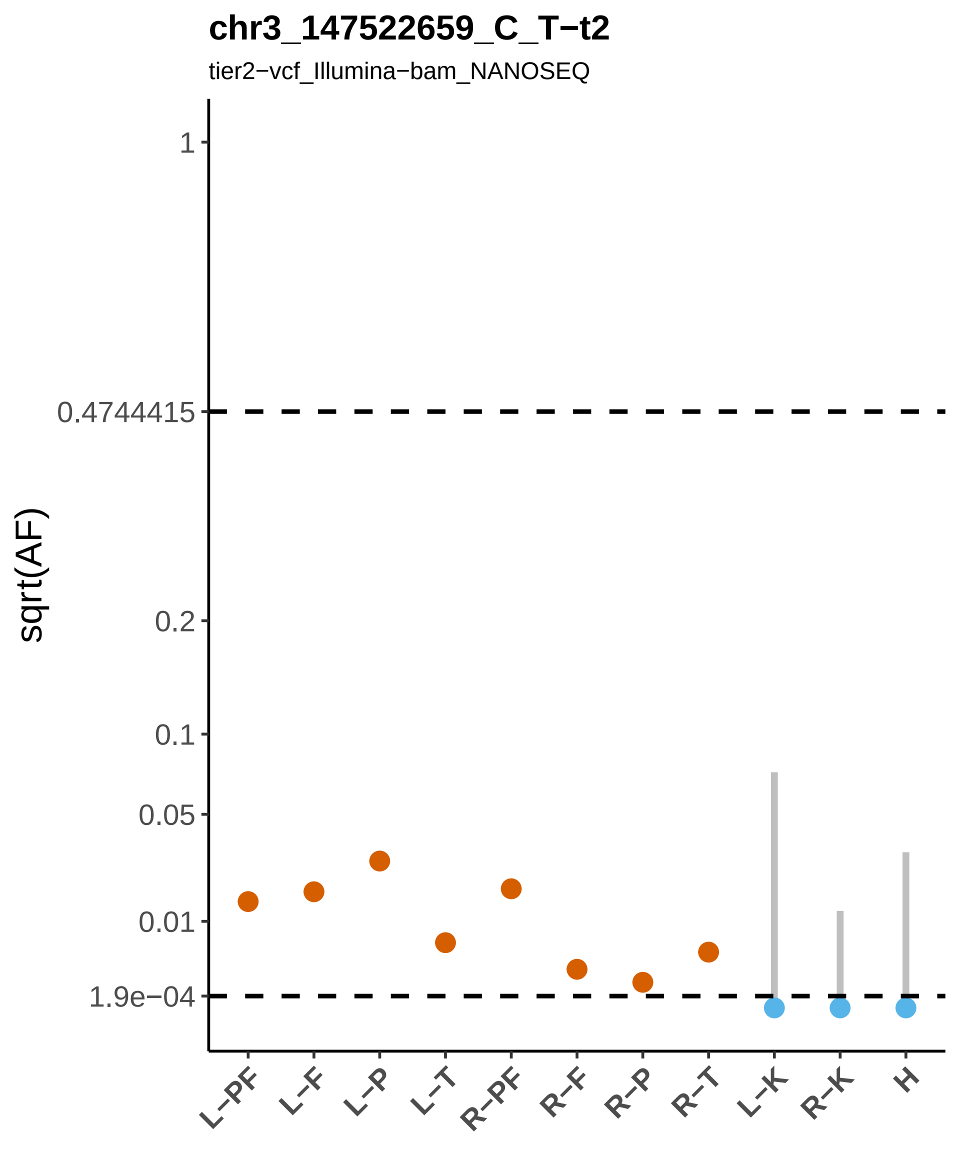
chr3_147522659_C_T−t2
tier2−vcf_Illumina−bam_NANOSEQ
sqrt(AF)
1
0.4744415
0.2
0.1
0.05
0.01
1.9e−04
L−PF
L−F
L−P
L−T
R−PF
R−F
R−P
R−T
L−K
R−K
H

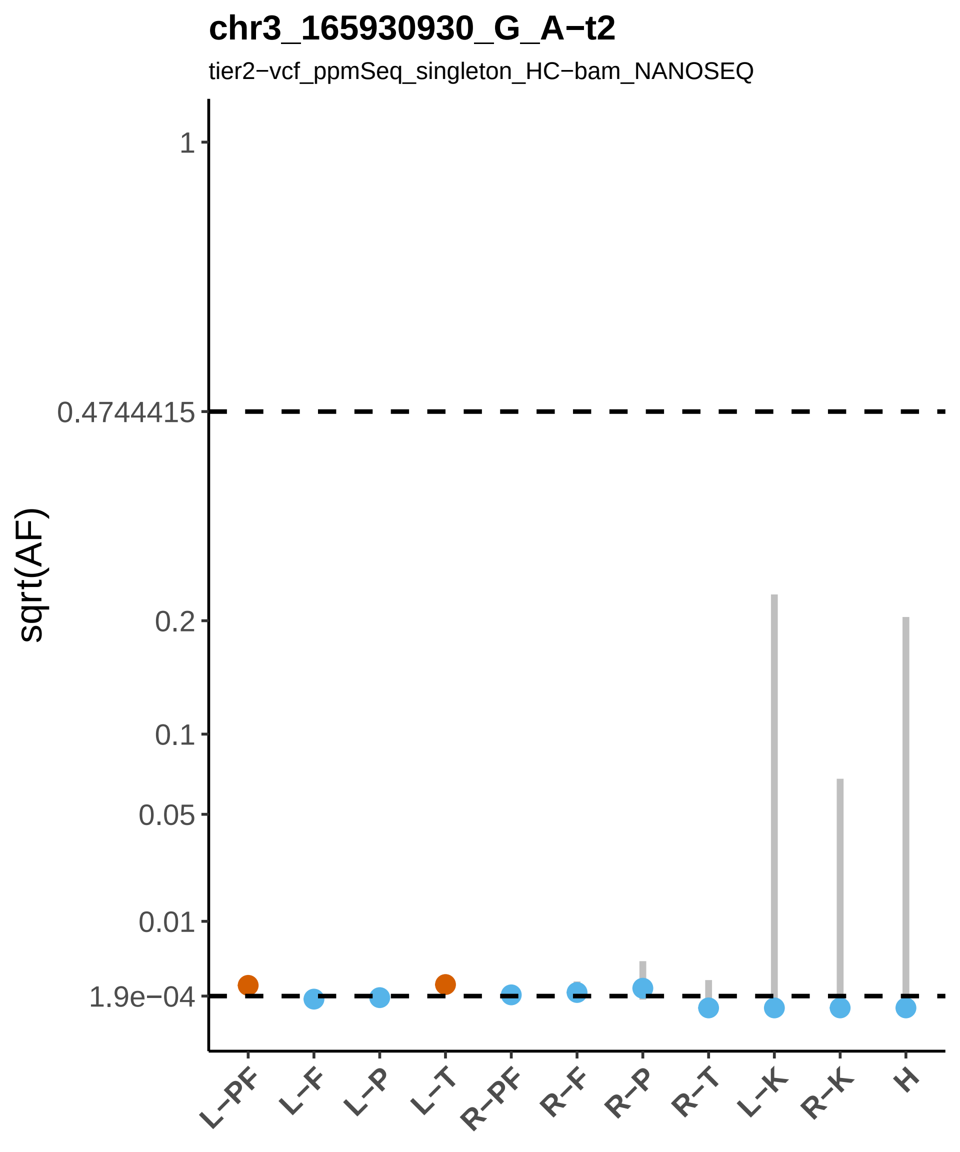

chr3_165930930_G_A−t2
tier2−vcf_ppmSeq_singleton_HC−bam_NANOSEQ
sqrt(AF)
1
0.4744415
0.2
0.1
0.05
0.01
1.9e−04
L−PF
L−F
L−P
L−T
R−PF
R−F
R−P
R−T
L−K
R−K
H

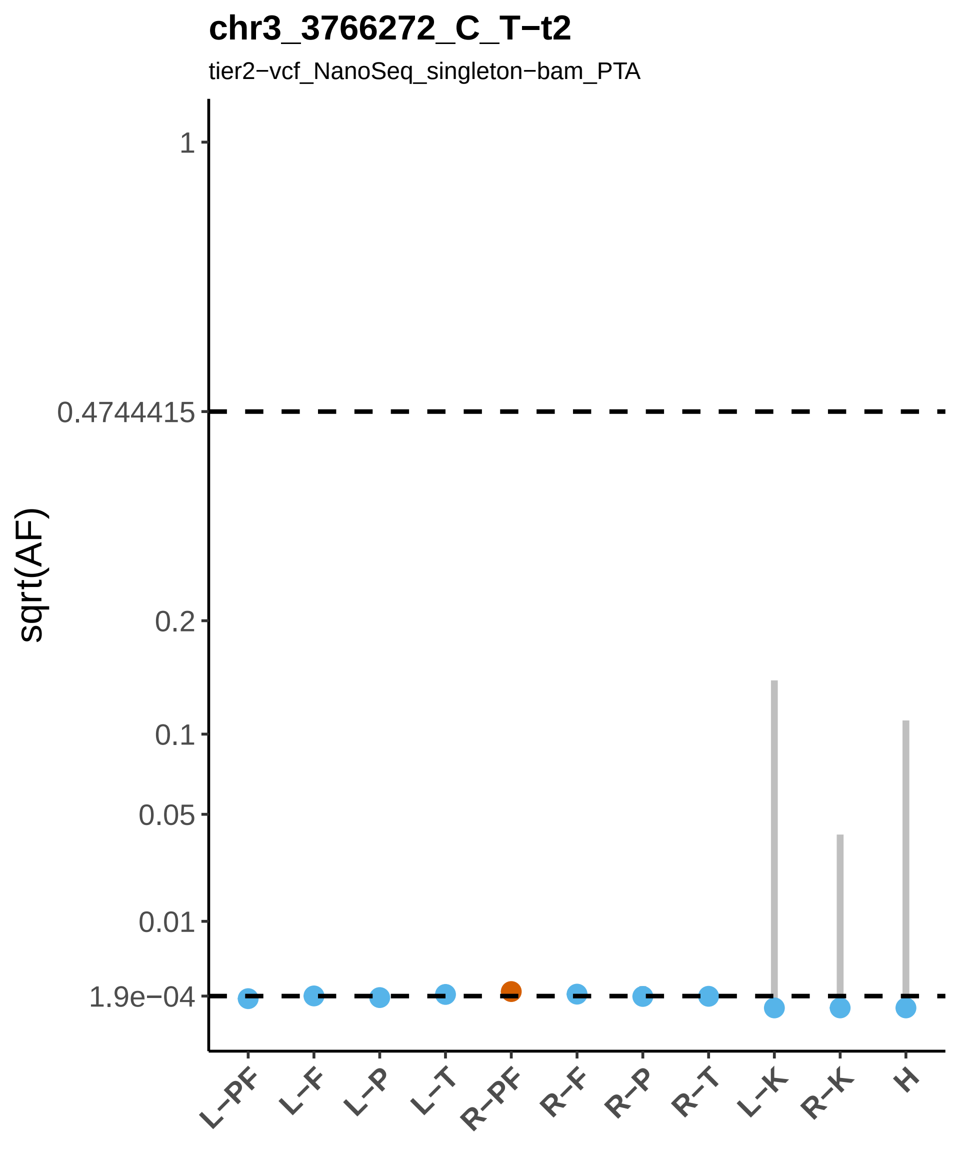
chr3_3766272_C_T−t2
tier2−vcf_NanoSeq_singleton−bam_PTA
sqrt(AF)
1
0.4744415
0.2
0.1
0.05
0.01
1.9e−04
L−PF
L−F
L−P
L−T
R−PF
R−F
R−P
R−T
L−K
R−K
H

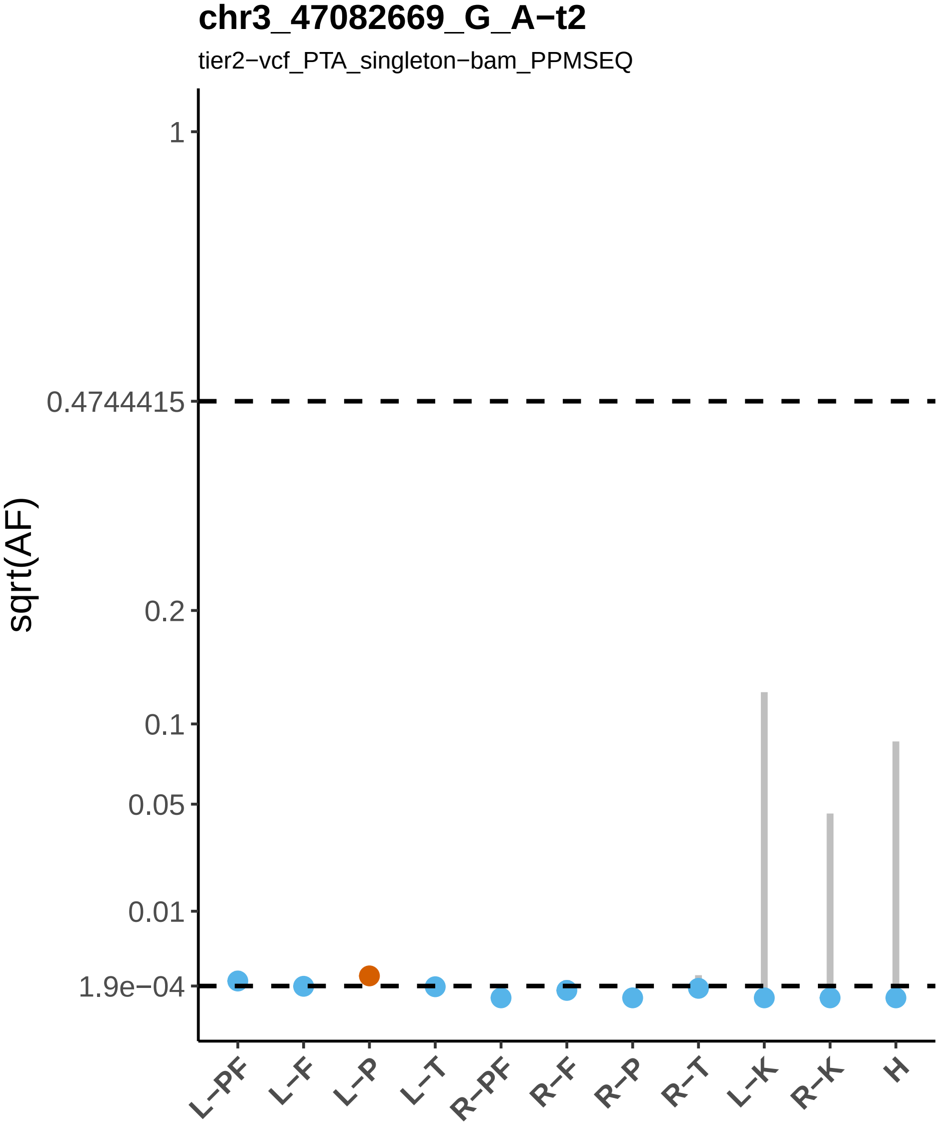

chr3_47082669_G_A−t2
tier2−vcf_PTA_singleton−bam_PPMSEQ
sqrt(AF)
1
0.4744415
0.2
0.1
0.05
0.01
1.9e−04
L−PF
L−F
L−P
L−T
R−PF
R−F
R−P
R−T
L−K
R−K
H

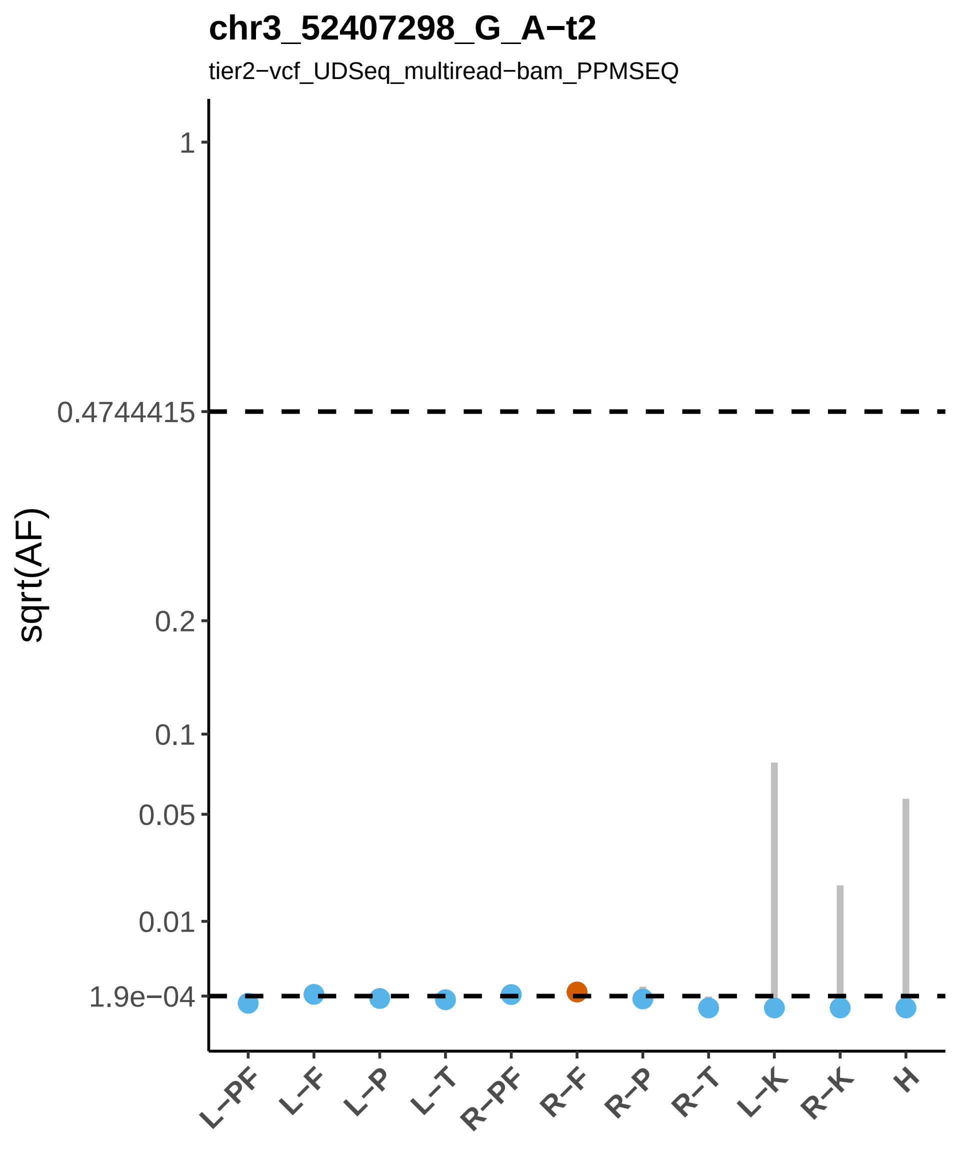

chr3_52407298_G_A−t2
tier2−vcf_UDSeq_multiread−bam_PPMSEQ
sqrt(AF)
1
0.4744415
0.2
0.1
0.05
0.01
1.9e−04
L−PF
L−F
L−P
L−T
R−PF
R−F
R−P
R−T
L−K
R−K
H

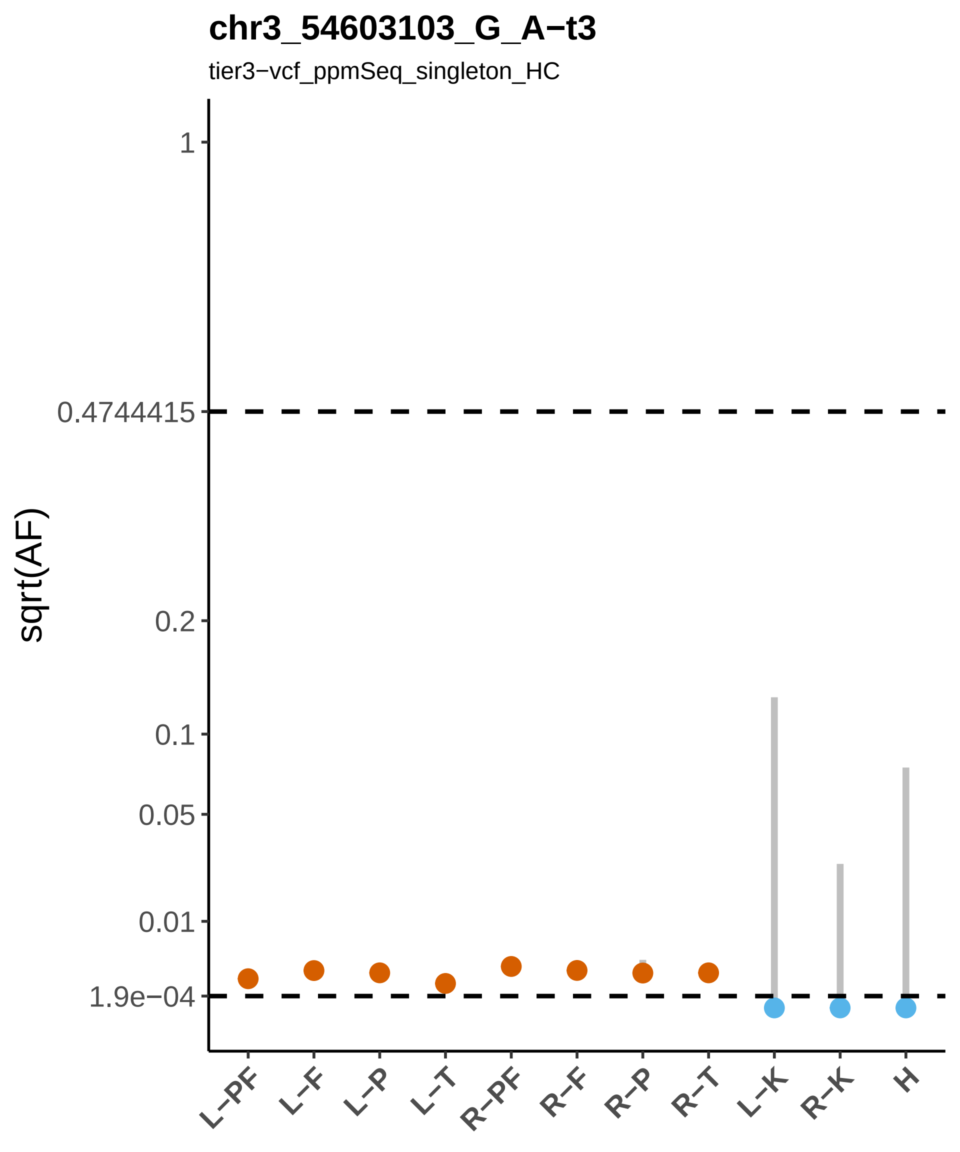
chr3_54603103_G_A−t3
tier3−vcf_ppmSeq_singleton_HC
sqrt(AF)
1
0.4744415
0.2
0.1
0.05
0.01
1.9e−04
L−PF
L−F
L−P
L−T
R−PF
R−F
R−P
R−T
L−K
R−K
H

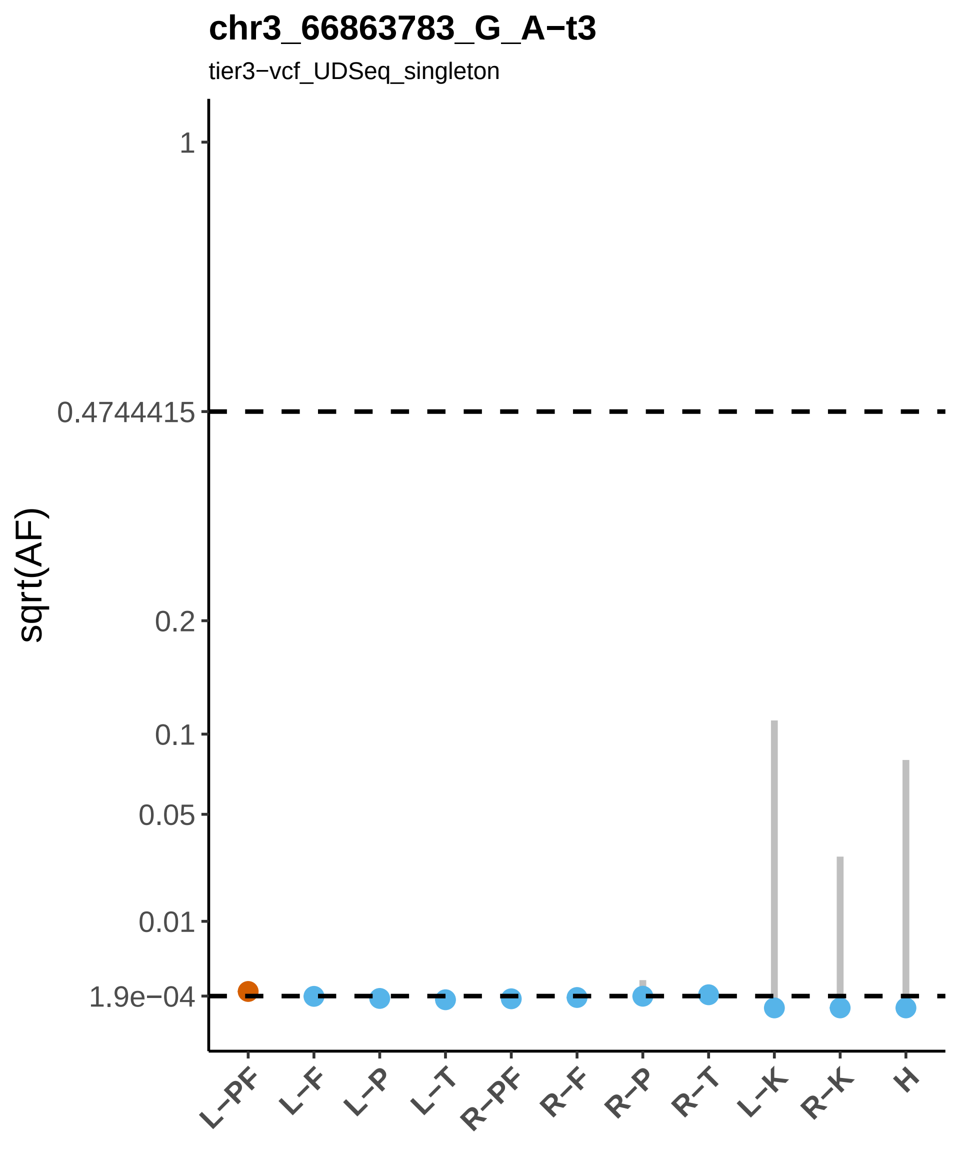
chr3_66863783_G_A−t3
tier3−vcf_UDSeq_singleton
sqrt(AF)
1
0.4744415
0.2
0.1
0.05
0.01
1.9e−04
L−PF
L−F
L−P
L−T
R−PF
R−F
R−P
R−T
L−K
R−K
H

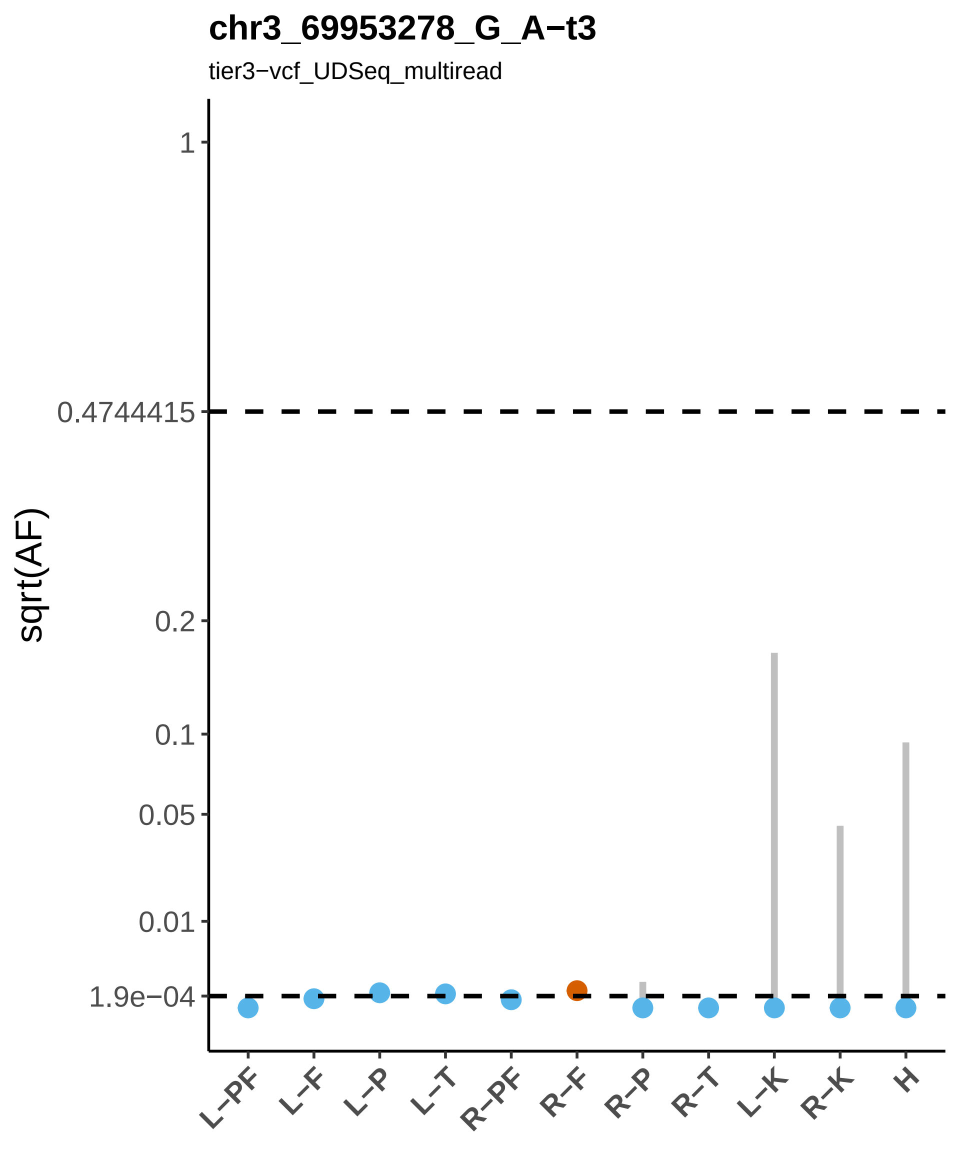
chr3_69953278_G_A−t3
tier3−vcf_UDSeq_multiread
sqrt(AF)
1
0.4744415
0.2
0.1
0.05
0.01
1.9e−04
L−PF
L−F
L−P
L−T
R−PF
R−F
R−P
R−T
L−K
R−K
H

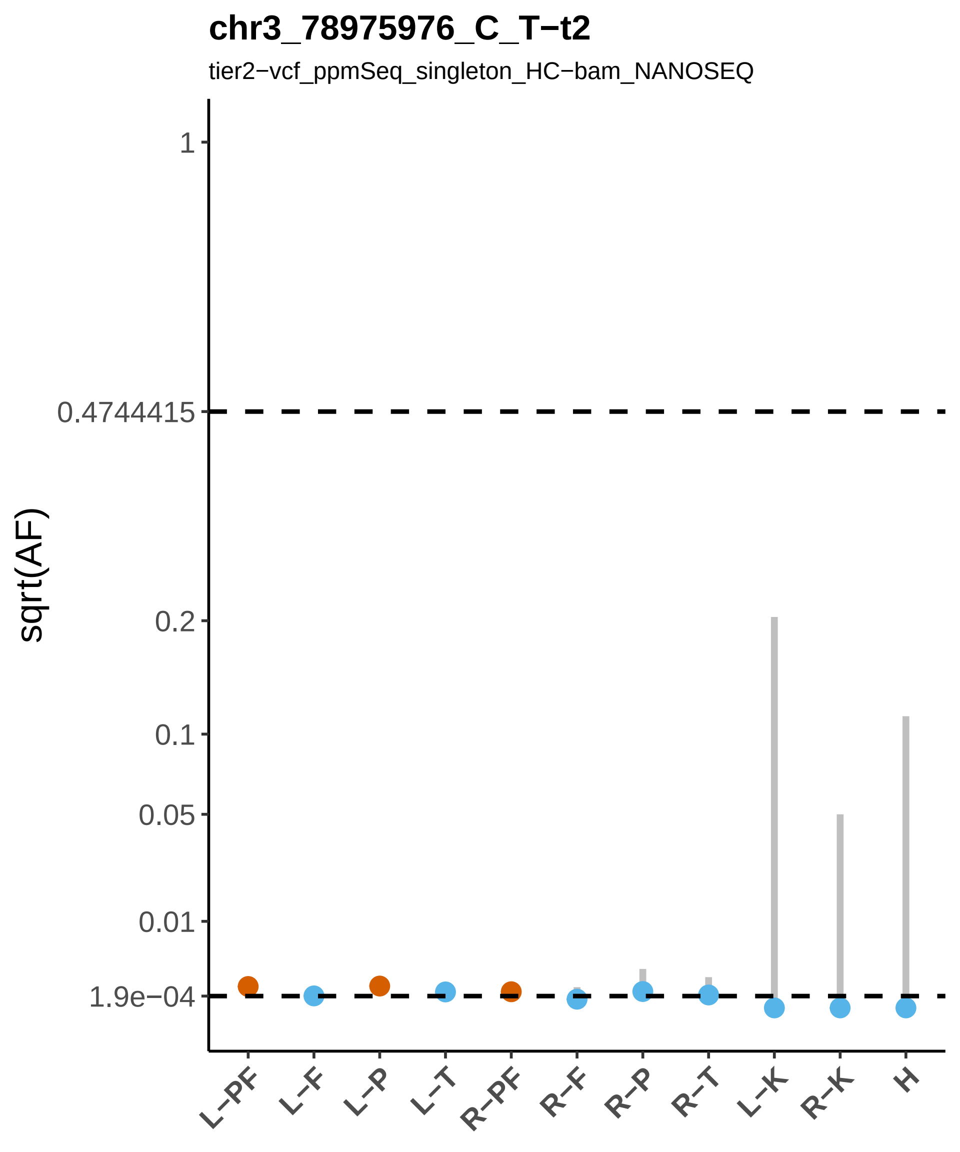
chr3_78975976_C_T−t2
tier2−vcf_ppmSeq_singleton_HC−bam_NANOSEQ
sqrt(AF)
1
0.4744415
0.2
0.1
0.05
0.01
1.9e−04
L−PF
L−F
L−P
L−T
R−PF
R−F
R−P
R−T
L−K
R−K
H

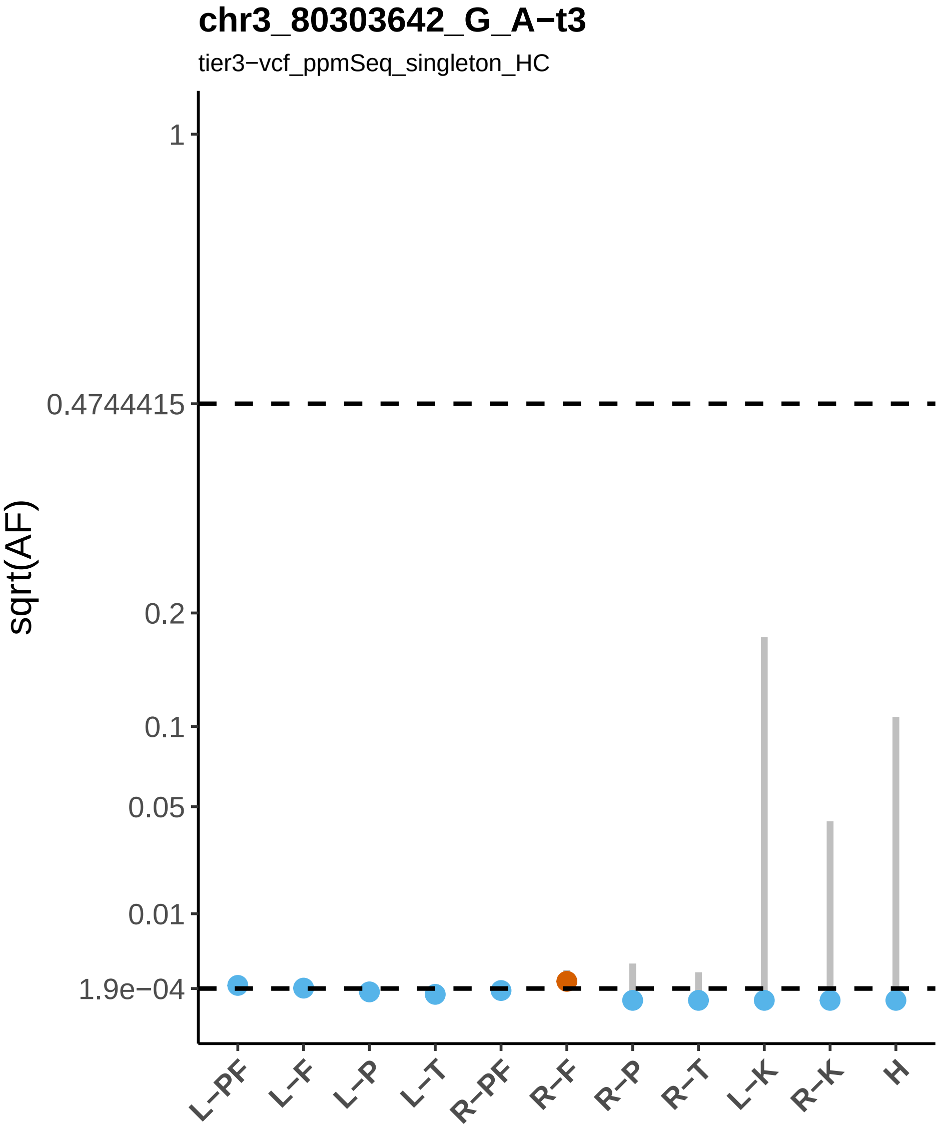

chr3_80303642_G_A−t3
tier3−vcf_ppmSeq_singleton_HC
sqrt(AF)
1
0.4744415
0.2
0.1
0.05
0.01
1.9e−04
L−PF
L−F
L−P
L−T
R−PF
R−F
R−P
R−T
L−K
R−K
H

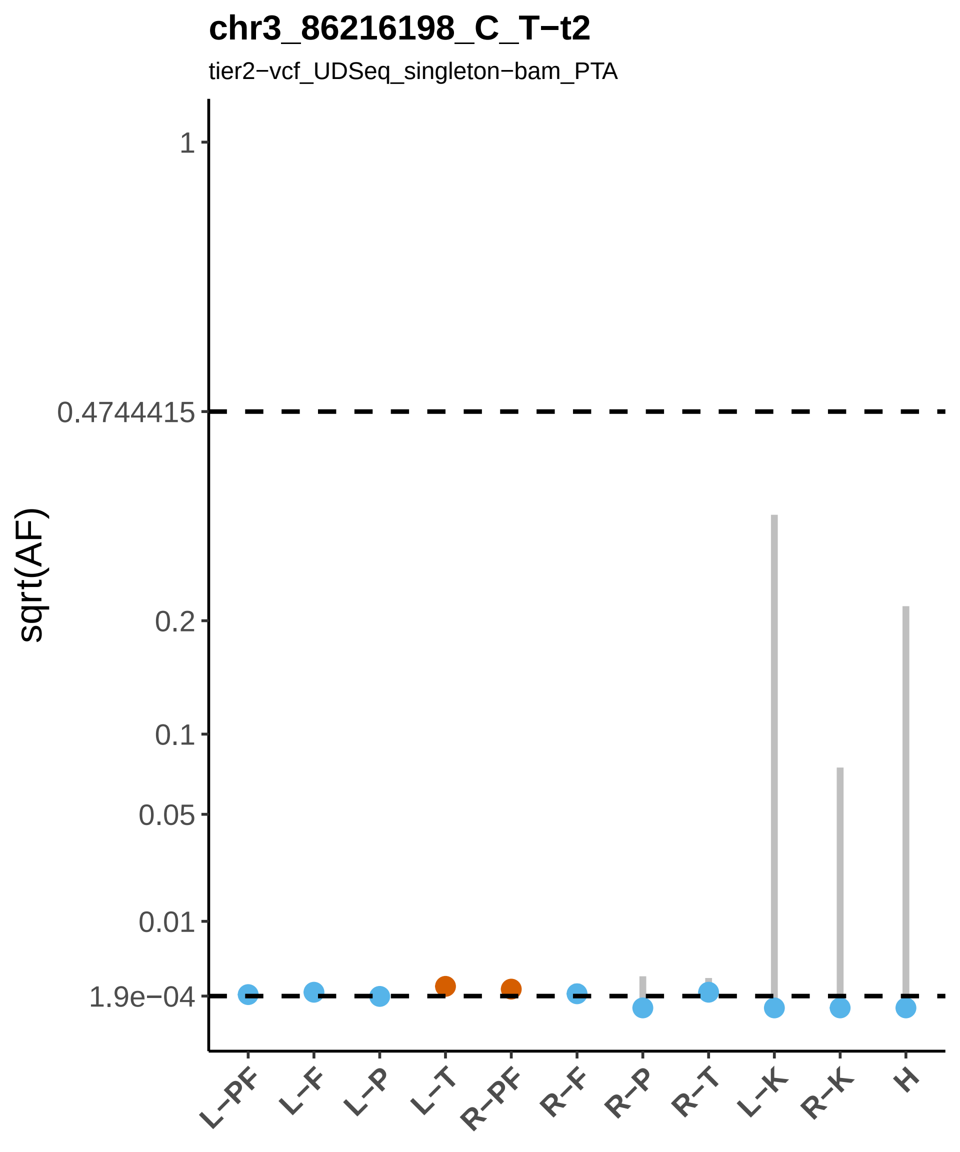

chr3_86216198_C_T−t2
tier2−vcf_UDSeq_singleton−bam_PTA
sqrt(AF)
1
0.4744415
0.2
0.1
0.05
0.01
1.9e−04
L−PF
L−F
L−P
L−T
R−PF
R−F
R−P
R−T
L−K
R−K
H

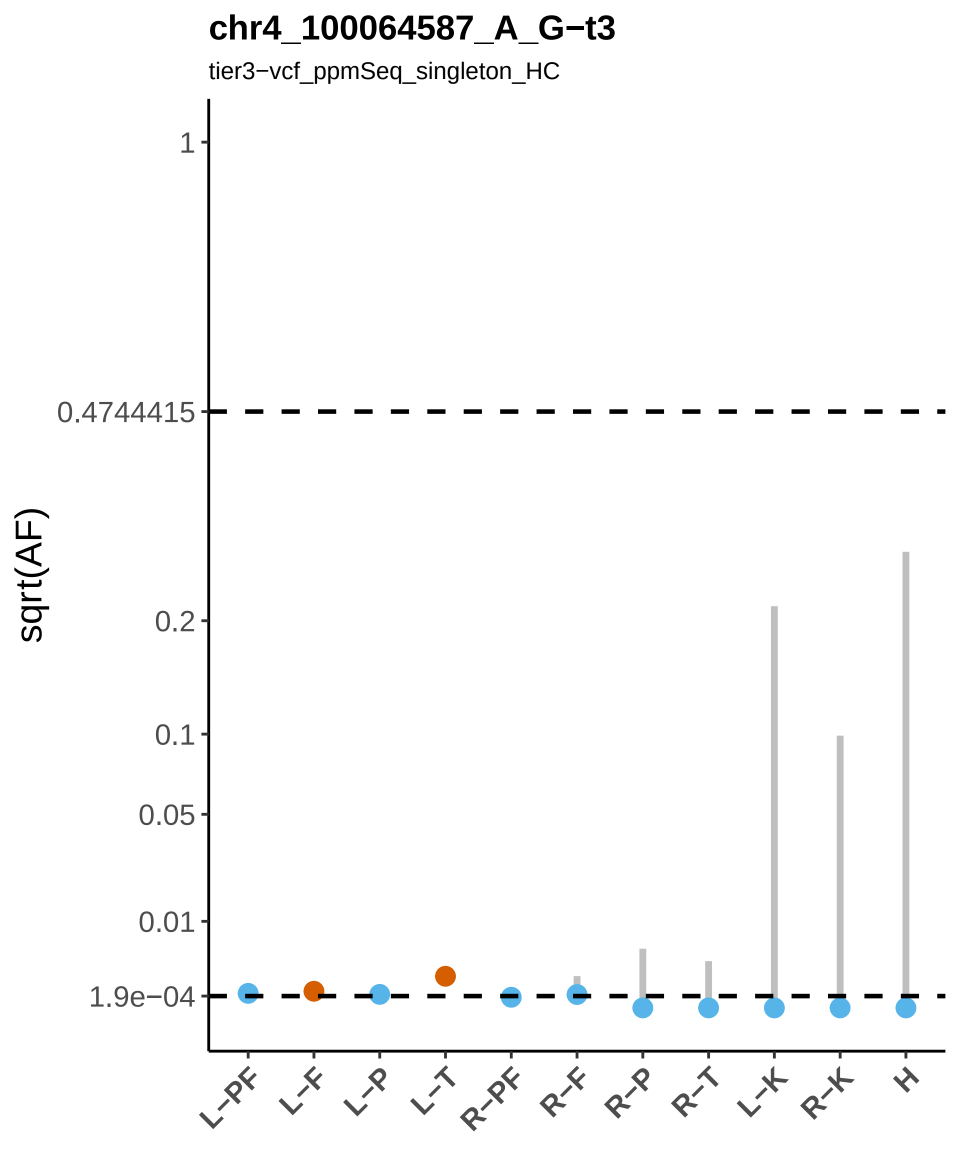

chr4_100064587_A_G−t3
tier3−vcf_ppmSeq_singleton_HC
sqrt(AF)
1
0.4744415
0.2
0.1
0.05
0.01
1.9e−04
L−PF
L−F
L−P
L−T
R−PF
R−F
R−P
R−T
L−K
R−K
H

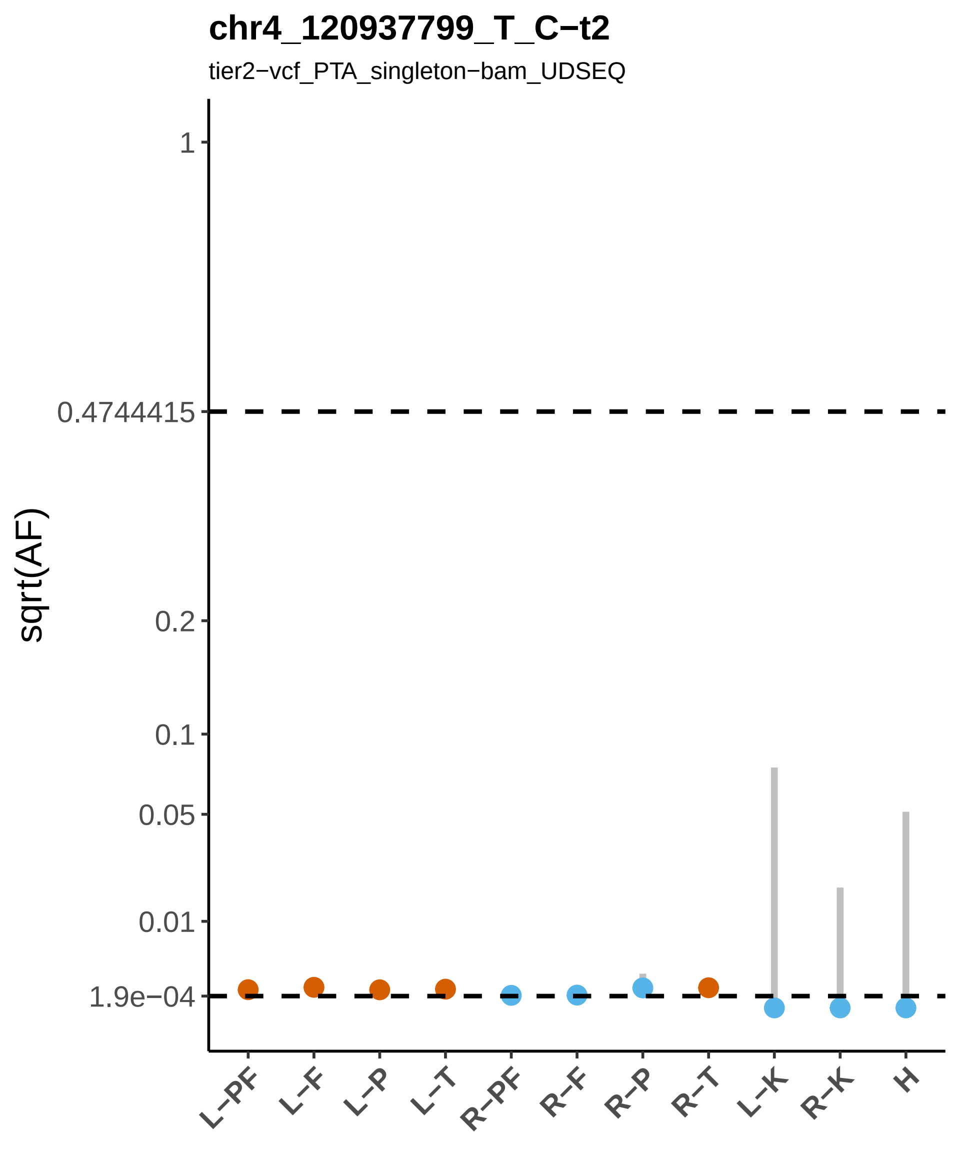
chr4_120937799_T_C−t2
tier2−vcf_PTA_singleton−bam_UDSEQ
sqrt(AF)
1
0.4744415
0.2
0.1
0.05
0.01
1.9e−04
L−PF
L−F
L−P
L−T
R−PF
R−F
R−P
R−T
L−K
R−K
H

#### chr4_132395095_C_T−t3

tier3−vcf_NanoSeq_singleton

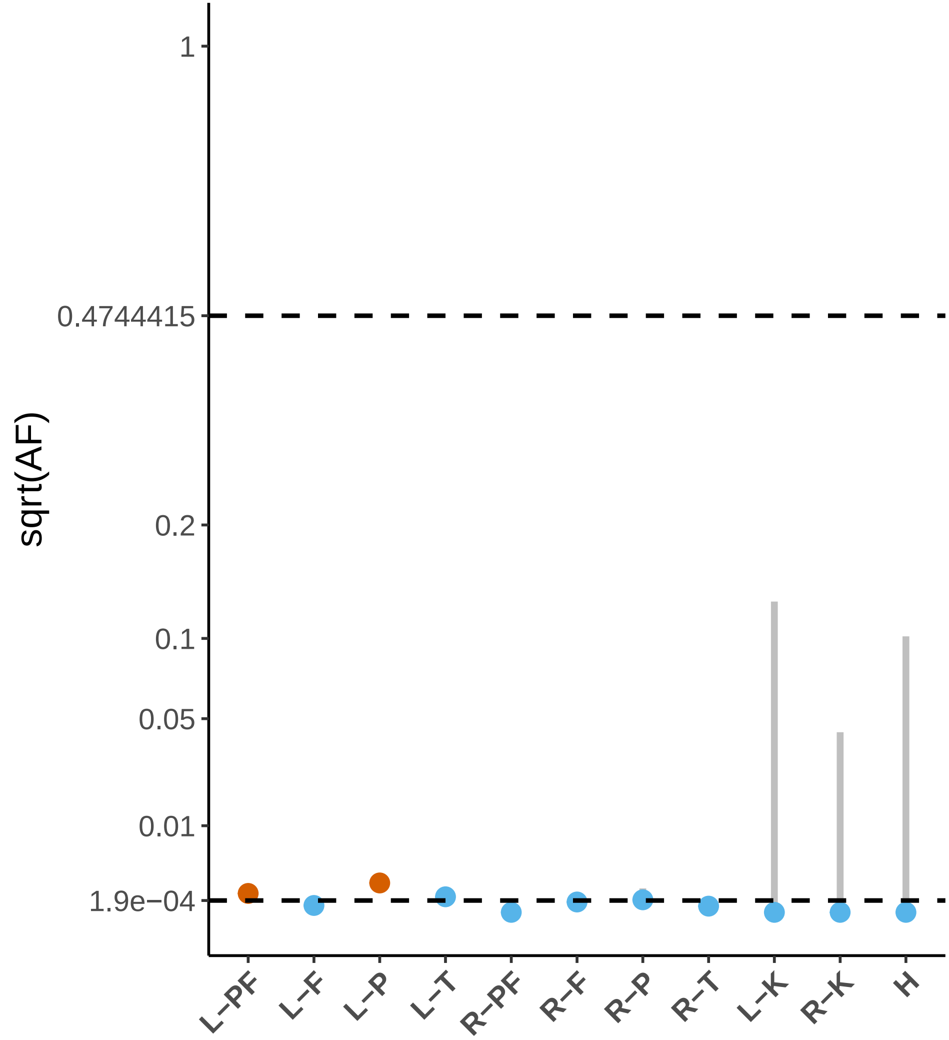

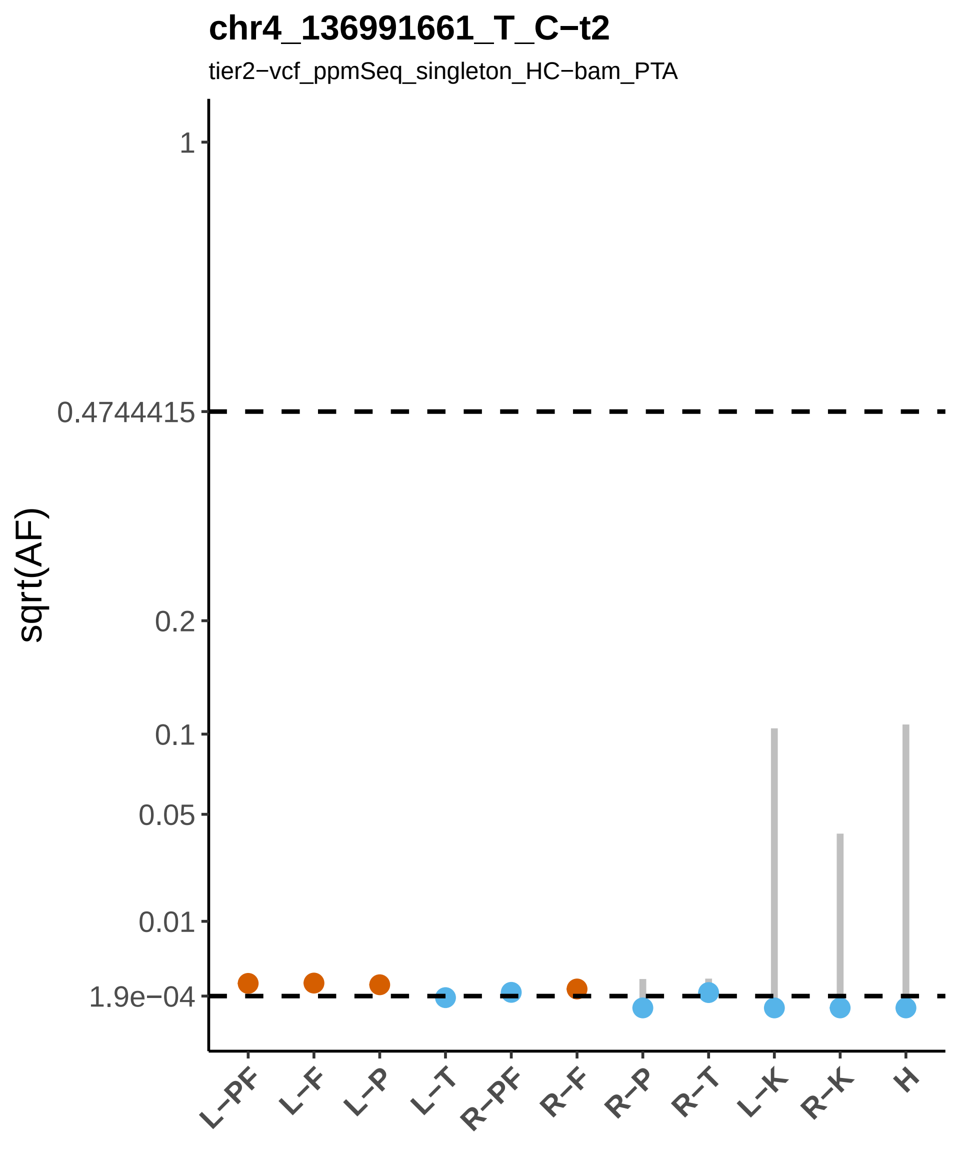

chr4_136991661_T_C−t2
tier2−vcf_ppmSeq_singleton_HC−bam_PTA
sqrt(AF)
1
0.4744415
0.2
0.1
0.05
0.01
1.9e−04
L−PF
L−F
L−P
L−T
R−PF
R−F
R−P
R−T
L−K
R−K
H

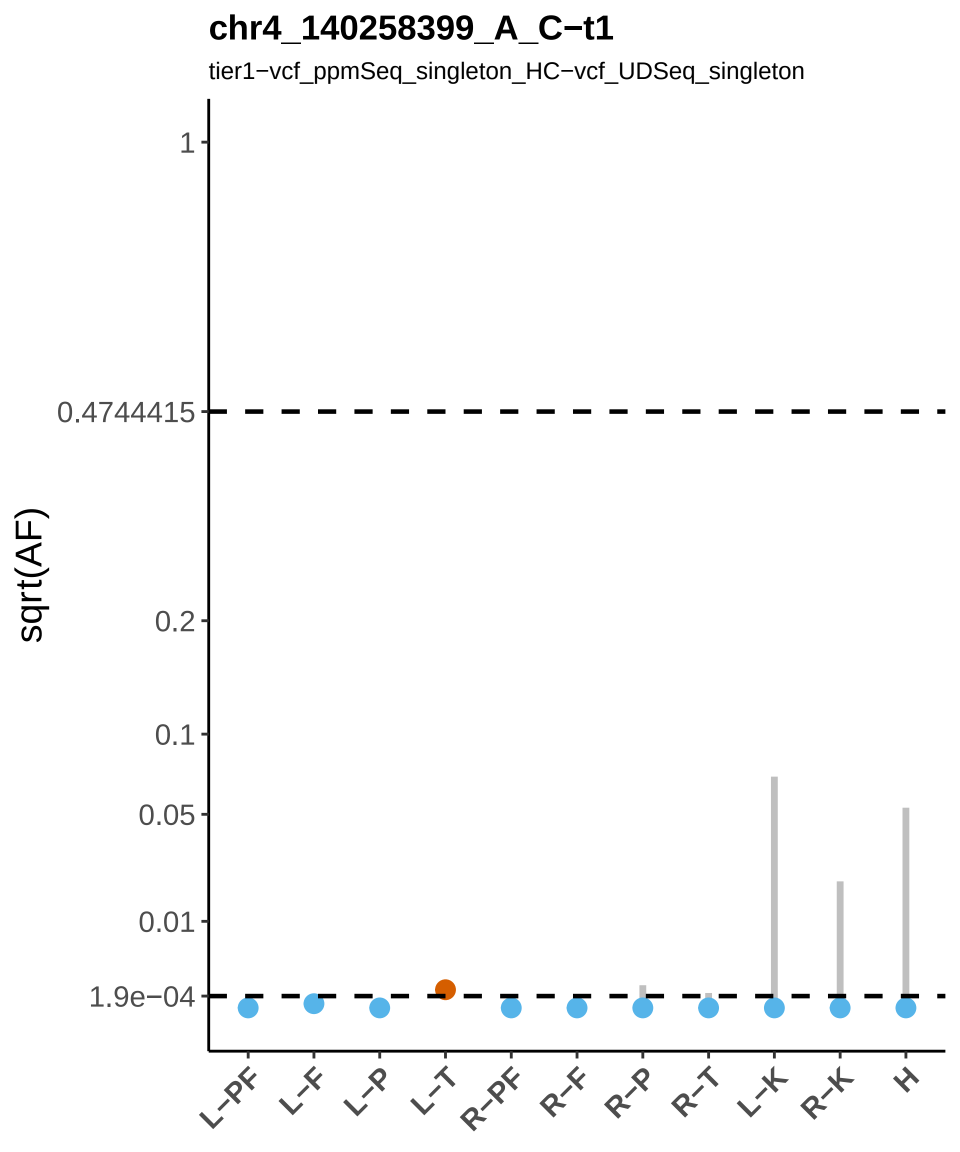

chr4_140258399_A_C−t1
tier1−vcf_ppmSeq_singleton_HC−vcf_UDSeq_singleton
sqrt(AF)
1
0.4744415
0.2
0.1
0.05
0.01
1.9e−04
L−PF
L−F
L−P
L−T
R−PF
R−F
R−P
R−T
L−K
R−K
H

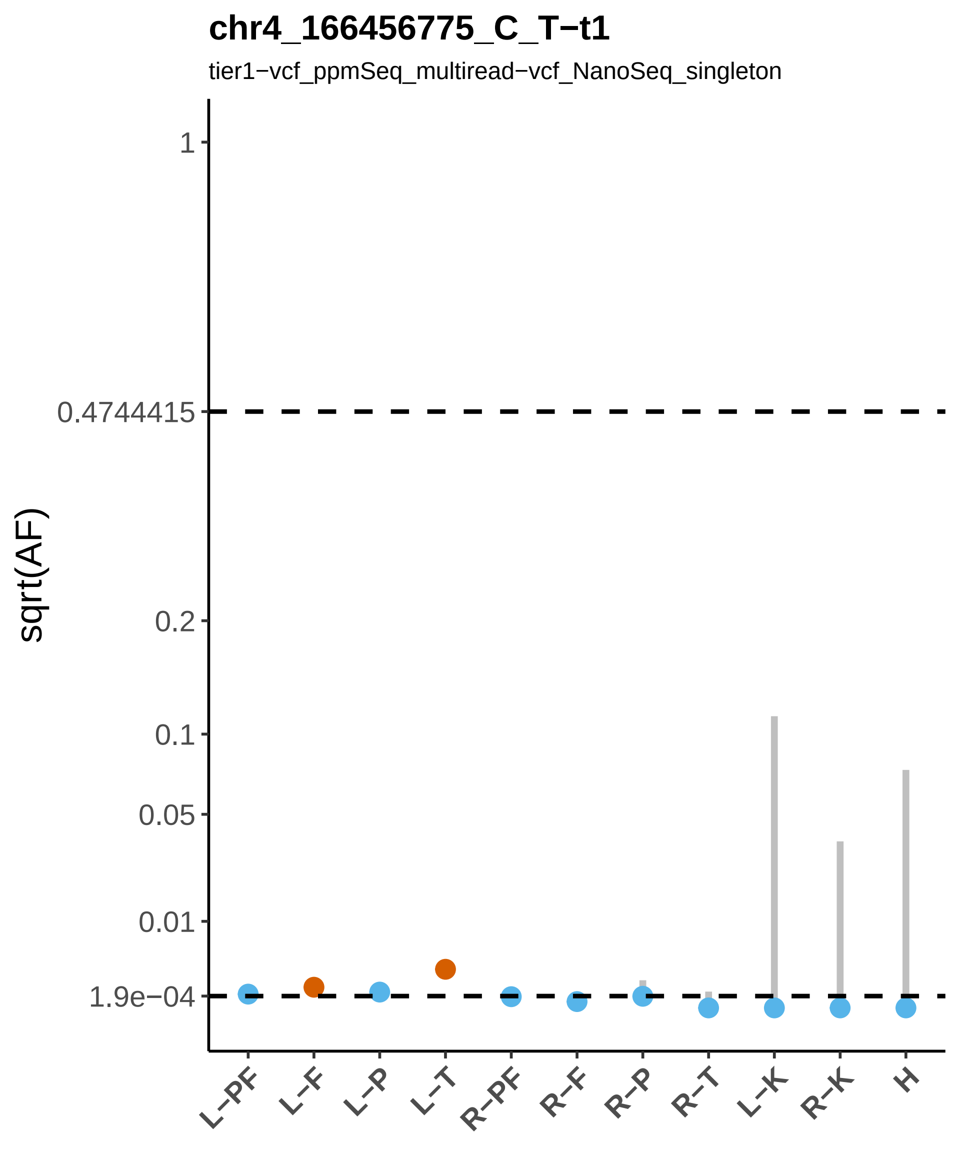
chr4_166456775_C_T−t1
tier1−vcf_ppmSeq_multiread−vcf_NanoSeq_singleton
sqrt(AF)
1
0.4744415
0.2
0.1
0.05
0.01
1.9e−04
L−PF
L−F
L−P
L−T
R−PF
R−F
R−P
R−T
L−K
R−K
H

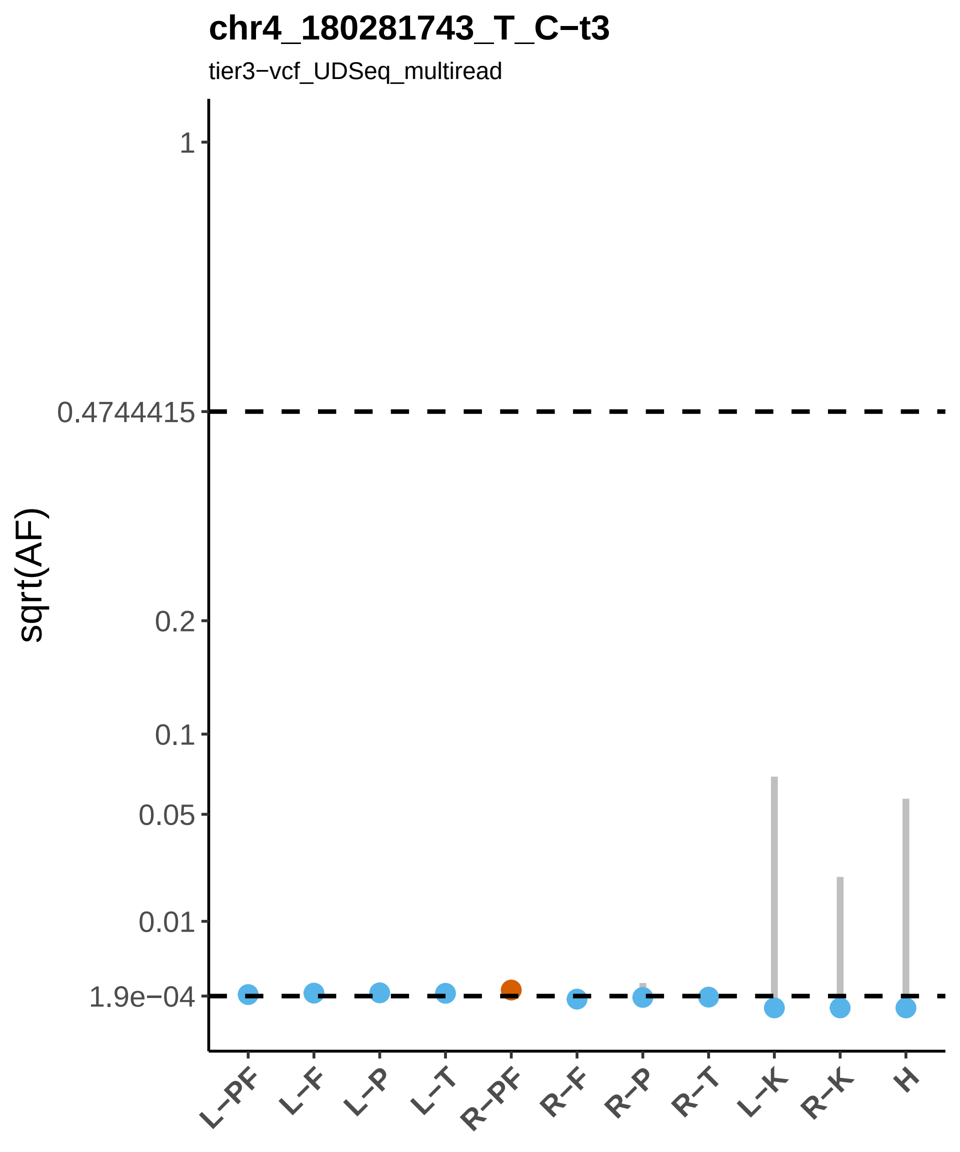
chr4_180281743_T_C−t3
tier3−vcf_UDSeq_multiread
sqrt(AF)
1
0.4744415
0.2
0.1
0.05
0.01
1.9e−04
L−PF
L−F
L−P
L−T
R−PF
R−F
R−P
R−T
L−K
R−K
H

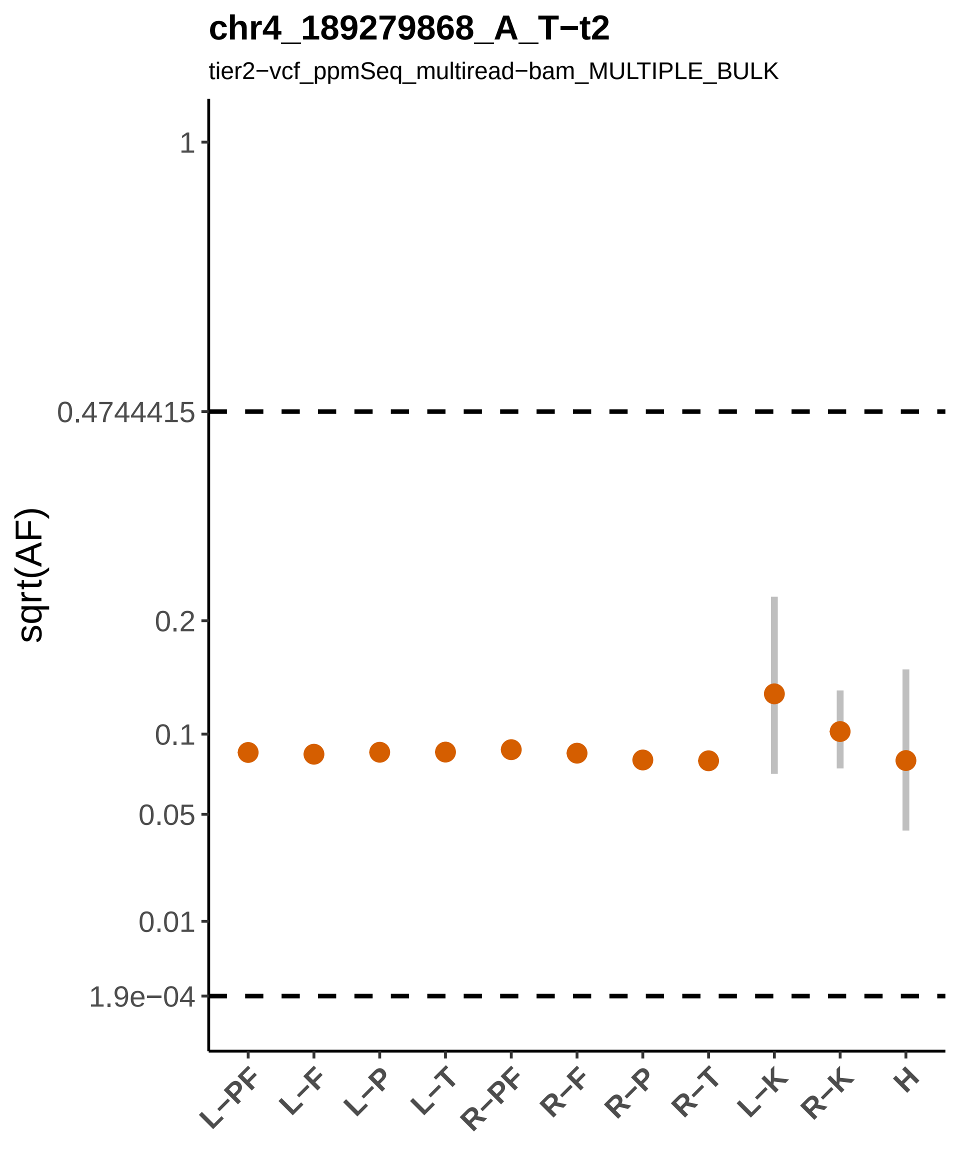
chr4_189279868_A_T−t2
tier2−vcf_ppmSeq_multiread−bam_MULTIPLE_BULK
sqrt(AF)
1
0.4744415
0.2
0.1
0.05
0.01
1.9e−04
L−PF
L−F
L−P
L−T
R−PF
R−F
R−P
R−T
L−K
R−K
H

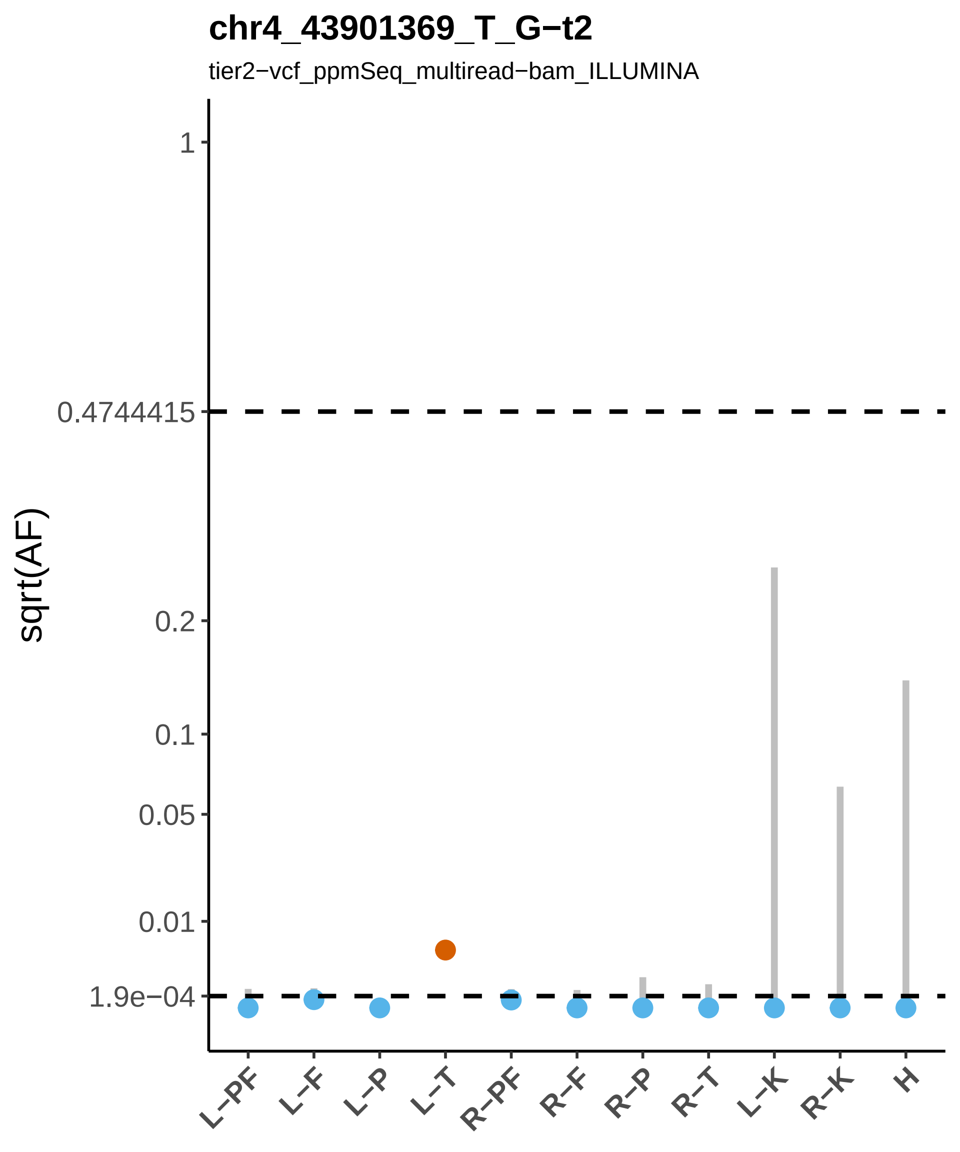

chr4_43901369_T_G−t2
tier2−vcf_ppmSeq_multiread−bam_ILLUMINA
sqrt(AF)
1
0.4744415
0.2
0.1
0.05
0.01
1.9e−04
L−PF
L−F
L−P
L−T
R−PF
R−F
R−P
R−T
L−K
R−K
H

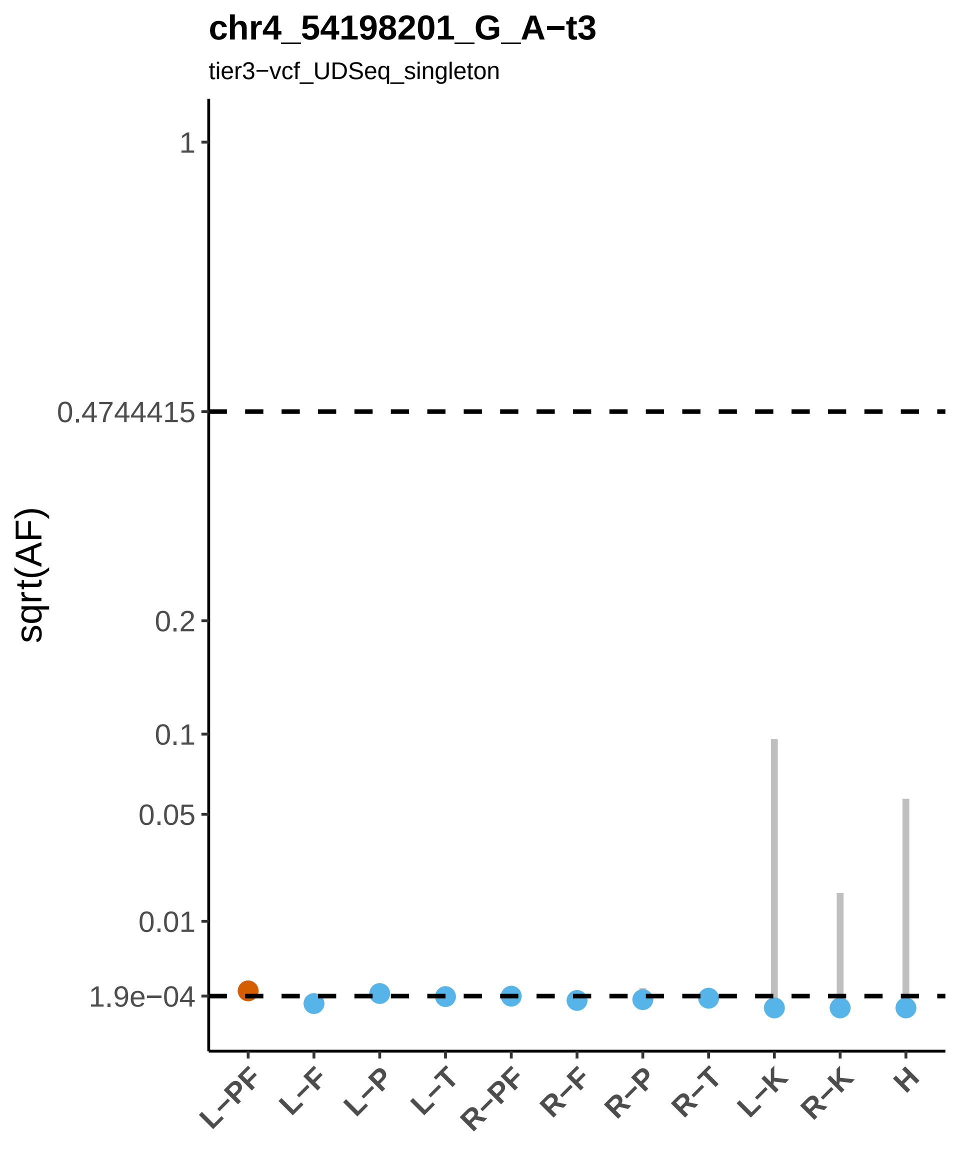
chr4_54198201_G_A−t3
tier3−vcf_UDSeq_singleton
sqrt(AF)
1
0.4744415
0.2
0.1
0.05
0.01
1.9e−04
L−PF
L−F
L−P
L−T
R−PF
R−F
R−P
R−T
L−K
R−K
H

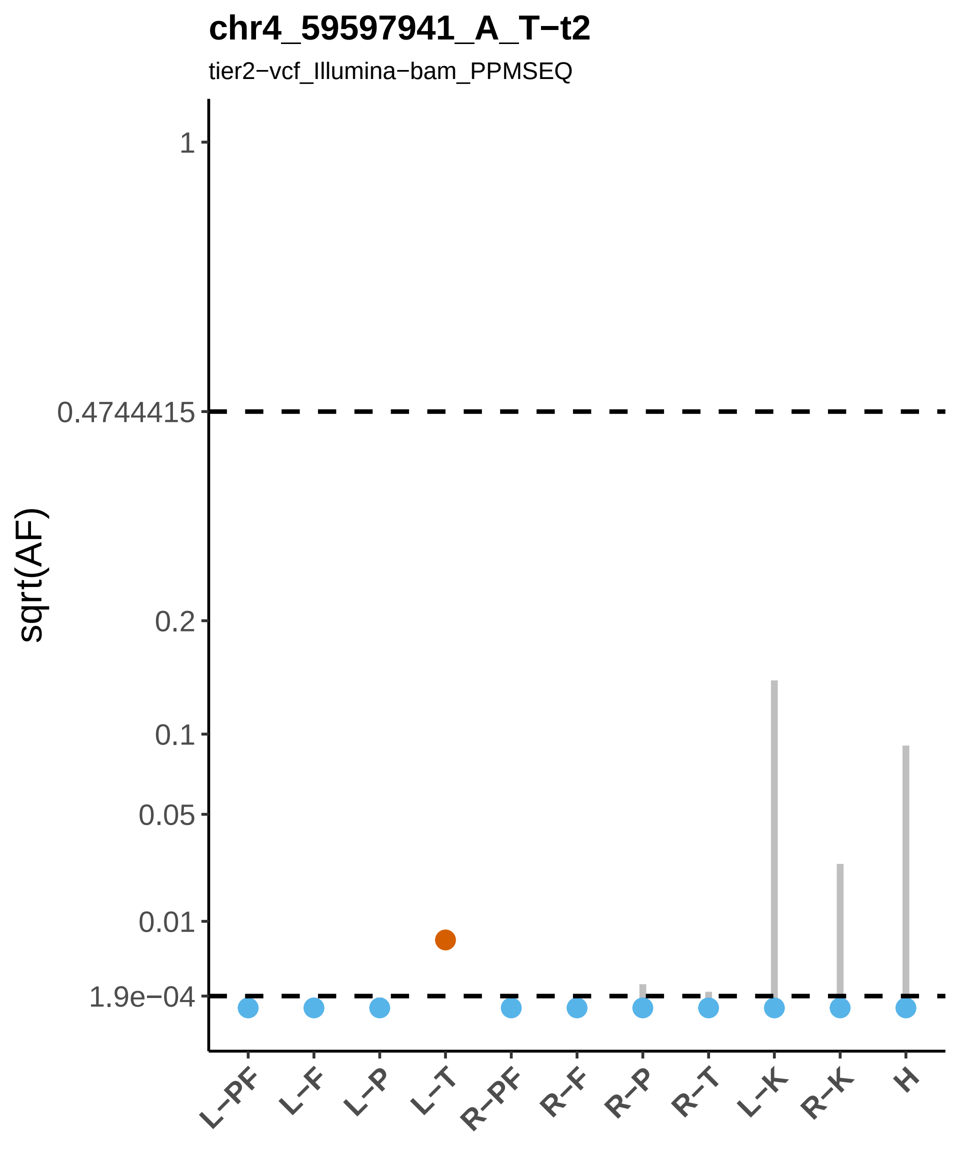

chr4_59597941_A_T−t2
tier2−vcf_Illumina−bam_PPMSEQ
sqrt(AF)
1
0.4744415
0.2
0.1
0.05
0.01
1.9e−04
L−PF
L−F
L−P
L−T
R−PF
R−F
R−P
R−T
L−K
R−K
H

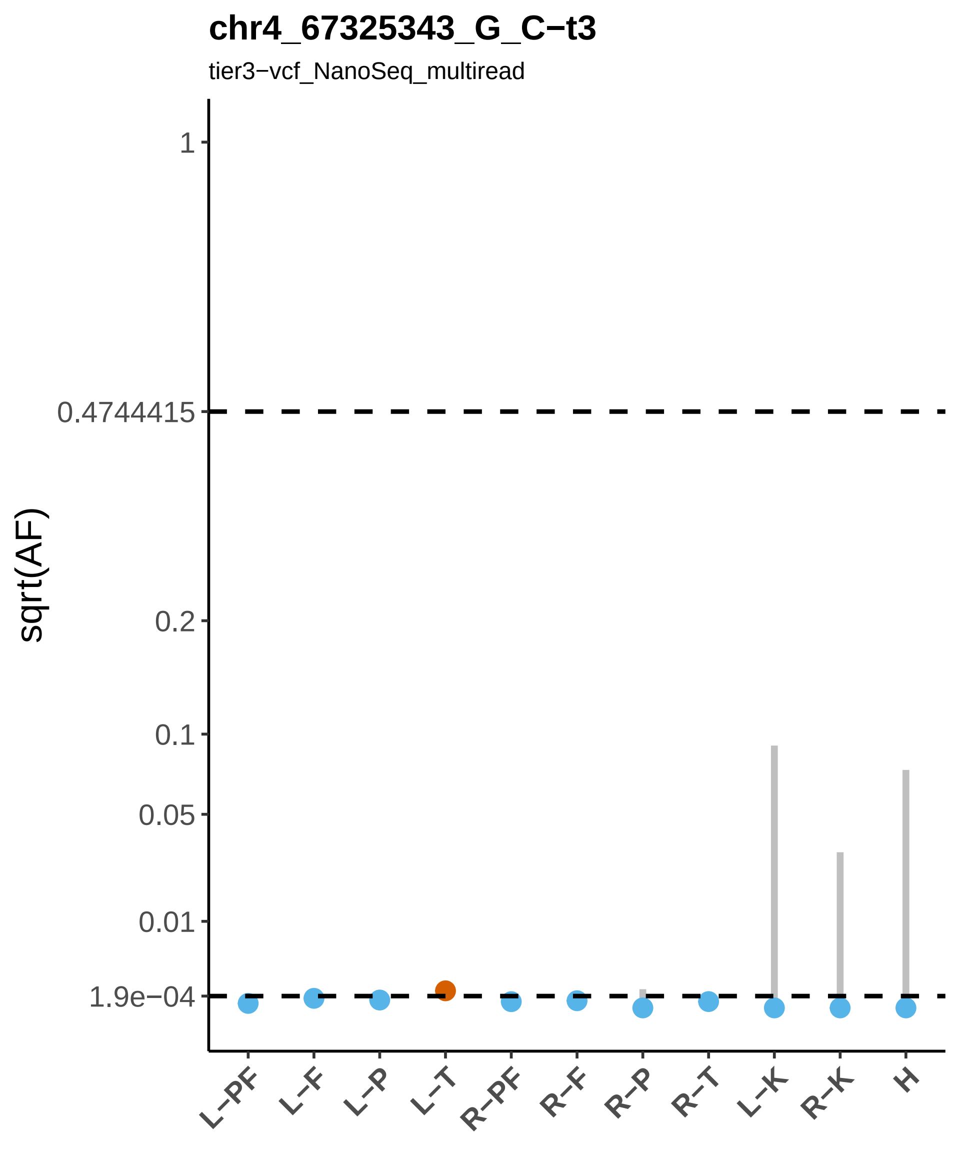
chr4_67325343_G_C−t3
tier3−vcf_NanoSeq_multiread
sqrt(AF)
1
0.4744415
0.2
0.1
0.05
0.01
1.9e−04
L−PF
L−F
L−P
L−T
R−PF
R−F
R−P
R−T
L−K
R−K
H

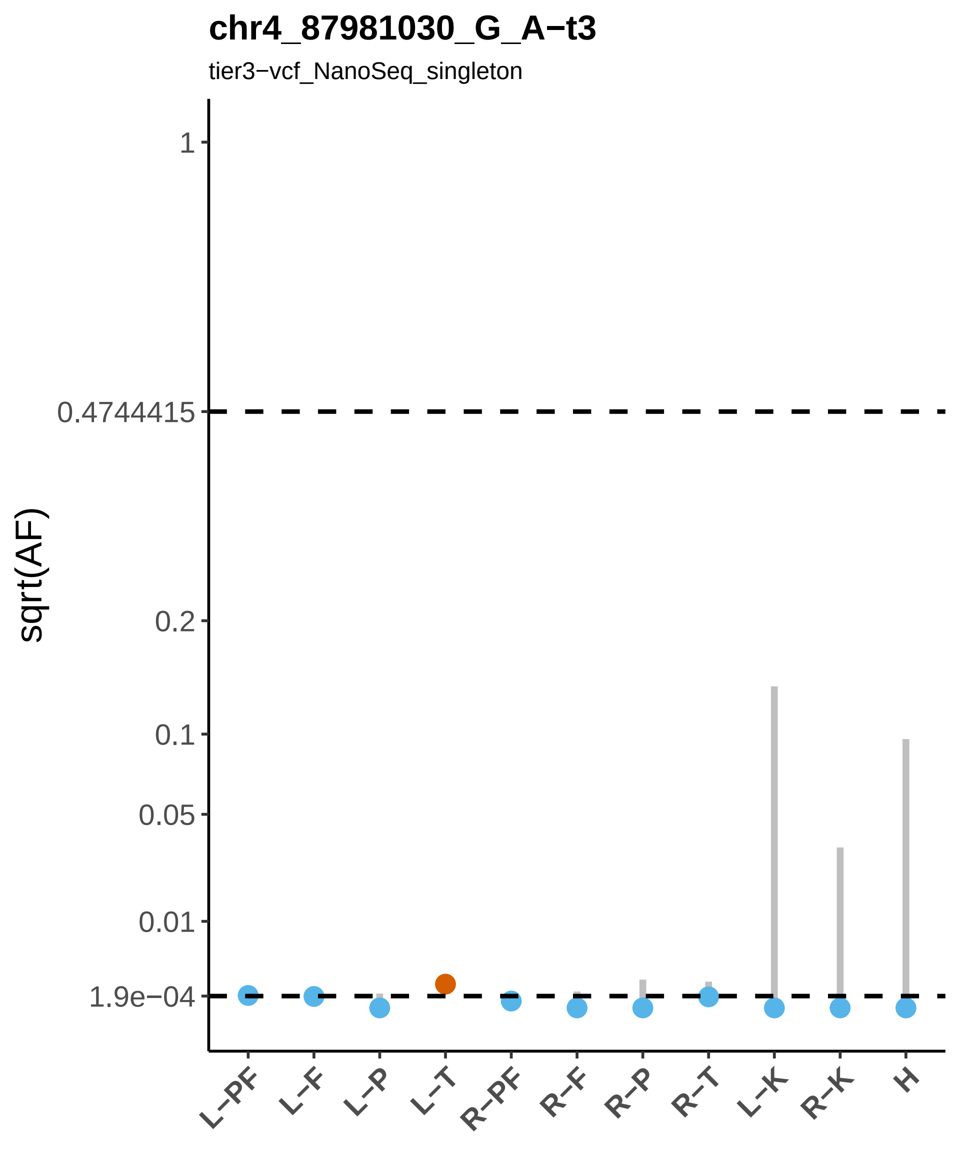

chr4_87981030_G_A−t3
tier3−vcf_NanoSeq_singleton
sqrt(AF)
1
0.4744415
0.2
0.1
0.05
0.01
1.9e−04
L−PF
L−F
L−P
L−T
R−PF
R−F
R−P
R−T
L−K
R−K
H

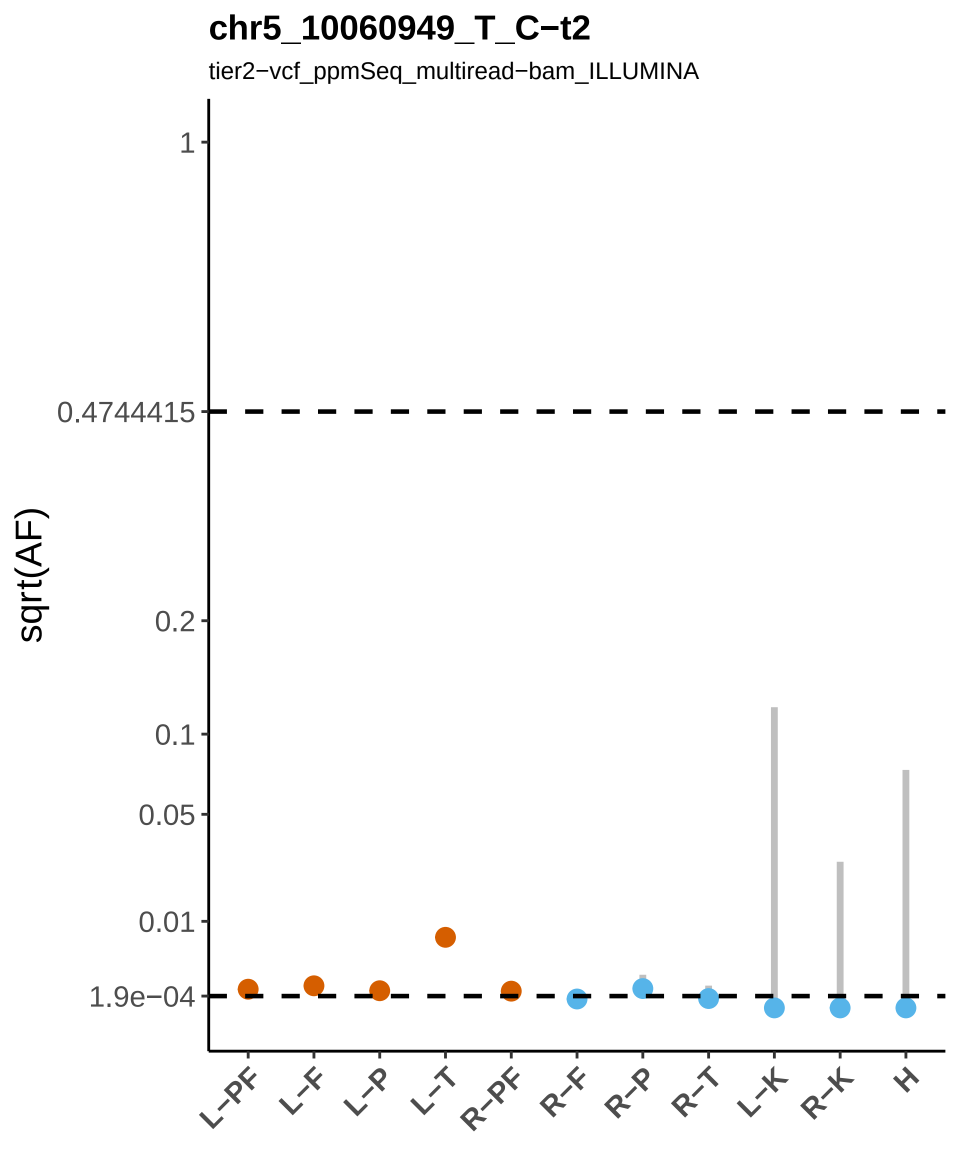
chr5_10060949_T_C−t2
tier2−vcf_ppmSeq_multiread−bam_ILLUMINA
sqrt(AF)
1
0.4744415
0.2
0.1
0.05
0.01
1.9e−04
L−PF
L−F
L−P
L−T
R−PF
R−F
R−P
R−T
L−K
R−K
H

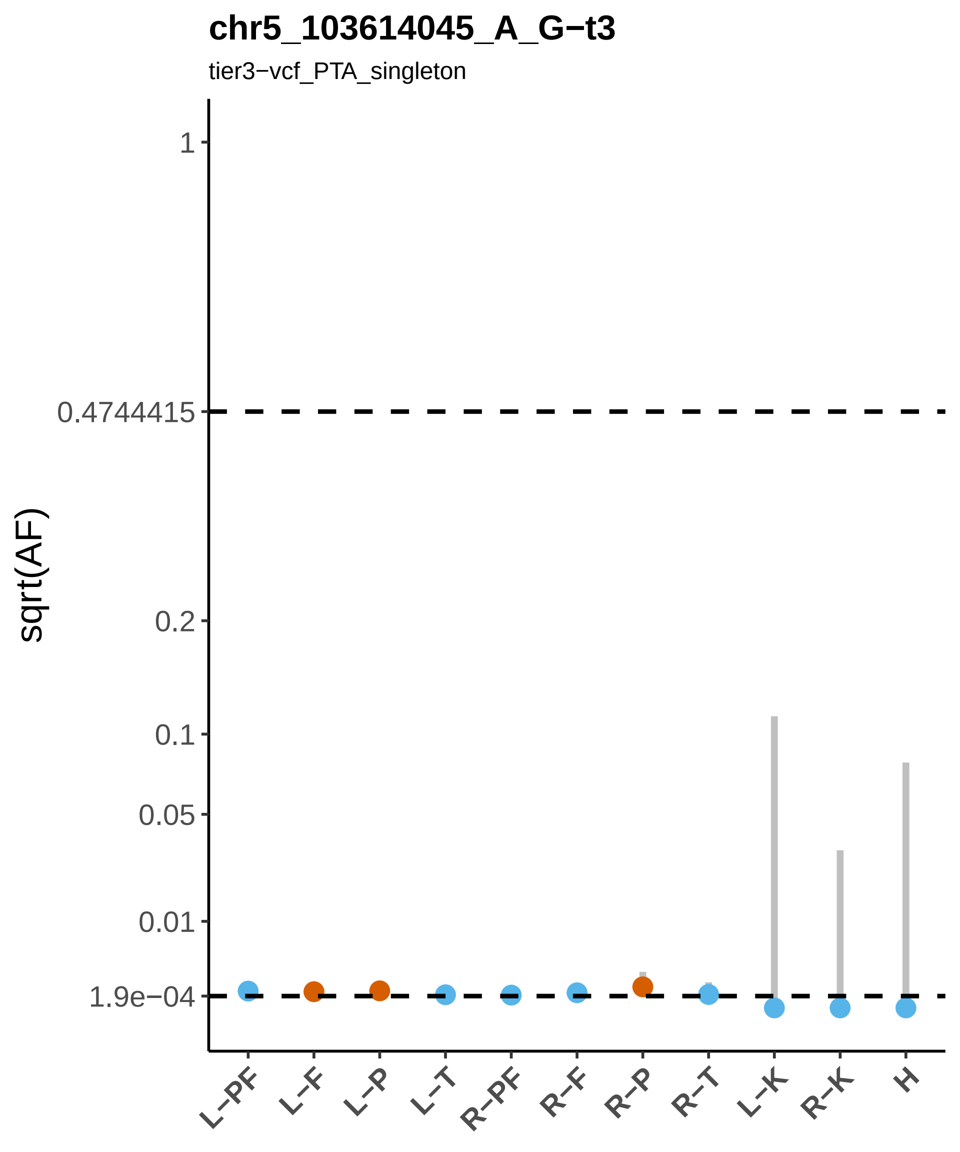
chr5_103614045_A_G−t3
tier3−vcf_PTA_singleton
sqrt(AF)
1
0.4744415
0.2
0.1
0.05
0.01
1.9e−04
L−PF
L−F
L−P
L−T
R−PF
R−F
R−P
R−T
L−K
R−K
H

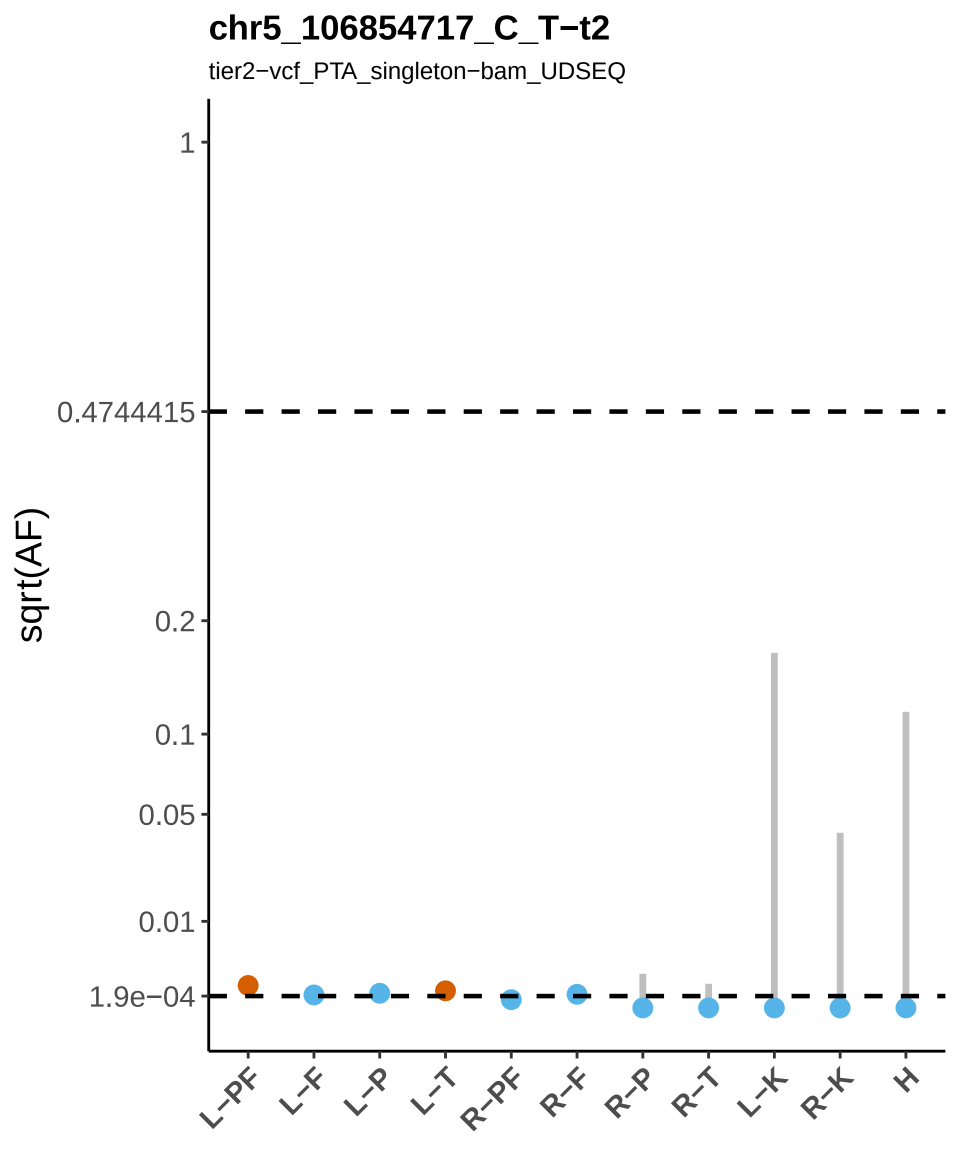
chr5_106854717_C_T−t2
tier2−vcf_PTA_singleton−bam_UDSEQ
sqrt(AF)
1
0.4744415
0.2
0.1
0.05
0.01
1.9e−04
L−PF
L−F
L−P
L−T
R−PF
R−F
R−P
R−T
L−K
R−K
H

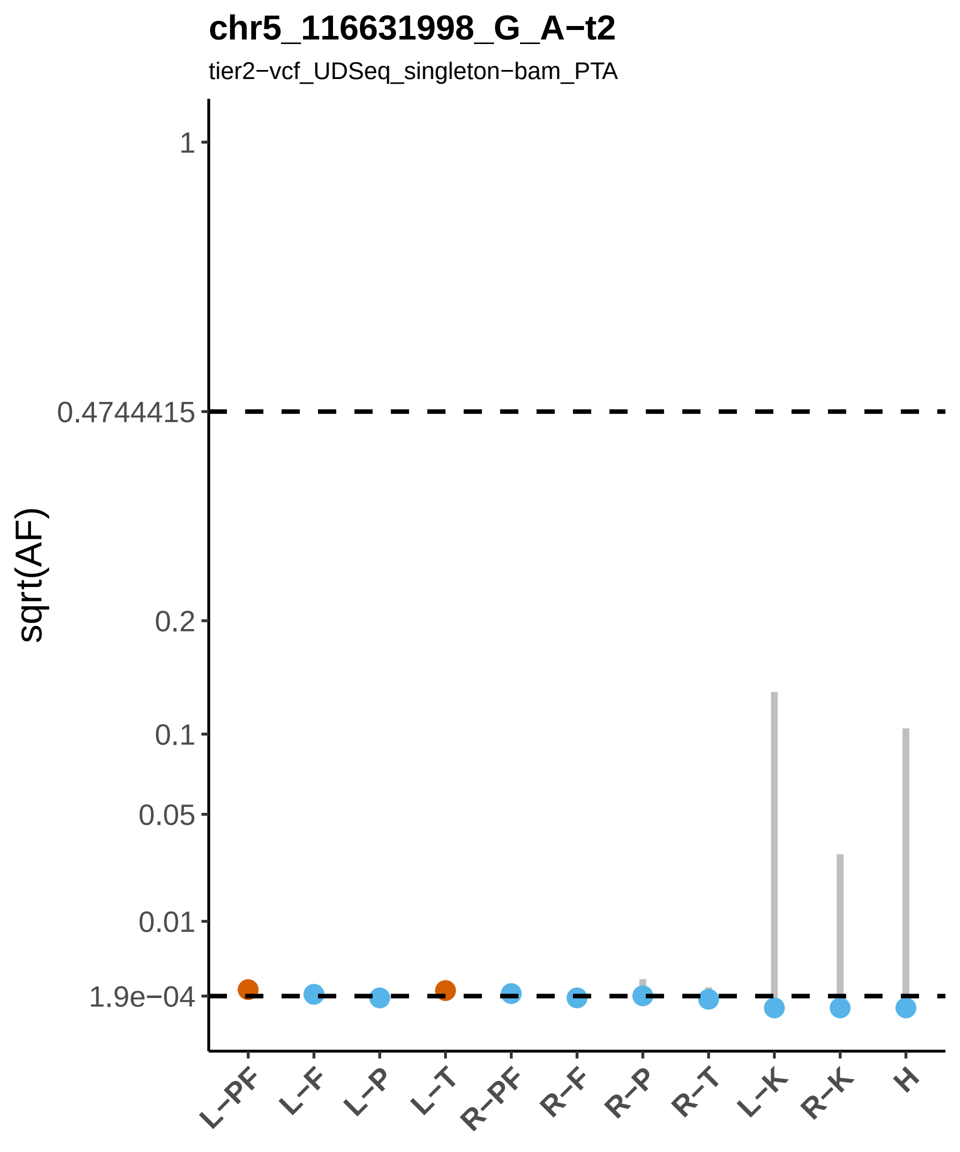

chr5_116631998_G_A−t2
tier2−vcf_UDSeq_singleton−bam_PTA
sqrt(AF)
1
0.4744415
0.2
0.1
0.05
0.01
1.9e−04
L−PF
L−F
L−P
L−T
R−PF
R−F
R−P
R−T
L−K
R−K
H

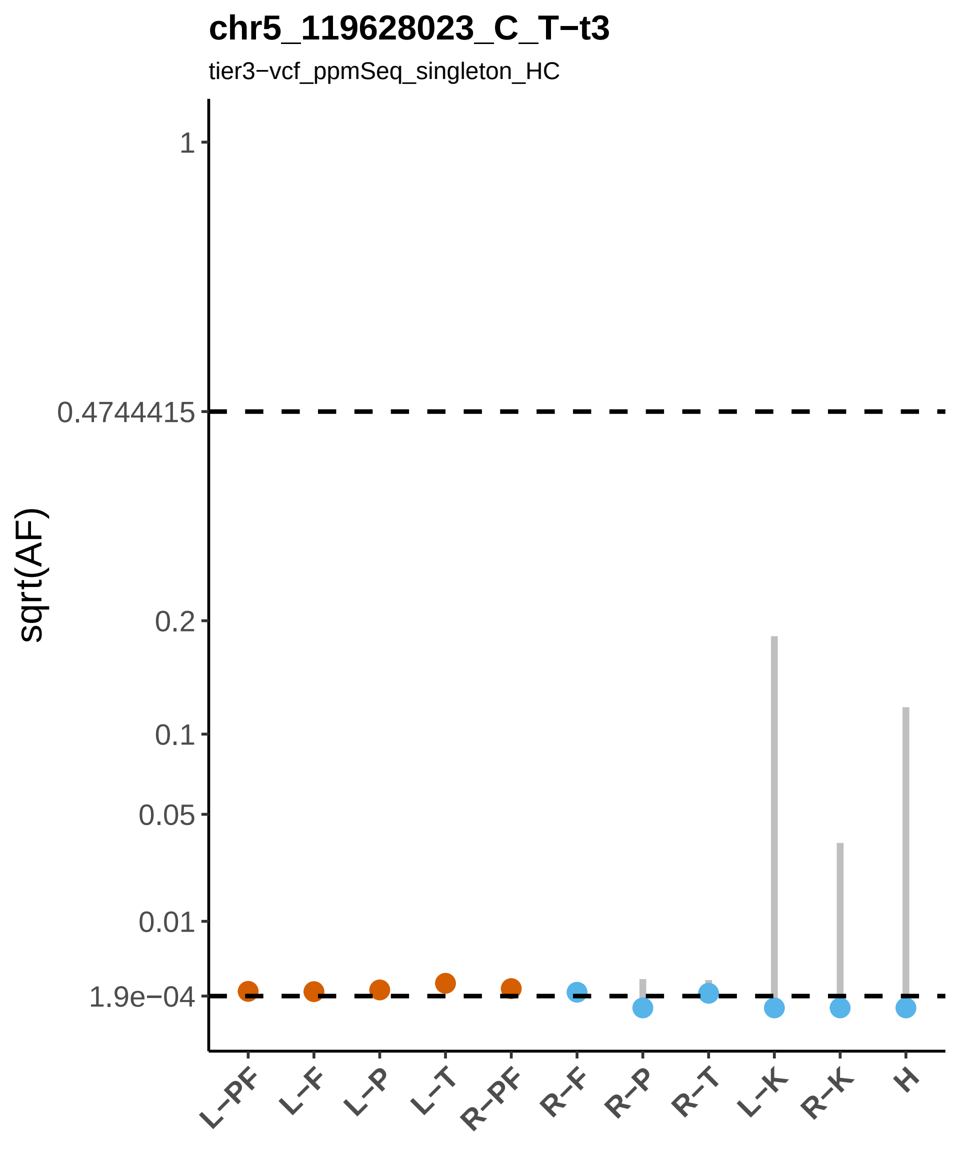

chr5_119628023_C_T−t3
tier3−vcf_ppmSeq_singleton_HC
sqrt(AF)
1
0.4744415
0.2
0.1
0.05
0.01
1.9e−04
L−PF
L−F
L−P
L−T
R−PF
R−F
R−P
R−T
L−K
R−K
H

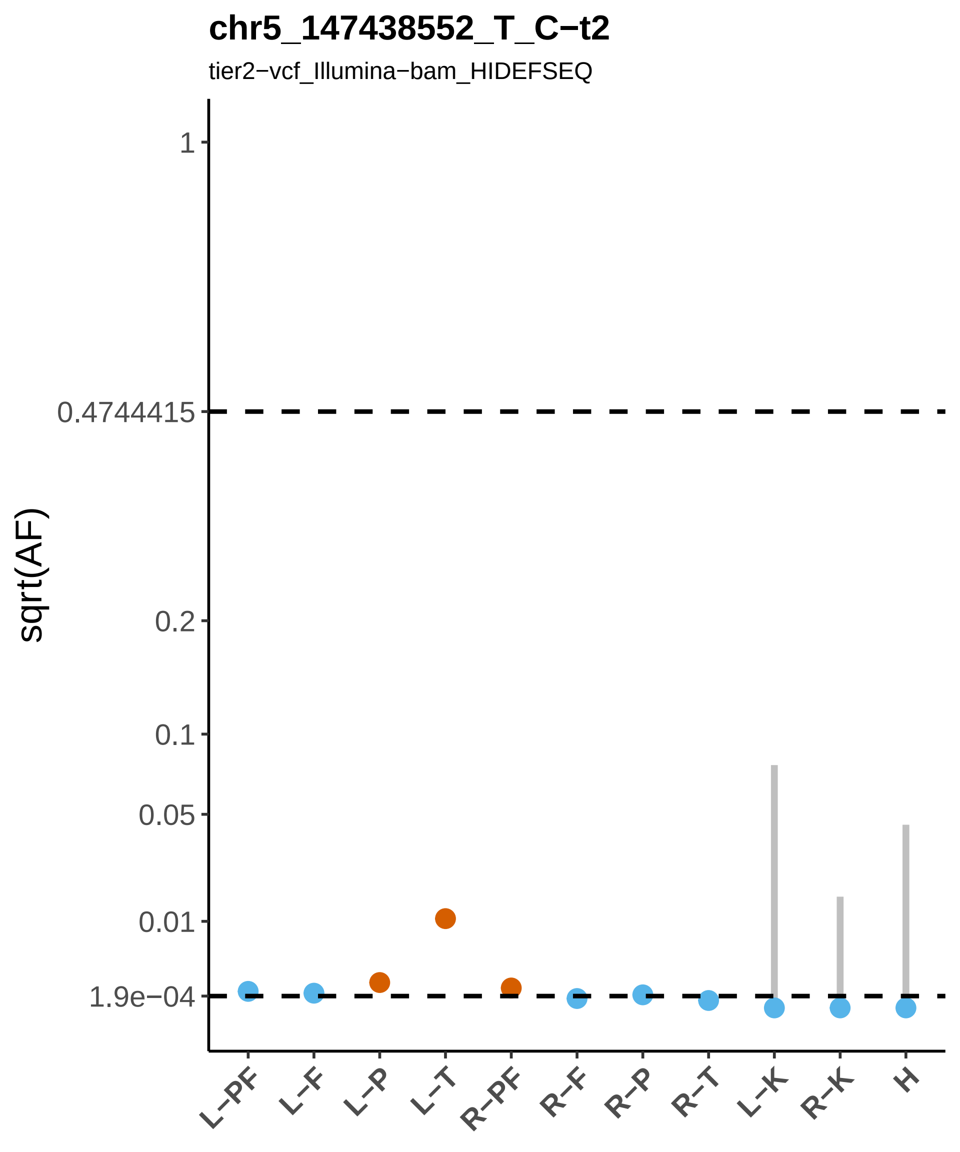

chr5_147438552_T_C−t2
tier2−vcf_Illumina−bam_HIDEFSEQ
sqrt(AF)
1
0.4744415
0.2
0.1
0.05
0.01
1.9e−04
L−PF
L−F
L−P
L−T
R−PF
R−F
R−P
R−T
L−K
R−K
H

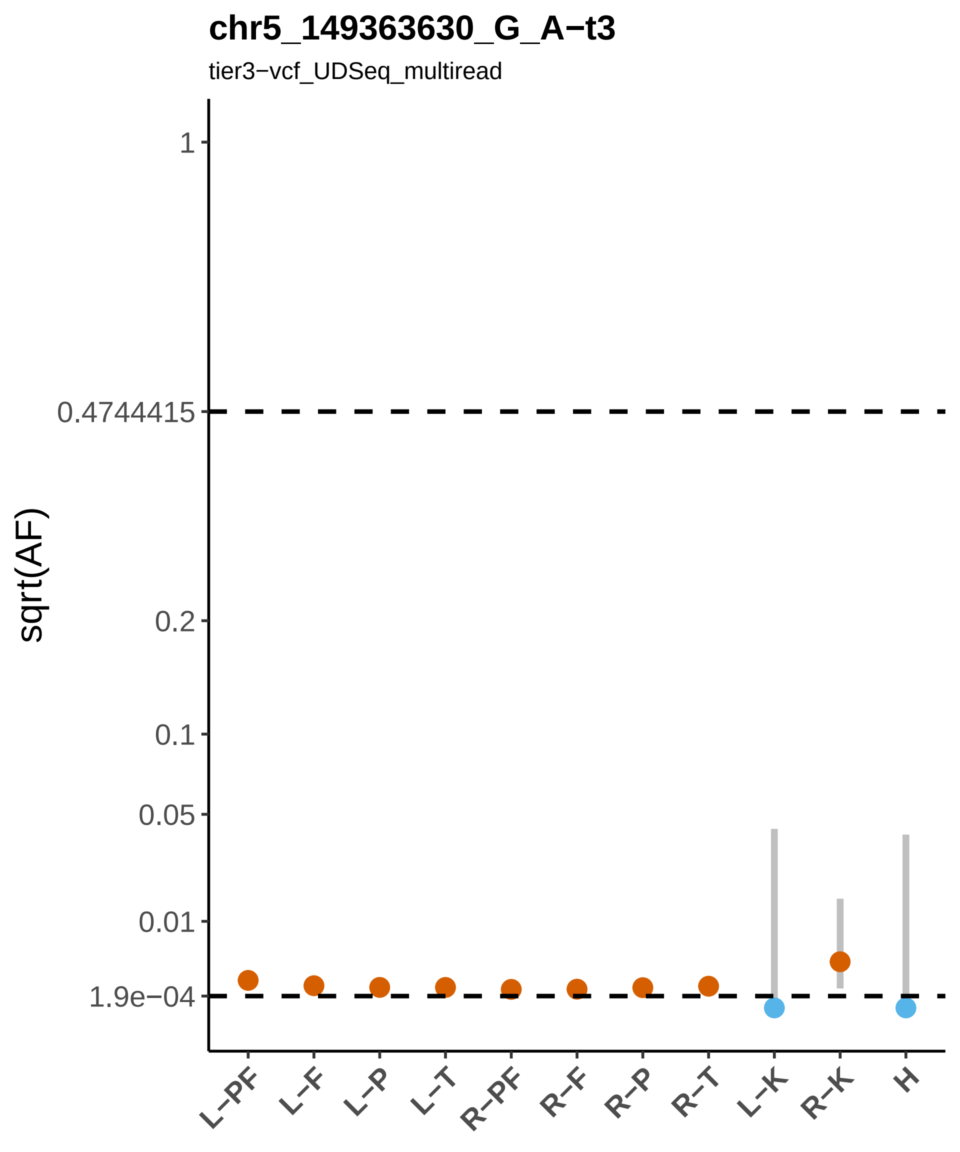
chr5_149363630_G_A−t3
tier3−vcf_UDSeq_multiread
sqrt(AF)
1
0.4744415
0.2
0.1
0.05
0.01
1.9e−04
L−PF
L−F
L−P
L−T
R−PF
R−F
R−P
R−T
L−K
R−K
H

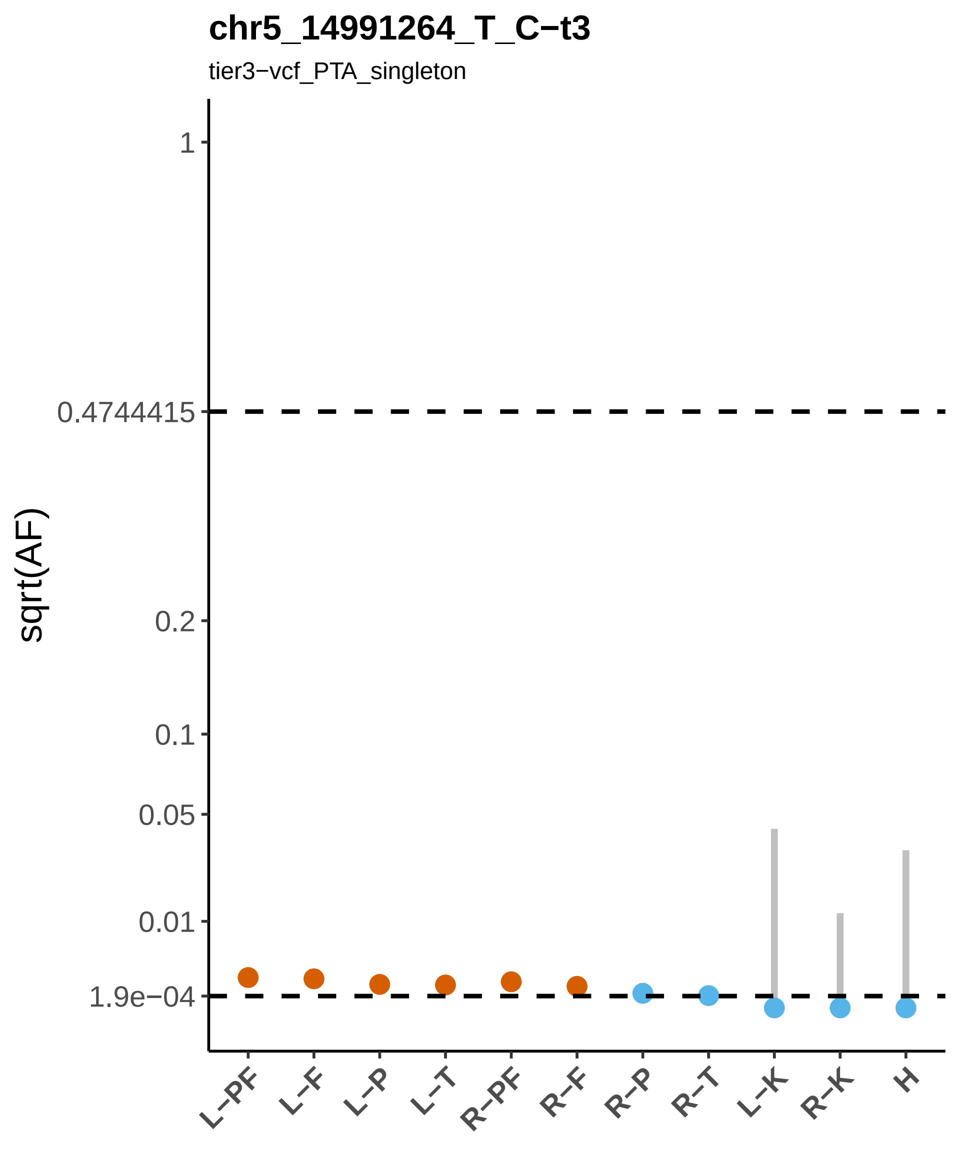
chr5_14991264_T_C−t3
tier3−vcf_PTA_singleton
sqrt(AF)
1
0.4744415
0.2
0.1
0.05
0.01
1.9e−04
L−PF
L−F
L−P
L−T
R−PF
R−F
R−P
R−T
L−K
R−K
H

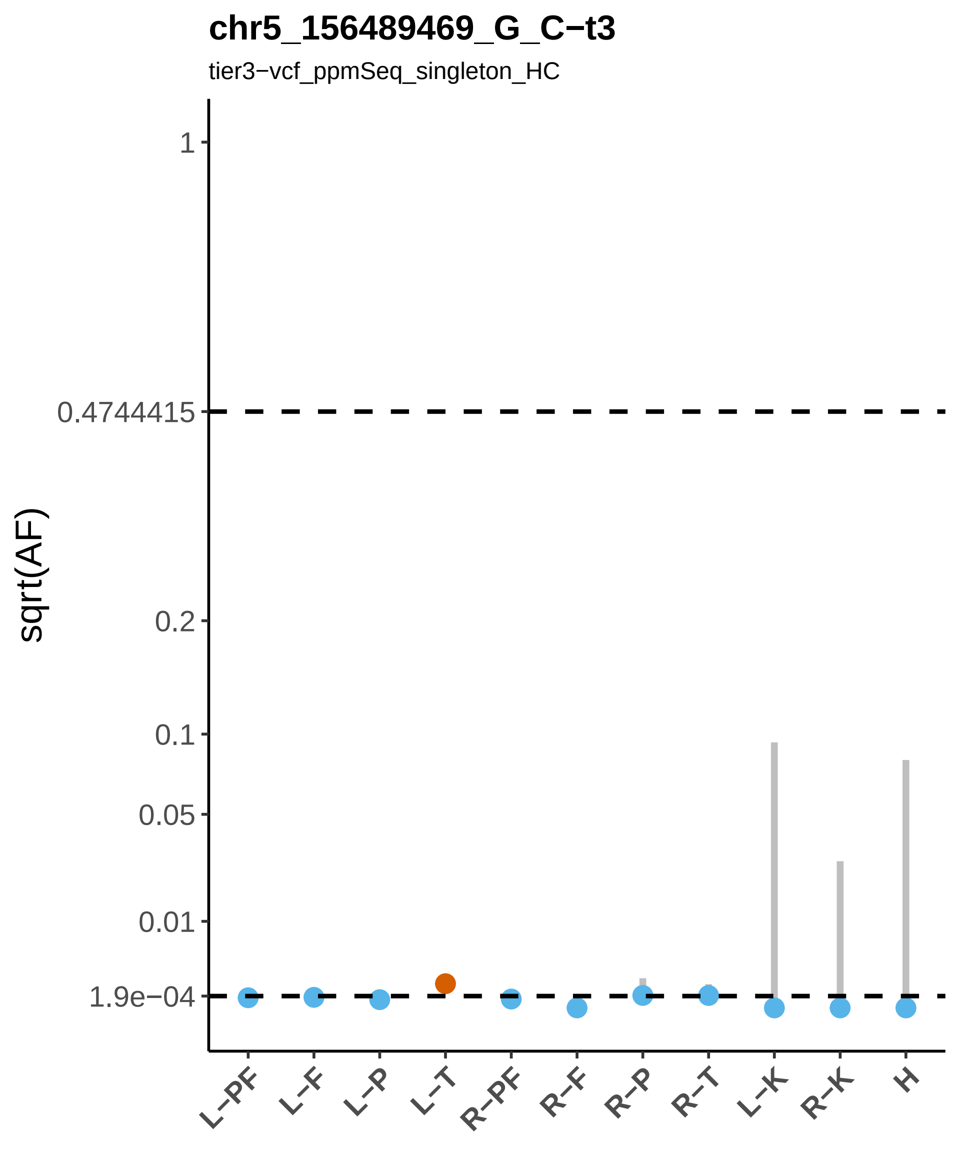

chr5_156489469_G_C−t3
tier3−vcf_ppmSeq_singleton_HC
sqrt(AF)
1
0.4744415
0.2
0.1
0.05
0.01
1.9e−04
L−PF
L−F
L−P
L−T
R−PF
R−F
R−P
R−T
L−K
R−K
H

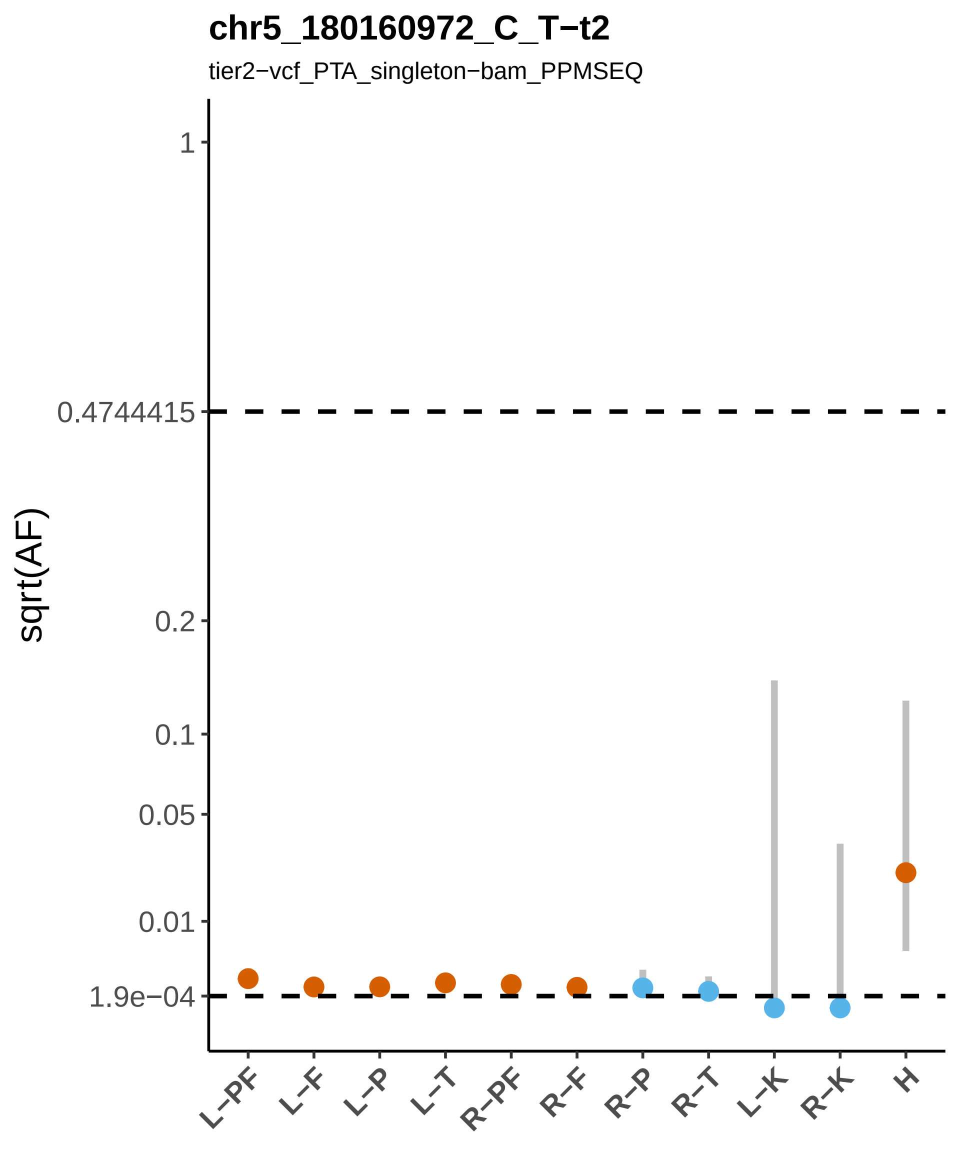

chr5_180160972_C_T−t2
tier2−vcf_PTA_singleton−bam_PPMSEQ
sqrt(AF)
1
0.4744415
0.2
0.1
0.05
0.01
1.9e−04
L−PF
L−F
L−P
L−T
R−PF
R−F
R−P
R−T
L−K
R−K
H

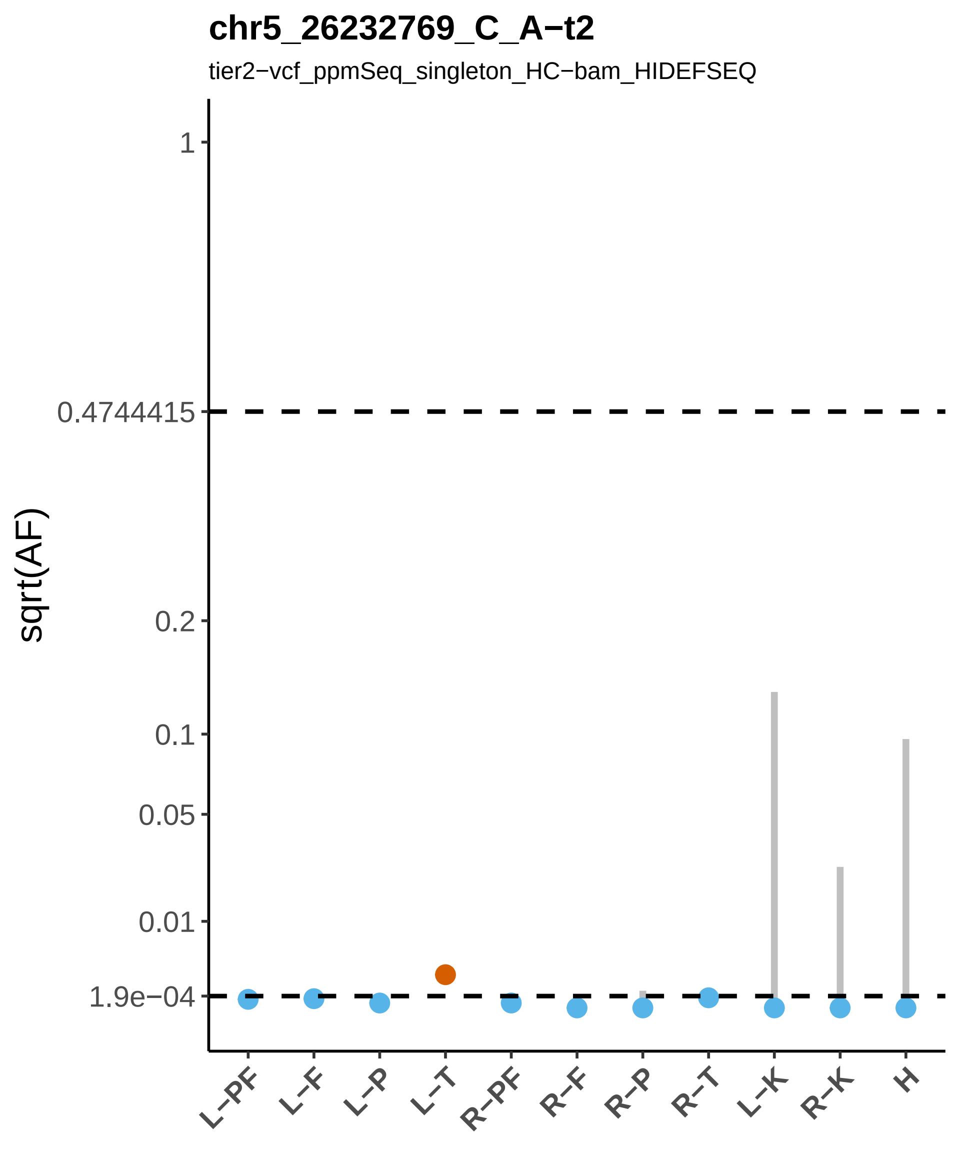

chr5_26232769_C_A−t2
tier2−vcf_ppmSeq_singleton_HC−bam_HIDEFSEQ
sqrt(AF)
1
0.4744415
0.2
0.1
0.05
0.01
1.9e−04
L−PF
L−F
L−P
L−T
R−PF
R−F
R−P
R−T
L−K
R−K
H

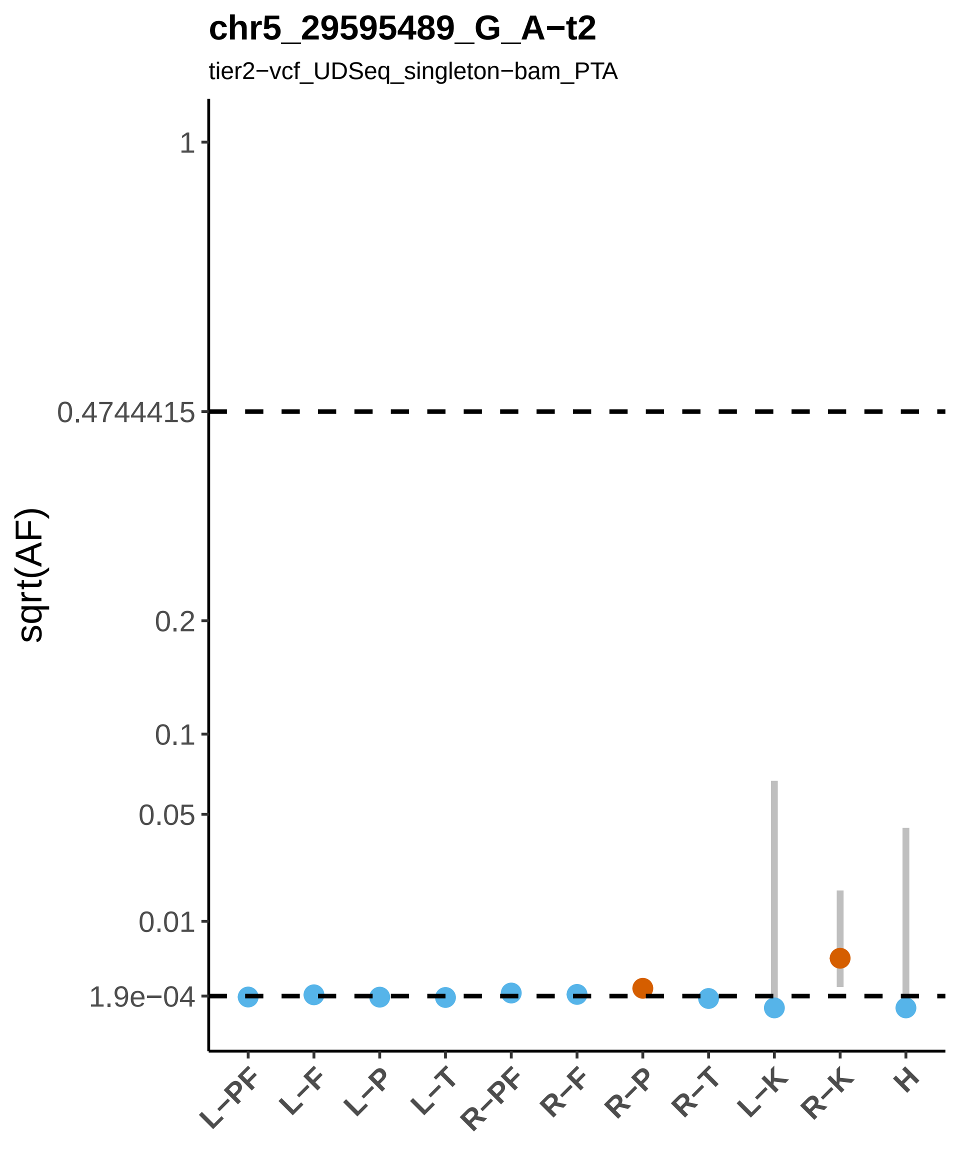

chr5_29595489_G_A−t2
tier2−vcf_UDSeq_singleton−bam_PTA
sqrt(AF)
1
0.4744415
0.2
0.1
0.05
0.01
1.9e−04
L−PF
L−F
L−P
L−T
R−PF
R−F
R−P
R−T
L−K
R−K
H

### chr5_36652430_T_C−t3

tier3−vcf_PTA_singleton

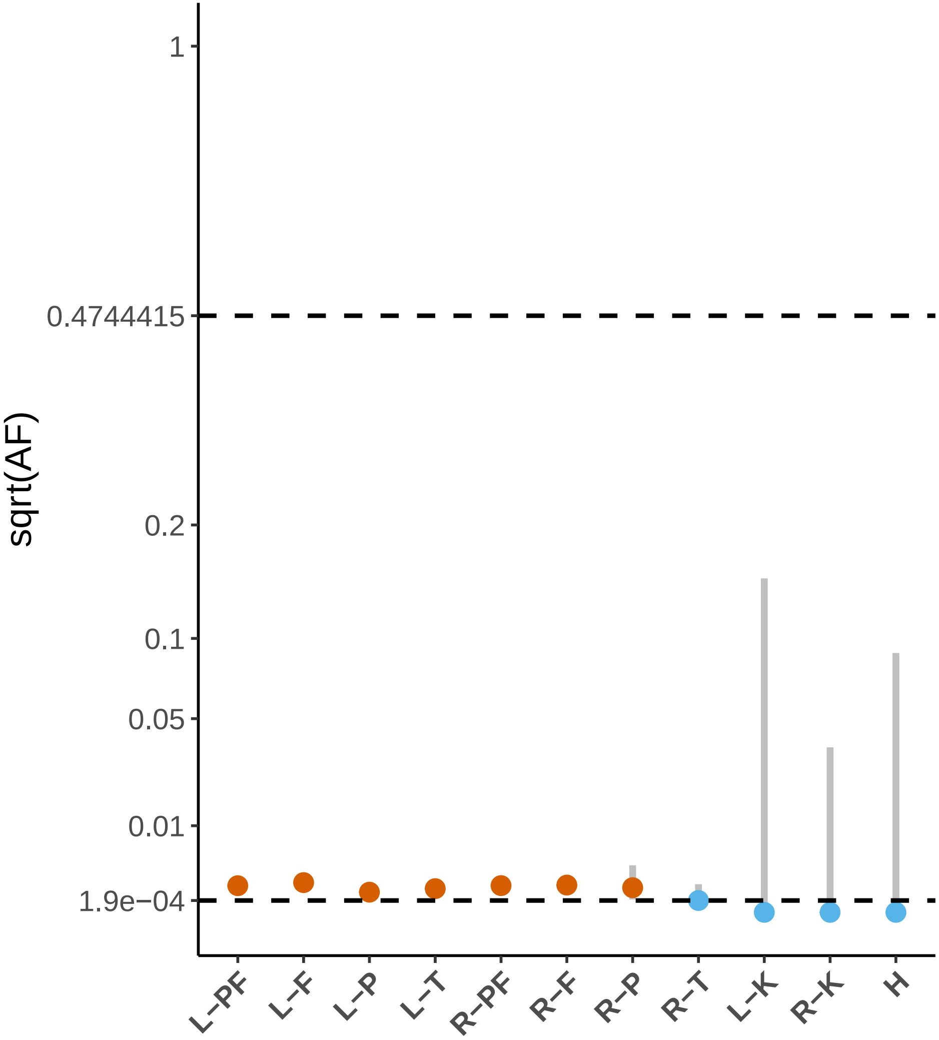

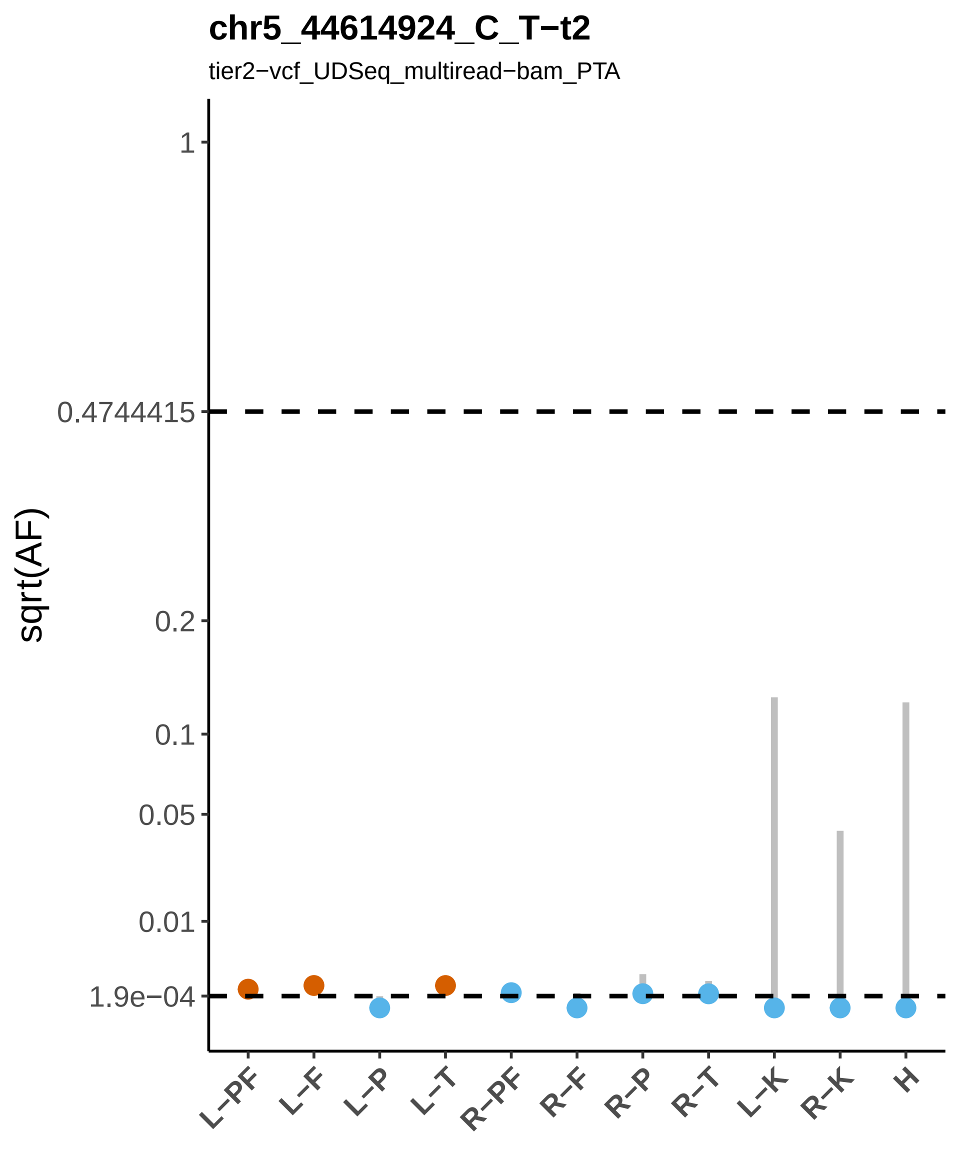

chr5_44614924_C_T−t2
tier2−vcf_UDSeq_multiread−bam_PTA
sqrt(AF)
1
0.4744415
0.2
0.1
0.05
0.01
1.9e−04
L−PF
L−F
L−P
L−T
R−PF
R−F
R−P
R−T
L−K
R−K
H

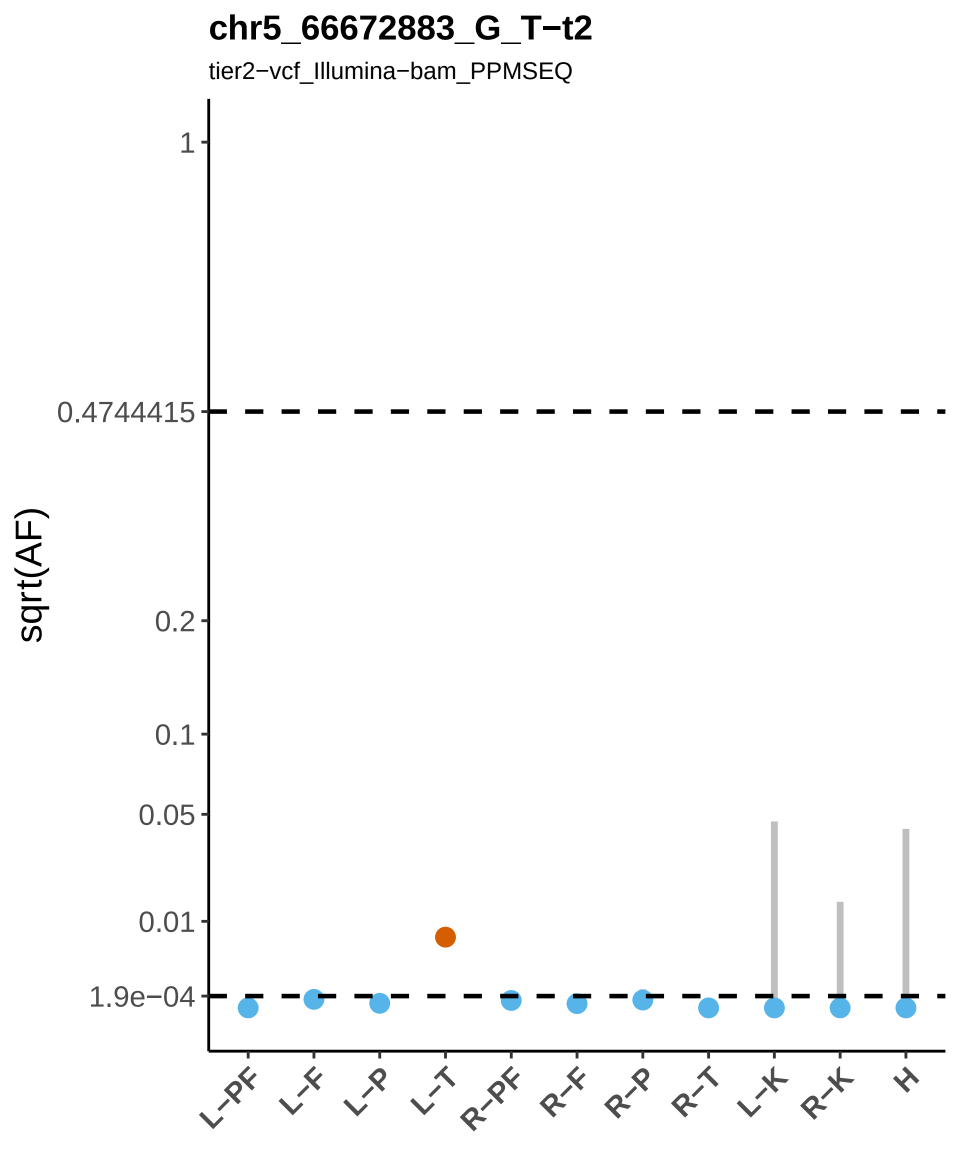
chr5_66672883_G_T−t2
tier2−vcf_Illumina−bam_PPMSEQ
sqrt(AF)
1
0.4744415
0.2
0.1
0.05
0.01
1.9e−04
L−PF
L−F
L−P
L−T
R−PF
R−F
R−P
R−T
L−K
R−K
H

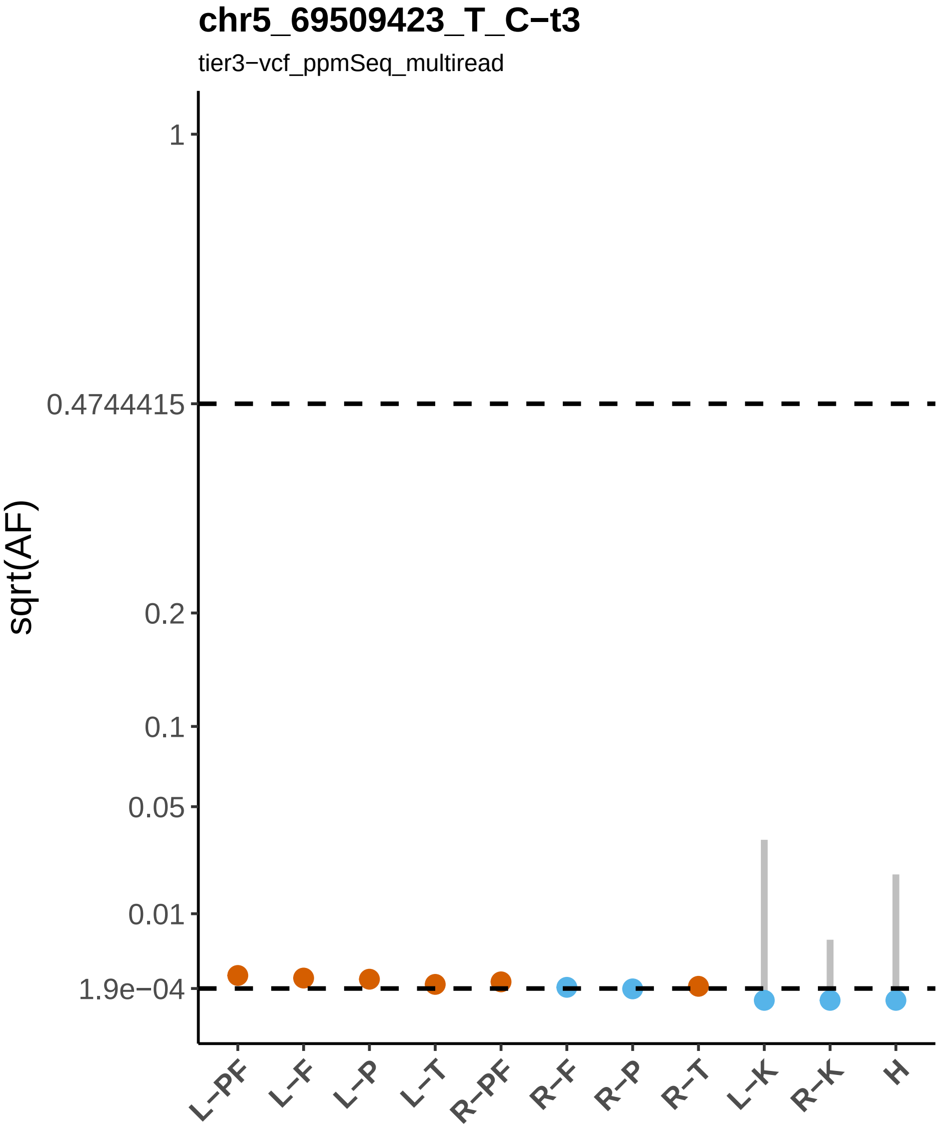
chr5_69509423_T_C−t3
tier3−vcf_ppmSeq_multiread
sqrt(AF)
1
0.4744415
0.2
0.1
0.05
0.01
1.9e−04
L−PF
L−F
L−P
L−T
R−PF
R−F
R−P
R−T
L−K
R−K
H

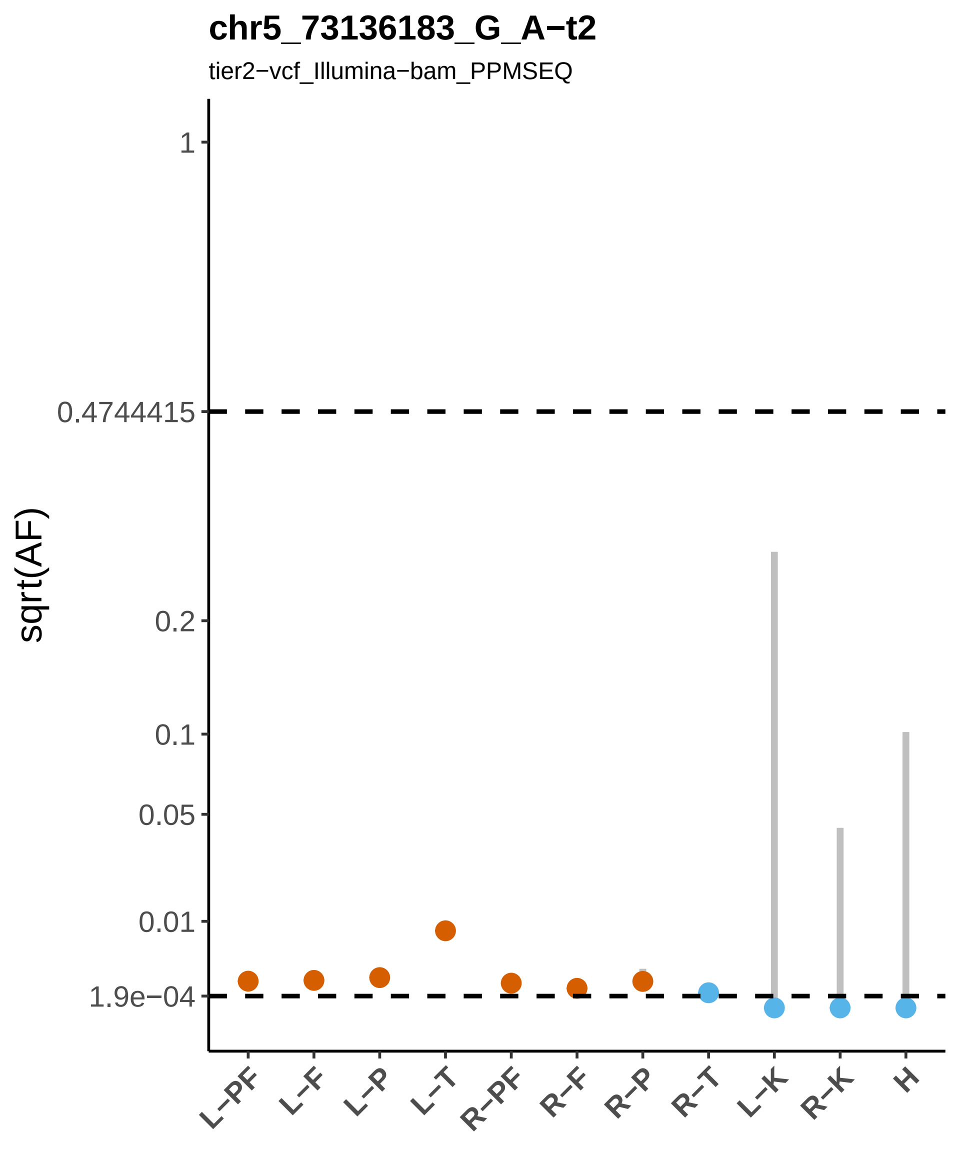

chr5_73136183_G_A−t2
tier2−vcf_Illumina−bam_PPMSEQ
sqrt(AF)
1
0.4744415
0.2
0.1
0.05
0.01
1.9e−04
L−PF
L−F
L−P
L−T
R−PF
R−F
R−P
R−T
L−K
R−K
H

### chr5_82993498_C_T−t2

tier2−vcf_ppmSeq_singleton_HC−bam_PTA

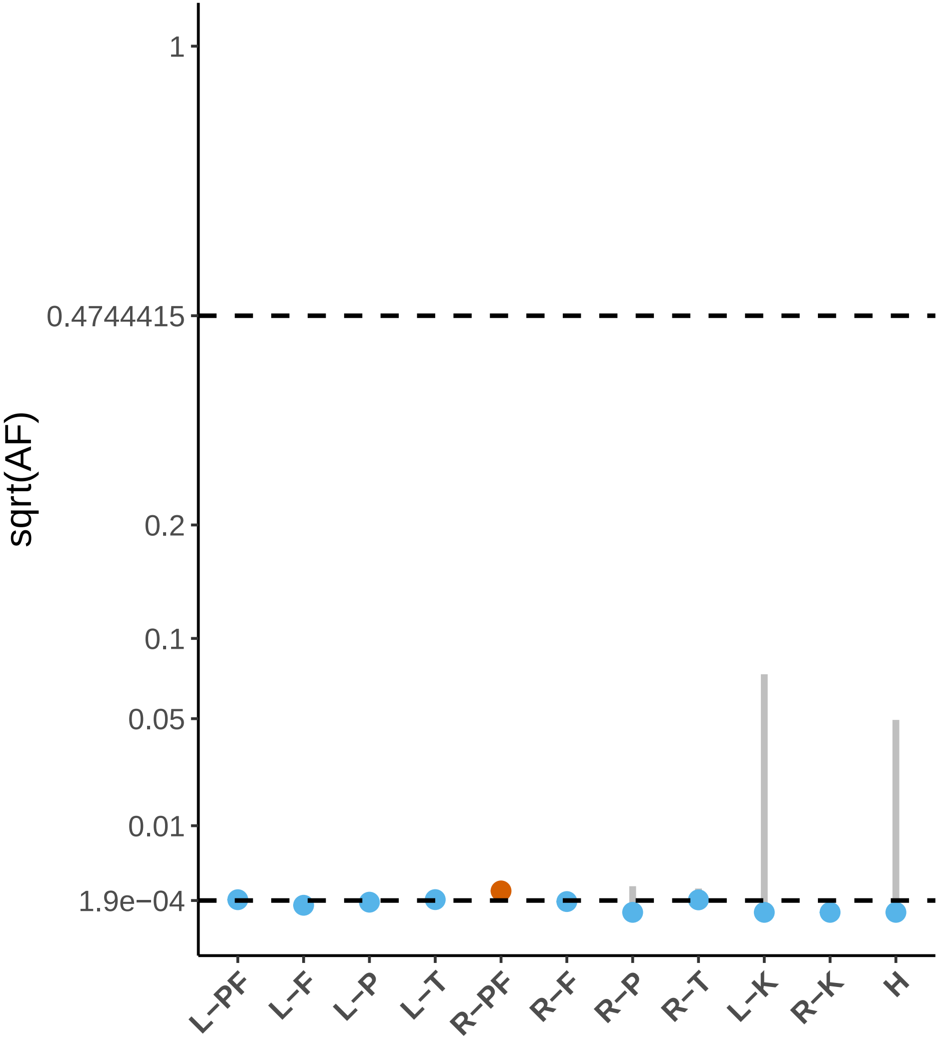

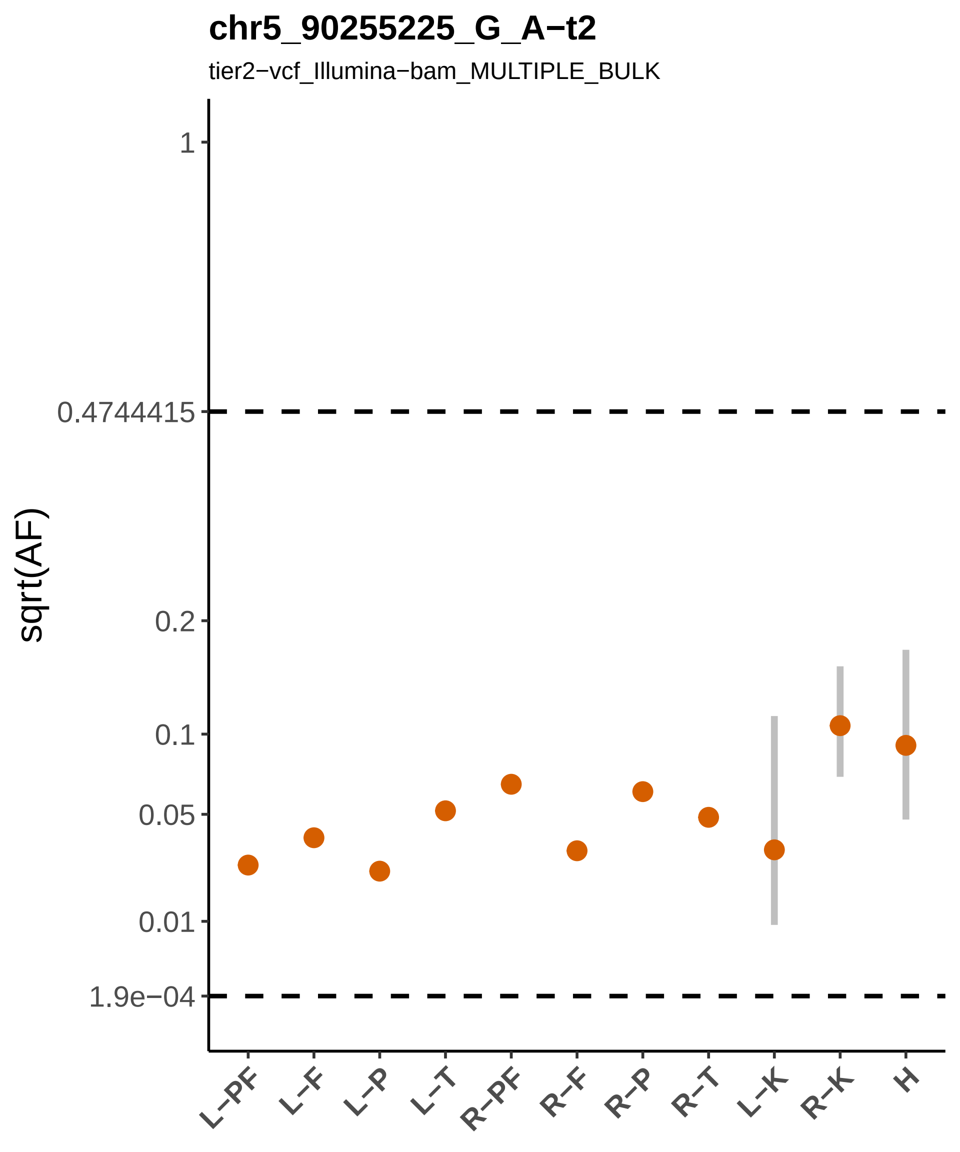

chr5_90255225_G_A−t2
tier2−vcf_Illumina−bam_MULTIPLE_BULK
sqrt(AF)
1
0.4744415
0.2
0.1
0.05
0.01
1.9e−04
L−PF
L−F
L−P
L−T
R−PF
R−F
R−P
R−T
L−K
R−K
H

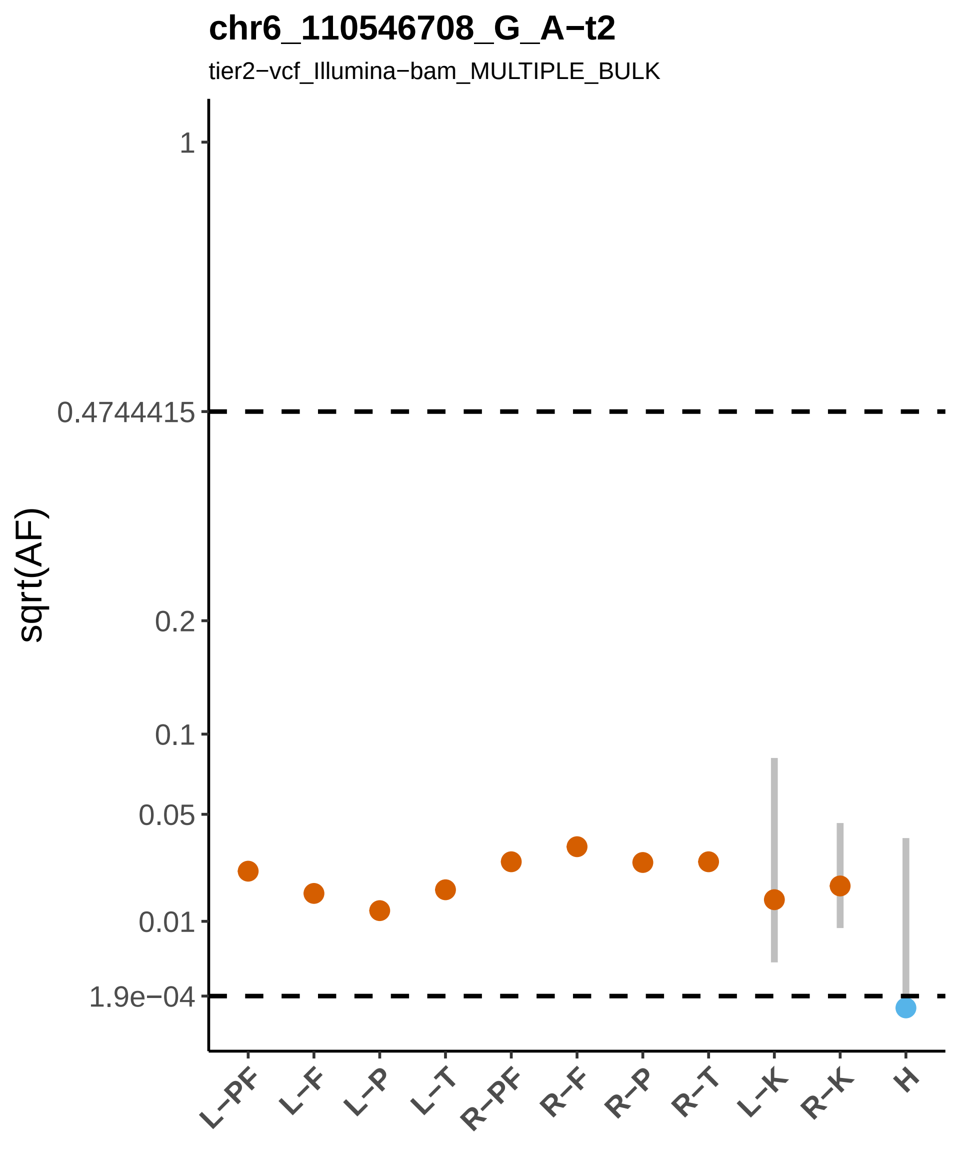

chr6_110546708_G_A−t2
tier2−vcf_Illumina−bam_MULTIPLE_BULK
sqrt(AF)
1
0.4744415
0.2
0.1
0.05
0.01
1.9e−04
L−PF
L−F
L−P
L−T
R−PF
R−F
R−P
R−T
L−K
R−K
H

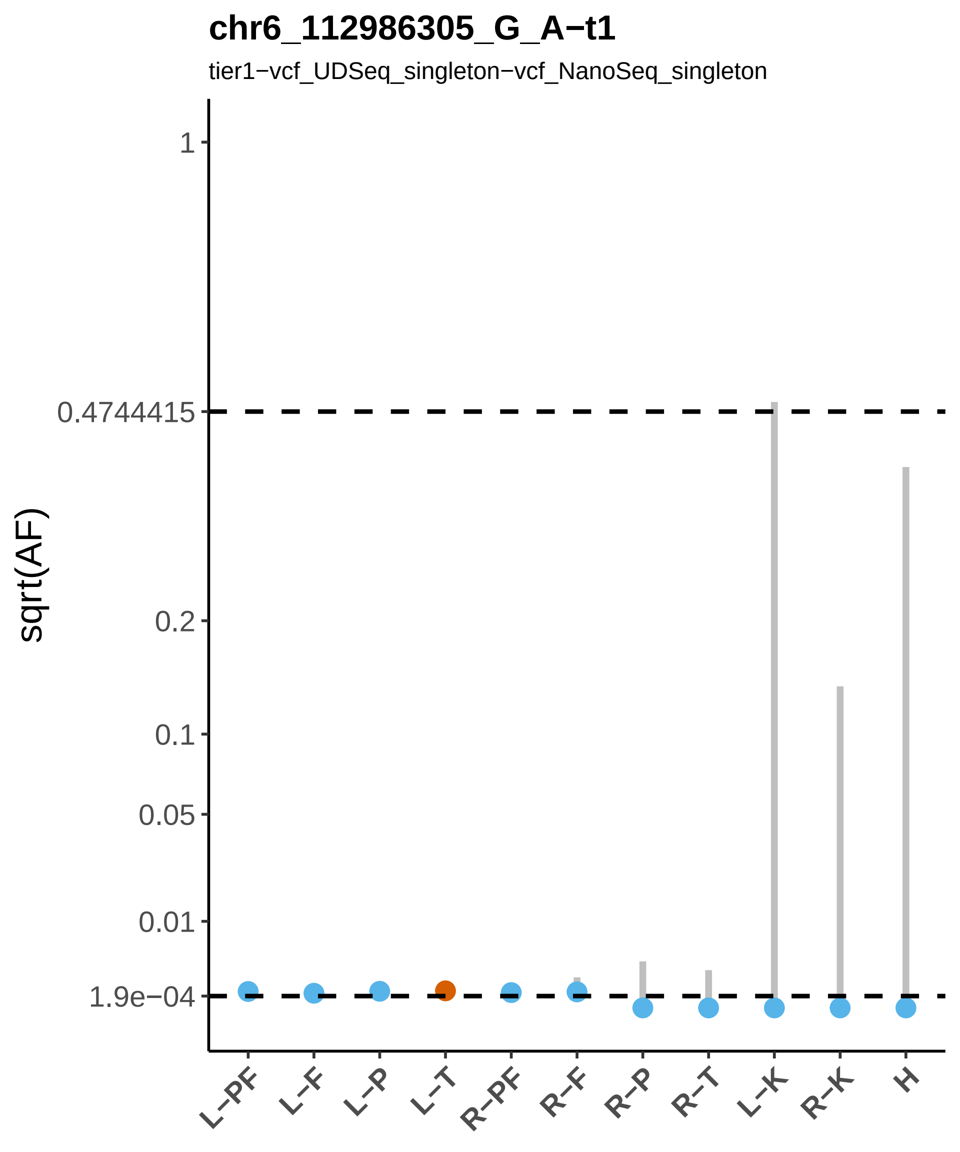
chr6_112986305_G_A−t1
tier1−vcf_UDSeq_singleton−vcf_NanoSeq_singleton
sqrt(AF)
1
0.4744415
0.2
0.1
0.05
0.01
1.9e−04
L−PF
L−F
L−P
L−T
R−PF
R−F
R−P
R−T
L−K
R−K
H

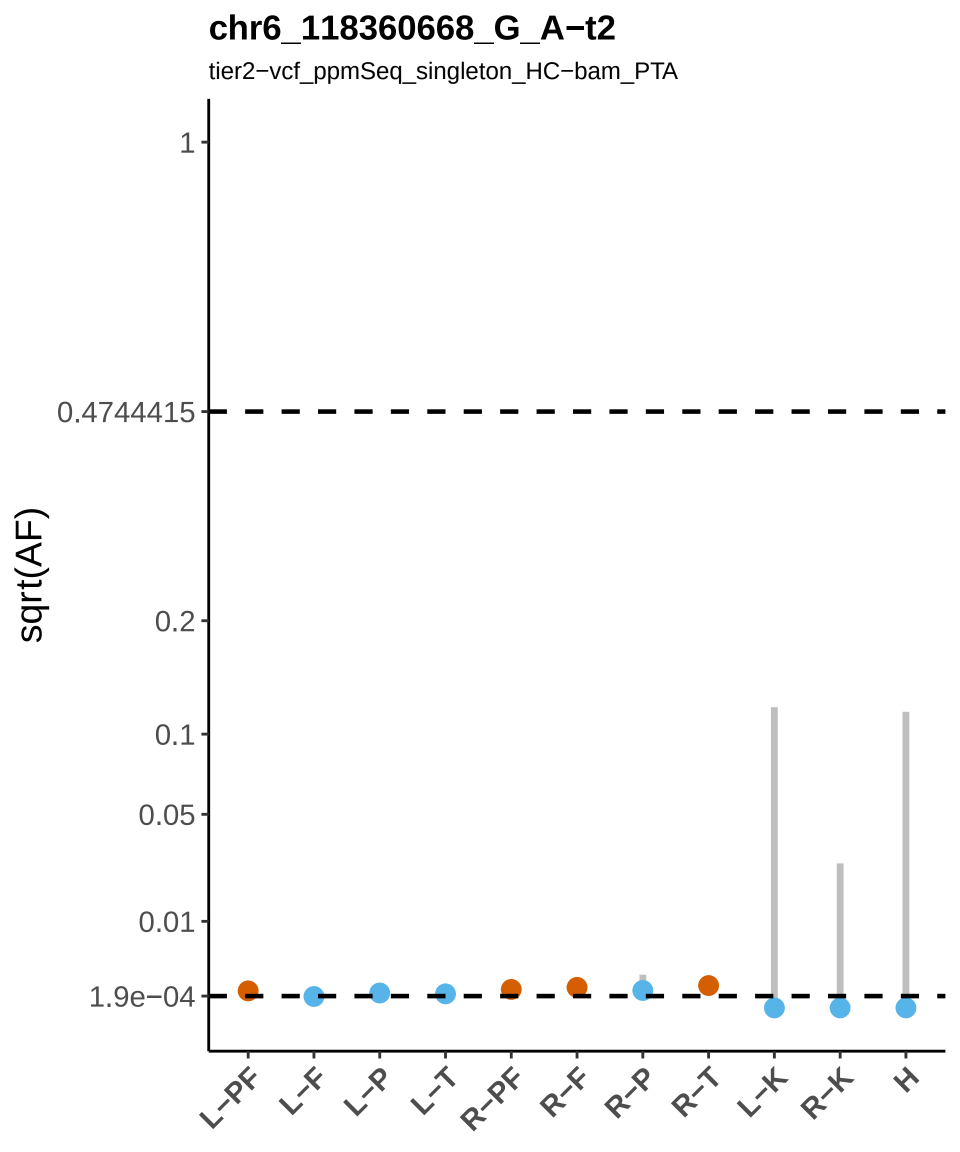
chr6_118360668_G_A−t2
tier2−vcf_ppmSeq_singleton_HC−bam_PTA
sqrt(AF)
1
0.4744415
0.2
0.1
0.05
0.01
1.9e−04
L−PF
L−F
L−P
L−T
R−PF
R−F
R−P
R−T
L−K
R−K
H

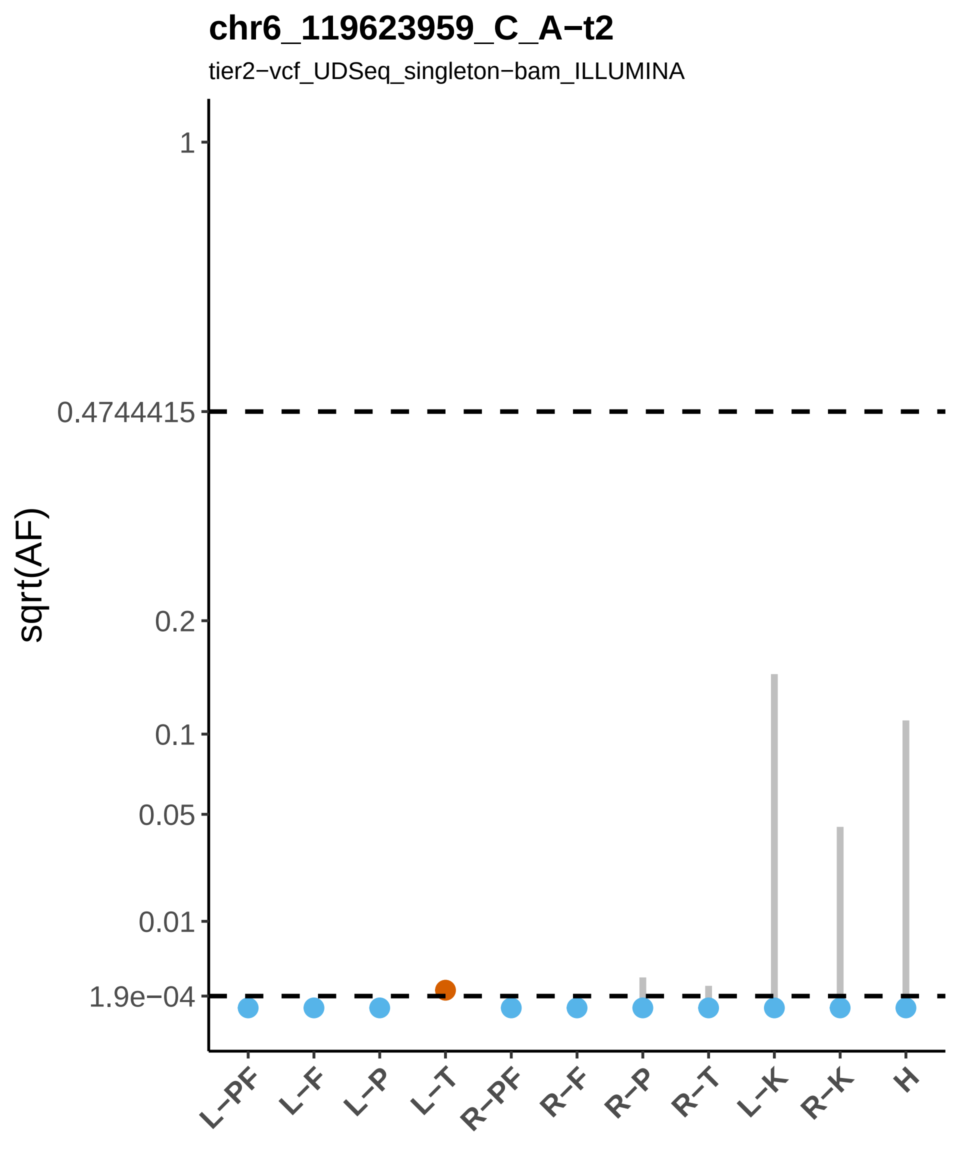

chr6_119623959_C_A−t2
tier2−vcf_UDSeq_singleton−bam_ILLUMINA
sqrt(AF)
1
0.4744415
0.2
0.1
0.05
0.01
1.9e−04
L−PF
L−F
L−P
L−T
R−PF
R−F
R−P
R−T
L−K
R−K
H

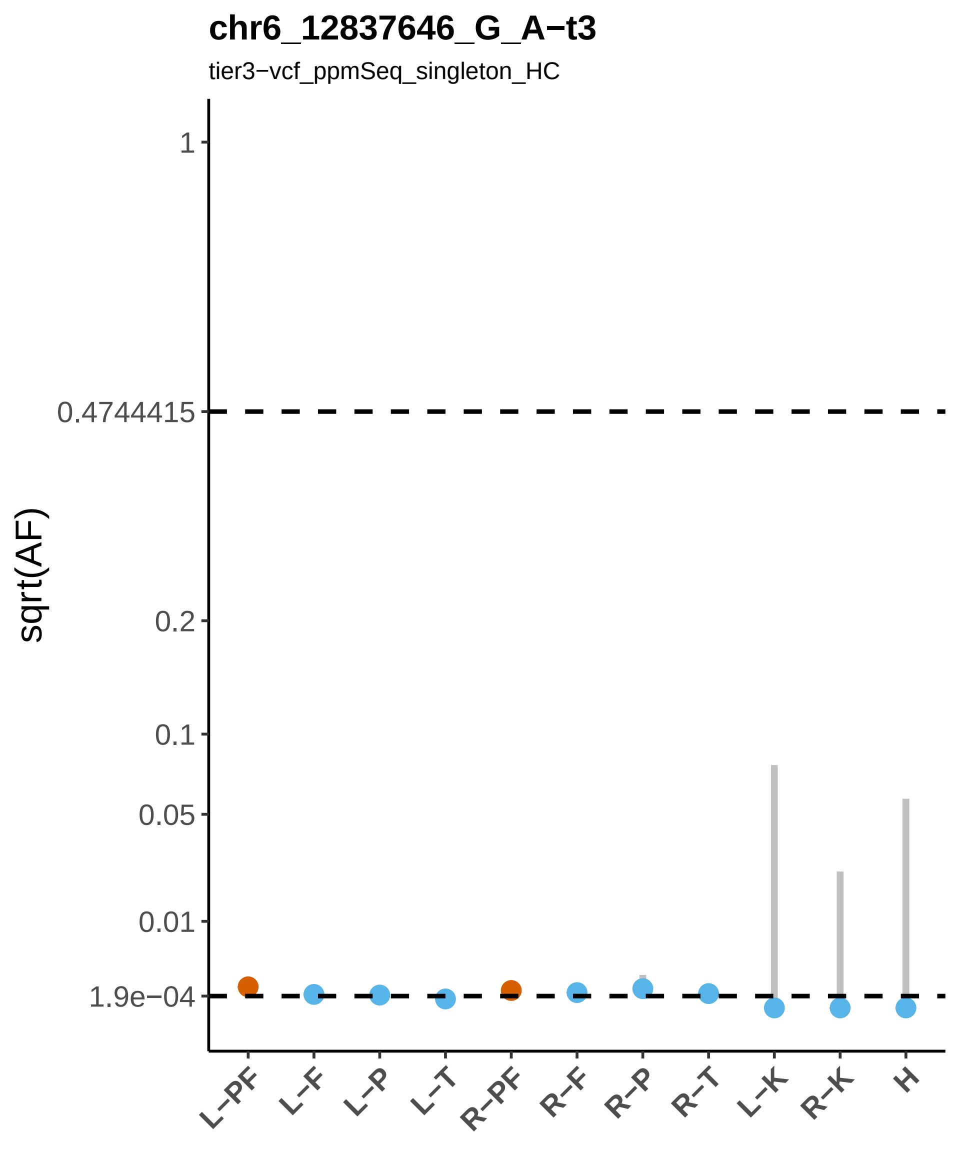
chr6_12837646_G_A−t3
tier3−vcf_ppmSeq_singleton_HC
sqrt(AF)
1
0.4744415
0.2
0.1
0.05
0.01
1.9e−04
L−PF
L−F
L−P
L−T
R−PF
R−F
R−P
R−T
L−K
R−K
H

#### chr6_130719209_T_C−t3

tier3−vcf_PTA_singleton

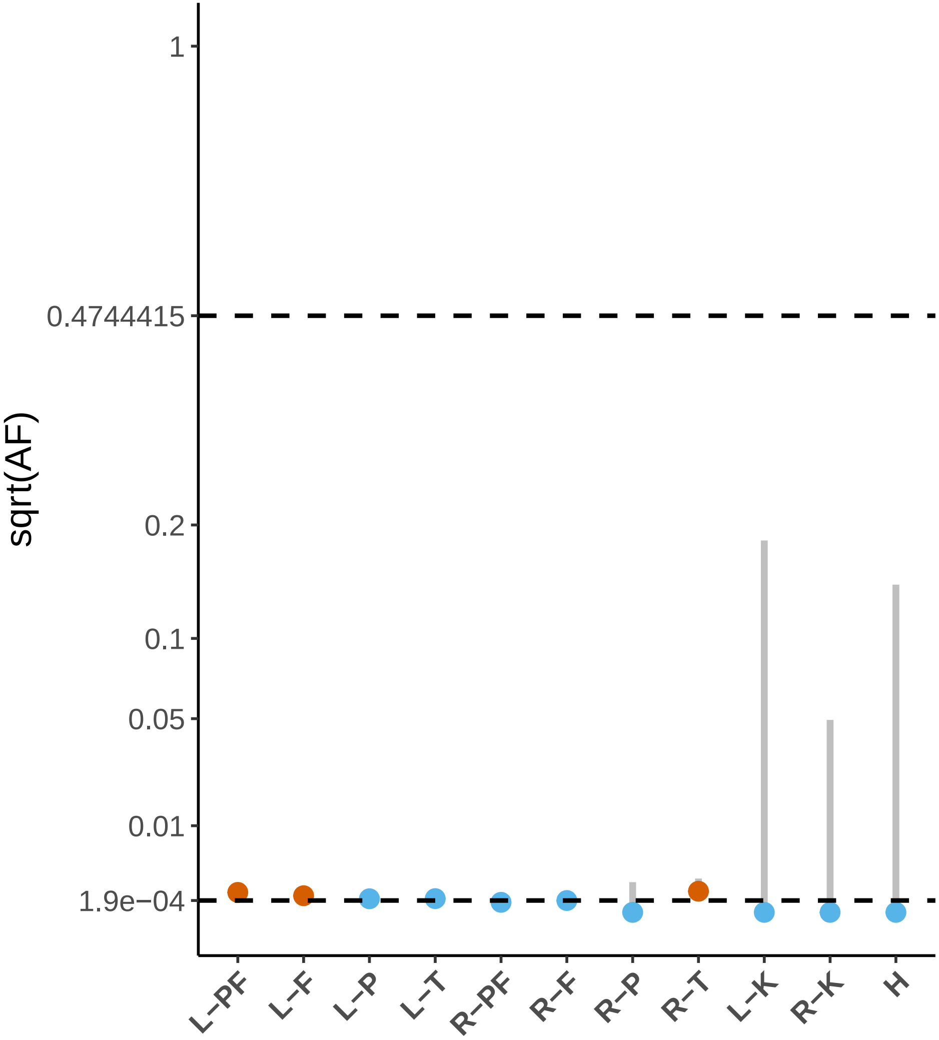

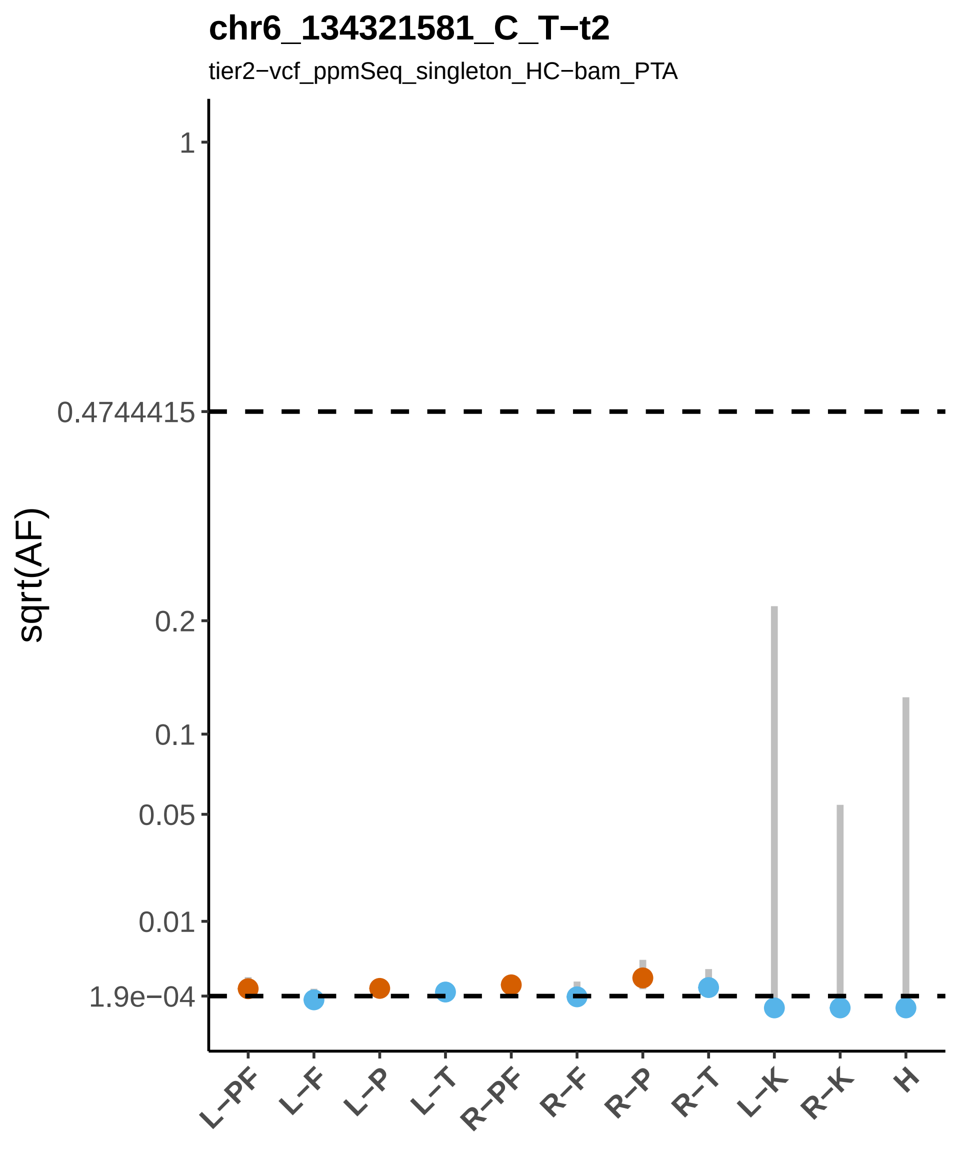

chr6_134321581_C_T−t2
tier2−vcf_ppmSeq_singleton_HC−bam_PTA
sqrt(AF)
1
0.4744415
0.2
0.1
0.05
0.01
1.9e−04
L−PF
L−F
L−P
L−T
R−PF
R−F
R−P
R−T
L−K
R−K
H

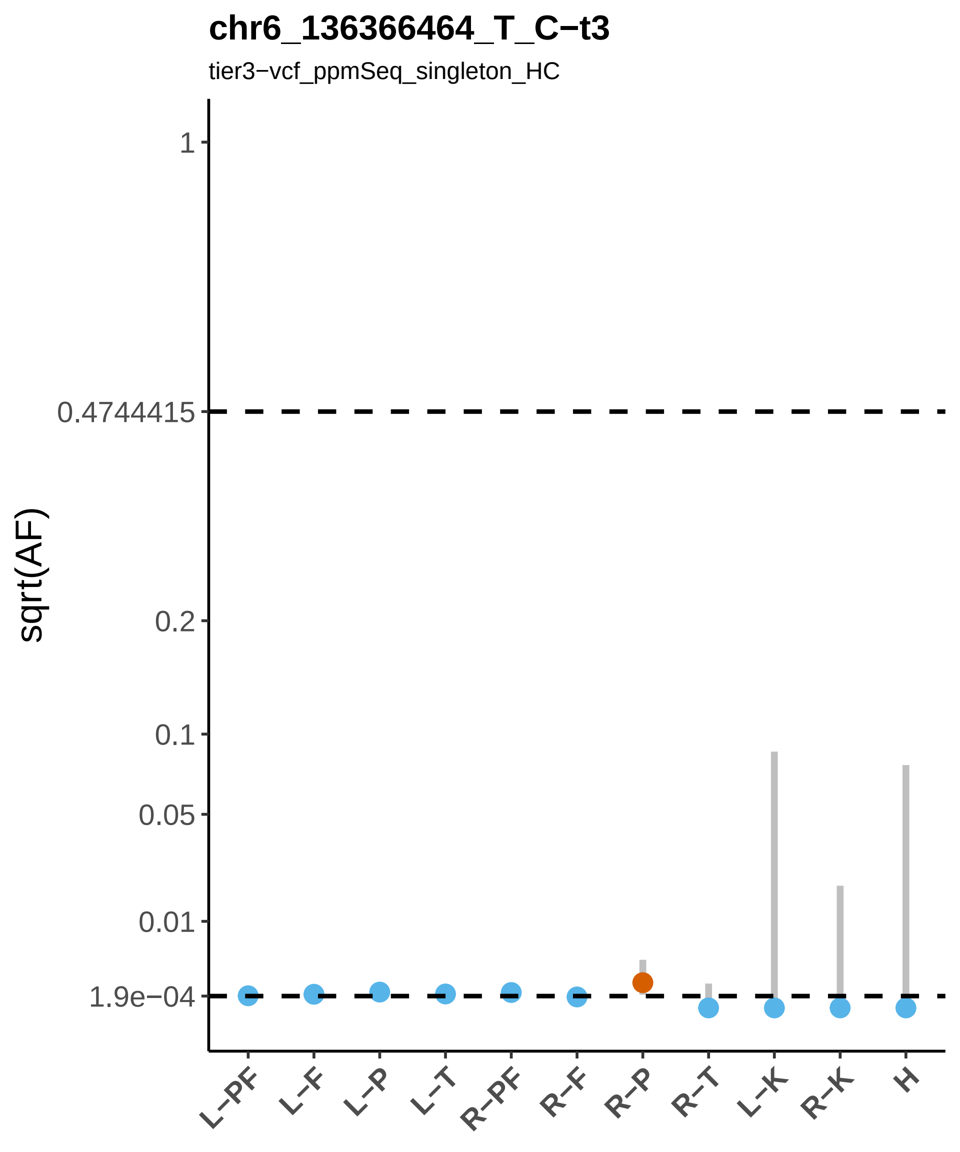
chr6_136366464_T_C−t3
tier3−vcf_ppmSeq_singleton_HC
sqrt(AF)
1
0.4744415
0.2
0.1
0.05
0.01
1.9e−04
L−PF
L−F
L−P
L−T
R−PF
R−F
R−P
R−T
L−K
R−K
H

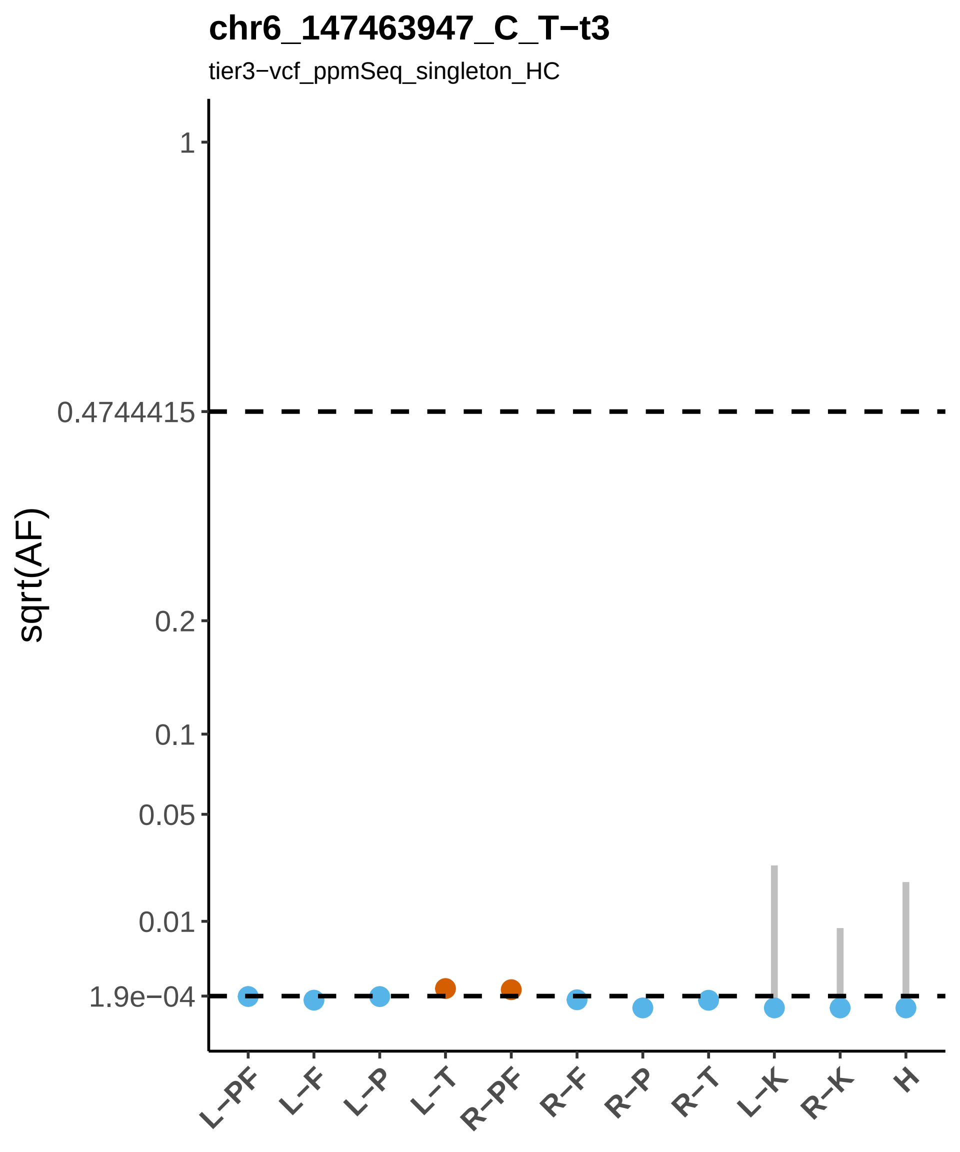

chr6_147463947_C_T−t3
tier3−vcf_ppmSeq_singleton_HC
sqrt(AF)
1
0.4744415
0.2
0.1
0.05
0.01
1.9e−04
L−PF
L−F
L−P
L−T
R−PF
R−F
R−P
R−T
L−K
R−K
H

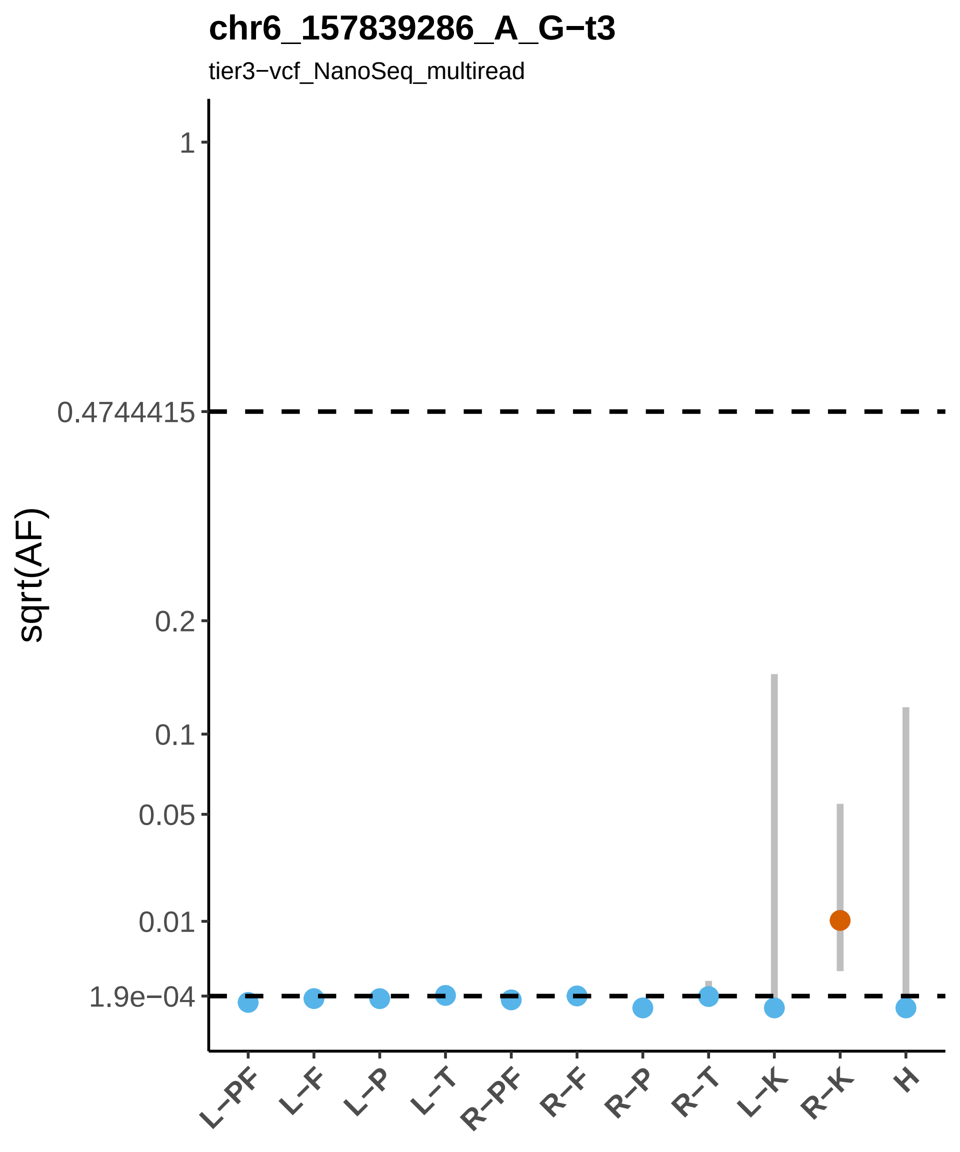
chr6_157839286_A_G−t3
tier3−vcf_NanoSeq_multiread
sqrt(AF)
1
0.4744415
0.2
0.1
0.05
0.01
1.9e−04
L−PF
L−F
L−P
L−T
R−PF
R−F
R−P
R−T
L−K
R−K
H

#### chr6_169061825_C_A−t2

tier2−vcf_ppmSeq_singleton_HC−bam_ILLUMINA

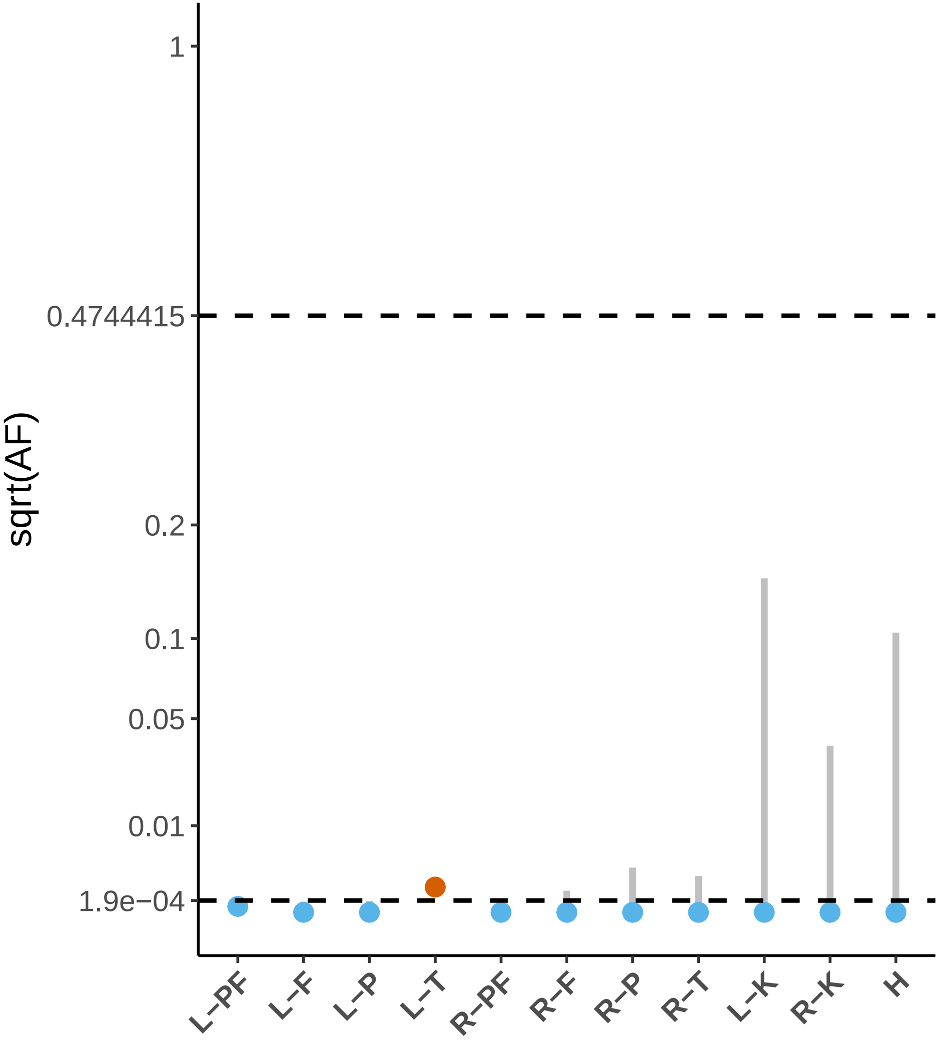

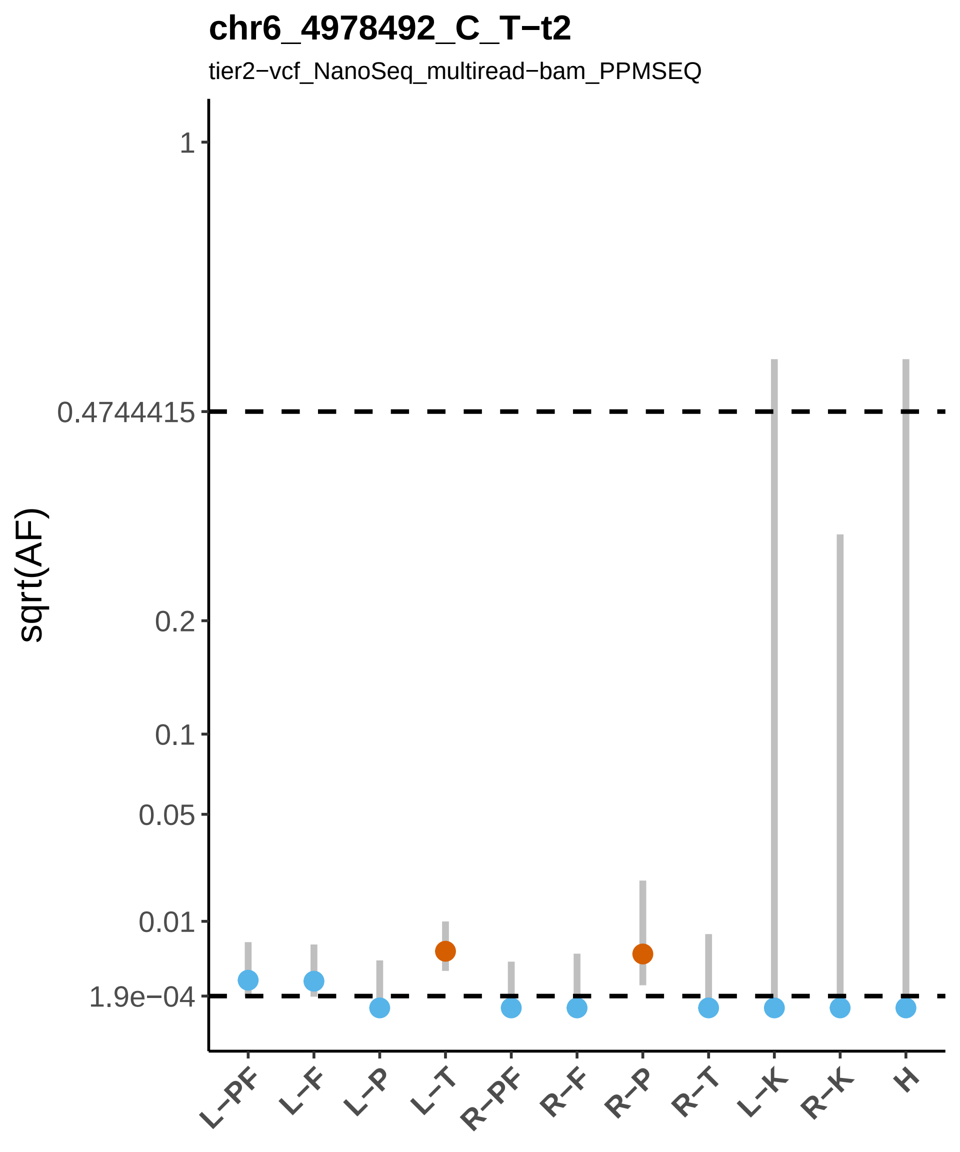
chr6_4978492_C_T−t2
tier2−vcf_NanoSeq_multiread−bam_PPMSEQ
sqrt(AF)
1
0.4744415
0.2
0.1
0.05
0.01
1.9e−04
L−PF
L−F
L−P
L−T
R−PF
R−F
R−P
R−T
L−K
R−K
H

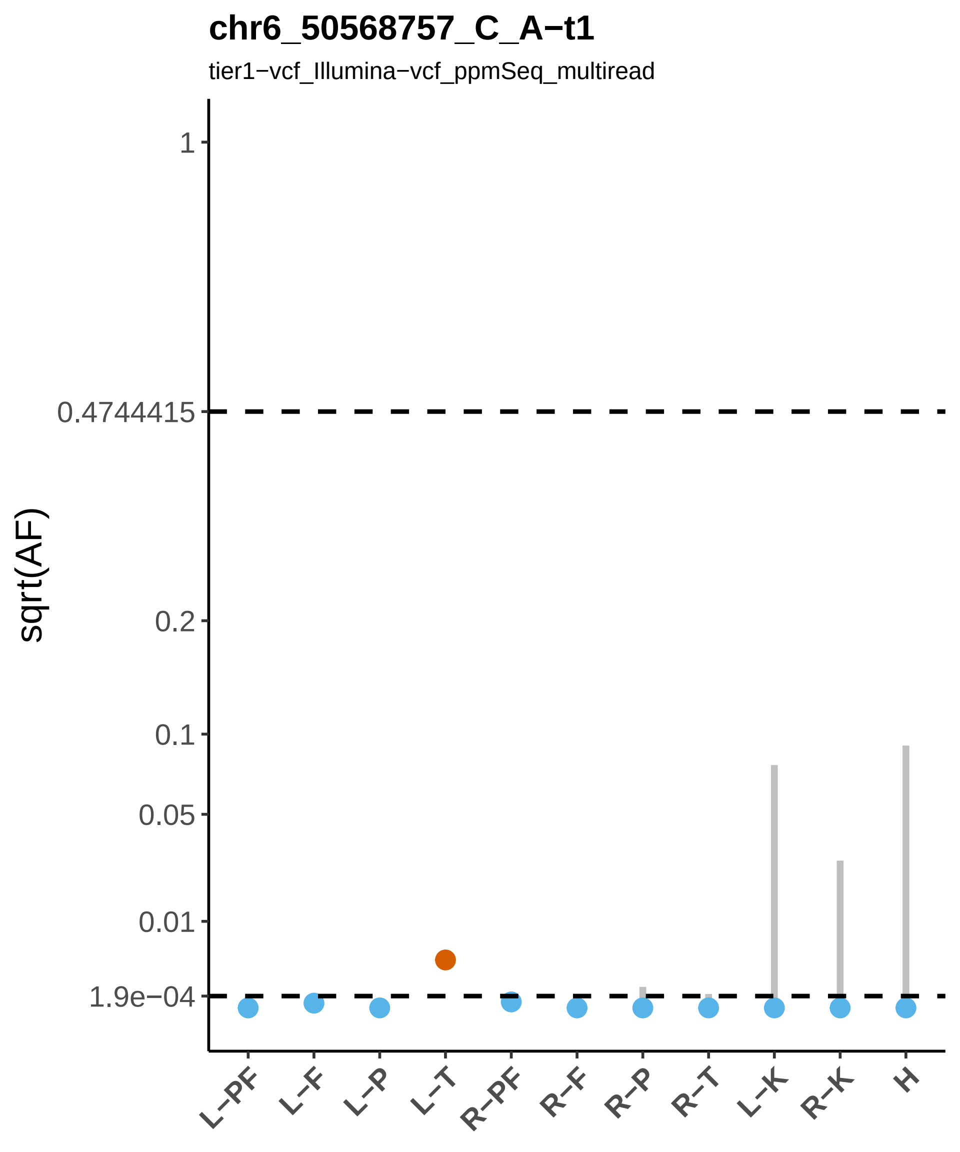

chr6_50568757_C_A−t1
tier1−vcf_Illumina−vcf_ppmSeq_multiread
sqrt(AF)
1
0.4744415
0.2
0.1
0.05
0.01
1.9e−04
L−PF
L−F
L−P
L−T
R−PF
R−F
R−P
R−T
L−K
R−K
H

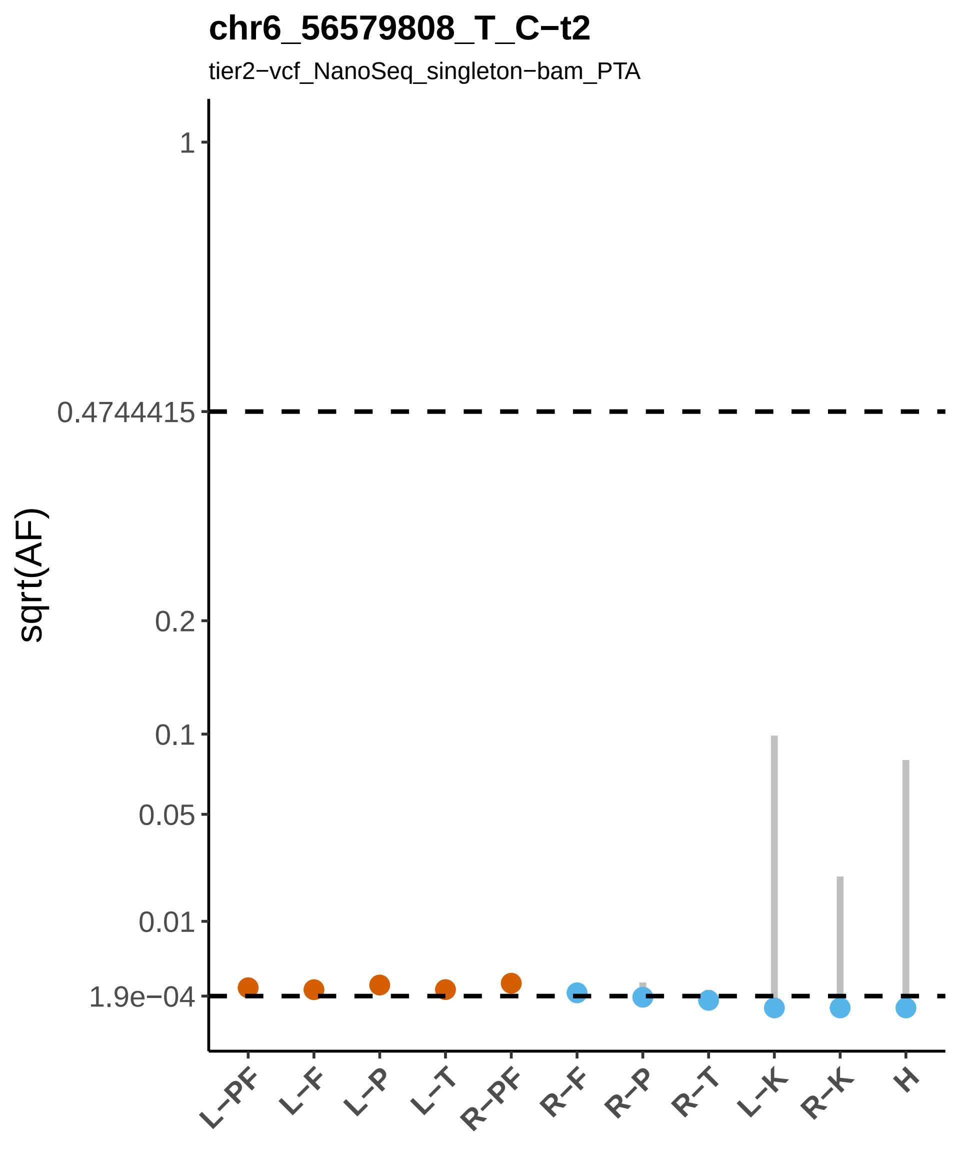

chr6_56579808_T_C−t2
tier2−vcf_NanoSeq_singleton−bam_PTA
sqrt(AF)
1
0.4744415
0.2
0.1
0.05
0.01
1.9e−04
L−PF
L−F
L−P
L−T
R−PF
R−F
R−P
R−T
L−K
R−K
H

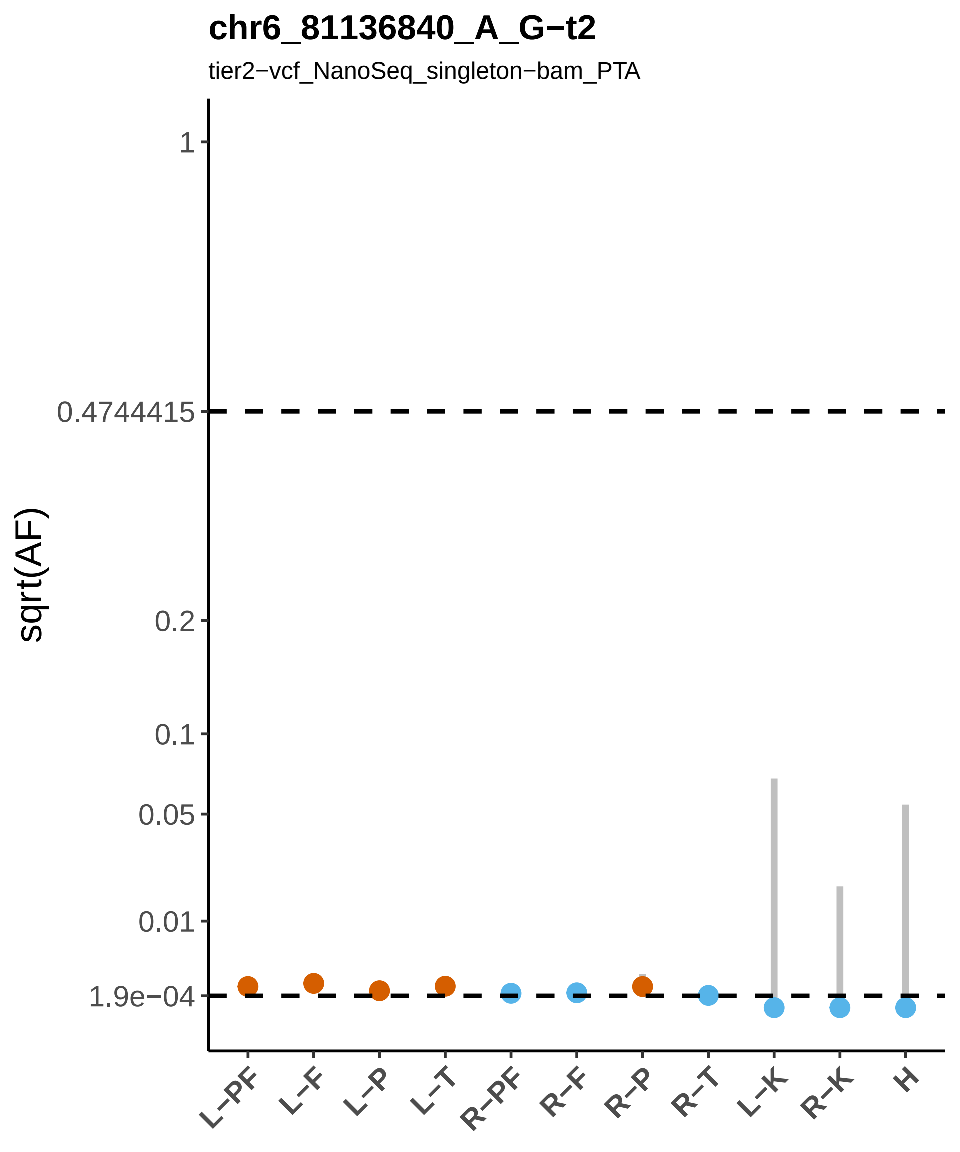

chr6_81136840_A_G−t2
tier2−vcf_NanoSeq_singleton−bam_PTA
sqrt(AF)
1
0.4744415
0.2
0.1
0.05
0.01
1.9e−04
L−PF
L−F
L−P
L−T
R−PF
R−F
R−P
R−T
L−K
R−K
H

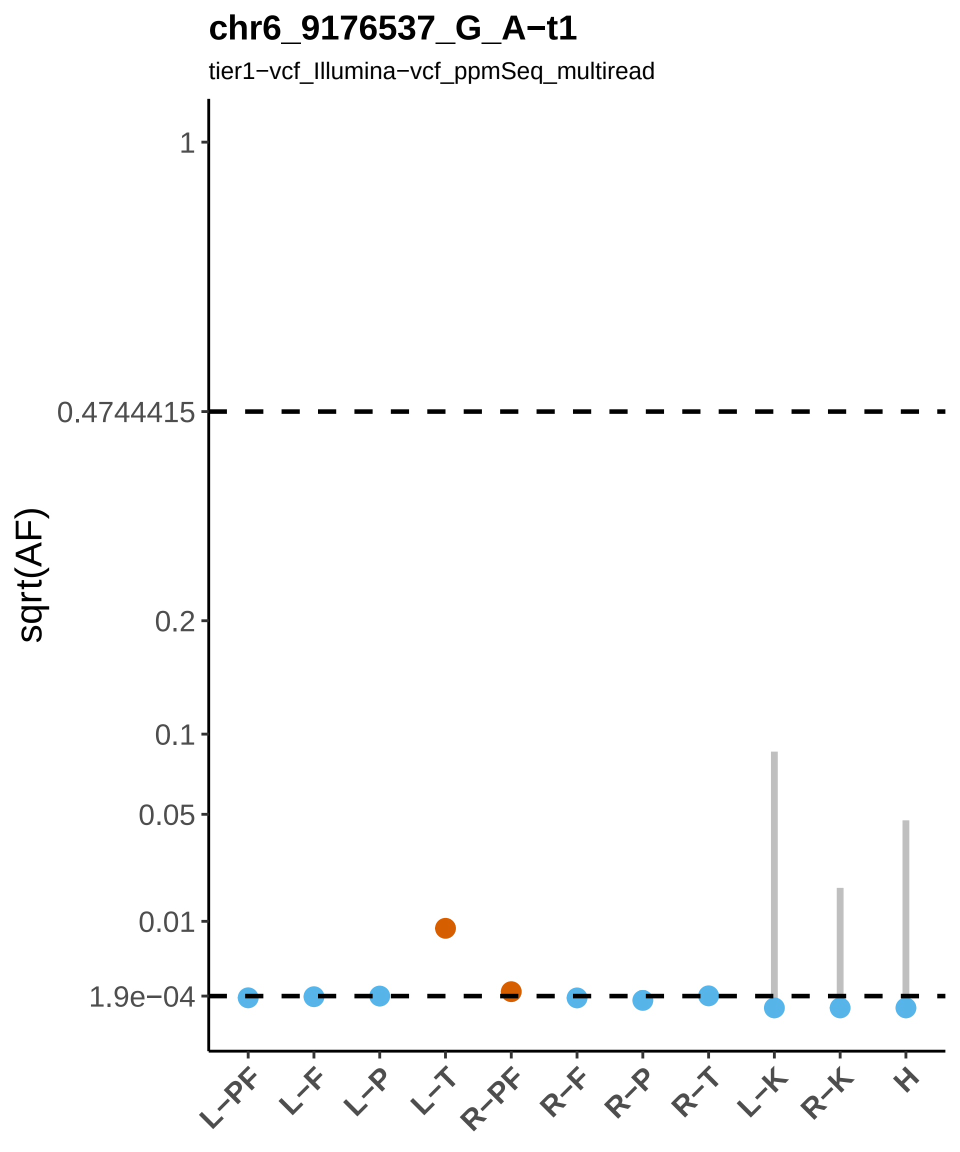

chr6_9176537_G_A−t1
tier1−vcf_Illumina−vcf_ppmSeq_multiread
sqrt(AF)
1
0.4744415
0.2
0.1
0.05
0.01
1.9e−04
L−PF
L−F
L−P
L−T
R−PF
R−F
R−P
R−T
L−K
R−K
H

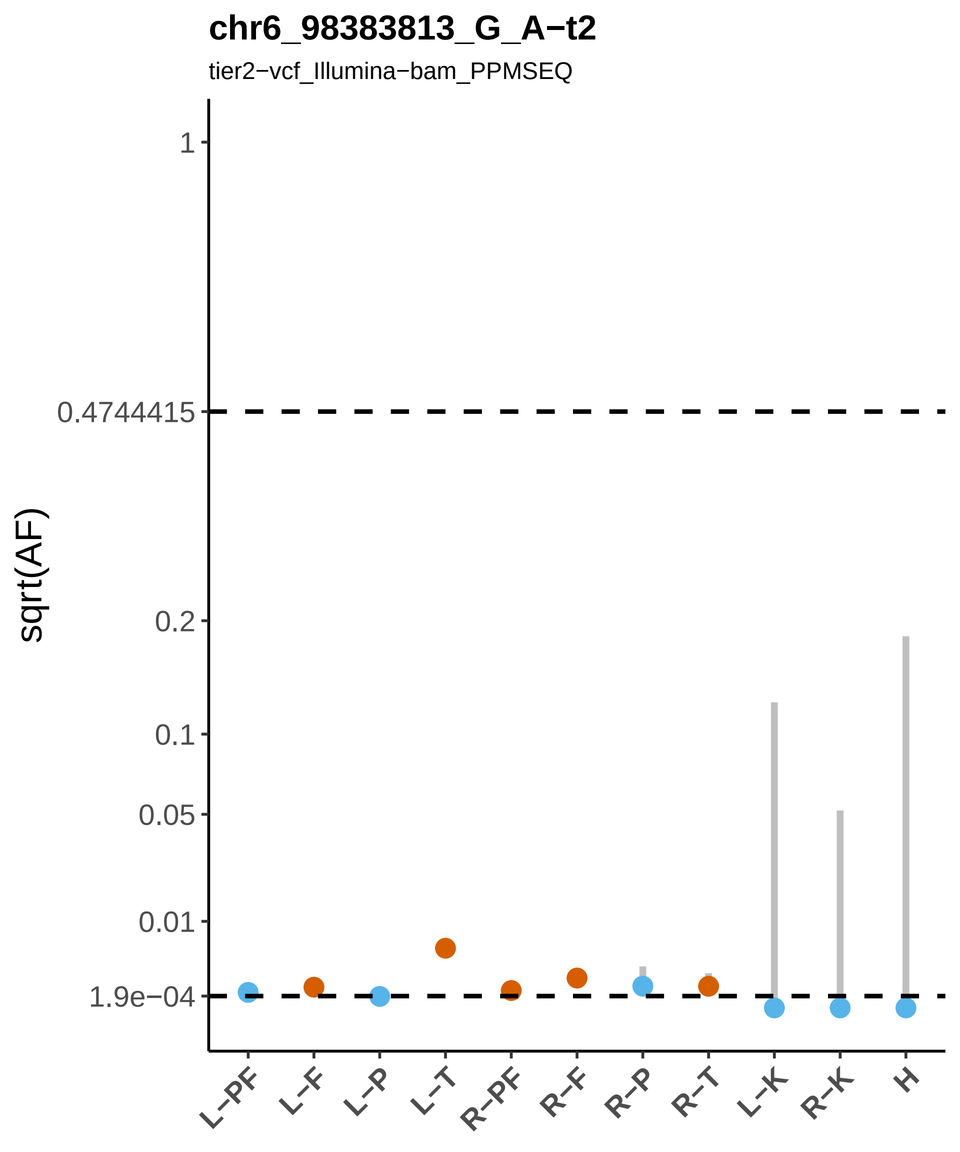
chr6_98383813_G_A−t2
tier2−vcf_Illumina−bam_PPMSEQ
sqrt(AF)
1
0.4744415
0.2
0.1
0.05
0.01
1.9e−04
L−PF
L−F
L−P
L−T
R−PF
R−F
R−P
R−T
L−K
R−K
H

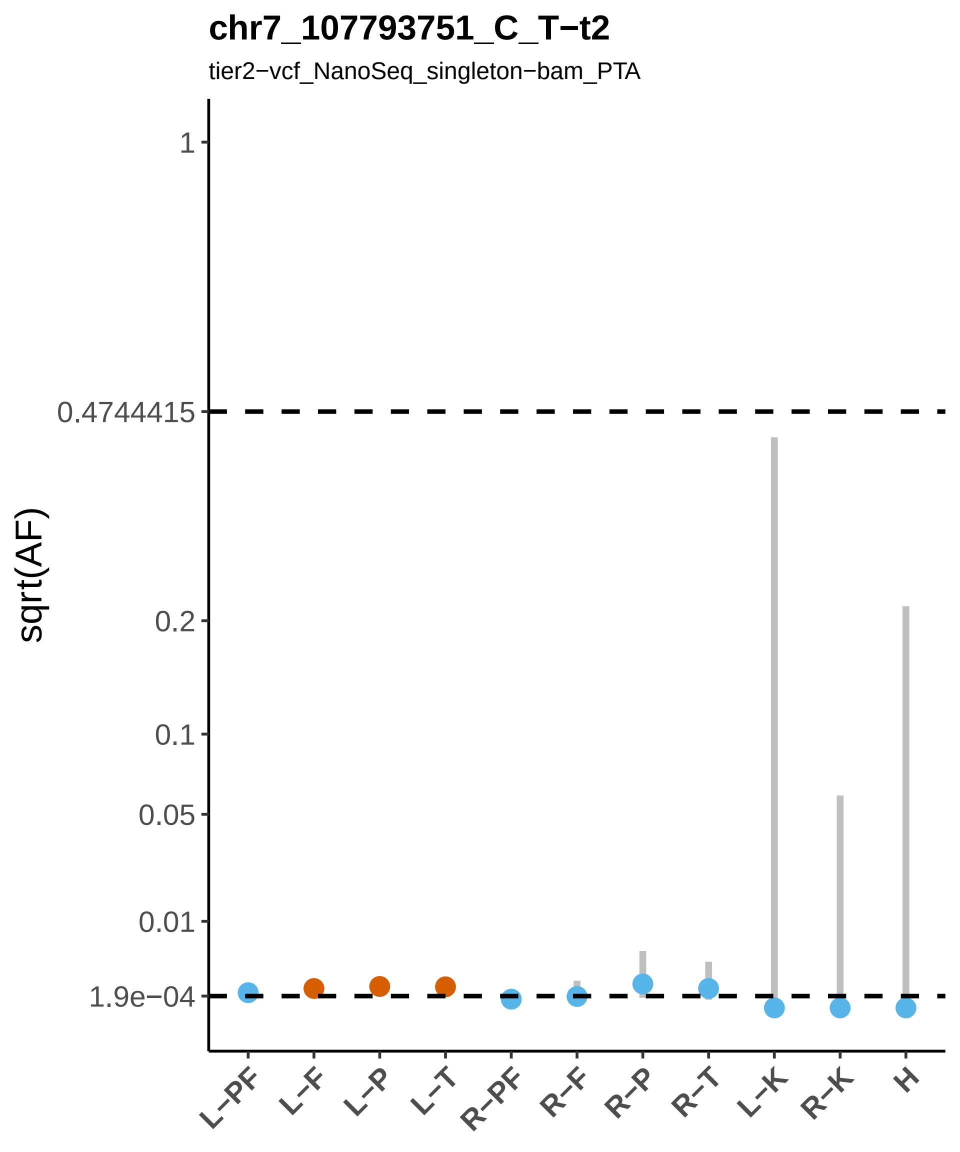
chr7_107793751_C_T−t2
tier2−vcf_NanoSeq_singleton−bam_PTA
sqrt(AF)
1
0.4744415
0.2
0.1
0.05
0.01
1.9e−04
L−PF
L−F
L−P
L−T
R−PF
R−F
R−P
R−T
L−K
R−K
H

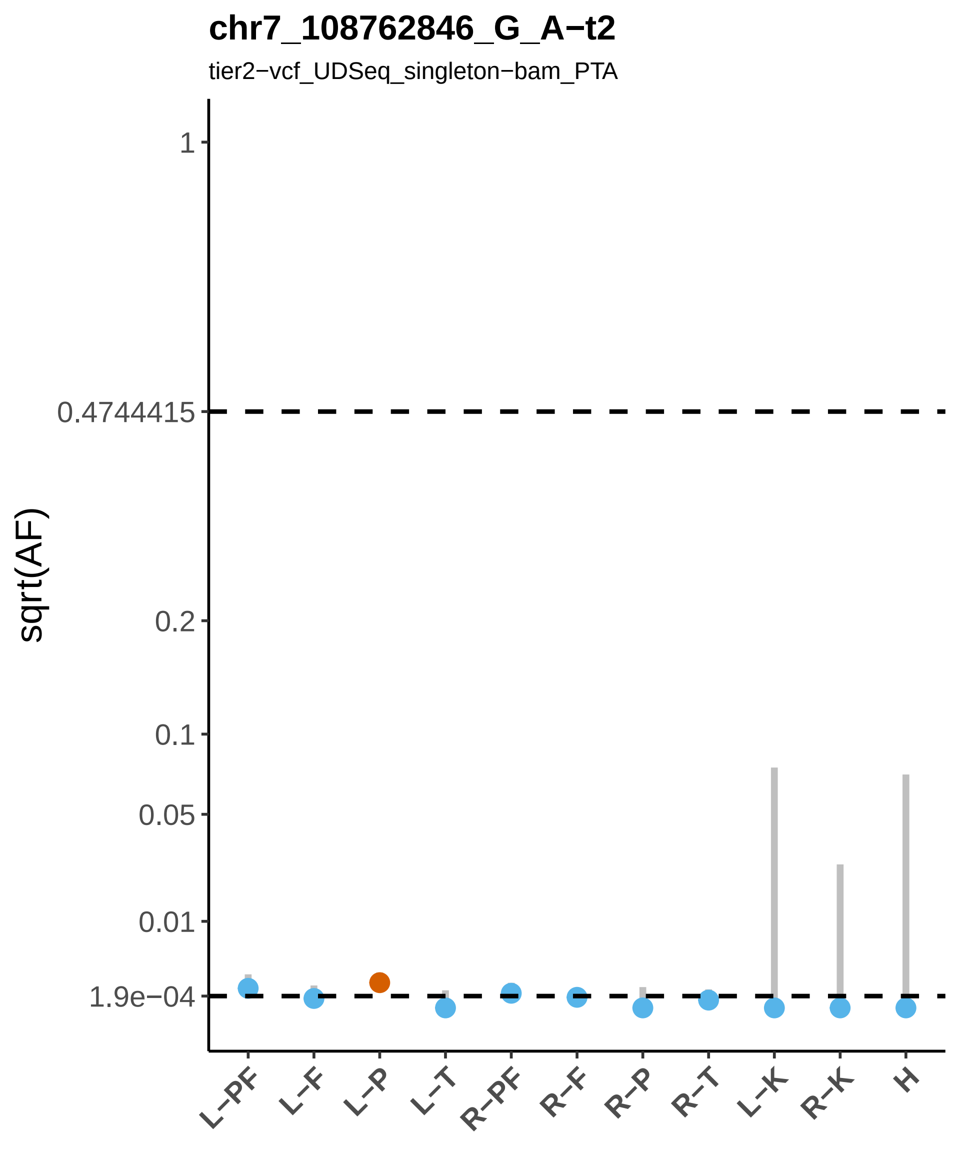
chr7_108762846_G_A−t2
tier2−vcf_UDSeq_singleton−bam_PTA
sqrt(AF)
1
0.4744415
0.2
0.1
0.05
0.01
1.9e−04
L−PF
L−F
L−P
L−T
R−PF
R−F
R−P
R−T
L−K
R−K
H

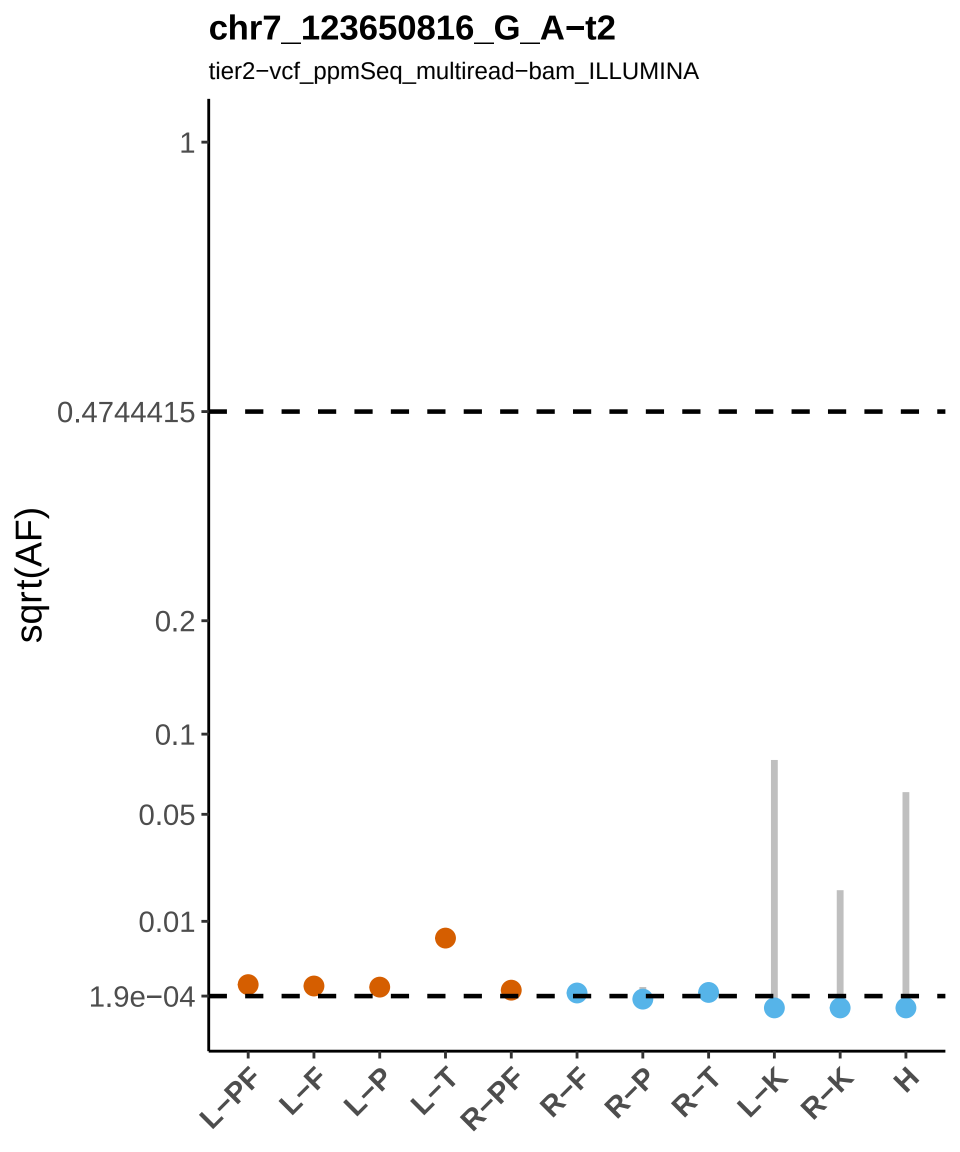

chr7_123650816_G_A−t2
tier2−vcf_ppmSeq_multiread−bam_ILLUMINA
sqrt(AF)
1
0.4744415
0.2
0.1
0.05
0.01
1.9e−04
L−PF
L−F
L−P
L−T
R−PF
R−F
R−P
R−T
L−K
R−K
H

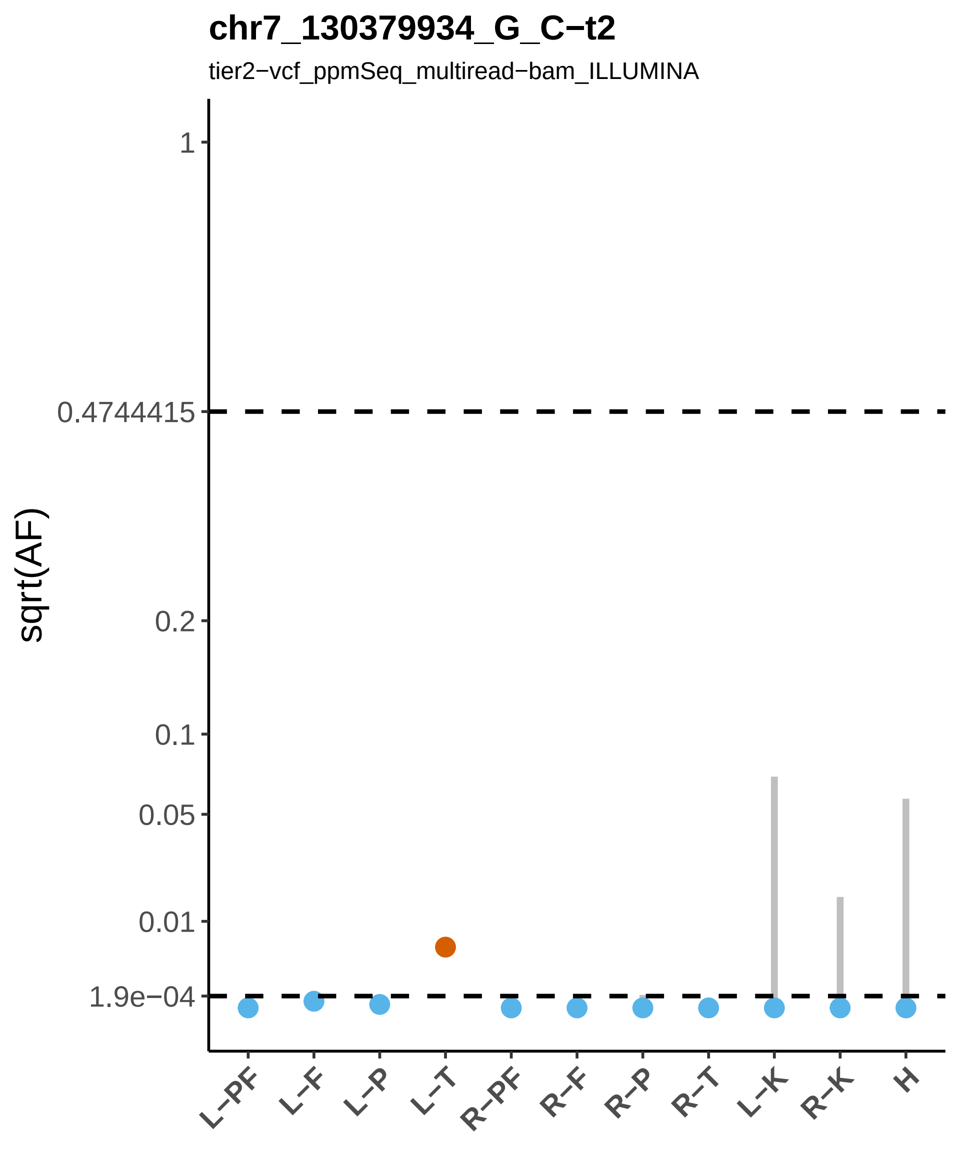

chr7_130379934_G_C−t2
tier2−vcf_ppmSeq_multiread−bam_ILLUMINA
sqrt(AF)
1
0.4744415
0.2
0.1
0.05
0.01
1.9e−04
L−PF
L−F
L−P
L−T
R−PF
R−F
R−P
R−T
L−K
R−K
H

### chr7_155172864_A_G−t3

tier3−vcf_ppmSeq_multiread

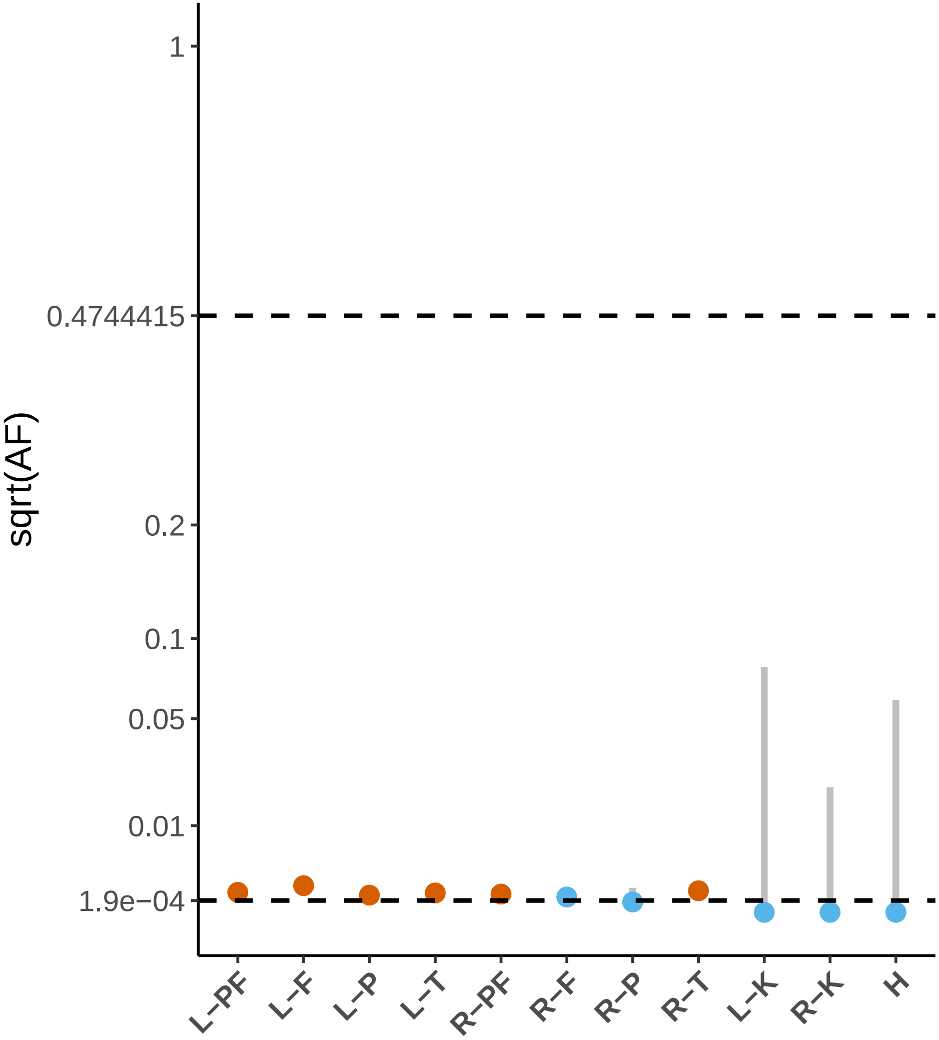

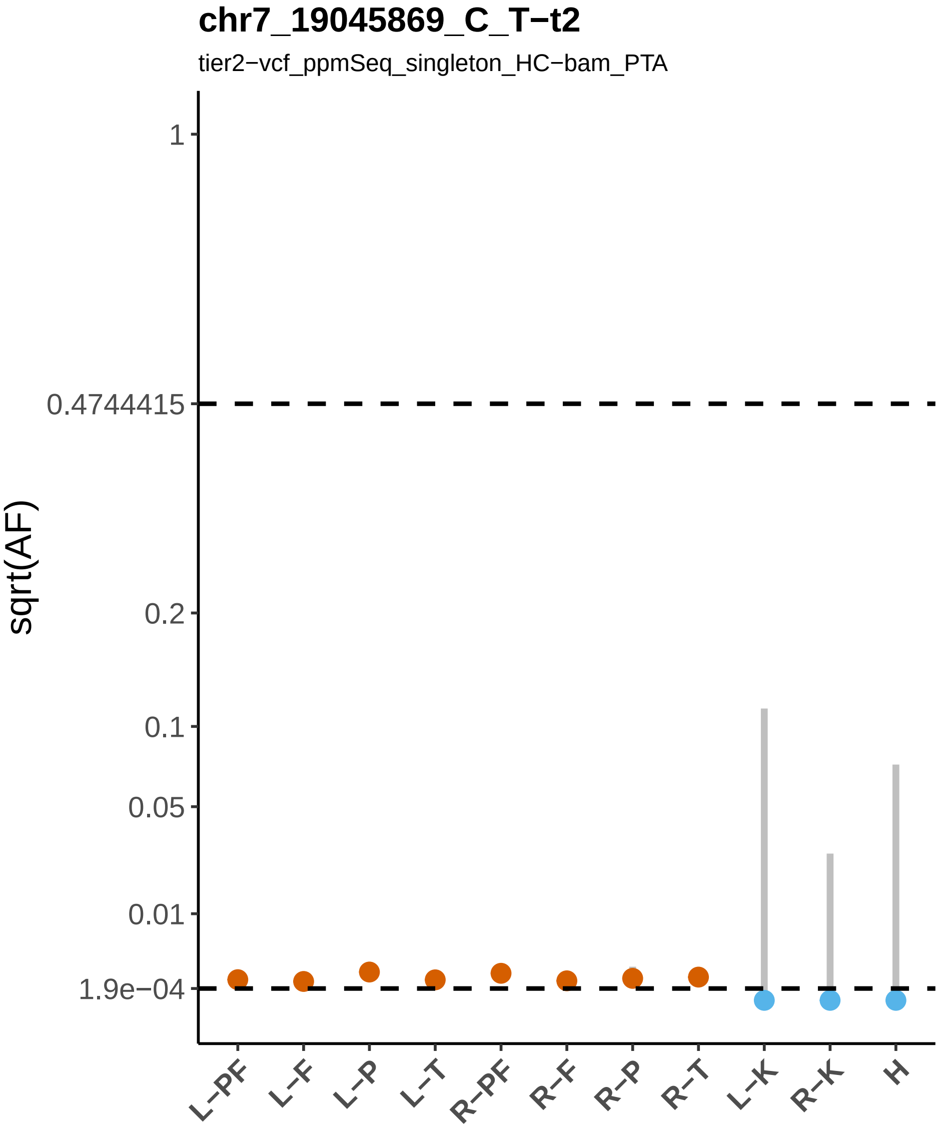

chr7_19045869_C_T−t2
tier2−vcf_ppmSeq_singleton_HC−bam_PTA
sqrt(AF)
1
0.4744415
0.2
0.1
0.05
0.01
1.9e−04
L−PF
L−F
L−P
L−T
R−PF
R−F
R−P
R−T
L−K
R−K
H

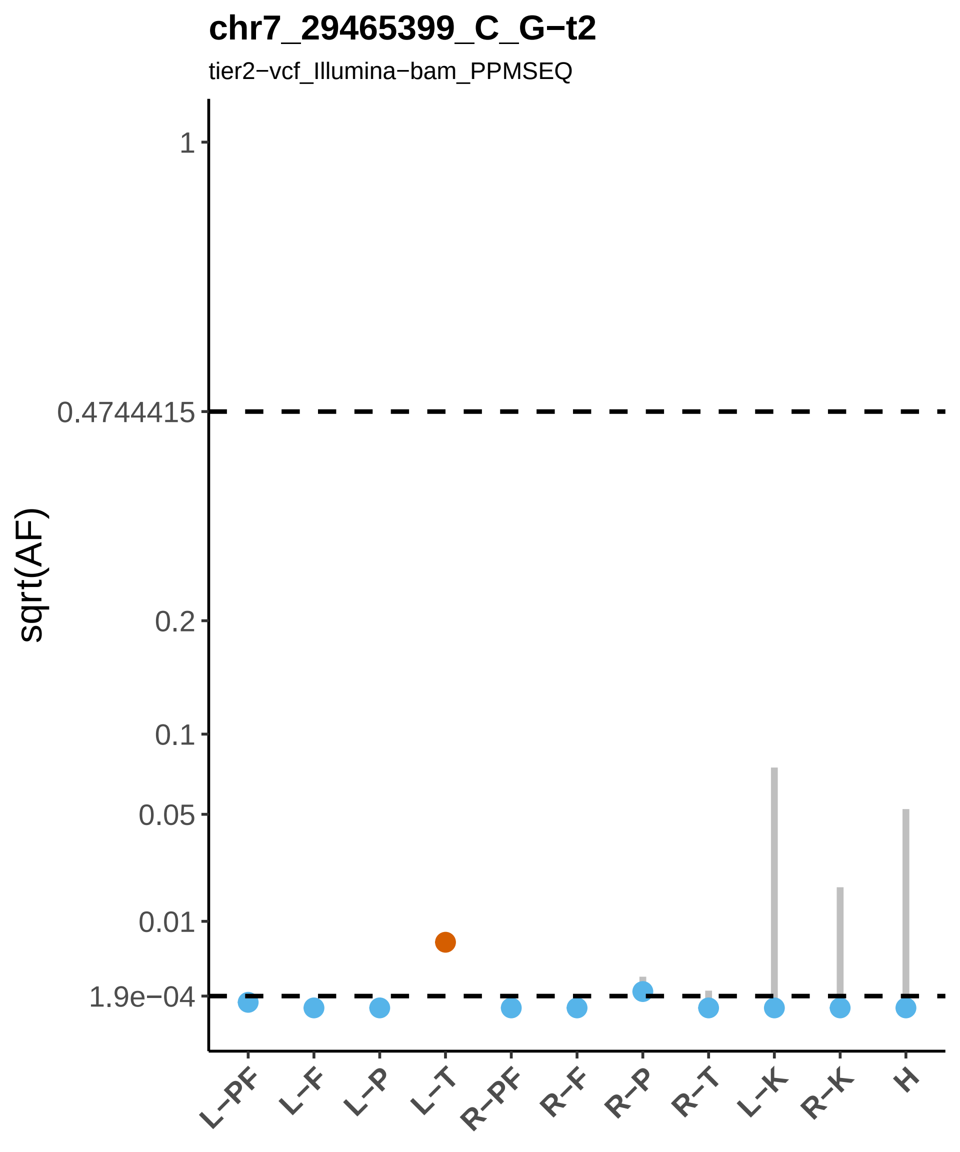

chr7_29465399_C_G−t2
tier2−vcf_Illumina−bam_PPMSEQ
sqrt(AF)
1
0.4744415
0.2
0.1
0.05
0.01
1.9e−04
L−PF
L−F
L−P
L−T
R−PF
R−F
R−P
R−T
L−K
R−K
H

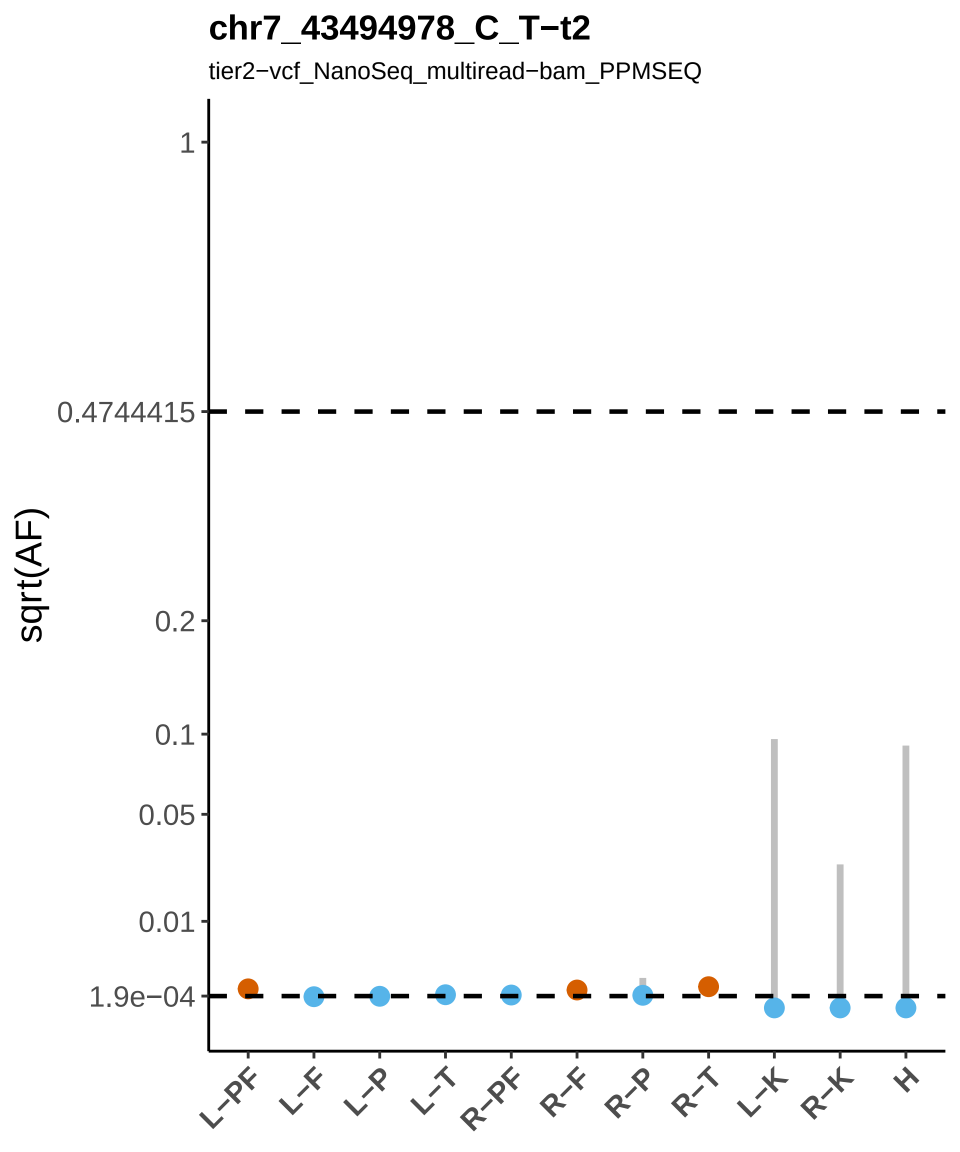
chr7_43494978_C_T−t2
tier2−vcf_NanoSeq_multiread−bam_PPMSEQ
sqrt(AF)
1
0.4744415
0.2
0.1
0.05
0.01
1.9e−04
L−PF
L−F
L−P
L−T
R−PF
R−F
R−P
R−T
L−K
R−K
H

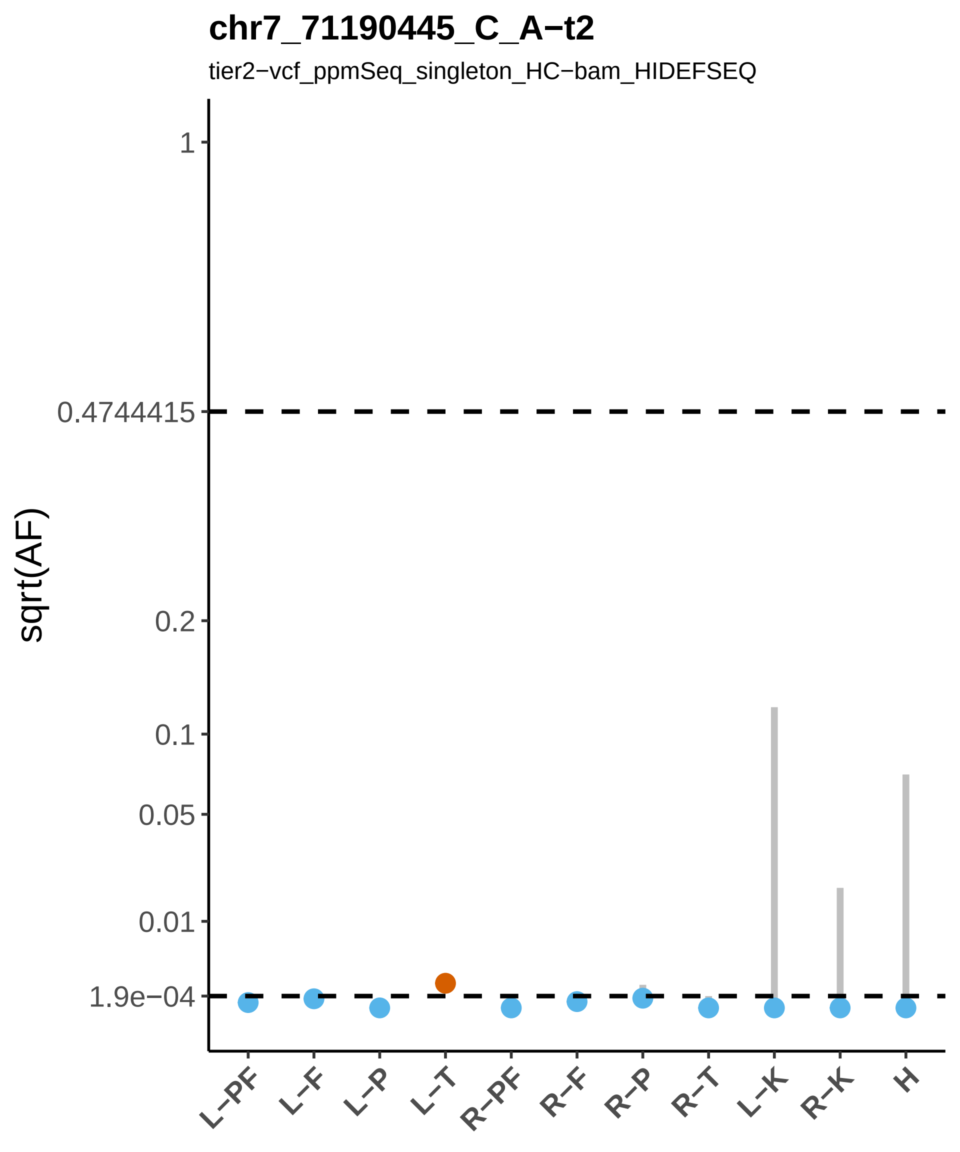

chr7_71190445_C_A−t2
tier2−vcf_ppmSeq_singleton_HC−bam_HIDEFSEQ
sqrt(AF)
1
0.4744415
0.2
0.1
0.05
0.01
1.9e−04
L−PF
L−F
L−P
L−T
R−PF
R−F
R−P
R−T
L−K
R−K
H

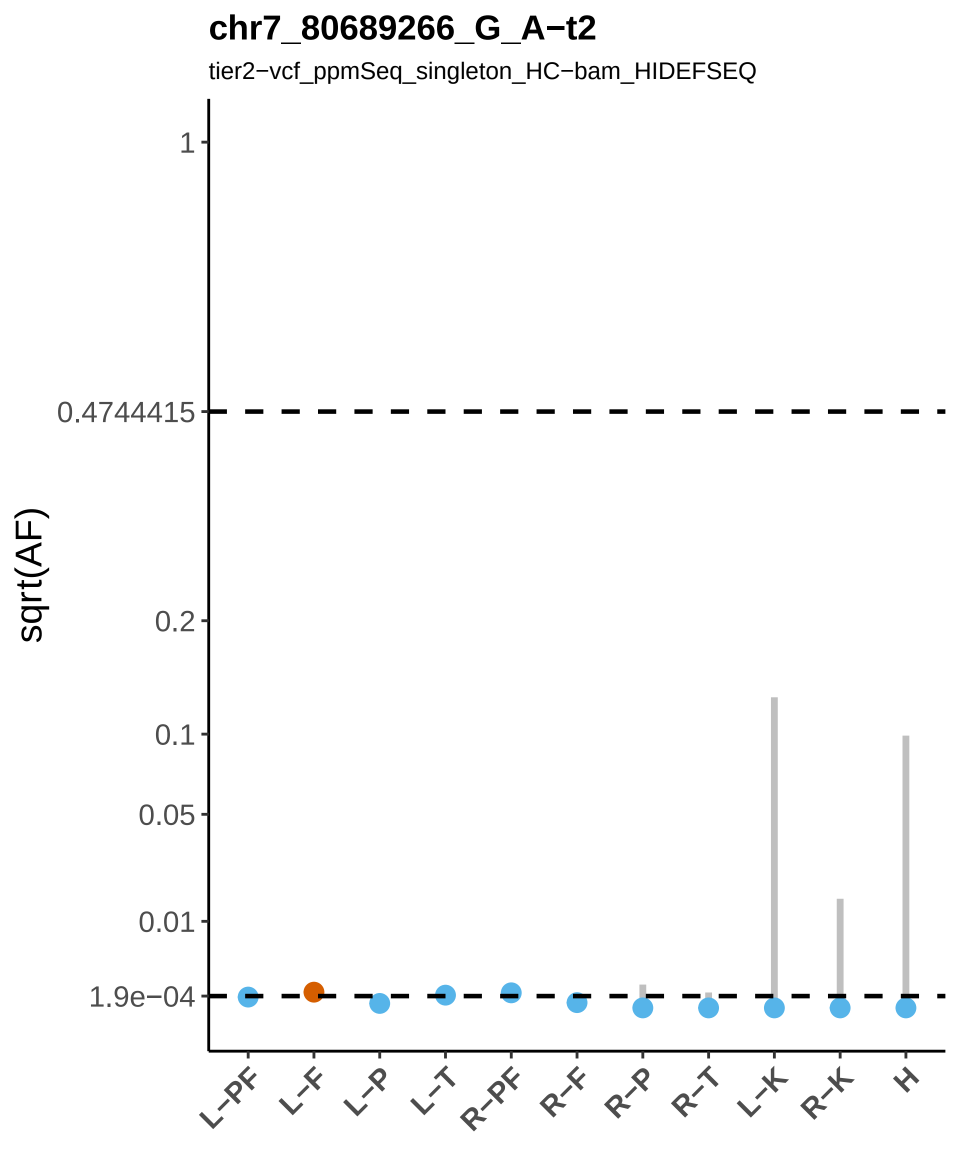
chr7_80689266_G_A−t2
tier2−vcf_ppmSeq_singleton_HC−bam_HIDEFSEQ
sqrt(AF)
1
0.4744415
0.2
0.1
0.05
0.01
1.9e−04
L−PF
L−F
L−P
L−T
R−PF
R−F
R−P
R−T
L−K
R−K
H

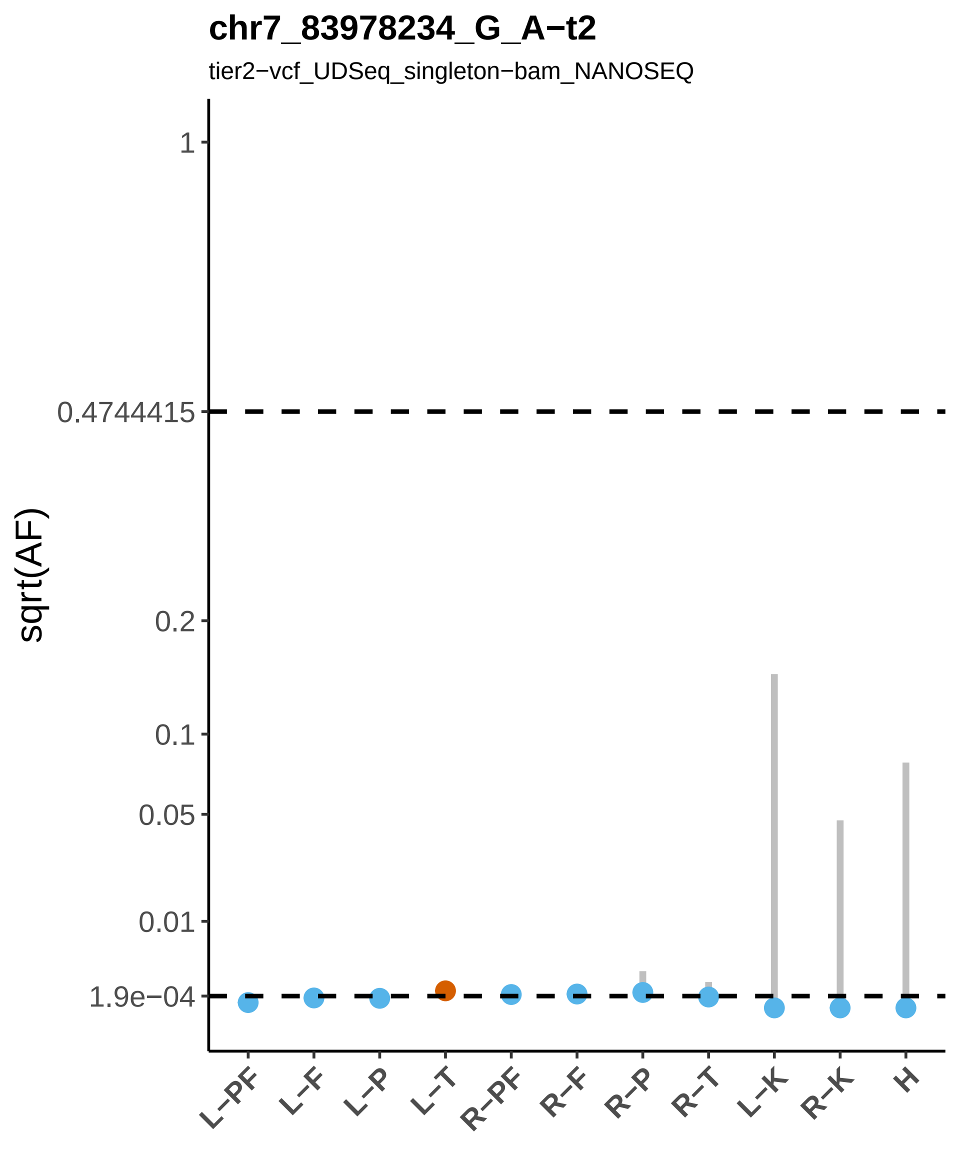

chr7_83978234_G_A−t2
tier2−vcf_UDSeq_singleton−bam_NANOSEQ
sqrt(AF)
1
0.4744415
0.2
0.1
0.05
0.01
1.9e−04
L−PF
L−F
L−P
L−T
R−PF
R−F
R−P
R−T
L−K
R−K
H

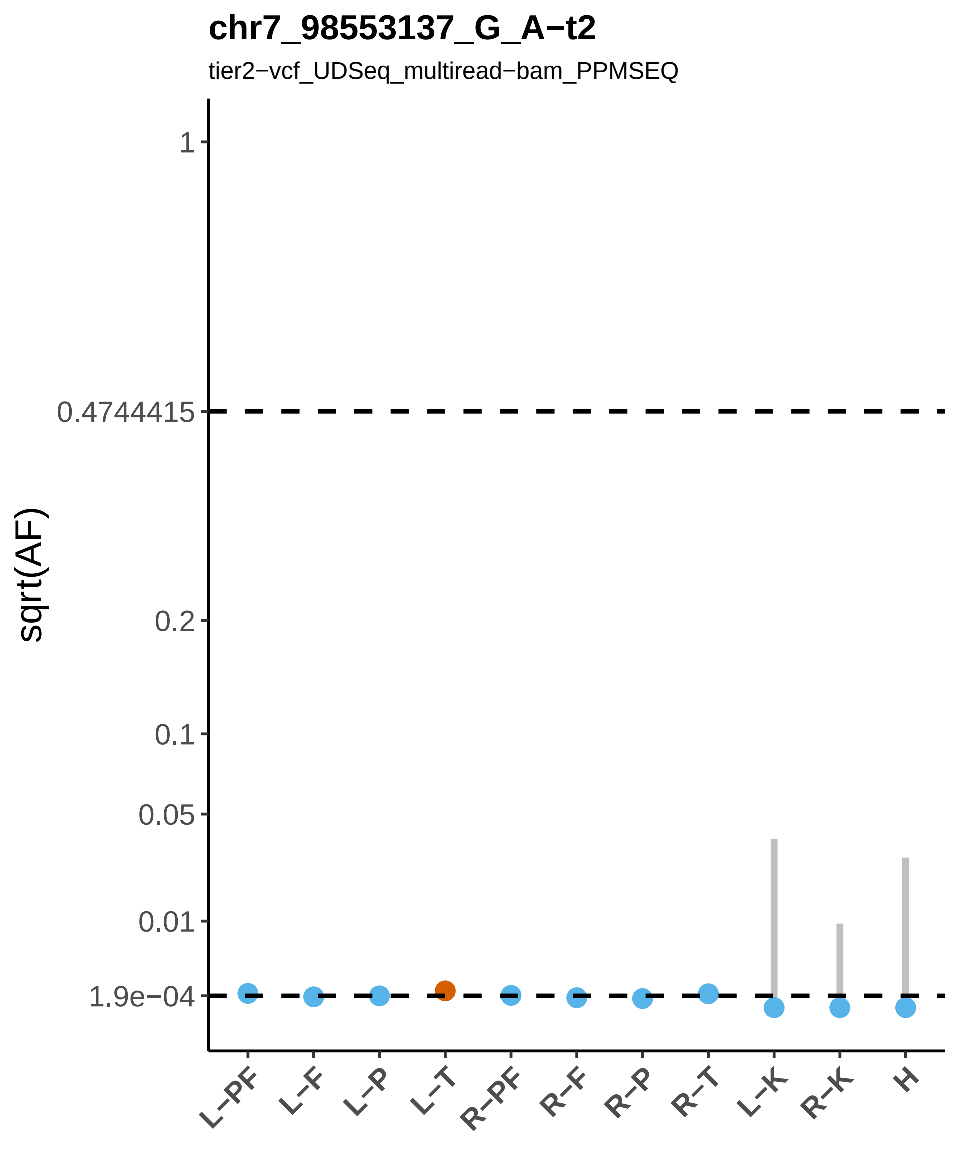
chr7_98553137_G_A−t2
tier2−vcf_UDSeq_multiread−bam_PPMSEQ
sqrt(AF)
1
0.4744415
0.2
0.1
0.05
0.01
1.9e−04
L−PF
L−F
L−P
L−T
R−PF
R−F
R−P
R−T
L−K
R−K
H

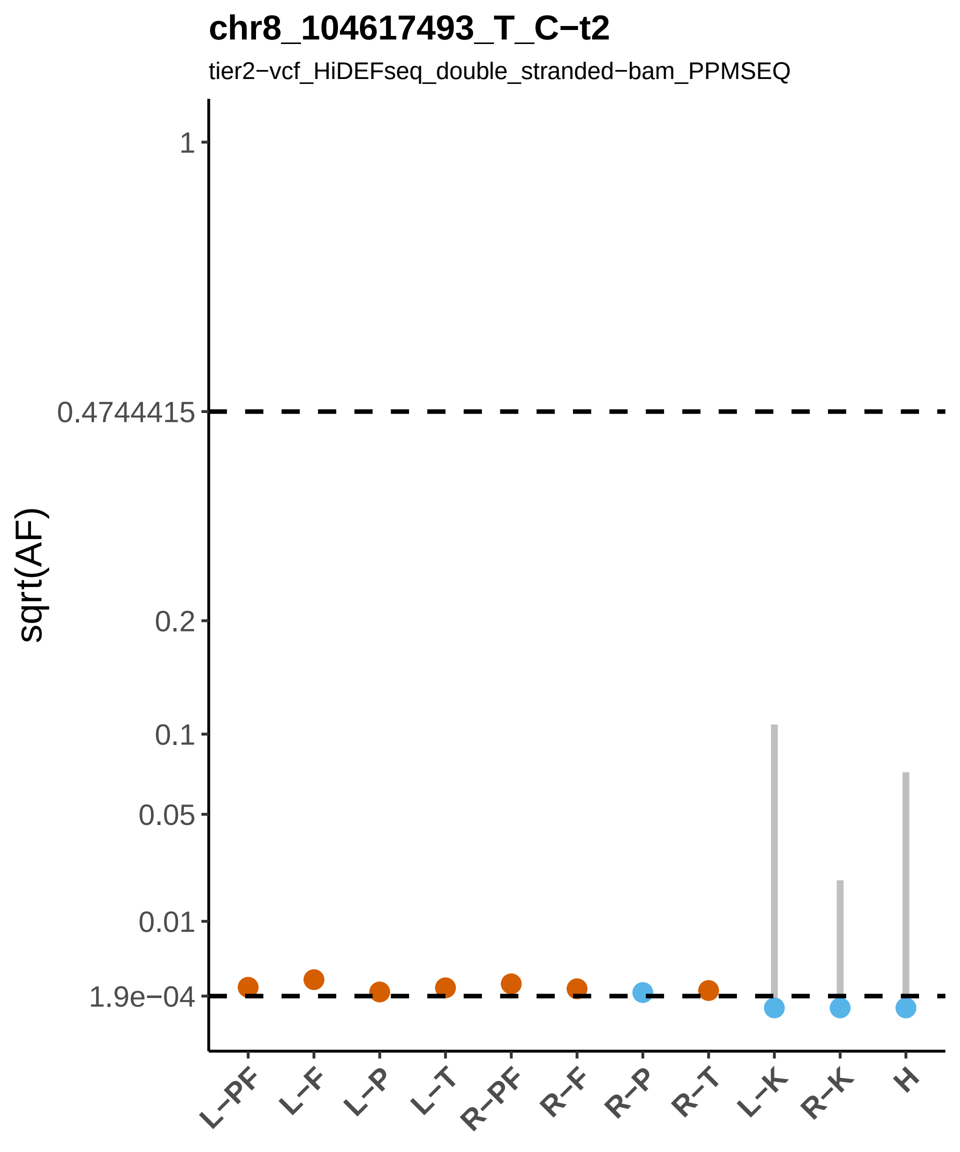
chr8_104617493_T_C−t2
tier2−vcf_HiDEFseq_double_stranded−bam_PPMSEQ
sqrt(AF)
1
0.4744415
0.2
0.1
0.05
0.01
1.9e−04
L−PF
L−F
L−P
L−T
R−PF
R−F
R−P
R−T
L−K
R−K
H

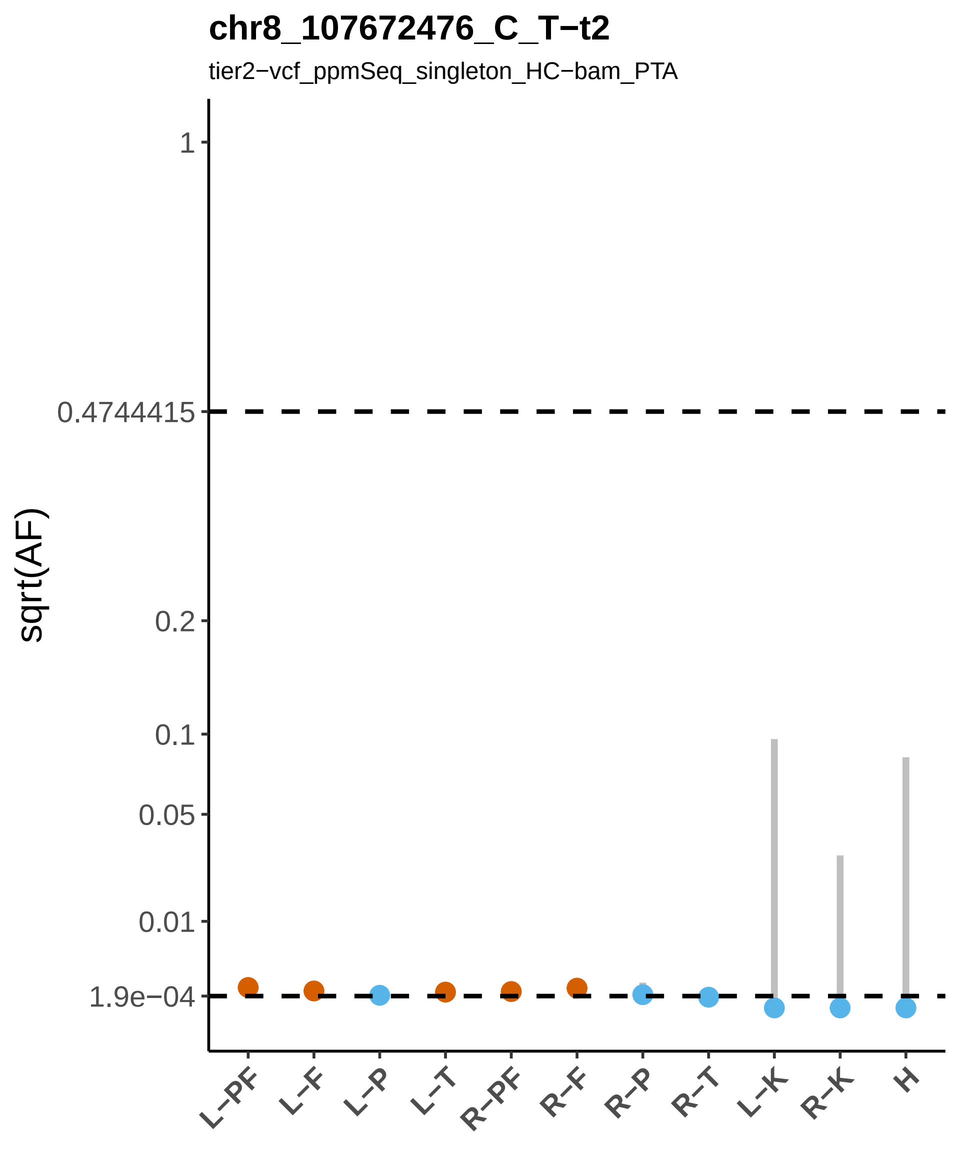
chr8_107672476_C_T−t2
tier2−vcf_ppmSeq_singleton_HC−bam_PTA
sqrt(AF)
1
0.4744415
0.2
0.1
0.05
0.01
1.9e−04
L−PF
L−F
L−P
L−T
R−PF
R−F
R−P
R−T
L−K
R−K
H

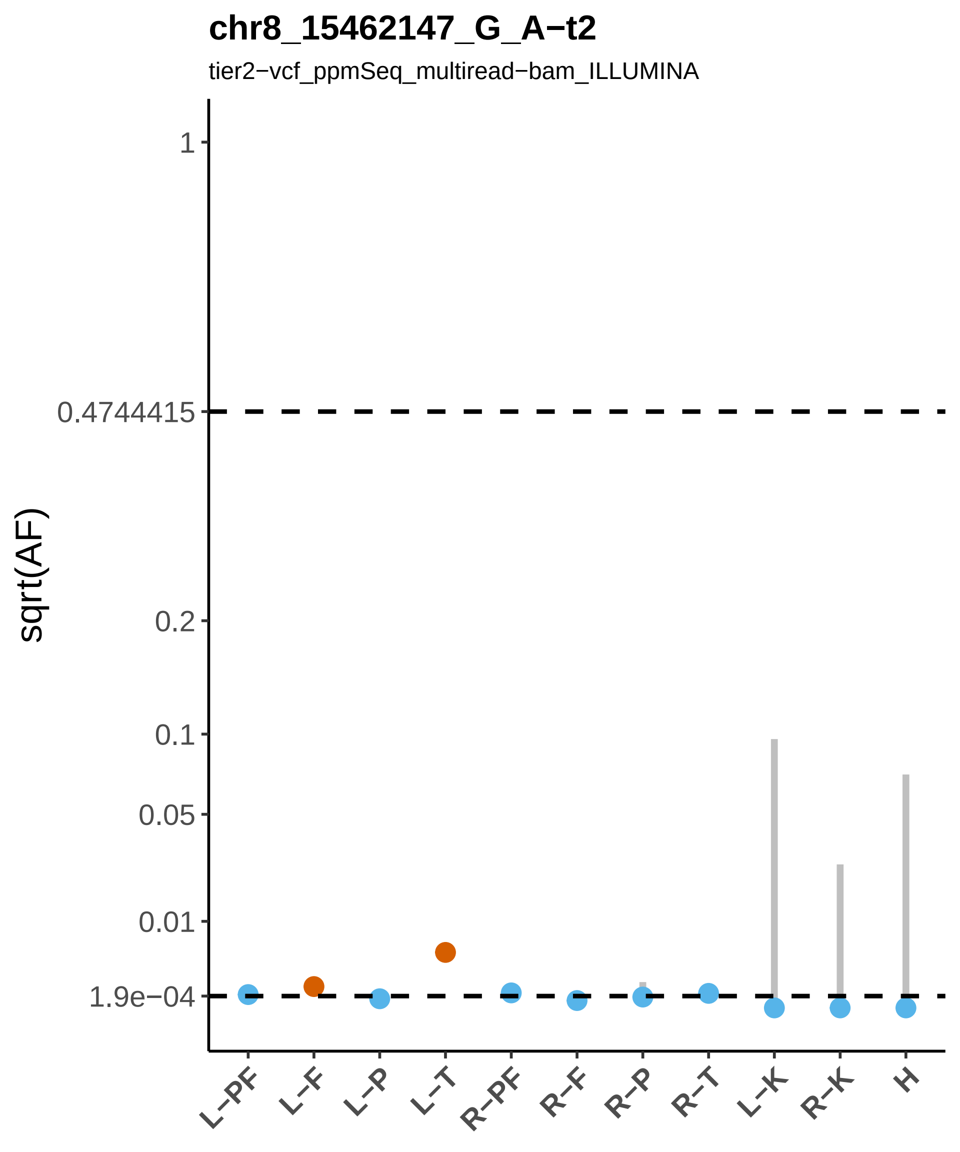
chr8_15462147_G_A−t2
tier2−vcf_ppmSeq_multiread−bam_ILLUMINA
sqrt(AF)
1
0.4744415
0.2
0.1
0.05
0.01
1.9e−04
L−PF
L−F
L−P
L−T
R−PF
R−F
R−P
R−T
L−K
R−K
H

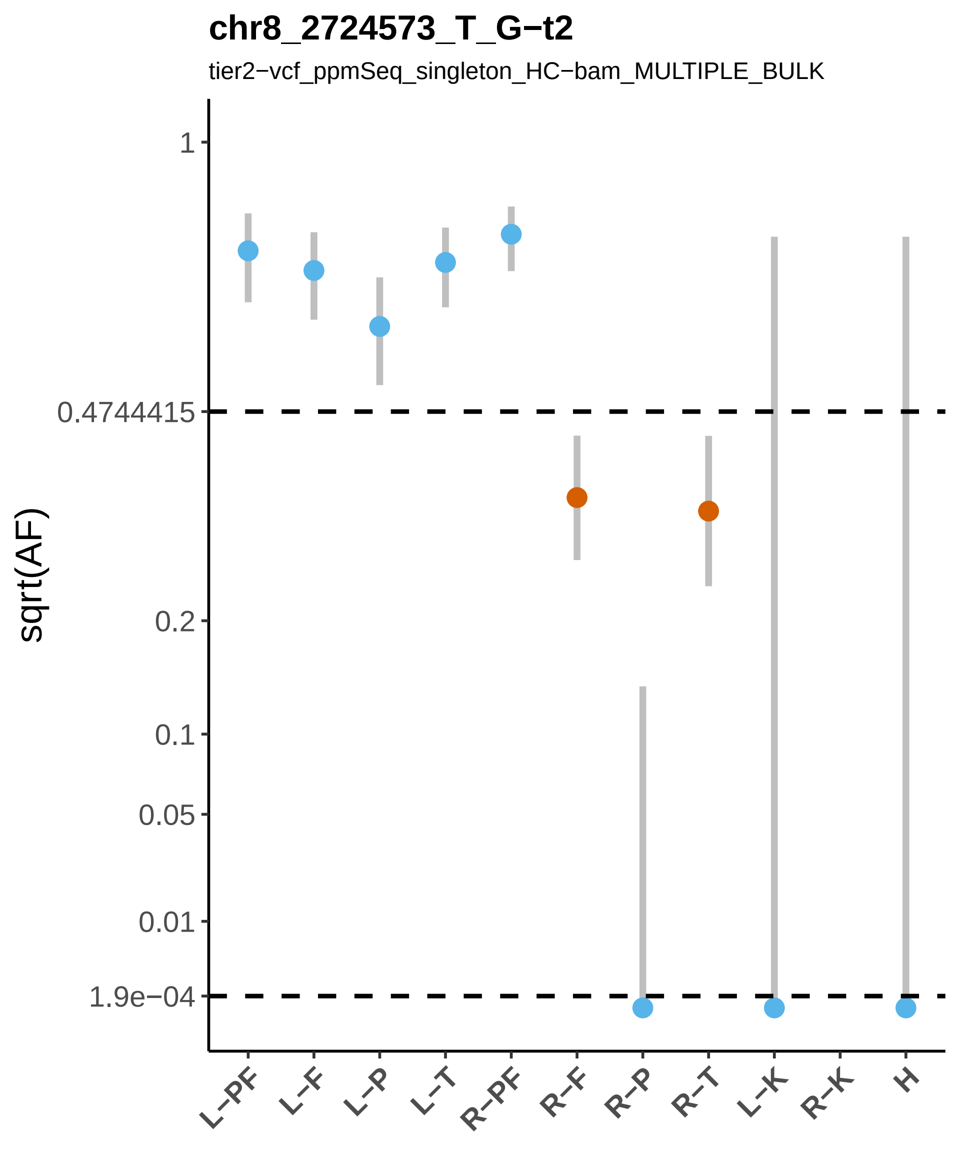
chr8_2724573_T_G−t2
tier2−vcf_ppmSeq_singleton_HC−bam_MULTIPLE_BULK
sqrt(AF)
1
0.4744415
0.2
0.1
0.05
0.01
1.9e−04
L−PF
L−F
L−P
L−T
R−PF
R−F
R−P
R−T
L−K
R−K
H

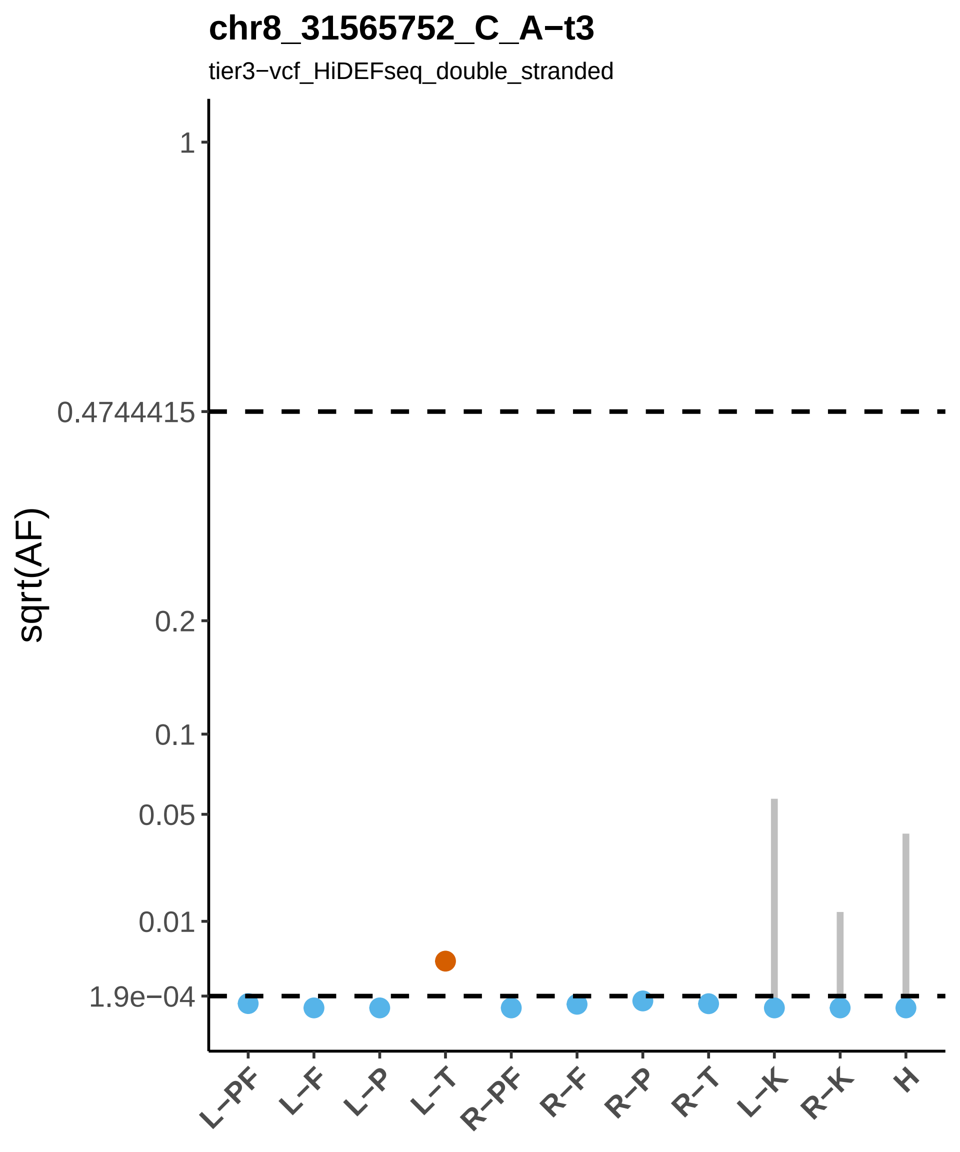
chr8_31565752_C_A−t3
tier3−vcf_HiDEFseq_double_stranded
sqrt(AF)
1
0.4744415
0.2
0.1
0.05
0.01
1.9e−04
L−PF
L−F
L−P
L−T
R−PF
R−F
R−P
R−T
L−K
R−K
H

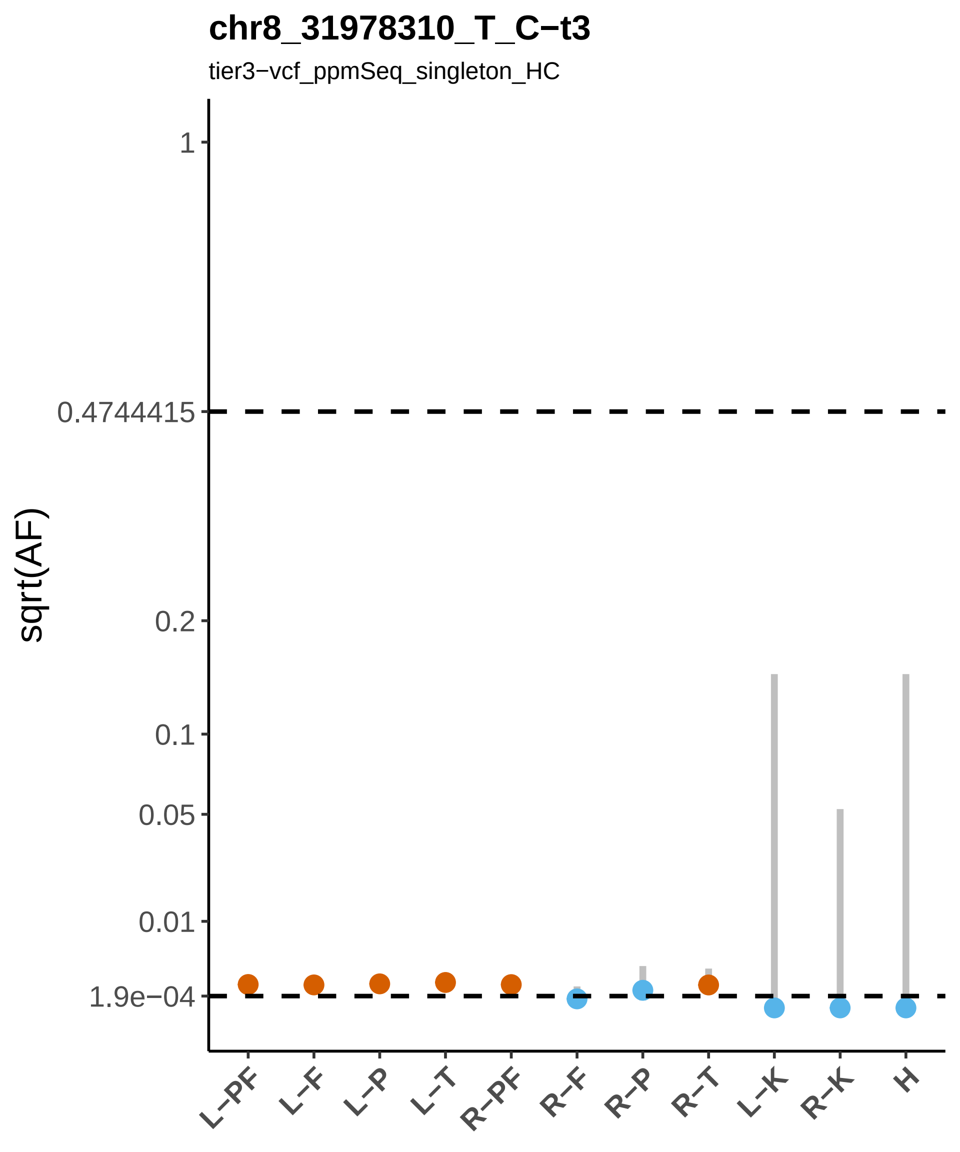
chr8_31978310_T_C−t3
tier3−vcf_ppmSeq_singleton_HC
sqrt(AF)
1
0.4744415
0.2
0.1
0.05
0.01
1.9e−04
L−PF
L−F
L−P
L−T
R−PF
R−F
R−P
R−T
L−K
R−K
H

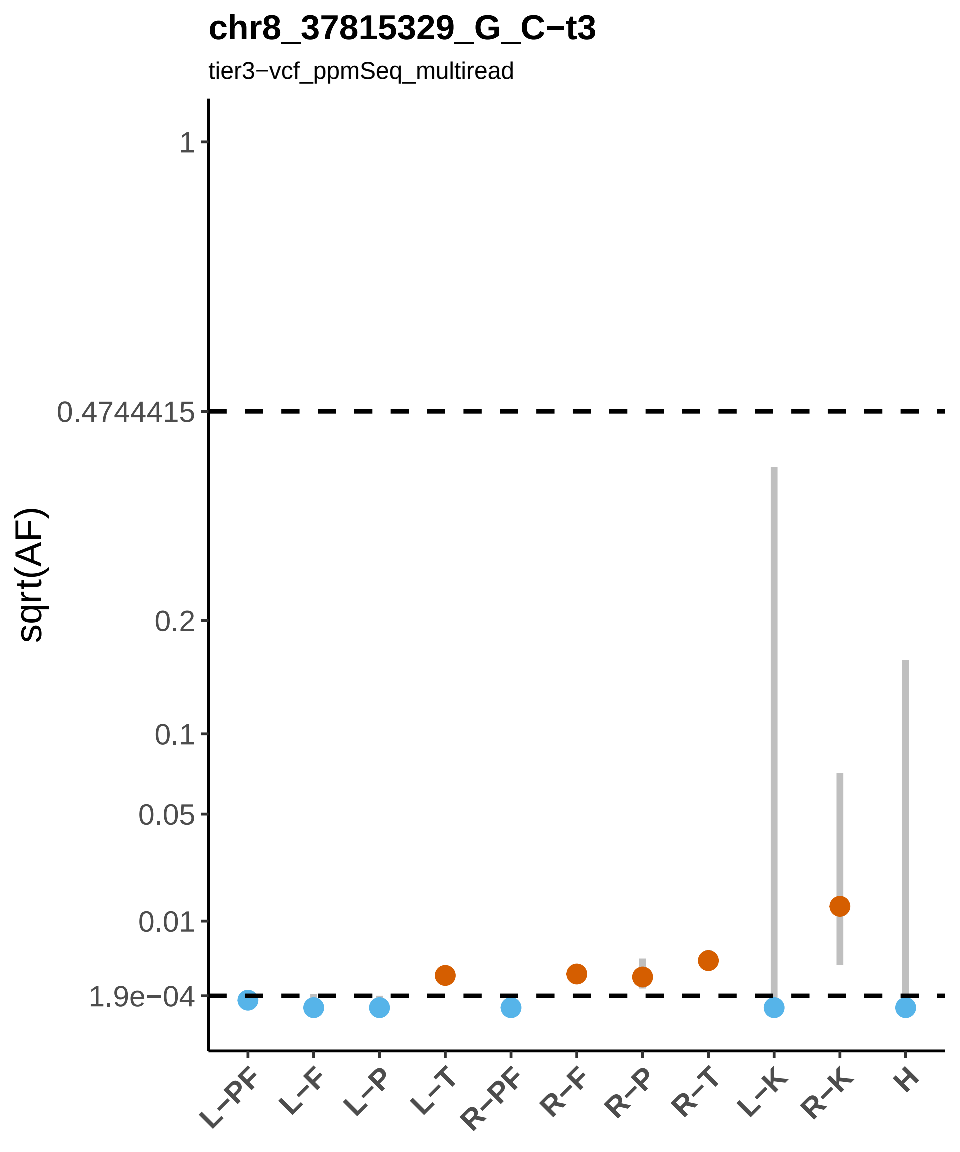

chr8_37815329_G_C−t3
tier3−vcf_ppmSeq_multiread
sqrt(AF)
1
0.4744415
0.2
0.1
0.05
0.01
1.9e−04
L−PF
L−F
L−P
L−T
R−PF
R−F
R−P
R−T
L−K
R−K
H

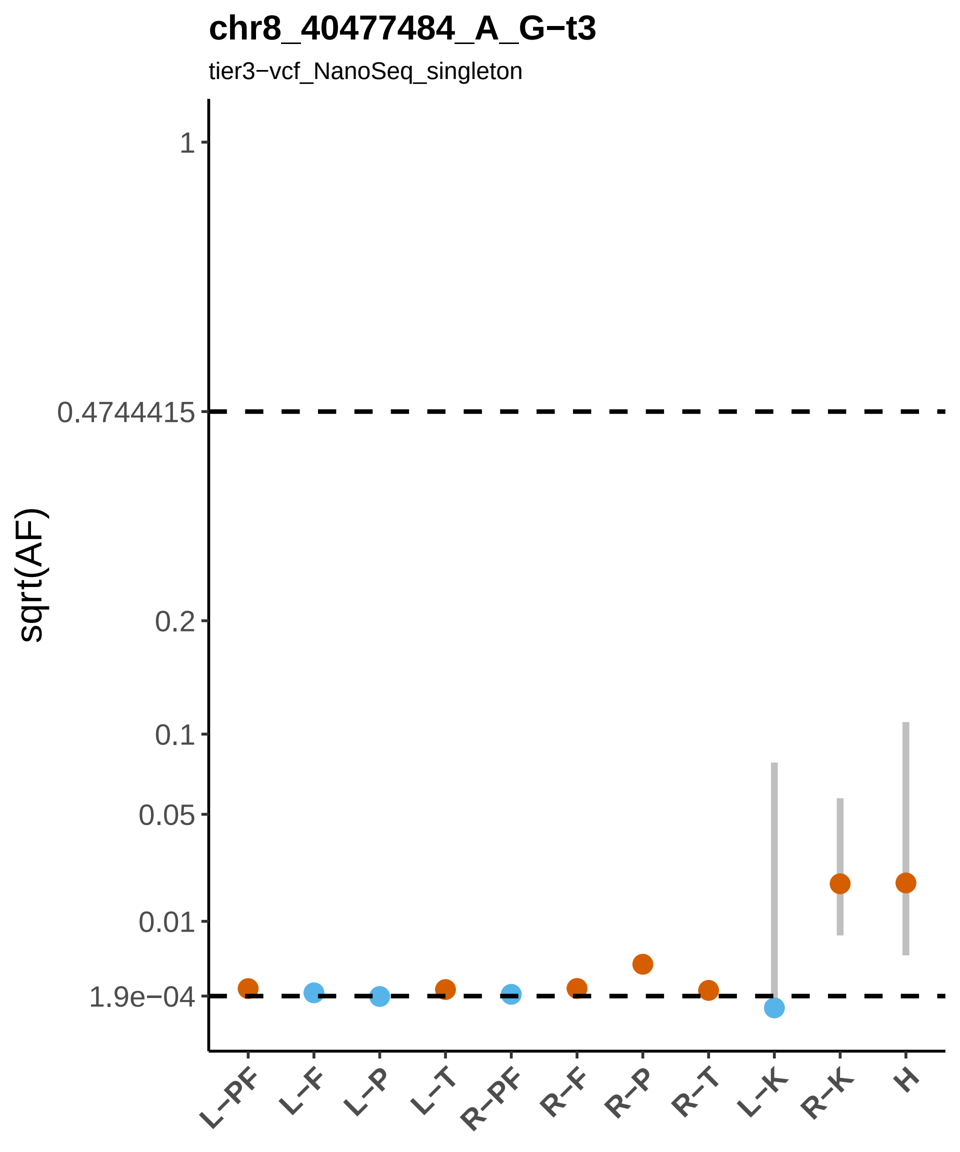
chr8_40477484_A_G−t3
tier3−vcf_NanoSeq_singleton
sqrt(AF)
1
0.4744415
0.2
0.1
0.05
0.01
1.9e−04
L−PF
L−F
L−P
L−T
R−PF
R−F
R−P
R−T
L−K
R−K
H

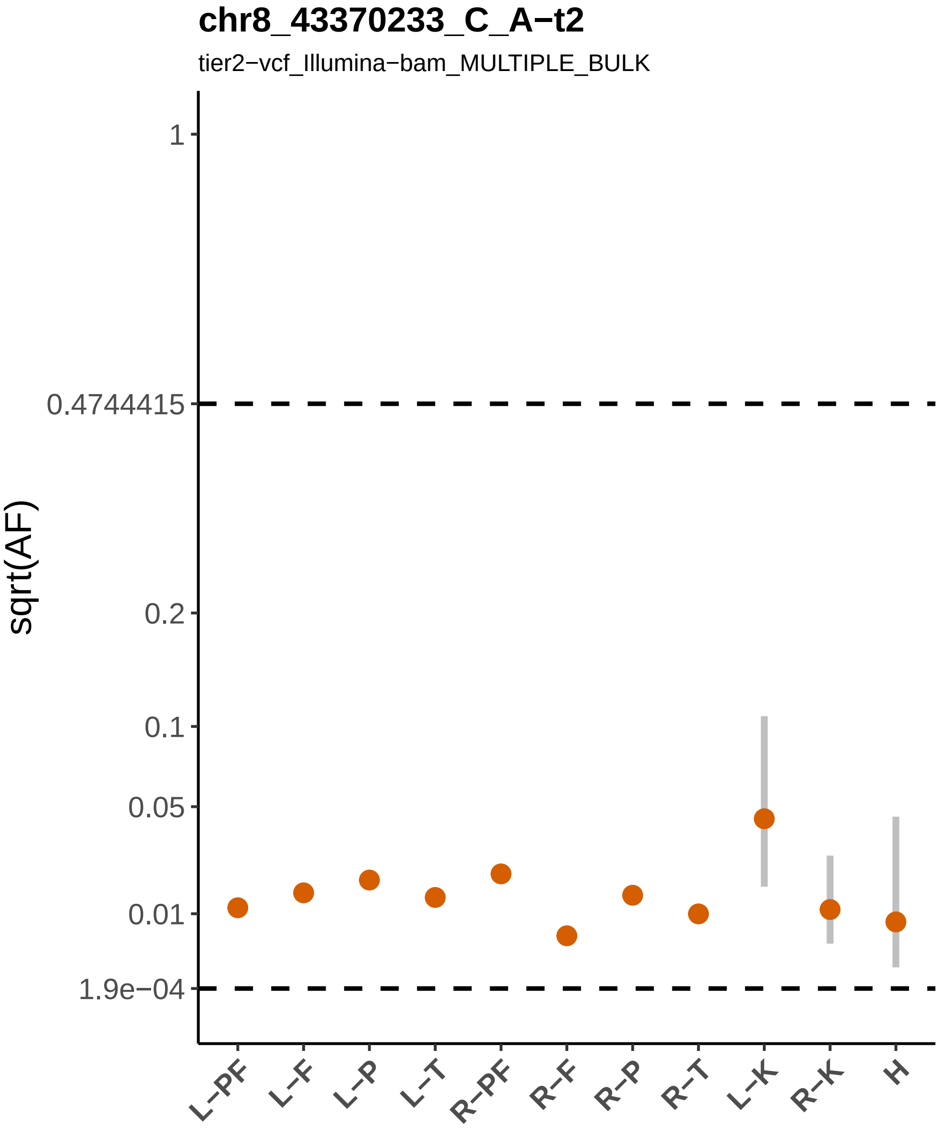
chr8_43370233_C_A−t2
tier2−vcf_Illumina−bam_MULTIPLE_BULK
sqrt(AF)
1
0.4744415
0.2
0.1
0.05
0.01
1.9e−04
L−PF
L−F
L−P
L−T
R−PF
R−F
R−P
R−T
L−K
R−K
H

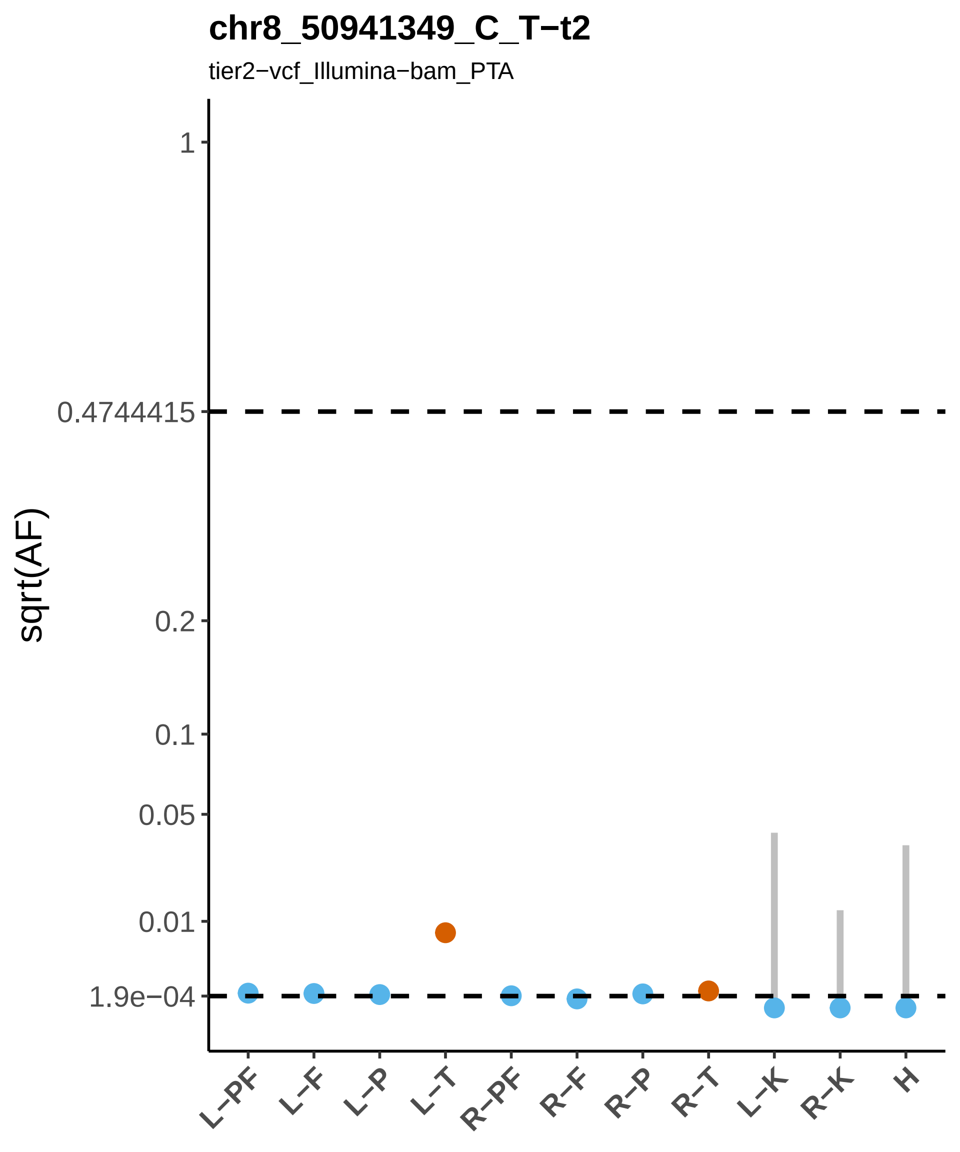

chr8_50941349_C_T−t2
tier2−vcf_Illumina−bam_PTA
sqrt(AF)
1
0.4744415
0.2
0.1
0.05
0.01
1.9e−04
L−PF
L−F
L−P
L−T
R−PF
R−F
R−P
R−T
L−K
R−K
H

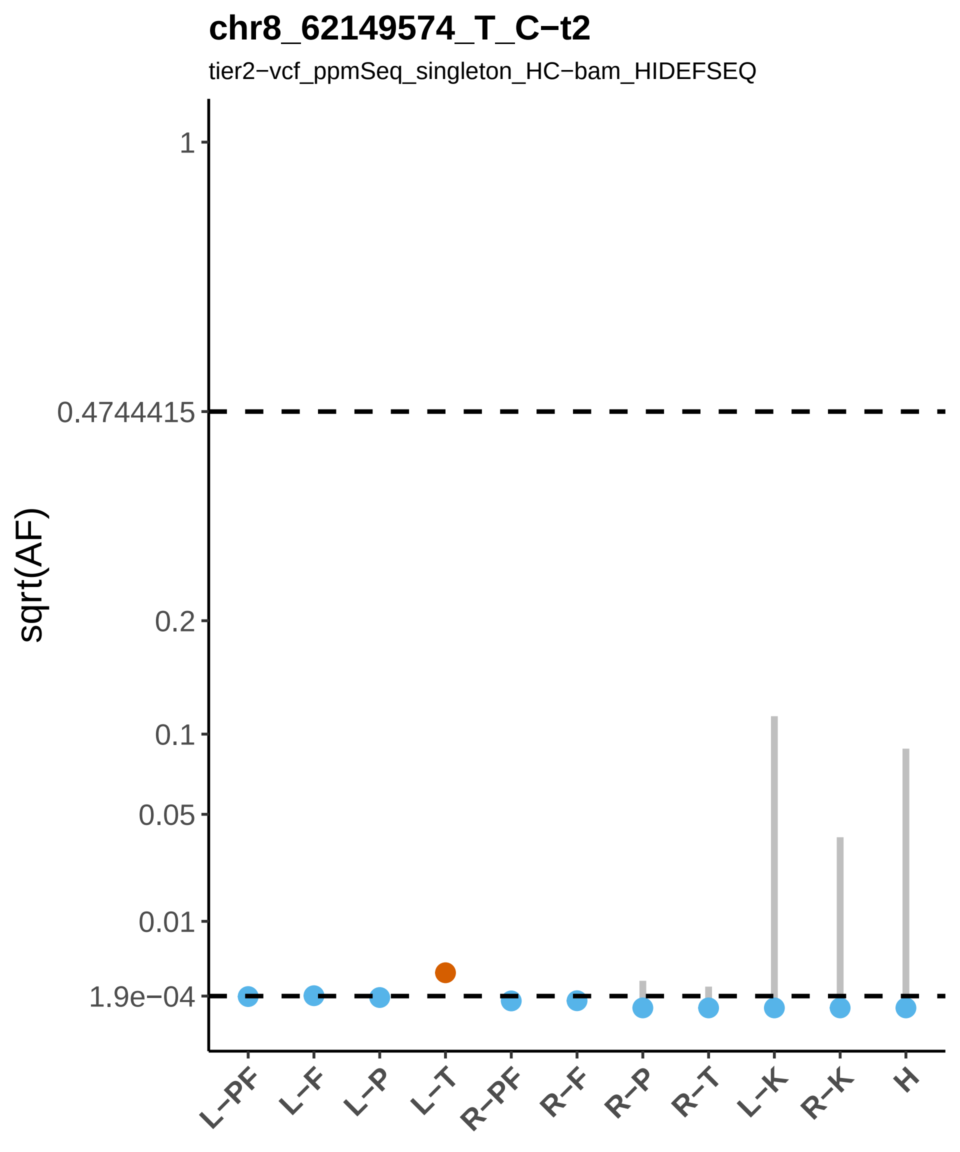
chr8_62149574_T_C−t2
tier2−vcf_ppmSeq_singleton_HC−bam_HIDEFSEQ
sqrt(AF)
1
0.4744415
0.2
0.1
0.05
0.01
1.9e−04
L−PF
L−F
L−P
L−T
R−PF
R−F
R−P
R−T
L−K
R−K
H

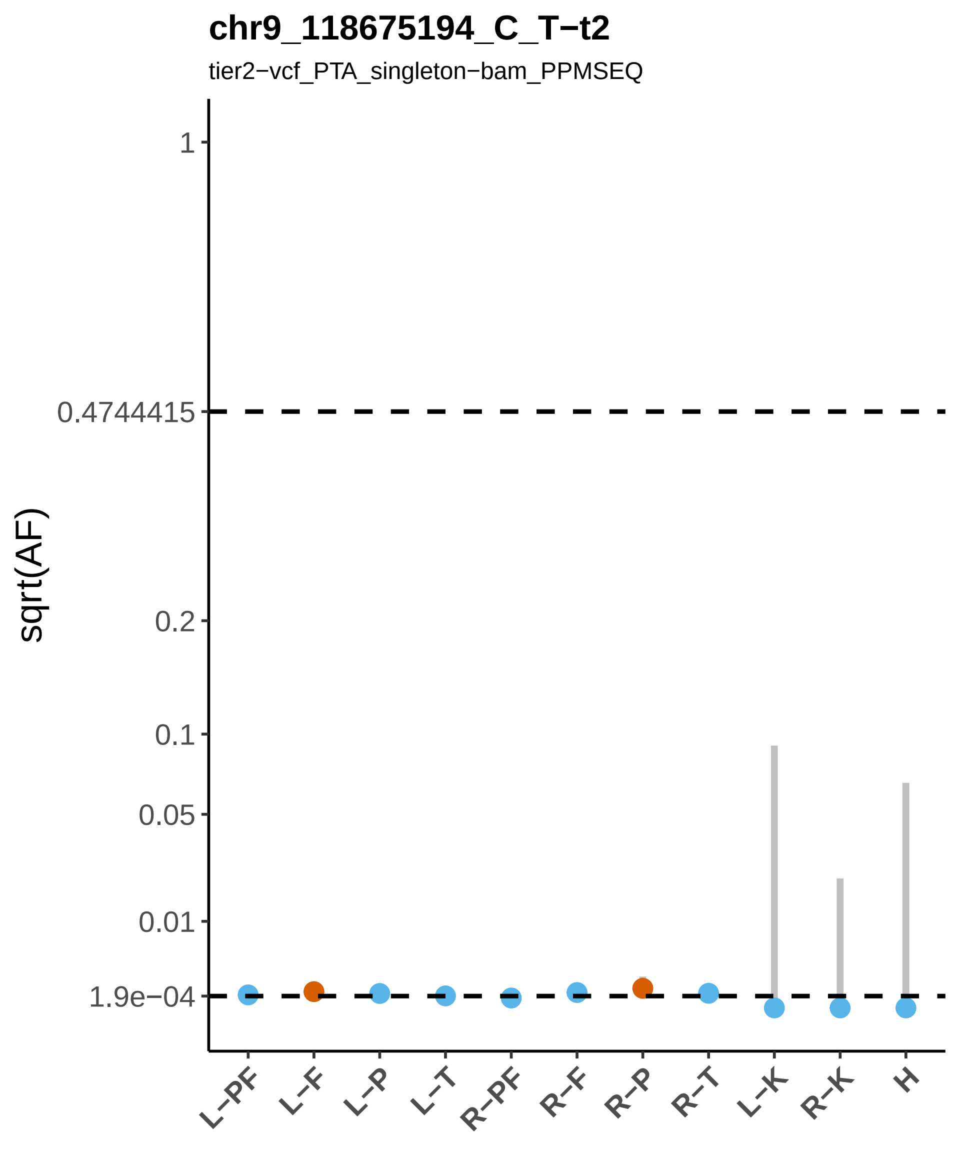
chr9_118675194_C_T−t2
tier2−vcf_PTA_singleton−bam_PPMSEQ
sqrt(AF)
1
0.4744415
0.2
0.1
0.05
0.01
1.9e−04
L−PF
L−F
L−P
L−T
R−PF
R−F
R−P
R−T
L−K
R−K
H

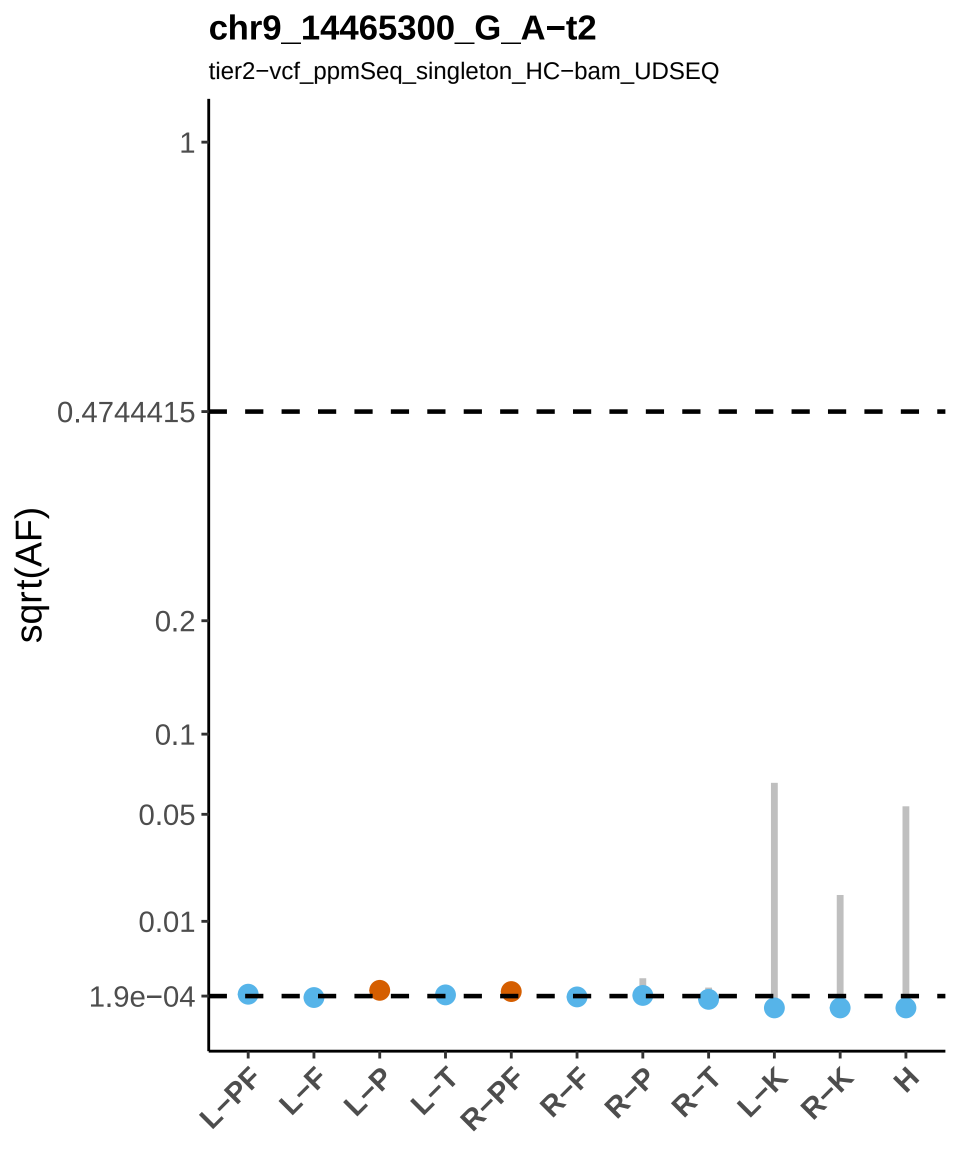

chr9_14465300_G_A−t2
tier2−vcf_ppmSeq_singleton_HC−bam_UDSEQ
sqrt(AF)
1
0.4744415
0.2
0.1
0.05
0.01
1.9e−04
L−PF
L−F
L−P
L−T
R−PF
R−F
R−P
R−T
L−K
R−K
H

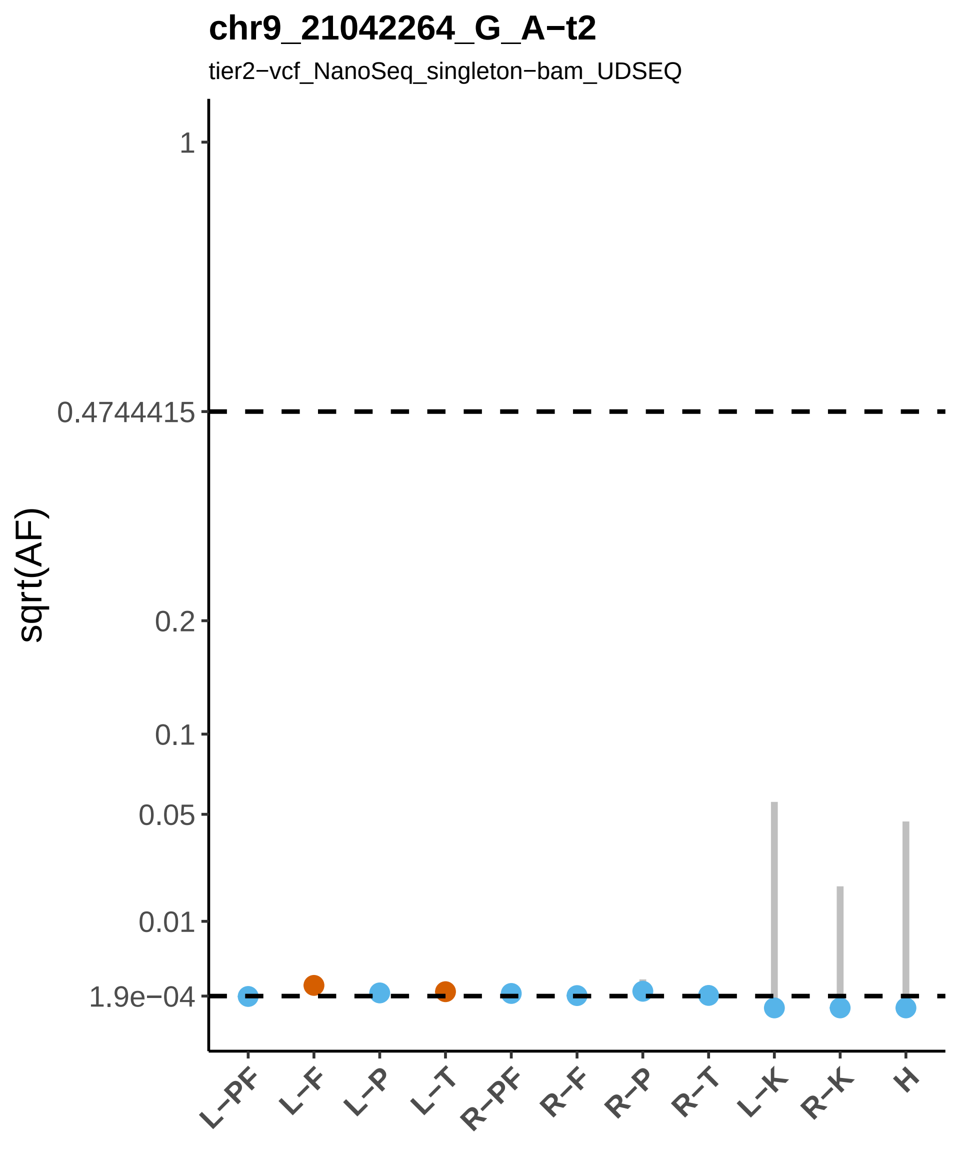

chr9_21042264_G_A−t2
tier2−vcf_NanoSeq_singleton−bam_UDSEQ
sqrt(AF)
1
0.4744415
0.2
0.1
0.05
0.01
1.9e−04
L−PF
L−F
L−P
L−T
R−PF
R−F
R−P
R−T
L−K
R−K
H

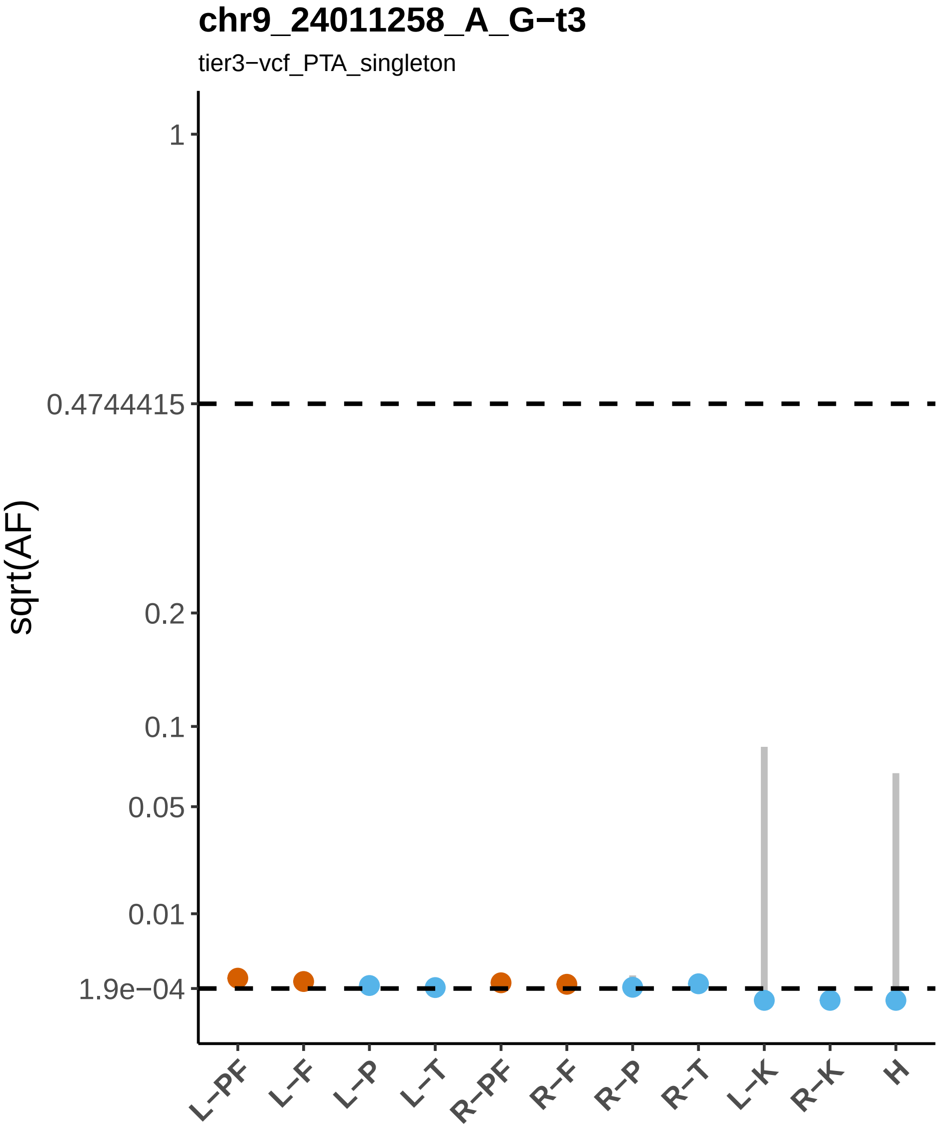

chr9_24011258_A_G−t3
tier3−vcf_PTA_singleton
sqrt(AF)
1
0.4744415
0.2
0.1
0.05
0.01
1.9e−04
L−PF
L−F
L−P
L−T
R−PF
R−F
R−P
R−T
L−K
R−K
H

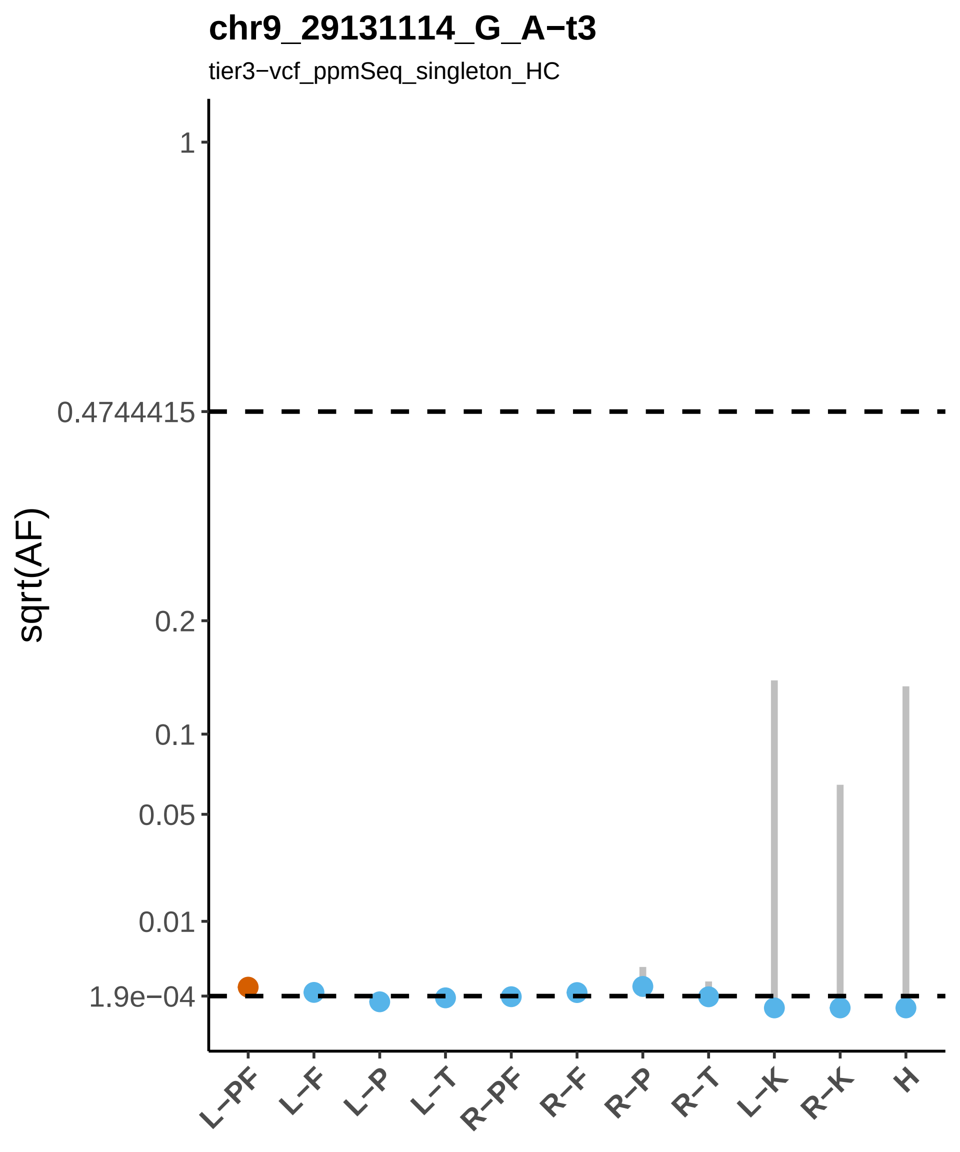

chr9_29131114_G_A−t3
tier3−vcf_ppmSeq_singleton_HC
sqrt(AF)
1
0.4744415
0.2
0.1
0.05
0.01
1.9e−04
L−PF
L−F
L−P
L−T
R−PF
R−F
R−P
R−T
L−K
R−K
H

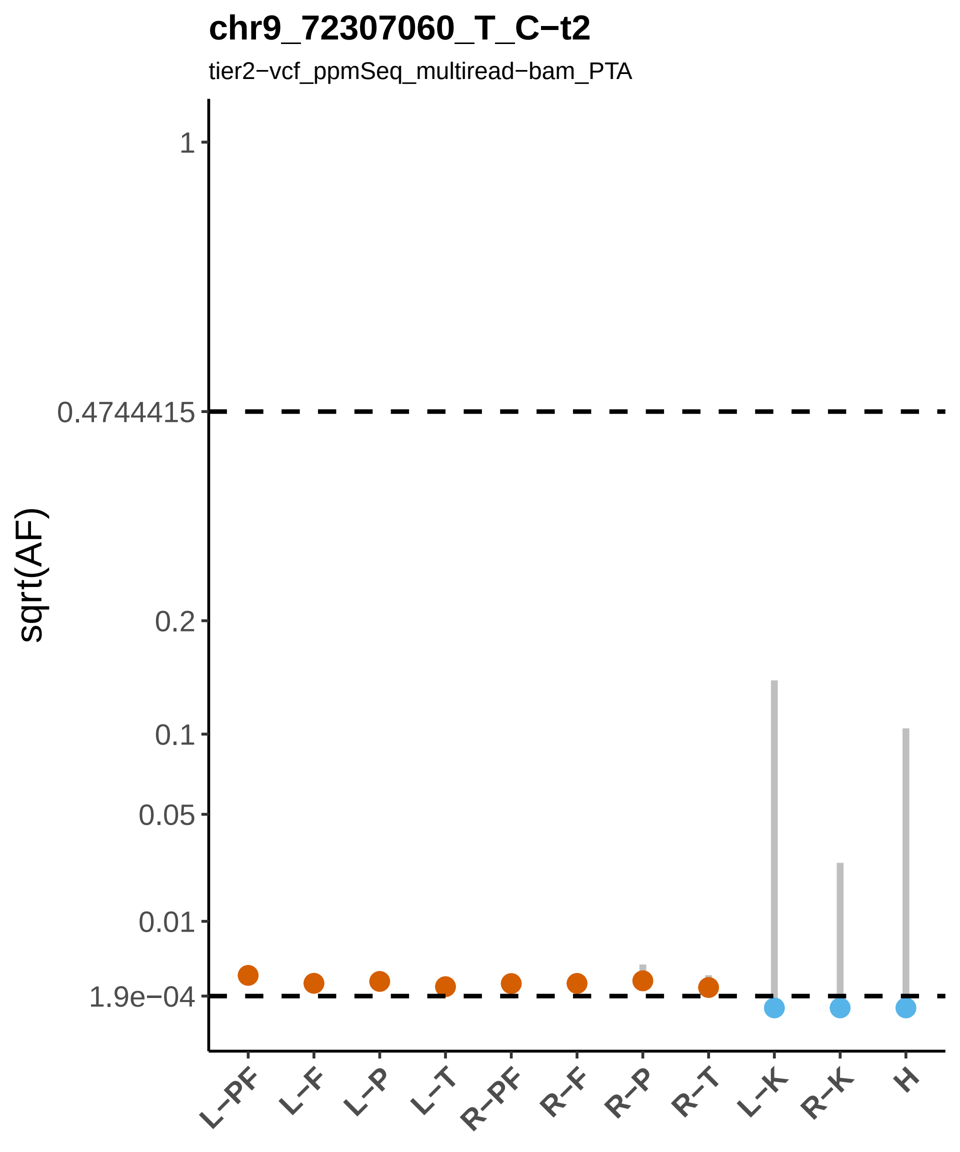

chr9_72307060_T_C−t2
tier2−vcf_ppmSeq_multiread−bam_PTA
sqrt(AF)
1
0.4744415
0.2
0.1
0.05
0.01
1.9e−04
L−PF
L−F
L−P
L−T
R−PF
R−F
R−P
R−T
L−K
R−K
H

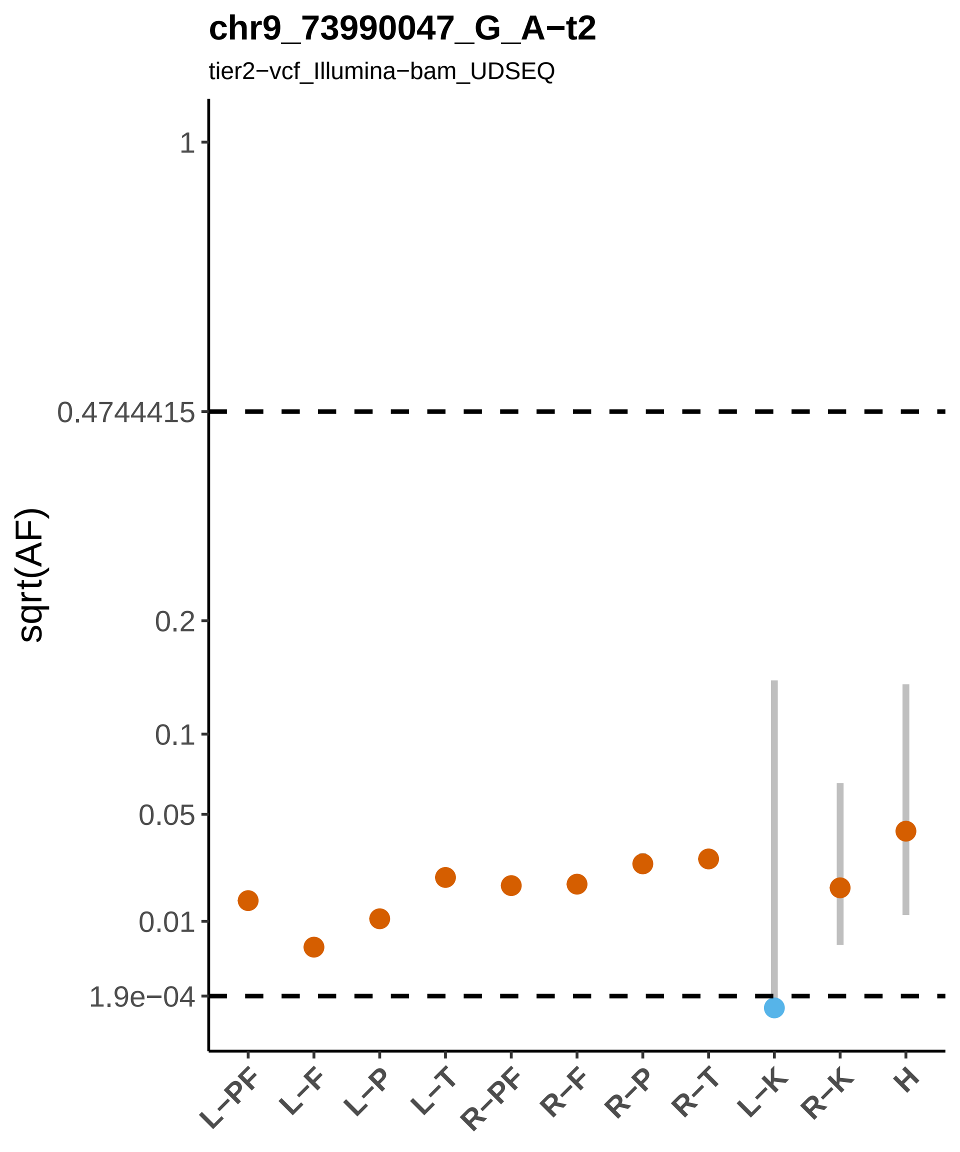

chr9_73990047_G_A−t2
tier2−vcf_Illumina−bam_UDSEQ
sqrt(AF)
1
0.4744415
0.2
0.1
0.05
0.01
1.9e−04
L−PF
L−F
L−P
L−T
R−PF
R−F
R−P
R−T
L−K
R−K
H

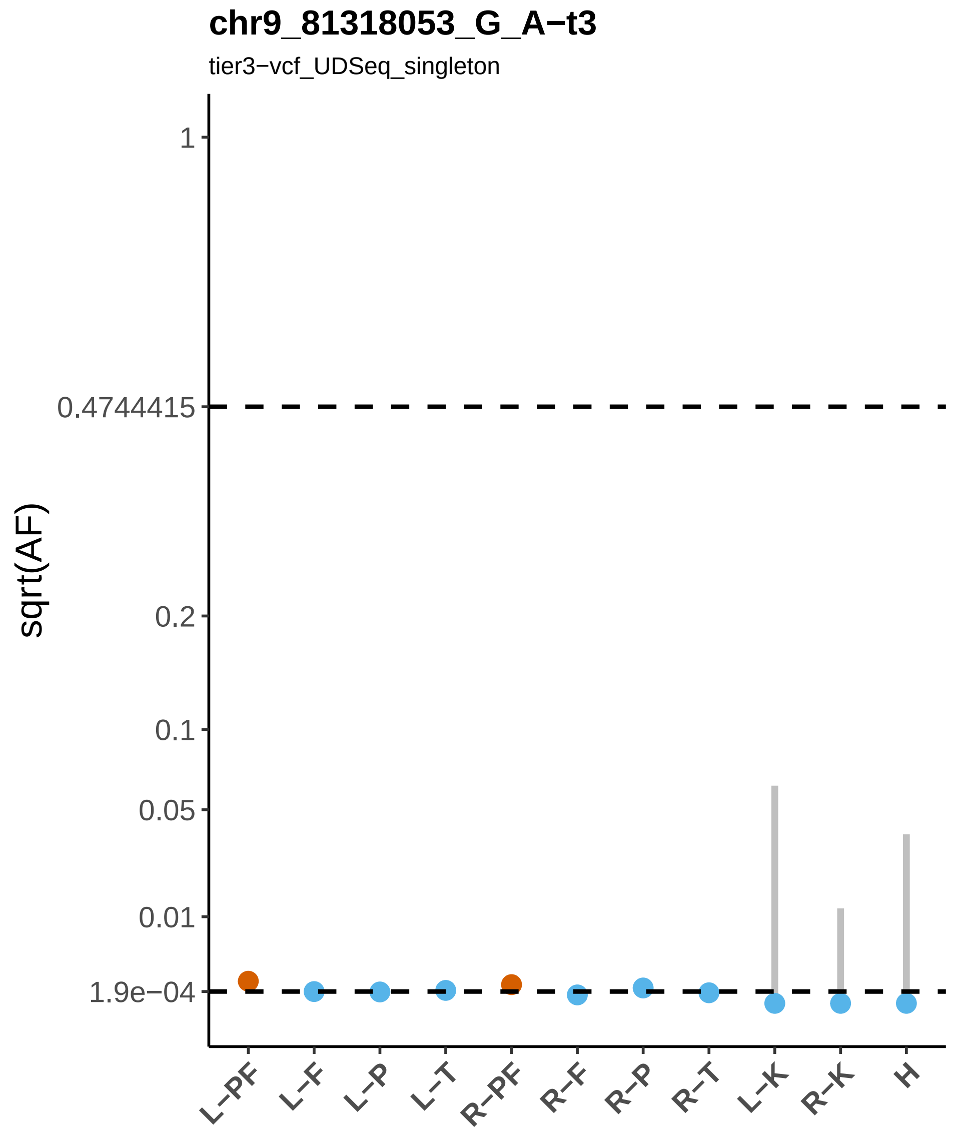
chr9_81318053_G_A−t3
tier3−vcf_UDSeq_singleton
sqrt(AF)
1
0.4744415
0.2
0.1
0.05
0.01
1.9e−04
L−PF
L−F
L−P
L−T
R−PF
R−F
R−P
R−T
L−K
R−K
H

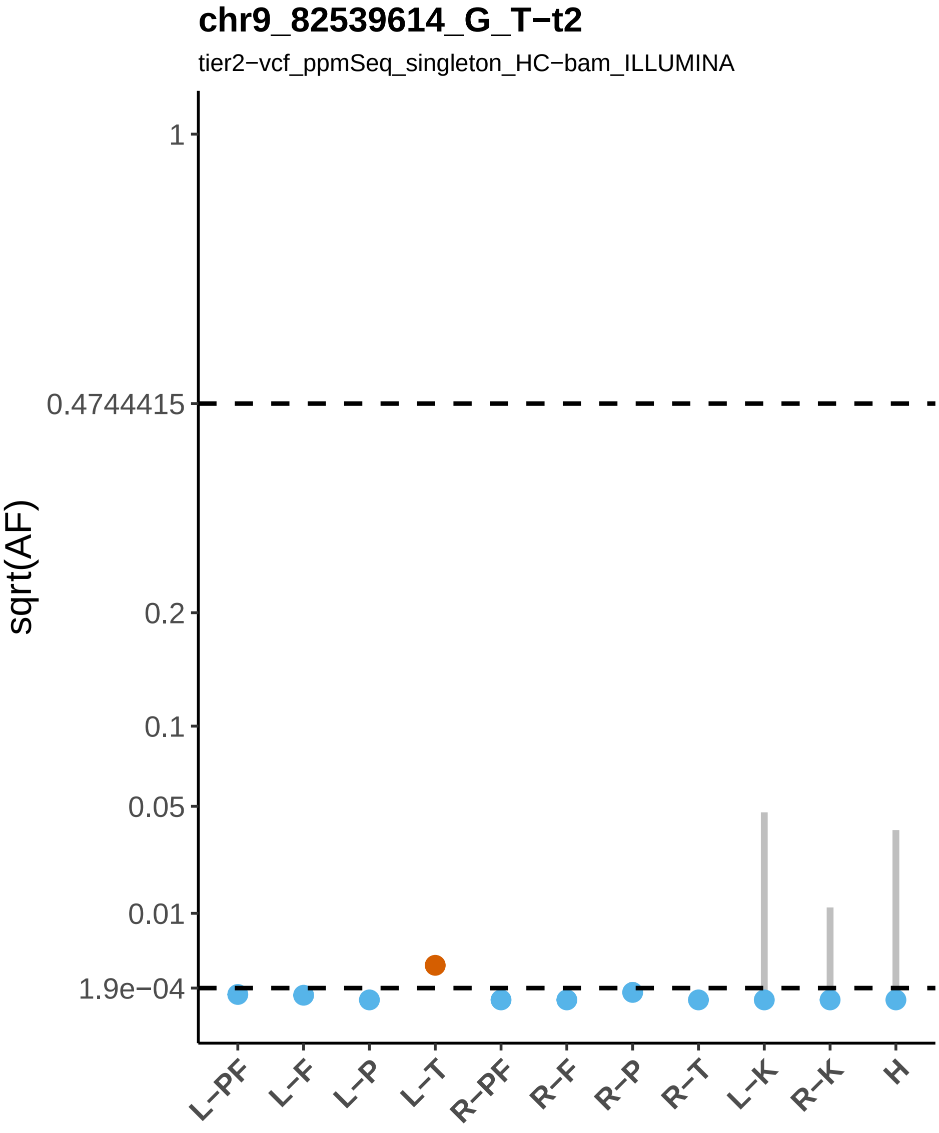
chr9_82539614_G_T−t2
tier2−vcf_ppmSeq_singleton_HC−bam_ILLUMINA
sqrt(AF)
1
0.4744415
0.2
0.1
0.05
0.01
1.9e−04
L−PF
L−F
L−P
L−T
R−PF
R−F
R−P
R−T
L−K
R−K
H

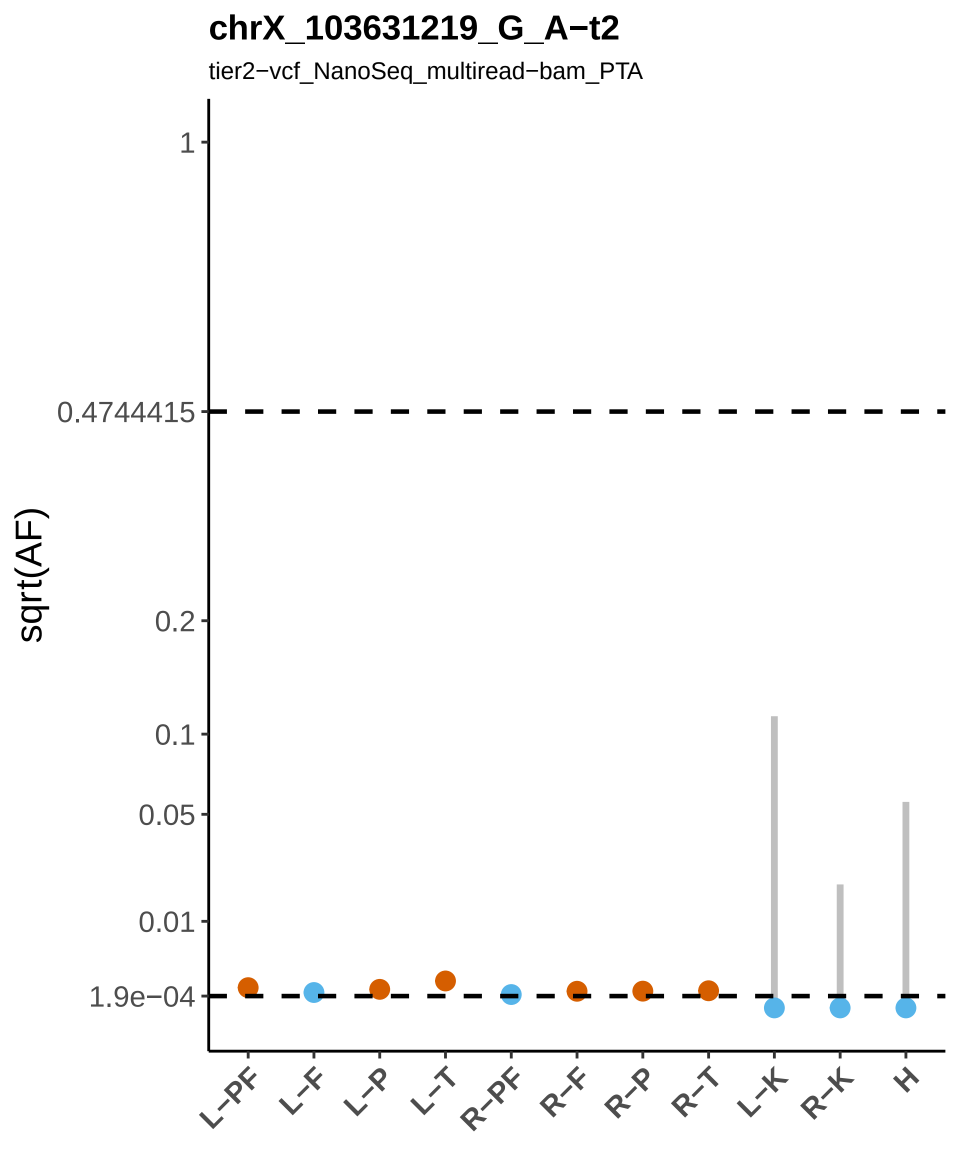
chrX_103631219_G_A−t2
tier2−vcf_NanoSeq_multiread−bam_PTA
sqrt(AF)
1
0.4744415
0.2
0.1
0.05
0.01
1.9e−04
L−PF
L−F
L−P
L−T
R−PF
R−F
R−P
R−T
L−K
R−K
H

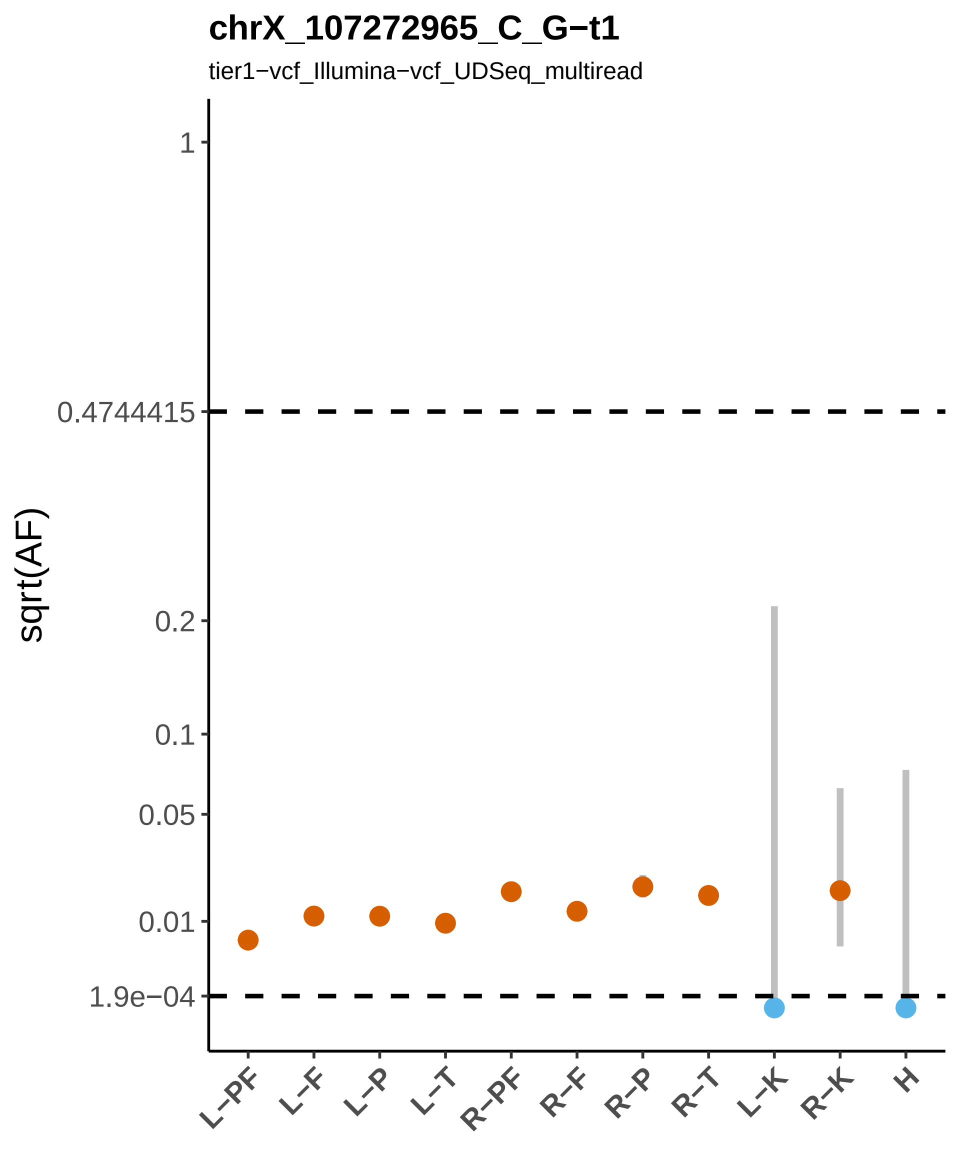
chrX_107272965_C_G−t1
tier1−vcf_Illumina−vcf_UDSeq_multiread
sqrt(AF)
1
0.4744415
0.2
0.1
0.05
0.01
1.9e−04
L−PF
L−F
L−P
L−T
R−PF
R−F
R−P
R−T
L−K
R−K
H

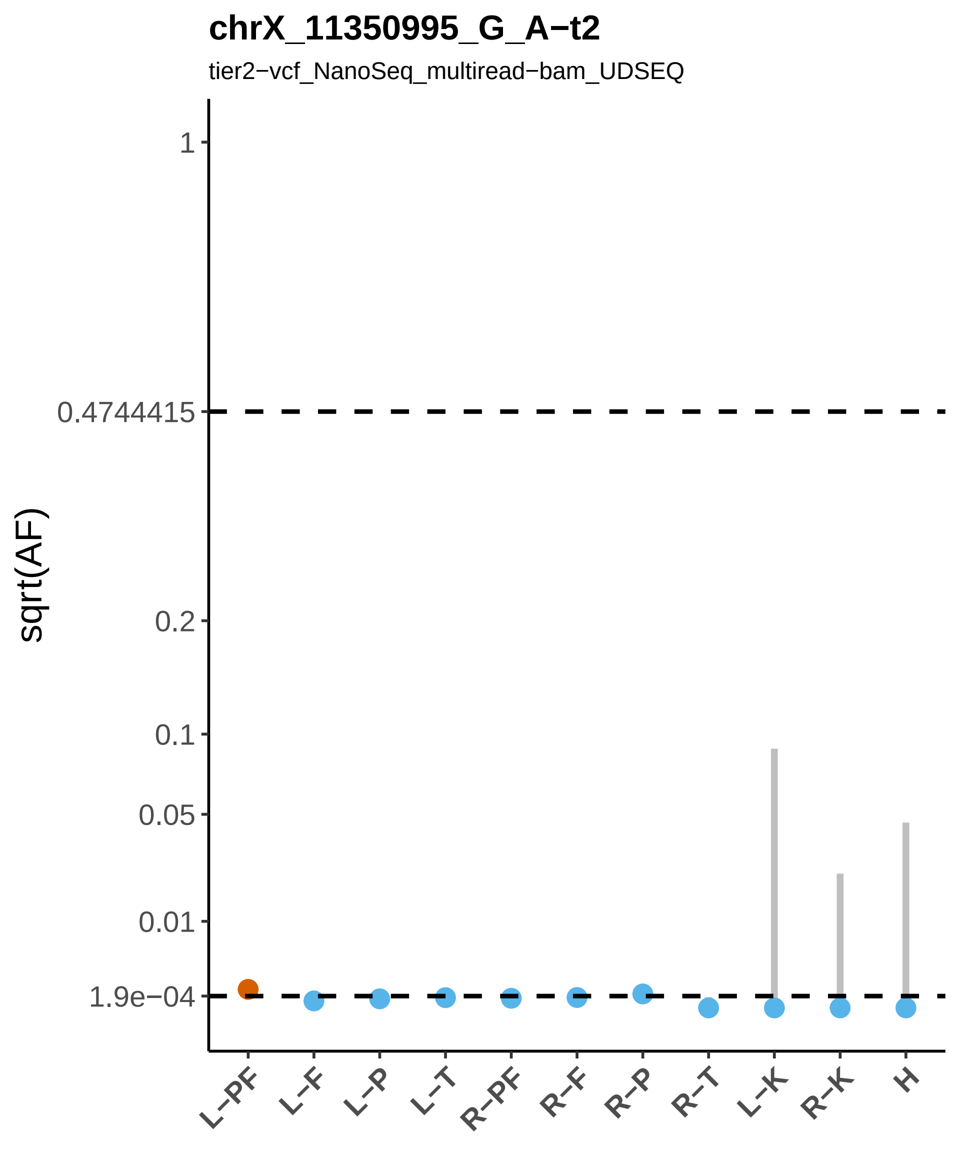

chrX_11350995_G_A−t2
tier2−vcf_NanoSeq_multiread−bam_UDSEQ
sqrt(AF)
1
0.4744415
0.2
0.1
0.05
0.01
1.9e−04
L−PF
L−F
L−P
L−T
R−PF
R−F
R−P
R−T
L−K
R−K
H

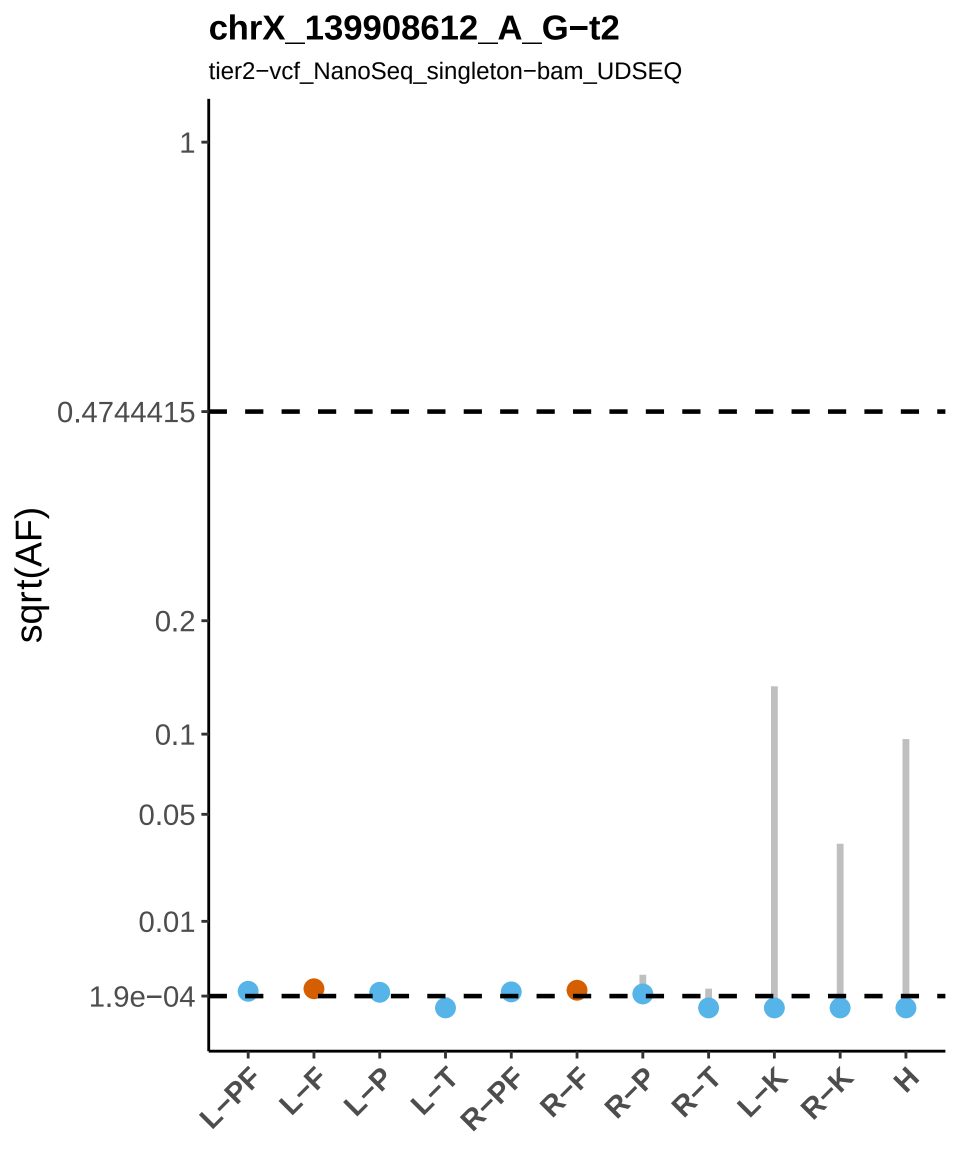
chrX_139908612_A_G−t2
tier2−vcf_NanoSeq_singleton−bam_UDSEQ
sqrt(AF)
1
0.4744415
0.2
0.1
0.05
0.01
1.9e−04
L−PF
L−F
L−P
L−T
R−PF
R−F
R−P
R−T
L−K
R−K
H

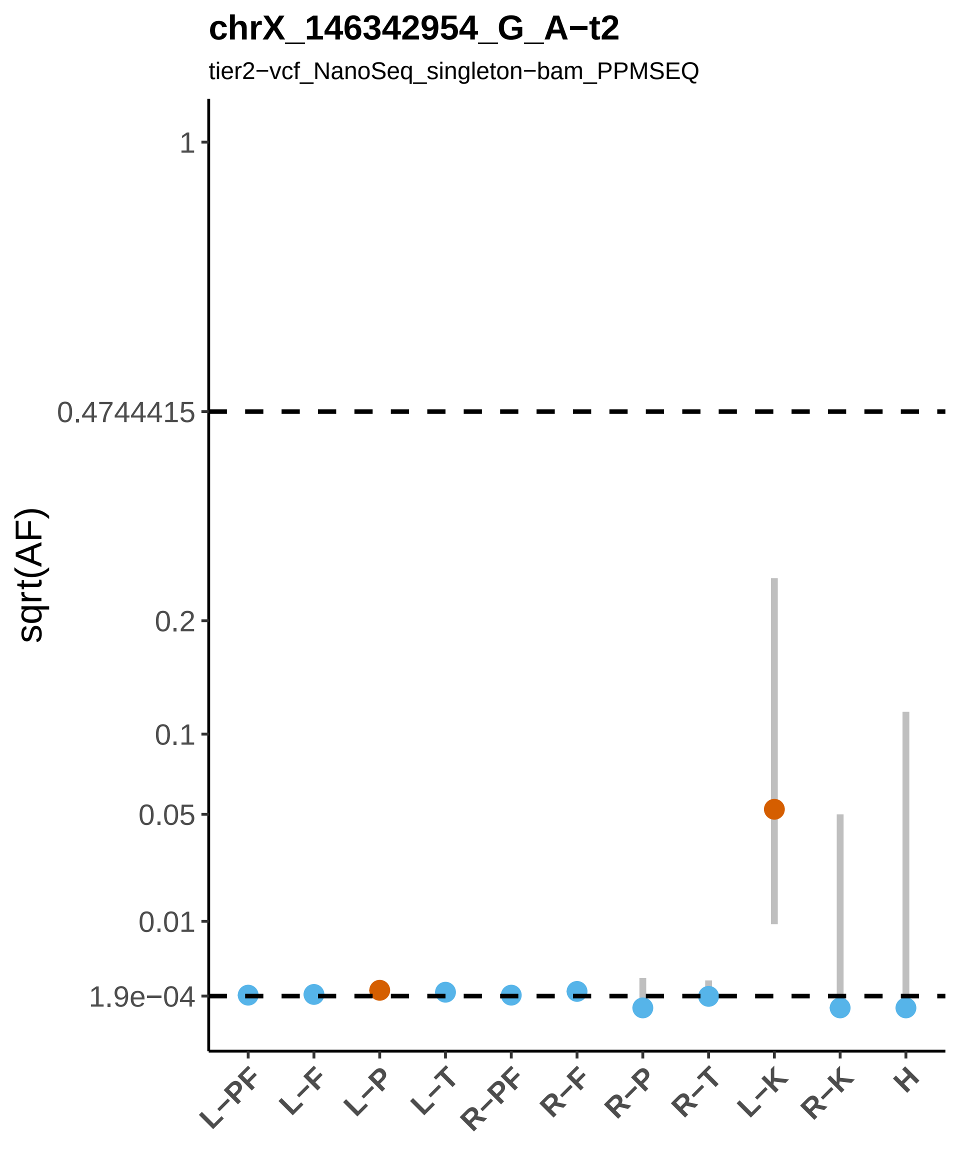

chrX_146342954_G_A−t2
tier2−vcf_NanoSeq_singleton−bam_PPMSEQ
sqrt(AF)
1
0.4744415
0.2
0.1
0.05
0.01
1.9e−04
L−PF
L−F
L−P
L−T
R−PF
R−F
R−P
R−T
L−K
R−K
H

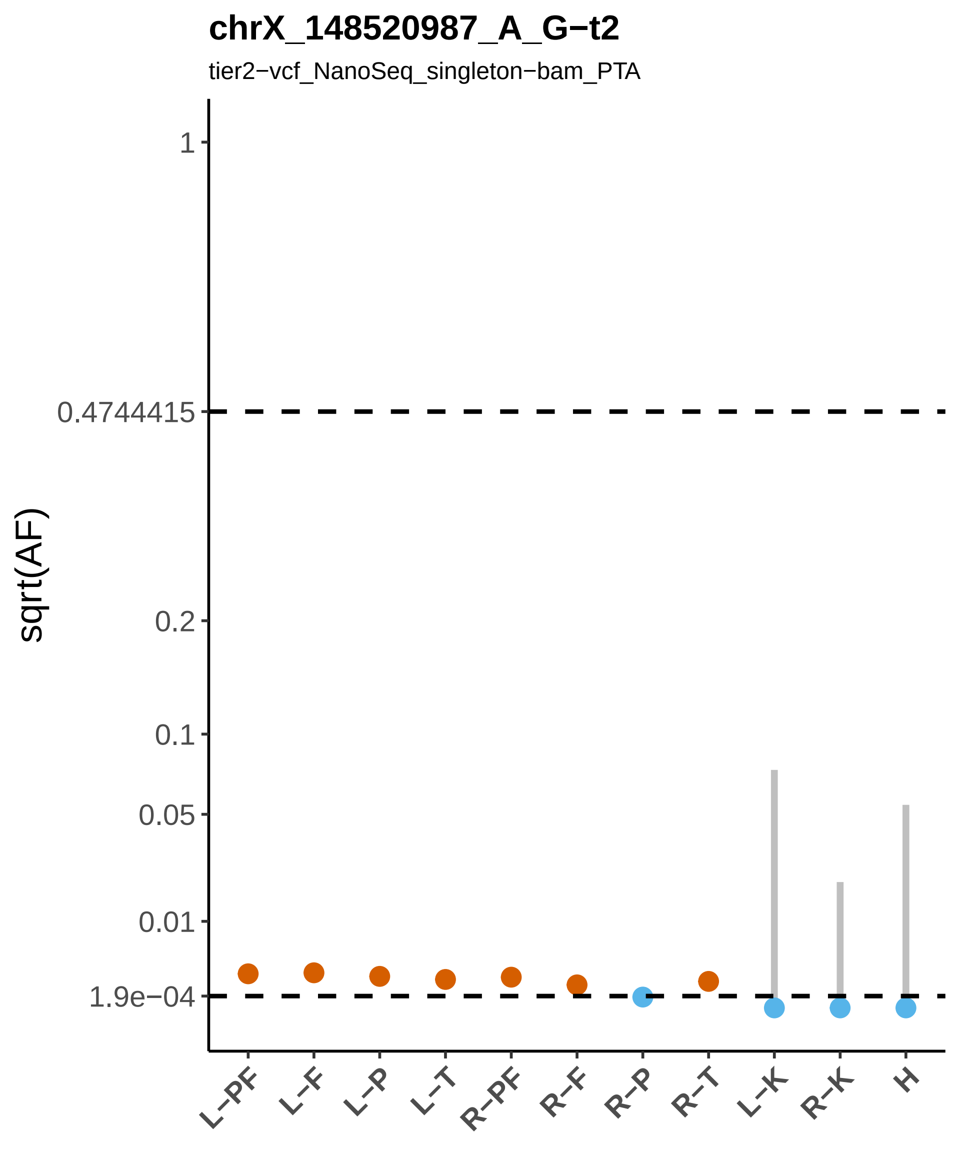
chrX_148520987_A_G−t2
tier2−vcf_NanoSeq_singleton−bam_PTA
sqrt(AF)
1
0.4744415
0.2
0.1
0.05
0.01
1.9e−04
L−PF
L−F
L−P
L−T
R−PF
R−F
R−P
R−T
L−K
R−K
H

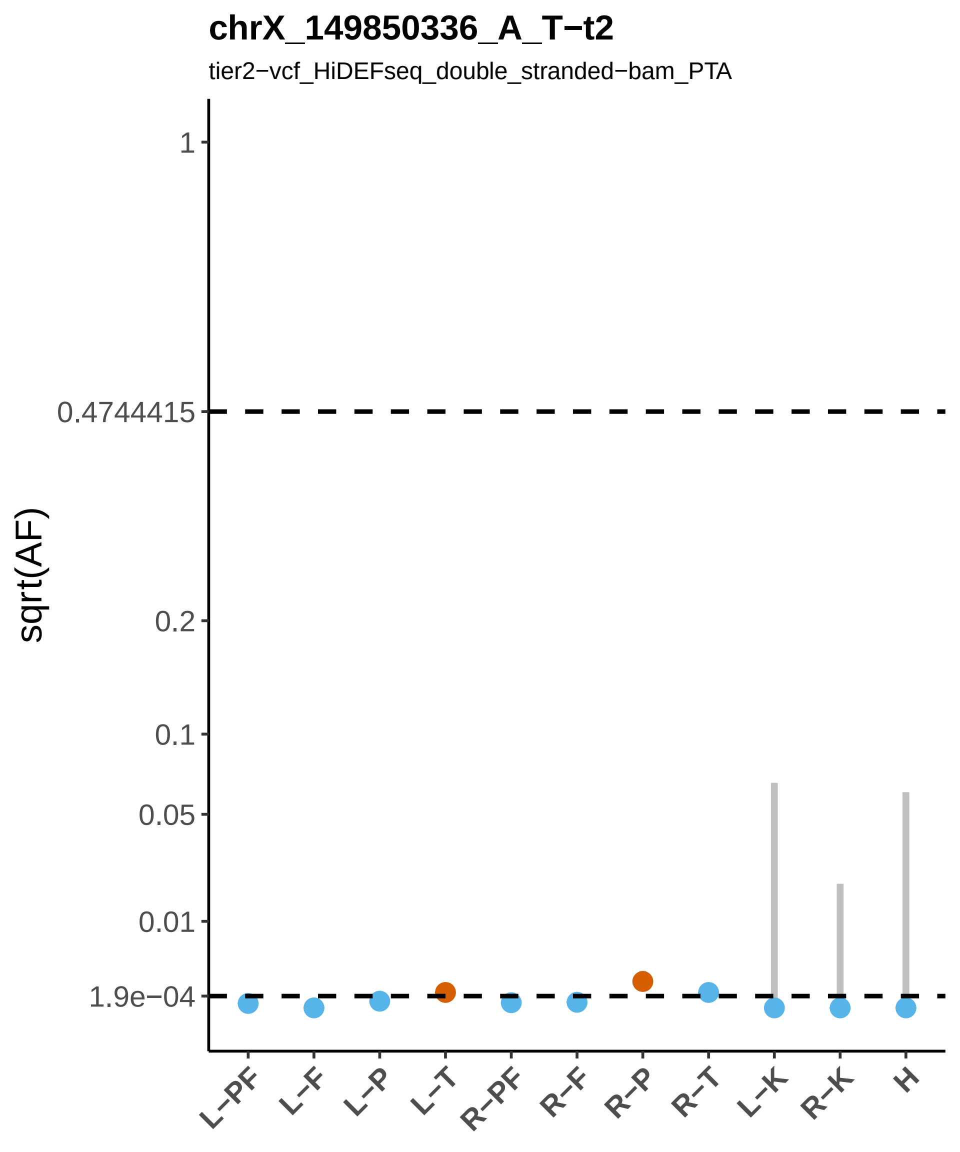
chrX_149850336_A_T−t2
tier2−vcf_HiDEFseq_double_stranded−bam_PTA
sqrt(AF)
1
0.4744415
0.2
0.1
0.05
0.01
1.9e−04
L−PF
L−F
L−P
L−T
R−PF
R−F
R−P
R−T
L−K
R−K
H

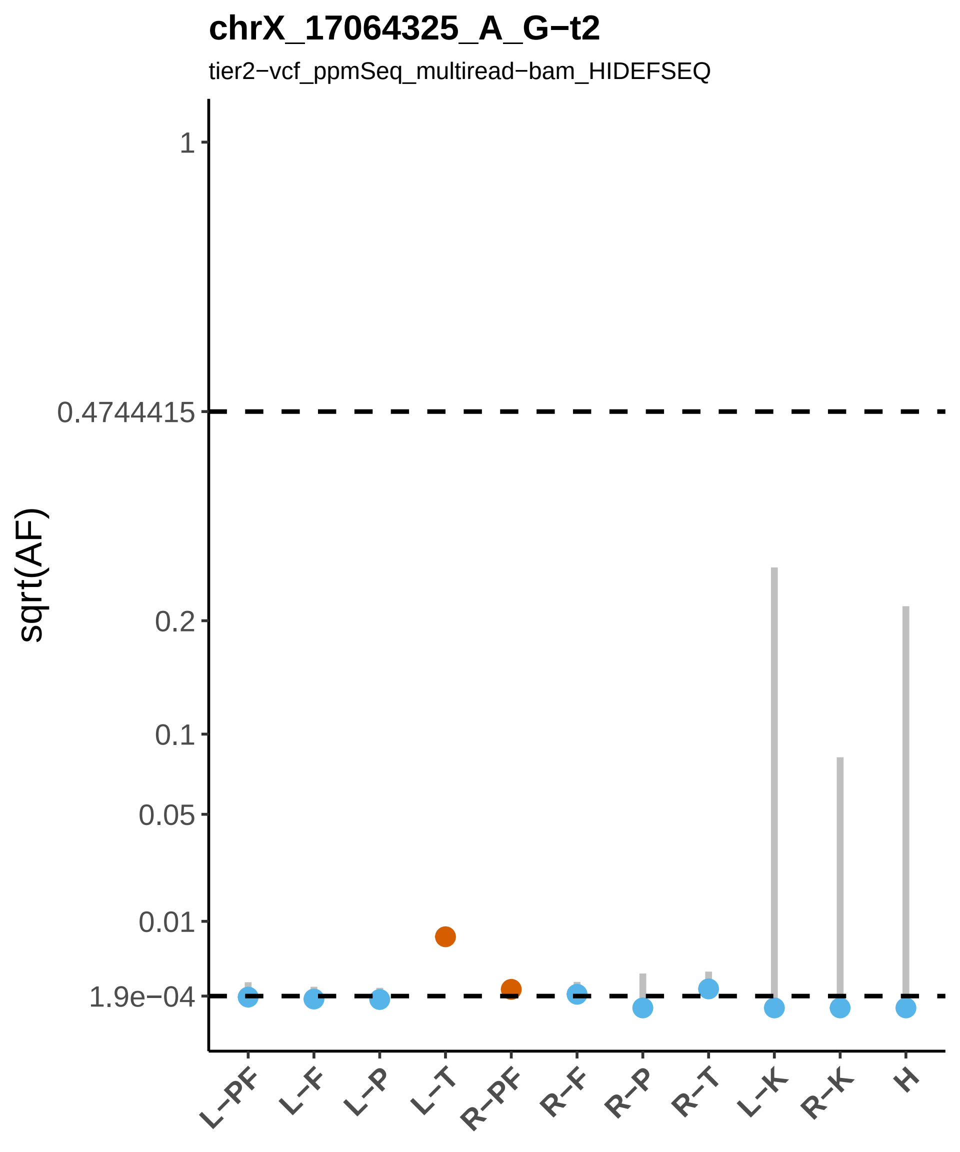

chrX_17064325_A_G−t2
tier2−vcf_ppmSeq_multiread−bam_HIDEFSEQ
sqrt(AF)
1
0.4744415
0.2
0.1
0.05
0.01
1.9e−04
L−PF
L−F
L−P
L−T
R−PF
R−F
R−P
R−T
L−K
R−K
H

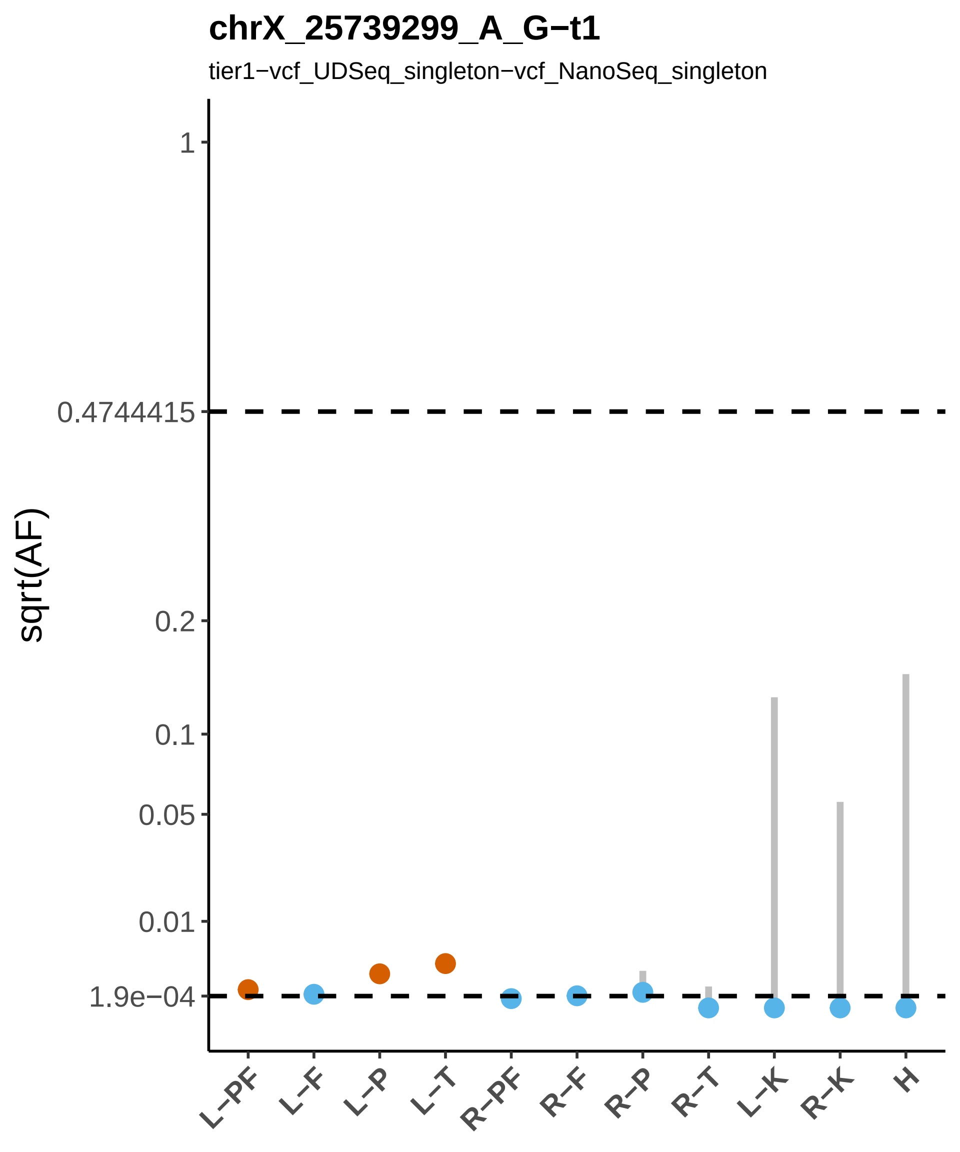

chrX_25739299_A_G−t1
tier1−vcf_UDSeq_singleton−vcf_NanoSeq_singleton
sqrt(AF)
1
0.4744415
0.2
0.1
0.05
0.01
1.9e−04
L−PF
L−F
L−P
L−T
R−PF
R−F
R−P
R−T
L−K
R−K
H

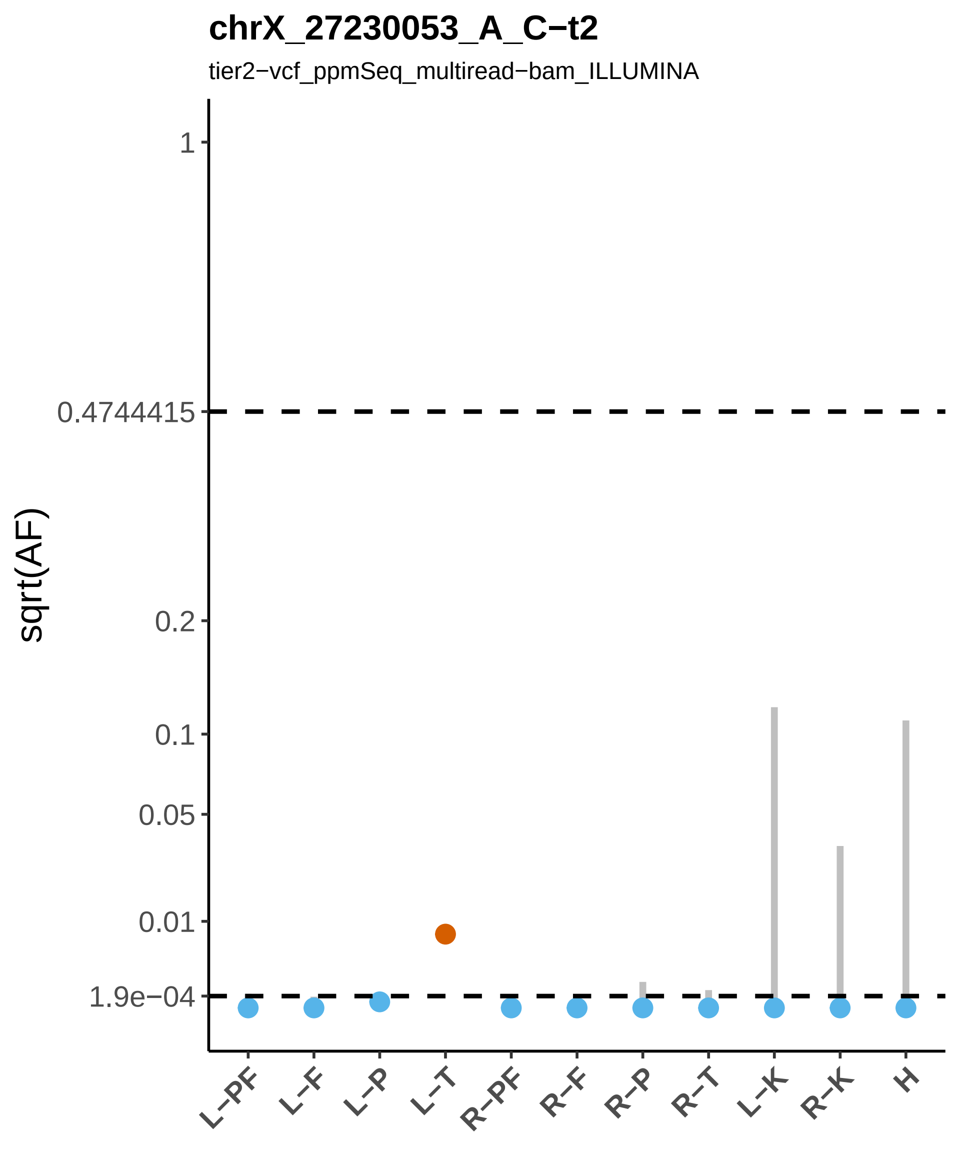
chrX_27230053_A_C−t2
tier2−vcf_ppmSeq_multiread−bam_ILLUMINA
sqrt(AF)
1
0.4744415
0.2
0.1
0.05
0.01
1.9e−04
L−PF
L−F
L−P
L−T
R−PF
R−F
R−P
R−T
L−K
R−K
H

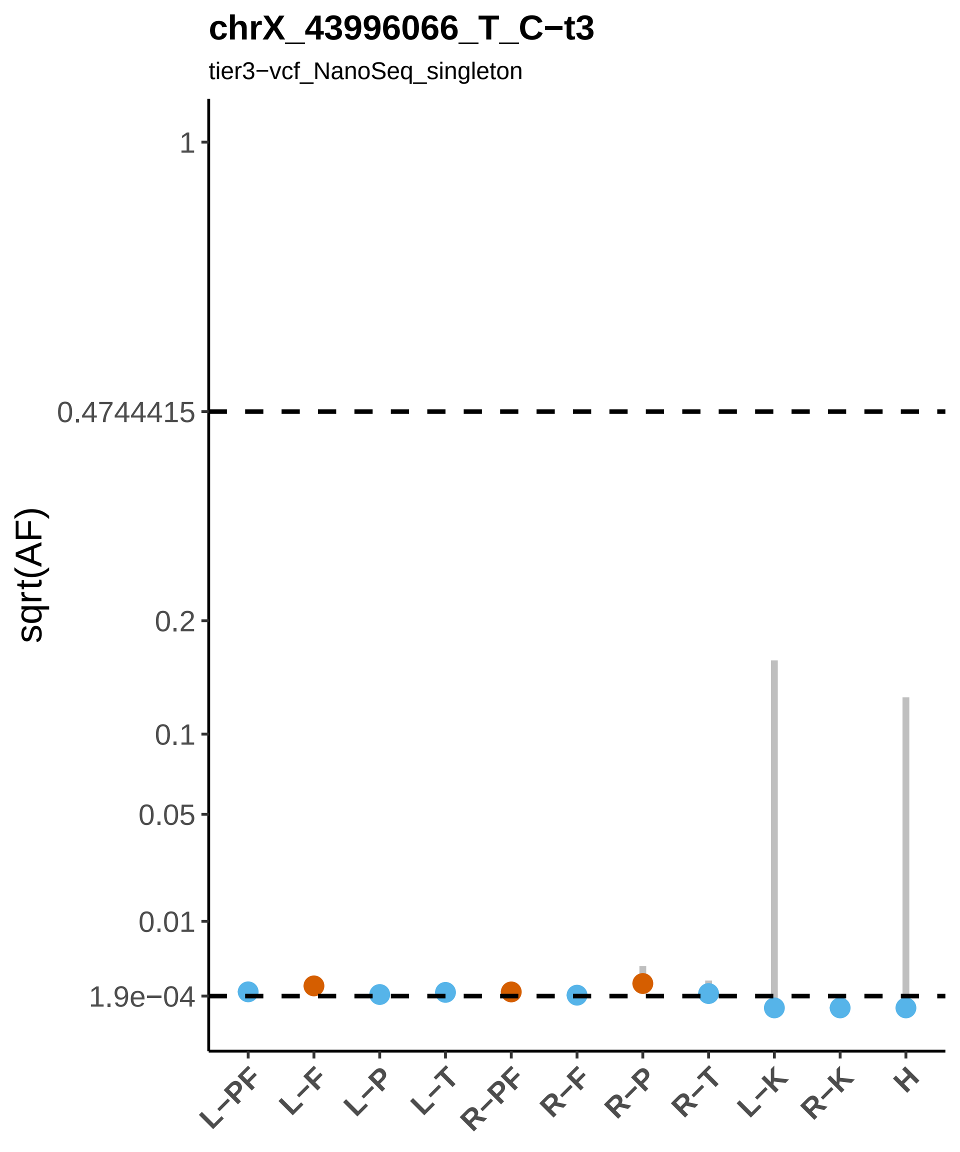
chrX_43996066_T_C−t3
tier3−vcf_NanoSeq_singleton
sqrt(AF)
1
0.4744415
0.2
0.1
0.05
0.01
1.9e−04
L−PF
L−F
L−P
L−T
R−PF
R−F
R−P
R−T
L−K
R−K
H

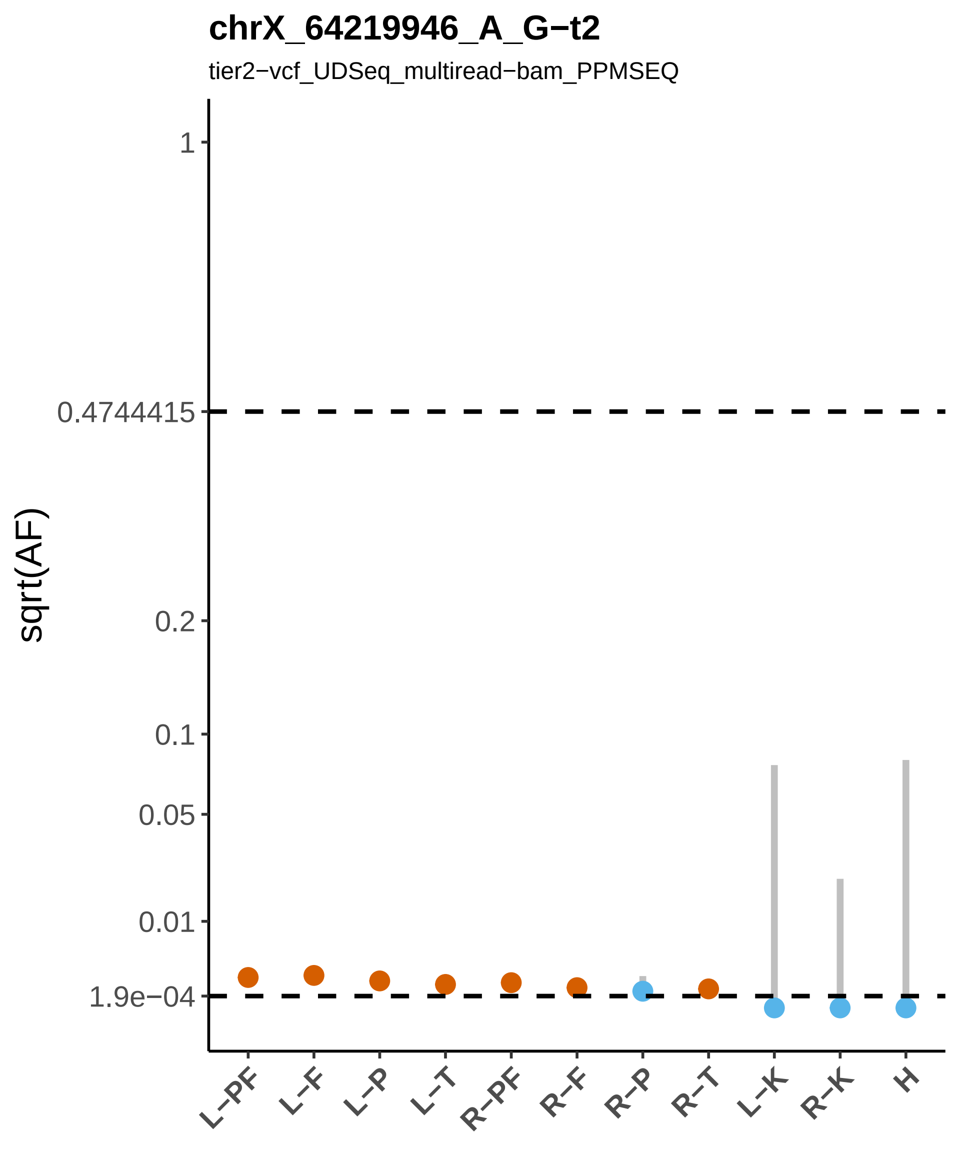

chrX_64219946_A_G−t2
tier2−vcf_UDSeq_multiread−bam_PPMSEQ
sqrt(AF)
1
0.4744415
0.2
0.1
0.05
0.01
1.9e−04
L−PF
L−F
L−P
L−T
R−PF
R−F
R−P
R−T
L−K
R−K
H

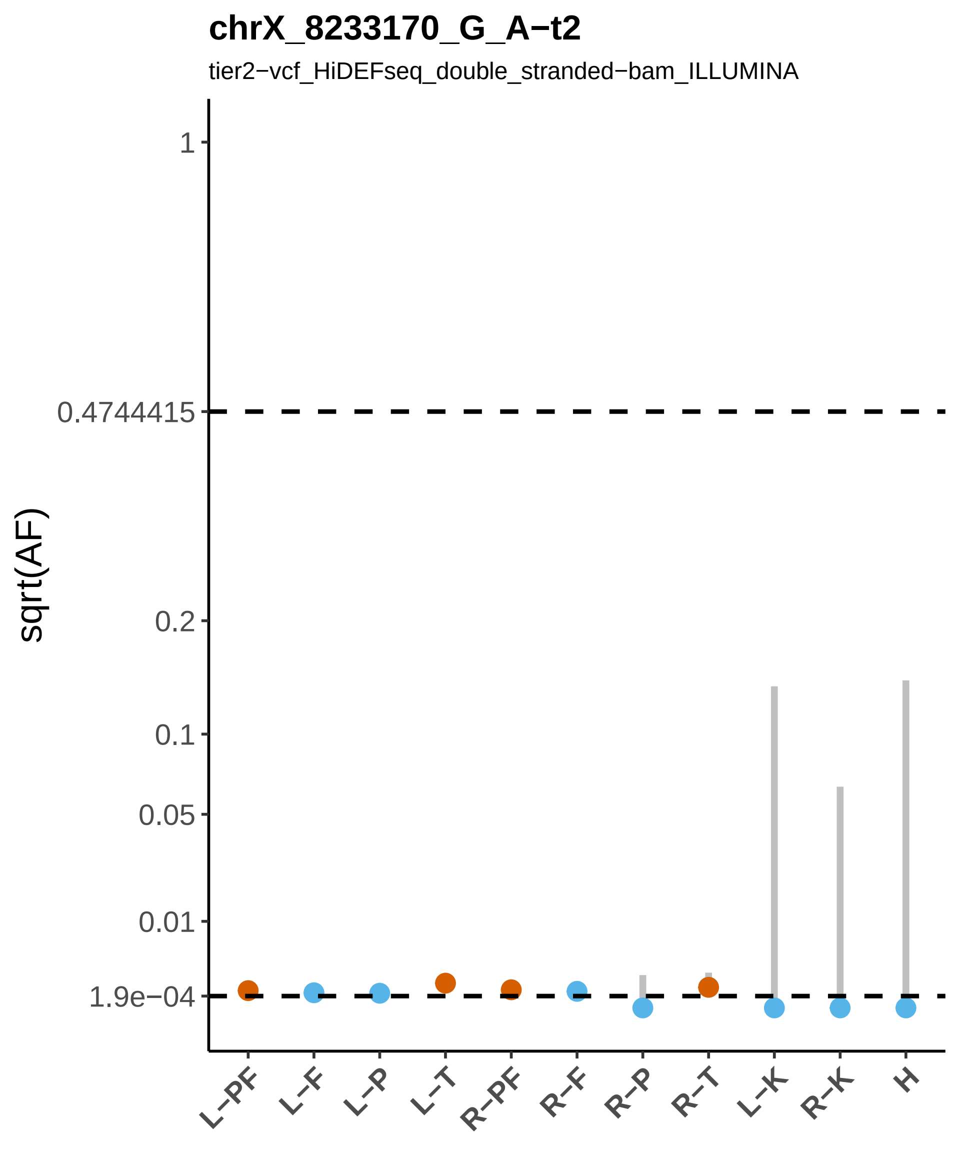

chrX_8233170_G_A−t2
tier2−vcf_HiDEFseq_double_stranded−bam_ILLUMINA
sqrt(AF)
1
0.4744415
0.2
0.1
0.05
0.01
1.9e−04
L−PF
L−F
L−P
L−T
R−PF
R−F
R−P
R−T
L−K
R−K
H

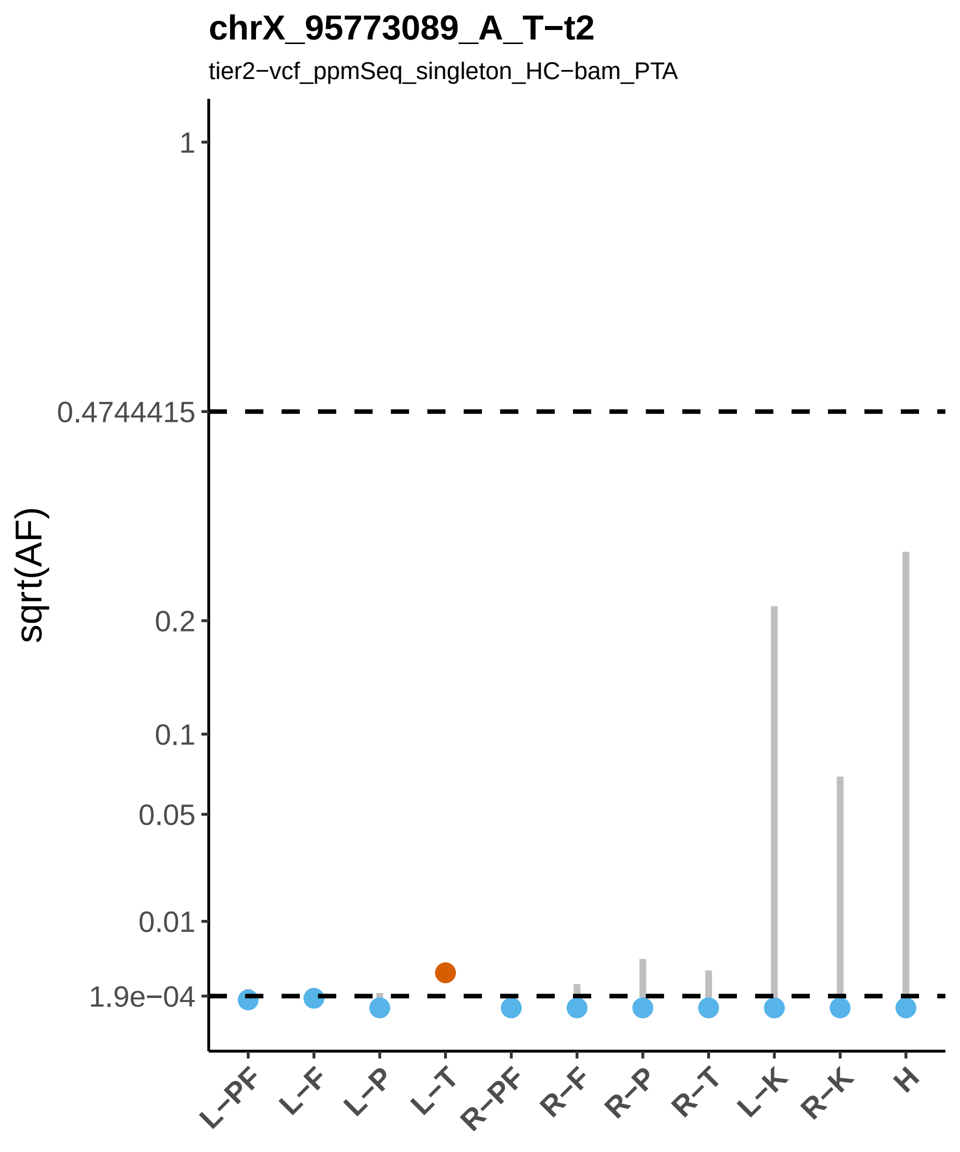

chrX_95773089_A_T−t2
tier2−vcf_ppmSeq_singleton_HC−bam_PTA
sqrt(AF)
1
0.4744415
0.2
0.1
0.05
0.01
1.9e−04
L−PF
L−F
L−P
L−T
R−PF
R−F
R−P
R−T
L−K
R−K
H
